# The western redcedar genome reveals low genetic diversity in a self-compatible conifer

**DOI:** 10.1101/2022.08.26.505492

**Authors:** Tal J. Shalev, Omnia Gamal El-Dien, Macaire M.S. Yuen, Shu Shengqiang, Shaun D. Jackman, René L. Warren, Lauren Coombe, Lise van der Merwe, Ada Stewart, Lori B. Boston, Christopher Plott, Jerry Jenkins, Guifen He, Juying Yan, Mi Yan, Jie Guo, Jesse W. Breinholt, Leandro G. Neves, Jane Grimwood, Loren H. Rieseberg, Jeremy Schmutz, Inanc Birol, Matias Kirst, Alvin D. Yanchuk, Carol Ritland, John H. Russell, Joerg Bohlmann

## Abstract

We assembled the 9.8 Gbp genome of western redcedar (WRC, *Thuja plicata*), an ecologically and economically important conifer species of the Cupressaceae. The genome assembly, derived from a uniquely inbred tree produced through five generations of self-fertilization (selfing), was determined to be 86% complete by BUSCO analysis – one of the most complete genome assemblies for a conifer. Population genomic analysis revealed WRC to be one of the most genetically depauperate wild plant species, with an effective population size of approximately 300 and no significant genetic differentiation across its geographic range. Nucleotide diversity, π, is low for a continuous tree species, with many loci exhibiting zero diversity, and the ratio of π at zero-to four-fold degenerate sites is relatively high (∼ 0.33), suggestive of weak purifying selection. Using an array of genetic lines derived from up to five generations of selfing, we explored the relationship between genetic diversity and mating system. While overall heterozygosity was found to decline faster than expected during selfing, heterozygosity persisted at many loci, and nearly 100 loci were found to deviate from expectations of genetic drift, suggestive of associative overdominance. Non-reference alleles at such loci often harbor deleterious mutations and are rare in natural populations, implying that balanced polymorphisms are maintained by linkage to dominant beneficial alleles. This may account for how WRC remains responsive to natural and artificial selection, despite low genetic diversity.

## Introduction

Gymnosperms are an ancient group of plants, with fossil records dating over 300 million years ago (MYA) (Stewart 1983). Conifers are by far the largest group of gymnosperms with approximately 615 known species (Christenhusz et al. 2011; Farjon 2018). The Pinaceae form the largest conifer family, and genomes of numerous members of the Pinaceae, such as white spruce (*Picea glauca*), Norway spruce (*Picea abies*), loblolly pine (*Pinus taeda*), sugar pine (*Pinus lambertiana*), and Douglas-fir (*Pseudotsuga menziesii*), have been sequenced (Birol et al. 2013; Nystedt et al. 2013; De La Torre et al. 2014; Zimin et al. 2014, 2017; Warren et al. 2015; Stevens et al. 2016; Neale et al. 2017). Such efforts revealed the notoriously complex nature of their immense genomes, which are rife with repetitive sequences, transposable elements, long introns, gene duplications, pseudogenes, and gene fragments.

However, little genomic research has been completed on conifers of other families. In particular, the Cupressaceae, such as cypresses, junipers, and redwoods, are thought to have undergone a whole genome duplication unique from the Pinaceae (Li et al. 2015). There is also evidence for substantial rearrangements of orthologous linkage groups during the evolutionary history of the two families, that resulted in differences in karyotype (*n* = 11 in Cupressaceae; *n* = 12 in Pinaceae) (De Miguel et al. 2015), genome size (9 – 20 Gbp in Cupressaceae; 18 – 31 Gbp in Pinaceae) (Hizume et al. 2001; De La Torre et al. 2014; Stevens et al. 2016), and likely other genomic differences. However, the genomes of only two Cupressaceae species, giant sequoia (*Sequoiadendron giganteum*) and the hexaploid coast redwood (*Sequoia sempervirens*), have been published (Scott et al. 2020; Neale et al. 2022).

Western redcedar (WRC, *Thuja plicata*) is an ecologically, economically, and culturally important species in the Cupressaceae. Endemic to the Pacific Northwest of North America and ranging from Northern California to Southern Alaska, WRC is a stress-tolerant, slow-growing tree prized for its durable, lightweight and rot-resistant wood (Grime 1977). WRC is one of only five extant *Thuja* species and is estimated to have diverged from its North American sister species *Thuja occidentalis* around 26 MYA (Li and Xiang 2005). Genetic studies have indicated low diversity in WRC (Copes 1981; Glaubitz et al. 2000; O’Connell et al. 2008). Microsatellite data suggest that all WRC originated from an isolated refugium near the southern end of its current distribution and radiated north and inland following the last glacial period (O’Connell et al. 2008). Current climate models predict that its range will increase over the next century, particularly in the interior of British Columbia (BC) (Gray and Hamann 2013), thus making it a priority for genome analysis and expediting of traditional breeding cycles via genomic selection (GS).

Uniquely among conifers, WRC employs a mixed mating system of outcrossing and self-fertilization (selfing), with a mean outcrossing rate of around 70% (El-Kassaby et al. 1994; O’Connell et al. 2001, 2004), and appears to suffer very little inbreeding depression for fitness growth traits (Wang and Russell 2006; Russell and Ferguson 2008). Mating systems in plants, particularly rates of selfing (*s*) and its complement, outcrossing (1 –*s*), are of interest to evolutionary biologists due to their implications for genetic diversity and fitness and have been investigated extensively over the past century (Stebbins 1957; Lande and Schemske 1985; Barrett and Eckert 1990; Barrett et al. 2003; Wright et al. 2013). Though inbreeding depression resulting from selfing can lead to negative fitness impacts, a benefit of selfing may include reproductive assurance (Fisher 1941; Baker 1955), which, in the absence of strong inbreeding depression, can allow self-compatible populations to expand their geographic range faster than obligate outcrossers (Lande and Schemske 1985). Research on inbreeding in plants has mostly focused on mating strategies in angiosperms (Barrett and Eckert 1990; Jarne and Charlesworth 1993; Vogler and Kalisz 2001; Barrett et al. 2003; Kalisz et al. 2004; Wright et al. 2013). Characterization of mixed mating systems in conifers has been limited, largely by their long generation times and generally high self-incompatibility (Sorensen 1982; Bishir and Namkoong 1987; Remington and O’Malley 2000; Williams et al. 2003; Williams 2008). The exceptional ability of WRC to maintain such a mating system has allowed for successful selfing for up to five generations in experimental trials, making WRC a potential model for the study of inbreeding in conifers and more broadly in gymnosperms (Russell and Ferguson 2008).

Here we introduce the first genome sequence for WRC and present unique features of the genome in the context of genetic diversity and the evolutionary history of WRC. We further explore the effects of extreme selfing on heterozygosity and selective pressures in multiple selfing lines (SLs).

## Results

### The WRC genome assembly represents a highly complete conifer genome

The assembly of conifer genomes remains challenging partly due to their large size (Nystedt et al. 2013; Warren et al. 2015; Stevens et al. 2016; Zimin et al. 2017) and high heterozygosity (Prunier et al. 2016). Given WRC’s unique selfing abilities, we were able to facilitate assembly of a WRC reference genome using a fifth-generation SL tree (2323-211-S5; **Table S1**) expected to be > 98% homozygous. The S5 reference genome assembly was generated from a combination of short fragment paired-end reads, large fragment mate-pair reads, and linked-reads from large molecules, using 13 libraries and 28 lanes of Illumina sequencing, with a sequencing read length of 2×151 bp (**Table S2**). Overall genome depth of coverage was estimated to be 77×.

The WRC genome was previously estimated to be 12.5 Gbp in size across 11 chromosomes (Ohri and Khoshoo 1986; Hizume et al. 2001). GenomeScope (Vurture et al. 2017) estimated the genome size at 9.8 Gbp (**Figure S1**). We calculated approximately one single nucleotide variant (SNV) every 4.6 kbp, for an estimated genome-wide heterozygosity of 0.000216 – an exceptionally low estimate, highlighting the value of SLs for genome sequencing and assembly.

We assembled 7.95 Gbp of the estimated 9.8 Gbp genome to produce a draft assembly with an N50 of 2.31 Mbp, the largest scaffold being 16.3 Mbp. This assembly comprises 67,895 scaffolds > 1 kbp (**Table 1; Table S3**). Benchmarking Universal Single Copy Ortholog (BUSCO) analysis (Simão et al. 2015) determined the genome assembly to be 86% complete in the gene space. This is one of the highest completeness estimates for a conifer genome (**Table 2; Table S4A**).

**Table 1.** Assembly metrics and statistics for each version of the WRC draft genome.

| Assembly version | N50 (Mbp) | NG50 (Mbp) | Largest scaffold (Mbp) | Size (Gbp) | L50 (bp) | LG50 (bp) | Scaffolds > 1kbp |
| --- | --- | --- | --- | --- | --- | --- | --- |
| redcedar-v1 | 1.45 | 1.07 | 9.79 | 7.95 | 1,642 | 2,463 | 94,166 |
| redcedar-v2 | 2.23 | 1.63 | 15.3 | 7.95 | 1,067 | 1,605 | 90,083 |
| redcedar-v3 | 2.31 | 1.71 | 16.3 | 7.95 | 1,035 | 1,551 | 67,895 |

Genome annotation using evidence from Iso-Seq full-length cDNAs resulted in the identification of 39,659 gene models supported by the alignment of unique primary transcripts (**Dataset S1**), and an additional 26,150 alternative transcripts. A total of 25,984 gene models had Pfam protein family annotation, 31,537 had transcriptome support over their full length (100%), and 19,506 had peptide homology coverage support 90% or greater (**Table S5; Dataset S2**). Intron length ranged from 20 bp to 148.3 kbp, which is consistent with estimates from other conifers, with maximum lengths ranging from 68 kbp (Norway spruce) (Nystedt et al. 2013) up to 579 kbp (sugar pine) (Stevens et al. 2016). Scott et al. (2020) reported a maximum intron length of 1.4 Mb in the highly contiguous giant sequoia genome assembly. Repeat elements comprised 60% of the WRC genome, which is low compared to other conifers. Repeats comprised 79% of the sugar pine (Stevens et al. 2016) and giant sequoia genomes (Scott et al. 2020). Single copy orthologs (SCOs) were detected by orthogroup comparison to the giant sequoia gene set, yielding 11,937 SCOs (**Dataset S3**).

BUSCO analysis found the predicted gene set to be 90.5% complete, much higher than any other conifer gene set to date (**Table 2; Table S4B**). We further validated the completeness of the genome assembly and annotation using a panel of 59 full-length WRC sequences from GenBank, of which 48 were reliably identified in the genome annotation. We also searched for a set of 33 WRC terpene synthase (TPS) transcripts, of which we reliably (>90% identity) identified 15 (Shalev et al. 2018) **(Table S6; Dataset S4**). This confirms the completeness of the gene space and quality of the draft genome annotation, while suggesting that BUSCO core genes may somewhat overestimate gene space completeness when considering family or species-specific genes.

**Table 2.** BUSCO genome assembly and predicted gene set completeness of seven currently available conifer genome assemblies.

| Taxon | *Thuja plicata* | *Sequoiadendron giganteum* (Scott et al. 2020) | *Picea glauca* (Warren et al. 2015) | *Picea abies* (Nystedt et al. 2013) | *Pinus lambertiana* (Stevens et al. 2016) | *Pinus taeda* (Zimin et al. 2014) | *Pseudotsuga menziesii* (Neale et al. 2017) |
| --- | --- | --- | --- | --- | --- | --- | --- |
|  | Western redcedar | Giant sequoia | White spruce | Norway spruce | Sugar pine | Loblolly pine | Douglas- fir |
| Family | Cupressaceae | Cupressaceae | Pinaceae | Pinaceae | Pinaceae | Pinaceae | Pinaceae |
| Genome completeness (%) | 86.0 | 84.3 | 49.7 | 34.9 | 61.5 | 49.7 | 74.1 |
| Gene set completeness (%) | 90.5 | 49.9 | 18.0 | 28.1 | 73.3 | 41.7 | 68.5 |

Genome completeness and gene set completeness were estimated in genome mode and protein mode, respectively, on the Embryophyta OrthoDB v10 database. MetaEuk was used for gene prediction in genome mode.

### Population genomic analysis reveals extremely low levels of genetic diversity in WRC

We estimated nucleotide diversity, short-range linkage disequilibrium (LD), population structure, genetic differentiation, and effective population size (*N_e_*) in *n* =112 unrelated trees from across the geographic range of WRC (range-wide population; RWP) (**Table S7)**. Trees were grouped into three subpopulations: Northern-Coastal (*n* = 77), Central (*n* = 26), and Southern-Interior (*n* = 9) (**Figure 1A**). Using a panel of single nucleotide polymorphisms (SNPs) that were genotyped via targeted sequence capture approach, we identified 2,454,925 variant and invariant sites, which were filtered separately and resulted in sets of 18,371 SNPs (**Dataset S5**) and 2,186,998 invariant sites (see **Materials and Methods**). Total mean SNP depth was 34.3×.

**Figure 1:**
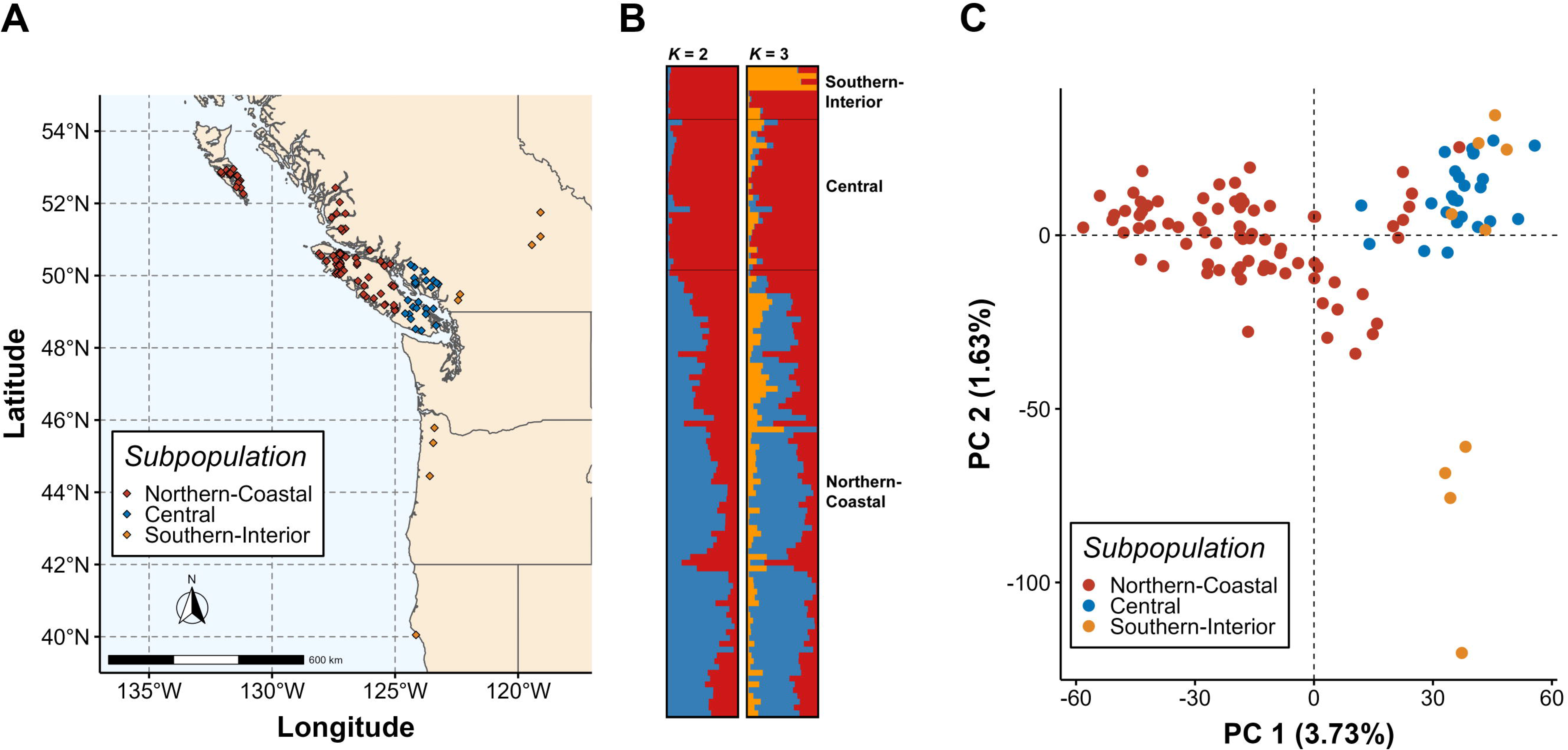
Genetic structure is weak across the geographic range of WRC. A) Map of geographic origin for trees in the range-wide population (RWP) (*n* = 112). Subpopulations were defined *a priori* based on analysis outcomes of O’Connell et al. (2008). Trees were separated into three main subpopulations: Northern-Coastal (*n* = 77); Central (*n* = 26); and Southern-Interior (*n* = 9). **B) STRUCTURE plot of the RWP for K = 2 and K = 3**. Optimal K was determined by evaluating STRUCTURE results using the methods of Evanno et al. (2005) and Puechmaille (2016), and by the approach of fastStructure (Raj et al. 2014). Gene flow is present throughout all three subpopulations. **C) Principal component analysis (PCA) of genetic distance between trees in the RWP.** Latitudinal separation of trees from different s can be observed, although each principal component only explains a very small proportion of the variation between individuals, indicating that genetic differentiation is low.

We annotated 17,728 SNPs across 2,886 genomic scaffolds using the Ensembl Variant Effect Predictor (VEP) (McLaren et al. 2016) (**Dataset S6**). We detected 13,097 SNPs within 5,045 genes, 3,288 of which were SCOs. Intergenic loci made up 25.2% of all annotated SNPs (4,631), 1,105 of which were in regions 0.5 to 2 kbp up- or downstream of coding regions. Within coding regions, 50.0% (3,002) were missense variants while 46.7% (2,807) were synonymous variants (**Table S8**).

### Linkage Disequilibrium

Decay of linkage disequilibrium (LD), the non-random association of alleles at different loci in a population, can inform on how likely different loci are to be assorted together during recombination. We assessed short-range LD as represented by the squared correlation coefficient *r*^2^ in the RWP, at a minor allele frequency (MAF) threshold of 0.05 to avoid bias due to rare alleles (*n* = 16,202 SNPs). The mean of all pairwise *r*^2^ estimates was 0.299 with a median of 0.151. The half-decay value (the distance in which *r*^2^ decays to half of the 90^th^ percentile value) was 0.118 Mbp. LD decayed to an *r*^2^ of 0.2 at 0.751 Mbp, and an *r*^2^ of 0.1 at 2.17 Mbp (**Figure 2A**). Further, high LD (*r*^2^ > 0.8) appears to exist for SNPs millions of bp apart (**Figure 2B**). These estimates for LD decay are several orders of magnitude greater than those found in other conifers, as well as many other tree species, where LD has been reported to decay rapidly within tens to a few thousand bp (Krutovsky and Neale 2005; Heuertz et al. 2006; Pyhäjärvi et al. 2011; Pavy et al. 2012; Fahrenkrog et al. 2017).

**Figure 2:**
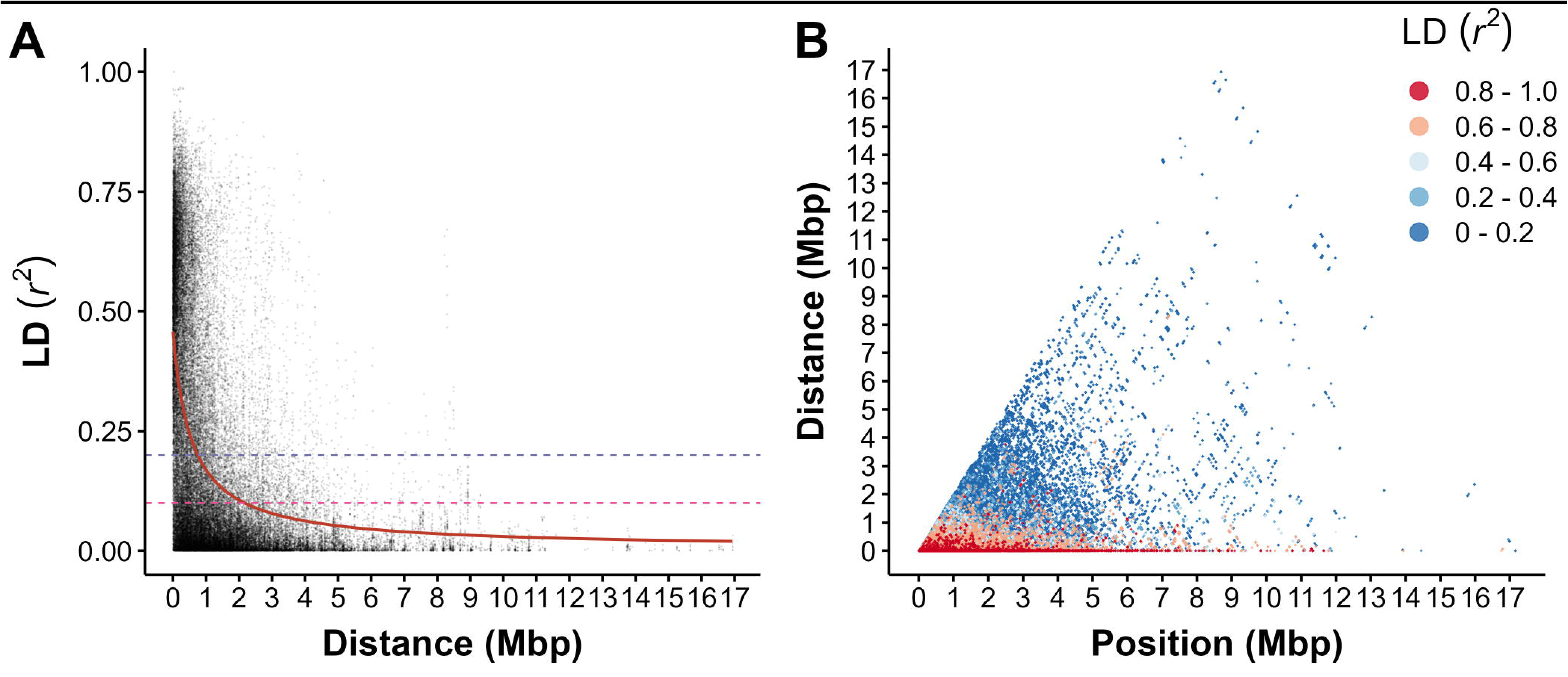
Within-scaffold linkage disequilibrium (LD) in the range-wide population. **A)** LD was assessed using SNPs with a minor allele frequency cutoff of 0.05 to reduce error associated with rare alleles (*n* = 16,202 SNPs). Decay was estimated using a non-linear model (red line); *r*^2^ decayed to baseline thresholds of 0.2 (purple dotted line) and 0.1 (pink dotted line) at 0.751 and 2.17 Mbp, respectively. **B)** Pairwise LD for all pairs of SNPs (*n* = 16,202). Each point on the plot represents the LD between two SNPs at a given distance from one another and relative position on the scaffold. Colour indicates LD range, with red indicating SNPs in strong LD.

### Population structure and genetic differentiation

We analysed STRUCTURE (Pritchard et al. 2000) results using two post-hoc cluster identification methods on a filtered set of *n* = 4,765 SNPs (see **Methods and Materials**). The ΔK method of Evanno et al. (2005) identified an optimal K of two; this approach may return a K of two more often than expected when genetic structure is weak (Janes et al. 2017). The approach of Puechmaille (2016), which can help resolve K when subsampling is uneven, identified an optimal K of two as well. Analysis of fastStructure (Raj et al. 2014) results using cross-validation suggested that optimal K may lie between one and three. These results suggest genetic structure is exceptionally weak in our RWP. Indeed, there is apparent gene flow between trees in all three subpopulations across all three STRUCTURE clusters (**Figure 1B**).

We applied non-parametric approaches of Discriminant Analysis of Principal Components (DAPC) and Principal Component Analysis (PCA) using a set of *n* = 13,427 SNPs from SCO and intergenic regions. Cross-validation for DAPC with *a priori* cluster definitions optimally retained 22 PCs and two discriminant functions capturing 30.3% of the conserved variance; however, *de novo k*-means clustering failed to resolve any clusters, identifying an optimal K of one (**Figure S2**). PCA revealed a latitudinal gradient of differentiation along the first principal component (PC), with some separation of the Southern-Interior subpopulation along the second PC (**Figure 1C**), mostly for trees originating from California and Oregon. However, the first PC only explains 3.73% of the variance in the data, and the second explains 1.63%. These results are consistent with DAPC and suggest that gene flow has been prevalent across the range of WRC. No significant differentiation was found between trees from different subpopulations based on a hierarchical *F*_ST_ test (*F*_ST_ = 0.0334, *p* = 0.726) (**Table S9**), and no significant isolation by distance was found by our Mantel test for subpopulations (*r* = -0.241, *p* = 0.672) and individuals (*r* = 0.0833, *p* = 0.121) (**Figure S3**).

### Nucleotide diversity in WRC

We estimated nucleotide diversity π (Nei and Li 1979) in the RWP and absolute nucleotide divergence *d*_XY_ between subpopulations using all SCO and intergenic SNPs. Average π (SD) across 10,631 SCOs was 0.00272 (0.0122) (**Figure 3A**; **Figure S4; Table S10**); 1,411 genes had a π of zero. Across 10 kb windows, average π was 0.00204 (0.0141), indicating diversity is similar in coding and noncoding regions. Average *d*_XY_ was not significantly different between any pair of subpopulations nor was it significantly different from π in SCOs (*p* > 0.05, Kruskal-Wallis rank sum test) (**Figure 3B**; **Table S11**).

**Figure 3:**
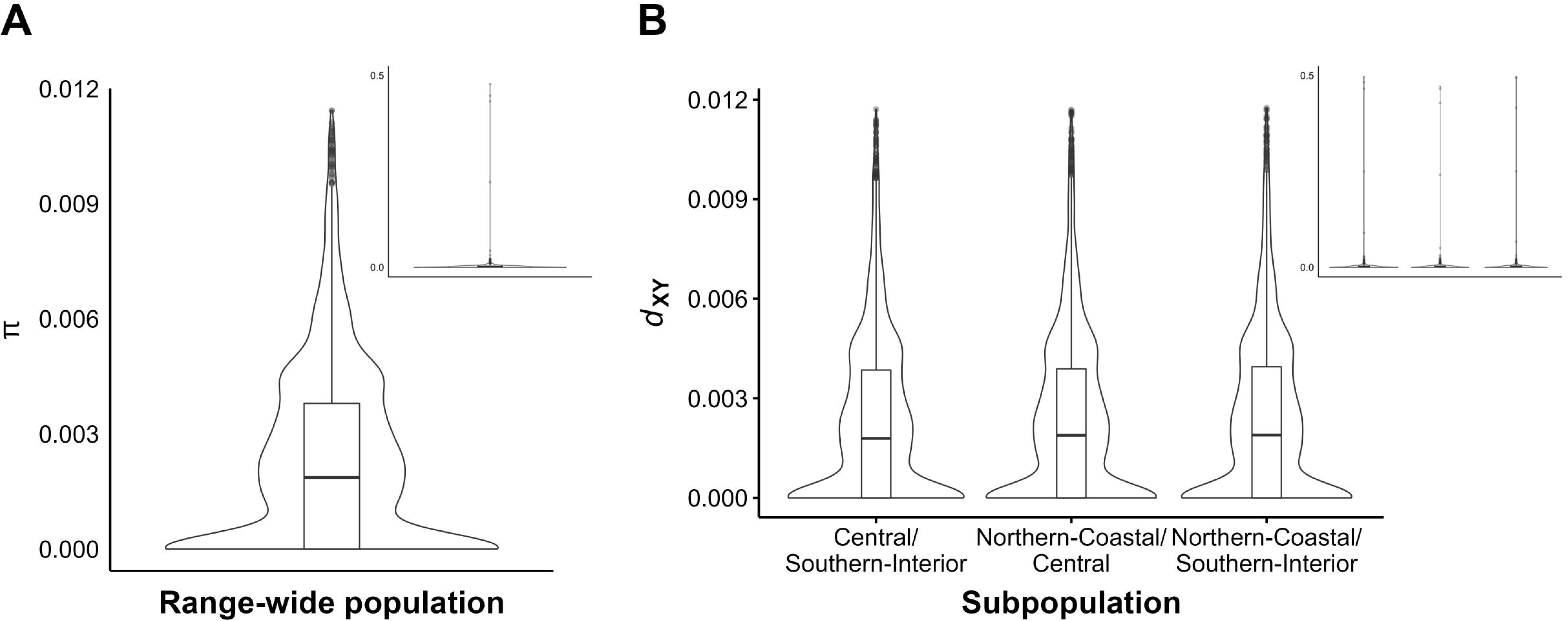
A) Overall distribution of average π of the range-wide population (RWP) in SCOs. We detected 1,411 SCOs with a π of zero, with an average π of 0.00272. B) Overall distribution of average *d*_XY_ between each pair of geographic subpopulations. No significant differences were observed between comparisons of different subpopulations. **Inlays** show all π estimates; **main plots** show π estimates with outliers in the top one percentile removed for clarity. The top 1% of π estimates account for 3% of the total estimated diversity, and the top 5% account for over 13% of the total.

To assess the efficacy of purifying selection, we estimated π_0_/π_4_, the ratio of π in 0-fold to 4-fold degenerate sites. We found a π_0_ of 0.00158 (0.0146) and a π_4_ of 0.00485 (0.0147), yielding a π_0_/π_4_ of 0.325. The site frequency spectrum (SFS) for 4-fold SNPs appeared to decay slower than the SFS for 0-fold and for all SNPs, supporting evidence of a recent bottleneck and indicating that there may be stronger positive selection at these sites (**Figure S5**).

### Effective population size (N_e_)

*N_e_* can be defined as the idealized population size expected to experience the same rate of loss of genetic diversity as the population under observation (Wright 1931). We estimated *N_e_* using the LD method of NeEstimator (Do et al. 2014). SNPs were mapped to the giant sequoia genome (Scott et al. 2020) (**Dataset S7**) and SNPs estimated to be at least 2.17 MB apart were isolated for the analysis, resulting in a set of *n* = 412 SNPs. *N_e_* was estimated to be 270.3 (JackKnife 95% CI: 205.5, 384.6).

We further explored demography using Stairway Plot 2 (Liu and Fu 2020). We observe a decline in *N_e_* from ∼ 500,000 beginning ca. 2 MYA, accelerating from ∼40 KYA down to under 300 by present day, consistent with one or more bottleneck events during the recent glacial maximum (**Figure S6**). Our estimates of *N_e_* are extremely low for a continuous tree population; for example, species in *Picea* (Chen et al. 2010), *Pinus* (Brown et al. 2004), and *Populus* (Fahrenkrog et al. 2017) have estimated *N_e_* in the range of 10^4^ – 10^5^.

### Persistent heterozygosity during complete selfing highlights genomic regions under selection

To examine the effect of complete selfing on heterozygosity and selection in WRC, we selected 189 trees from the 15 FS families, forming 41 SLs for SNP genotyping. The process of SNP calling is error-prone, and despite filtering for multiple quality criteria, errors are likely to remain in any SNP data set. Using SLs, we were able to correct for erroneous genotyping calls and impute missing genotypes for SLs up to S4 (*n* = 28) or S5 (*n* = 11), retaining *n* = 151 trees (**Dataset S8**). We used all filtered SNPs for these analyses (*n* = 18,371 SNPs).

Under selfing in diploids, heterozygosity is expected to decline by 50% in each generation purely through genetic drift. Mean heterozygosity declined slower than expected (**Figure 4A**; **Table S12; Table S13A**), while median observed heterozygosity was significantly lower than expected beginning in generation S3 (*p* < 0.05, pairwise Sign test; **Table S13B**). Mean heterozygosity at FS was 0.296, while in the RWP it was 0.219 (**Table S12**). In comparison, Chen et al. (2013) found mean heterozygosities of 0.33 and 0.36 in lodgepole pine (*Pinus contorta*) and white spruce, respectively.

**Figure 4:**
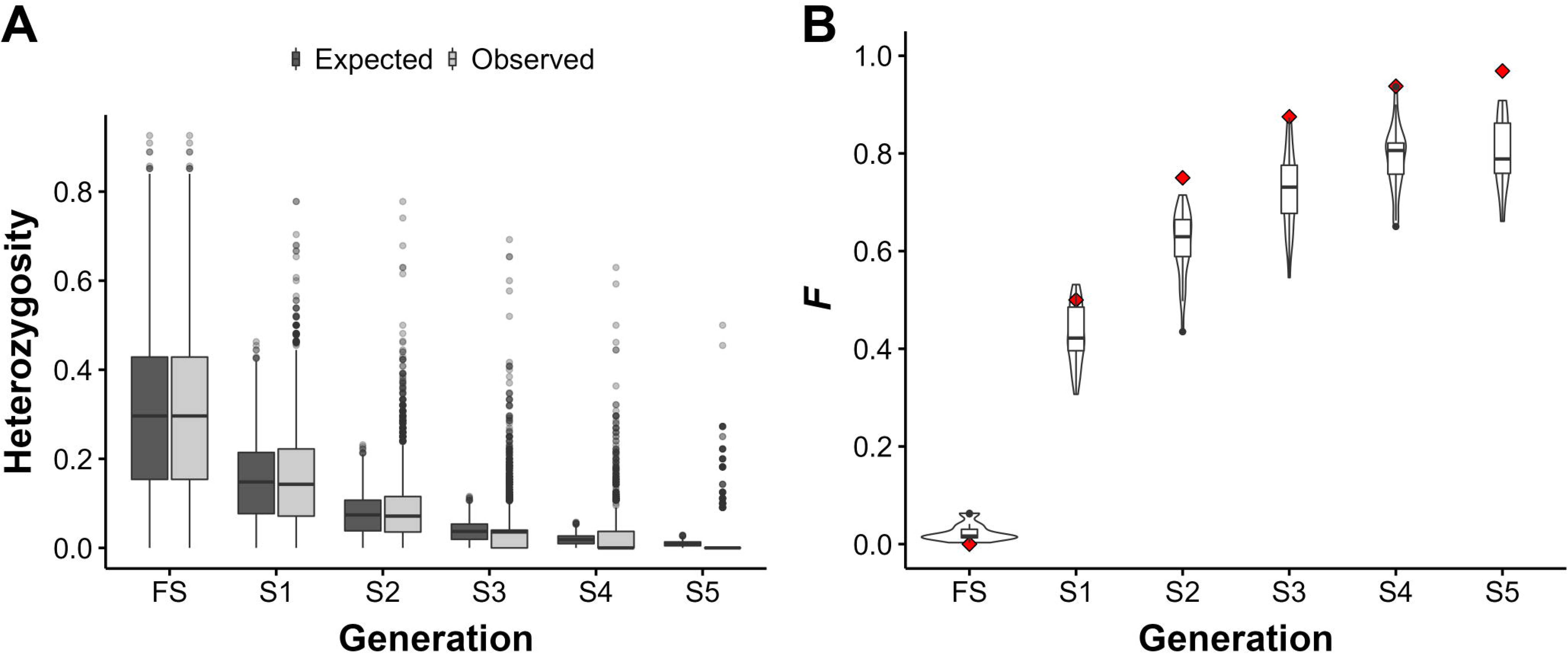
Change in heterozygosity (H) and inbreeding coefficients (*F*) over five successive generations of complete selfing in WRC. **A)** Observed vs. expected change in heterozygosity over five successive generations of complete selfing in *n* = 28 (FS – S4) and *n* = 11 (S5) different selfing lines (SLs), at *n* = 18,371 SNP loci, after manual error correction. Each line at each generation is represented by a single tree. Black points indicate boxplot outliers. Observed median heterozygosity declines faster than expected under theoretical expectations during complete selfing, despite many SNP loci remaining heterozygous across all generations. **B)** Inbreeding coefficients (*F*) for *n* = 28 samples (FS – S4) and *n* = 11 samples (S5). Black points indicate boxplot outliers. Under complete selfing, *F* is expected to increase by a factor of ½(1+*F*) in the previous generation (red diamonds). *F* increases at a slower rate than expected in our SLs.

The inbreeding coefficient *F* is the probability of any two alleles being identical by descent (IBD) and is a measure of reduction in heterozygosity due to inbreeding. We estimated *F* for each sample in the SLs and the RWP using the approach of Yang et al. (2011) (*F*_UNI_). Mean *F* increased from 0.00569 at FS to 0.801 at S5 – significantly less than expected (*p* < 0.05, one-sample *t*-test), indicating that the observed reduction in heterozygosity cannot entirely be attributed to inbreeding (**Figure 4B**; **Table S12; Table S13C**). Mean *F* in the RWP was 0.331, further emphasizing the degree of inbreeding in wild populations **(Table S12**).

Following the expectation of a 50% decline in heterozygosity per generation under complete selfing and assuming a model of only genetic drift, we anticipated that 25% of SLs would become fixed for one allele at any given locus and 25% for the other in each generation. Thus, by generation S4, 46.875% of SLs should fix for each allele, and 6.25% should remain heterozygous. We identified 83 SNPs that deviated from expected proportions of fixation at a false discovery rate threshold of 0.05 (hereafter: outlier SNPs) (**Dataset S9**). Of these, 15 fixed for the reference allele and 2 fixed for the alternate allele more often than would be expected under drift alone; meanwhile, all outlier SNPs had a higher proportion of heterozygous alleles by S4 than expected under drift. Outlier SNPs were present on all putative LGs in the genome, and mean depth was similar to the mean depth of the total SNP set (29.3× vs 34.3×, respectively). VEP predicted effects for 67 outlier SNPs, 14 (16.9%) of which were in coding regions (**Dataset S10**). Gene Ontology (GO) annotation was available for 30 genes containing outlier SNPs; ten GO categories were over-represented in outliers when compared to the entire SNP set (*p* < 0.05, Fisher’s Exact Test) (**Table S14**).

When comparing SNP effect categories, we found intergenic variants (1.76×10^-5^) to be over-represented in outlier SNPs, while synonymous variants (*p* = 0.0274) and 3’ UTR variants (*p* = 0.00993) were under-represented (**Figure S7**). In the RWP, the minor allele for these SNPs was nearly always the alternate allele, i.e., the allele inducing the change. This pattern suggests that balanced polymorphisms may be maintained by selection favouring linked dominant alleles, i.e., associative overdominance (Bierne et al. 2000).

## Discussion

Genome analysis of WRC, a self-compatible conifer, revealed low genetic diversity, high levels of LD, and low *N_e_* across its geographic range. WRC emerges as a genetically depauperate wild plant species, providing insight into how selfing may have facilitated its expansion across its current geographic range, but at the expense of genetic variation.

### The WRC genome

Sequence assembly of large and repetitive conifer genomes is becoming more feasible, with new technologies such as Single Molecule Real Time (SMRT) long-read sequencing or linked-reads (Zimin et al. 2017). WRC is one of only two conifers outside of the Pinaceae whose genome sequence has been published. Recently, Scott et al. (2020) reported the first genome sequence in Cupressaceae, giant sequoia, with a near-chromosome-scale assembly of 8.125 Gbp of the estimated 9 Gbp genome using a combination of Oxford Nanopore long reads and Illumina short reads together with a Dovetail HiRise Chicago and Hi-C statistical scaffolding and assembly approach. Though future chromosome-level assembly would be of value to improve contiguity, BUSCO completeness scores (86.0% and 84.3% for WRC and giant sequoia, respectively) and the very high completeness of the annotated gene set suggest that the WRC assembly is currently of very high quality for the gene space. Previous studies using flow cytometry estimated WRC’s genome size at 12.0 – 12.5 Gbp (Ohri and Khoshoo 1986; Hizume et al. 2001). The WRC genome assembled at 7.95 Gbp. This discrepancy may be partially explained by filtering of *k*-mers with very high depth of coverage in GenomeScope to remove organelle-derived reads, which may also remove other heterochromatic sequences such as centromeres and telomeres; however, a recent study in maize (*Zea mays*) found that selfing over several generations can reduce genome size by up to 7.9% (Roessler et al. 2019), which may suggest that we are observing genome loss in WRC as well given the genome assembly source. Flow cytometry to assess genome loss during selfing would be a valuable future endeavour.

### Genetic diversity in WRC

We estimated π to be 0.0027 in SCOs and 0.0020 across all sequenced space. These estimates are lower than many other plant species using comparable methods, for example, Norway spruce (π = 0.0049 – 0.0063) (Wang et al. 2020), weedy broomcorn millet (π = 0.14; *Panicum miliaceum*) (Li et al. 2021), and most *Populus* species (π = 0.0041 – 0.011) (Liu et al. 2022). We found lower π estimates only in highly cultivated plants, such as soybean (π = 0.0015; *Glycine* spp.) (Bayer et al. 2022), or rare, isolated species, such as *Populus qiongdaoensis* (π = 0.0014), which is restricted to a single small island and has an estimated *N_e_* of ∼500 (Liu et al. 2022). Additionally, our probe selection strategy for genotyping targeted regions of high variability due to the very low levels of polymorphism in initial sequencing runs. This may have led to inflated estimates of π, suggesting that genome-wide diversity may be even lower.

The relatively high observed π_0_/π_4_ ratio (0.33) may suggest weak purifying selection in WRC (Chen et al. 2017); it could also be indicative of demography, as π_0_ returns to equilibrium quicker than π_4_ following a bottleneck event (Brandvain and Wright 2016; Chen et al. 2019). The low *N_e_* (∼ 300) and general lack of population structure, genetic differentiation, or nucleotide divergence between geographic subpopulations despite a relatively wide geographic range and continuous population suggest that much of the variation in WRC was likely eliminated due to bottlenecks following the last glacial period, a pattern confirmed by our Stairway Plot results and affirming the conclusions of previous studies (Copes 1981; Glaubitz et al. 2000; O’Connell et al. 2008). Mating system likely plays a role as well in WRC’s low diversity. It has been argued that selfing species should generally have a lower nucleotide diversity due to a reduction of the effective recombination rate (Buckler IV and Thornsberry 2002). Thus, the exceptionally slow rate of LD decay observed in our RWP is further evidence of a recent population bottleneck or long-term effects of inbreeding (Golding and Strobeck 1980; Zhang et al. 2004; Slatkin 2008). In future studies, more extensive sampling, in particular for the Southern-Interior region, could help in gaining more accurate estimates of genetic differentiation across populations of WRC.

### Selfing in WRC

Heterozygosity declined faster than expected under complete selfing. Despite starting at an *F* of nearly 0, our FS generation had a low mean heterozygosity (0.296), and with each successive generation, *F* increased slower than expected, indicating that IBD does not fully explain the reduction in heterozygosity in WRC (Wright 1922; Slate et al. 2004). The lack of strong fitness costs associated with selfing in WRC (Wang and Russell 2006; Russell and Ferguson 2008) suggests that most strongly deleterious alleles have been purged from the genome, presumably due to past population bottlenecks and inbreeding. Nonetheless, even weak purifying selection could explain the faster than expected decline in heterozygosity.

Of greater interest, however, is that the majority of loci deviating from expectations of drift during selfing remained heterozygous, suggestive of balancing selection or associative overdominance at these loci, with high levels of LD promoting genetic hitch-hiking near loci under selection to remain heterozygous. The presence of missense variants in outlier loci coupled with the general rarity of missense mutations in natural populations offers further support for associative overdominance as an explanation for the retention of heterozygosity at these loci. This is congruent with relatively high π_0_/π_4_ in the RWP, suggesting strong positive selection is maintaining current allele frequencies. No outlier loci remained heterozygous in all lines, which suggests that these loci do not harbour strongly deleterious or lethal mutations. Further, all three genotypes exist for many of these loci in the RWP.

Excess heterozygosity can also occur from genotyping error due to the presence of paralogs. To address this source of error, we employed stringent filters for maximum mean depth, allele balance, excess heterozygosity, read-ratio deviations, and deviations from HWE. Further, the low estimated π in the RWP as well as similar mean depths (∼30×) for outlier SNPs and the total SNP set suggest paralog content is minimal. Higher than expected heterozygosity was observed during selfing in eucalypts (*Eucalyptus grandis*) (Hedrick et al. 2016) and maize (Roessler et al. 2019). However, the average heterozygosity in these species is notably much higher (∼ 0.65 in each for S1, compared with 0.15 at S1 for WRC). It is also possible that our genotyping strategy, in which probes were designed to capture highly variable sites, may have influenced heterozygosity estimates. Future analyses using whole-genome sequencing or comprehensive genotyping-by-sequencing (GBS) for comparison may be of value.

We recognize that use of SLs of single seed descent makes differentiating between patterns of selection and genetic drift difficult, as genetic drift is stronger when there are fewer individuals in a population. The use of multiple cloned seedlings for each SL in future studies could help improve our analysis, with the potential to find more SNPs under selection.

### Implications for conservation, adaptation to climate change and breeding with genomic selection (GS)

Current breeding of WRC focuses on traits such as growth and herbivore and disease resistance; thus, low genetic diversity may have considerable ecological and potential economic consequences. When low genetic diversity is observed in plant or animal populations, conservation strategies may become necessary to maintain existing genetic variation and reduce the risk of extreme inbreeding depression, especially when census population size in the wild is small. Although ours and previous results (O’Connell et al. 2008) indicate its range was likely reduced to a single refugium during the last glaciation, WRC has since greatly expanded throughout the Pacific Northwest. We found genetic isolation by distance to be small, consistent with the low observed variation. Yet, successful selection of genetically superior families for these traits has been possible. Provenance trials have revealed significant local adaptation among natural populations of WRC (Cherry 1995), and WRC can be found in a variety of different climates, moisture levels, elevations, and light availabilities (Grime 1977; Antos et al. 2016). Resistance to cedar leaf blight, a foliar fungal pathogen, has been observed to be an adaptation to native climate, with trees from wetter climates showing greater resistance than those from drier climates, regardless of geographical distance (Russell et al. 2007). These observations suggest sufficient genetic variation exists within and between natural populations upon which selection can act. Furthermore, WRC is well known for its high phenotypic plasticity (El-Kassaby 1999), possibly due to epigenetic variation (Zhang et al. 2013), although the fraction of plasticity that is adaptive remains unknown. Our observation of balanced polymorphisms, due in part to associative overdominance, offers a potential explanation for WRC’s reported adaptability and response to selection. Together with self-compatibility, which is known to facilitate range expansion (Baker 1955), WRC may be less threatened by climate change and other anthropogenic pressures than might be expected based on its low genetic diversity.

WRC’s apparent adaptability and potential for range expansion make it an important forest tree in a time of changing climate and environments (Gray and Hamann 2013). Low genetic diversity and unique mating system need to be considered as WRC breeding adopts strategies of GS that largely rely on controlling relatedness in the population (Ritland et al. 2020). Recombination rate is another important consideration, as GS relies on the presence of LD between SNPs and causal regions for traits, in addition to relatedness between individuals (Meuwissen et al. 2001). WRC’s high LD may be an advantage for finding linked SNPs, but may also increase the risk of unintentional selection for correlated traits not under selection. This could be mitigated by whole genome sequencing across breeding populations, similar to GS approaches in livestock breeding (Raymond et al. 2018; Georges et al. 2019).

WRC is a fascinating example of adaptation in a long-lived conifer, despite very low levels of genetic variation. As our understanding of the genome improves, we will be able to improve prospects for survival and maintenance of this tree as an ecologically and economically significant species and better understand and test how selfing behaviour evolves and can be advantageous in wild plant populations.

## Methods and Materials

### Plant materials

The WRC RWP represented *n* = 112 individuals originating from across the geographic range growing at the Cowichan Lake Research Station (CLRS) at Mesachie Lake, BC, Canada. Trees were separated into three geographic subpopulations, Northern-Coastal, Central, and Southern-Interior, based on UPGMA clustering of genetic distances (O’Connell et al. 2008). SLs were produced over 12 years (1995 – 2007) using an accelerated breeding approach (Russell and Ferguson 2008) at CLRS. Briefly, 15 pairs (30 individuals) of unrelated parents from across coastal BC and Vancouver Island (**Table S15**) were crossed to create 15 FS families and ensure an initial inbreeding coefficient of *F* = 0. Each FS line was then selfed for up to five generations (S1 – S5) with one generation every two years, facilitated by GA_3_ hormone treatment. A single S5 seedling of SL 23 (2323-211-S5) was used for genome sequencing. We selected 189 individuals from the 15 FS selfing families for genotyping for the SL analysis (**Table S1**).

### Genome sequencing and assembly

Foliar tissue was used for DNA extraction for genome sequencing. Purified nuclear genomic DNA was extracted at BioS&T (http://www.biost.com/, Montreal, Canada) (Birol et al. 2013) and sequenced at the Joint Genome Institute (JGI; Berkeley, USA).

Genome sequencing was executed using three types of libraries: short fragment paired-end, large fragment mate-pair, and linked-reads from large molecules using 10× Genomics Chromium. Depth of *k*-mer coverage profiles were computed for multiple values of *k* using ntCard v1.0.1 (Mohamadi et al. 2017) (**Figure S8**). The largest value of *k* providing a *k*-mer coverage of at least 15 was selected, based on an estimated coverage of > 99.9%, yielding *k* = 128 (Lander and Waterman 1988). We analyzed and visualized *k*-mer profiles using GenomeScope v1.0.0 (Vurture et al. 2017). Paired-end reads were assembled using ABySS v2.1.4 (parameters: k=128; kc=3) and scaffolded using the mate-pair reads with ABySS-Scaffold (Jackman et al. 2017) (**Figure S9**). Linked-reads were aligned and misassemblies were identified and corrected with Tigmint v1.1.2 (Jackman et al. 2018). The assembly was scaffolded using the linked-reads with ARCS v1.0.5 (Yeo et al. 2018) (-c 2; -m 4-20000) and ABySS-Scaffold (-n 5-7; -s 5000-20000). Molecule size of the linked read libraries was estimated using ChromeQC v1.0.4 (https://bcgsc.github.io/chromeqc). Detailed DNA extraction, sequencing, and assembly methods can be found in **Supplemental Methods**.

We estimated completeness of the WRC genome assembly and other conifer genome assemblies using BUSCO v5.0.0 in genome mode, on OrthoDB Embryophyta v10 (Simão et al. 2015; Waterhouse et al. 2018; Kriventseva et al. 2019), which determines the proportion and completeness of single-copy genes from the Embryophtya database (1,614 models) present in the genome.

### Genome annotation

For PacBio Iso-Seq, full-length cDNAs were synthesized from total RNA. We then generated transcript assemblies from 1.4B 2×150 and 50M 2×100 stranded paired-end Illumina RNA-seq reads using PERTRAN (Shengqiang et al. 2013), 18M PacBio Iso-Seq Circular Consensus Sequences (CCS), and previous RNA-seq assemblies (Shalev et al. 2018) (<u>NCBI PRJNA704616</u>). We determined gene loci by transcript assembly alignments and EXONERATE v2.4.0 (Slater and Birney 2005) alignments of proteins from *Arabidopsis thaliana*, *Glycine max*, *Populus trichocarpa*, *Oryza sativa*, *Vitis vinifera*, *Aquilegia coerulea*, *Solanum lycopersicum*, *Amborella trichopoda*, *Physcomitrella patens*, *Selaginella moellendorffii*, *Sphagnum magellanicum*, UniProt Pinales and Cupressales, and Swiss-Prot proteomes to the repeat-soft-masked WRC genome using RepeatMasker v4.0.8 (Smit et al. 2015) with up to 2 kbp extension on both ends. Gene models were predicted using FGENESH+ v3.1.1 (Salamov and Solovyev 2000), FGENESH_EST v2.6, and EXONERATE and PASA (Haas et al. 2003) assembly ORFs. The best-scored predictions for each locus were selected and improved by PASA, adding untranslated regions (UTRs), splicing correction, and alternative transcripts. All software was run using default parameters. Detailed RNA extraction, sequencing, and genome annotation methods can be found in **Supplemental Methods**.

We estimated completeness of the WRC primary transcript gene set and other conifer gene sets using BUSCO v5.0.0 in protein mode on OrthoDB Embryophyta v10. We also assessed coverage of 59 complete WRC sequences found on NCBI and 33 WRC terpene synthase sequences (Shalev et al. 2018) . Sequences were searched against the genome using BLAST+ v2.10.0 (Altschul et al. 1990; Camacho et al. 2009), and presence or absence analyzed using EXONERATE. SCOs were identified using OrthoFinder v2.5.4 (Emms and Kelly 2019), isolating genes identified in one copy in the WRC gene set when compared against the giant sequoia gene set (Scott et al. 2020).

### SNP genotyping

DNA was isolated from lyophilized tissue with a modified protocol of Xin and Chen (2012). Targeted sequencing-based genotyping was done by Capture-Seq methodology at Rapid Genomics (Gainesville FL, USA). Initially, probes were designed using only limited publicly transcriptome data and database matches for functionally characterized conifer genes, an approach that has worked previously for other organisms (e.g., Mukrimin et al. 2018; Vidalis et al. 2018; Acosta et al. 2019; Telfer et al. 2019); however, this approach yielded less than 2,000 polymorphic sites. Thus, we developed a specialized probe design approach targeting regions of putative high variability, specifically: previously identified differentially expressed regions from cold-tolerance, deer browse, wood durability, leaf blight, and growth trials, database matches for functionally characterized conifer genes, and whole transcriptome and genome data (NCBI Umbrella BioProject <u>PRJNA704616</u>). A set of 57,630 probes, 37,294 targeting genic regions and 19,706 targeting intergenic regions, was designed for marker discovery, from which a panel of 20,858 probes was selected for genotyping.

Putative SNPs were identified using FreeBayes v1.2.0 (Garrison and Marth 2012) in 150bp on either side of the probes and filtered probes that had more than 17 SNPs per 420 bp target region to prevent over-capture. Sequencing depth was used to select the final set, removing probes with low and high sequencing depth for Capture-Seq on the remainder of the samples. Detailed methods for SNP genotyping can be found in **Supplemental Methods**.

### SNP filtering and annotation

Variant sites were filtered using VCFtools v0.1.17 (Danecek et al. 2011) with the following flags: --max-missing 0.95; --minQ 30; --min-meanDP 15; --max-meanDP 60. SNPs with an allele balance > 0.2 and < 0.8 or < 0.01 were retained to eliminate incorrectly called heterozygotes using vcffilter in vcflib v1.0.1 (Garrison 2016). To eliminate paralogous loci, we excluded: SNPs with a read-ratio deviation score *D* (McKinney et al. 2017) > 5 and < -5, SNPs with a heterozygosity greater than 0.55, and SNPs with excess heterozygosity and deviations from HWE in the RWP at a *p*-value cutoff of 0.05 and 1e-5, respectively. We also excluded SNPs with negative inbreeding coefficients (*F*_IS_), using the formula:

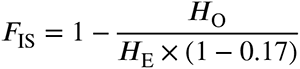

where *H*_O_ is the observed heterozygosity of the locus, *H*_E_ is the expected heterozygosity of the locus under HWE, and the factor of (1 – 0.17) accounts for the expected equilibrium fixation index of 0.17 in WRC based on the average outcrossing rate of 0.7 (El-Kassaby et al. 1994; O’Connell et al. 2001, 2004). Invariant sites were filtered using the following flags: --max-missing 0.95; --min-meanDP 15; --max-meanDP 60. Variant effect prediction was carried out using the Ensembl Variant Effect Predictor r103; one effect per SNP was selected, and for compound effects, only the most severe consequence was retained (McLaren et al. 2016). Relationships between trees were estimated by generating a genomic realized relationship matrix for all individuals using the ‘A.mat’ function of rrBLUP v4.6.1 in R (Endelman 2011; R Core Team 2021). For the RWP, five trees with a relatedness coefficient > 0.2 were excluded from analyses for a total of *n* = 112 trees. For the SLs, nine individuals whose relationships did not match the *a priori* pedigree were removed from analysis (**Table S1**).

### Linkage disequilibrium

Pairwise LD (*r*^2^) was estimated in the RWP using PLINK v1.9 (Chang et al. 2015). LD was calculated for all scaffolds containing at least two SNPs (--r2; --ld-window-r2 0; --ld-window-kb 999999; --ld-window 999999; --maf 0.05). Under drift-recombination equilibrium, our expectation of LD decay over distance will be:

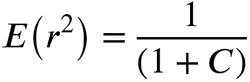

Where *C* is the product of the population recombination parameter ρ = 4*N_e_r* and the distance in bp, *N_e_* is the effective population size and *r* is the recombination rate per bp (Sved 1971). Adjusting for population size *n* and a low level of mutation, decay of LD was estimated as a factor of *n*, and *C* (Hill and Weir 1988; Remington et al. 2001; Marroni et al. 2011; Fahrenkrog et al. 2017).

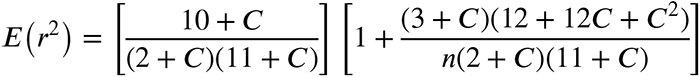

*C* was estimated by non-linear regression, using the nls function in R.

### Population structure and genetic differentiation

For STRUCTURE (Pritchard et al. 2000; Falush et al. 2007) analysis, SNPs were further filtered to include only SNPs with an *r*^2^ threshold of 0.1 using a 2.17 Mb window, following the outcome of our LD analysis, a MAC threshold of 3 was used to remove singletons, and only SCO and intergenic SNPs were retained (*n* = 4,765). For all other genetic differentiation analyses, all SCO and intergenic SNPs were used (*n* = 13,427). Population structure of the RWP was estimated using STRUCTURE v2.3.4 and a DAPC using the adegenet package (v2.1.1) in R (Jombart 2008; Jombart et al. 2010; Jombart and Ahmed 2011). A hierarchical *F*_ST_ test as implemented in the hierfstat package (v0.04-22) in R (Goudet 2005) was used to assess genetic differentiation between subpopulations in the RWP and a Mantel test was executed using mantel.randtest for 9,999 permutations in ade4 v1.7-15 in R (Dray and Dufour 2007) to assess isolation by distance. PCA was performed on the genotype matrix of each subpopulation to visualise genetic distance between individuals using ade4.

STRUCTURE software was run using 10,000 MCMC repetitions with 10,000 repetitions of burn-in and 10 iterations of each K. Results were analyzed using methods of Evanno et al. (2005) and Puechmaille (2016) of cluster and admixture estimation and selection; fastStructure v1.0 (Raj et al. 2014) was also used with 10-fold cross-validation. For DAPC, find.clusters was used to select the optimal number of clusters based on Bayesian Information Criterion (BIC), and 10-fold cross-validation with 1,000 replicates was performed with xvaldapc to select the number of PCs and discriminant functions to retain.

### Nucleotide diversity

To avoid downward bias due to missing data, SCO and intergenic variant sites (*n* = 13,427) and all invariant sites were used to estimate π and *d*_XY_ for *n* = 10,631 SCO genes and 10 kb windows in the RWP using pixy v1.2.7.beta1 (Korunes and Samuk 2021). Zero- and four-fold degenerate variant and invariant sites were identified using the NewAnnotateRef.py script (Williamson et al. 2014), and π_0_/π_4_ was calculated over all SCO genes. SFS was estimated using easySFS (https://github.com/isaacovercast/easySFS).

### Effective population size (N_e_)

We estimated *N_e_* using the LD model estimation method under random mating as implemented in NeEstimator v2.1 (Do et al. 2014). This method uses background LD shared among samples to estimate *N_e_*; thus, SNPs with as little LD as possible are required (Waples and Do 2008; Gilbert and Whitlock 2015). Due to the high LD observed in WRC, we first generated putative linkage groups (LGs) for the WRC genome by aligning all genomic scaffolds containing SNPs to the giant sequoia genome (Scott et al. 2020) using BLAST+. Scaffolds were assigned to their most likely LG based on bitscore. We then used the nucmer command from MUMmer v4 (Marçais et al. 2018) to determine the most likely alignment region for each scaffold in each LG. We retained SNPs estimated to be at least 2.17 Mbp apart. A MAF threshold of 0.05 was established to eliminate bias that may be introduced by rare alleles, and a 95% nonparametric JackKnife confidence interval was taken for the estimated value, as recommended by Waples and Do (2008) and Gilbert and Whitlock (2015).

Stairway Plot 2 (Liu and Fu 2020) was used on the folded SFS from intergenic and 4-fold degenerate positions to further assess *N_e_* changes over time. We used the following parameters: nseq = 222; L = 238,557; pct_training = 0.67; nrand = 55, 110, 165, 220; ninput = 200; mu = 3.74e-9; year_per_generation = 50.

### Genotype correction for SLs

Genotype correction in continuous SLs used two criteria: Individuals with homozygous calls in at least two consecutive generations were considered to be homozygous for that allele in all subsequent generations; and individuals with heterozygous calls in at least two consecutive generations were considered to be heterozygous for all preceding generations, up to and including the FS generation. SNPs that could not be corrected following these criteria were removed. We manually corrected genotypes for SLs which had been completely genotyped from either S1 – S4 or S1 – S5 (**Table S1**). For seven SLs in which only the S3 generation had not been sequenced, we imputed the genotypes for the S3 generation at each locus for SNPs where no other genotype was possible, and marked the rest as missing.

### Change in heterozygosity and inbreeding coefficients over time

Corrected genotypes for all filtered SNPs (*n* = 18,371) were used to calculate the observed and expected changes in heterozygosity and inbreeding coefficients over time in the SLs, and in the RWP. Observed heterozygosity was calculated at each SNP locus for each generation across all SLs using the adegenet package in R. Expected heterozygosities in the S1 – S5 generations were calculated for each SNP locus as half the observed heterozygosity of the previous generation. We calculated inbreeding coefficients in PLINK using the --ibc flag to obtain a measure for inbreeding based on the correlation between uniting gametes (*F*_UNI_). This metric is defined by Yang et al. (2011) for each *i*th SNP and each *j*th individual as

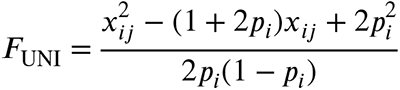

Where *x* is the number of copies of the reference allele and *p* is the population-wide allele frequency at that locus. Calculations of *F*_UNI_ do not consider LD; thus, we used SNPs filtered for LD and MAC (*n* = 6,123). In a diploid population, *F* should increase as 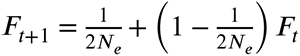 each generation; thus, under complete selfing we expect *F* to increase by a factor of 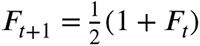. *F* was calculated using corrected genotypes in SLs and uncorrected genotypes in the RWP.

### Isolating SNPs significantly deviating from expectations of drift

To identify loci that diverge from patterns expected under genetic drift, we evaluated all SNPs for which the FS generation was heterozygous and then observed whether the SNP went to fixation or not by the S4 generation in our 28 complete, corrected SLs. The S5 generation was excluded from this analysis due to small sample size. For statistical analysis, each SL was considered an independent replicate. SLs were categorized at each generation as ‘fixed for reference allele’, ‘fixed for alternate allele’, or ‘not fixed’. The observed number of SLs in each category was tabulated for the S4 generation. The expected number of SLs in each category was calculated following the expectation of a 50% reduction in heterozygosity in each generation, resulting in an expectation of 6.25% of the SLs being heterozygous, 46.875% being homozygous for the reference allele, and 46.875% being homozygous for the alternate allele in the S4 generation. A χ^2^ test was performed for SNPs with genotyping data present in at least 3 SLs to test for significant differentiation from this expectation, and a Benjamini-Hochberg false discovery rate of 0.05 was used to correct for multiple hypothesis testing across all SNPs. Variant effects were predicted for significant SNPs, and a Fisher’s Exact Test was used to determine the presence of over or under-representation of significant SNPs and over-representation of GO categories in the significant SNPs.

### Data access

The genome sequence reads, assembly and annotation, and transcriptomes used in annotation generated in this study have been submitted to the NCBI BioProject database (https://www.ncbi.nlm.nih.gov/bioproject/) under accession number <u>PRJNA704616</u>.

The SNP data generated in this study has been submitted to the Zenodo data repository under DOI <u>10.5281/zenodo.6562381</u>, and is available in the **Supplemental Datasets**.

**Supplemental Code** and raw data used for generating data files and figures, including all filtered SNP sets for each step of the study, are available as **Supplemental Code** files, and have been uploaded together with copies of the genome annotation and **Supplemental Dataset** files to the following GitHub repository: https://github.com/tshalev/WRC-genome-paper. Summaries of **Supplemental Code** are available in the **Supplemental Information.**

## Acknowledgements

The research was supported with funds from Genome Canada and Genome British Columbia (CEDaR GAPP [184CED] to JB, ADY, JHR; 281ANV to IB); the Natural Sciences and Engineering Research Council of Canada (to JB, TJS, IB, OG); and the Province of British Columbia Land-Based Investment Strategy Tree Improvement Program to the western redcedar breeding program (JHR, ADY, LvdM). The work conducted by the U.S. Department of Energy Joint Genome Institute is supported by the Office of Science of the U.S. Department of Energy under Contract No. DE-AC02-05CH11231. We thank Allyson Miscampbell for SL DNA extraction, Angela Chiang for total RNA extraction, Lori Handley at HudsonAlpha for genomic data quality control and the HudsonAlpha Genomic Services Laboratory for the 10x library preparation, Eugene Goltsman at the JGI for assistance with HipMer, Kermit Ritland and Michael Whitlock for their careful help with the analyses and review of the paper, Annette Fahrenkrog for population genomics data analysis support.

## Author Contributions

TJS conceived the project, performed research, analyzed data, and wrote the manuscript. SS and SDJ contributed to data analysis and writing of the manuscript. OG, MMSY, RLW and LC contributed to data analysis. AS, LBB, CP, JJ, GH, JY, MY, JGu and JWB generated materials and data. LvdM contributed to study design and provided essential materials. LGN and JGr contributed to study design and generated materials and data. LHR contributed to interpretation of the results and writing of the manuscript. JS, IB, MK and ADY contributed to study design and interpretation of the results. CR contributed to study design, generated materials and data, contributed to interpretation of the results, and managed and coordinated the overall project. JHR conceived the project and provided essential materials. JB conceived the project, managed and coordinated the overall project, and wrote the manuscript. All authors reviewed and edited the manuscript.

## Competing Interest Statement

The authors declare no competing interests.

## Notes

https://www.ncbi.nlm.nih.gov/bioproject/PRJNA684624

https://zenodo.org/record/6562381

https://github.com/tshalev/WRC-genome-paper

### Competing Interest Statement

The authors have declared no competing interest.

