## Supplemental Information for "The western redcedar genome reveals low genetic diversity in a self-compatible conifer"

**This PDF file includes:**

Supplemental Methods
Figures S1 to S9
Tables S1 to S15
Summaries for Datasets S1 to S10
Summaries for Code S1 to S8
SI References

**Other supplementary materials for this manuscript include the following:**

Datasets S1 to S10
Code S1 to S8

### Supplemental Methods

#### *DNA extraction for genome sequencing and SNP genotyping*

Foliar tissue was used for DNA extraction for genome sequencing and genotyping. For genome sequencing, purified nuclear genomic DNA was extracted at BioS&T (http://www.biost.com/, Montreal, Canada) (Birol et al. 2013) and sequenced at the Joint Genome Institute (JGI; Berkeley, USA). For SNP genotyping, DNA was isolated from lyophilized tissue with a modified protocol of Xin and Chen (Xin and Chen 2012). DNA extraction protocol modifications included: the addition of 10% polyvinylpyrrolidone to the CTAB extraction buffer, all centrifugation steps were performed for 10-12 min at 10,000 rpm, the amount of Qiagen MagAttract solution was increased to 7µl, one-minute wait time for binding on the magnetic block was added, and 115µl TE buffer was used for DNA dilution. The concentration and quality of DNA was verified with a Nanodrop 2000c (Fisher Scientific, Toronto, ON, Canada), Quantiflor (Promega Corporate, Madison, WI, USA) and 0.8% agarose gel to cross check for the quantity and quality of the DNA.

#### *Genome sequencing and assembly*

Genome sequencing was executed using three types of libraries: short fragment paired-end, large fragment mate-pair, and linked-reads from large molecules using 10× Genomics Chromium. Illumina paired-end adapters were trimmed using Trimadap v0.1r11 (https://github.com/lh3/trimadap). Mate-pair sequencing adapters were removed using NxTrim v0.4.3 (O'Connell et al. 2015). For 10× Genomics libraries, adapters were trimmed using Long Ranger Basic v2.1.6 (https://github.com/10XGenomics/longranger). Depth of *k*-mer coverage profiles were computed for multiple values of *k* using ntCard v1.0.1 (Mohamadi et al. 2017) (**Figure S8**). The largest value of *k* providing a *k*-mer coverage of at least 15 was selected, based on an estimated coverage of > 99.9%, yielding $k$ = 128 (Lander and Waterman 1988). We analyzed and visualized *k*-mer profiles using GenomeScope v1.0.0 (Vurture et al. 2017). Paired-end reads were assembled using ABySS v2.1.4 (parameters: k=128; kc=3) and scaffolded using the mate-pair reads with ABySS-Scaffold (Jackman et al. 2017) (**Figure S9**). Linked-reads were aligned and misassemblies were identified and corrected with Tigmint v1.1.2 (Jackman et al. 2018). The assembly was scaffolded using the linked-reads with ARCS v1.0.5 (Yeo et al. 2018) (-c 2; -m 4-20000) and ABySS-Scaffold (-n 5-7; -s 5000-20000). Molecule size of the linked read libraries was estimated using ChromeQC v1.0.4 (https://bcgsc.github.io/chromeqc). Assembled scaffolds were screened against bacterial proteins, organelle (chloroplast and mitochondria) sequences, and adapters on submission to NCBI and contaminants were removed.

***Genome annotation***

*RNA extraction*

Young and mature foliage, xylem and bark samples from a variety of samples (NCBI Umbrella BioProject NCBI PRJNA704616) were ground to a fine powder and total RNA extracted using Purelink ®Plant RNAReagent (Life Technologies, Carlsbad, CA, USA). RNA concentration was determined using a NanoDrop 1000 (Thermo Scientific, Waltham, MA, USA). RNA integrity (RIN) was assessed using Bioanalyzer 2100 RNA Nano chip assays (Agilent, Santa Clara, CA, USA). The minimum RIN was 8.

*Transcriptome sequencing and gene space annotation*

For PacBio Iso-Seq, full-length cDNAs were synthesized from total RNA using template-switching technology with SMARTer PCR cDNA Synthesis kit (Clontech, CA, USA). First-strand cDNA was amplified with PrimeSTAR GL DNA Polymerase (Clontech) using template-switching oligos to produce double-stranded cDNA, which was purified with AMPure PB beads and either non-size selected or selected for 2-10 kb cutoff by BluePippin (Sage Science, MA, USA). Amplified cDNA was end-repaired and ligated with blunt end PacBio sequencing adaptors using SMRTbell Template Prep Kit 1.0 (Pacific Biosciences, CA, USA), exonuclease treated to remove unligated products, and purified using AMPure PB beads. PacBio Sequencing primer was annealed to the SMRTbell template library and sequencing polymerase was bound using Sequel II Binding kit 1.0. SMRTbell template libraries were sequenced on a Pacific Biosciences' Sequel II sequencer using v4 sequencing primer, 8M v1 SMRT cells, and v1.0 sequencing chemistry with 1×1440 (24-hour) sequencing run times. For Illumina RNA-seq libraries, sample prep was performed on the PerkinElmer Sciclone NGS robotic liquid handling system using Illumina TruSeq Stranded mRNA HT sample prep kit utilizing poly-A mRNA selection (https://support.illumina.com/sequencing/sequencing_kits/truseq-stranded-mrna.html). Libraries were quantified using KAPA Biosystem's next-generation sequencing library qPCR kit and run on a Roche LightCycler 480 real-time PCR instrument. Libraries were multiplexed and the pool of libraries was prepared for sequencing on the Illumina NovaSeq 6000 sequencing platform using NovaSeq XP v1 reagent kits, S4 flow cell, following a 2×150 indexed run recipe.

Gene space annotation was carried out using transcript assemblies from 1) 1.4B 2×150 and 50M 2×100 stranded paired-end Illumina RNA-seq reads using PERTRAN (Shengqiang et al. 2013); 2) 18M PacBio Iso-Seq Circular Consensus Sequences (CCS) which were corrected and collapsed by genome guided correction pipeline to obtain 530K putative full-length transcripts; and 3) previous RNA-seq assemblies (Shalev et al. 2018). PASA v2.0.2 (Haas et al. 2003) was used to assemble 375,524 transcripts from these three sources. Briefly, PERTRAN executes genome-guided transcriptome short-read (Illumina) assembly via GSNAP (Wu and Nacu 2010) and builds splice alignment graphs after alignment validation, realignment and correction; Iso-Seq CCS correction and clustering aligns CCS reads to the genome with GMAP (Wu and Nacu 2010) while performing intron correction for small indels in splice junctions, if present, and clusters alignments when all introns are the same or have at least 95% overlap for single exon. Gene loci were determined by transcript assembly alignments and EXONERATE v2.4.0 (Slater and Birney 2005) alignments of proteins from *Arabidopsis thaliana*, *Glycine max*, *Populus trichocarpa*, *Oryza sativa*, *Vitis vinifera*, *Aquilegia coerulea*, *Solanum lycopersicum*, *Amborella trichopoda*, *Physcomitrella patens*, *Selaginella moellendorffii*, *Sphagnum magellanicum*, UniProt Pinales and Cupressales, and Swiss-Prot proteomes to the repeat-soft-masked WRC genome using RepeatMasker v4.0.8 (Smit et al. 2015) with up to 2 kbp extension on both ends unless extending into another locus on the same strand. For EXONERATE alignments, an intron size cut-off of 90 kb was used. The repeat library consisted of *de novo* repeats identified by RepeatModeler v1.0.11 (Smit and Hubley 2015) from the WRC genome and repeats in RepBase. Gene models were predicted by homology-based predictors, FGENESH+ v3.1.1 (Salamov and Solovyev 2000), FGENESH_EST v2.6, and EXONERATE and PASA assembled ORFs (in-house homology constrained ORF finder). The best-scored predictions for each locus were selected using multiple positive factors, including transcriptome, EST and protein support, and one negative factor; overlap with repeats. The selected gene predictions were improved by PASA, adding untranslated regions (UTRs), splicing correction, and alternative transcripts. PASA-improved gene models were subjected to protein homology analysis to the proteomes mentioned above to obtain Cscore (BLASTP score ratio to mutual best hit BLASTP score) and protein coverage (highest percentage of protein aligned to the best homologs). PASA-improved transcripts were selected based on the Cscore, protein coverage, transcriptome and EST coverage, and the overlap of coding sequence (CDS) with repeats. Transcripts were selected with Cscore ≥ 0.5 and protein coverage ≥ 0.5, or if they had transcriptome or EST coverage, but their CDS overlap with repeats was less than 20%. For gene models with CDS overlap with repeats of more than 20%, Cscore was required to be at least 0.9 and homology coverage at least 70% to be selected. Selected gene models were subjected to Pfam analysis. Gene models > 30% of the protein in Pfam TE domains were removed, along with weak gene models (HMMER e-value < $10^{-5}$). Incomplete gene models, low homology support without fully transcriptome supported gene models and short single exons (< 300 bp CDS) without protein domains nor strong expression gene models were manually removed. All software was run using default parameters.

#### *SNP genotyping*

Targeted sequencing-based genotyping was done by Capture-Seq methodology at Rapid Genomics (Gainesville FL, USA). A set of 57,630 probes was designed for initial marker discovery, from which a panel of 20,858 probes was selected for genotyping. A set of transcriptomes (Shalev et al. 2018) (PRJNA704616) was aligned to the reference genome to identify SNPs.

Candidate probes (120 nt) were initially designed *in silico* and were selected by removing candidates with poor base composition for hybridization (GC content <0.2 and >0.6, high G content >0.2 and long homopolymers >7), followed by removing probes aligning to more than one position on the reference genome (≥90% identity and length). A final set of 57,630 probes was selected, representing 14,517 scaffolds (average 3.9 probes/scaffold), with 37,294 targeting at least one SNP and 19,706 mapping to intergenic regions not containing pre-identified SNPs.

A set of 128 individuals were selected to validate the probe panel and associated polymorphisms. Genomic DNA (0.5 μg) was fragmented (mean size 300 bp), followed by repair of ends, phosphorylation, adenylation, ligation of Illumina compatible adapters containing 8bp indexes and 5' T-overhang, and 10 cycles PCR amplification with universal primers to produce sequencing-ready libraries. Libraries were quantified using PicoGreen. Libraries from 16 samples were pooled, hybridized to the 120 nt RNA probes following Agilent's SureSelect Target Enrichment System (Agilent Technologies) and sequenced on an Illumina HiSeq X machine with paired-end 150 bp cycle for an average sequencing depth per sample of 15×. Sequence data were aligned to the reference genome with BWA-MEM v0.7.17 (Li 2013) and sets of four samples were combined to increase sequencing depth for identifying markers.

Putative SNPs were identified using FreeBayes v1.2.0 (Garrison and Marth 2012) in 150bp on either side of the 57,630 probes and filtered probes that had more than 17 SNPs per 420 bp target region (150 bp + 120 bp + 150 bp) to avoid over-capture. The sequencing depth of the probes was used to select the final set of 20,858 probes, removing probes on both sides of the distribution (low and high sequencing depth), for Capture-Seq on the remainder of the samples.

SNP discovery was carried out for a large group of 4,832 trees, which included our range-wide population (RWP), selfing lines (SLs), and training and target populations for genomic selection; the latter two were not included in analyses for this study.

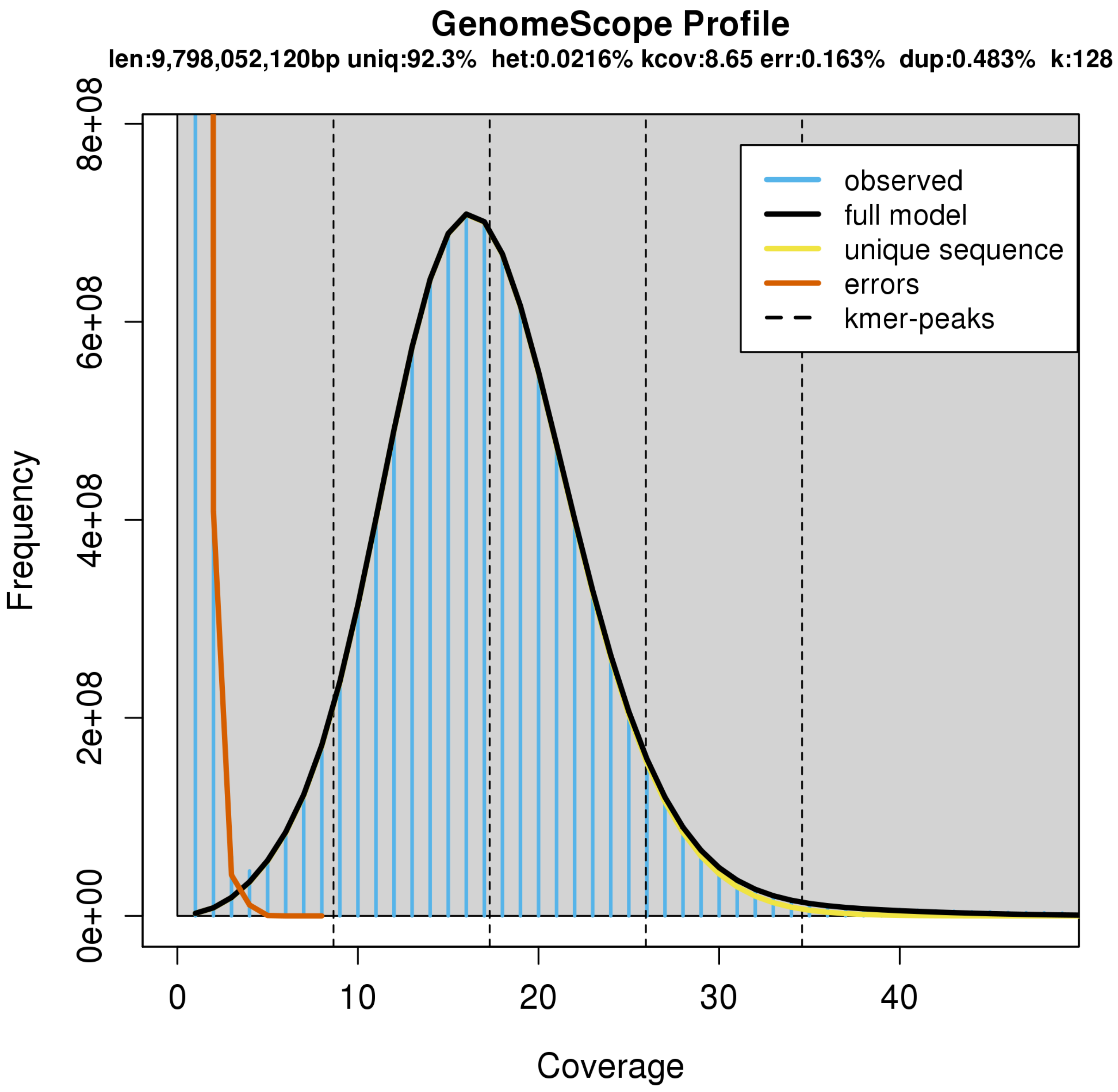

**Figure S1. GenomeScope** (Vurture et al. 2017) **estimation of western redcedar (WRC) genome size.** Genomescope estimated the size of the WRC genome at 9.8 Gbp.

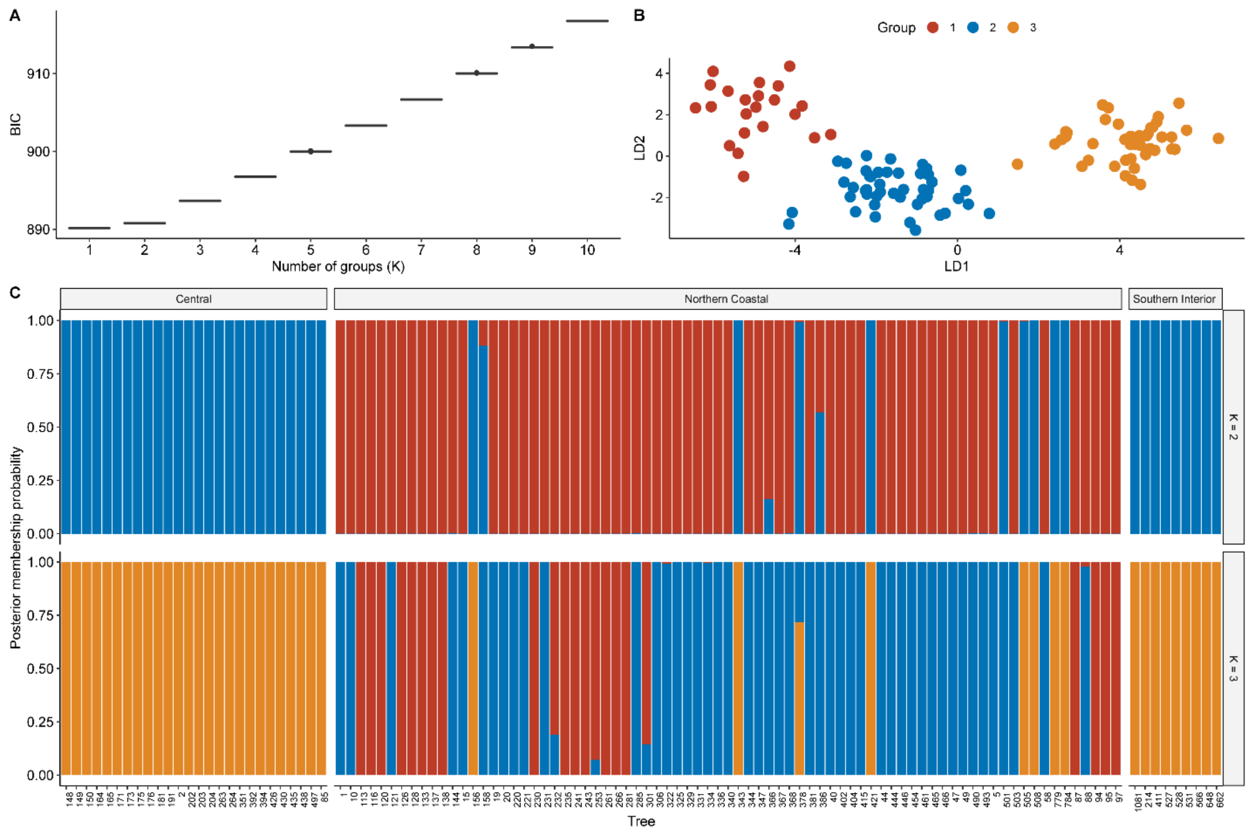

**Figure S2. Discriminant Analysis of Principal Components (DAPC) assessment of optimal K for trees in the range-wide population (RWP) (*n* = 112). A) Selection of optimal K using the find.clusters approach.** Optimal K is typically determined by selecting the K with lowest Bayesian Information Criterion (BIC) score. However, when the lowest BIC is assigned to K = 1, this is interpreted as an inability to resolve clusters (Miller et al. 2020). **B) DAPC clustering using the first 22 principal components and K = 3. C) Composition plots of the reassignment of individual trees to clusters based on DAPC for K = 2-3.** DAPC was not able to properly reassign trees from the RWP, placing all trees from the Central and Southern-Interior subpopulations together while showing complete mixing in the Northern-Coastal subpopulation.

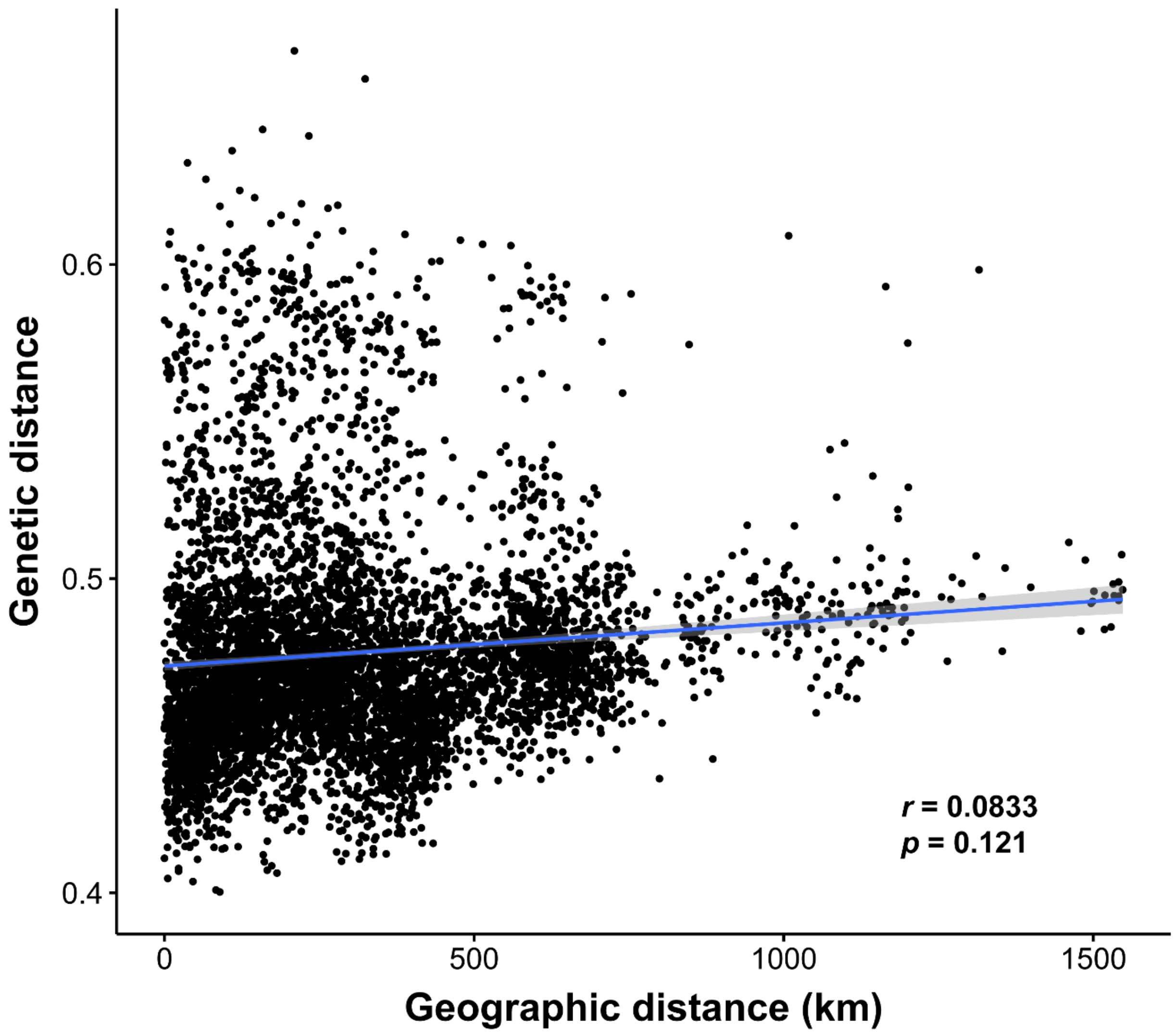

**Figure S3. Individual genetic distance as a factor of geographic distance in the range-wide population ($n$ = 112)**. No significant association was observed between geographic distance and genetic distance when grouping trees based on geographic origin. Surprisingly, no significant association was observed between individual trees either (as shown in the figure), which would normally be expected in continuous populations expanding their range from one source.

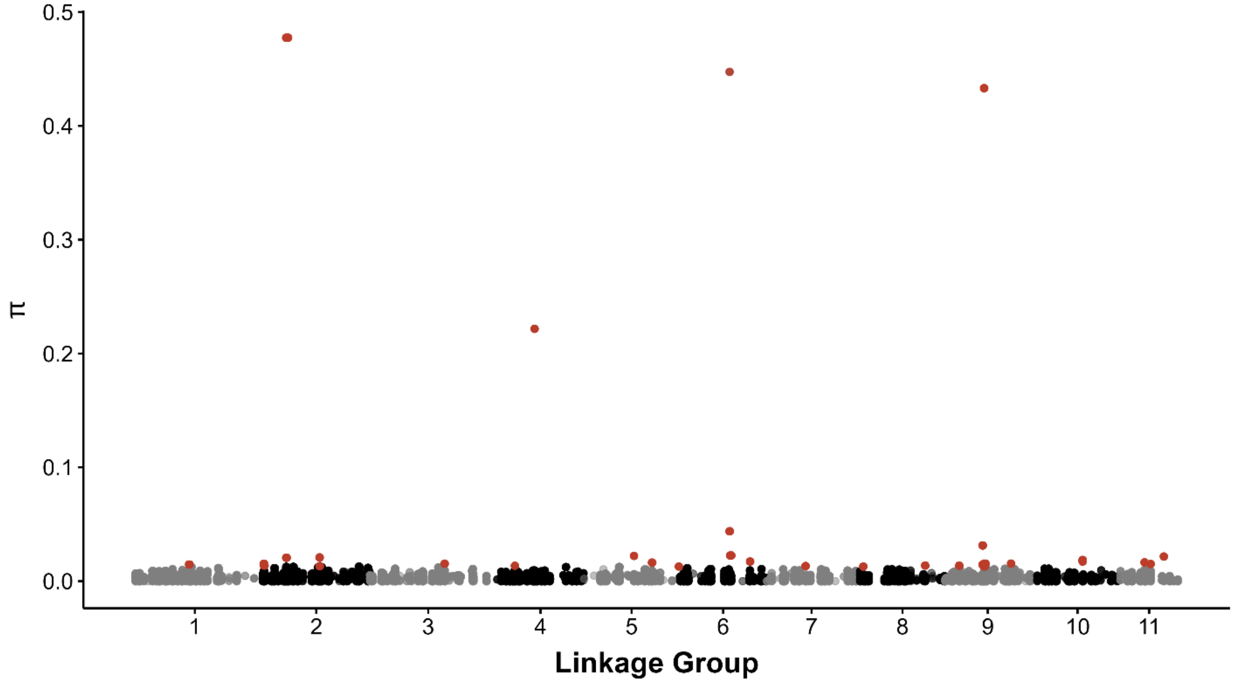

**Figure S4. Nucleotide diversity (π) in the range-wide population (*n* = 112).** Average π in Single Copy Ortholog (SCO) genes across each putative linkage group (see **Methods and Materials**). The top 1% of SCOs with highest π, which account for 3% of total nucleotide diversity, are shown in red (*n* = 47; **Table S10**).

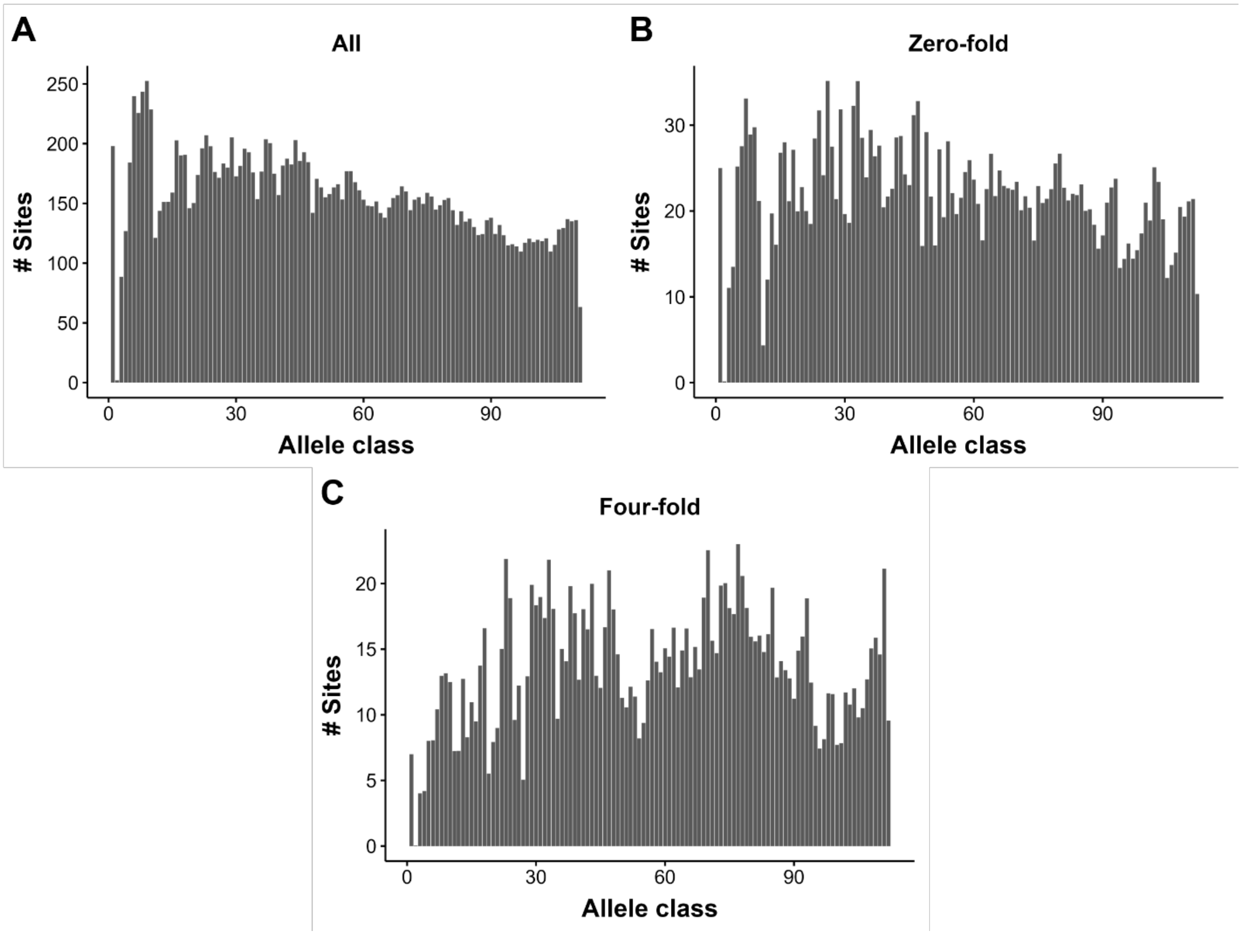

**Figure S5. Folded Site Frequency Spectrum (SFS) plots for A)** All SNPs in the range-wide population (*n* = 18,371); **B)** All zero-fold degenerate SNPs (*n* = 2,674); and **C)** All four-fold degenerate SNPs (*n* = 1,651s).

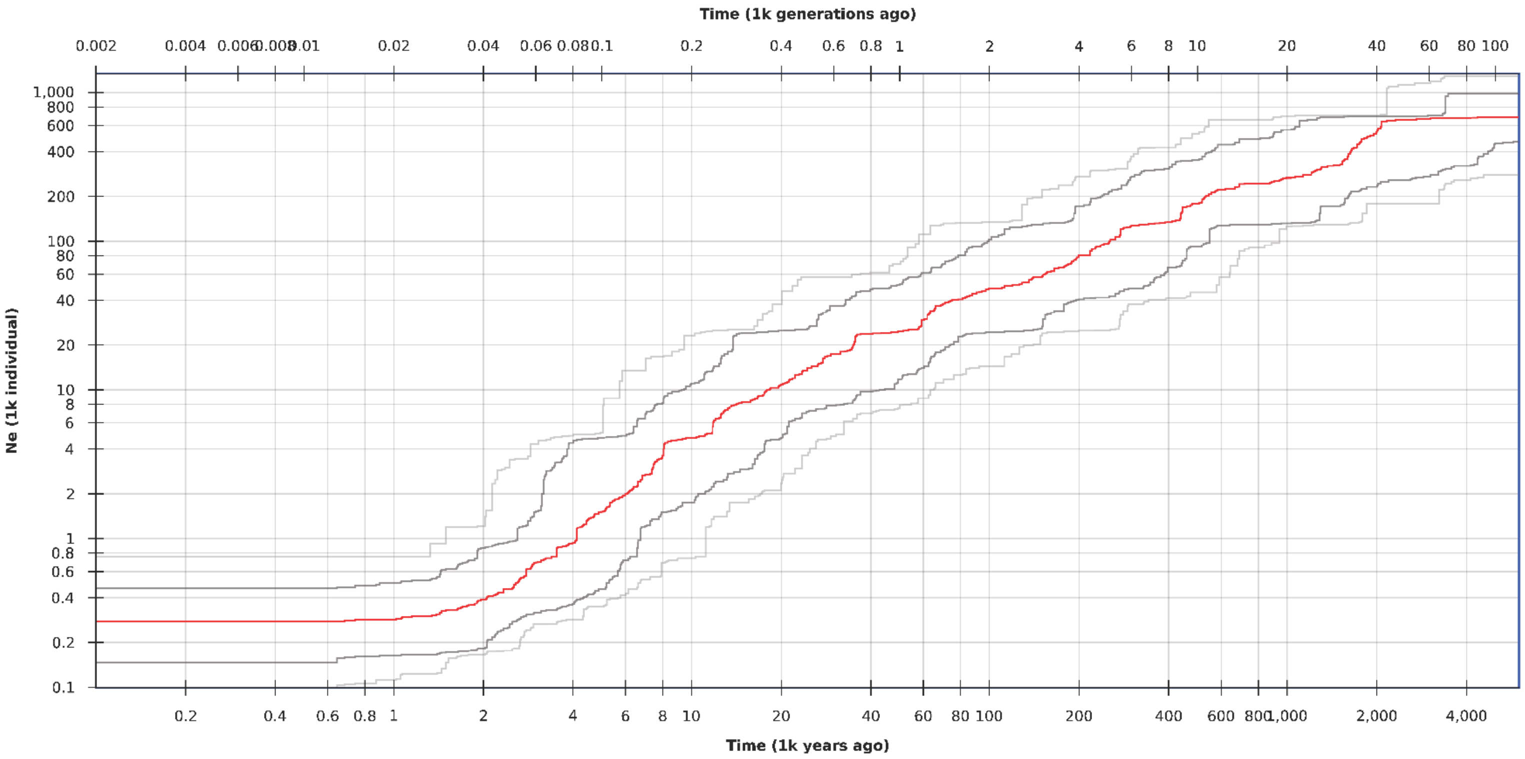

**Figure S6. Stairway Plot (Liu and Fu 2020) on folded SFS of intergenic and four-fold degenerate SNPs in the range-wide population.** $N_e$ appears to enter a period of decline starting ca. 2 MYA. At the peak of the last glacial period ca. 30 KYA $N_e$ had declined to ~ 20,000 individuals, and continued to drop up until it reached its current level of ~ 300 individuals ca. 2 KYA.

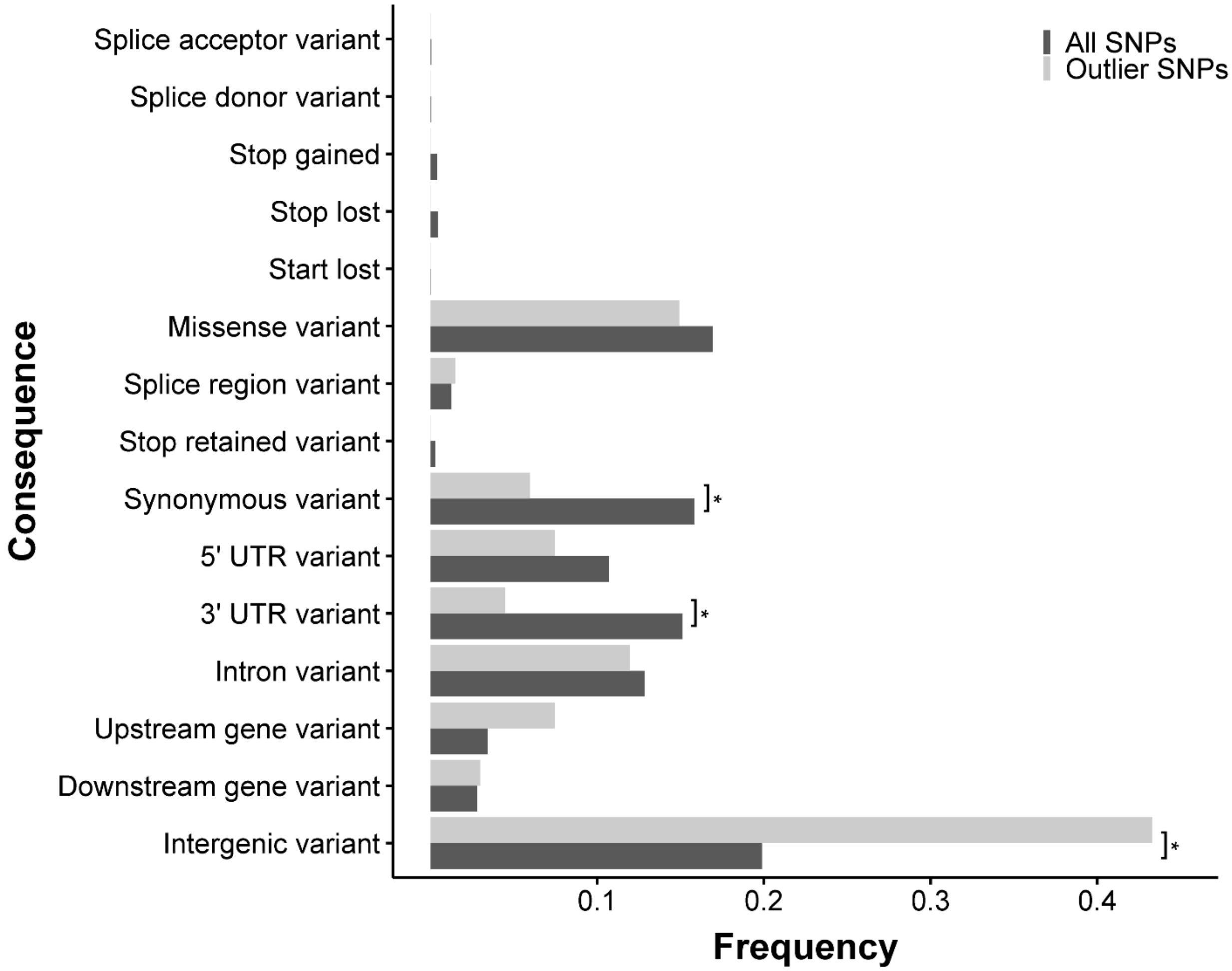

**Figure S7. Consequences of SNPs deviating from expectations of genetic drift.** Comparison of the consequences of all SNPs with effects predicted by Variant Effect Predictor (VEP) (McLaren et al. 2016) ($n$ = 17,728) and SNPs deviating from expectations of genetic drift with predicted effects (outlier SNPs; $n$ = 83). "*" indicates a significant difference between the two sets of SNPs for proportion of SNPs for a certain consequence (Fisher's Exact Test, $p < 0.05$).

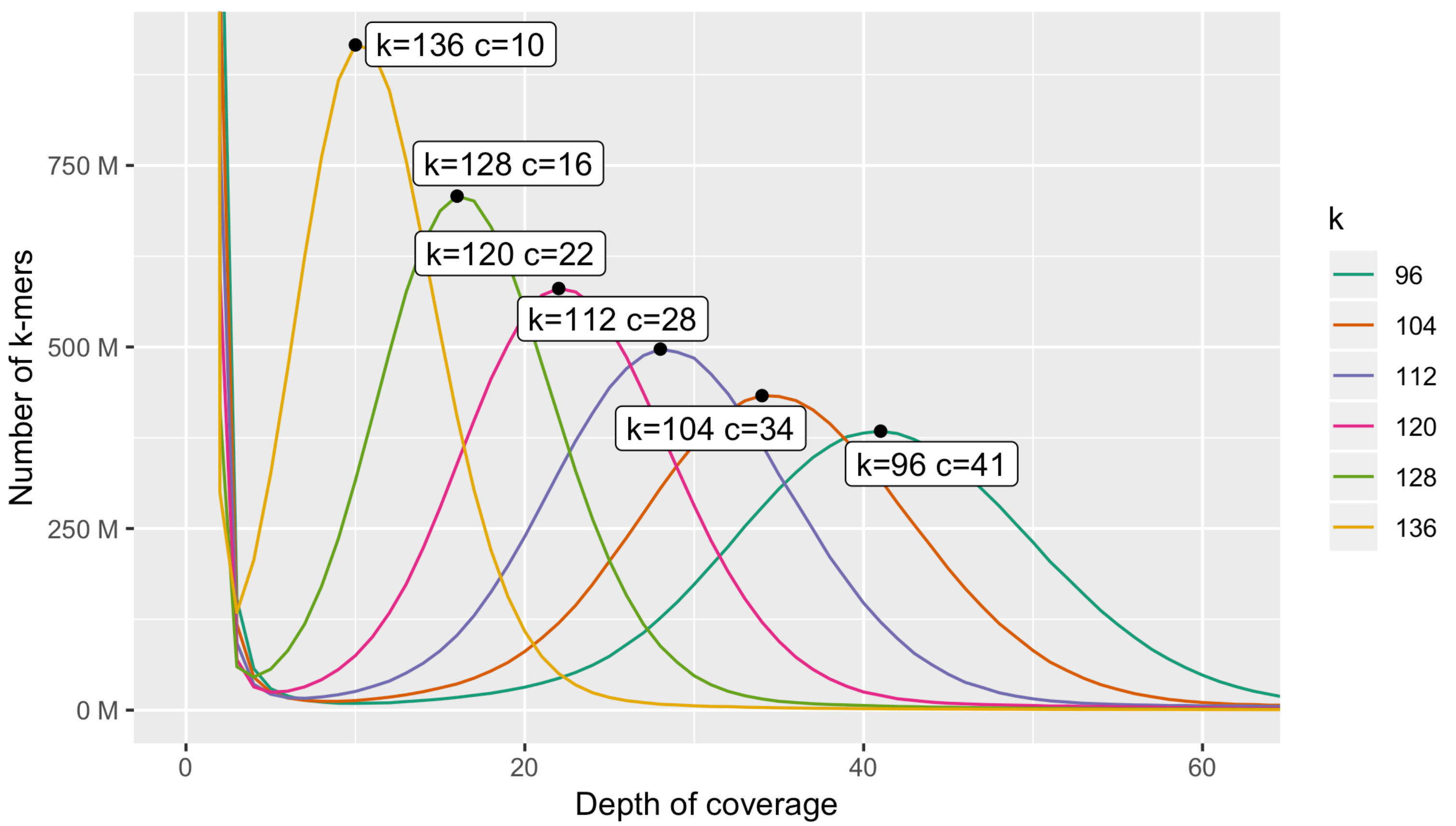

**Figure S8. Depth of *k*-mer coverage profiles for multiple values of *k* estimated in ntCard (Mohamadi et al. 2017).** Mode of depth of coverage *c* is labeled for each value of *k*.

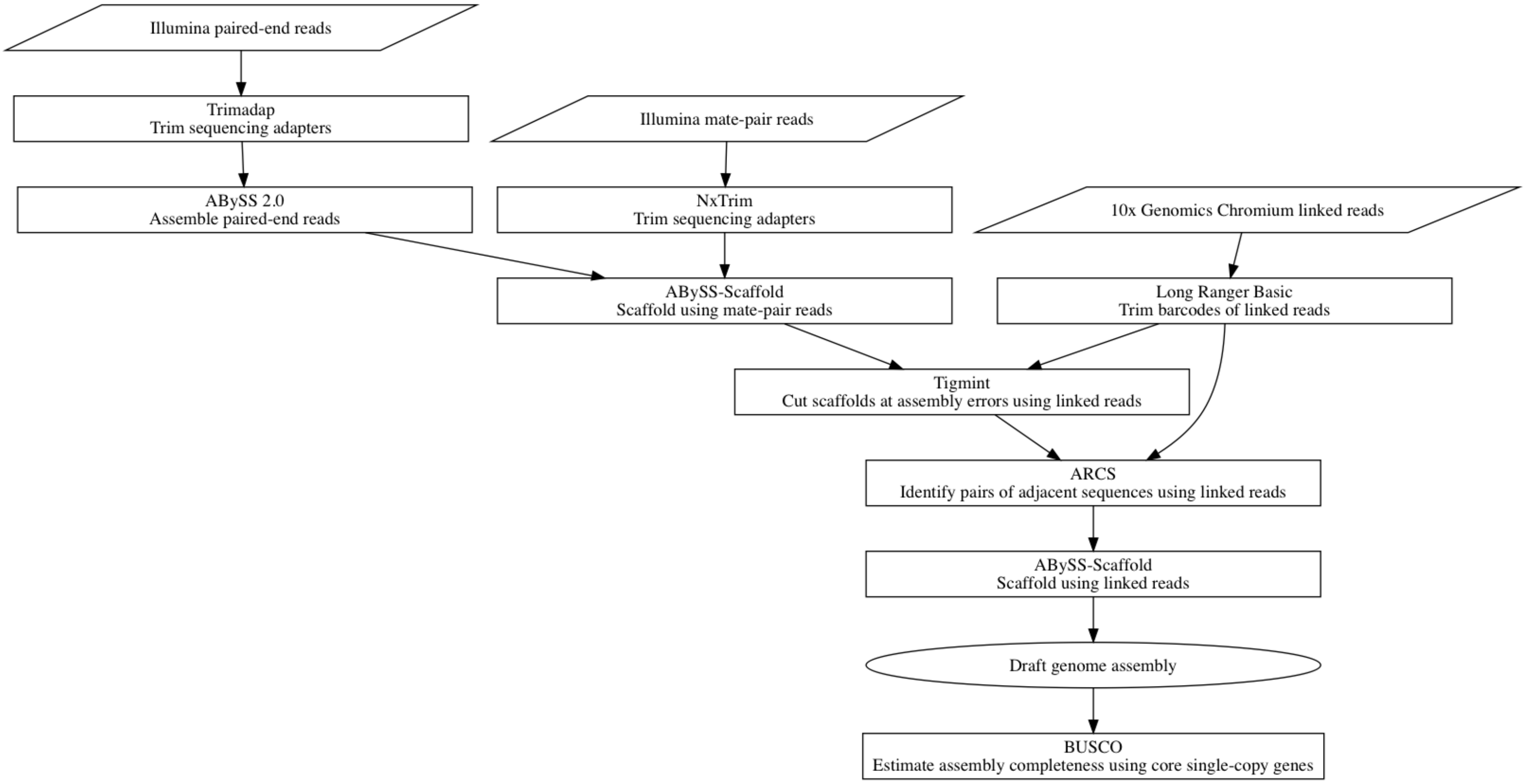

**Figure S9. Flowchart of the assembly process of the WRC draft genome.**

**Table S1. Overview of selfing lines (SLs) selected for the study**

| | Selfing Line | FS | S1 | S2 | S3 | S4 | S5 |
|---|---|---|---|---|---|---|---|
| *** | 1 | 1_1 | 111 | 111-2 | 111-22 | 111-221 | 111-221-S5 |
| | 1 | 1_2 | 122 | 122-1 | 122-12 | 122-121 | |
| | 1 | 1_4 | 145 | 145-5 | 145-54 | 145-545 | 145-545-S5 |
| *** | 6 | 6_1 | 611 | 611-1 | 611-11 | 611-111 | |
| | 6 | 6_2 | 623 | 623-2 | 623-21 | 623-211 | |
| *** | 6 | 6_4 | 646 | 646-4 | 646-45 | 646-454 | 646-454-S5 |
| | 7 | 7_1 | 712 | 712-1 | 712-11 | 712-111 | |
| *** | 7 | 7_2 | 721 | 721-2 | 721-21 | 721-211 | |
| *** | 7 | 7_4 | 745 | 745-4 | 745-44 | 745-444 | |
| *** | 8* | 8_2 | 821 | 821-1 | 821-11 | 821-111 | |
| *** | 8 | 8_4 | 845 | 845-4 | 845-44 | 845-444 | |
| | 8 | 8_5 | 854 | 854-5 | 854-54 | 854-544 | |
| | 10 | 10_1 | 1011 | 1011-1 | 1011-12 | 1011-121 | |
| *** | 10 | 10_2 | 1022 | 1022-1 | 1022-11 | 1022-113 | 1022-113-S5 |
| *** | 10 | 10_5 | 1056 | 1056-4 | 1056-45 | 1056-454 | |
| | 13 | 13_2 | 1323 | 1323-2 | 1323-21 | 1323-211 | |
| *** | 13 | 13_3 | 1332 | 1332-1 | 1332-11 | 1332-111 | |
| *** | 13 | 13_4 | 1345 | 1345-5 | 1345-54 | 1345-544 | |
| *** | 16 | 16_1 | 1611 | 1611-2 | 1611-21 | 1611-211 | |
| | 16 | 16_2 | 1621 | 1621-2 | 1621-22 | 1621-221 | |
| *** | 16 | 16_5 | 1654 | 1654-4 | 1654-45 | 1654-454 | 1654-454-S5 |
| *** | 17 | 17_2 | 1721 | 1721-1 | 1721-11 | 1721-111 | 1721-111-S5 |
| | 17 | 17_4 | 1744 | 1744-5 | 1744-54 | 1744-544 | |
| *** | 17 | 17_5 | 1755 | 1755-5 | 1755-54 | 1755-545 | 1755-545-S5 |
| | 19 | 19_1 | 1912 | 1912-1 | 1912-13 | 1912-131 | |
| *** | 19 | 19_2 | 1922 | 1922-1 | 1922-11 | 1922-111 | |
| *** | 19 | 19_5 | 1955 | 1955-5 | 1955-54 | 1955-544 | 1955-544-S5 |
| *** | 20 | 20_1 | 2013 | 2013-1 | 2013-13 | 2013-131 | |
| *** | 20 | 20_4 | 2045 | 2045-5 | 2045-54 | 2045-544 | |
| | 21 | 21_1 | 2111 | 2111-1 | 2111-11 | 2111-111 | |
| *** | 21 | 21_2 | 2121 | 2121-1 | 2121-11 | 2121-111 | 2121-111-S5 |
| *** | 21 | 21_6 | 2165 | 2165-4 | 2165-44 | 2165-444 | 2165-444-S5 |
| *** | 23 | 23_2 | 2323 | 2323-2 | 2323-21 | 2323-211 | 2323-211-S5** |
| *** | 23 | 23_4 | 2344 | 2344-4 | 2344-46 | 2344-464 | |

| | | | | | | | |
|---|---|---|---|---|---|---|---|
| *** | 26 | 26_1 | 2612 | 2612-1 | 2612-12 | 2612-121 | |
| *** | 26 | 26_4 | 2645 | 2645-4 | 2645-46 | 2645-464 | |
| | 27 | 27_1 | 2712 | 2712-2 | 2712-21 | 2712-211 | |
| *** | 27 | 27_2 | 2721 | 2721-1 | 2721-12 | 2721-121 | |
| | 27 | 27_6 | 2765 | 2765-5 | 2765-55 | 2765-554 | |
| *** | 29 | 29_2 | 2923 | 2923-1 | 2923-12 | 2923-121 | 2923-121-S5 |
| *** | 29 | 29_4 | 2944 | 2944-5 | 2944-56 | 2944-565 | 2944-565-S5 |

| |
|---|
| Sequenced; Correctly Labeled |
| Manually imputed |
| Sequenced; Mislabeled |
| Not Sequenced; Dead/Missing |
| Not Sequenced; not imputed |

The first digit is the line identifier. Each subsequent digit identifies the seedling that was chose to continue that SL. Every generation adds an additional digit to assign a unique identifier to each individual up until the S4 generation.

S5 is indicated by adding S5 to the end of the identifier.

* Entire FS line was incorrectly labeled and is unrelated to the other lines starting with "8".
** S5 tree that was used for genome sequencing and assembly.
*** Lines used in heterozygosity and selection under selfing analysis ($n$ = 28).

**Table S2. Overview of sequencing data generated for the western redcedar draft genome.**

| *Library type* | *Fragment* | *Lanes* | *Sequence* | *Depth* |
|---|---|---|---|---|
| *Paired end* | 431 bp | 3 | 437 Gbp | 35 |
| | 433 bp | 3 | 412 Gbp | 33 |
| | 430 bp | 2 | 257 Gbp | 20.6 |
| | 419 bp | 2 | 252 Gbp | 20.2 |
| ***Total*** | | 10 | 1,358 Gbp | 108.6 |
| *Mate pair* | 2.7 kbp | 2 | 239 Gbp | 19.1 |
| | 5.4 kbp | 2 | 283 Gbp | 22.6 |
| | 9.0 kbp | 2 | 283 Gbp | 22.6 |
| | 10.9 kbp | 2 | 282 Gbp | 22.6 |
| | 15.5 kbp | 2 | 259 Gbp | 20.7 |
| ***Total*** | | 10 | 1,346 Gbp | 107.7 |
| *Linked-reads* | 35 kbp | 2 | 236 Gbp | 18.9 |
| | 35 kbp | 2 | 241 Gbp | 19.3 |
| | 20 kbp | 2 | 241 Gbp | 19.3 |
| | 22 kbp | 2 | 241 Gbp | 19.3 |
| ***Total*** | | 8 | 959 Gbp | 76.7 |

**The sequencing read length is 2×151 bp. Depth of coverage is calculated using raw sequencing data before trimming adapters and an estimated genome size of 12.8 Gbp.**

**Table S3. Metrics for each stage of the western redcedar draft genome assembly.**

| Stage | N50 (Mbp) | NG50 (Mbp) | Largest scaffold (Mbp) | Size (Gbp) | L50 (bp) | LG50 (bp) | Scaffolds |
|---|---|---|---|---|---|---|---|
| **Unitigs** | 0.00698 | 0.00420 | 0.0119 | 7.52 | 294,835 | 524,000 | 1,811,335 |
| **Contigs (PE)** | 0.0178 | 0.0129 | 166 | 7.97 | 128,893 | 196,169 | 882,457 |
| **Scaffolds (MP)** | 0.377 | 0.280 | 2.77 | 7.95 | 6,320 | 9,479 | 95,754 |
| **Tigmint** | 0.375 | 0.278 | 2.77 | 7.95 | 6,355 | 9,531 | 96,557 |
| **ARCS** | 2.04 | 1.50 | 16.3 | 7.95 | 1,176 | 1,761 | 68,083 |
| **Tigmint + ARCS** | 2.31 | 1.71 | 16.3 | 7.95 | 1,035 | 1,551 | 67,895 |

NG50 is calculated using a genome size of 10 Gbp, rounded up from the 9.8 Gbp estimated by GenomeScope.

Number of scaffolds and draft genome size includes sequences that are 1 kbp or larger.

"Scaffolds (MP)" shows the assembly contiguity after scaffolding with Illumina mate-pair sequencing.

"Tigmint + ARCS" shows the assembly contiguity after correcting misassemblies using Tigmint and then scaffolding with ARCS. "ARCS" scaffolds the assembly without first correcting misassemblies using Tigmint. Comparing these two rows shows the effects of Tigmint on improving contiguity.

**Table S4. A) Genome assembly completeness of seven currently available conifer genome assemblies as estimated by BUSCO v5.0.0 in genome mode (Embryophyta OrthoDB v10, Metaeuk). B) Predicted gene set completeness of seven sequenced conifer genomes as estimated by BUSCO v5.0.0 in protein mode (Embryophyta OrthoDB v10).**

**A**

| Taxon | ***Thuja plicata*** | ***Sequoiadendron giganteum*** (Scott et al. 2020) | ***Picea glauca*** (Warren et al. 2015) | ***Picea abies*** (Nystedt et al. 2013) | ***Pinus lambertiana*** (Stevens et al. 2016) | ***Pinus taeda*** (Zimin et al. 2014) | ***Pseudotsuga menziesii*** (Neale et al. 2017) |
|---|---|---|---|---|---|---|---|
| | **Western redcedar** | **Giant sequoia** | **White spruce** | **Norway spruce** | **Sugar pine** | **Loblolly pine** | **Douglas- fir** |
| Family | Cupressaceae | Cupressaceae | Pinaceae | Pinaceae | Pinaceae | Pinaceae | Pinaceae |
| Complete BUSCOs (C) | 1388 | 1361 | 802 | 564 | 995 | 803 | 1196 |
| Complete and single-copy BUSCOs (S) | 1293 | 1272 | 678 | 468 | 881 | 664 | 1067 |
| Complete and duplicated BUSCOs (D) | 95 | 89 | 124 | 96 | 114 | 139 | 129 |
| Fragmented BUSCOs (F) | 128 | 116 | 387 | 433 | 321 | 454 | 242 |
| Missing BUSCOs (M) | 98 | 137 | 425 | 617 | 298 | 357 | 176 |
| Total BUSCO groups searched | 1614 | 1614 | 1614 | 1614 | 1614 | 1614 | 1614 |
| Percentage complete | 86.0% | 84.3% | 49.7% | 34.9% | 61.5% | 49.7% | 74.1% |

**B**

| Taxon | *Thuja plicata* | *Sequoiadendron giganteum* | *Picea glauca* | *Picea abies* | *Pinus lambertiana* | *Pinus taeda* | *Pseudotsuga menziesii* |
|---|---|---|---|---|---|---|---|
| | **Western redcedar** | **Giant sequoia** | **White spruce** | **Norway spruce** | **Sugar pine** | **Loblolly pine** | **Douglas- fir** |
| Family | Cupressaceae | Cupressaceae | Pinaceae | Pinaceae | Pinaceae | Pinaceae | Pinaceae |
| Complete BUSCOs (C) | 1461 | 806 | 291 | 454 | 1183 | 673 | 1106 |
| Complete and single-copy BUSCOs (S) | 1367 | 747 | 141 | 381 | 1090 | 579 | 1014 |
| Complete and duplicated BUSCOs (D) | 94 | 59 | 150 | 73 | 93 | 94 | 92 |
| Fragmented BUSCOs (F) | 69 | 260 | 172 | 441 | 121 | 313 | 190 |
| Missing BUSCOs (M) | 84 | 548 | 1151 | 719 | 310 | 628 | 318 |
| Total BUSCO groups searched | 1614 | 1614 | 1614 | 1614 | 1614 | 1614 | 1614 |
| Percentage complete | 90.5% | 49.9% | 18.0% | 28.1% | 73.3% | 41.7% | 68.5% |

**Table S5. Annotation metrics and statistics for the western redcedar draft genome.**

| | |
|---|---|
| **Primary transcripts (loci)** | 39,659 |
| **Alternative transcripts (loci)** | 26,150 |
| **Total transcripts (loci)** | 65,809 |
| **For primary transcripts:** | |
| Average number of exons per transcript | 4.1 |
| Median exon length (bp) | 188 |
| Median intron length (bp) | 235 |
| Minimum intron length (bp) | 20 |
| Maximum intron length (bp) | 148,235 |
| **Gene model support (value is number of gene models):** | |
| Any EST support | 33,377 |
| EST support over 100% of their lengths | 31,537 |
| EST support over 95% of their lengths | 32,083 |
| EST support over 90% of their lengths | 32,352 |
| EST support over 75% of their lengths | 32,732 |
| EST support over 50% of their lengths | 32,983 |
| Peptide homology coverage of 100% | 1,388 |
| Peptide homology coverage of over 95% | 13,980 |
| Peptide homology coverage of over 90% | 19,506 |
| Peptide homology coverage of over 75% | 27,030 |
| Peptide homology coverage of over 50% | 31,979 |
| Pfam annotation | 25,894 |
| Panther annotation | 30,847 |
| KOG annotation | 12,416 |
| KEGG Orthology annotation | 7,088 |
| E.C. number annotation | 11,362 |

**Table S6. Verification of the western redcedar (WRC) draft genome annotation.** Of a set of 59 WRC transcripts from NCBI and 33 WRC terpene synthase transcripts, 48 and 15 were reliably identified (>90% BLASTN identity, best match in EXONERATE), respectively, in the WRC draft genome.

| | Gene | Genomic scaffold | Sequence source |
|---|---|---|---|
| 1 | gi\|1001017507\|gb\|KU701356.1\| Thuja plicata putative LOV domain-containing protein mRNA, complete cds | 29379333 | NCBI |
| 2 | gi\|1241840804\|gb\|KY214317.1\| Thuja plicata putative glycoside hydrolase family 2 protein mRNA, complete cds | 29378743 | NCBI |
| 3 | gi\|1242943498\|gb\|KY233467.1\| Thuja plicata putative MNS3 mRNA, complete cds | 29381436 | NCBI |
| 4 | gi\|1242943576\|gb\|KY233506.1\| Thuja plicata putative sphingomyelin synthetase family protein mRNA, complete cds | 29378174 | NCBI |
| 5 | gi\|1242943687\|gb\|KY233562.1\| Thuja plicata putative SPPA1 mRNA, complete cds | 29378649 | NCBI |
| 6 | gi\|1242943771\|gb\|KY249831.1\| Thuja plicata putative ABA hypersensitive 1 mRNA, complete cds | 29380247 | NCBI |
| 7 | gi\|1242943809\|gb\|KY249738.1\| Thuja plicata putative ABCG7 mRNA, complete cds | 29378727 | NCBI |
| 8 | gi\|1242943845\|gb\|KY249850.1\| Thuja plicata putative ATRRP44A mRNA, complete cds | 29382381 | NCBI |
| 9 | gi\|1242944211\|gb\|KY296273.1\| Thuja plicata cyclophilin-like peptidyl-prolyl cis-trans isomerase family protein mRNA, complete cds | 29377565 | NCBI |
| 10 | gi\|1242944351\|gb\|KY296166.1\| Thuja plicata putative OWL1 mRNA, complete cds | 29377407 | NCBI |
| 11 | gi\|1248694559\|gb\|KY314818.1\| Thuja plicata ATTRM11 mRNA, complete cds | 29380894 | NCBI |
| 12 | gi\|1248694675\|gb\|KY314876.1\| Thuja plicata FG-GAP repeat-containing protein mRNA, complete cds | 29377050 | NCBI |
| 13 | gi\|1248694816\|gb\|KY314948.1\| Thuja plicata INO80 mRNA, complete cds | 29379961 | NCBI |
| 14 | gi\|1248694885\|gb\|KY314984.1\| Thuja plicata ATCCH1 fatty acid oxygenation upregulated 2 mRNA, complete cds | 29377034 | NCBI |
| 15 | gi\|1248694997\|gb\|KY367630.1\| Thuja plicata cytochrome P450 mRNA, complete cds | 29382440 | NCBI |
| 16 | gi\|1248695181\|gb\|KY367685.1\| Thuja plicata FGGY family of carbohydrate kinase mRNA, complete cds | 29378653 | NCBI |
| 17 | gi\|1248695219\|gb\|KY367704.1\| Thuja plicata importin alpha isoform 9 mRNA, complete cds | 29381189 | NCBI |
| 18 | gi\|1248695293\|gb\|KY367741.1\| Thuja plicata HCF145 mRNA, complete cds | 29377609 | NCBI |
| 19 | gi\|1248695329\|gb\|KY367759.1\| Thuja plicata AP4B mRNA, complete cds | 29377919 | NCBI |
| 20 | gi\|1248695367\|gb\|KY367796.1\| Thuja plicata no exine fromation 1 mRNA, complete cds | 29379245 | NCBI |

| | | | |
|---|---|---|---|
| **21** | gi\|1248695439\|gb\|KY367851.1\| Thuja plicata Spc97 / Spc98 family mRNA, complete cds | 29379997 | NCBI |
| **22** | gi\|1248695589\|gb\|KY367926.1\| Thuja plicata saccharopine dehydrogenase mRNA, complete cds | 29381726 | NCBI |
| **23** | gi\|1248695663\|gb\|KY367963.1\| Thuja plicata ATDUR3 mRNA, complete cds | 29377833 | NCBI |
| **24** | gi\|1248695843\|gb\|KY368053.1\| Thuja plicata Zn-dependent exopeptidases superfamily protein mRNA, complete cds | 29382329 | NCBI |
| **25** | gi\|1248695921\|gb\|KY368092.1\| Thuja plicata ATDPE1 mRNA, complete cds | 29377587 | NCBI |
| **26** | gi\|1786237212\|gb\|MK817116.1\| Thuja plicata DXS1 mRNA, complete cds | 29379753 | NCBI |
| **27** | gi\|1786237229\|gb\|MK817117.1\| Thuja plicata DXS2B mRNA, complete cds | 29381756 | NCBI |
| **28** | gi\|1786237253\|gb\|MK817118.1\| Thuja plicata DXS2A mRNA, complete cds | 29378340 | NCBI |
| **29** | gi\|1786237278\|gb\|MK817119.1\| Thuja plicata DXS3 mRNA, complete cds | 29377322 | NCBI |
| **30** | gi\|1890528724\|gb\|MT468207.1\| Thuja plicata diterpene synthase (diTPS1) mRNA, complete cds | 29379660 | NCBI |
| **31** | gi\|1890528727\|gb\|MT468208.1\| Thuja plicata diterpene synthase (diTPS2) mRNA, complete cds | 29381898 | NCBI |
| **32** | gi\|1890528729\|gb\|MT468209.1\| Thuja plicata diterpene synthase (diTPS3) mRNA, complete cds | 29382319 | NCBI |
| **33** | gi\|510793902\|gb\|KC767281.1\| Thuja plicata sabinene synthase mRNA, complete cds | 29377789 | NCBI |
| **34** | gi\|635148812\|gb\|KJ195050.1\| Thuja plicata phototropin (PHOT1) mRNA, complete cds | 29378985 | NCBI |
| **35** | gi\|635148826\|gb\|KJ195057.1\| Thuja plicata phototropin (PHOT2) mRNA, complete cds | 29378009 | NCBI |
| **36** | gi\|6694696\|gb\|AF210063.1\| Thuja plicata dirigent-like protein (psd_Tp1) mRNA, complete cds | 29378666 | NCBI |
| **37** | gi\|6694698\|gb\|AF210064.1\| Thuja plicata dirigent protein (psd_Tp2) mRNA, complete cds_Tp4) mRNA, complete cds | 29377524 | NCBI |
| **38** | gi\|6694702\|gb\|AF210066.1\| Thuja plicata dirigent-like protein (psd_Tp4) mRNA, complete cds | 29380546 | NCBI |
| **39** | gi\|6694706\|gb\|AF210068.1\| Thuja plicata dirigent-like protein (psd_Tp6) mRNA, complete cds | 29377169 | NCBI |
| **40** | gi\|6694708\|gb\|AF210069.1\| Thuja plicata dirigent protein (psd_Tp7) mRNA, complete cds | 29207864 | NCBI |
| **41** | gi\|7542587\|gb\|AF242506.1\| Thuja plicata clone 4 pinoresinol-lariciresinol reductase mRNA, complete cds | 29377837 | NCBI |
| **42** | gi\|7578912\|gb\|AF242500.1\| Thuja plicata phenylcoumaran benzylic ether reductase homolog Tp1 mRNA, complete cds | 29381391 | NCBI |
| **43** | gi\|821372382\|gb\|KP004988.1\| Thuja plicata (+)-sabinene-3-hydroxylase mRNA, complete cds | 29379924 | NCBI |
| **44** | gi\|821372384\|gb\|KP015848.1\| Thuja plicata cytochrome P450 CYP76AA20 mRNA, complete cds | 29381501 | NCBI |
| **45** | gi\|821372386\|gb\|KP015849.1\| Thuja plicata cytochrome P450 CYP76AA21 mRNA, complete cds | 29380148 | NCBI |

| | | | |
|---|---|---|---|
| **46** | gi\|821372394\|gb\|KP015853.1\| Thuja plicata cytochrome P450 CYP76AA25 mRNA, complete cds | 29377693 | NCBI |
| **47** | gi\|821372396\|gb\|KP015854.1\| Thuja plicata cytochrome P450 CYP76AA26 mRNA, complete cds | 29377042 | NCBI |
| **48** | gi\|821372398\|gb\|KP015855.1\| Thuja plicata cytochrome P450 CYP76Z2 mRNA, complete cds | 29377249 | NCBI |
| **1** | TpTPS2_evgvelvLoc955t9 | 29194648 | Putative WRC terpene synthase transcripts |
| **2** | TpTPS3_evgvelvLoc2319t10 | 29382319 | Putative WRC terpene synthase transcripts |
| **3** | TpTPS10_evgvelvLoc9436t1 | 29381898 | Putative WRC terpene synthase transcripts |
| **4** | TpTPS13_evgvelvLoc5170t3 | 29378510 | Putative WRC terpene synthase transcripts |
| **5** | TpTPS21_evgvelvLoc4915t2 | 29379564 | Putative WRC terpene synthase transcripts |
| **6** | TpTPS24_evgvelvLoc16400d36129t1 | 29381804 | Putative WRC terpene synthase transcripts |
| **7** | TpTPS25_evgvelvLoc2138t7 | 29382319 | Putative WRC terpene synthase transcripts |
| **8** | TpTPS26_evgvelvLoc4951t5 | 29380592 | Putative WRC terpene synthase transcripts |
| **9** | TpTPS27_evgvelvLoc1076t3 | 29377789 | Putative WRC terpene synthase transcripts |
| **10** | TpTPS28_evgvelvLoc842d111924t1 | 29379713 | Putative WRC terpene synthase transcripts |
| **11** | TpTPS29_evgvelvLoc7235d121309t1 | 29380776 | Putative WRC terpene synthase transcripts |
| **12** | TpTPS30_evgvelvLoc16257d162311t1 | 29381146 | Putative WRC terpene synthase transcripts |
| **13** | TpTPS31_evgR1187289 | 29380228 | Putative WRC terpene synthase transcripts |
| **14** | TpTPS32_evgR741119 | 29379604 | Putative WRC terpene synthase transcripts |
| **15** | TpTPS33_evgS303450 | 29380592 | Putative WRC terpene synthase transcripts |

**Table S7. The range-wide population (*n* = 112) of western redcedar used in this study.** The population was stratified into three subpopulations based on the analysis of O'Connell et al. (2008).

| Tree ID | Subpopulation | Location | Latitude (° N) | Longitude (° W) | Elevation (m) |
|---|---|---|---|---|---|
| **2** | Central | Jordan River | 48.477 | -123.929 | 50 |
| **85** | Central | Powell Lake | 50.233 | -124.22 | 15 |
| **148** | Central | Tzoonie River | 49.867 | -123.44 | 425 |
| **149** | Central | Tzoonie River | 49.867 | -123.44 | 365 |
| **150** | Central | Tzoonie River | 49.867 | -123.733 | 450 |
| **164** | Central | Powell Lake | 50.30 | -124.37 | 15 |
| **165** | Central | Lois Lake | 49.817 | -124.19 | 250 |
| **171** | Central | Lewis Lake | 49.933 | -124.22 | 470 |
| **173** | Central | Labour Day Lake | 49.133 | -124.27 | 975 |
| **175** | Central | Francis Lake | 48.933 | -124.42 | 503 |
| **176** | Central | San Mateo Bay | 48.95 | -124.59 | 90 |
| **181** | Central | Cameron Lake | 49.32 | -124.48 | 15 |
| **191** | Central | Jervis Inlet | 50.117 | -123.783 | 98 |
| **202** | Central | Red Tusk | 49.767 | -123.24 | 240 |
| **203** | Central | Sechelt Creek | 49.676 | -123.535 | 400 |
| **204** | Central | Bear Creek | 49.8 | -123.32 | 580 |
| **263** | Central | Narrows Inlet | 49.083 | -123.733 | 120 |
| **264** | Central | Narrows Inlet | 49.083 | -123.44 | 210 |
| **351** | Central | Second Lake | 49.1 | -124.13 | 300 |
| **392** | Central | Hardy Island | 49.75 | -124.18 | 37 |
| **394** | Central | Lois Lake | 49.8 | -124.17 | 262 |
| **426** | Central | Jordan River | 48.52 | -124.17 | 730 |
| **430** | Central | Errington | 49.267 | -124.05 | 152 |
| **435** | Central | Chemainus | 48.933 | -123.75 | 115 |
| **438** | Central | Bamberton | 48.617 | -123.33 | 168 |
| **497** | Central | Caycuse Lake | 48.8 | -124.36 | 230 |
| **1** | Northern-Coastal | Keogh River | 50.617 | -127.19 | 61 |
| **5** | Northern-Coastal | Mahatta Creek | 50.4 | -127.783 | 122 |
| **10** | Northern-Coastal | Mahatta Creek | 50.317 | -127.39 | 396 |
| **15** | Northern-Coastal | Mahatta Creek | 50.4 | -127.783 | 155 |
| **19** | Northern-Coastal | Benson Lake | 50.4 | -127.26 | 637 |
| **20** | Northern-Coastal | Snowsaddle Mountain | 50.25 | -127.33 | 305 |
| **40** | Northern-Coastal | Keogh River | 50.633 | -127.23 | 91 |
| **44** | Northern-Coastal | Waukwaas Creek | 50.55 | -127.18 | 183 |
| **47** | Northern-Coastal | Waukwaas Creek | 50.583 | -127.21 | 30 |

| | | | | | |
|---|---|---|---|---|---|
| **49** | Northern-Coastal | Misty Lake | 50.633 | -127.283 | 91 |
| **58** | Northern-Coastal | MacJack River | 50.617 | -128.09 | 60 |
| **87** | Northern-Coastal | Louise Island | 52.92 | -131.85 | 600 |
| **88** | Northern-Coastal | Skedans Islands | 52.95 | -131.57 | 395 |
| **94** | Northern-Coastal | Talunkwan Island | 52.829 | -131.752 | 335 |
| **95** | Northern-Coastal | Tasu | 52.817 | -132.04 | 275 |
| **97** | Northern-Coastal | Tasu | 52.817 | -132.04 | 565 |
| **113** | Northern-Coastal | Gogit Passage | 52.633 | -131.29 | 245 |
| **116** | Northern-Coastal | Sedgwick Bay | 52.6 | -131.34 | 305 |
| **120** | Northern-Coastal | Tanu Passage | 52.767 | -131.44 | 365 |
| **121** | Northern-Coastal | Tanu Island | 52.767 | -131.41 | 305 |
| **126** | Northern-Coastal | Ramsay Island | 52.564 | -131.379 | 150 |
| **128** | Northern-Coastal | Huxley Island | 52.432 | -131.268 | 185 |
| **133** | Northern-Coastal | Huston Inlet | 52.267 | -131.17 | 185 |
| **137** | Northern-Coastal | Tangil Peninsula | 52.791 | -131.705 | 305 |
| **138** | Northern-Coastal | Talunkwan Island | 52.829 | -131.692 | 215 |
| **144** | Northern-Coastal | Stewardson Inlet | 49.417 | -126.18 | 410 |
| **156** | Northern-Coastal | Waring Creek | 49.95 | -126.08 | 350 |
| **158** | Northern-Coastal | Cougar Creek | 49.717 | -126.25 | 355 |
| **220** | Northern-Coastal | Roselle Lake | 50.517 | -127 | 120 |
| **221** | Northern-Coastal | Nimpkish Lake | 50.483 | -126.59 | 75 |
| **230** | Northern-Coastal | Long Lake | 51.3 | -127.02 | 317 |
| **231** | Northern-Coastal | Long Lake | 51.233 | -127.14 | 268 |
| **232** | Northern-Coastal | Naysash Inlet | 51.3 | -127.2 | 110 |
| **235** | Northern-Coastal | Amback Creek | 51.717 | -127.02 | 189 |
| **241** | Northern-Coastal | Rivers Inlet | 51.60 | -127.58 | 300 |
| **243** | Northern-Coastal | Hardy Inlet | 51.717 | -127.38 | 159 |
| **253** | Northern-Coastal | Kwatna | 52.033 | -127.25 | 985 |
| **261** | Northern-Coastal | Link Lake | 52.433 | -127.42 | 950 |
| **266** | Northern-Coastal | Slim Creek | 52.87 | -132.08 | 75 |
| **281** | Northern-Coastal | Gwaii Haanas | 52.45 | -131.45 | 45 |
| **285** | Northern-Coastal | Misty Lake | 50.616 | -127.24 | 90 |
| **301** | Northern-Coastal | Waukwaas River | 50.6 | -127.23 | 100 |
| **306** | Northern-Coastal | Denad Creek | 50.55 | -128 | 25 |
| **322** | Northern-Coastal | Koprino River | 50.55 | -127.5 | 280 |
| **325** | Northern-Coastal | Hathaway Creek | 50.55 | -127.51 | 95 |
| **329** | Northern-Coastal | Victoria Lake | 50.43 | -127.40 | 123 |
| **331** | Northern-Coastal | Victoria Lake | 50.317 | -127.22 | 200 |
| **334** | Northern-Coastal | Snowsaddle Mountain | 50.25 | -127.2 | 213 |
| **336** | Northern-Coastal | Victoria Lake | 50.35 | -127.22 | 122 |
| **340** | Northern-Coastal | Utlah Creek | 50.3 | -127.26 | 183 |

| | | | | | |
|---|---|---|---|---|---|
| **343** | Northern-Coastal | Victoria Lake | 50.05 | -127.417 | 457 |
| **344** | Northern-Coastal | Victoria Lake | 50.05 | -127.417 | 457 |
| **347** | Northern-Coastal | Utlah Creek | 50.283 | -127.25 | 244 |
| **366** | Northern-Coastal | Henderson Lake | 49.1 | -125.02 | 300 |
| **367** | Northern-Coastal | Henderson Lake | 49.1 | -125.017 | 200 |
| **368** | Northern-Coastal | Henderson Lake | 49.10 | -125.03 | 280 |
| **378** | Northern-Coastal | Tranquil Inlet | 49.2 | -125.4 | 3 |
| **381** | Northern-Coastal | Sydney Inlet | 49.483 | -126.283 | 65 |
| **386** | Northern-Coastal | Cypre River | 49.367 | -125.867 | 180 |
| **402** | Northern-Coastal | Tahsish River | 50.15 | -127.13 | 15 |
| **404** | Northern-Coastal | Chamiss Bay | 50.08 | -127.35 | 350 |
| **415** | Northern-Coastal | Lull Creek | 50.7 | -126.02 | 183 |
| **421** | Northern-Coastal | Wood Bay | 50.317 | -125.2 | 200 |
| **444** | Northern-Coastal | Anutz Lake | 50.3 | -126.55 | 80 |
| **446** | Northern-Coastal | Kinman Creek | 50.35 | -126.54 | 250 |
| **454** | Northern-Coastal | Mt. Bate | 49.85 | -126.51 | 1000 |
| **461** | Northern-Coastal | Artlish Caves | 50.133 | -127.083 | 100 |
| **465** | Northern-Coastal | Union Island | 50.029 | -127.277 | 50 |
| **466** | Northern-Coastal | Hohoae Island | 50.048 | -127.207 | 50 |
| **490** | Northern-Coastal | Mitla Creek | 49.5 | -125.58 | 220 |
| **493** | Northern-Coastal | Fortune Creek | 49.183 | -125.44 | 365 |
| **501** | Northern-Coastal | Snug Bay | 49.033 | -125.01 | 60 |
| **503** | Northern-Coastal | Nahmint Lake | 49.183 | -125.05 | 150 |
| **505** | Northern-Coastal | Bigtree Creek | 50.267 | -125.43 | 150 |
| **508** | Northern-Coastal | Brasseau Bay | 50.4 | -125.59 | 45 |
| **779** | Northern-Coastal | Mt. Washington | 49.733 | -125.14 | 950 |
| **784** | Northern-Coastal | Mt. Washington | 49.7 | -125.05 | 110 |
| **214** | Southern-Interior | Burke Mountain | 49.317 | -122.44 | 440 |
| **411** | Southern-Interior | Debeck Creek | 49.483 | -122.36 | 275 |
| **527** | Southern-Interior | Tumtum Lake | 51.75 | -119.09 | 960 |
| **528** | Southern-Interior | Tumtum Lake | 51.083 | -119.09 | 910 |
| **531** | Southern-Interior | East Barrier Lake | 50.85 | -119.44 | 610 |
| **566** | Southern-Interior | Price Creek | 40.05 | -124.15 | 50 |
| **648** | Southern-Interior | Rector Ridge | 45.783 | -123.4 | 246 |
| **662** | Southern-Interior | Simmons Loop | 45.367 | -123.45 | 246 |
| **1081** | Southern-Interior | Alsea River | 44.45 | -123.59 | 49 |

**Table S8. SNP effects as predicted by Ensembl Variant Effect Predictor (McLaren et al. 2016) for 17,728 SNPs out of the 18,371 SNPs in our dataset.**

| SNP effect | # of SNPs | % of total | Region | IMPACT |
|---|---|---|---|---|
| *Splice acceptor variant* | 9 | 0.0490 | Non-coding | High |
| *Splice donor variant* | 7 | 0.0381 | Non-coding | High |
| *Stop gained* | 69 | 0.376 | Coding | High |
| *Stop lost* | 78 | 0.425 | Coding | High |
| *Start lost* | 4 | 0.0218 | Coding | High |
| *Missense variant* | 3002 | 16.34 | Coding | Moderate |
| *Splice region variant* | 221 | 1.20 | Non-coding | Low |
| *Stop retained variant* | 50 | 0.272 | Coding | Low |
| *Synonymous variant* | 2807 | 15.3 | Coding | Low |
| *5′ UTR variant* | 1896 | 10.3 | Non-coding | Modifier |
| *3′ UTR variant* | 2680 | 14.6 | Non-coding | Modifier |
| *Intron variant* | 2274 | 12.4 | Non-coding | Modifier |
| *Upstream gene variant* | 607 | 3.30 | Intergenic | Modifier |
| *Downstream gene variant* | 498 | 2.71 | Intergenic | Modifier |
| *Intergenic variant* | 3526 | 19.2 | Intergenic | Modifier |
| *Unknown* | 643 | 3.50 | - | - |
| **Total** | **18371** | | | |

Effects are listed from top to bottom in order of severity from most to least severe.

IMPACTs are defined by Ensembl as follows:
**High:** The variant is assumed to have high (disruptive) impact in the protein, probably causing protein truncation, loss of function or triggering nonsense mediated decay.
**Moderate:** A non-disruptive variant that might change protein effectiveness.
**Low:** A variant that is assumed to be mostly harmless or unlikely to change protein behaviour.
**Modifier:** Usually non-coding variants or variants affecting non-coding genes, where predictions are difficult or there is no evidence of impact.

**Table S9. Pairwise $F_{ST}$ for all subpopulations.** $F_{ST}$ values are based on Weir and Cockerham's estimator (Weir and Cockerham 1984)**.**

| Subpopulation | Northern-Coastal | Central | Southern-Interior |
|---|---|---|---|
| **Northern-Coastal** | - | - | - |
| **Central** | 0.0335 | - | - |
| **Southern-Interior** | 0.0412 | 0.00961 | - |

**Table S10. Annotation for single-copy ortholog genes with highest 1% of π estimates of nucleotide diversity (*n* = 47).**

| Gene | Annotation source | Annotation access code | Annotation |
|---|---|---|---|
| **Thupl.293114653s0001** | PANTHER | PTHR11926:SF500 | - |
| **Thupl.293114653s0001** | KEGGORTH | K08237 | hydroquinone glucosyltransferase [EC:2.4.1.218] |
| **Thupl.293114653s0001** | PANTHER | PTHR11926 | GLUCOSYL/GLUCURONOSYL TRANSFERASES |
| **Thupl.293114653s0001** | GO | GO:0008152 | metabolic process |
| **Thupl.293114653s0001** | GO | GO:0016758 | transferase activity, transferring hexosyl groups |
| **Thupl.293114653s0001** | EC | 2.4.1.170 | Isoflavone 7-O-glucosyltransferase |
| **Thupl.293114653s0001** | PFAM | PF00201 | UDP-glucoronosyl and UDP-glucosyl transferase |
| **Thupl.29343737s0001** | GO | GO:0005618 | cell wall |
| **Thupl.29343737s0001** | PFAM | PF04043 | Plant invertase/pectin methylesterase inhibitor |
| **Thupl.29343737s0001** | PANTHER | PTHR31707:SF25 | - |
| **Thupl.29343737s0001** | EC | 3.1.1.11 | Pectinesterase |
| **Thupl.29343737s0001** | GO | GO:0004857 | enzyme inhibitor activity |
| **Thupl.29343737s0001** | PFAM | PF01095 | Pectinesterase |
| **Thupl.29343737s0001** | GO | GO:0042545 | cell wall modification |
| **Thupl.29343737s0001** | KEGGORTH | K01051 | pectinesterase [EC:3.1.1.11] |
| **Thupl.29343737s0001** | GO | GO:0030599 | pectinesterase activity |
| **Thupl.29343737s0001** | PANTHER | PTHR31707 | FAMILY NOT NAMED |
| **Thupl.29376925s0001** | KEGGORTH | K08762 | diazepam-binding inhibitor (GABA receptor modulator, acyl-CoA-binding protein) |
| **Thupl.29376925s0001** | PANTHER | PTHR23310 | ACYL-COA-BINDING PROTEIN, ACBP |
| **Thupl.29376925s0001** | PFAM | PF00887 | Acyl CoA binding protein |
| **Thupl.29376925s0001** | GO | GO:0000062 | fatty-acyl-CoA binding |
| **Thupl.29376925s0001** | PANTHER | PTHR23310:SF67 | - |
| **Thupl.29377001s0021** | GO | GO:0016616 | oxidoreductase activity, acting on the CH-OH group of donors, NAD or NADP as acceptor |
| **Thupl.29377001s0021** | KOG | KOG1494 | NAD-dependent malate dehydrogenase |
| **Thupl.29377001s0021** | PFAM | PF00056 | lactate/malate dehydrogenase, NAD binding domain |
| **Thupl.29377001s0021** | PANTHER | PTHR11540:SF21 | L-LACTATE DEHYDROGENASE |
| **Thupl.29377001s0021** | PFAM | PF02866 | lactate/malate dehydrogenase, alpha/beta C-terminal domain |
| **Thupl.29377001s0021** | GO | GO:0055114 | oxidation-reduction process |
| **Thupl.29377001s0021** | GO | GO:0016491 | oxidoreductase activity |
| **Thupl.29377001s0021** | KEGGORTH | K00026 | malate dehydrogenase [EC:1.1.1.37] |
| **Thupl.29377001s0021** | PANTHER | PTHR11540 | MALATE AND LACTATE DEHYDROGENASE |
| **Thupl.29377001s0021** | EC | 1.1.1.37 | Malate dehydrogenase |
| **Thupl.29377148s0010** | KOG | KOG3328 | HGG motif-containing thioesterase |
| **Thupl.29377148s0010** | EC | 3.1.2.20 | Acyl-CoA hydrolase |

| | | | |
|---|---|---|---|
| **Thupl.29377148s0010** | KEGGORTH | K17362 | - |
| **Thupl.29377148s0010** | PANTHER | PTHR21660:SF8 | - |
| **Thupl.29377148s0010** | PANTHER | PTHR21660 | THIOESTERASE SUPERFAMILY MEMBER-RELATED |
| **Thupl.29377148s0010** | PFAM | PF03061 | Thioesterase superfamily |
| **Thupl.29377170s0054** | GO | GO:0045263 | proton-transporting ATP synthase complex, coupling factor F(o) |
| **Thupl.29377170s0054** | GO | GO:0015078 | hydrogen ion transmembrane transporter activity |
| **Thupl.29377170s0054** | KEGGORTH | K02109 | F-type H+-transporting ATPase subunit b [EC:3.6.3.14] |
| **Thupl.29377170s0054** | PANTHER | PTHR33445 | - |
| **Thupl.29377170s0054** | PFAM | PF00430 | ATP synthase B/B' CF(0) |
| **Thupl.29377170s0054** | GO | GO:0015986 | ATP synthesis coupled proton transport |
| **Thupl.29377251s0003** | PANTHER | PTHR33918 | - |
| **Thupl.29377251s0003** | PANTHER | PTHR33918:SF2 | - |
| **Thupl.29377336s0006** | EC | 6.3.5.7 | Glutaminyl-tRNA synthase (glutamine-hydrolyzing) |
| **Thupl.29377336s0006** | GO | GO:0016874 | ligase activity |
| **Thupl.29377336s0006** | KOG | KOG2438 | Glutamyl-tRNA amidotransferase subunit B |
| **Thupl.29377336s0006** | PFAM | PF02637 | GatB domain |
| **Thupl.29377336s0006** | PFAM | PF02934 | GatB/GatE catalytic domain |
| **Thupl.29377336s0006** | PANTHER | PTHR11659 | GLUTAMYL-TRNA(GLN) AMIDOTRANSFERASE SUBUNIT B (MITOCHONDRIAL AND PROKARYOTIC) PET112-RELATED |
| **Thupl.29377336s0006** | GO | GO:0016884 | carbon-nitrogen ligase activity, with glutamine as amido-N-donor |
| **Thupl.29377336s0006** | KEGGORTH | K02434 | aspartyl-tRNA(Asn)/glutamyl-tRNA (Gln) amidotransferase subunit B [EC:6.3.5.6 6.3.5.7] |
| **Thupl.29377336s0006** | PANTHER | PTHR11659:SF3 | - |
| **Thupl.29377352s0001** | GO | GO:0005789 | endoplasmic reticulum membrane |
| **Thupl.29377352s0001** | SIGNALP | SignalP-noTM | - |
| **Thupl.29377352s0001** | PFAM | PF03896 | Translocon-associated protein (TRAP), alpha subunit |
| **Thupl.29377352s0001** | KOG | KOG1631 | Translocon-associated complex TRAP, alpha subunit |
| **Thupl.29377352s0001** | KEGGORTH | K13249 | translocon-associated protein subunit alpha |
| **Thupl.29377352s0001** | PANTHER | PTHR12924 | TRANSLOCON-ASSOCIATED PROTEIN, ALPHA SUBUNIT |
| **Thupl.29377414s0008** | PFAM | PF07145 | Ataxin-2 C-terminal region |
| **Thupl.29377414s0008** | PANTHER | PTHR33790 | - |
| **Thupl.29377534s0013** | PANTHER | PTHR21257 | STEROL REDUCTASE/LAMIN B RECEPTOR |
| **Thupl.29377534s0013** | EC | 1.3.1.21 | 7-dehydrocholesterol reductase |
| **Thupl.29377534s0013** | GO | GO:0016020 | membrane |
| **Thupl.29377534s0013** | KOG | KOG1435 | Sterol reductase/lamin B receptor |

| | | | |
|---|---|---|---|
| **Thupl.29377534s0013** | PFAM | PF01222 | Ergosterol biosynthesis ERG4/ERG24 family |
| **Thupl.29377534s0013** | KEGGORTH | K00213 | 7-dehydrocholesterol reductase [EC:1.3.1.21] |
| **Thupl.29377534s0013** | PANTHER | PTHR21257:SF38 | - |
| **Thupl.29377763s0003** | PANTHER | PTHR10643 | KINETOCHORE PROTEIN NDC80 |
| **Thupl.29377763s0003** | PFAM | PF03801 | HEC/Ndc80p family |
| **Thupl.29377763s0003** | KEGGORTH | K11547 | kinetochore protein NDC80 |
| **Thupl.29377763s0003** | KOG | KOG0995 | Centromere-associated protein HEC1 |
| **Thupl.29377905s0004** | PANTHER | PTHR11630:SF66 | - |
| **Thupl.29377905s0004** | PFAM | PF00493 | MCM2/3/5 family |
| **Thupl.29377905s0004** | GO | GO:0042555 | MCM complex |
| **Thupl.29377905s0004** | GO | GO:0003678 | DNA helicase activity |
| **Thupl.29377905s0004** | KOG | KOG0478 | DNA replication licensing factor, MCM4 component |
| **Thupl.29377905s0004** | PFAM | PF17207 | - |
| **Thupl.29377905s0004** | PFAM | PF14551 | MCM N-terminal domain |
| **Thupl.29377905s0004** | KEGGORTH | K02212 | minichromosome maintenance protein 4 (cell division control protein 54) |
| **Thupl.29377905s0004** | PANTHER | PTHR11630 | DNA REPLICATION LICENSING FACTOR |
| **Thupl.29377905s0004** | GO | GO:0003677 | DNA binding |
| **Thupl.29377905s0004** | EC | 3.6.4.12 | DNA helicase |
| **Thupl.29377905s0004** | GO | GO:0006270 | DNA replication initiation |
| **Thupl.29377905s0004** | GO | GO:0005524 | ATP binding |
| **Thupl.29378116s0023** | GO | GO:0016020 | membrane |
| **Thupl.29378116s0023** | GO | GO:0015297 | antiporter activity |
| **Thupl.29378116s0023** | PFAM | PF01554 | MatE |
| **Thupl.29378116s0023** | GO | GO:0015238 | drug transmembrane transporter activity |
| **Thupl.29378116s0023** | PANTHER | PTHR11206 | MULTIDRUG RESISTANCE PROTEIN |
| **Thupl.29378116s0023** | GO | GO:0055085 | transmembrane transport |
| **Thupl.29378116s0023** | KEGGORTH | K03327 | multidrug resistance protein, MATE family |
| **Thupl.29378116s0023** | PANTHER | PTHR11206:SF85 | - |
| **Thupl.29378116s0023** | KOG | KOG1347 | Uncharacterized membrane protein, predicted efflux pump |
| **Thupl.29378116s0023** | GO | GO:0006855 | drug transmembrane transport |
| **Thupl.29378169s0021** | PANTHER | PTHR36000 | - |
| **Thupl.29378269s0005** | PANTHER | PTHR31391 | FAMILY NOT NAMED |
| **Thupl.29378269s0005** | GO | GO:0003677 | DNA binding |
| **Thupl.29378269s0005** | PFAM | PF02362 | B3 DNA binding domain |
| **Thupl.29378329s0001** | PANTHER | PTHR31048:SF81 | - |
| **Thupl.29378329s0001** | PFAM | PF00314 | Thaumatin family |
| **Thupl.29378329s0001** | PANTHER | PTHR31048 | FAMILY NOT NAMED |
| **Thupl.29378467s0019** | KOG | KOG2014 | SMT3/SUMO-activating complex, AOS1/RAD31 component |
| **Thupl.29378467s0019** | PANTHER | PTHR10953 | UBIQUITIN-ACTIVATING ENZYME E1 |

| | | | |
|---|---|---|---|
| **Thupl.29378467s0019** | KEGGORTH | K10684 | ubiquitin-like 1-activating enzyme E1 A [EC:6.3.2.19] |
| **Thupl.29378467s0019** | GO | GO:0008641 | small protein activating enzyme activity |
| **Thupl.29378467s0019** | PANTHER | PTHR10953:SF162 | - |
| **Thupl.29378467s0019** | PFAM | PF00899 | ThiF family |
| **Thupl.29378604s0006** | NA | NA | NA |
| **Thupl.29378682s0038** | KEGGORTH | K00472 | prolyl 4-hydroxylase [EC:1.14.11.2] |
| **Thupl.29378682s0038** | PFAM | PF13640 | 2OG-Fe(II) oxygenase superfamily |
| **Thupl.29378682s0038** | KOG | KOG1591 | Prolyl 4-hydroxylase alpha subunit |
| **Thupl.29378682s0038** | PANTHER | PTHR10869:SF133 | - |
| **Thupl.29378682s0038** | GO | GO:0055114 | oxidation-reduction process |
| **Thupl.29378682s0038** | SIGNALP | SignalP-noTM | - |
| **Thupl.29378682s0038** | GO | GO:0016491 | oxidoreductase activity |
| **Thupl.29378682s0038** | PANTHER | PTHR10869 | PROLYL 4-HYDROXYLASE ALPHA SUBUNIT |
| **Thupl.29378682s0038** | EC | 1.14.11.2 | Procollagen-proline dioxygenase |
| **Thupl.29378750s0001** | PFAM | PF07847 | Protein of unknown function (DUF1637) |
| **Thupl.29378750s0001** | PANTHER | PTHR22966:SF1 | gb def: CG7550-PA |
| **Thupl.29378750s0001** | EC | 1.13.11.19 | Cysteamine dioxygenase |
| **Thupl.29378750s0001** | KOG | KOG4281 | Uncharacterized conserved protein |
| **Thupl.29378750s0001** | GO | GO:0055114 | oxidation-reduction process |
| **Thupl.29378750s0001** | KEGGORTH | K10712 | cysteamine dioxygenase [EC:1.13.11.19] |
| **Thupl.29378750s0001** | GO | GO:0016702 | oxidoreductase activity, acting on single donors with incorporation of molecular oxygen, incorporation of two atoms of oxygen |
| **Thupl.29378750s0001** | PANTHER | PTHR22966 | UNCHARACTERIZED |
| **Thupl.29378953s0001** | PFAM | PF01535 | PPR repeat |
| **Thupl.29378953s0001** | PANTHER | PTHR44149 | - |
| **Thupl.29378953s0001** | PANTHER | PTHR44149:SF1 | - |
| **Thupl.29378953s0001** | PFAM | PF13041 | PPR repeat family |
| **Thupl.29378953s0001** | PFAM | PF12854 | PPR repeat |
| **Thupl.29379280s0001** | PANTHER | PTHR22814:SF78 | - |
| **Thupl.29379280s0001** | PFAM | PF00403 | Heavy-metal-associated domain |
| **Thupl.29379280s0001** | GO | GO:0046872 | metal ion binding |
| **Thupl.29379280s0001** | KOG | KOG1603 | Copper chaperone |
| **Thupl.29379280s0001** | PANTHER | PTHR22814 | COPPER TRANSPORT PROTEIN ATOX1-RELATED |
| **Thupl.29379280s0001** | GO | GO:0030001 | metal ion transport |
| **Thupl.29379302s0001** | PANTHER | PTHR44566 | - |
| **Thupl.29379302s0001** | GO | GO:0005515 | protein binding |
| **Thupl.29379302s0001** | KOG | KOG0265 | U5 snRNP-specific protein-like factor and related proteins |
| **Thupl.29379302s0001** | PFAM | PF00400 | WD domain, G-beta repeat |
| **Thupl.29379321s0005** | KEGGORTH | K09838 | zeaxanthin epoxidase [EC:1.14.13.90] |

| | | | |
|---|---|---|---|
| **Thupl.29379321s0005** | PANTHER | PTHR13789:SF245 | - |
| **Thupl.29379321s0005** | PFAM | PF00498 | FHA domain |
| **Thupl.29379321s0005** | KOG | KOG2614 | Kynurenine 3-monooxygenase and related flavoprotein monooxygenases |
| **Thupl.29379321s0005** | EC | 1.14.13.90 | Zeaxanthin epoxidase |
| **Thupl.29379321s0005** | GO | GO:0005515 | protein binding |
| **Thupl.29379321s0005** | PFAM | PF01494 | FAD binding domain |
| **Thupl.29379321s0005** | GO | GO:0071949 | FAD binding |
| **Thupl.29379321s0005** | PANTHER | PTHR13789 | MONOOXYGENASE |
| **Thupl.29379402s0007** | PFAM | PF02668 | Taurine catabolism dioxygenase TauD, TfdA family |
| **Thupl.29379402s0007** | GO | GO:0055114 | oxidation-reduction process |
| **Thupl.29379402s0007** | PANTHER | PTHR10696 | GAMMA-BUTYROBETAINE HYDROXYLASE-RELATED |
| **Thupl.29379402s0007** | PANTHER | PTHR10696:SF29 | - |
| **Thupl.29379402s0007** | GO | GO:0016491 | oxidoreductase activity |
| **Thupl.29379488s0007** | PFAM | PF13862 | p21-C-terminal region-binding protein |
| **Thupl.29379488s0007** | KEGGORTH | K15262 | - |
| **Thupl.29379488s0007** | PANTHER | PTHR13261 | CDK INHIBITOR P21 BINDING PROTEIN |
| **Thupl.29379488s0007** | KOG | KOG3034 | Isoamyl acetate-hydrolyzing esterase and related enzymes |
| **Thupl.29379722s0024** | PANTHER | PTHR10209:SF251 | - |
| **Thupl.29379722s0024** | PANTHER | PTHR10209 | OXIDOREDUCTASE, 2OG-FE(II) OXYGENASE FAMILY PROTEIN |
| **Thupl.29379722s0024** | PFAM | PF03171 | 2OG-Fe(II) oxygenase superfamily |
| **Thupl.29379722s0024** | GO | GO:0016491 | oxidoreductase activity |
| **Thupl.29379722s0024** | PFAM | PF14226 | non-haem dioxygenase in morphine synthesis N-terminal |
| **Thupl.29379722s0024** | EC | 1.14.11.9 | Flavanone 3-dioxygenase |
| **Thupl.29379722s0024** | KOG | KOG0143 | Iron/ascorbate family oxidoreductases |
| **Thupl.29379722s0024** | GO | GO:0055114 | oxidation-reduction process |
| **Thupl.29379747s0003** | PANTHER | PTHR33648 | - |
| **Thupl.29379747s0003** | SIGNALP | SignalP-noTM | - |
| **Thupl.29379768s0005** | KEGGORTH | K19307 | - |
| **Thupl.29379768s0005** | PANTHER | PTHR11538 | PHENYLALANYL-TRNA SYNTHETASE |
| **Thupl.29379768s0005** | KOG | KOG4174 | Uncharacterized conserved protein |
| **Thupl.29379768s0005** | PFAM | PF10354 | Domain of unknown function (DUF2431) |
| **Thupl.29379768s0005** | PANTHER | PTHR11538:SF54 | - |
| **Thupl.29379839s0006** | PANTHER | PTHR12169 | ATPASE N2B |
| **Thupl.29379839s0006** | GO | GO:0005524 | ATP binding |
| **Thupl.29379839s0006** | PANTHER | PTHR12169:SF6 | - |
| **Thupl.29379839s0006** | EC | 2.7.1.1 | Hexokinase |
| **Thupl.29379933s0022** | GO | GO:0009247 | glycolipid biosynthetic process |

| | | | |
|---|---|---|---|
| **Thupl.29379933s0022** | PFAM | PF04101 | Glycosyltransferase family 28 C-terminal domain |
| **Thupl.29379933s0022** | GO | GO:0016758 | transferase activity, transferring hexosyl groups |
| **Thupl.29379933s0022** | PFAM | PF06925 | Monogalactosyldiacylglycerol (MGDG) synthase |
| **Thupl.29379933s0022** | EC | 2.4.1.46 | Monogalactosyldiacylglycerol synthase |
| **Thupl.29379933s0022** | PANTHER | PTHR43025 | - |
| **Thupl.29379933s0022** | PANTHER | PTHR43025:SF1 | - |
| **Thupl.29379933s0022** | KEGGORTH | K03715 | 1,2-diacylglycerol 3-beta-galactosyltransferase [EC:2.4.1.46] |
| **Thupl.29380045s0002** | PANTHER | PTHR23417:SF16 | - |
| **Thupl.29380045s0002** | GO | GO:0008176 | tRNA (guanine-N7-)-methyltransferase activity |
| **Thupl.29380045s0002** | KEGGORTH | K03439 | tRNA (guanine-N7-)-methyltransferase [EC:2.1.1.33] |
| **Thupl.29380045s0002** | KOG | KOG3115 | Methyltransferase-like protein |
| **Thupl.29380045s0002** | EC | 2.1.1.33 | tRNA (guanine(46)-N(7))-methyltransferase |
| **Thupl.29380045s0002** | PANTHER | PTHR23417 | 3-DEOXY-D-MANNO-OCTULOSONIC-ACID TRANSFERASE/TRNA (GUANINE-N(7)-)-METHYLTRANSFERASE |
| **Thupl.29380045s0002** | GO | GO:0006400 | tRNA modification |
| **Thupl.29380045s0002** | PFAM | PF02390 | Putative methyltransferase |
| **Thupl.29380174s0008** | PFAM | PF01195 | Peptidyl-tRNA hydrolase |
| **Thupl.29380174s0008** | PANTHER | PTHR17224:SF4 | - |
| **Thupl.29380174s0008** | SIGNALP | SignalP-noTM | - |
| **Thupl.29380174s0008** | PANTHER | PTHR17224 | PEPTIDYL-TRNA HYDROLASE |
| **Thupl.29380174s0008** | KEGGORTH | K01056 | peptidyl-tRNA hydrolase, PTH1 family [EC:3.1.1.29] |
| **Thupl.29380174s0008** | KOG | KOG2255 | Peptidyl-tRNA hydrolase |
| **Thupl.29380174s0008** | EC | 3.1.1.29 | Aminoacyl-tRNA hydrolase |
| **Thupl.29380174s0008** | GO | GO:0004045 | aminoacyl-tRNA hydrolase activity |
| **Thupl.29380345s0016** | GO | GO:0003723 | RNA binding |
| **Thupl.29380345s0016** | PANTHER | PTHR17408:SF0 | SUBFAMILY NOT NAMED |
| **Thupl.29380345s0016** | PANTHER | PTHR17408 | HISTONE RNA HAIRPIN-BINDING PROTEIN |
| **Thupl.29380345s0016** | PFAM | PF15247 | Histone RNA hairpin-binding protein RNA-binding domain |
| **Thupl.29380345s0016** | GO | GO:0003729 | mRNA binding |
| **Thupl.29380839s0001** | SIGNALP | SignalP-TM | - |
| **Thupl.29380839s0001** | PANTHER | PTHR34132:SF1 | - |
| **Thupl.29380839s0001** | PANTHER | PTHR34132 | - |
| **Thupl.29380857s0017** | PANTHER | PTHR16019:SF13 | - |
| **Thupl.29380857s0017** | PFAM | PF03909 | BSD domain |
| **Thupl.29380857s0017** | PANTHER | PTHR16019 | SYNAPSE-ASSOCIATED PROTEIN |
| **Thupl.29380857s0017** | KOG | KOG2690 | Uncharacterized conserved protein, contains BSD domain |
| **Thupl.29380857s0020** | PANTHER | PTHR31793:SF24 | - |

| | | | |
|---|---|---|---|
| **Thupl.29380857s0020** | PANTHER | PTHR31793 | FAMILY NOT NAMED |
| **Thupl.29380857s0020** | PFAM | PF03061 | Thioesterase superfamily |
| **Thupl.29380989s0001** | EC | 2.4.2.30 | NAD(+) ADP-ribosyltransferase |
| **Thupl.29380989s0001** | PFAM | PF00644 | Poly(ADP-ribose) polymerase catalytic domain |
| **Thupl.29380989s0001** | PANTHER | PTHR21328:SF2 | POLY (ADP-RIBOSE) POLYMERASE FAMILY, MEMBER 6 (PARP6) |
| **Thupl.29380989s0001** | GO | GO:0003950 | NAD+ ADP-ribosyltransferase activity |
| **Thupl.29380989s0001** | PANTHER | PTHR21328 | POLY (ADP-RIBOSE) POLYMERASE FAMILY, MEMBER (PARP) |
| **Thupl.29381152s0001** | PANTHER | PTHR12458:SF8 | - |
| **Thupl.29381152s0001** | KOG | KOG3213 | Transcription factor IIB |
| **Thupl.29381152s0001** | PANTHER | PTHR12458 | ORF PROTEIN |
| **Thupl.29381152s0001** | PFAM | PF05018 | Protein of unknown function (DUF667) |
| **Thupl.29381153s0002** | PANTHER | PTHR33384 | - |
| **Thupl.29381693s0002** | GO | GO:0004553 | hydrolase activity, hydrolyzing O-glycosyl compounds |
| **Thupl.29381693s0002** | PFAM | PF00332 | Glycosyl hydrolases family 17 |
| **Thupl.29381693s0002** | EC | 3.2.1.39 | Glucan endo-1,3-beta-D-glucosidase |
| **Thupl.29381693s0002** | SIGNALP | SignalP-noTM | - |
| **Thupl.29381693s0002** | PANTHER | PTHR32227 | FAMILY NOT NAMED |
| **Thupl.29381693s0002** | GO | GO:0005975 | carbohydrate metabolic process |
| **Thupl.29381693s0002** | PANTHER | PTHR32227:SF72 | - |
| **Thupl.29381804s0017** | PFAM | PF09493 | Tryptophan-rich protein (DUF2389) |
| **Thupl.29382099s0019** | PANTHER | PTHR35302 | - |
| **Thupl.29382099s0019** | PFAM | PF12046 | Protein of unknown function (DUF3529) |
| **Thupl.29382102s0017** | NA | NA | NA |
| **Thupl.29382330s0004** | EC | 2.5.1.75 | tRNA dimethylallyltransferase |
| **Thupl.29382330s0004** | KEGGORTH | K00791 | tRNA dimethylallyltransferase [EC:2.5.1.75] |
| **Thupl.29382330s0004** | KOG | KOG1384 | tRNA delta(2)-isopentenylpyrophosphate transferase |
| **Thupl.29382330s0004** | PANTHER | PTHR11088 | TRNA DELTA(2)-ISOPENTENYLPYROPHOSPHATE TRANSFERASE-RELATED |
| **Thupl.29382330s0004** | PFAM | PF01715 | IPP transferase |
| **Thupl.29382330s0004** | PANTHER | PTHR11088:SF50 | - |
| **Thupl.29382330s0004** | GO | GO:0008033 | tRNA processing |
| **Thupl.29382473s0007** | KOG | KOG1746 | Defender against cell death protein/oligosaccharyltransferase, epsilon subunit |
| **Thupl.29382473s0007** | EC | 2.4.99.18 | Dolichyl-diphosphooligosaccharide--protein glycotransferase |
| **Thupl.29382473s0007** | GO | GO:0008250 | oligosaccharyltransferase complex |
| **Thupl.29382473s0007** | PFAM | PF02109 | DAD family |

| | | | |
|---|---|---|---|
| **Thupl.29382473s0007** | KEGGORTH | K12668 | oligosaccharyltransferase complex subunit epsilon |
| **Thupl.29382473s0007** | GO | GO:0004579 | dolichyl-diphosphooligosaccharide-protein glycotransferase activity |
| **Thupl.29382473s0007** | PANTHER | PTHR10705 | DOLICHYL-DIPHOSPHOOLIGOSACCHARIDE--PROTEIN GLYCOSYLTRANSFERASE SUBUNIT DAD1 |
| **Thupl.29382473s0007** | GO | GO:0016021 | integral to membrane |

**Table S11. Kruskal-Wallis analysis of variance in π and $d_{XY}$ estimates in single-copy orthologs across subpopulations.**

| Subpopulation | Average π (SD) | Kruskal-Wallis $\chi^2$ | *p*-value |
|---|---|---|---|
| Northern-Coastal | 0.00267 (0.0120) | 5.79 | 0.0554 |
| Central | 0.00267 (0.0119) | | |
| Southern-Interior | 0.00266 (0.0135) | | |

| Subpop. comparison | Mean $d_{XY}$ (SD) | Kruskal-Wallis $\chi^2$ | *p*-value |
|---|---|---|---|
| Northern-Coastal/Central | 0.00272 (0.0130) | 2.12 | 0.347 |
| Northern-Coastal/Southern-Interior | 0.00277 (0.0123) | | |
| Central/Southern-Interior | 0.00280 (0.0127) | | |

**Table S12. Mean (SD) observed heterozygosity (H) and inbreeding coefficient (*F*) in the range-wide population (RWP) (*n* = 112) and each generation of selfing lines (FS – S4: *n* = 28; S5: *n* = 11).**

| | H | *F* |
|---|---|---|
| **RWP** | 0.219 (0.110) | 0.331 (0.181) |
| **FS** | 0.296 (0.174) | 0.00569 (0.0312) |
| **S1** | 0.148 (0.103) | 0.428 (0.0610) |
| **S2** | 0.0777 (0.0659) | 0.616 (0.0678) |
| **S3** | 0.0328 (0.0422) | 0.724 (0.0785) |
| **S4** | 0.0194 (0.0314) | 0.789 (0.0669) |
| **S5** | 0.00786 (0.0281) | 0.801 (0.0733) |

**Table S13. A) Pairwise Sign test for variance of median observed ($H_O$) and expected ($H_E$) heterozygosities in each generation of selfing lines (SLs), testing whether median $H_O$ is significantly lower than median $H_E$. B) One-tailed Wilcoxon signed rank test for variance of mean $H_O$ and $H_E$ in each generation of SLs, testing whether mean $H_O$ is significantly lower than mean $H_E$. C) One-sample *t*-test for variance of mean observed ($F_O$) and expected ($F_E$) inbreeding coefficients in each generation of SLs.**

**A**

| Generation | $H_O$ (median) | $H_E$ (median) | *S* | *p*-value |
|---|---|---|---|---|
| FS | 0.296 | - | - | - |
| S1 | 0.143 | 0.148 | 7338 | 0.0710 |
| S2 | 0.0714 | 0.0741 | 7843 | 0.000223 |
| S3 | 0.0357 | 0.0370 | 6072 | $< 2.2\times10^{-16}$ |
| S4 | 0 | 0.0185 | 6641 | $< 2.2\times10^{-16}$ |
| S5 | 0 | 0.00926 | 4544 | $< 2.2\times10^{-16}$ |

**B**

| Generation | $H_O$ (mean) | $H_E$ (mean) | *V* | *p*-value |
|---|---|---|---|---|
| FS | 0.296 | - | - | - |
| S1 | 0.148 | 0.148 | 54445812 | 0.0803 |
| S2 | 0.0777 | 0.0741 | 68311222 | 1 |
| S3 | 0.0328 | 0.0371 | 53919636 | $< 2.2\times10^{-16}$ |
| S4 | 0.0194 | 0.0185 | 65921119 | $< 2.2\times10^{-16}$ |
| S5 | 0.00786 | 0.00926 | 23642536 | $< 2.2\times10^{-16}$ |

**C**

| Generation | $F_O$ (mean) | $F_E$ (mean) | *t* | 95% CI | *p*-value |
|---|---|---|---|---|---|
| FS | 0.00569 | 0 | 0.966 | -0.00640, 0.0178 | 0.343 |
| S1 | 0.428 | 0.5 | -6.24 | 0.404, 0.452 | $1.12\times10^{-6}$ |
| S2 | 0.616 | 0.75 | -10.5 | 0.590, 0.642 | $5.29\times10^{-11}$ |
| S3 | 0.724 | 0.875 | -10.1 | 0.694, 0.755 | $1.05\times10^{-10}$ |
| S4 | 0.789 | 0.9375 | -11.7 | 0.763, 0.815 | $4.29\times10^{-12}$ |
| S5 | 0.801 | 0.96875 | -7.59 | 0.752, 0.850 | $1.87\times10^{-5}$ |

**Table S14. Gene Ontology (GO) categories for genes containing outlier SNPs (*n* = 83).**

| | GO category | GO ID | # of genes with GO category | Frequency (out of 13 genes) |
|---|---|---|---|---|
| * | GO:0016021 | integral to membrane | 3 | 0.230769 |
| | GO:0006355 | regulation of transcription, DNA-dependent | 2 | 0.153846 |
| | GO:0000287 | magnesium ion binding | 1 | 0.076923 |
| * | GO:0003690 | double-stranded DNA binding | 1 | 0.076923 |
| | GO:0003700 | sequence-specific DNA binding transcription factor activity | 1 | 0.076923 |
| * | GO:0004012 | phospholipid-translocating ATPase activity | 1 | 0.076923 |
| | GO:0004252 | serine-type endopeptidase activity | 1 | 0.076923 |
| * | GO:0004518 | nuclease activity | 1 | 0.076923 |
| | GO:0005515 | protein binding | 1 | 0.076923 |
| | GO:0005524 | ATP binding | 1 | 0.076923 |
| | GO:0005525 | GTP binding | 1 | 0.076923 |
| * | GO:0005743 | mitochondrial inner membrane | 1 | 0.076923 |
| | GO:0006281 | DNA repair | 1 | 0.076923 |
| | GO:0006508 | proteolysis | 1 | 0.076923 |
| | GO:0006629 | lipid metabolic process | 1 | 0.076923 |
| * | GO:0006850 | mitochondrial pyruvate transport | 1 | 0.076923 |
| | GO:0007165 | signal transduction | 1 | 0.076923 |
| | GO:0008270 | zinc ion binding | 1 | 0.076923 |
| * | GO:0010112 | regulation of systemic acquired resistance | 1 | 0.076923 |
| * | GO:0015095 | magnesium ion transmembrane transporter activity | 1 | 0.076923 |
| * | GO:0015693 | magnesium ion transport | 1 | 0.076923 |
| * | GO:0015914 | phospholipid transport | 1 | 0.076923 |
| | GO:0016491 | oxidoreductase activity | 1 | 0.076923 |
| | GO:0043531 | ADP binding | 1 | 0.076923 |
| | GO:0043565 | sequence-specific DNA binding | 1 | 0.076923 |
| | GO:0055085 | transmembrane transport | 1 | 0.076923 |
| | GO:0055114 | oxidation-reduction process | 1 | 0.076923 |

* Over-represented in outlier SNPS when compared to the full SNP set (*n* = 18,371) ($p < 0.05$, Fisher's Exact Test)

**Table S15. Geographic locations of origin of parent trees for selfing lines.**

| Parent | Selfing Line | Location | Latitude (° N) | Longitude (° W) | Elevation (m) |
|---|---|---|---|---|---|
| 491 | 1 | Bacchante Bay | 49.45 | -126.10 | 305 |
| 13 | 1 | O'Connell Lake | 50.35 | -127.70 | 122 |
| 150 | 6 | Tzoonie River | 49.87 | -123.73 | 450 |
| 49 | 6 | Misty Lake | 50.63 | -127.28 | 91 |
| 213 | 7 | Norrish Creek | 49.22 | -122.15 | 400 |
| 76 | 7 | Hathaway Creek | 50.58 | -127.82 | 105 |
| 142 | 8 | Bedwell Sound | 49.37 | -125.75 | 490 |
| 289 | 8 | Misty Lake | 50.63 | -127.30 | 60 |
| 263 | 10 | Narrows Inlet | 49.08 | -123.73 | 120 |
| 306 | 10 | Denad Creek | 50.55 | -128.00 | 25 |
| 341 | 13 | Utlah Creek | 50.28 | -127.43 | 183 |
| 323 | 13 | Koprino River | 50.55 | -127.85 | 280 |
| 367 | 16 | Henderson Lake | 49.10 | -125.02 | 200 |
| 381 | 16 | Sydney Inlet | 49.48 | -126.28 | 65 |
| 386 | 17 | Cypre River | 49.37 | -125.87 | 180 |
| 253 | 17 | Labouchere Channel | 52.03 | -127.25 | 152 |
| 403 | 19 | Eelstow Passage | 50.10 | -127.17 | 60 |
| 191 | 19 | Jervis Inlet | 50.12 | -123.78 | 98 |
| 424 | 20 | Chatham Channel | 50.52 | -126.25 | 75 |
| 318 | 20 | Ronning Creek | 50.57 | -128.27 | 105 |
| 435 | 21 | Chemainus | 48.93 | -123.75 | 115 |
| 15 | 21 | Mahatta Creek | 50.40 | -127.78 | 155 |
| 461 | 23 | Artlish Caves | 50.13 | -127.08 | 100 |
| 5 | 23 | Mahatta Creek | 50.40 | -127.78 | 122 |
| 344 | 26 | Victoria Lake | 50.05 | -127.42 | 457 |
| 313 | 26 | Brink Lake | 50.68 | -128.27 | 213 |
| 343 | 27 | Victoria Lake | 50.05 | -127.42 | 457 |
| 319 | 27 | Ronning Creek | 50.57 | -128.27 | 115 |
| 333 | 29 | Marble River | 50.25 | -127.33 | 244 |
| 312 | 29 | Brink Lake | 50.70 | -128.27 | 122 |

**Supplemental Datasets**

**Dataset S1. Primary transcripts in FASTA format.**

**Dataset S2. Predicted gene functions for annotated genes.**

**Dataset S3. List of western redcedar (WRC) single-copy ortholog (SCO) genes.**

**Dataset S4. Sequences used for validation of the genome annotation in FASTA format.**

**Dataset S5. Filtered SNP genotypes for the range-wide population (RWP) in VCF format (*n* = 18,371 SNPs).**

**Dataset S6. VEP SNP annotation for all annotated SNPS.**

**Dataset S7. WRC SNPs successfully mapped to putative linkage groups (LGs) on the *Sequoiadendron giganteum* genome (*n* = 26,140 SNPs).**

**Dataset S8. Corrected SNP genotypes for manually corrected and imputed selfing lines in PLINK binary format (*n* = 18,371 SNPs).**

**Dataset S9. Statistical analysis and outcomes for SNP fate based on expectations of genetic drift for all SNPs after four generations of selfing.**

**Dataset S10. Annotation of outlier SNPs deviating from expectations of genetic drift during selfing.**

**Supplemental Code**

**Code S1. Geographic map used in Figure 1.**

**Code S2. $F_{st}$ analysis, DAPC, PCA plots used in Figure 1.**

**Code S3. LD plots used in Figure 1. Code derived and adapted from a post made here: https://www.biostars.org/p/347796/.**

**Code S4. Analysis of pi, dxy in pixy, plots used in Figure 3.**

**Code S5. Manual correction of SNP data for complete selfing lines. Further fine-scale manual correction was carried out by hand.**

**Code S6. Analysis of heterozygosity, statistical tests, chi-squared test for analysis of loci under selection during selfing (outlier SNPs), plots used in Figure 4.**

**Code S7. Analysis and plots for inbreeding coefficients in Figure 4.**

**Code S8. Analysis of outlier SNPs during selfing, variant effects, statistical analyses of SNP consequences and GO categories, annotation of SNPs.**
