## Supplementary material for "The western redcedar genome reveals low genetic diversity in a self-compatible conifer": Code S1: geographic_maps.R

```
library(sf)
library(rnaturalearth)
library(rnaturalearthdata)
library(rnaturalearthhires)
library(ggrepel)
library(ggmap)
library(maps)
library(ggsci)
library(ggspatial)
library(elevatr)
library(tidyverse)
source("publication_theme.r")

### Load data and set up map properties ####
# Typically, load from NaturalEarth, but lately the database has been down so can also download and load it manually
land <- ne_download(scale = "large", returnclass = "sf", category = "physical", type = "land")
# land <- ne_load(scale = "large", returnclass = "sf", category = "physical", type = "land", destdir = getwd())
# rivers <- ne_download(scale = "large", returnclass = "sf", category = "physical", type = "rivers_lake_centerlines")
# lakes <- ne_download(scale = "large", returnclass = "sf", category = "physical", type = "lakes")
# ocean <- ne_download(scale = "large", returnclass = "sf", category = "physical", type = "ocean")
canada <- ne_states(country = "canada", returnclass = "sf")
usa <- ne_states(country = "united states of america", returnclass = "sf")

# Load in geographic coordinates and cluster associations
sites <- read.delim("RWP_subpops_geog_locations.txt", header = T)
RWP_popmap_subpops <- read.table("popmap_RWP_subpops.txt")

RWP_popmap_subpops <- RWP_popmap_subpops %>% 
  arrange(V1) %>% 
  rename(Parent = V1)

sites_RWP_popmap_subpops <- merge(sites, RWP_popmap_subpops, by = "Parent")

### Plots ####
# Colour palette
col <- c(pal_nejm()(8))

# Figure 1A
map <- ggplot() +
  theme_Publication() +
  geom_sf(data = land, fill ="antiquewhite") +
  geom_sf(data = canada, fill ="antiquewhite") +
  geom_sf(data = usa, fill ="antiquewhite") +
  # geom_sf(data = lakes, fill = "lightblue") +
  # geom_sf(data = rivers, colour = "lightblue") +
  # geom_sf(data = ocean, fill = "aliceblue") +
  # geom_text_repel(data = sites, aes(x = Longitude, y = Latitude), size = 2.5, label = sites$Parent, max.overlaps = 50) +
  geom_point(data = sites_RWP_popmap_subpops, aes(x = Longitude, y = Latitude, fill = subpop), shape = 23, size = 2) +
  scale_fill_manual(name="Subpopulation",
                    breaks=c("Northern_Coastal", "Central", "Southern_Interior"),
                    labels=c("Northern-Coastal", "Central", "Southern-Interior"),
                    values = col) +
  annotation_scale(location = "bl", width_hint = 0.5) +
  annotation_north_arrow(location = "bl", which_north = "true", pad_x = unit(0.75, "in"), pad_y = unit(0.5, "in"), style = north_arrow_fancy_orienteering) +
  coord_sf(xlim = c(-137, -117), ylim = c(39, 55), expand = FALSE) +
  theme(panel.grid.major = element_line(color = gray(0.5), linetype = "dashed",
                                        size = 0.5), panel.background = element_rect(fill = "aliceblue")) +
  theme(legend.position = c(0.24, 0.31), legend.background = element_rect(colour = "black"))


ggsave("RWP_geog_locations.tiff", map, width = 7, height = 8)
```
