## Supplementary material for "The western redcedar genome reveals low genetic diversity in a self-compatible conifer": Code S2: genetic_differentiation.R

```
library(adegenet)
library(geodist)
library(hierfstat)
library(vcfR)
library(pegas)
library(reshape2)
library(ggsci)
library(apex)
library(mmod)
library(factoextra)
library(FactoMineR)
library(poppr)
library(ggpubr)
library(rstatix)
library(tidyverse)

source("publication_theme.r")

# Read in VCF
RWP_sco_intergenic_vcf <- read.vcfR("sco_intergenic_snps.recode.vcf")


# Read in popmap
subpops_popmap <- read.table("popmap_RWP_subpops.txt")

# Convert VCF to genind

RWP_sco_intergenic_genind <- vcfR2genind(RWP_sco_intergenic_vcf, pop = subpops_popmap$V2)
RWP_sco_intergenic_genind_no_pop <- vcfR2genind(RWP_sco_intergenic_vcf, pop = subpops_popmap$V1)


# Geographic locations for Mantel test
all_geo_dist <- read.table("RWP_subpops_geog_locations.txt", header = T, sep = "\t")
subpops_geo_dist <- read.table("RWP_subpops_average_coord.txt", header = T, row.names = 1)

# Stats

RWP_sco_intergenic_stats <- genind2hierfstat(RWP_sco_intergenic_genind)
RWP_sco_intergenic_stats_no_pop <- genind2hierfstat(RWP_sco_intergenic_genind_no_pop)

### FST tests ####
subpops <- subpops_popmap$V2

basic.stats(RWP_sco_intergenic_genind)
wc(RWP_sco_intergenic_stats)
parents_div <- summary(RWP_sco_intergenic_genind)
loci <- RWP_sco_intergenic_stats[, -1]
# global_varcomp_RWP_mac_3_r2_01_sco <- varcomp.glob(levels = data.frame(subpops, pops), loci, diploid = T)
g_test_sco <- test.g(loci, level = subpops, nperm = 1000)
# between_test_RWP_mac_3_r2_01_sco <- test.between(loci, test.lev = subpops, rand.unit = pops, nperm = 999)
genet.dist(RWP_sco_intergenic_genind, method = "WC84")


### Mantel tests ####
all_geo_dist <- all_geo_dist %>% 
  select(Parent, Longitude, Latitude) %>% 
  column_to_rownames(var = "Parent")

# Exact geo dist
all_geo_dist_mat <- geodist(all_geo_dist, measure = "geodesic", )
rownames(all_geo_dist_mat) <- row.names(all_geo_dist)
colnames(all_geo_dist_mat) <- row.names(all_geo_dist)

all_geo_dist_mat[upper.tri(all_geo_dist_mat)] <- ""
all_geo_dist_mat[upper.tri(all_geo_dist_mat,diag=TRUE)] <- ""

# Euclidean gen dist
all_gen_dist_mat <- dist(RWP_sco_intergenic_genind, method = "euclidean", diag = F, upper = F)

all_geo_dist_vector <- c(all_geo_dist_mat)

RWP_sco_intergenic_genpop <- genind2genpop(RWP_sco_intergenic_genind_no_pop)
Dgen <- dist.genpop(RWP_sco_intergenic_genpop, method = 2)
Dgeo <- dist(cbind(all_geo_dist$Longitude, all_geo_dist$Latitude))

all_gen_dist_vector <- c(Dgen)

all_geo_dist_vector <- as.numeric(all_geo_dist_vector[all_geo_dist_vector != ""])

geo_gen_dist_df <- data.frame(geo = all_geo_dist_vector/1000, gen = all_gen_dist_vector)

ibd <- mantel.randtest(Dgen, Dgeo, nrepet = 9999)

ibd

# Plot individual correlation ge gen dist
ggplot(geo_gen_dist_df, aes(x = geo, y = gen)) +
  theme_Publication() +
  geom_point() +
  xlab("Geographic distance (km)") +
  ylab("Genetic distance") +
  geom_smooth(method = "lm")

# ggsave("mantel_indiv_plot.svg", dpi = 300, width = 10, height = 9)  

m <- lm(gen ~ geo, geo_gen_dist_df)
summary(m)
anova(m)

# Mantel test subpops
RWP_sco_intergenic_genpop <- genind2genpop(RWP_sco_intergenic_genind)

Dgen <- dist.genpop(RWP_sco_intergenic_genpop, method = 2)
Dgeo <- dist(cbind(subpops_geo_dist$Longitude, subpops_geo_dist$Latitude))

ibd <- mantel.randtest(Dgen, Dgeo, nrepet = 9999)

ibd
plot(ibd)

mcor <- vegan::mantel.correlog(Dgen, Dgeo)
mcor
plot(mcor)

Dgen <- as.numeric(Dgen)
Dgeo <- as.numeric(Dgeo)

par(mar=c(4,4,0,0))
dens <- MASS::kde2d(Dgeo, Dgen, n=300)
myPal <- colorRampPalette(c("white","blue","gold","orange","red"))
plot(Dgeo, Dgen, pch=20, cex=1,  
     xlab="Geographic Distance", ylab="Genetic Distance")
image(dens, col=transp(myPal(300), 0.7), add=TRUE)
abline(lm(Dgen ~ Dgeo))

lines(loess.smooth(log(Dgeo), Dgen), col="red")

# Generating PCA for Figure 1C

RWP_sco_intergenic_scaled <- scaleGen(RWP_sco_intergenic_genind, NA.method = "mean")
pca_RWP_sco_intergenic <- dudi.pca(RWP_sco_intergenic_scaled,cent=FALSE,scale=FALSE,scannf=FALSE,nf=3)
barplot(pca_RWP_sco_intergenic$eig[1:50],main="PCA eigenvalues", col=heat.colors(50))

col <- c(pal_nejm()(8))

get_eigenvalue(pca_RWP_sco_intergenic)
fviz_eig(pca_RWP_sco_intergenic, addlabels = T)
p <- fviz_pca_ind(pca_RWP_sco_intergenic,
                  axes = c(1, 2),
                  col.ind = subpops_popmap$V2, # color by groups
                  pointsize = 4,
                  palette = col,
                  # addEllipses = TRUE, # Concentration ellipse
                  # ellipse.type = "confidence",
                  # legend.title = "Population",
                  # repel = T,
                  label = "none",
                  mean.point = FALSE,
                  pointshape = 19
)

pca_plot <- p + 
  theme_Publication() +
  labs(title = NULL) +
  xlab("PC 1 (3.73%)") + 
  ylab("PC 2 (1.63%)") +
  scale_color_manual(name="Subpopulation",
                     breaks=c("Northern_Coastal", "Central", "Southern_Interior"),
                     labels=c("Northern-Coastal", "Central", "Southern-Interior"),
                     values = col) +
  theme(legend.position = c(0.24, 0.16), legend.background = element_rect(colour = "black"))

pca_plot
# ggsave("pca_plot_pops_all_snps_subpops.tiff", pca_plot, width = 7, height = 8, dpi = 300)


### DAPC ###########################################

## DAPC cross-validation ####
RWP_sco_intergenic_cluster_xdapc <- xvalDapc(tab(RWP_sco_intergenic_genind, NA.method = "mean"), pop(RWP_sco_intergenic_genind), n.pca = 1:70, n.rep = 1000, parallel = "snow", ncpus = 4L)

RWP_sco_intergenic_cluster_xdapc$DAPC


### DAPC K selection ####
# Adapted from https://grunwaldlab.github.io/Population_Genetics_in_R/
maxK <- 10
myMat <- matrix(nrow=10, ncol=maxK)
colnames(myMat) <- 1:ncol(myMat)
for(i in 1:nrow(myMat)){
  grp <- find.clusters(RWP_sco_intergenic_genind, n.pca = 200, choose.n.clust = FALSE,  max.n.clust = maxK)
  myMat[i,] <- grp$Kstat
}

my_df <- melt(myMat)
colnames(my_df)[1:3] <- c("Group", "K", "BIC")
my_df$K <- as.factor(my_df$K)
head(my_df)

p1 <- ggplot(my_df, aes(x = K, y = BIC))
p1 <- p1 + geom_boxplot()
p1 <- p1 + theme_pubr()
p1 <- p1 + xlab("Number of groups (K)")
p1

my_k <- 2:3

grp_l <- vector(mode = "list", length = length(my_k))
dapc_l <- vector(mode = "list", length = length(my_k))

for(i in 1:length(dapc_l)){
  set.seed(200)
  grp_l[[i]] <- find.clusters(RWP_sco_intergenic_genind, n.pca = 200, n.clust = my_k[i])
  dapc_l[[i]] <- dapc(RWP_sco_intergenic_genind, pop = grp_l[[i]]$grp, n.pca = 22, n.da = my_k[i])
  #  dapc_l[[i]] <- dapc(gl_rubi, pop = grp_l[[i]]$grp, n.pca = 3, n.da = 2)
}

my_df <- as.data.frame(dapc_l[[ length(dapc_l) ]]$ind.coord)
my_df$Group <- dapc_l[[ length(dapc_l) ]]$grp
head(my_df)

my_pal <- RColorBrewer::brewer.pal(n=8, name = "Dark2")

col <- c(pal_nejm()(3), "#7876B1FF", "#6F99ADFF")

p2 <- ggplot(my_df, aes(x = LD1, y = LD2, color = Group, fill = Group))
p2 <- p2 + geom_point(size = 4, shape = 21)
p2 <- p2 + theme_pubr()
p2 <- p2 + scale_color_manual(values=col)
p2 <- p2 + scale_fill_manual(values=col)
p2

tmp <- as.data.frame(dapc_l[[1]]$posterior)
tmp$K <- my_k[1]
tmp$Isolate <- rownames(tmp)
tmp <- melt(tmp, id = c("Isolate", "K"))
names(tmp)[3:4] <- c("Group", "Posterior")
tmp$Region <- subpops_popmap$V2
my_df <- tmp

for(i in 2:length(dapc_l)){
  set.seed(200)
  tmp <- as.data.frame(dapc_l[[i]]$posterior)
  tmp$K <- my_k[i]
  tmp$Isolate <- rownames(tmp)
  tmp <- melt(tmp, id = c("Isolate", "K"))
  names(tmp)[3:4] <- c("Group", "Posterior")
  tmp$Region <- subpops_popmap$V2
  
  my_df <- rbind(my_df, tmp)
}

grp.labs <- paste("K =", my_k)
names(grp.labs) <- my_k

p3 <- ggplot(my_df, aes(x = Isolate, y = Posterior, fill = Group))
p3 <- p3 + geom_bar(stat = "identity")
p3 <- p3 + facet_grid(K ~ Region, scales = "free_x", space = "free", 
                      labeller = labeller(K = grp.labs))
p3 <- p3 + theme_pubr()
p3 <- p3 + ylab("Posterior membership probability")
p3 <- p3 + theme(legend.position='none')
#p3 <- p3 + scale_color_brewer(palette="Dark2")
p3 <- p3 + scale_fill_manual(values=col)
p3 <- p3 + theme(axis.text.x = element_text(angle = 90, hjust = 1, size = 8))
p3 <- p3 + xlab("Tree")
p3

library(ggpubr)
ggarrange(ggarrange(p1,
                    p2,
                    ncol = 2, labels = c("A", "B")),
          p3,
          nrow = 2,
          labels = c("", "C"),
          heights = c(1, 2)
)

# ggsave("dapc_multiplot.svg", dpi = 300, width = 15, height = 10)
```
