## Supplementary material for "The western redcedar genome reveals low genetic diversity in a self-compatible conifer": Code S3: LD_plots.R

```
rm(list=ls())

# load libraries
library(tidyverse)

source("publication_theme.r")
library(extrafont)

# define plotting functions
#' @title plotPairwiseLD
#' @description Plots R2 heatmap across the chromosome (like Haploview)
#' @param dfr A data.frame with minimum CHR_A, BP_A, CHR_B, BP_B and R2.
#' An output from tomahawk works.
#' @param chr A chromosome name.
#' @param xlim A two number vector specifying min and max x-axis limits. Any one or both can be defaulted by specifying NA.
#' @param ylim A two number vector specifying min and max y-axis limits. Any one or both can be defaulted by specifying NA.
#' @param minr2 A value between 0 and 1. All SNPs with R2 value below this 
#' value is excluded from plot.
#' 
#' #' Adapted from BioStars forum user rmf
plotPairwiseLD <- function(dfr,chr,xlim=c(NA,NA),ylim=c(NA,NA),minr2) {
  if(missing(dfr)) stop("Input data.frame 'dfr' missing.")
  
  if(!missing(chr)) {
    ld <- filter(ld,CHR_A==get("chr") & CHR_B==get("chr"))
  }
  ld <- filter(ld,BP_A<BP_B)
  
  if(!missing(minr2)) {
    ld <- filter(ld,R2>get("minr2"))
  }
  
  ld <- ld %>% arrange(R2)
  
  ld$x <- ld$BP_A+((ld$BP_B-ld$BP_A)/2)
  ld$y <- ld$BP_B-ld$BP_A
  ld$r2c <- cut(ld$R2,breaks=seq(0,1,0.2),labels=c("0 - 0.2","0.2 - 0.4",
                                                   "0.4 - 0.6","0.6 - 0.8",
                                                   "0.8 - 1.0"))
  ld$r2c <- factor(ld$r2c,levels=rev(c("0 - 0.2","0.2 - 0.4",
                                       "0.4 - 0.6","0.6 - 0.8",
                                       "0.8 - 1.0")))
  
  ggplot(ld,aes(x=x,y=y,col=r2c))+
    geom_point(shape=18,size=0.6,alpha=0.8)+
    scale_color_manual(values=c("#ca0020","#f4a582","#d1e5f0","#67a9cf","#2166ac"))+
    scale_x_continuous(limits=xlim, breaks = c(0, 1e06, 2e06, 3e06, 4e06, 5e06, 6e06, 7e06, 8e06, 9e06,
                                               10e06, 11e06, 12e06, 13e06, 14e06, 15e06, 16e06, 17e06),
                       labels = c(0, 1, 2, 3, 4, 5, 6, 7, 8, 9, 10, 11, 12, 13, 14, 15, 16, 17)) +
    scale_y_continuous(limits=ylim, breaks = c(0, 1e06, 2e06, 3e06, 4e06, 5e06, 6e06, 7e06, 8e06, 9e06,
                                             10e06, 11e06, 12e06, 13e06, 14e06, 15e06, 16e06, 17e06),
                       labels = c(0, 1, 2, 3, 4, 5, 6, 7, 8, 9, 10, 11, 12, 13, 14, 15, 16, 17)) +
    guides(colour=guide_legend(title=expression(LD~(italic(r^{2}))),override.aes=list(shape=20,size=8)))+
    labs(x="Position (Mbp)",y="Distance (Mbp)")+
    theme_Publication()+
    # theme(panel.border=element_blank(),
          # axis.ticks=element_blank()) +
    theme(panel.grid.major = element_blank(), panel.grid.minor = element_blank()) +
    theme(legend.position = c(0.9,0.85)) %>%
    return()
}

#' @title plotDecayLD
#' @description Plots R2 heatmap across the chromosome (like Haploview)
#' @param dfr A data.frame with minimum CHR_A, BP_A, CHR_B, BP_B and R2.
#' An output from tomahawk works.
#' @param chr A chromosome name.
#' @param xlim A two number vector specifying min and max x-axis limits. Any one or both can be defaulted by specifying NA.
#' @param ylim A two number vector specifying min and max y-axis limits.  Any one or both can be defaulted by specifying NA.
#' @param avgwin An integer specifying window size. Mean R2 is computed within windows.
#' @param minr2 A value between 0 and 1. All SNPs with R2 value below this 
#' value is excluded from plot.
#' 
#' Adapted from BioStars forum user rmf
plotDecayLD <- function(dfr,chr,xlim=c(NA,NA),ylim=c(NA,NA),avgwin=0,minr2) {
  if(missing(dfr)) stop("Input data.frame 'dfr' missing.")
  
  if(!missing(chr)) {
    ld <- filter(ld,CHR_A==get("chr") & CHR_B==get("chr"))
  }
  ld <- filter(ld,BP_A<BP_B)
  
  if(!missing(minr2)) {
    ld <- filter(ld,R2>get("minr2"))
  }
  
  ld <- ld %>% arrange(R2)
  
  ld$dist <- abs(ld$BP_B-ld$BP_A)
  
  if(avgwin>0) {
    ld$distc <- cut(ld$dist,breaks=seq(from=min(ld$dist),to=max(ld$dist),by=avgwin))
    ld <- ld %>% group_by(distc) %>% summarise(dist=mean(dist),R2=mean(R2))
  }
  
  ggplot(ld,aes(x=dist,y=R2))+
    theme_Publication()+
    geom_point(shape=20,size=0.15,alpha=0.1)+
    geom_path(data = ld.df, aes(x = distance, y = fpoints), colour = "#BC3C29FF", size= 0.9) +
    # geom_path(data = ld.df_05, aes(x = distance, y = fpoints), colour = "#0072B5FF", size= 0.9) +
    # geom_path(data = ld.df_1, aes(x = distance, y = fpoints), colour = "#E18727FF", size= 0.9) +
    geom_hline(yintercept = 0.2, linetype = "dashed", color = "#7876B1FF") +
    geom_hline(yintercept = 0.1, linetype = "dashed", color ="#EE4C97FF") +
    # geom_smooth() +
    scale_x_continuous(limits=xlim, breaks = c(0, 1e06, 2e06, 3e06, 4e06, 5e06, 6e06, 7e06, 8e06, 9e06,
                                               10e06, 11e06, 12e06, 13e06, 14e06, 15e06, 16e06, 17e06),
                       labels = c(0, 1, 2, 3, 4, 5, 6, 7, 8, 9, 10, 11, 12, 13, 14, 15, 16, 17)) +
    scale_y_continuous(limits=ylim)+
    labs(x="Distance (Mbp)",y=expression(bold(LD~(italic(r^{2})))))+
   # theme(panel.border=element_blank(),
    #      axis.ticks=element_blank()) +
    theme(panel.grid.major = element_blank(), panel.grid.minor = element_blank()) %>%
    return()
}

# read file

ld <- read.table("rwp_maf05.ld", header = T)
ld$CHR_A <- factor(ld$CHR_A)
ld$CHR_B <- factor(ld$CHR_B)

distance<-abs(ld$BP_B - ld$BP_A)
LD.data<-ld$R2
n<-224
HW.st<-c(C=0.1)
HW.nonlinear<-nls(LD.data~((10+C*distance)/((2+C*distance)*(11+C*distance)))*(1+((3+C*distance)*(12+12*C*distance+(C*distance)^2))/(n*(2+C*distance)*(11+C*distance))),start=HW.st,control=nls.control(maxiter=100))
tt<-summary(HW.nonlinear)
new.rho<-tt$parameters[1]
fpoints<-((10+new.rho*distance)/((2+new.rho*distance)*(11+new.rho*distance)))*(1+((3+new.rho*distance)*(12+12*new.rho*distance+(new.rho*distance)^2))/(n*(2+new.rho*distance)*(11+new.rho*distance)))

ld.df<-data.frame(distance, fpoints)
ld.df<-ld.df[order(ld.df$distance),]

## Plots for Figure 2
pairwise_LD <- plotPairwiseLD(ld, minr2 = 0)
# ggsave("pairwise_ld.tiff", pairwise_LD, height = 6, width = 7)

LD_decay <- plotDecayLD(ld, avgwin = 10)
# ggsave("ld_decay_avg_win_10_1_decay.tiff", LD_decay, height = 6, width = 7)

## LD decay thresholds, half-decay ####

summary(LD.data)
filter(ld.df, fpoints < 0.60001 & fpoints > 0.49997)
filter(ld.df, fpoints < 0.20001 & fpoints > 0.19997)


filter(ld.df, fpoints < 0.10011 & fpoints > 0.09999)


df<-data.frame(distance,fpoints)
maxld<-quantile(LD.data, 0.9)

h.decay<-maxld/2
half.decay.distance<-df$distance[which.min(abs(df$fpoints-h.decay))]
half.decay.distance
```
