## Supplementary material for "The western redcedar genome reveals low genetic diversity in a self-compatible conifer": Code S4: nucleotide_diversity_analysis.R

```
rm(list=ls())

library(rstatix)
library(gghighlight)
library(tidyverse)


### Reading in pi files ####
# Pi 10 Kb window for whole RWP
all_snps_10k_window_one_pop_pi <- read.table("sco_intergenic_10k_window/sco_intergenic_10k_window_pi.txt", header = T)

# Pi SCO genes for whole pop, subpops
all_snps_primary_genes_sco_one_pop_pi <- read.table("sco_intergenic_all_one_pop/sco_intergenic_all_one_pop_pi.txt", header = T)
all_snps_primary_genes_sco_subpops_pi <- read.table("sco_intergenic_subpops/sco_intergenic_subpops_pi.txt", header = T)

# Pi SCO genes for zero- and four-fold degenerate sit
all_snps_4_fold_gff_one_pop_pi <- read.table("sco_intergenic_four_fold/sco_intergenic_four_fold_pi.txt", header = T)
all_snps_0_fold_gff_one_pop_pi <- read.table("sco_intergenic_zero_fold/sco_intergenic_zero_fold_pi.txt", header = T) 

# DXY SCO genes subpops
all_snps_primary_genes_sco_subpops_dxy <- read.table("sco_intergenic_subpops/sco_intergenic_subpops_dxy.txt", header = T)


### Data summaries ####
# 10k window
sum(all_snps_10k_window_one_pop_pi$count_diffs, na.rm = T)/sum(all_snps_10k_window_one_pop_pi$count_comparisons, na.rm = T)
sd(all_snps_10k_window_one_pop_pi$count_diffs/all_snps_10k_window_one_pop_pi$count_comparisons, na.rm = T)

# Primary gene SCO
sum(all_snps_primary_genes_sco_one_pop_pi$count_diffs, na.rm = T)/sum(all_snps_primary_genes_sco_one_pop_pi$count_comparisons, na.rm = T)
sd(all_snps_primary_genes_sco_one_pop_pi$count_diffs/all_snps_primary_genes_sco_one_pop_pi$count_comparisons, na.rm = T)

table(all_snps_primary_genes_sco_one_pop_pi$avg_pi)

# Primary gene SCO subpops
all_snps_primary_genes_sco_subpops_pi %>% 
  group_by(pop) %>%
  group_map(~ sum(.x$count_diffs, na.rm = T)/sum(.x$count_comparisons, na.rm = T))

all_snps_primary_genes_sco_subpops_pi %>% 
  group_by(pop) %>%
  group_map(~ sd(.x$count_diffs/.x$count_comparisons, na.rm = T))
  
# Dxy
all_snps_primary_genes_sco_subpops_dxy <- all_snps_primary_genes_sco_subpops_dxy %>% 
  filter(no_sites >= 1) %>%
  unite(pop_comparison, c("pop1", "pop2"))

all_snps_primary_genes_sco_subpops_dxy %>% 
  group_by(pop_comparison) %>% 
  group_map(~ sum(.x$count_diffs)/sum(.x$count_comparisons))

all_snps_primary_genes_sco_subpops_dxy %>% 
  group_by(pop_comparison) %>% 
  group_map(~ sd(.x$count_diffs/.x$count_comparisons))

# 0-fold
sum(all_snps_0_fold_gff_one_pop_pi$count_diffs, na.rm = T)/sum(all_snps_0_fold_gff_one_pop_pi$count_comparisons, na.rm = T)
sd(all_snps_0_fold_gff_one_pop_pi$count_diffs/all_snps_0_fold_gff_one_pop_pi$count_comparisons, na.rm = T)

# 4 -fold

sum(all_snps_4_fold_gff_one_pop_pi$count_diffs, na.rm = T)/sum(all_snps_4_fold_gff_one_pop_pi$count_comparisons, na.rm = T)
sd(all_snps_4_fold_gff_one_pop_pi$count_diffs/all_snps_4_fold_gff_one_pop_pi$count_comparisons, na.rm = T)

### Plots #####

# Make scaffolds a factor
all_snps_primary_genes_sco_one_pop_pi$chromosome <- factor(all_snps_primary_genes_sco_one_pop_pi$chromosome)

# Remove NA
all_pi_no_NA <- all_snps_primary_genes_sco_one_pop_pi %>% 
  filter(!(avg_pi == "NA"))

# Outliers
pi_no_outliers <- all_snps_primary_genes_sco_one_pop_pi[all_snps_primary_genes_sco_one_pop_pi$avg_pi < quantile(all_snps_primary_genes_sco_one_pop_pi$avg_pi, probs = 0.99, na.rm = T),]
pi_no_outliers <- pi_no_outliers[complete.cases(pi_no_outliers),]

dxy_no_outliers <- all_snps_primary_genes_sco_subpops_dxy[all_snps_primary_genes_sco_subpops_dxy$avg_dxy < quantile(all_snps_primary_genes_sco_subpops_dxy$avg_dxy, probs = 0.99, na.rm = T),]
dxy_no_outliers <- dxy_no_outliers[complete.cases(dxy_no_outliers),]

# Top 1% of pi estimates
top_1 <- all_pi_no_NA[all_pi_no_NA$avg_pi > quantile(all_pi_no_NA$avg_pi,prob=1-1/100, na.rm = T),]

# Figure S4
## Linkage groups
LG_old_snps <- read.csv("WRC_linkage_group_SNPs_nucmer.csv")
snps <- read.table("all_filtered_snp_ids.txt")
LG_new_snps <- LG_old_snps %>% 
  filter(SNP %in% snps$V1)

LG_new_snps$chr <- factor(LG_new_snps$chr)
LG_new_snps$WRC_scaffold <- factor(LG_new_snps$WRC_scaffold)

LG_new_snps <- LG_new_snps %>% 
  arrange(chr,accurate_pos) %>% 
  rename(chromosome = WRC_scaffold)

LG_new_snps$SNP <- factor(LG_new_snps$SNP, levels=unique(LG_new_snps$SNP))

merged_pi_LG <- merge(LG_new_snps, all_snps_primary_genes_sco_one_pop_pi, by = "chromosome")

snps_in_LG <- merged_pi_LG %>% 
  
  # Compute chromosome size
  group_by(chr) %>% 
  summarise(chr_len=max(accurate_pos)) %>% 
  
  # Calculate cumulative position of each chromosome
  mutate(tot=cumsum(as.numeric(chr_len))-as.numeric(chr_len)) %>%
  select(-chr_len) %>%
  
  # Add this info to the initial dataset
  left_join(merged_pi_LG, ., by=c("chr"="chr")) %>%
  
  # Add a cumulative position of each SNP
  arrange(chr, accurate_pos) %>%
  mutate(BPcum=accurate_pos+tot) %>% 
  mutate(chrom_color_group = case_when(as.numeric(chr) %% 2 != 0 ~ "even",
                                       chr == "X" ~ "even",
                                       TRUE ~ "odd" )) %>%
  mutate(chromosome = factor(chr))


axisdf = snps_in_LG %>% group_by(chr) %>% summarize(center=( max(BPcum) + min(BPcum)) / 2 )

LG_plot <- ggplot(snps_in_LG, aes(x=BPcum, y = avg_pi, color = chrom_color_group)) +
  
  # Show all points
  geom_point(color = "#BC3C29FF", alpha = 0.8, size = 2.5) +
  # scale_color_nejm() +
  # custom X axis:
  scale_x_continuous(label = axisdf$chr, breaks= axisdf$center) +
  # scale_y_continuous(expand = c(0, 0) ) +     # remove space between plot area and x axis
  # geom_vline(xintercept = c(888232888, 1683028619, 2539001849, 3225909214, 3875153489, 4496253278, 5139632346, 5771913333, 6413980409, 7022253572), linetype = "dashed") +
  xlab("Linkage Group") +
  ylab(expression(bold(pi))) +
  
  # Custom the theme:
  theme_Publication() +
  theme( 
    legend.position="top",
    panel.border = element_blank(),
    panel.grid.major.x = element_blank(),
    panel.grid.minor.x = element_blank()
  ) +
  guides(colour = "none") +
  gghighlight(avg_pi > quantile(avg_pi,prob=1-1/100, na.rm = T), keep_scales = T,
              unhighlighted_params = list(colour = ifelse(snps_in_LG$chrom_color_group == "even", "grey50", "black"), alpha = 0.5)) +
  scale_color_manual(values = c("grey50", "black"))

ggsave("test.tiff", height = 8, width = 14, dpi = 300)

# Figure 3A - inlay
ggplot(all_snps_primary_genes_sco_one_pop_pi, aes(x = pop, y = avg_pi)) +
  geom_violin() +
  geom_boxplot(outlier.alpha = 0.3, width = 0.15) +
  theme_Publication() +
  theme(axis.text.x = element_blank(), axis.ticks = element_blank(), axis.title.x = element_blank(), axis.title.y = element_blank()) +
  scale_y_continuous(breaks = c(0, 0.5), limits = c(0,0.5))

# ggsave("all_pi_violinplot_inlay.tiff", dpi = 300, height = 6, width = 6)

# Figure 3A - main
ggplot(pi_no_outliers, aes(x = pop, y = avg_pi)) +
  geom_violin(scale = "area") +
  geom_boxplot(outlier.alpha = 0.3, width = 0.15) +
  theme_Publication() +
  xlab("Range-wide population") +
  ylab(expression(bold(pi))) +
  theme(axis.text.x = element_blank(), axis.ticks = element_blank())

# ggsave("all_pi_violinplot.tiff", dpi = 300, height = 6, width = 6)

# Figure 3B - inlay
ggplot(all_snps_primary_genes_sco_subpops_dxy, aes(x = pop_comparison, y = avg_dxy)) +
  geom_violin() +
  geom_boxplot(outlier.alpha = 0.3, width = 0.15) +
  theme_Publication() +
  ylab(expression(bold(italic(d)[XY]))) +
  scale_x_discrete(name="Subpopulation",
                   breaks=c("Northern_Coastal_Central", "Central_Southern_Interior", "Northern_Coastal_Southern_Interior"),
                   labels=c("Northern-Coastal/\nCentral", "Central/\nSouthern-Interior", "Northern-Coastal/\nSouthern-Interior")) +
  theme(axis.text.x = element_blank(), axis.ticks = element_blank(), axis.title.x = element_blank(), axis.title.y = element_blank()) +
  scale_y_continuous(breaks = c(0, 0.5), limits = c(0,0.5))

# ggsave("all_dxy_violinplot_inlay.tiff", dpi = 300, height = 6, width = 7)

# Figure 3B - main
ggplot(dxy_no_outliers, aes(x = pop_comparison, y = avg_dxy)) +
  geom_violin() +
  geom_boxplot(outlier.alpha = 0.3, width = 0.15) +
  theme_Publication() +
  ylab(expression(bold(italic(d)[XY]))) +
  scale_x_discrete(name="Subpopulation",
                   breaks=c("Northern_Coastal_Central", "Central_Southern_Interior", "Northern_Coastal_Southern_Interior"),
                   labels=c("Northern-Coastal/\nCentral", "Central/\nSouthern-Interior", "Northern-Coastal/\nSouthern-Interior"))

# ggsave("all_dxy_violinplot.tiff", dpi = 300, height = 6, width = 8)


# # Combine zero- and four-fold estimates
# all_snps_0_4_gff <- rbind(data.frame(pi = all_snps_0_fold_gff_one_pop_pi$avg_pi, degen = rep("zero", 9259)), data.frame(pi = all_snps_4_fold_gff_one_pop_pi$avg_pi, degen = rep("four", 7131)))
# 
# # unused plot
# ggplot(all_snps_0_4_gff, aes(x = degen, y = pi)) +
#   geom_violin() +
#   geom_boxplot(outlier.alpha = 0.3, width = 0.15) +
#   theme_Publication() +
#   ylab(expression(bold(pi))) +
#   scale_x_discrete(name="Sites",
#                    breaks=c("four", "zero"),
#                    labels=c(expression(paste(bold(pi[4]))), expression(paste(bold(pi[0]))))) +
#   scale_y_continuous(limits = c(0,0.0015))
# ggsave("0-4-fold_pi.tiff", height = 6, width = 7)

### Annotations for top 1% ####  
# Load annotations
gene_annotations <- read.delim("gene_functions.txt", skip = 1, header = F,
                               sep = "\t")

# Load GFF
gff <- read.table("Tplicatav3.1.primaryTrs.gff3.gz")

# Create variable for "window"
gff_genes <- gff %>% 
  filter(V3 == "gene") %>% 
  unite(window, c(V4, V5))

# Simplify gene names
gff_genes[, "V9"] <- str_extract_all(gff_genes[, "V9"], "Name=Thupl.*", simplify = T)[,1]
gff_genes[, "V9"] <- str_extract_all(gff_genes[, "V9"], "Thupl.*", simplify = TRUE)[,1]

# Match GFF windows to pi windows
top_1_bed_window_adjusted <- top_1 %>% 
  mutate(window_pos_1_adjusted = (window_pos_1 + 1)) %>% 
  unite(window, c(window_pos_1_adjusted, window_pos_2))
  
top_1_gff <- gff_genes %>% 
  filter(window %in% top_1_bed_window_adjusted$window) %>% 
  rename(Gene = V9)

top_1_gene_annotations <- gene_annotations %>% 
  rename(Gene = V1) %>% 
  filter(Gene %in% top_1_gff$Gene)
  
top_1_annotations <- merge(top_1_gff, top_1_gene_annotations, by = "Gene", all = T) 

# write.csv(top_1_annotations, "top_1_annotations.csv")

### subpop differences - statistical tests ####

# Pi
NC_subpop_pi <- all_snps_primary_genes_sco_subpops_pi %>% 
  filter(pop == "Northern_Coastal")

C_subpop_pi <- all_snps_primary_genes_sco_subpops_pi %>% 
  filter(pop == "Central")

SI_subpop_pi <- all_snps_primary_genes_sco_subpops_pi %>% 
  filter(pop == "Southern_Interior")

mean(NC_subpop_pi$avg_pi, na.rm = T)
sd(NC_subpop_pi$avg_pi, na.rm = T)

mean(C_subpop_pi$avg_pi, na.rm = T)
mean(SI_subpop_pi$avg_pi, na.rm = T)

subpops_df <- data.frame(NC_subpop_pi$avg_pi, C_subpop_pi$avg_pi, SI_subpop_pi$avg_pi)

wilcox.test((sum(NC_subpop_pi$count_diffs, na.rm = T)/sum(NC_subpop_pi$count_comparisons, na.rm = T)), (sum(C_subpop_pi$count_diffs, na.rm = T)/sum(C_subpop_pi$count_comparisons, na.rm = T)))

wilcox.test(NC_subpop_pi$avg_pi, C_subpop_pi$avg_pi)
wilcox.test(NC_subpop_pi$avg_pi, SI_subpop_pi$avg_pi)
wilcox.test(SI_subpop_pi$avg_pi, C_subpop_pi$avg_pi)

kruskal.test(subpops_df)

# dxy
NC_C_subpop_dxy <- all_snps_primary_genes_sco_subpops_dxy %>% 
  filter(pop_comparison == "Northern_Coastal_Central")

C_SI_subpop_dxy <- all_snps_primary_genes_sco_subpops_dxy %>% 
  filter(pop_comparison == "Central_Southern_Interior")

NC_SI_subpop_dxy <- all_snps_primary_genes_sco_subpops_dxy %>% 
  filter(pop_comparison == "Northern_Coastal_Southern_Interior")


mean(NC_C_subpop_dxy$avg_dxy, na.rm = T)
sd(NC_C_subpop_dxy$avg_dxy, na.rm = T)

mean(C_SI_subpop_dxy$avg_dxy, na.rm = T)
mean(NC_SI_subpop_dxy$avg_dxy, na.rm = T)

subpops_df_dxy <- data.frame(NC_C_subpop_dxy$avg_dxy, C_SI_subpop_dxy$avg_dxy, NC_SI_subpop_dxy$avg_dxy)

kruskal.test(subpops_df_dxy)

## Top pi
sum(top_1$count_diffs, na.rm = T)/sum(top_1$count_comparisons, na.rm = T)

all_pi_bed_window_adjusted <- all_pi_no_NA %>% 
  mutate(window_pos_1_adjusted = (window_pos_1 + 1)) %>% 
  unite(window, c(window_pos_1_adjusted, window_pos_2))

pi_without_top_1 <- all_pi_bed_window_adjusted %>% 
  filter(!(window %in% top_1_bed_window_adjusted$window))

sum(pi_without_top_1$count_diffs, na.rm = T)/sum(pi_without_top_1$count_comparisons, na.rm = T)

top_5 <- all_pi_bed_window_adjusted[all_pi_bed_window_adjusted$avg_pi > quantile(all_pi_bed_window_adjusted$avg_pi,prob=1-5/100, na.rm = T),]

pi_without_top_5 <- all_pi_bed_window_adjusted %>% 
  filter(!(window %in% top_5$window))

sum(pi_without_top_5$count_diffs, na.rm = T)/sum(pi_without_top_5$count_comparisons, na.rm = T)
```
