## Supplementary material for "The western redcedar genome reveals low genetic diversity in a self-compatible conifer": Code S5: S_line_genotype_correction.R

```
rm(list=ls())

library(sommer)
library(reshape2)
library(RColorBrewer)
library(pheatmap)
library(tidyverse)
source("publication_theme.r")

## Zygosity correction functions #######################################################################################
zygosity_correction_fun_F_S5 <- function(x){
  for(i in 1:nrow(x)){
    if(x[i, 1] == 0 && x[i, 2] == 0 && is.na(x[i, 1]) == FALSE && is.na(x[i, 2]) == FALSE){
      x[i, 3:6] <- 0
    }
    if(x[i, 2] == 0 && x[i, 3] == 0 && is.na(x[i, 2]) == FALSE && is.na(x[i, 3]) == FALSE){
      x[i, 4:6] <- 0
    }
    if(x[i, 3] == 0 && x[i, 4] == 0 && is.na(x[i, 3]) == FALSE && is.na(x[i, 4]) == FALSE){
      x[i, 5:6] <- 0
    }
    if(x[i, 4] == 0 && x[i, 5] == 0 && is.na(x[i, 4]) == FALSE && is.na(x[i, 5]) == FALSE){
      x[i, 6] <- 0
    }
    if(x[i, 2] == 2 && x[i, 2] == 2 && is.na(x[i, 2]) == FALSE && is.na(x[i, 2]) == FALSE){
      x[i, 3:6] <- 2
    }
    if(x[i, 2] == 2 && x[i, 3] == 2 && is.na(x[i, 2]) == FALSE && is.na(x[i, 3]) == FALSE){
      x[i, 4:6] <- 2
    }
    if(x[i, 3] == 2 && x[i, 4] == 2 && is.na(x[i, 3]) == FALSE && is.na(x[i, 4]) == FALSE){
      x[i, 5:6] <- 2
    }
    if(x[i, 4] == 2 && x[i, 5] == 2 && is.na(x[i, 4]) == FALSE && is.na(x[i, 5]) == FALSE){
      x[i, 6] <- 2
    }
    if(x[i, 2] == 1 && x[i, 3] == 1 && is.na(x[i, 2]) == FALSE && is.na(x[i, 3]) == FALSE){
      x[i, 1] <- 1
    }
    if(x[i, 3] == 1 && x[i, 4] == 1 && is.na(x[i, 3]) == FALSE && is.na(x[i, 4]) == FALSE){
      x[i, 1:2] <- 1
    }
    if(x[i, 4] == 1 && x[i, 5] == 1 && is.na(x[i, 4]) == FALSE && is.na(x[i, 5]) == FALSE){
      x[i, 1:3] <- 1
    }
    if(x[i, 5] == 1 && x[i, 6] == 1 && is.na(x[i, 5]) == FALSE && is.na(x[i, 6]) == FALSE){
      x[i, 1:4] <- 1
    }
    if(x[i, 6] == 1 && x[i, 1:4] == 1 && is.na(x[i, 6]) == FALSE && is.na(x[i, 1:4]) == FALSE){
      x[i, 5] <- 1
    }
    if(x[i, 5:6] == 1 && x[i, 1:3] == 1 && is.na(x[i, 5:6]) == FALSE && is.na(x[i, 1:3]) == FALSE){
      x[i, 4] <- 1
    }
    if(x[i, 4:6] == 1 && x[i, 1:2] == 1 && is.na(x[i, 4:6]) == FALSE && is.na(x[i, 1:2]) == FALSE){
      x[i,3] <- 1
    }
    if(x[i, 3:6] == 1 && x[i, 1] == 1 && is.na(x[i, 3:6]) == FALSE && is.na(x[i, 1]) == FALSE){
      x[i, 2] <- 1
    }
    else if(x[i, 1] == 0 && x[i, 2] == 2 && is.na(x[i, 1]) == FALSE && is.na(x[i, 2]) == FALSE){
      x[i,1:6] <- "remove"
    }
    else if(x[i, 2] == 0 && x[i, 3] == 2 && is.na(x[i, 2]) == FALSE && is.na(x[i, 3]) == FALSE){
      x[i,1:6] <- "remove"
    }
    else if(x[i, 3] == 0 && x[i, 4] == 2 && is.na(x[i, 3]) == FALSE && is.na(x[i, 4]) == FALSE){
      x[i,1:6] <- "remove"
    }
    else if(x[i, 4] == 0 && x[i, 5] == 2 && is.na(x[i, 4]) == FALSE && is.na(x[i, 5]) == FALSE){
      x[i, 1:6] <- "remove"
    }
    else if(x[i, 5] == 0 && x[i, 6] == 2 && is.na(x[i, 5]) == FALSE && is.na(x[i, 6]) == FALSE){
      x[i,1:6] <- "remove" 
    }
    else if(x[i, 1] == 2 && x[i, 2] == 0 && is.na(x[i, 1]) == FALSE && is.na(x[i, 2]) == FALSE){
      x[i,1:6] <- "remove"
    }
    else if(x[i, 2] == 2 && x[i, 3] == 0 && is.na(x[i, 2]) == FALSE && is.na(x[i, 3]) == FALSE){
      x[i,1:6] <- "remove"
    }
    else if(x[i, 3] == 2 && x[i, 4] == 0 && is.na(x[i, 3]) == FALSE && is.na(x[i, 4]) == FALSE){
      x[i,1:6] <- "remove"
    }
    else if(x[i, 4] == 2 && x[i, 5] == 0 && is.na(x[i, 4]) == FALSE && is.na(x[i, 5]) == FALSE){
      x[i, 1:6] <- "remove"
    }
    else if(x[i, 5] == 2 && x[i, 6] == 0 && is.na(x[i, 5]) == FALSE && is.na(x[i, 6]) == FALSE){
      x[i,1:6] <- "remove" 
    }
  }
  return(x)
}

zygosity_correction_fun_P_S5 <- function(x){
  for(i in 1:nrow(x)){
    if(x[i, 1] == 0 && x[i, 2] == 0 && is.na(x[i, 1] == FALSE) && is.na(x[i, 2]) == FALSE){
      x[i, 3:8]
    }
    if(x[i, 3] == 0 && x[i, 4] == 0 && is.na(x[i, 3]) == FALSE && is.na(x[i, 4]) == FALSE){
      x[i, 5:8] <- 0
    }
    if(x[i, 4] == 0 && x[i, 5] == 0 && is.na(x[i, 4]) == FALSE && is.na(x[i, 5]) == FALSE){
      x[i, 6:8] <- 0
    }
    if(x[i, 5] == 0 && x[i, 6] == 0 && is.na(x[i, 5]) == FALSE && is.na(x[i, 6]) == FALSE){
      x[i, 7:8] <- 0
    }
    if(x[i, 6] == 0 && x[i, 7] == 0 && is.na(x[i, 6]) == FALSE && is.na(x[i, 7]) == FALSE){
      x[i, 8] <- 0
    }
    if(x[i, 4] == 2 && x[i, 4] == 2 && is.na(x[i, 4]) == FALSE && is.na(x[i, 4]) == FALSE){
      x[i, 5:8] <- 2
    }
    if(x[i, 4] == 2 && x[i, 5] == 2 && is.na(x[i, 4]) == FALSE && is.na(x[i, 5]) == FALSE){
      x[i, 6:8] <- 2
    }
    if(x[i, 5] == 2 && x[i, 6] == 2 && is.na(x[i, 5]) == FALSE && is.na(x[i, 6]) == FALSE){
      x[i, 7:8] <- 2
    }
    if(x[i, 6] == 2 && x[i, 7] == 2 && is.na(x[i, 6]) == FALSE && is.na(x[i, 7]) == FALSE){
      x[i, 8] <- 2
    }
    if(x[i, 4] == 1 && x[i, 5] == 1 && is.na(x[i, 4]) == FALSE && is.na(x[i, 5]) == FALSE){
      x[i, 3] <- 1
    }
    if(x[i, 5] == 1 && x[i, 6] == 1 && is.na(x[i, 5]) == FALSE && is.na(x[i, 6]) == FALSE){
      x[i, 3:2] <- 1
    }
    if(x[i, 6] == 1 && x[i, 7] == 1 && is.na(x[i, 6]) == FALSE && is.na(x[i, 7]) == FALSE){
      x[i, 3:5] <- 1
    }
    if(x[i, 7] == 1 && x[i, 8] == 1 && is.na(x[i, 7]) == FALSE && is.na(x[i, 8]) == FALSE){
      x[i, 3:6] <- 1
    }
    if(x[i, 8] == 1 && x[i, 3:6] == 1 && is.na(x[i, 8]) == FALSE && is.na(x[i, 3:6]) == FALSE){
      x[i, 7] <- 1
    }
    if(x[i, 7:8] == 1 && x[i, 3:5] == 1 && is.na(x[i, 7:8]) == FALSE && is.na(x[i, 3:5]) == FALSE){
      x[i, 6] <- 1
    }
    if(x[i, 6:8] == 1 && x[i, 3:2] == 1 && is.na(x[i, 6:8]) == FALSE && is.na(x[i, 3:2]) == FALSE){
      x[i,5] <- 1
    }
    if(x[i, 5:8] == 1 && x[i, 3] == 1 && is.na(x[i, 5:8]) == FALSE && is.na(x[i, 3]) == FALSE){
      x[i, 4] <- 1
    }
    else if(x[i, 3] == 0 && x[i, 4] == 2 && is.na(x[i, 3]) == FALSE && is.na(x[i, 4]) == FALSE){
      x[i,1:8] <- "remove"
    }
    else if(x[i, 4] == 0 && x[i, 5] == 2 && is.na(x[i, 4]) == FALSE && is.na(x[i, 5]) == FALSE){
      x[i,1:8] <- "remove"
    }
    else if(x[i, 5] == 0 && x[i, 6] == 2 && is.na(x[i, 5]) == FALSE && is.na(x[i, 6]) == FALSE){
      x[i,1:8] <- "remove"
    }
    else if(x[i, 6] == 0 && x[i, 7] == 2 && is.na(x[i, 6]) == FALSE && is.na(x[i, 7]) == FALSE){
      x[i, 3:8] <- "remove"
    }
    else if(x[i, 7] == 0 && x[i, 8] == 2 && is.na(x[i, 7]) == FALSE && is.na(x[i, 8]) == FALSE){
      x[i,1:8] <- "remove" 
    }
    else if(x[i, 3] == 2 && x[i, 4] == 0 && is.na(x[i, 3]) == FALSE && is.na(x[i, 4]) == FALSE){
      x[i,1:8] <- "remove"
    }
    else if(x[i, 4] == 2 && x[i, 5] == 0 && is.na(x[i, 4]) == FALSE && is.na(x[i, 5]) == FALSE){
      x[i,1:8] <- "remove"
    }
    else if(x[i, 5] == 2 && x[i, 6] == 0 && is.na(x[i, 5]) == FALSE && is.na(x[i, 6]) == FALSE){
      x[i,1:8] <- "remove"
    }
    else if(x[i, 6] == 2 && x[i, 7] == 0 && is.na(x[i, 6]) == FALSE && is.na(x[i, 7]) == FALSE){
      x[i, 3:8] <- "remove"
    }
    else if(x[i, 7] == 2 && x[i, 8] == 0 && is.na(x[i, 7]) == FALSE && is.na(x[i, 8]) == FALSE){
      x[i,1:8] <- "remove" 
    }
  }
  return(x)
}

zygosity_correction_fun_F_S4 <- function(x){
  for(i in 1:nrow(x)){
    if(x[i, 1] == 0 && x[i, 2] == 0 && is.na(x[i, 1]) == FALSE && is.na(x[i, 2]) == FALSE){
      x[i, 3:5] <- 0
    }
    if(x[i, 2] == 0 && x[i, 3] == 0 && is.na(x[i, 2]) == FALSE && is.na(x[i, 3]) == FALSE){
      x[i, 4:5] <- 0
    }
    if(x[i, 3] == 0 && x[i, 4] == 0 && is.na(x[i, 3]) == FALSE && is.na(x[i, 4]) == FALSE){
      x[i, 5] <- 0
    }
    if(x[i, 2] == 2 && x[i, 2] == 2 && is.na(x[i, 2]) == FALSE && is.na(x[i, 2]) == FALSE){
      x[i, 3:5] <- 2
    }
    if(x[i, 2] == 2 && x[i, 3] == 2 && is.na(x[i, 2]) == FALSE && is.na(x[i, 3]) == FALSE){
      x[i, 4:5] <- 2
    }
    if(x[i, 3] == 2 && x[i, 4] == 2 && is.na(x[i, 3]) == FALSE && is.na(x[i, 4]) == FALSE){
      x[i, 5] <- 2
    }
    if(x[i, 2] == 1 && x[i, 3] == 1 && is.na(x[i, 2]) == FALSE && is.na(x[i, 3]) == FALSE){
      x[i, 1] <- 1
    }
    if(x[i, 3] == 1 && x[i, 4] == 1 && is.na(x[i, 3]) == FALSE && is.na(x[i, 4]) == FALSE){
      x[i, 1:2] <- 1
    }
    if(x[i, 4] == 1 && x[i, 5] == 1 && is.na(x[i, 4]) == FALSE && is.na(x[i, 5]) == FALSE){
      x[i, 1:3] <- 1
    }
    if(x[i, 5] == 1 && x[i, 1:3] == 1 && is.na(x[i, 5]) == FALSE && is.na(x[i, 1:3]) == FALSE){
      x[i, 4] <- 1
    }
    if(x[i, 4:5] == 1 && x[i, 1:2] == 1 && is.na(x[i, 4:5]) == FALSE && is.na(x[i, 1:2]) == FALSE){
      x[i,3] <- 1
    }
    if(x[i, 3:5] == 1 && x[i, 1] == 1 && is.na(x[i, 3:5]) == FALSE && is.na(x[i, 1]) == FALSE){
      x[i, 2] <- 1
    }
    else if(x[i, 1] == 0 && x[i, 2] == 2 && is.na(x[i, 1]) == FALSE && is.na(x[i, 2]) == FALSE){
      x[i,1:5] <- "remove"
    }
    else if(x[i, 2] == 0 && x[i, 3] == 2 && is.na(x[i, 2]) == FALSE && is.na(x[i, 3]) == FALSE){
      x[i,1:5] <- "remove"
    }
    else if(x[i, 3] == 0 && x[i, 4] == 2 && is.na(x[i, 3]) == FALSE && is.na(x[i, 4]) == FALSE){
      x[i,1:5] <- "remove"
    }
    else if(x[i, 4] == 0 && x[i, 5] == 2 && is.na(x[i, 4]) == FALSE && is.na(x[i, 5]) == FALSE){
      x[i, 1:5] <- "remove"
    }
    else if(x[i, 1] == 2 && x[i, 2] == 0 && is.na(x[i, 1]) == FALSE && is.na(x[i, 2]) == FALSE){
      x[i,1:5] <- "remove"
    }
    else if(x[i, 2] == 2 && x[i, 3] == 0 && is.na(x[i, 2]) == FALSE && is.na(x[i, 3]) == FALSE){
      x[i,1:5] <- "remove"
    }
    else if(x[i, 3] == 2 && x[i, 4] == 0 && is.na(x[i, 3]) == FALSE && is.na(x[i, 4]) == FALSE){
      x[i,1:5] <- "remove"
    }
    else if(x[i, 4] == 2 && x[i, 5] == 0 && is.na(x[i, 4]) == FALSE && is.na(x[i, 5]) == FALSE){
      x[i, 1:5] <- "remove" 
    }
  }
  return(x)
}
#Line 1##################################################################################################################

# Read in Line 1 012 file
alleles <- read.csv("Line_1_samples.012",
                    header = F, sep = "\t", na.strings = -1)
str(alleles)
alleles <- alleles[,-1]

# Read in Line 1 SNP ids
snps <- read.csv("Line_1_samples.012.pos",
                 header = F, sep  = "\t")
snps$V1 <- as.character(snps$V1)
snps$V2 <- as.character(snps$V2)

# Merge columns and transpose
snps <- snps %>% 
  unite(col = snp, V1:V2, sep = '_')
t_snps <- t(snps)

# Read Line 1 sample ids
indivs <- read.csv("Line_1_samples.012.indv", header = F, sep = "\t")

# Put row and column names into 012 file
row.names(alleles) <- indivs$V1
colnames(alleles) <- t_snps

# Convert to matrix
g.mat_line_1 <- as.matrix(alleles)

t_g.mat_line_1 <- t(g.mat_line_1)
col.order <- c("1_1-F", "111-S1", "111-2-S2", "111-22-S3", 
               "111-221-S4", "111-221-S5", "1_2-F", "122-S1",
               "145-S1", "145-5-S2",
               "145-54-S3", "145-545-S4", "145-545-S5")
t_g.mat_line_1 <- t_g.mat_line_1[,col.order]

my_col <- brewer.pal(5, "Spectral")
pheatmap(t_g.mat_line_1[201:400,], cluster_rows = F, cluster_cols = F, color = my_col)

df_gmat_line_1 <- data.frame(t_g.mat_line_1, check.names = F)

Line_1_1 <- df_gmat_line_1 %>% 
  rownames_to_column() %>%
  select(1:7) %>%
  #filter((`1_1-F` == 1 & `111-S1` == 1 & `111-2-S2` == 1 & `111-22-S3` == 1 & `111-221-S4` == 1 & `111-221-S5` == 1)) %>%
  column_to_rownames()

# Line_1_4 <- df_gmat_line_1 %>% 
#   rownames_to_column() %>%
#   select(1, 11:16) %>%
#   #filter((`1_1-F` == 1 & `111-S1` == 1 & `111-2-S2` == 1 & `111-22-S3` == 1 & `111-221-S4` == 1 & `111-221-S5` == 1)) %>%
#   column_to_rownames()

new_1_1 <- zygosity_correction_fun_F_S5(Line_1_1)

new_1_1_cor <- new_1_1 %>%
  rownames_to_column() %>% 
  filter(!(`1_1-F` == "remove")) %>% 
  column_to_rownames()

new_1_1_cor[1:6] <- lapply(new_1_1_cor[1:6], as.numeric)

test_1_1 <- new_1_1_cor %>% 
  rownames_to_column() %>% 
  filter((.[[2]] == 0 | .[[2]] == 2) & .[[3]] == 1 |
           (.[[2]] == 0 | .[[2]] == 2) & .[[4]] == 1 |
           (.[[2]] == 0 | .[[2]] == 2) & .[[5]] == 1 |
           (.[[2]] == 0 | .[[2]] == 2) & .[[6]] == 1 |
           (.[[2]] == 0 | .[[2]] == 2) & .[[7]] == 1 |
           (.[[3]] == 0 | .[[3]] == 2) & .[[4]] == 1 |
           (.[[3]] == 0 | .[[3]] == 2) & .[[5]] == 1 |
           (.[[3]] == 0 | .[[3]] == 2) & .[[6]] == 1 |
           (.[[3]] == 0 | .[[3]] == 2) & .[[7]] == 1 |
           (.[[4]] == 0 | .[[4]] == 2) & .[[5]] == 1 |
           (.[[4]] == 0 | .[[4]] == 2) & .[[6]] == 1 |
           (.[[4]] == 0 | .[[4]] == 2) & .[[7]] == 1 |
           (.[[5]] == 0 | .[[5]] == 2) & .[[6]] == 1 |
           (.[[5]] == 0 | .[[5]] == 2) & .[[7]] == 1 |
           (.[[6]] == 0 | .[[6]] == 2) & .[[7]] == 1) %>% 
  column_to_rownames()

Line_1_1_corrected <- new_1_1_cor %>%
  rownames_to_column() %>% 
  filter(!(row.names(new_1_1_cor) %in% row.names(test_1_1))) %>% 
  column_to_rownames()


# new_1_4 <- zygosity_correction_fun_F_S5(Line_1_4)
# new_1_4_cor <- new_1_4 %>%
#   rownames_to_column() %>% 
#   filter(!(`1_4-F` == "remove")) %>% 
#   column_to_rownames()
# 
# new_1_4_cor[1:6] <- lapply(new_1_4_cor[1:6], as.numeric)
#   
# test_1_4 <- new_1_4_cor %>% 
#   rownames_to_column() %>%
#   filter((.[[2]] == 0 | .[[2]] == 2) & .[[3]] == 1 |
#            (.[[2]] == 0 | .[[2]] == 2) & .[[4]] == 1 |
#            (.[[2]] == 0 | .[[2]] == 2) & .[[5]] == 1 |
#            (.[[2]] == 0 | .[[2]] == 2) & .[[6]] == 1 |
#            (.[[2]] == 0 | .[[2]] == 2) & .[[7]] == 1 |
#            (.[[3]] == 0 | .[[3]] == 2) & .[[4]] == 1 |
#            (.[[3]] == 0 | .[[3]] == 2) & .[[5]] == 1 |
#            (.[[3]] == 0 | .[[3]] == 2) & .[[6]] == 1 |
#            (.[[3]] == 0 | .[[3]] == 2) & .[[7]] == 1 |
#            (.[[4]] == 0 | .[[4]] == 2) & .[[5]] == 1 |
#            (.[[4]] == 0 | .[[4]] == 2) & .[[6]] == 1 |
#            (.[[4]] == 0 | .[[4]] == 2) & .[[7]] == 1 |
#            (.[[5]] == 0 | .[[5]] == 2) & .[[6]] == 1 |
#            (.[[5]] == 0 | .[[5]] == 2) & .[[7]] == 1 |
#            (.[[6]] == 0 | .[[6]] == 2) & .[[7]] == 1) %>%  
#   column_to_rownames()
# 
# Line_1_4_corrected <- new_1_4_cor %>% 
#   rownames_to_column() %>% 
#   filter(!(row.names(new_1_4_cor) %in% row.names(test_1_4))) %>% 
#   column_to_rownames()


# how many heterozygotes
table(Line_1_1$`1_1-F`)
table(Line_1_4$`1_4-F`)

table(Line_1_1$`111-S1`)
table(Line_1_4$`145-S1`)

table(Line_1_1$`111-2-S2`)
table(Line_1_4$`145-5-S2`)

table(Line_1_1$`111-22-S3`)
table(Line_1_4$`145-54-S3`)

table(Line_1_1$`111-221-S4`)
table(Line_1_4$`145-545-S4`)

table(Line_1_1$`111-221-S5`)
table(Line_1_4$`145-545-S5`)


# pdf("Line_1_4_homozygotes_becoming_heterozygotes_heatmap.pdf", onefile = T, width = 15, height = 10)
pheatmap(Line_1_4_corrected[1200:1300,], cluster_rows = F, cluster_cols = F, color = my_col, show_rownames = F)
dev.off()

#write.csv(rownames(Line_1_1), "snp_zygosity_analysis/Line_11_1_heterozygote_persistence_no_parents_snps.txt", row.names = F, col.names = F)
##Line 6#################################################################################################################

# Read in Line 6 012 file
alleles <- read.csv("Line_6_samples.012",
                    header = F, sep = "\t", na.strings = -1)
str(alleles)
alleles <- alleles[,-1]

# Read in Line 6 SNP ids
snps <- read.csv("Line_6_samples.012.pos",
                 header = F, sep  = "\t")
snps$V1 <- as.character(snps$V1)
snps$V2 <- as.character(snps$V2)

# Merge columns and transpose
snps <- snps %>% 
  unite(col = snp, V1:V2, sep = '_')
t_snps <- t(snps)

# Read Line 6 sample ids
indivs <- read.csv("Line_6_samples.012.indv", header = F, sep = "\t")

# Put row and column names into 012 file
row.names(alleles) <- indivs$V1
colnames(alleles) <- t_snps

# Convert to matrix
g.mat_Line_6 <- as.matrix(alleles)

t_g.mat_Line_6 <- t(g.mat_Line_6)
col.order <- c("6_1-F", "611-S1","611-1-S2", "611-11-S3", "611-111-S4",
               "6_2-F", "623-S1", "623-211-S4",
               "6_4-F", "646-S1", "646-4-S2", "646-45-S3", "646-454-S4", "646-454-S5")
t_g.mat_Line_6 <- t_g.mat_Line_6[,col.order]

my_col <- brewer.pal(5, "Spectral")
pheatmap(t_g.mat_Line_6[1:25,], cluster_rows = F, cluster_cols = F, color = my_col)

df_gmat_line_6 <- data.frame(t_g.mat_Line_6, check.names = F)

# Filter Line 6_1
Line_6_1 <- df_gmat_line_6 %>% 
  rownames_to_column() %>%
  select(1:6) %>%
  #filter((`49-P` == 1 & `150-P` == 1 & `6_1-F` == 1 & `611-S1` == 1 & `611-1-S2` == 1 & `611-11-S3` == 1 & `611-111-S4` == 1)) %>%
  column_to_rownames()


# Correcting Line 6_1
new_6_1 <- zygosity_correction_fun_F_S4(Line_6_1)

new_6_1_cor <- new_6_1 %>%
  rownames_to_column() %>% 
  filter(!(`6_1-F` == "remove")) %>% 
  column_to_rownames()

new_6_1_cor[1:5] <- lapply(new_6_1_cor[1:5], as.numeric)

test_6_1 <- new_6_1_cor %>% 
  rownames_to_column() %>% 
  filter((.[[2]] == 0 | .[[2]] == 2) & .[[3]] == 1 |
           (.[[2]] == 0 | .[[2]] == 2) & .[[4]] == 1 |
           (.[[2]] == 0 | .[[2]] == 2) & .[[5]] == 1 |
           (.[[2]] == 0 | .[[2]] == 2) & .[[6]] == 1 |
           (.[[3]] == 0 | .[[3]] == 2) & .[[4]] == 1 |
           (.[[3]] == 0 | .[[3]] == 2) & .[[5]] == 1 |
           (.[[3]] == 0 | .[[3]] == 2) & .[[6]] == 1 |
           (.[[4]] == 0 | .[[4]] == 2) & .[[5]] == 1 |
           (.[[4]] == 0 | .[[4]] == 2) & .[[6]] == 1 |
           (.[[5]] == 0 | .[[5]] == 2) & .[[6]] == 1) %>%  
  column_to_rownames()

Line_6_1_corrected <- new_6_1_cor %>%
  rownames_to_column() %>% 
  filter(!(row.names(new_6_1_cor) %in% row.names(test_6_1))) %>% 
  column_to_rownames()

# Filter Line 6_2
Line_6_2 <- df_gmat_line_6 %>% 
  rownames_to_column() %>%
  select(1, 7:9) %>%
  #filter((`49-P` == 1 & `150-P` == 1 & `6_1-F` == 1 & `611-S1` == 1 & `611-1-S2` == 1 & `611-11-S3` == 1 & `611-111-S4` == 1)) %>%
  column_to_rownames()

# Filter Line 6_4
Line_6_4 <- df_gmat_line_6 %>%
  rownames_to_column() %>%
  select(1, 10:15) %>%
  #filter((`49-P` == 1 & `150-P` == 1 & `6_4-F` == 1 & `646-S1` == 1 & `646-4-S2` == 1 & `646-45-S3` == 1 & `646-454-S4` == 1 & `646-454-S5` == 1)) %>%
  column_to_rownames()

# Correcting Line 6_4
new_6_4 <- zygosity_correction_fun_F_S5(Line_6_4)
new_6_4_cor <- new_6_4 %>%
  rownames_to_column() %>% 
  filter(!(`6_4-F` == "remove")) %>% 
  column_to_rownames()

new_6_4_cor[1:6] <- lapply(new_6_4_cor[1:6], as.numeric)

test_6_4 <- new_6_4_cor %>% 
  rownames_to_column() %>%
  filter((.[[2]] == 0 | .[[2]] == 2) & .[[3]] == 1 |
           (.[[2]] == 0 | .[[2]] == 2) & .[[4]] == 1 |
           (.[[2]] == 0 | .[[2]] == 2) & .[[5]] == 1 |
           (.[[2]] == 0 | .[[2]] == 2) & .[[6]] == 1 |
           (.[[2]] == 0 | .[[2]] == 2) & .[[7]] == 1 |
           (.[[3]] == 0 | .[[3]] == 2) & .[[4]] == 1 |
           (.[[3]] == 0 | .[[3]] == 2) & .[[5]] == 1 |
           (.[[3]] == 0 | .[[3]] == 2) & .[[6]] == 1 |
           (.[[3]] == 0 | .[[3]] == 2) & .[[7]] == 1 |
           (.[[4]] == 0 | .[[4]] == 2) & .[[5]] == 1 |
           (.[[4]] == 0 | .[[4]] == 2) & .[[6]] == 1 |
           (.[[4]] == 0 | .[[4]] == 2) & .[[7]] == 1 |
           (.[[5]] == 0 | .[[5]] == 2) & .[[6]] == 1 |
           (.[[5]] == 0 | .[[5]] == 2) & .[[7]] == 1 |
           (.[[6]] == 0 | .[[6]] == 2) & .[[7]] == 1) %>%  
  column_to_rownames()

Line_6_4_corrected <- new_6_4_cor %>% 
  rownames_to_column() %>% 
  filter(!(row.names(new_6_4_cor) %in% row.names(test_6_4))) %>% 
  column_to_rownames()


# Line_6_1_het_parents_minor_allele_homozygote <- df_gmat_line_6 %>% 
#   rownames_to_column() %>%
#   select(1:8) %>%
#   filter((`49-P` == 1 & `150-P` == 1 & `6_1-F` == 0 & `611-S1` == 0 & `611-1-S2` == 0 & `611-11-S3` == 0 & `611-111-S4` == 0) |
#           (`49-P` == 1 & `150-P` == 1 & `6_1-F` == 1 & `611-S1` == 0 & `611-1-S2` == 0 & `611-11-S3` == 0 & `611-111-S4` == 0) |
#           (`49-P` == 1 & `150-P` == 1 & `6_1-F` == 1 & `611-S1` == 1 & `611-1-S2` == 0 & `611-11-S3` == 0 & `611-111-S4` == 0) |
#           (`49-P` == 1 & `150-P` == 1 & `6_1-F` == 1 & `611-S1` == 1 & `611-1-S2` == 1 & `611-11-S3` == 0 & `611-111-S4` == 0) |
#            (`49-P` == 1 & `150-P` == 1 & `6_1-F` == 1 & `611-S1` == 1 & `611-1-S2` == 1 & `611-11-S3` == 1 & `611-111-S4` == 0)) %>%
#   column_to_rownames()
# 
# Line_6_1_het_parents_major_allele_homozygote <- df_gmat_line_6 %>% 
#   rownames_to_column() %>%
#   select(1:8) %>%
#   filter((`49-P` == 1 & `150-P` == 1 & `6_1-F` == 2 & `611-S1` == 2 & `611-1-S2` == 2 & `611-11-S3` == 2 & `611-111-S4` == 2) |
#            (`49-P` == 1 & `150-P` == 1 & `6_1-F` == 1 & `611-S1` == 2 & `611-1-S2` == 2 & `611-11-S3` == 2 & `611-111-S4` == 2) |
#            (`49-P` == 1 & `150-P` == 1 & `6_1-F` == 1 & `611-S1` == 1 & `611-1-S2` == 2 & `611-11-S3` == 2 & `611-111-S4` == 2) |
#            (`49-P` == 1 & `150-P` == 1 & `6_1-F` == 1 & `611-S1` == 1 & `611-1-S2` == 1 & `611-11-S3` == 2 & `611-111-S4` == 2) |
#            (`49-P` == 1 & `150-P` == 1 & `6_1-F` == 1 & `611-S1` == 1 & `611-1-S2` == 1 & `611-11-S3` == 1 & `611-111-S4` == 2)) %>%
#   column_to_rownames()
# 
# Line_6_4 <- df_gmat_line_6 %>% 
#   rownames_to_column() %>%
#   select(1:3, 12:17) %>%
#   #filter((`49-P` == 1 & `150-P` == 1 & `6_4-F` == 1 & `646-S1` == 1 & `646-4-S2` == 1 & `646-45-S3` == 1 & `646-454-S4` == 1 & `646-454-S5` == 1)) %>%
#   column_to_rownames()
# 
# Line_6_4_het_parents_minor_allele_homozygote <- df_gmat_line_6 %>% 
#   rownames_to_column() %>%
#   select(1:3, 12:17) %>%
#   filter((`49-P` == 1 & `150-P` == 1 & `6_4-F` == 0 & `646-S1` == 0 & `646-4-S2` == 0 & `646-45-S3` == 0 & `646-454-S4` == 0 & `646-454-S5` == 0) |
#            (`49-P` == 1 & `150-P` == 1 & `6_4-F` == 1 & `646-S1` == 0 & `646-4-S2` == 0 & `646-45-S3` == 0 & `646-454-S4` == 0 & `646-454-S5` == 0) |
#            (`49-P` == 1 & `150-P` == 1 & `6_4-F` == 1 & `646-S1` == 1 & `646-4-S2` == 0 & `646-45-S3` == 0 & `646-454-S4` == 0 & `646-454-S5` == 0) |
#            (`49-P` == 1 & `150-P` == 1 & `6_4-F` == 1 & `646-S1` == 1 & `646-4-S2` == 1 & `646-45-S3` == 0 & `646-454-S4` == 0 & `646-454-S5` == 0) |
#            (`49-P` == 1 & `150-P` == 1 & `6_4-F` == 1 & `646-S1` == 1 & `646-4-S2` == 1 & `646-45-S3` == 1 & `646-454-S4` == 0 & `646-454-S5` == 0) |
#            (`49-P` == 1 & `150-P` == 1 & `6_4-F` == 1 & `646-S1` == 1 & `646-4-S2` == 1 & `646-45-S3` == 1 & `646-454-S4` == 1 & `646-454-S5` == 0)) %>%
#   column_to_rownames()
# 
# Line_6_4_het_parents_major_allele_homozygote <- df_gmat_line_6 %>% 
#   rownames_to_column() %>%
#   select(1:3, 12:17) %>%
#   filter((`49-P` == 1 & `150-P` == 1 & `6_4-F` == 2 & `646-S1` == 2 & `646-4-S2` == 2 & `646-45-S3` == 2 & `646-454-S4` == 2 & `646-454-S5` == 2) |
#            (`49-P` == 1 & `150-P` == 1 & `6_4-F` == 1 & `646-S1` == 2 & `646-4-S2` == 2 & `646-45-S3` == 2 & `646-454-S4` == 2 & `646-454-S5` == 2) |
#            (`49-P` == 1 & `150-P` == 1 & `6_4-F` == 1 & `646-S1` == 1 & `646-4-S2` == 2 & `646-45-S3` == 2 & `646-454-S4` == 2 & `646-454-S5` == 2) |
#            (`49-P` == 1 & `150-P` == 1 & `6_4-F` == 1 & `646-S1` == 1 & `646-4-S2` == 1 & `646-45-S3` == 2 & `646-454-S4` == 2 & `646-454-S5` == 2) |
#            (`49-P` == 1 & `150-P` == 1 & `6_4-F` == 1 & `646-S1` == 1 & `646-4-S2` == 1 & `646-45-S3` == 1 & `646-454-S4` == 2 & `646-454-S5` == 2) |
#            (`49-P` == 1 & `150-P` == 1 & `6_4-F` == 1 & `646-S1` == 1 & `646-4-S2` == 1 & `646-45-S3` == 1 & `646-454-S4` == 1 & `646-454-S5` == 2)) %>%
#   column_to_rownames()


# how many heterozygotes
table(Line_6_1_corrected$`6_1-F`)
table(Line_6_4_corrected$`6_4-F`)

table(Line_6_1_corrected$`611-S1`)
table(Line_6_4_corrected$`646-S1`)

table(Line_6_1_corrected$`611-1-S2`)
table(Line_6_4_corrected$`646-4-S2`)

table(Line_6_1_corrected$`611-11-S3`)
table(Line_6_4_corrected$`646-45-S3`)

table(Line_6_1_corrected$`611-111-S4`)
table(Line_6_4_corrected$`646-454-S4`)

table(Line_6_4_corrected$`646-454-S5`)

pheatmap(Line_6_1_corrected[1:100,], cluster_rows = F, cluster_cols = F, color = my_col)

# write.csv(Line_6_1, "snp_zygosity_analysis/Line_6_1_heterozygote_persistence.txt")
# write.csv(Line_6_4, "snp_zygosity_analysis/Line_6_4_heterozygote_persistence.txt")
# 
# write.csv(rownames(Line_6_1), "snp_zygosity_analysis/Line_6_1_heterozygote_persistence_parents_any_zygosity_snps.txt", row.names = F, col.names = F)
# write.csv(rownames(Line_6_4), "snp_zygosity_analysis/Line_6_4_heterozygote_persistence_parents_any_zygosity_snps.txt", row.names = F, col.names = F)

#Line 7##################################################################################################################

# Read in Line 7 012 file
alleles <- read.csv("Line_7_samples.012",
                    header = F, sep = "\t", na.strings = -1)
str(alleles)
alleles <- alleles[,-1]

# Read in Line 7 SNP ids
snps <- read.csv("Line_7_samples.012.pos",
                 header = F, sep  = "\t")
snps$V1 <- as.character(snps$V1)
snps$V2 <- as.character(snps$V2)

# Merge columns and transpose
snps <- snps %>% 
  unite(col = snp, V1:V2, sep = '_')
t_snps <- t(snps)

# Read Line 7 sample ids
indivs <- read.csv("Line_7_samples.012.indv", header = F, sep = "\t")

# Put row and column names into 012 file
row.names(alleles) <- indivs$V1
colnames(alleles) <- t_snps

# Convert to matrix
g.mat_Line_7 <- as.matrix(alleles)

t_g.mat_Line_7 <- t(g.mat_Line_7)
t_g.mat_Line_7 <- cbind(t_g.mat_Line_7, `721-21-S3` = rep(NA, 51638))
col.order <- c("7_1-F", "712-S1","712-111-S4",
               "7_2-F", "721-S1", "721-2-S2", "721-21-S3","721-211-S4",
               "7_4-F", "745-S1", "745-4-S2", "745-44-S3", "745-444-S4")
t_g.mat_Line_7 <- t_g.mat_Line_7[,col.order]

df_gmat_line_7 <- data.frame(t_g.mat_Line_7, check.names = F)

# Filter Line 7_2
Line_7_2 <- df_gmat_line_7 %>% 
  rownames_to_column() %>%
  select(1, 5:9) %>%
  column_to_rownames()

# Correcting Line 7_2
new_7_2 <- zygosity_correction_fun_F_S4(Line_7_2)

new_7_2_cor <- new_7_2 %>%
  rownames_to_column() %>% 
  filter(!(`7_2-F` == "remove")) %>% 
  column_to_rownames()

new_7_2_cor[1:5] <- lapply(new_7_2_cor[1:5], as.numeric)

test_7_2 <- new_7_2_cor %>% 
  rownames_to_column() %>% 
  filter((.[[2]] == 0 | .[[2]] == 2) & .[[3]] == 1 |
           (.[[2]] == 0 | .[[2]] == 2) & .[[4]] == 1 |
           (.[[2]] == 0 | .[[2]] == 2) & .[[5]] == 1 |
           (.[[2]] == 0 | .[[2]] == 2) & .[[6]] == 1 |
           (.[[3]] == 0 | .[[3]] == 2) & .[[4]] == 1 |
           (.[[3]] == 0 | .[[3]] == 2) & .[[5]] == 1 |
           (.[[3]] == 0 | .[[3]] == 2) & .[[6]] == 1 |
           (.[[4]] == 0 | .[[4]] == 2) & .[[5]] == 1 |
           (.[[4]] == 0 | .[[4]] == 2) & .[[6]] == 1 |
           (.[[5]] == 0 | .[[5]] == 2) & .[[6]] == 1) %>%  
  column_to_rownames()

Line_7_2_corrected <- new_7_2_cor %>%
  rownames_to_column() %>% 
  filter(!(row.names(new_7_2_cor) %in% row.names(test_7_2))) %>% 
  column_to_rownames()


# Filter Line 7_4
Line_7_4 <- df_gmat_line_7 %>% 
  rownames_to_column() %>%
  select(1, 10:14) %>%
  column_to_rownames()

# Correcting Line 7_4
new_7_4 <- zygosity_correction_fun_F_S4(Line_7_4)

new_7_4_cor <- new_7_4 %>%
  rownames_to_column() %>% 
  filter(!(`7_4-F` == "remove")) %>% 
  column_to_rownames()

new_7_4_cor[1:5] <- lapply(new_7_4_cor[1:5], as.numeric)

test_7_4 <- new_7_4_cor %>% 
  rownames_to_column() %>% 
  filter((.[[2]] == 0 | .[[2]] == 2) & .[[3]] == 1 |
           (.[[2]] == 0 | .[[2]] == 2) & .[[4]] == 1 |
           (.[[2]] == 0 | .[[2]] == 2) & .[[5]] == 1 |
           (.[[2]] == 0 | .[[2]] == 2) & .[[6]] == 1 |
           (.[[3]] == 0 | .[[3]] == 2) & .[[4]] == 1 |
           (.[[3]] == 0 | .[[3]] == 2) & .[[5]] == 1 |
           (.[[3]] == 0 | .[[3]] == 2) & .[[6]] == 1 |
           (.[[4]] == 0 | .[[4]] == 2) & .[[5]] == 1 |
           (.[[4]] == 0 | .[[4]] == 2) & .[[6]] == 1 |
           (.[[5]] == 0 | .[[5]] == 2) & .[[6]] == 1) %>%  
  column_to_rownames()

Line_7_4_corrected <- new_7_4_cor %>%
  rownames_to_column() %>% 
  filter(!(row.names(new_7_4_cor) %in% row.names(test_7_4))) %>% 
  column_to_rownames()


# how many heterozygotes
table(Line_7_4$`7_4-F`)

table(Line_7_4$`745-S1`)

table(Line_7_4$`745-4-S2`)

table(Line_7_4$`745-44-S3`)

table(Line_7_4$`745-444-S4`)

my_col <- brewer.pal(5, "Spectral")
pheatmap(t_g.mat_Line_7[1:50,], cluster_rows = F, cluster_cols = F, color = my_col)


#Line 8##################################################################################################################

# Read in Line 8 012 file
alleles <- read.csv("Line_8_samples.012",
                    header = F, sep = "\t", na.strings = -1)
str(alleles)
alleles <- alleles[,-1]

# Read in Line 8 SNP ids
snps <- read.csv("Line_8_samples.012.pos",
                 header = F, sep  = "\t")
snps$V1 <- as.character(snps$V1)
snps$V2 <- as.character(snps$V2)

# Merge columns and transpose
snps <- snps %>% 
  unite(col = snp, V1:V2, sep = '_')
t_snps <- t(snps)

# Read Line 8 sample ids
indivs <- read.csv("Line_8_samples.012.indv", header = F, sep = "\t")

# Put row and column names into 012 file
row.names(alleles) <- indivs$V1
colnames(alleles) <- t_snps

# Convert to matrix
g.mat_Line_8 <- as.matrix(alleles)

t_g.mat_Line_8 <- t(g.mat_Line_8)
col.order <- c("8_2-F", "821-S1", "821-1-S2", "821-11-S3", "821-111-S4",
               "8_4-F", "845-S1", "845-4-S2", "845-44-S3", "845-444-S4",
               "8_5-F", "854-S1", "854-544-S4")
t_g.mat_Line_8 <- t_g.mat_Line_8[,col.order]

df_gmat_line_8 <- data.frame(t_g.mat_Line_8, check.names = F)

# Filter Line 8_2
Line_8_2 <- df_gmat_line_8 %>% 
  rownames_to_column() %>%
  select(1:6) %>%
  column_to_rownames()

# Correcting Line 8_2
new_8_2 <- zygosity_correction_fun_F_S4(Line_8_2)

new_8_2_cor <- new_8_2 %>%
  rownames_to_column() %>% 
  filter(!(`8_2-F` == "remove")) %>% 
  column_to_rownames()

new_8_2_cor[1:5] <- lapply(new_8_2_cor[1:5], as.numeric)

test_8_2 <- new_8_2_cor %>% 
  rownames_to_column() %>% 
  filter((.[[2]] == 0 | .[[2]] == 2) & .[[3]] == 1 |
           (.[[2]] == 0 | .[[2]] == 2) & .[[4]] == 1 |
           (.[[2]] == 0 | .[[2]] == 2) & .[[5]] == 1 |
           (.[[2]] == 0 | .[[2]] == 2) & .[[6]] == 1 |
           (.[[3]] == 0 | .[[3]] == 2) & .[[4]] == 1 |
           (.[[3]] == 0 | .[[3]] == 2) & .[[5]] == 1 |
           (.[[3]] == 0 | .[[3]] == 2) & .[[6]] == 1 |
           (.[[4]] == 0 | .[[4]] == 2) & .[[5]] == 1 |
           (.[[4]] == 0 | .[[4]] == 2) & .[[6]] == 1 |
           (.[[5]] == 0 | .[[5]] == 2) & .[[6]] == 1) %>%  
  column_to_rownames()

Line_8_2_corrected <- new_8_2_cor %>%
  rownames_to_column() %>% 
  filter(!(row.names(new_8_2_cor) %in% row.names(test_8_2))) %>% 
  column_to_rownames()

# Filter Line 8_4
Line_8_4 <- df_gmat_line_8 %>% 
  rownames_to_column() %>%
  select(1, 7:11) %>%
  column_to_rownames()

# Correcting Line 8_4
new_8_4 <- zygosity_correction_fun_F_S4(Line_8_4)

new_8_4_cor <- new_8_4 %>%
  rownames_to_column() %>% 
  filter(!(`8_4-F` == "remove")) %>% 
  column_to_rownames()

new_8_4_cor[1:5] <- lapply(new_8_4_cor[1:5], as.numeric)

test_8_4 <- new_8_4_cor %>% 
  rownames_to_column() %>% 
  filter((.[[2]] == 0 | .[[2]] == 2) & .[[3]] == 1 |
           (.[[2]] == 0 | .[[2]] == 2) & .[[4]] == 1 |
           (.[[2]] == 0 | .[[2]] == 2) & .[[5]] == 1 |
           (.[[2]] == 0 | .[[2]] == 2) & .[[6]] == 1 |
           (.[[3]] == 0 | .[[3]] == 2) & .[[4]] == 1 |
           (.[[3]] == 0 | .[[3]] == 2) & .[[5]] == 1 |
           (.[[3]] == 0 | .[[3]] == 2) & .[[6]] == 1 |
           (.[[4]] == 0 | .[[4]] == 2) & .[[5]] == 1 |
           (.[[4]] == 0 | .[[4]] == 2) & .[[6]] == 1 |
           (.[[5]] == 0 | .[[5]] == 2) & .[[6]] == 1) %>%  
  column_to_rownames()

Line_8_4_corrected <- new_8_4_cor %>%
  rownames_to_column() %>% 
  filter(!(row.names(new_8_4_cor) %in% row.names(test_8_4))) %>% 
  column_to_rownames()


# how many heterozygotes
table(Line_8_2$`8_2-F`)

table(Line_8_2$`821-S1`)

table(Line_8_2$`821-1-S2`)

table(Line_8_2$`821-11-S3`)

table(Line_8_2$`821-111-S4`)

table(Line_8_4$`8_4-F`)

table(Line_8_4$`845-S1`)

table(Line_8_4$`845-4-S2`)

table(Line_8_4$`845-44-S3`)

table(Line_8_4$`845-444-S4`)

my_col <- brewer.pal(5, "Spectral")
pheatmap(t_g.mat_Line_8[1:50,], cluster_rows = F, cluster_cols = F, color = my_col)


##Line 10#################################################################################################################

# Read in Line 10 012 file
alleles <- read.csv("Line_10_samples.012",
                    header = F, sep = "\t", na.strings = -1)
str(alleles)
alleles <- alleles[,-1]

# Read in Line 10 SNP ids
snps <- read.csv("Line_10_samples.012.pos",
                 header = F, sep  = "\t")
snps$V1 <- as.character(snps$V1)
snps$V2 <- as.character(snps$V2)

# Merge columns and transpose
snps <- snps %>% 
  unite(col = snp, V1:V2, sep = '_')
t_snps <- t(snps)

# Read Line 10 sample ids
indivs <- read.csv("Line_10_samples.012.indv", header = F, sep = "\t")

# Put row and column names into 012 file
row.names(alleles) <- indivs$V1
colnames(alleles) <- t_snps

# Convert to matrix
g.mat_Line_10 <- as.matrix(alleles)

t_g.mat_Line_10 <- t(g.mat_Line_10)
t_g.mat_Line_10 <- cbind(t_g.mat_Line_10, `1022-11-S3` = rep(NA, 51638), `1056-45-S3` = rep(NA, 51638))

col.order <- c("10_1-F", "1011-S1", "1011-121-S4",
               "10_2-F", "1022-S1", "1022-1-S2", "1022-11-S3", "1022-113-S4", "1022-113-S5",
               "10_5-F", "1056-S1", "1056-4-S2", "1056-45-S3", "1056-454-S4")
t_g.mat_Line_10 <- t_g.mat_Line_10[,col.order]

my_col <- brewer.pal(5, "Spectral")
pheatmap(t_g.mat_Line_10[1:50,], cluster_rows = F, cluster_cols = F, color = my_col)

df_gmat_line_10 <- data.frame(t_g.mat_Line_10, check.names = F)

# Filter Line 10_2
Line_10_2 <- df_gmat_line_10 %>% 
  rownames_to_column() %>%
  select(1, 5:10) %>%
  # filter(((`263-P` == 1 & `306-P` == 1) | (`263-P` == 1 & `306-P` == 0) | (`263-P` == 1 & `306-P` == 2) | 
  #         (`263-P` == 0 & `306-P` == 1) | (`263-P` == 0 & `306-P` == 2) | (`263-P` == 2 & `306-P` == 0) | (`263-P` == 2 & `306-P` == 1)) & 
  #          `10_2-F` == 1 & `1022-S1` == 1 & `1022-1-S2` == 1 & `1022-113-S4` == 1 & `1022-113-S5` == 1) %>%
  column_to_rownames()

# Correcting Line 10_2
new_10_2 <- zygosity_correction_fun_F_S5(Line_10_2)

new_10_2_cor <- new_10_2 %>%
  rownames_to_column() %>% 
  filter(!(`10_2-F` == "remove")) %>% 
  column_to_rownames()

new_10_2_cor[1:6] <- lapply(new_10_2_cor[1:6], as.numeric)

test_10_2 <- new_10_2_cor %>% 
  rownames_to_column() %>%
  filter((.[[2]] == 0 | .[[2]] == 2) & .[[3]] == 1 |
           (.[[2]] == 0 | .[[2]] == 2) & .[[4]] == 1 |
           (.[[2]] == 0 | .[[2]] == 2) & .[[5]] == 1 |
           (.[[2]] == 0 | .[[2]] == 2) & .[[6]] == 1 |
           (.[[2]] == 0 | .[[2]] == 2) & .[[7]] == 1 |
           (.[[3]] == 0 | .[[3]] == 2) & .[[4]] == 1 |
           (.[[3]] == 0 | .[[3]] == 2) & .[[5]] == 1 |
           (.[[3]] == 0 | .[[3]] == 2) & .[[6]] == 1 |
           (.[[3]] == 0 | .[[3]] == 2) & .[[7]] == 1 |
           (.[[4]] == 0 | .[[4]] == 2) & .[[5]] == 1 |
           (.[[4]] == 0 | .[[4]] == 2) & .[[6]] == 1 |
           (.[[4]] == 0 | .[[4]] == 2) & .[[7]] == 1 |
           (.[[5]] == 0 | .[[5]] == 2) & .[[6]] == 1 |
           (.[[5]] == 0 | .[[5]] == 2) & .[[7]] == 1 |
           (.[[6]] == 0 | .[[6]] == 2) & .[[7]] == 1) %>%  
  column_to_rownames()

Line_10_2_corrected <- new_10_2_cor %>% 
  rownames_to_column() %>% 
  filter(!(row.names(new_10_2_cor) %in% row.names(test_10_2))) %>% 
  column_to_rownames()

# Filter Line 10_5
Line_10_5 <- df_gmat_line_10 %>% 
  rownames_to_column() %>%
  select(1, 11:15) %>%
  #filter((`263-P` == 1 & `306-P` == 1 & `10_5-F` == 1 & `1056-S1` == 1 & `1056-4-S2` == 1 & `1056-45-S3` == 1 & `1056-454-S4` == 1)) %>%
  column_to_rownames()

# Correcting Line 10_5
new_10_5 <- zygosity_correction_fun_F_S4(Line_10_5)

new_10_5_cor <- new_10_5 %>%
  rownames_to_column() %>% 
  filter(!(`10_5-F` == "remove")) %>% 
  column_to_rownames()

new_10_5_cor[1:5] <- lapply(new_10_5_cor[1:5], as.numeric)

test_10_5 <- new_10_5_cor %>% 
  rownames_to_column() %>% 
  filter((.[[2]] == 0 | .[[2]] == 2) & .[[3]] == 1 |
           (.[[2]] == 0 | .[[2]] == 2) & .[[4]] == 1 |
           (.[[2]] == 0 | .[[2]] == 2) & .[[5]] == 1 |
           (.[[2]] == 0 | .[[2]] == 2) & .[[6]] == 1 |
           (.[[3]] == 0 | .[[3]] == 2) & .[[4]] == 1 |
           (.[[3]] == 0 | .[[3]] == 2) & .[[5]] == 1 |
           (.[[3]] == 0 | .[[3]] == 2) & .[[6]] == 1 |
           (.[[4]] == 0 | .[[4]] == 2) & .[[5]] == 1 |
           (.[[4]] == 0 | .[[4]] == 2) & .[[6]] == 1 |
           (.[[5]] == 0 | .[[5]] == 2) & .[[6]] == 1) %>%  
  column_to_rownames()

Line_10_5_corrected <- new_10_5_cor %>%
  rownames_to_column() %>% 
  filter(!(row.names(new_10_5_cor) %in% row.names(test_10_5))) %>% 
  column_to_rownames()

# how many heterozygotes
table(Line_10_5$`10_5-F`)

table(Line_10_5$`1056-S1`)

table(Line_10_5$`1056-4-S2`)

table(Line_10_5$`1056-45-S3`)

table(Line_10_5$`1056-454-S4`)

pheatmap(Line_10_2, cluster_rows = F, cluster_cols = F, color = my_col)

table(Line_10_5$`10_5-F`)

# write.csv(Line_10_1, "snp_zygosity_analysis/Line_10_1_heterozygote_persistence.txt")
# write.csv(Line_10_2, "snp_zygosity_analysis/Line_10_2_heterozygote_persistence.txt")
# write.csv(Line_10_5, "snp_zygosity_analysis/Line_10_5_heterozygote_persistence.txt")
# 
# write.csv(rownames(Line_10_1), "snp_zygosity_analysis/Line_10_1_heterozygote_persistence_parents_any_zygosity_snps.txt", row.names = F, col.names = F)
# write.csv(rownames(Line_10_2), "snp_zygosity_analysis/Line_10_2_heterozygote_persistence_parents_any_zygosity_snps.txt", row.names = F, col.names = F)
# write.csv(rownames(Line_10_5), "snp_zygosity_analysis/Line_10_5_heterozygote_persistence_parents_any_zygosity_snps.txt", row.names = F, col.names = F)

#Line 13##################################################################################################################

# Read in Line 13 012 file
alleles <- read.csv("Line_13_samples.012",
                    header = F, sep = "\t", na.strings = -1)
str(alleles)
alleles <- alleles[,-1]

# Read in Line 13 SNP ids
snps <- read.csv("Line_13_samples.012.pos",
                 header = F, sep  = "\t")
snps$V1 <- as.character(snps$V1)
snps$V2 <- as.character(snps$V2)

# Merge columns and transpose
snps <- snps %>% 
  unite(col = snp, V1:V2, sep = '_')
t_snps <- t(snps)

# Read Line 13 sample ids
indivs <- read.csv("Line_13_samples.012.indv", header = F, sep = "\t")

# Put row and column names into 012 file
row.names(alleles) <- indivs$V1
colnames(alleles) <- t_snps

# Convert to matrix
g.mat_Line_13 <- as.matrix(alleles)

t_g.mat_Line_13 <- t(g.mat_Line_13)
t_g.mat_Line_13 <- cbind(t_g.mat_Line_13, `1332-11-S3` = rep(NA, 51638))

col.order <- c("13_2-F", "1323-S1", "1323-211-S4", 
               "13_3-F", "1332-S1", "1332-1-S2", "1332-11-S3", "1332-111-S4",
               "13_4-F", "1345-S1", "1345-5-S2", "1345-54-S3", "1345-544-S4")
t_g.mat_Line_13 <- t_g.mat_Line_13[,col.order]

my_col <- brewer.pal(5, "Spectral")
pheatmap(t_g.mat_Line_13[1:200,], cluster_rows = F, cluster_cols = F, color = my_col)

df_gmat_Line_13 <- data.frame(t_g.mat_Line_13, check.names = F)

# Filtering Line 13_3
Line_13_3 <- df_gmat_Line_13 %>% 
  rownames_to_column() %>%
  select(1, 5:9) %>% 
  #filter(`1_1-F` == 1, `111-221-S5` == 1, `1_2-F` == 1, `122-122-S4` == 1, `1_4-F` == 1, `145-545-S5` == 1) %>% 
  #filter_all(any_vars (. == 1)) %>% 
  column_to_rownames()

# Correcting Line 13_3
new_13_3 <- zygosity_correction_fun_F_S4(Line_13_3)

new_13_3_cor <- new_13_3 %>%
  rownames_to_column() %>% 
  filter(!(`13_3-F` == "remove")) %>% 
  column_to_rownames()

new_13_3_cor[1:5] <- lapply(new_13_3_cor[1:5], as.numeric)

test_13_3 <- new_13_3_cor %>% 
  rownames_to_column() %>% 
  filter((.[[2]] == 0 | .[[2]] == 2) & .[[3]] == 1 |
           (.[[2]] == 0 | .[[2]] == 2) & .[[4]] == 1 |
           (.[[2]] == 0 | .[[2]] == 2) & .[[5]] == 1 |
           (.[[2]] == 0 | .[[2]] == 2) & .[[6]] == 1 |
           (.[[3]] == 0 | .[[3]] == 2) & .[[4]] == 1 |
           (.[[3]] == 0 | .[[3]] == 2) & .[[5]] == 1 |
           (.[[3]] == 0 | .[[3]] == 2) & .[[6]] == 1 |
           (.[[4]] == 0 | .[[4]] == 2) & .[[5]] == 1 |
           (.[[4]] == 0 | .[[4]] == 2) & .[[6]] == 1 |
           (.[[5]] == 0 | .[[5]] == 2) & .[[6]] == 1) %>%  
  column_to_rownames()

Line_13_3_corrected <- new_13_3_cor %>%
  rownames_to_column() %>% 
  filter(!(row.names(new_13_3_cor) %in% row.names(test_13_3))) %>% 
  column_to_rownames()

# Filtering Line 13_4
Line_13_4 <- df_gmat_Line_13 %>% 
  rownames_to_column() %>%
  select(1, 10:14) %>% 
  #filter(`1_1-F` == 1, `111-221-S5` == 1, `1_2-F` == 1, `122-122-S4` == 1, `1_4-F` == 1, `145-545-S5` == 1) %>% 
  #filter_all(any_vars (. == 1)) %>% 
  column_to_rownames()

# Correcting Line 13_4
new_13_4 <- zygosity_correction_fun_F_S4(Line_13_4)

new_13_4_cor <- new_13_4 %>%
  rownames_to_column() %>% 
  filter(!(`13_4-F` == "remove")) %>% 
  column_to_rownames()

new_13_4_cor[1:5] <- lapply(new_13_4_cor[1:5], as.numeric)

test_13_4 <- new_13_4_cor %>% 
  rownames_to_column() %>% 
  filter((.[[2]] == 0 | .[[2]] == 2) & .[[3]] == 1 |
           (.[[2]] == 0 | .[[2]] == 2) & .[[4]] == 1 |
           (.[[2]] == 0 | .[[2]] == 2) & .[[5]] == 1 |
           (.[[2]] == 0 | .[[2]] == 2) & .[[6]] == 1 |
           (.[[3]] == 0 | .[[3]] == 2) & .[[4]] == 1 |
           (.[[3]] == 0 | .[[3]] == 2) & .[[5]] == 1 |
           (.[[3]] == 0 | .[[3]] == 2) & .[[6]] == 1 |
           (.[[4]] == 0 | .[[4]] == 2) & .[[5]] == 1 |
           (.[[4]] == 0 | .[[4]] == 2) & .[[6]] == 1 |
           (.[[5]] == 0 | .[[5]] == 2) & .[[6]] == 1) %>%  
  column_to_rownames()

Line_13_4_corrected <- new_13_4_cor %>%
  rownames_to_column() %>% 
  filter(!(row.names(new_13_4_cor) %in% row.names(test_13_4))) %>% 
  column_to_rownames()


# how many heterozygotes
table(Line_13_4$`13_4-F`)

table(Line_13_4$`1345-S1`)

table(Line_13_4$`1345-5-S2`)

table(Line_13_4$`1345-54-S3`)

table(Line_13_4$`1345-544-S4`)

#pdf("Line_13_heterozygote_persistence_heatmap.pdf", onefile = T, width = 15, height = 10)
pheatmap(inbred_heterozygotes, cluster_rows = F, cluster_cols = F, color = my_col, show_rownames = F)
dev.off()

#Line 16##################################################################################################################

# Read in Line 16 012 file
alleles <- read.csv("Line_16_samples.012",
                    header = F, sep = "\t", na.strings = -1)
str(alleles)
alleles <- alleles[,-1]

# Read in Line 16 SNP ids
snps <- read.csv("Line_16_samples.012.pos",
                 header = F, sep  = "\t")
snps$V1 <- as.character(snps$V1)
snps$V2 <- as.character(snps$V2)

# Merge columns and transpose
snps <- snps %>% 
  unite(col = snp, V1:V2, sep = '_')
t_snps <- t(snps)

# Read Line 16 sample ids
indivs <- read.csv("Line_16_samples.012.indv", header = F, sep = "\t")

# Put row and column names into 012 file
row.names(alleles) <- indivs$V1
colnames(alleles) <- t_snps

# Convert to matrix
g.mat_Line_16 <- as.matrix(alleles)

t_g.mat_Line_16 <- t(g.mat_Line_16)
t_g.mat_Line_16 <- cbind(t_g.mat_Line_16, `1654-45-S3` = rep(NA, 51638))

col.order <- c("16_1-F", "1611-S1", "1611-2-S2", "1611-21-S3", "1611-211-S4", 
               "16_2-F", "1621-S1", "1621-221-S4",
               "16_5-F", "1654-S1", "1654-4-S2", "1654-45-S3", "1654-454-S4", "1654-454-S5")
t_g.mat_Line_16 <- t_g.mat_Line_16[,col.order]

my_col <- brewer.pal(5, "Spectral")
pheatmap(t_g.mat_Line_16[1:200,], cluster_rows = F, cluster_cols = F, color = my_col)

df_gmat_Line_16 <- data.frame(t_g.mat_Line_16, check.names = F)

# Filtering Line 16_1
Line_16_1 <- df_gmat_Line_16 %>% 
  rownames_to_column() %>%
  select(1:6) %>% 
# filter(((`381-P` == 1 & `367-P` == 1) | (`381-P` == 1 & `367-P` == 0) | (`381-P` == 1 & `367-P` == 2) | 
  #          (`381-P` == 0 & `367-P` == 1) | (`381-P` == 0 & `367-P` == 2) | (`381-P` == 2 & `367-P` == 0) | (`381-P` == 2 & `367-P` == 1)) &
   #        `16_1-F` == 1 & `1611-S1` == 1 & `1611-2-S2` == 1 & `1611-21-S3` == 1 & `1611-211-S4` == 1) %>% 
  column_to_rownames()

# Correcting Line 16_1
new_16_1 <- zygosity_correction_fun_F_S4(Line_16_1)

new_16_1_cor <- new_16_1 %>%
  rownames_to_column() %>% 
  filter(!(`16_1-F` == "remove")) %>% 
  column_to_rownames()

new_16_1_cor[1:5] <- lapply(new_16_1_cor[1:5], as.numeric)

test_16_1 <- new_16_1_cor %>% 
  rownames_to_column() %>% 
  filter((.[[2]] == 0 | .[[2]] == 2) & .[[3]] == 1 |
           (.[[2]] == 0 | .[[2]] == 2) & .[[4]] == 1 |
           (.[[2]] == 0 | .[[2]] == 2) & .[[5]] == 1 |
           (.[[2]] == 0 | .[[2]] == 2) & .[[6]] == 1 |
           (.[[3]] == 0 | .[[3]] == 2) & .[[4]] == 1 |
           (.[[3]] == 0 | .[[3]] == 2) & .[[5]] == 1 |
           (.[[3]] == 0 | .[[3]] == 2) & .[[6]] == 1 |
           (.[[4]] == 0 | .[[4]] == 2) & .[[5]] == 1 |
           (.[[4]] == 0 | .[[4]] == 2) & .[[6]] == 1 |
           (.[[5]] == 0 | .[[5]] == 2) & .[[6]] == 1) %>%  
  column_to_rownames()

Line_16_1_corrected <- new_16_1_cor %>%
  rownames_to_column() %>% 
  filter(!(row.names(new_16_1_cor) %in% row.names(test_16_1))) %>% 
  column_to_rownames()

# Filter Line 16_5
Line_16_5 <- df_gmat_Line_16 %>% 
  rownames_to_column() %>%
  select(1, 10:15) %>%
  # filter(((`263-P` == 1 & `306-P` == 1) | (`263-P` == 1 & `306-P` == 0) | (`263-P` == 1 & `306-P` == 2) | 
  #         (`263-P` == 0 & `306-P` == 1) | (`263-P` == 0 & `306-P` == 2) | (`263-P` == 2 & `306-P` == 0) | (`263-P` == 2 & `306-P` == 1)) & 
  #          `16_5-F` == 1 & `1022-S1` == 1 & `1022-1-S2` == 1 & `1022-113-S4` == 1 & `1022-113-S5` == 1) %>%
  column_to_rownames()

# Correcting Line 16_5
new_16_5 <- zygosity_correction_fun_F_S5(Line_16_5)

new_16_5_cor <- new_16_5 %>%
  rownames_to_column() %>% 
  filter(!(`16_5-F` == "remove")) %>% 
  column_to_rownames()

new_16_5_cor[1:6] <- lapply(new_16_5_cor[1:6], as.numeric)

test_16_5 <- new_16_5_cor %>% 
  rownames_to_column() %>%
  filter((.[[2]] == 0 | .[[2]] == 2) & .[[3]] == 1 |
           (.[[2]] == 0 | .[[2]] == 2) & .[[4]] == 1 |
           (.[[2]] == 0 | .[[2]] == 2) & .[[5]] == 1 |
           (.[[2]] == 0 | .[[2]] == 2) & .[[6]] == 1 |
           (.[[2]] == 0 | .[[2]] == 2) & .[[7]] == 1 |
           (.[[3]] == 0 | .[[3]] == 2) & .[[4]] == 1 |
           (.[[3]] == 0 | .[[3]] == 2) & .[[5]] == 1 |
           (.[[3]] == 0 | .[[3]] == 2) & .[[6]] == 1 |
           (.[[3]] == 0 | .[[3]] == 2) & .[[7]] == 1 |
           (.[[4]] == 0 | .[[4]] == 2) & .[[5]] == 1 |
           (.[[4]] == 0 | .[[4]] == 2) & .[[6]] == 1 |
           (.[[4]] == 0 | .[[4]] == 2) & .[[7]] == 1 |
           (.[[5]] == 0 | .[[5]] == 2) & .[[6]] == 1 |
           (.[[5]] == 0 | .[[5]] == 2) & .[[7]] == 1 |
           (.[[6]] == 0 | .[[6]] == 2) & .[[7]] == 1) %>%  
  column_to_rownames()

Line_16_5_corrected <- new_16_5_cor %>% 
  rownames_to_column() %>% 
  filter(!(row.names(new_16_5_cor) %in% row.names(test_16_5))) %>% 
  column_to_rownames()


# how many heterozygotes
table(Line_16_1$`16_1-F`)

table(Line_16_1$`1611-S1`)

table(Line_16_1$`1611-2-S2`)

table(Line_16_1$`1611-21-S3`)

table(Line_16_1$`1611-211-S4`)

# Line_16_2 <- df_gmat_Line_16 %>% 
#   rownames_to_column() %>%
#   select(1:3, 9:11) %>% 
#   filter(((`381-P` == 1 & `367-P` == 1) | (`381-P` == 1 & `367-P` == 0) | (`381-P` == 1 & `367-P` == 2) | 
#             (`381-P` == 0 & `367-P` == 1) | (`381-P` == 0 & `367-P` == 2) | (`381-P` == 2 & `367-P` == 0) | (`381-P` == 2 & `367-P` == 1)) & 
#            `16_2-F` == 1 & `1621-S1` == 1 & `1621-221-S4` == 1) %>% 
#   column_to_rownames()
# 
# Line_16_5 <- df_gmat_Line_16 %>% 
#   rownames_to_column() %>%
#   select(1:3, 12:16) %>% 
#   filter(((`381-P` == 1 & `367-P` == 1) | (`381-P` == 1 & `367-P` == 0) | (`381-P` == 1 & `367-P` == 2) | 
#             (`381-P` == 0 & `367-P` == 1) | (`381-P` == 0 & `367-P` == 2) | (`381-P` == 2 & `367-P` == 0) | (`381-P` == 2 & `367-P` == 1)) &
#            `16_5-F` == 1 & `1654-S1` == 1 & `1654-4-S2` == 1 & `1654-454-S4` == 1 & `1654-454-S5` == 1) %>% 
#   column_to_rownames()

table(Line_16_1$`16_1-F`)
# pdf("Line_16_heterozygote_persistence_heatmap.pdf", onefile = T, width = 15, height = 10)
pheatmap(inbred_heterozygotes, cluster_rows = F, cluster_cols = F, color = my_col, show_rownames = F)
dev.off()

# write.csv(Line_16_1, "snp_zygosity_analysis/Line_16_1_heterozygote_persistence.txt")
# write.csv(Line_16_2, "snp_zygosity_analysis/Line_16_2_heterozygote_persistence.txt")
# write.csv(Line_16_5, "snp_zygosity_analysis/Line_16_5_heterozygote_persistence.txt")
# 
# write.csv(rownames(Line_16_1), "snp_zygosity_analysis/Line_16_1_heterozygote_persistence_parents_any_zygosity_snps.txt", row.names = F, col.names = F)
# write.csv(rownames(Line_16_2), "snp_zygosity_analysis/Line_16_2_heterozygote_persistence_parents_any_zygosity_snps.txt", row.names = F, col.names = F)
# write.csv(rownames(Line_16_5), "snp_zygosity_analysis/Line_16_5_heterozygote_persistence_parents_any_zygosity_snps.txt", row.names = F, col.names = F)
#Line 17##################################################################################################################

# Read in Line 17 012 file
alleles <- read.csv("Line_17_samples.012",
                    header = F, sep = "\t", na.strings = -1)
str(alleles)
alleles <- alleles[,-1]

# Read in Line 17 SNP ids
snps <- read.csv("Line_17_samples.012.pos",
                 header = F, sep  = "\t")
snps$V1 <- as.character(snps$V1)
snps$V2 <- as.character(snps$V2)

# Merge columns and transpose
snps <- snps %>% 
  unite(col = snp, V1:V2, sep = '_')
t_snps <- t(snps)

# Read Line 17 sample ids
indivs <- read.csv("Line_17_samples.012.indv", header = F, sep = "\t")

# Put row and column names into 012 file
row.names(alleles) <- indivs$V1
colnames(alleles) <- t_snps

# Convert to matrix
g.mat_Line_17 <- as.matrix(alleles)

t_g.mat_Line_17 <- t(g.mat_Line_17)
col.order <- c("17_2-F", "1721-S1", "1721-1-S2", "1721-11-S3", "1721-111-S4", "1721-111-S5",
               "17_4-F", "1744-S1", "1744-544-S4",
               "17_5-F", "1755-S1", "1755-5-S2", "1755-54-S3", "1755-545-S4", "1755-545-S5")
t_g.mat_Line_17 <- t_g.mat_Line_17[,col.order]

my_col <- brewer.pal(5, "Spectral")
pheatmap(t_g.mat_Line_17[1:200,], cluster_rows = F, cluster_cols = F, color = my_col)

df_gmat_Line_17 <- data.frame(t_g.mat_Line_17, check.names = F)

# Filtering Line 17_2
Line_17_2 <- df_gmat_Line_17 %>% 
  rownames_to_column() %>%
  select(1:7) %>% 
  # filter(((`386-P` == 1 & `253-P` == 1) | (`386-P` == 1 & `253-P` == 0) | (`386-P` == 1 & `253-P` == 2) | 
  #           (`386-P` == 0 & `253-P` == 1) | (`386-P` == 0 & `253-P` == 2) | (`386-P` == 2 & `253-P` == 0) | (`386-P` == 2 & `253-P` == 1)) & 
  #          `17_2-F` == 1 & `1721-S1` == 1 & `1721-1-S2` == 1 & `1721-11-S3` == 1 & `1721-111-S4` == 1 & `1721-111-S5` == 1) %>% 
  column_to_rownames()


# Correcting Line 17_2
new_17_2 <- zygosity_correction_fun_F_S5(Line_17_2)

new_17_2_cor <- new_17_2 %>%
  rownames_to_column() %>% 
  filter(!(`17_2-F` == "remove")) %>% 
  column_to_rownames()

new_17_2_cor[1:6] <- lapply(new_17_2_cor[1:6], as.numeric)

test_17_2 <- new_17_2_cor %>% 
  rownames_to_column() %>% 
  filter((.[[2]] == 0 | .[[2]] == 2) & .[[3]] == 1 |
           (.[[2]] == 0 | .[[2]] == 2) & .[[4]] == 1 |
           (.[[2]] == 0 | .[[2]] == 2) & .[[5]] == 1 |
           (.[[2]] == 0 | .[[2]] == 2) & .[[6]] == 1 |
           (.[[2]] == 0 | .[[2]] == 2) & .[[7]] == 1 |
           (.[[3]] == 0 | .[[3]] == 2) & .[[4]] == 1 |
           (.[[3]] == 0 | .[[3]] == 2) & .[[5]] == 1 |
           (.[[3]] == 0 | .[[3]] == 2) & .[[6]] == 1 |
           (.[[3]] == 0 | .[[3]] == 2) & .[[7]] == 1 |
           (.[[4]] == 0 | .[[4]] == 2) & .[[5]] == 1 |
           (.[[4]] == 0 | .[[4]] == 2) & .[[6]] == 1 |
           (.[[4]] == 0 | .[[4]] == 2) & .[[7]] == 1 |
           (.[[5]] == 0 | .[[5]] == 2) & .[[6]] == 1 |
           (.[[5]] == 0 | .[[5]] == 2) & .[[7]] == 1 |
           (.[[6]] == 0 | .[[6]] == 2) & .[[7]] == 1) %>% 
  column_to_rownames()

Line_17_2_corrected <- new_17_2_cor %>%
  rownames_to_column() %>% 
  filter(!(row.names(new_17_2_cor) %in% row.names(test_17_2))) %>% 
  column_to_rownames()


# Filtering Line 17_5
Line_17_5 <- df_gmat_Line_17 %>% 
  rownames_to_column() %>%
  select(1, 11:16) %>% 
  # filter(((`386-P` == 1 & `253-P` == 1) | (`386-P` == 1 & `253-P` == 0) | (`386-P` == 1 & `253-P` == 2) | 
  #           (`386-P` == 0 & `253-P` == 1) | (`386-P` == 0 & `253-P` == 2) | (`386-P` == 2 & `253-P` == 0) | (`386-P` == 2 & `253-P` == 1)) & 
  #          `17_5-F` == 1 & `1721-S1` == 1 & `1721-1-S2` == 1 & `1721-11-S3` == 1 & `1721-111-S4` == 1 & `1721-111-S5` == 1) %>% 
  column_to_rownames()


# Correcting Line 17_5
new_17_5 <- zygosity_correction_fun_F_S5(Line_17_5)

new_17_5_cor <- new_17_5 %>%
  rownames_to_column() %>% 
  filter(!(`17_5-F` == "remove")) %>% 
  column_to_rownames()

new_17_5_cor[1:6] <- lapply(new_17_5_cor[1:6], as.numeric)

test_17_5 <- new_17_5_cor %>% 
  rownames_to_column() %>% 
  filter((.[[2]] == 0 | .[[2]] == 2) & .[[3]] == 1 |
           (.[[2]] == 0 | .[[2]] == 2) & .[[4]] == 1 |
           (.[[2]] == 0 | .[[2]] == 2) & .[[5]] == 1 |
           (.[[2]] == 0 | .[[2]] == 2) & .[[6]] == 1 |
           (.[[2]] == 0 | .[[2]] == 2) & .[[7]] == 1 |
           (.[[3]] == 0 | .[[3]] == 2) & .[[4]] == 1 |
           (.[[3]] == 0 | .[[3]] == 2) & .[[5]] == 1 |
           (.[[3]] == 0 | .[[3]] == 2) & .[[6]] == 1 |
           (.[[3]] == 0 | .[[3]] == 2) & .[[7]] == 1 |
           (.[[4]] == 0 | .[[4]] == 2) & .[[5]] == 1 |
           (.[[4]] == 0 | .[[4]] == 2) & .[[6]] == 1 |
           (.[[4]] == 0 | .[[4]] == 2) & .[[7]] == 1 |
           (.[[5]] == 0 | .[[5]] == 2) & .[[6]] == 1 |
           (.[[5]] == 0 | .[[5]] == 2) & .[[7]] == 1 |
           (.[[6]] == 0 | .[[6]] == 2) & .[[7]] == 1) %>% 
  column_to_rownames()

Line_17_5_corrected <- new_17_5_cor %>%
  rownames_to_column() %>% 
  filter(!(row.names(new_17_5_cor) %in% row.names(test_17_5))) %>% 
  column_to_rownames()


# how many heterozygotes
table(Line_17_2$`17_2-F`)

table(Line_17_2$`1721-S1`)

table(Line_17_2$`1721-1-S2`)

table(Line_17_2$`1721-11-S3`)

table(Line_17_2$`1721-111-S4`)

table(Line_17_2$`1721-111-S5`)

# Line_17_5 <- df_gmat_Line_17 %>% 
#   rownames_to_column() %>%
#   select(1:3, 10:13, 15) %>% 
#   filter(((`386-P` == 1 & `253-P` == 1) | (`386-P` == 1 & `253-P` == 0) | (`386-P` == 1 & `253-P` == 2) | 
#             (`386-P` == 0 & `253-P` == 1) | (`386-P` == 0 & `253-P` == 2) | (`386-P` == 2 & `253-P` == 0) | (`386-P` == 2 & `253-P` == 1)) & 
#            `17_5-F` == 1 & `1755-S1` == 1 & `1755-5-S2` == 1 & `1755-54-S3` == 1 & `1755-545-S5` == 1) %>% 
#   column_to_rownames()


table(Line_17_2$`17_2-F`)

# pdf("Line_17_heterozygote_persistence_heatmap.pdf", onefile = T, width = 15, height = 10)
pheatmap(inbred_heterozygotes, cluster_rows = F, cluster_cols = F, color = my_col, show_rownames = F)
dev.off()

# write.csv(Line_17_2, "snp_zygosity_analysis/Line_17_2_heterozygote_persistence.txt")
# write.csv(Line_17_5, "snp_zygosity_analysis/Line_17_5_heterozygote_persistence.txt")
# 
# write.csv(rownames(Line_17_2), "snp_zygosity_analysis/Line_17_2_heterozygote_persistence_parents_any_zygosity_snps.txt", row.names = F, col.names = F)
# write.csv(rownames(Line_17_5), "snp_zygosity_analysis/Line_17_5_heterozygote_persistence_parents_any_zygosity_snps.txt", row.names = F, col.names = F)
#Line 19##################################################################################################################

# Read in Line 19 012 file
alleles <- read.csv("Line_19_samples.012",
                    header = F, sep = "\t", na.strings = -1)
str(alleles)
alleles <- alleles[,-1]

# Read in Line 19 SNP ids
snps <- read.csv("Line_19_samples.012.pos",
                 header = F, sep  = "\t")
snps$V1 <- as.character(snps$V1)
snps$V2 <- as.character(snps$V2)

# Merge columns and transpose
snps <- snps %>% 
  unite(col = snp, V1:V2, sep = '_')
t_snps <- t(snps)

# Read Line 19 sample ids
indivs <- read.csv("Line_19_samples.012.indv", header = F, sep = "\t")

# Put row and column names into 012 file
row.names(alleles) <- indivs$V1
colnames(alleles) <- t_snps

# Convert to matrix
g.mat_Line_19 <- as.matrix(alleles)

t_g.mat_Line_19 <- t(g.mat_Line_19)
t_g.mat_Line_19 <- cbind(t_g.mat_Line_19, `1922-11-S3` = rep(NA, 51638))

col.order <- c("19_1-F", "1912-S1", "1912-131-S4", 
               "19_2-F", "1922-S1", "1922-1-S2", "1922-11-S3", "1922-111-S4",
               "19_5-F", "1955-S1", "1955-5-S2", "1955-54-S3", "1955-544-S4")
t_g.mat_Line_19 <- t_g.mat_Line_19[,col.order]

my_col <- brewer.pal(5, "Spectral")
pheatmap(t_g.mat_Line_19[1:200,], cluster_rows = F, cluster_cols = F, color = my_col)

df_gmat_Line_19 <- data.frame(t_g.mat_Line_19, check.names = F)

# Filtering Line 19_2
Line_19_2 <- df_gmat_Line_19 %>% 
  rownames_to_column() %>%
  select(1, 5:9) %>% 
  # filter(((`386-P` == 1 & `253-P` == 1) | (`386-P` == 1 & `253-P` == 0) | (`386-P` == 1 & `253-P` == 2) | 
  #           (`386-P` == 0 & `253-P` == 1) | (`386-P` == 0 & `253-P` == 2) | (`386-P` == 2 & `253-P` == 0) | (`386-P` == 2 & `253-P` == 1)) & 
  #          `17_2-F` == 1 & `1721-S1` == 1 & `1721-1-S2` == 1 & `1721-11-S3` == 1 & `1721-111-S4` == 1 & `1721-111-S5` == 1) %>% 
  column_to_rownames()

# Correcting Line 19_2
new_19_2 <- zygosity_correction_fun_F_S4(Line_19_2)

new_19_2_cor <- new_19_2 %>%
  rownames_to_column() %>% 
  filter(!(`19_2-F` == "remove")) %>% 
  column_to_rownames()

new_19_2_cor[1:5] <- lapply(new_19_2_cor[1:5], as.numeric)

test_19_2 <- new_19_2_cor %>% 
  rownames_to_column() %>% 
  filter((.[[2]] == 0 | .[[2]] == 2) & .[[3]] == 1 |
           (.[[2]] == 0 | .[[2]] == 2) & .[[4]] == 1 |
           (.[[2]] == 0 | .[[2]] == 2) & .[[5]] == 1 |
           (.[[2]] == 0 | .[[2]] == 2) & .[[6]] == 1 |
           (.[[3]] == 0 | .[[3]] == 2) & .[[4]] == 1 |
           (.[[3]] == 0 | .[[3]] == 2) & .[[5]] == 1 |
           (.[[3]] == 0 | .[[3]] == 2) & .[[6]] == 1 |
           (.[[4]] == 0 | .[[4]] == 2) & .[[5]] == 1 |
           (.[[4]] == 0 | .[[4]] == 2) & .[[6]] == 1 |
           (.[[5]] == 0 | .[[5]] == 2) & .[[6]] == 1) %>%  
  column_to_rownames()

Line_19_2_corrected <- new_19_2_cor %>%
  rownames_to_column() %>% 
  filter(!(row.names(new_19_2_cor) %in% row.names(test_19_2))) %>% 
  column_to_rownames()

# Filtering Line 19_5
Line_19_5 <- df_gmat_Line_19 %>% 
  rownames_to_column() %>%
  select(1, 10:14) %>% 
  # filter(((`386-P` == 1 & `253-P` == 1) | (`386-P` == 1 & `253-P` == 0) | (`386-P` == 1 & `253-P` == 2) | 
  #           (`386-P` == 0 & `253-P` == 1) | (`386-P` == 0 & `253-P` == 2) | (`386-P` == 2 & `253-P` == 0) | (`386-P` == 2 & `253-P` == 1)) & 
  #          `17_2-F` == 1 & `1721-S1` == 1 & `1721-1-S2` == 1 & `1721-11-S3` == 1 & `1721-111-S4` == 1 & `1721-111-S5` == 1) %>% 
  column_to_rownames()

# Correcting Line 19_5
new_19_5 <- zygosity_correction_fun_F_S4(Line_19_5)

new_19_5_cor <- new_19_5 %>%
  rownames_to_column() %>% 
  filter(!(`19_5-F` == "remove")) %>% 
  column_to_rownames()

new_19_5_cor[1:5] <- lapply(new_19_5_cor[1:5], as.numeric)

test_19_5 <- new_19_5_cor %>% 
  rownames_to_column() %>% 
  filter((.[[2]] == 0 | .[[2]] == 2) & .[[3]] == 1 |
           (.[[2]] == 0 | .[[2]] == 2) & .[[4]] == 1 |
           (.[[2]] == 0 | .[[2]] == 2) & .[[5]] == 1 |
           (.[[2]] == 0 | .[[2]] == 2) & .[[6]] == 1 |
           (.[[3]] == 0 | .[[3]] == 2) & .[[4]] == 1 |
           (.[[3]] == 0 | .[[3]] == 2) & .[[5]] == 1 |
           (.[[3]] == 0 | .[[3]] == 2) & .[[6]] == 1 |
           (.[[4]] == 0 | .[[4]] == 2) & .[[5]] == 1 |
           (.[[4]] == 0 | .[[4]] == 2) & .[[6]] == 1 |
           (.[[5]] == 0 | .[[5]] == 2) & .[[6]] == 1) %>%  
  column_to_rownames()

Line_19_5_corrected <- new_19_5_cor %>%
  rownames_to_column() %>% 
  filter(!(row.names(new_19_5_cor) %in% row.names(test_19_5))) %>% 
  column_to_rownames()


# how many heterozygotes
table(Line_19_5$`19_5-F`)

table(Line_19_5$`1955-S1`)

table(Line_19_5$`1955-5-S2`)

table(Line_19_5$`1955-54-S3`)

table(Line_19_5$`1955-544-S4`)

inbred_heterozygotes <- df_gmat_Line_19 %>% 
  rownames_to_column() %>%
  #select(1:7) %>% 
  filter(`1_1-F` == 1, `111-221-S5` == 1, `1_2-F` == 1, `122-122-S4` == 1, `1_4-F` == 1, `145-545-S5` == 1) %>% 
  #filter_all(any_vars (. == 1)) %>% 
  column_to_rownames()

table(Line_19_5$`19_5-F`)

# pdf("Line_19_heterozygote_persistence_heatmap.pdf", onefile = T, width = 15, height = 10)
pheatmap(inbred_heterozygotes, cluster_rows = F, cluster_cols = F, color = my_col, show_rownames = F)
dev.off()

#Line 20##################################################################################################################

# Read in Line 20 012 file
alleles <- read.csv("Line_20_samples.012",
                    header = F, sep = "\t", na.strings = -1)
str(alleles)
alleles <- alleles[,-1]

# Read in Line 20 SNP ids
snps <- read.csv("Line_20_samples.012.pos",
                 header = F, sep  = "\t")
snps$V1 <- as.character(snps$V1)
snps$V2 <- as.character(snps$V2)

# Merge columns and transpose
snps <- snps %>% 
  unite(col = snp, V1:V2, sep = '_')
t_snps <- t(snps)

# Read Line 20 sample ids
indivs <- read.csv("Line_20_samples.012.indv", header = F, sep = "\t")

# Put row and column names into 012 file
row.names(alleles) <- indivs$V1
colnames(alleles) <- t_snps

# Convert to matrix
g.mat_Line_20 <- as.matrix(alleles)

t_g.mat_Line_20 <- t(g.mat_Line_20)
col.order <- c("20_1-F", "2013-S1", "2013-1-S2", "2013-13-S3", "2013-131-S4",
               "20_4-F", "2045-S1", "2045-5-S2", "2045-54-S3", "2045-544-S4")
t_g.mat_Line_20 <- t_g.mat_Line_20[,col.order]

my_col <- brewer.pal(5, "Spectral")
pheatmap(t_g.mat_Line_20[1:200,], cluster_rows = F, cluster_cols = F, color = my_col)

df_gmat_Line_20 <- data.frame(t_g.mat_Line_20, check.names = F)

# Filter Line 20_1
Line_20_1 <- df_gmat_Line_20 %>% 
  rownames_to_column() %>%
  select(1:6) %>% 
  column_to_rownames()

# Correcting Line 20_1
new_20_1 <- zygosity_correction_fun_F_S4(Line_20_1)

new_20_1_cor <- new_20_1 %>%
  rownames_to_column() %>% 
  filter(!(`20_1-F` == "remove")) %>% 
  column_to_rownames()

new_20_1_cor[1:5] <- lapply(new_20_1_cor[1:5], as.numeric)

test_20_1 <- new_20_1_cor %>% 
  rownames_to_column() %>% 
  filter((.[[2]] == 0 | .[[2]] == 2) & .[[3]] == 1 |
           (.[[2]] == 0 | .[[2]] == 2) & .[[4]] == 1 |
           (.[[2]] == 0 | .[[2]] == 2) & .[[5]] == 1 |
           (.[[2]] == 0 | .[[2]] == 2) & .[[6]] == 1 |
           (.[[3]] == 0 | .[[3]] == 2) & .[[4]] == 1 |
           (.[[3]] == 0 | .[[3]] == 2) & .[[5]] == 1 |
           (.[[3]] == 0 | .[[3]] == 2) & .[[6]] == 1 |
           (.[[4]] == 0 | .[[4]] == 2) & .[[5]] == 1 |
           (.[[4]] == 0 | .[[4]] == 2) & .[[6]] == 1 |
           (.[[5]] == 0 | .[[5]] == 2) & .[[6]] == 1) %>%  
  column_to_rownames()

Line_20_1_corrected <- new_20_1_cor %>%
  rownames_to_column() %>% 
  filter(!(row.names(new_20_1_cor) %in% row.names(test_20_1))) %>% 
  column_to_rownames()

# Filter Line 20_4
Line_20_4 <- df_gmat_Line_20 %>% 
  rownames_to_column() %>%
  select(1, 7:11) %>% 
  column_to_rownames()

# Correcting Line 20_4
new_20_4 <- zygosity_correction_fun_F_S4(Line_20_4)

new_20_4_cor <- new_20_4 %>%
  rownames_to_column() %>% 
  filter(!(`20_4-F` == "remove")) %>% 
  column_to_rownames()

new_20_4_cor[1:5] <- lapply(new_20_4_cor[1:5], as.numeric)

test_20_4 <- new_20_4_cor %>% 
  rownames_to_column() %>% 
  filter((.[[2]] == 0 | .[[2]] == 2) & .[[3]] == 1 |
           (.[[2]] == 0 | .[[2]] == 2) & .[[4]] == 1 |
           (.[[2]] == 0 | .[[2]] == 2) & .[[5]] == 1 |
           (.[[2]] == 0 | .[[2]] == 2) & .[[6]] == 1 |
           (.[[3]] == 0 | .[[3]] == 2) & .[[4]] == 1 |
           (.[[3]] == 0 | .[[3]] == 2) & .[[5]] == 1 |
           (.[[3]] == 0 | .[[3]] == 2) & .[[6]] == 1 |
           (.[[4]] == 0 | .[[4]] == 2) & .[[5]] == 1 |
           (.[[4]] == 0 | .[[4]] == 2) & .[[6]] == 1 |
           (.[[5]] == 0 | .[[5]] == 2) & .[[6]] == 1) %>%  
  column_to_rownames()

Line_20_4_corrected <- new_20_4_cor %>%
  rownames_to_column() %>% 
  filter(!(row.names(new_20_4_cor) %in% row.names(test_20_4))) %>% 
  column_to_rownames()


# how many heterozygotes
table(Line_20_1$`20_1-F`)

table(Line_20_1$`2013-S1`)

table(Line_20_1$`2013-1-S2`)

table(Line_20_1$`2013-13-S3`)

table(Line_20_1$`2013-131-S4`)

table(Line_20_4$`20_4-F`)

table(Line_20_4$`2045-S1`)

table(Line_20_4$`2045-5-S2`)

table(Line_20_4$`2045-54-S3`)

table(Line_20_4$`2045-544-S4`)

# pdf("Line_20_heterozygote_persistence_heatmap.pdf", onefile = T, width = 15, height = 10)
pheatmap(inbred_heterozygotes, cluster_rows = F, cluster_cols = F, color = my_col, show_rownames = F)
dev.off()

#Line 21##################################################################################################################

# Read in Line 21 012 file
alleles <- read.csv("Line_21_samples.012",
                    header = F, sep = "\t", na.strings = -1)
str(alleles)
alleles <- alleles[,-1]

# Read in Line 21 SNP ids
snps <- read.csv("Line_21_samples.012.pos",
                 header = F, sep  = "\t")
snps$V1 <- as.character(snps$V1)
snps$V2 <- as.character(snps$V2)

# Merge columns and transpose
snps <- snps %>% 
  unite(col = snp, V1:V2, sep = '_')
t_snps <- t(snps)

# Read Line 21 sample ids
indivs <- read.csv("Line_21_samples.012.indv", header = F, sep = "\t")

# Put row and column names into 012 file
row.names(alleles) <- indivs$V1
colnames(alleles) <- t_snps

# Convert to matrix
g.mat_Line_21 <- as.matrix(alleles)

t_g.mat_Line_21 <- t(g.mat_Line_21)
col.order <- c("2111-S1", "2111-111-S4", 
               "21_2-F", "2121-S1", "2121-1-S2", "2121-11-S3", "2121-111-S4", "2121-111-S5",
               "21_6-F", "2165-S1", "2165-4-S2", "2165-44-S3", "2165-444-S4", "2165-444-S5")
t_g.mat_Line_21 <- t_g.mat_Line_21[,col.order]

my_col <- brewer.pal(5, "Spectral")
pheatmap(t_g.mat_Line_21[1:200,], cluster_rows = F, cluster_cols = F, color = my_col)

df_gmat_Line_21 <- data.frame(t_g.mat_Line_21, check.names = F)

# Line_21_1 <- df_gmat_Line_21 %>% 
#   rownames_to_column() %>%
#   select(1:6) %>% 
#   filter(((`435-P` == 1 & `15-P` == 1) | (`435-P` == 1 & `15-P` == 0) | (`435-P` == 1 & `15-P` == 2) | 
#             (`435-P` == 0 & `15-P` == 1) | (`435-P` == 0 & `15-P` == 2) | (`435-P` == 2 & `15-P` == 0) | (`435-P` == 2 & `15-P` == 1)) & 
#            `21_1-F` == 1 & `2111-S1` == 1 & `2111-111-S4`) %>% 
#   column_to_rownames()

# Filtering Line 21_2
Line_21_2 <- df_gmat_Line_21 %>% 
  rownames_to_column() %>%
  select(1, 4:9) %>% 
  # filter(((`435-P` == 1 & `15-P` == 1) | (`435-P` == 1 & `15-P` == 0) | (`435-P` == 1 & `15-P` == 2) | 
  #           (`435-P` == 0 & `15-P` == 1) | (`435-P` == 0 & `15-P` == 2) | (`435-P` == 2 & `15-P` == 0) | (`435-P` == 2 & `15-P` == 1)) & 
  #          `21_2-F` == 1 & `2121-S1` == 1 & `2121-1-S2` == 1 & `2121-11-S3` == 1 & `2121-111-S4` & `2121-111-S5` == 1) %>% 
  column_to_rownames()

# Correcting Line 21_2
new_21_2 <- zygosity_correction_fun_F_S5(Line_21_2)

new_21_2_cor <- new_21_2 %>%
  rownames_to_column() %>% 
  filter(!(`21_2-F` == "remove")) %>% 
  column_to_rownames()

new_21_2_cor[1:6] <- lapply(new_21_2_cor[1:6], as.numeric)

test_21_2 <- new_21_2_cor %>% 
  rownames_to_column() %>% 
  filter((.[[2]] == 0 | .[[2]] == 2) & .[[3]] == 1 |
           (.[[2]] == 0 | .[[2]] == 2) & .[[4]] == 1 |
           (.[[2]] == 0 | .[[2]] == 2) & .[[5]] == 1 |
           (.[[2]] == 0 | .[[2]] == 2) & .[[6]] == 1 |
           (.[[2]] == 0 | .[[2]] == 2) & .[[7]] == 1 |
           (.[[3]] == 0 | .[[3]] == 2) & .[[4]] == 1 |
           (.[[3]] == 0 | .[[3]] == 2) & .[[5]] == 1 |
           (.[[3]] == 0 | .[[3]] == 2) & .[[6]] == 1 |
           (.[[3]] == 0 | .[[3]] == 2) & .[[7]] == 1 |
           (.[[4]] == 0 | .[[4]] == 2) & .[[5]] == 1 |
           (.[[4]] == 0 | .[[4]] == 2) & .[[6]] == 1 |
           (.[[4]] == 0 | .[[4]] == 2) & .[[7]] == 1 |
           (.[[5]] == 0 | .[[5]] == 2) & .[[6]] == 1 |
           (.[[5]] == 0 | .[[5]] == 2) & .[[7]] == 1 |
           (.[[6]] == 0 | .[[6]] == 2) & .[[7]] == 1) %>% 
  column_to_rownames()

Line_21_2_corrected <- new_21_2_cor %>%
  rownames_to_column() %>% 
  filter(!(row.names(new_21_2_cor) %in% row.names(test_21_2))) %>% 
  column_to_rownames()

# Filtering Line 21_6
Line_21_6 <- df_gmat_Line_21 %>% 
  rownames_to_column() %>%
  select(1, 10:15) %>% 
  # filter(((`435-P` == 1 & `15-P` == 1) | (`435-P` == 1 & `15-P` == 0) | (`435-P` == 1 & `15-P` == 2) | 
  #           (`435-P` == 0 & `15-P` == 1) | (`435-P` == 0 & `15-P` == 2) | (`435-P` == 2 & `15-P` == 0) | (`435-P` == 2 & `15-P` == 1)) & 
  #          `21_6-F` == 1 & `2165-S1` == 1 & `2165-4-S2` == 1 & `2165-44-S3` == 1 & `2165-444-S4` & `2165-444-S5` == 1) %>%
  column_to_rownames()

# Correcting Line 21_6
new_21_6 <- zygosity_correction_fun_F_S5(Line_21_6)

new_21_6_cor <- new_21_6 %>%
  rownames_to_column() %>% 
  filter(!(`21_6-F` == "remove")) %>% 
  column_to_rownames()

new_21_6_cor[1:6] <- lapply(new_21_6_cor[1:6], as.numeric)

test_21_6 <- new_21_6_cor %>% 
  rownames_to_column() %>% 
  filter((.[[2]] == 0 | .[[2]] == 2) & .[[3]] == 1 |
           (.[[2]] == 0 | .[[2]] == 2) & .[[4]] == 1 |
           (.[[2]] == 0 | .[[2]] == 2) & .[[5]] == 1 |
           (.[[2]] == 0 | .[[2]] == 2) & .[[6]] == 1 |
           (.[[2]] == 0 | .[[2]] == 2) & .[[7]] == 1 |
           (.[[3]] == 0 | .[[3]] == 2) & .[[4]] == 1 |
           (.[[3]] == 0 | .[[3]] == 2) & .[[5]] == 1 |
           (.[[3]] == 0 | .[[3]] == 2) & .[[6]] == 1 |
           (.[[3]] == 0 | .[[3]] == 2) & .[[7]] == 1 |
           (.[[4]] == 0 | .[[4]] == 2) & .[[5]] == 1 |
           (.[[4]] == 0 | .[[4]] == 2) & .[[6]] == 1 |
           (.[[4]] == 0 | .[[4]] == 2) & .[[7]] == 1 |
           (.[[5]] == 0 | .[[5]] == 2) & .[[6]] == 1 |
           (.[[5]] == 0 | .[[5]] == 2) & .[[7]] == 1 |
           (.[[6]] == 0 | .[[6]] == 2) & .[[7]] == 1) %>% 
  column_to_rownames()

Line_21_6_corrected <- new_21_6_cor %>%
  rownames_to_column() %>% 
  filter(!(row.names(new_21_6_cor) %in% row.names(test_21_6))) %>% 
  column_to_rownames()


# how many heterozygotes
table(Line_21_2$`21_2-F`)

table(Line_21_2$`2121-S1`)

table(Line_21_2$`2121-1-S2`)

table(Line_21_2$`2121-11-S3`)

table(Line_21_2$`2121-111-S4`)

table(Line_21_2$`2121-111-S5`)

table(Line_21_6$`21_6-F`)

table(Line_21_6$`2165-S1`)

table(Line_21_6$`2165-4-S2`)

table(Line_21_6$`2165-44-S3`)

table(Line_21_6$`2165-444-S4`)

table(Line_21_6$`2165-444-S5`)

# pdf("Line_21_heterozygote_persistence_heatmap.pdf", onefile = T, width = 15, height = 10)
pheatmap(inbred_heterozygotes, cluster_rows = F, cluster_cols = F, color = my_col, show_rownames = F)
dev.off()

# write.csv(Line_21_1, "snp_zygosity_analysis/Line_21_1_heterozygote_persistence.txt")
# write.csv(Line_21_2, "snp_zygosity_analysis/Line_21_2_heterozygote_persistence.txt")
# write.csv(Line_21_6, "snp_zygosity_analysis/Line_21_6_heterozygote_persistence.txt")
# 
# write.csv(rownames(Line_21_1), "snp_zygosity_analysis/Line_21_1_heterozygote_persistence_parents_any_zygosity_snps.txt", row.names = F, col.names = F)
# write.csv(rownames(Line_21_2), "snp_zygosity_analysis/Line_21_2_heterozygote_persistence_parents_any_zygosity_snps.txt", row.names = F, col.names = F)
# write.csv(rownames(Line_21_6), "snp_zygosity_analysis/Line_21_6_heterozygote_persistence_parents_any_zygosity_snps.txt", row.names = F, col.names = F)
##Line 23#################################################################################################################

# Read in Line 23 012 file
alleles <- read.csv("Line_23_samples.012",
                    header = F, sep = "\t", na.strings = -1)
str(alleles)
alleles <- alleles[,-1]

# Read in Line 23 SNP ids
snps <- read.csv("Line_23_samples.012.pos",
                 header = F, sep  = "\t")
snps$V1 <- as.character(snps$V1)
snps$V2 <- as.character(snps$V2)

# Merge columns and transpose
snps <- snps %>% 
  unite(col = snp, V1:V2, sep = '_')
t_snps <- t(snps)

# Read Line 23 sample ids
indivs <- read.csv("Line_23_samples.012.indv", header = F, sep = "\t")

# Put row and column names into 012 file
row.names(alleles) <- indivs$V1
colnames(alleles) <- t_snps

# Convert to matrix
g.mat_Line_23 <- as.matrix(alleles)

t_g.mat_Line_23 <- t(g.mat_Line_23)
col.order <- c("23_2-F", "2323-S1", "2323-2-S2", "2323-21-S3", "2323-211-S4", "2323-211-S5",
               "23_4-F", "2344-S1", "2344-4-S2", "2344-46-S3", "2344-464-S4")
t_g.mat_Line_23 <- t_g.mat_Line_23[,col.order]

my_col <- brewer.pal(5, "Spectral")
pheatmap(t_g.mat_Line_23[1:200,], cluster_rows = F, cluster_cols = F, color = my_col)

df_gmat_Line_23 <- data.frame(t_g.mat_Line_23, check.names = F)

# Filtering Line 23_2
Line_23_2 <- df_gmat_Line_23 %>% 
  rownames_to_column() %>%
  select(1:7) %>% 
  # filter(((`461-P` == 1 & `5-P` == 1) | (`461-P` == 1 & `5-P` == 0) | (`461-P` == 1 & `5-P` == 2) | 
  #           (`461-P` == 0 & `5-P` == 1) | (`461-P` == 0 & `5-P` == 2) | (`461-P` == 2 & `5-P` == 0) | (`461-P` == 2 & `5-P` == 1)) & 
  #          `23_2-F` == 1 & `2323-S1` == 1 & `2323-2-S2` == 1 & `2323-21-S3` == 1 & `2323-211-S4` & `2323-211-S5` == 1) %>% 
  column_to_rownames()

# Correcting Line 23_2
new_23_2 <- zygosity_correction_fun_F_S5(Line_23_2)

new_23_2_cor <- new_23_2 %>%
  rownames_to_column() %>% 
  filter(!(`23_2-F` == "remove")) %>% 
  column_to_rownames()

new_23_2_cor[1:6] <- lapply(new_23_2_cor[1:6], as.numeric)

test_23_2 <- new_23_2_cor %>% 
  rownames_to_column() %>% 
  filter((.[[2]] == 0 | .[[2]] == 2) & .[[3]] == 1 |
           (.[[2]] == 0 | .[[2]] == 2) & .[[4]] == 1 |
           (.[[2]] == 0 | .[[2]] == 2) & .[[5]] == 1 |
           (.[[2]] == 0 | .[[2]] == 2) & .[[6]] == 1 |
           (.[[2]] == 0 | .[[2]] == 2) & .[[7]] == 1 |
           (.[[3]] == 0 | .[[3]] == 2) & .[[4]] == 1 |
           (.[[3]] == 0 | .[[3]] == 2) & .[[5]] == 1 |
           (.[[3]] == 0 | .[[3]] == 2) & .[[6]] == 1 |
           (.[[3]] == 0 | .[[3]] == 2) & .[[7]] == 1 |
           (.[[4]] == 0 | .[[4]] == 2) & .[[5]] == 1 |
           (.[[4]] == 0 | .[[4]] == 2) & .[[6]] == 1 |
           (.[[4]] == 0 | .[[4]] == 2) & .[[7]] == 1 |
           (.[[5]] == 0 | .[[5]] == 2) & .[[6]] == 1 |
           (.[[5]] == 0 | .[[5]] == 2) & .[[7]] == 1 |
           (.[[6]] == 0 | .[[6]] == 2) & .[[7]] == 1) %>% 
  column_to_rownames()

Line_23_2_corrected <- new_23_2_cor %>%
  rownames_to_column() %>% 
  filter(!(row.names(new_23_2_cor) %in% row.names(test_23_2))) %>% 
  column_to_rownames()

Line_23_4 <- df_gmat_Line_23 %>% 
  rownames_to_column() %>%
  select(1, 8:12) %>% 
  # filter(((`461-P` == 1 & `5-P` == 1) | (`461-P` == 1 & `5-P` == 0) | (`461-P` == 1 & `5-P` == 2) | 
  #           (`461-P` == 0 & `5-P` == 1) | (`461-P` == 0 & `5-P` == 2) | (`461-P` == 2 & `5-P` == 0) | (`461-P` == 2 & `5-P` == 1)) & 
  #          `23_4-F` == 1 & `2344-S1` == 1 & `2344-4-S2` == 1 & `2344-46-S3` == 1 & `2344-464-S4`) %>% 
  column_to_rownames()

# Correcting Line 23_4
new_23_4 <- zygosity_correction_fun_F_S4(Line_23_4)

new_23_4_cor <- new_23_4 %>%
  rownames_to_column() %>% 
  filter(!(`23_4-F` == "remove")) %>% 
  column_to_rownames()

new_23_4_cor[1:5] <- lapply(new_23_4_cor[1:5], as.numeric)

test_23_4 <- new_23_4_cor %>% 
  rownames_to_column() %>% 
  filter((.[[2]] == 0 | .[[2]] == 2) & .[[3]] == 1 |
           (.[[2]] == 0 | .[[2]] == 2) & .[[4]] == 1 |
           (.[[2]] == 0 | .[[2]] == 2) & .[[5]] == 1 |
           (.[[2]] == 0 | .[[2]] == 2) & .[[6]] == 1 |
           (.[[3]] == 0 | .[[3]] == 2) & .[[4]] == 1 |
           (.[[3]] == 0 | .[[3]] == 2) & .[[5]] == 1 |
           (.[[3]] == 0 | .[[3]] == 2) & .[[6]] == 1 |
           (.[[4]] == 0 | .[[4]] == 2) & .[[5]] == 1 |
           (.[[4]] == 0 | .[[4]] == 2) & .[[6]] == 1 |
           (.[[5]] == 0 | .[[5]] == 2) & .[[6]] == 1) %>%  
  column_to_rownames()

Line_23_4_corrected <- new_23_4_cor %>%
  rownames_to_column() %>% 
  filter(!(row.names(new_23_4_cor) %in% row.names(test_23_4))) %>% 
  column_to_rownames()


# how many heterozygotes
table(Line_23_2$`23_2-F`)

table(Line_23_2$`2323-S1`)

table(Line_23_2$`2323-2-S2`)

table(Line_23_2$`2323-21-S3`)

table(Line_23_2$`2323-211-S4`)

table(Line_23_2$`2323-211-S5`)

table(Line_23_4$`23_4-F`)

table(Line_23_4$`2344-S1`)

table(Line_23_4$`2344-4-S2`)

table(Line_23_4$`2344-46-S3`)

table(Line_23_4$`2344-464-S4`)


inbred_heterozygotes_23_2 <- df_gmat_Line_23 %>% 
  rownames_to_column() %>%
  select(1, 4:9) %>% 
  filter((`23_2-F` == 0 | `23_2-F` == 2) & `2323-S1` == 1 |
           (`23_2-F` == 0 | `23_2-F` == 2) & `2323-2-S2` == 1 |
           (`23_2-F` == 0 | `23_2-F` == 2) & `2323-21-S3` == 1 |
           (`23_2-F` == 0 | `23_2-F` == 2) & `2323-211-S4` == 1 |
           (`23_2-F` == 0 | `23_2-F` == 2) & `2323-211-S5` == 1 |
           (`2323-S1` == 0 | `2323-S1` == 2) & `2323-2-S2` == 1 |
           (`2323-S1` == 0 | `2323-S1` == 2) & `2323-21-S3` == 1 |
           (`2323-S1` == 0 | `2323-S1` == 2) & `2323-211-S4` == 1 |
           (`2323-S1` == 0 | `2323-S1` == 2) & `2323-211-S5` == 1 |
           (`2323-2-S2` == 0 | `2323-2-S2` == 2) & `2323-21-S3` == 1 |
           (`2323-2-S2` == 0 | `2323-2-S2` == 2) & `2323-211-S4` == 1 |
           (`2323-2-S2` == 0 | `2323-2-S2` == 2) & `2323-211-S5` == 1 |
           (`2323-21-S3` == 0 | `2323-21-S3` == 2) & `2323-211-S4` == 1 |
           (`2323-21-S3` == 0 | `2323-21-S3` == 2) & `2323-211-S5` == 1 |
           (`2323-211-S4` == 0 | `2323-211-S4` == 2) & `2323-211-S5` == 1) %>% 
  column_to_rownames()

# pdf("Line_23_2_homozygotes_becoming_heterozygotes_heatmap.pdf", onefile = T, width = 15, height = 10)
pheatmap(inbred_heterozygotes_23_2, cluster_rows = F, cluster_cols = F, color = my_col, show_rownames = F)
dev.off()

# write.csv(Line_23_2, "snp_zygosity_analysis/Line_23_2_heterozygote_persistence.txt")
# write.csv(Line_23_4, "snp_zygosity_analysis/Line_23_4_heterozygote_persistence.txt")
# 
# write.csv(rownames(Line_23_2), "snp_zygosity_analysis/Line_23_2_heterozygote_persistence_parents_any_zygosity_snps.txt", row.names = F, col.names = F)
# write.csv(rownames(Line_23_4), "snp_zygosity_analysis/Line_23_4_heterozygote_persistence_parents_any_zygosity_snps.txt", row.names = F, col.names = F)

#Line 26##################################################################################################################

# Read in Line 26 012 file
alleles <- read.csv("Line_26_samples.012",
                    header = F, sep = "\t", na.strings = -1)
str(alleles)
alleles <- alleles[,-1]

# Read in Line 26 SNP ids
snps <- read.csv("Line_26_samples.012.pos",
                 header = F, sep  = "\t")
snps$V1 <- as.character(snps$V1)
snps$V2 <- as.character(snps$V2)

# Merge columns and transpose
snps <- snps %>% 
  unite(col = snp, V1:V2, sep = '_')
t_snps <- t(snps)

# Read Line 26 sample ids
indivs <- read.csv("Line_26_samples.012.indv", header = F, sep = "\t")

# Put row and column names into 012 file
row.names(alleles) <- indivs$V1
colnames(alleles) <- t_snps

# Convert to matrix
g.mat_Line_26 <- as.matrix(alleles)

t_g.mat_Line_26 <- t(g.mat_Line_26)
t_g.mat_Line_26 <- cbind(t_g.mat_Line_26, `2612-12-S3` = rep(NA, 51638))

col.order <- c("26_4-F", "2645-S1", "2645-4-S2", "2645-46-S3", "2645-464-S4",
               "26_1-F", "2612-S1", "2612-1-S2", "2612-12-S3",  "2612-121-S4", 
               "26_2-F", "2621-S1", "2622-121-S4")
t_g.mat_Line_26 <- t_g.mat_Line_26[,col.order]

my_col <- brewer.pal(5, "Spectral")
pheatmap(t_g.mat_Line_26[1:200,], cluster_rows = F, cluster_cols = F, color = my_col)

df_gmat_Line_26 <- data.frame(t_g.mat_Line_26, check.names = F)

# Filtering Line 26_1
Line_26_1 <- df_gmat_Line_26 %>% 
  rownames_to_column() %>%
  select(1, 7:11) %>% 
  # filter(((`461-P` == 1 & `5-P` == 1) | (`461-P` == 1 & `5-P` == 0) | (`461-P` == 1 & `5-P` == 2) | 
  #           (`461-P` == 0 & `5-P` == 1) | (`461-P` == 0 & `5-P` == 2) | (`461-P` == 2 & `5-P` == 0) | (`461-P` == 2 & `5-P` == 1)) & 
  #          `23_4-F` == 1 & `2344-S1` == 1 & `2344-4-S2` == 1 & `2344-46-S3` == 1 & `2344-464-S4`) %>% 
  column_to_rownames()

# Correcting Line 26_1
new_26_1 <- zygosity_correction_fun_F_S4(Line_26_1)

new_26_1_cor <- new_26_1 %>%
  rownames_to_column() %>% 
  filter(!(`26_1-F` == "remove")) %>% 
  column_to_rownames()

new_26_1_cor[1:5] <- lapply(new_26_1_cor[1:5], as.numeric)

test_26_1 <- new_26_1_cor %>% 
  rownames_to_column() %>% 
  filter((.[[2]] == 0 | .[[2]] == 2) & .[[3]] == 1 |
           (.[[2]] == 0 | .[[2]] == 2) & .[[4]] == 1 |
           (.[[2]] == 0 | .[[2]] == 2) & .[[5]] == 1 |
           (.[[2]] == 0 | .[[2]] == 2) & .[[6]] == 1 |
           (.[[3]] == 0 | .[[3]] == 2) & .[[4]] == 1 |
           (.[[3]] == 0 | .[[3]] == 2) & .[[5]] == 1 |
           (.[[3]] == 0 | .[[3]] == 2) & .[[6]] == 1 |
           (.[[4]] == 0 | .[[4]] == 2) & .[[5]] == 1 |
           (.[[4]] == 0 | .[[4]] == 2) & .[[6]] == 1 |
           (.[[5]] == 0 | .[[5]] == 2) & .[[6]] == 1) %>%  
  column_to_rownames()

Line_26_1_corrected <- new_26_1_cor %>%
  rownames_to_column() %>% 
  filter(!(row.names(new_26_1_cor) %in% row.names(test_26_1))) %>% 
  column_to_rownames()


# Filtering Line 26_4
Line_26_4 <- df_gmat_Line_26 %>% 
  rownames_to_column() %>%
  select(1:6) %>% 
  # filter(((`461-P` == 1 & `5-P` == 1) | (`461-P` == 1 & `5-P` == 0) | (`461-P` == 1 & `5-P` == 2) | 
  #           (`461-P` == 0 & `5-P` == 1) | (`461-P` == 0 & `5-P` == 2) | (`461-P` == 2 & `5-P` == 0) | (`461-P` == 2 & `5-P` == 1)) & 
  #          `23_4-F` == 1 & `2344-S1` == 1 & `2344-4-S2` == 1 & `2344-46-S3` == 1 & `2344-464-S4`) %>% 
  column_to_rownames()

# Correcting Line 26_4
new_26_4 <- zygosity_correction_fun_F_S4(Line_26_4)

new_26_4_cor <- new_26_4 %>%
  rownames_to_column() %>% 
  filter(!(`26_4-F` == "remove")) %>% 
  column_to_rownames()

new_26_4_cor[1:5] <- lapply(new_26_4_cor[1:5], as.numeric)

test_26_4 <- new_26_4_cor %>% 
  rownames_to_column() %>% 
  filter((.[[2]] == 0 | .[[2]] == 2) & .[[3]] == 1 |
           (.[[2]] == 0 | .[[2]] == 2) & .[[4]] == 1 |
           (.[[2]] == 0 | .[[2]] == 2) & .[[5]] == 1 |
           (.[[2]] == 0 | .[[2]] == 2) & .[[6]] == 1 |
           (.[[3]] == 0 | .[[3]] == 2) & .[[4]] == 1 |
           (.[[3]] == 0 | .[[3]] == 2) & .[[5]] == 1 |
           (.[[3]] == 0 | .[[3]] == 2) & .[[6]] == 1 |
           (.[[4]] == 0 | .[[4]] == 2) & .[[5]] == 1 |
           (.[[4]] == 0 | .[[4]] == 2) & .[[6]] == 1 |
           (.[[5]] == 0 | .[[5]] == 2) & .[[6]] == 1) %>%  
  column_to_rownames()

Line_26_4_corrected <- new_26_4_cor %>%
  rownames_to_column() %>% 
  filter(!(row.names(new_26_4_cor) %in% row.names(test_26_4))) %>% 
  column_to_rownames()


# how many heterozygotes
table(Line_26_1$`26_1-F`)

table(Line_26_1$`2612-S1`)

table(Line_26_1$`2612-1-S2`)

table(Line_26_1$`2612-12-S3`)

table(Line_26_1$`2612-121-S4`)


# pdf("Line_26_heterozygote_persistence_heatmap.pdf", onefile = T, width = 15, height = 10)
pheatmap(inbred_heterozygotes, cluster_rows = F, cluster_cols = F, color = my_col, show_rownames = F)
dev.off()

#Line 27##################################################################################################################

# Read in Line 27 012 file
alleles <- read.csv("Line_27_samples.012",
                    header = F, sep = "\t", na.strings = -1)
str(alleles)
alleles <- alleles[,-1]

# Read in Line 27 SNP ids
snps <- read.csv("Line_27_samples.012.pos",
                 header = F, sep  = "\t")
snps$V1 <- as.character(snps$V1)
snps$V2 <- as.character(snps$V2)

# Merge columns and transpose
snps <- snps %>% 
  unite(col = snp, V1:V2, sep = '_')
t_snps <- t(snps)

# Read Line 27 sample ids
indivs <- read.csv("Line_27_samples.012.indv", header = F, sep = "\t")

# Put row and column names into 012 file
row.names(alleles) <- indivs$V1
colnames(alleles) <- t_snps

# Convert to matrix
g.mat_Line_27 <- as.matrix(alleles)

t_g.mat_Line_27 <- t(g.mat_Line_27)
t_g.mat_Line_27 <- cbind(t_g.mat_Line_27, `2721-12-S3` = rep(NA, 51638))

col.order <- c("27_1-F", "2712-S1", "2712-211-S4", 
               "27_2-F", "2721-S1", "2721-1-S2", "2721-12-S3", "2721-121-S4",
               "27_6-F", "2765-554-S4")
t_g.mat_Line_27 <- t_g.mat_Line_27[,col.order]

my_col <- brewer.pal(5, "Spectral")
pheatmap(t_g.mat_Line_27[1:200,], cluster_rows = F, cluster_cols = F, color = my_col)

df_gmat_Line_27 <- data.frame(t_g.mat_Line_27, check.names = F)

# Filtering Line 27_2
Line_27_2 <- df_gmat_Line_27 %>% 
  rownames_to_column() %>%
  select(1, 5:9) %>% 
  # filter(((`461-P` == 1 & `5-P` == 1) | (`461-P` == 1 & `5-P` == 0) | (`461-P` == 1 & `5-P` == 2) | 
  #           (`461-P` == 0 & `5-P` == 1) | (`461-P` == 0 & `5-P` == 2) | (`461-P` == 2 & `5-P` == 0) | (`461-P` == 2 & `5-P` == 1)) & 
  #          `23_4-F` == 1 & `2344-S1` == 1 & `2344-4-S2` == 1 & `2344-46-S3` == 1 & `2344-464-S4`) %>% 
  column_to_rownames()

# Correcting Line 27_2
new_27_2 <- zygosity_correction_fun_F_S4(Line_27_2)

new_27_2_cor <- new_27_2 %>%
  rownames_to_column() %>% 
  filter(!(`27_2-F` == "remove")) %>% 
  column_to_rownames()

new_27_2_cor[1:5] <- lapply(new_27_2_cor[1:5], as.numeric)

test_27_2 <- new_27_2_cor %>% 
  rownames_to_column() %>% 
  filter((.[[2]] == 0 | .[[2]] == 2) & .[[3]] == 1 |
           (.[[2]] == 0 | .[[2]] == 2) & .[[4]] == 1 |
           (.[[2]] == 0 | .[[2]] == 2) & .[[5]] == 1 |
           (.[[2]] == 0 | .[[2]] == 2) & .[[6]] == 1 |
           (.[[3]] == 0 | .[[3]] == 2) & .[[4]] == 1 |
           (.[[3]] == 0 | .[[3]] == 2) & .[[5]] == 1 |
           (.[[3]] == 0 | .[[3]] == 2) & .[[6]] == 1 |
           (.[[4]] == 0 | .[[4]] == 2) & .[[5]] == 1 |
           (.[[4]] == 0 | .[[4]] == 2) & .[[6]] == 1 |
           (.[[5]] == 0 | .[[5]] == 2) & .[[6]] == 1) %>%  
  column_to_rownames()

Line_27_2_corrected <- new_27_2_cor %>%
  rownames_to_column() %>% 
  filter(!(row.names(new_27_2_cor) %in% row.names(test_27_2))) %>% 
  column_to_rownames()


# how many heterozygotes
table(Line_27_6$`27_6-F`)

table(Line_27_6$`2765-S1`)

table(Line_27_6$`2765-5-S2`)

table(Line_27_6$`2765-55-S3`)

table(Line_27_6$`2765-554-S4`)

inbred_heterozygotes <- df_gmat_Line_27 %>% 
  rownames_to_column() %>%
  #select(1:7) %>% 
  filter(`1_1-F` == 1, `111-221-S5` == 1, `1_2-F` == 1, `122-122-S4` == 1, `1_4-F` == 1, `145-545-S5` == 1) %>% 
  #filter_all(any_vars (. == 1)) %>% 
  column_to_rownames()

# pdf("Line_27_heterozygote_persistence_heatmap.pdf", onefile = T, width = 15, height = 10)
pheatmap(inbred_heterozygotes, cluster_rows = F, cluster_cols = F, color = my_col, show_rownames = F)
dev.off()

#Line 29##################################################################################################################

# Read in Line 29 012 file
alleles <- read.csv("Line_29_samples.012",
                    header = F, sep = "\t", na.strings = -1)
str(alleles)
alleles <- alleles[,-1]

# Read in Line 29 SNP ids
snps <- read.csv("Line_29_samples.012.pos",
                 header = F, sep  = "\t")
snps$V1 <- as.character(snps$V1)
snps$V2 <- as.character(snps$V2)

# Merge columns and transpose
snps <- snps %>% 
  unite(col = snp, V1:V2, sep = '_')
t_snps <- t(snps)

# Read Line 29 sample ids
indivs <- read.csv("Line_29_samples.012.indv", header = F, sep = "\t")

# Put row and column names into 012 file
row.names(alleles) <- indivs$V1
colnames(alleles) <- t_snps

# Convert to matrix
g.mat_Line_29 <- as.matrix(alleles)

t_g.mat_Line_29 <- t(g.mat_Line_29)
col.order <- c("29_2-F", "2923-S1", "2923-1-S2","2923-12-S3", "2923-121-S4", "2923-121-S5",
               "29_4-F", "2944-S1", "2944-5-S2", "2944-56-S3", "2944-565-S4", "2944-565-S5")
t_g.mat_Line_29 <- t_g.mat_Line_29[,col.order]

my_col <- brewer.pal(5, "Spectral")
pheatmap(t_g.mat_Line_29[1:200,], cluster_rows = F, cluster_cols = F, color = my_col)

df_gmat_Line_29 <- data.frame(t_g.mat_Line_29, check.names = F)

# Filtering Line 29_2
Line_29_2 <- df_gmat_Line_29 %>% 
  rownames_to_column() %>%
  select(1:7) %>% 
  # filter(((`461-P` == 1 & `5-P` == 1) | (`461-P` == 1 & `5-P` == 0) | (`461-P` == 1 & `5-P` == 2) | 
  #           (`461-P` == 0 & `5-P` == 1) | (`461-P` == 0 & `5-P` == 2) | (`461-P` == 2 & `5-P` == 0) | (`461-P` == 2 & `5-P` == 1)) & 
  #          `23_4-F` == 1 & `2344-S1` == 1 & `2344-4-S2` == 1 & `2344-46-S3` == 1 & `2344-464-S4`) %>% 
  column_to_rownames()

# Correcting Line 29_2
new_29_2 <- zygosity_correction_fun_F_S5(Line_29_2)

new_29_2_cor <- new_29_2 %>%
  rownames_to_column() %>% 
  filter(!(`29_2-F` == "remove")) %>% 
  column_to_rownames()

new_29_2_cor[1:6] <- lapply(new_29_2_cor[1:6], as.numeric)

test_29_2 <- new_29_2_cor %>% 
  rownames_to_column() %>% 
  filter((.[[2]] == 0 | .[[2]] == 2) & .[[3]] == 1 |
           (.[[2]] == 0 | .[[2]] == 2) & .[[4]] == 1 |
           (.[[2]] == 0 | .[[2]] == 2) & .[[5]] == 1 |
           (.[[2]] == 0 | .[[2]] == 2) & .[[6]] == 1 |
           (.[[2]] == 0 | .[[2]] == 2) & .[[7]] == 1 |
           (.[[3]] == 0 | .[[3]] == 2) & .[[4]] == 1 |
           (.[[3]] == 0 | .[[3]] == 2) & .[[5]] == 1 |
           (.[[3]] == 0 | .[[3]] == 2) & .[[6]] == 1 |
           (.[[3]] == 0 | .[[3]] == 2) & .[[7]] == 1 |
           (.[[4]] == 0 | .[[4]] == 2) & .[[5]] == 1 |
           (.[[4]] == 0 | .[[4]] == 2) & .[[6]] == 1 |
           (.[[4]] == 0 | .[[4]] == 2) & .[[7]] == 1 |
           (.[[5]] == 0 | .[[5]] == 2) & .[[6]] == 1 |
           (.[[5]] == 0 | .[[5]] == 2) & .[[7]] == 1 |
           (.[[6]] == 0 | .[[6]] == 2) & .[[7]] == 1) %>% 
  column_to_rownames()

Line_29_2_corrected <- new_29_2_cor %>%
  rownames_to_column() %>% 
  filter(!(row.names(new_29_2_cor) %in% row.names(test_29_2))) %>% 
  column_to_rownames()


# Filtering Line 29_4
Line_29_4 <- df_gmat_Line_29 %>% 
  rownames_to_column() %>%
  select(1, 8:13) %>% 
  # filter(((`461-P` == 1 & `5-P` == 1) | (`461-P` == 1 & `5-P` == 0) | (`461-P` == 1 & `5-P` == 2) | 
  #           (`461-P` == 0 & `5-P` == 1) | (`461-P` == 0 & `5-P` == 2) | (`461-P` == 2 & `5-P` == 0) | (`461-P` == 2 & `5-P` == 1)) & 
  #          `23_4-F` == 1 & `2344-S1` == 1 & `2344-4-S2` == 1 & `2344-46-S3` == 1 & `2344-464-S4`) %>% 
  column_to_rownames()

# Correcting Line 29_4
new_29_4 <- zygosity_correction_fun_F_S5(Line_29_4)

new_29_4_cor <- new_29_4 %>%
  rownames_to_column() %>% 
  filter(!(`29_4-F` == "remove")) %>% 
  column_to_rownames()

new_29_4_cor[1:6] <- lapply(new_29_4_cor[1:6], as.numeric)

test_29_4 <- new_29_4_cor %>% 
  rownames_to_column() %>% 
  filter((.[[2]] == 0 | .[[2]] == 2) & .[[3]] == 1 |
           (.[[2]] == 0 | .[[2]] == 2) & .[[4]] == 1 |
           (.[[2]] == 0 | .[[2]] == 2) & .[[5]] == 1 |
           (.[[2]] == 0 | .[[2]] == 2) & .[[6]] == 1 |
           (.[[2]] == 0 | .[[2]] == 2) & .[[7]] == 1 |
           (.[[3]] == 0 | .[[3]] == 2) & .[[4]] == 1 |
           (.[[3]] == 0 | .[[3]] == 2) & .[[5]] == 1 |
           (.[[3]] == 0 | .[[3]] == 2) & .[[6]] == 1 |
           (.[[3]] == 0 | .[[3]] == 2) & .[[7]] == 1 |
           (.[[4]] == 0 | .[[4]] == 2) & .[[5]] == 1 |
           (.[[4]] == 0 | .[[4]] == 2) & .[[6]] == 1 |
           (.[[4]] == 0 | .[[4]] == 2) & .[[7]] == 1 |
           (.[[5]] == 0 | .[[5]] == 2) & .[[6]] == 1 |
           (.[[5]] == 0 | .[[5]] == 2) & .[[7]] == 1 |
           (.[[6]] == 0 | .[[6]] == 2) & .[[7]] == 1) %>% 
  column_to_rownames()

Line_29_4_corrected <- new_29_4_cor %>%
  rownames_to_column() %>% 
  filter(!(row.names(new_29_4_cor) %in% row.names(test_29_4))) %>% 
  column_to_rownames()


# how many heterozygotes
table(Line_29_2_corrected$`29_2-F`)

table(Line_29_2_corrected$`2923-S1`)

table(Line_29_2_corrected$`2923-1-S2`)

table(Line_29_2_corrected$`2923-12-S3`)

table(Line_29_2_corrected$`2923-121-S4`)

table(Line_29_2_corrected$`2923-211-S5`)

table(Line_29_4_corrected$`29_4-F`)

table(Line_29_4_corrected$`2944-S1`)

table(Line_29_4_corrected$`2944-5-S2`)

table(Line_29_4_corrected$`2944-56-S3`)

table(Line_29_4_corrected$`2944-565-S4`)

table(Line_29_4_corrected$`2944-565-S5`)

inbred_heterozygotes <- df_gmat_Line_29 %>% 
  rownames_to_column() %>%
  #select(1:7) %>% 
  filter(`1_1-F` == 1, `111-221-S5` == 1, `1_2-F` == 1, `122-122-S4` == 1, `1_4-F` == 1, `145-545-S5` == 1) %>% 
  #filter_all(any_vars (. == 1)) %>% 
  column_to_rownames()

# pdf("Line_29_heterozygote_persistence_heatmap.pdf", onefile = T, width = 15, height = 10)
pheatmap(inbred_heterozygotes, cluster_rows = F, cluster_cols = F, color = my_col, show_rownames = F)
dev.off()

### Write files #####
# write.csv(Line_1_1_corrected, "Line_1_1_corrected.csv")
# write.csv(Line_6_1_corrected, "Line_6_1_corrected.csv")
# write.csv(Line_6_4_corrected, "Line_6_4_corrected.csv")
# write.csv(Line_7_2_corrected, "Line_7_2_corrected.csv")
# write.csv(Line_7_4_corrected, "Line_7_4_corrected.csv")
# write.csv(Line_8_2_corrected, "Line_8_2_corrected.csv")
# write.csv(Line_8_4_corrected, "Line_8_4_corrected.csv")
# write.csv(Line_10_2_corrected, "Line_10_2_corrected.csv")
# write.csv(Line_10_5_corrected, "Line_10_5_corrected.csv")
# write.csv(Line_13_3_corrected, "Line_13_3_corrected.csv")
# write.csv(Line_13_4_corrected, "Line_13_4_corrected.csv")
# write.csv(Line_16_1_corrected, "Line_16_1_corrected.csv")
# write.csv(Line_16_5_corrected, "Line_16_5_corrected.csv")
# write.csv(Line_17_2_corrected, "Line_17_2_corrected.csv")
# write.csv(Line_17_5_corrected, "Line_17_5_corrected.csv")
# write.csv(Line_19_2_corrected, "Line_19_2_corrected.csv")
# write.csv(Line_19_5_corrected, "Line_19_5_corrected.csv")
# write.csv(Line_20_1_corrected, "Line_20_1_corrected.csv")
# write.csv(Line_20_4_corrected, "Line_20_4_corrected.csv")
# write.csv(Line_21_6_corrected, "Line_21_6_corrected.csv")
# write.csv(Line_21_2_corrected, "Line_21_2_corrected.csv")
# write.csv(Line_23_2_corrected, "Line_23_2_corrected.csv")
# write.csv(Line_23_4_corrected, "Line_23_4_corrected.csv")
# write.csv(Line_26_1_corrected, "Line_26_1_corrected.csv")
# write.csv(Line_26_4_corrected, "Line_26_4_corrected.csv")
# write.csv(Line_27_2_corrected, "Line_27_2_corrected.csv")
# write.csv(Line_29_2_corrected, "Line_29_2_corrected.csv")
# write.csv(Line_29_4_corrected, "Line_29_4_corrected.csv")
```
