## Supplementary material for "The western redcedar genome reveals low genetic diversity in a self-compatible conifer": Code S6: S_line_pop_gen_and_heterozygosity.R

```
rm(list=ls())

library(adegenet)
library(sommer)
library(reshape2)
library(RColorBrewer)
library(pheatmap)
library(hierfstat)
library(ggsci)
library(hierfstat)
library(vcfR)
library(pegas)
library(ggsci)
library(apex)
library(ggpubr)
library(mmod)
library(factoextra)
library(FactoMineR)
library(ggpubr)
library(rstatix)
library(tidyverse)

source("publication_theme.r")

### Loading manually corrected data and chi-squared analysis ###########################

all_snp_ids <- read.table("all_filtered_snp_ids.txt")


#Line 1##################################################################################################################

Line_1_1_corrected <- read.csv("Line_1_1_corrected_manually.csv", check.names = F, row.names = 1)

# Line_1_4_corrected <- read.csv("Line_1_4_corrected_manually.csv", check.names = F, row.names = 1)

Line_1_1_het_F <- Line_1_1_corrected %>% rownames_to_column() %>% 
  filter(`1_1-F` == 1) %>% filter(rowname %in% all_snp_ids$V1) %>% column_to_rownames()

# Line_1_4_het_F <- Line_1_4_corrected %>% rownames_to_column() %>% 
#   filter(`1_4-F` == 1) %>% filter(rowname %in% all_snp_ids$V1) %>% column_to_rownames()

Line_1_1_het_S4 <- Line_1_1_corrected %>% rownames_to_column() %>% 
  filter(`111-221-S4` == 1) %>% filter(rowname %in% all_snp_ids$V1) %>% column_to_rownames()

# Line_1_1_corrected <- Line_1_1_corrected %>% 
#   rownames_to_column() %>% 
#   filter(rowname %in% all_snp_ids$V1) %>% column_to_rownames()

##Line 6#################################################################################################################
Line_6_1_corrected <- read.csv("Line_6_1_corrected_manually.csv", check.names = F, row.names = 1)
Line_6_4_corrected <- read.csv("Line_6_4_corrected_manually.csv", check.names = F, row.names = 1)

Line_6_1_het_F <- Line_6_1_corrected %>% rownames_to_column() %>% 
  filter(`6_1-F` == 1) %>% filter(rowname %in% all_snp_ids$V1) %>% column_to_rownames()

Line_6_4_het_F <- Line_6_4_corrected %>% rownames_to_column() %>% 
  filter(`6_4-F` == 1) %>% filter(rowname %in% all_snp_ids$V1) %>% column_to_rownames()

Line_6_1_het_S4 <- Line_6_1_corrected %>% rownames_to_column() %>% 
  filter(`611-111-S4` == 1) %>% filter(rowname %in% all_snp_ids$V1) %>% column_to_rownames()

Line_6_4_het_S4 <- Line_6_4_corrected %>% rownames_to_column() %>% 
  filter(`646-454-S4` == 1) %>% filter(rowname %in% all_snp_ids$V1) %>% column_to_rownames()

# Line_6_1_corrected <- Line_6_1_corrected %>% 
#   rownames_to_column() %>% 
#   filter(rowname %in% all_snp_ids$V1) %>% column_to_rownames()
# 
# Line_6_4_corrected <- Line_6_4_corrected %>% 
#   rownames_to_column() %>% 
#   filter(rowname %in% all_snp_ids$V1) %>% column_to_rownames()

#Line 7##################################################################################################################

Line_7_2_corrected <- read.csv("Line_7_2_corrected_manually.csv", check.names = F, row.names = 1)
Line_7_4_corrected <- read.csv("Line_7_4_corrected_manually.csv", check.names = F, row.names = 1)

Line_7_2_het_F <- Line_7_2_corrected %>% rownames_to_column() %>% 
  filter(`7_2-F` == 1) %>% filter(rowname %in% all_snp_ids$V1) %>% column_to_rownames()

Line_7_4_het_F <- Line_7_4_corrected %>% rownames_to_column() %>% 
  filter(`7_4-F` == 1) %>% filter(rowname %in% all_snp_ids$V1) %>% column_to_rownames()

Line_7_4_het_S4 <- Line_7_4_corrected %>% rownames_to_column() %>% 
  filter(`745-444-S4` == 1) %>% filter(rowname %in% all_snp_ids$V1) %>% column_to_rownames()

# Line_7_2_corrected <- Line_7_2_corrected %>% 
#   rownames_to_column() %>% 
#   filter(rowname %in% all_snp_ids$V1) %>% column_to_rownames()
# 
# Line_7_4_corrected <- Line_7_4_corrected %>% 
#   rownames_to_column() %>% 
#   filter(rowname %in% all_snp_ids$V1) %>% column_to_rownames()

#Line 8##################################################################################################################
Line_8_2_corrected <- read.csv("Line_8_2_corrected_manually.csv", check.names = F, row.names = 1)
Line_8_4_corrected <- read.csv("Line_8_4_corrected_manually.csv", check.names = F, row.names = 1)


Line_8_2_het_F <- Line_8_2_corrected %>% rownames_to_column() %>% 
  filter(`8_2-F` == 1) %>% filter(rowname %in% all_snp_ids$V1) %>% column_to_rownames()

Line_8_4_het_F <- Line_8_4_corrected %>% rownames_to_column() %>% 
  filter(`8_4-F` == 1) %>% filter(rowname %in% all_snp_ids$V1) %>% column_to_rownames()

Line_8_2_het_S4 <- Line_8_2_corrected %>% rownames_to_column() %>% 
  filter(`821-111-S4` == 1) %>% filter(rowname %in% all_snp_ids$V1) %>% column_to_rownames()

Line_8_4_het_S4 <- Line_8_4_corrected %>% rownames_to_column() %>% 
  filter(`845-444-S4` == 1) %>% filter(rowname %in% all_snp_ids$V1) %>% column_to_rownames()

# Line_8_2_corrected <- Line_8_2_corrected %>% 
#   rownames_to_column() %>% 
#   filter(rowname %in% all_snp_ids$V1) %>% column_to_rownames()
# 
# Line_8_4_corrected <- Line_8_4_corrected %>% 
#   rownames_to_column() %>% 
#   filter(rowname %in% all_snp_ids$V1) %>% column_to_rownames()


##Line 10#################################################################################################################

Line_10_2_corrected <- read.csv("Line_10_2_corrected_manually.csv", check.names = F, row.names = 1)
Line_10_5_corrected <- read.csv("Line_10_5_corrected_manually.csv", check.names = F, row.names = 1)

Line_10_2_het_F <- Line_10_2_corrected %>% rownames_to_column() %>% 
  filter(`10_2-F` == 1) %>% filter(rowname %in% all_snp_ids$V1) %>% column_to_rownames()

Line_10_5_het_F <- Line_10_5_corrected %>% rownames_to_column() %>% 
  filter(`10_5-F` == 1) %>% filter(rowname %in% all_snp_ids$V1) %>% column_to_rownames()

Line_10_5_het_S4 <- Line_10_5_corrected %>% rownames_to_column() %>% 
  filter(`1056-454-S4` == 1) %>% filter(rowname %in% all_snp_ids$V1) %>% column_to_rownames()

# Line_10_2_corrected <- Line_10_2_corrected %>% 
#   rownames_to_column() %>% 
#   filter(rowname %in% all_snp_ids$V1) %>% column_to_rownames()
# 
# Line_10_5_corrected <- Line_10_5_corrected %>% 
#   rownames_to_column() %>% 
#   filter(rowname %in% all_snp_ids$V1) %>% column_to_rownames()

#Line 13##################################################################################################################

Line_13_3_corrected <- read.csv("Line_13_3_corrected_manually.csv", check.names = F, row.names = 1)
Line_13_4_corrected <- read.csv("Line_13_4_corrected_manually.csv", check.names = F, row.names = 1)

Line_13_3_het_F <- Line_13_3_corrected %>% rownames_to_column() %>% 
  filter(`13_3-F` == 1) %>% filter(rowname %in% all_snp_ids$V1) %>% column_to_rownames()

Line_13_4_het_F <- Line_13_4_corrected %>% rownames_to_column() %>% 
  filter(`13_4-F` == 1) %>% filter(rowname %in% all_snp_ids$V1) %>% column_to_rownames()

Line_13_4_het_S4 <- Line_13_4_corrected %>% rownames_to_column() %>% 
  filter(`1345-544-S4` == 1) %>% filter(rowname %in% all_snp_ids$V1) %>% column_to_rownames()

# Line_13_3_corrected <- Line_13_3_corrected %>% 
#   rownames_to_column() %>% 
#   filter(rowname %in% all_snp_ids$V1) %>% column_to_rownames()
# 
# Line_13_4_corrected <- Line_13_4_corrected %>% 
#   rownames_to_column() %>% 
#   filter(rowname %in% all_snp_ids$V1) %>% column_to_rownames()

#Line 16##################################################################################################################

Line_16_1_corrected <- read.csv("Line_16_1_corrected_manually.csv", check.names = F, row.names = 1)
Line_16_5_corrected <- read.csv("Line_16_5_corrected_manually.csv", check.names = F, row.names = 1)

Line_16_1_het_F <- Line_16_1_corrected %>% rownames_to_column() %>% 
  filter(`16_1-F` == 1) %>% filter(rowname %in% all_snp_ids$V1) %>% column_to_rownames()

Line_16_5_het_F <- Line_16_5_corrected %>% rownames_to_column() %>% 
  filter(`16_5-F` == 1) %>% filter(rowname %in% all_snp_ids$V1) %>% column_to_rownames()

Line_16_1_het_S4 <- Line_16_1_corrected %>% rownames_to_column() %>% 
  filter(`1611-211-S4` == 1) %>% filter(rowname %in% all_snp_ids$V1) %>% column_to_rownames()

# Line_16_1_corrected <- Line_16_1_corrected %>% 
#   rownames_to_column() %>% 
#   filter(rowname %in% all_snp_ids$V1) %>% column_to_rownames()
# 
# Line_16_5_corrected <- Line_16_5_corrected %>% 
#   rownames_to_column() %>% 
#   filter(rowname %in% all_snp_ids$V1) %>% column_to_rownames()

#Line 17##################################################################################################################

Line_17_2_corrected <- read.csv("Line_17_2_corrected_manually.csv", check.names = F, row.names = 1)
Line_17_5_corrected <- read.csv("Line_17_5_corrected_manually.csv", check.names = F, row.names = 1)

Line_17_2_het_F <- Line_17_2_corrected %>% rownames_to_column() %>% 
  filter(`17_2-F` == 1) %>% filter(rowname %in% all_snp_ids$V1) %>% column_to_rownames()

Line_17_5_het_F <- Line_17_5_corrected %>% rownames_to_column() %>% 
  filter(`17_5-F` == 1) %>% filter(rowname %in% all_snp_ids$V1) %>% column_to_rownames()

Line_17_2_het_S4 <- Line_17_2_corrected %>% rownames_to_column() %>% 
  filter(`1721-111-S4` == 1) %>% filter(rowname %in% all_snp_ids$V1) %>% column_to_rownames()

Line_17_5_het_S4 <- Line_17_5_corrected %>% rownames_to_column() %>% 
  filter(`1755-545-S4` == 1) %>% filter(rowname %in% all_snp_ids$V1) %>% column_to_rownames()

# Line_17_2_corrected <- Line_17_2_corrected %>% 
#   rownames_to_column() %>% 
#   filter(rowname %in% all_snp_ids$V1) %>% column_to_rownames()
# 
# Line_17_5_corrected <- Line_17_5_corrected %>% 
#   rownames_to_column() %>% 
#   filter(rowname %in% all_snp_ids$V1) %>% column_to_rownames()

#Line 19##################################################################################################################

Line_19_2_corrected <- read.csv("Line_19_2_corrected_manually.csv", check.names = F, row.names = 1)
Line_19_5_corrected <- read.csv("Line_19_5_corrected_manually.csv", check.names = F, row.names = 1)

Line_19_2_het_F <- Line_19_2_corrected %>% rownames_to_column() %>% 
  filter(`19_2-F` == 1) %>% filter(rowname %in% all_snp_ids$V1) %>% column_to_rownames()

Line_19_5_het_F <- Line_19_5_corrected %>% rownames_to_column() %>% 
  filter(`19_5-F` == 1) %>% filter(rowname %in% all_snp_ids$V1) %>% column_to_rownames()

Line_19_5_het_S4 <- Line_19_5_corrected %>% rownames_to_column() %>% 
  filter(`1955-544-S4` == 1) %>% filter(rowname %in% all_snp_ids$V1) %>% column_to_rownames()

# Line_19_2_corrected <- Line_19_2_corrected %>% 
#   rownames_to_column() %>% 
#   filter(rowname %in% all_snp_ids$V1) %>% column_to_rownames()
# 
# Line_19_5_corrected <- Line_19_5_corrected %>% 
#   rownames_to_column() %>% 
#   filter(rowname %in% all_snp_ids$V1) %>% column_to_rownames()

#Line 20##################################################################################################################

Line_20_1_corrected <- read.csv("Line_20_1_corrected_manually.csv", check.names = F, row.names = 1)
Line_20_4_corrected <- read.csv("Line_20_4_corrected_manually.csv", check.names = F, row.names = 1)

Line_20_1_het_F <- Line_20_1_corrected %>% rownames_to_column() %>% 
  filter(`20_1-F` == 1) %>% filter(rowname %in% all_snp_ids$V1) %>% column_to_rownames()

Line_20_4_het_F <- Line_20_4_corrected %>% rownames_to_column() %>% 
  filter(`20_4-F` == 1) %>% filter(rowname %in% all_snp_ids$V1) %>% column_to_rownames()

Line_20_1_het_S4 <- Line_20_1_corrected %>% rownames_to_column() %>% 
  filter(`2013-131-S4` == 1) %>% filter(rowname %in% all_snp_ids$V1) %>% column_to_rownames()

Line_20_4_het_S4 <- Line_20_4_corrected %>% rownames_to_column() %>% 
  filter(`2045-544-S4` == 1) %>% filter(rowname %in% all_snp_ids$V1) %>% column_to_rownames()
# 
# Line_20_1_corrected <- Line_20_1_corrected %>% 
#   rownames_to_column() %>% 
#   filter(rowname %in% all_snp_ids$V1) %>% column_to_rownames()
# 
# Line_20_4_corrected <- Line_20_4_corrected %>% 
#   rownames_to_column() %>% 
#   filter(rowname %in% all_snp_ids$V1) %>% column_to_rownames()

#Line 21##################################################################################################################

Line_21_2_corrected <- read.csv("Line_21_2_corrected_manually.csv", check.names = F, row.names = 1)
Line_21_6_corrected <- read.csv("Line_21_6_corrected_manually.csv", check.names = F, row.names = 1)


Line_21_2_het_F <- Line_21_2_corrected %>% rownames_to_column() %>% 
  filter(`21_2-F` == 1) %>% filter(rowname %in% all_snp_ids$V1) %>% column_to_rownames()

Line_21_6_het_F <- Line_21_6_corrected %>% rownames_to_column() %>% 
  filter(`21_6-F` == 1) %>% filter(rowname %in% all_snp_ids$V1) %>% column_to_rownames()

Line_21_2_het_S4 <- Line_21_2_corrected %>% rownames_to_column() %>% 
  filter(`2121-111-S4` == 1) %>% filter(rowname %in% all_snp_ids$V1) %>% column_to_rownames()

Line_21_6_het_S4 <- Line_21_6_corrected %>% rownames_to_column() %>% 
  filter(`2165-444-S4` == 1) %>% filter(rowname %in% all_snp_ids$V1) %>% column_to_rownames()

# Line_21_2_corrected <- Line_21_2_corrected %>% 
#   rownames_to_column() %>% 
#   filter(rowname %in% all_snp_ids$V1) %>% column_to_rownames()
# 
# Line_21_6_corrected <- Line_21_6_corrected %>% 
#   rownames_to_column() %>% 
#   filter(rowname %in% all_snp_ids$V1) %>% column_to_rownames()

##Line 23#################################################################################################################

Line_23_2_corrected <- read.csv("Line_23_2_corrected_manually.csv", check.names = F, row.names = 1)
Line_23_4_corrected <- read.csv("Line_23_4_corrected_manually.csv", check.names = F, row.names = 1)

Line_23_2_het_F <- Line_23_2_corrected %>% rownames_to_column() %>% 
  filter(`23_2-F` == 1) %>% filter(rowname %in% all_snp_ids$V1) %>% column_to_rownames()

Line_23_4_het_F <- Line_23_4_corrected %>% rownames_to_column() %>% 
  filter(`23_4-F` == 1) %>% filter(rowname %in% all_snp_ids$V1) %>% column_to_rownames()

Line_23_2_het_S4 <- Line_23_2_corrected %>% rownames_to_column() %>% 
  filter(`2323-211-S4` == 1) %>% filter(rowname %in% all_snp_ids$V1) %>% column_to_rownames()

Line_23_4_het_S4 <- Line_23_4_corrected %>% rownames_to_column() %>% 
  filter(`2344-464-S4` == 1) %>% filter(rowname %in% all_snp_ids$V1) %>% column_to_rownames()

# Line_23_2_corrected <- Line_23_2_corrected %>% 
#   rownames_to_column() %>% 
#   filter(rowname %in% all_snp_ids$V1) %>% column_to_rownames()
# 
# Line_23_4_corrected <- Line_23_4_corrected %>% 
#   rownames_to_column() %>% 
#   filter(rowname %in% all_snp_ids$V1) %>% column_to_rownames()

#Line 26##################################################################################################################

Line_26_1_corrected <- read.csv("Line_26_1_corrected_manually.csv", check.names = F, row.names = 1)
Line_26_4_corrected <- read.csv("Line_26_4_corrected_manually.csv", check.names = F, row.names = 1)

Line_26_1_het_F <- Line_26_1_corrected %>% rownames_to_column() %>% 
  filter(`26_1-F` == 1) %>% filter(rowname %in% all_snp_ids$V1) %>% column_to_rownames()

Line_26_4_het_F <- Line_26_4_corrected %>% rownames_to_column() %>% 
  filter(`26_4-F` == 1) %>% filter(rowname %in% all_snp_ids$V1) %>% column_to_rownames()

Line_26_1_het_S4 <- Line_26_1_corrected %>% rownames_to_column() %>% 
  filter(`2612-121-S4` == 1) %>% filter(rowname %in% all_snp_ids$V1) %>% column_to_rownames()

# Line_26_1_corrected <- Line_26_1_corrected %>% 
#   rownames_to_column() %>% 
#   filter(rowname %in% all_snp_ids$V1) %>% column_to_rownames()
# 
# Line_26_4_corrected <- Line_26_4_corrected %>% 
#   rownames_to_column() %>% 
#   filter(rowname %in% all_snp_ids$V1) %>% column_to_rownames()

#Line 27##################################################################################################################

Line_27_2_corrected <- read.csv("Line_27_2_corrected_manually.csv", check.names = F, row.names = 1)

Line_27_2_het_F <- Line_27_2_corrected %>% rownames_to_column() %>% 
  filter(`27_2-F` == 1) %>% filter(rowname %in% all_snp_ids$V1) %>% column_to_rownames()

# Line_27_2_corrected <- Line_27_2_corrected %>% 
#   rownames_to_column() %>% 
#   filter(rowname %in% all_snp_ids$V1) %>% column_to_rownames()

#Line 29##################################################################################################################

Line_29_2_corrected <- read.csv("Line_29_2_corrected_manually.csv", check.names = F, row.names = 1)
Line_29_4_corrected <- read.csv("Line_29_4_corrected_manually.csv", check.names = F, row.names = 1)

Line_29_2_het_F <- Line_29_2_corrected %>% rownames_to_column() %>% 
  filter(`29_2-F` == 1) %>% filter(rowname %in% all_snp_ids$V1) %>% column_to_rownames()

Line_29_4_het_F <- Line_29_4_corrected %>% rownames_to_column() %>% 
  filter(`29_4-F` == 1) %>% filter(rowname %in% all_snp_ids$V1) %>% column_to_rownames()

Line_29_2_het_S4 <- Line_29_2_corrected %>% rownames_to_column() %>% 
  filter(`2923-121-S4` == 1) %>% filter(rowname %in% all_snp_ids$V1) %>% column_to_rownames()

Line_29_4_het_S4 <- Line_29_4_corrected %>% rownames_to_column() %>% 
  filter(`2944-565-S4` == 1) %>% filter(rowname %in% all_snp_ids$V1) %>% column_to_rownames()

# Line_29_2_corrected <- Line_29_2_corrected %>% 
#   rownames_to_column() %>% 
#   filter(rowname %in% all_snp_ids$V1) %>% column_to_rownames()
# 
# Line_29_4_corrected <- Line_29_4_corrected %>% 
#   rownames_to_column() %>% 
#   filter(rowname %in% all_snp_ids$V1) %>% column_to_rownames()

# ### Fixation plots #################
# melted_1_1_het_F <- Line_1_1_het_F %>% 
#   rownames_to_column() %>% 
#   melt()
# 
# ggplot(melted_1_1_het_F, aes(x = variable, y = value)) +
#   geom_violin()
# 
# fixation_1_1_plot <- ggplot(melted_1_1_het_F, aes(x = value, fill = variable)) +
#   geom_histogram(binwidth = 0.5) +
#   scale_fill_nejm() +
#   theme_Publication()
# 
# melted_6_1_het_F <- melt(Line_6_1_het_F)
# 
# ggplot(melted_6_1_het_F, aes(x = variable, y = value)) +
#   geom_violin()
# 
# fixation_6_1_plot <- ggplot(melted_6_1_het_F, aes(x = value, fill = variable)) +
#   geom_histogram(binwidth = 0.5) +
#   scale_fill_nejm() +
#   theme_Publication()
# 
# melted_6_4_het_F <- melt(Line_6_4_het_F)
# 
# ggplot(melted_6_4_het_F, aes(x = variable, y = value)) +
#   geom_violin()
# 
# fixation_6_4_plot <- ggplot(melted_6_4_het_F, aes(x = value, fill = variable)) +
#   geom_histogram(binwidth = 0.5) +
#   scale_fill_nejm() +
#   theme_Publication()
# 
# melted_7_2_het_F <- Line_7_2_het_F %>% 
#   rownames_to_column() %>% 
#   melt()
# 
# ggplot(melted_7_2_het_F, aes(x = variable, y = value)) +
#   geom_violin()
# 
# fixation_7_2_plot <- ggplot(melted_7_2_het_F, aes(x = value, fill = variable)) +
#   geom_histogram(binwidth = 0.5) +
#   scale_fill_nejm() +
#   theme_Publication()
# 
# melted_7_4_het_F <- Line_7_4_het_F %>% 
#   rownames_to_column() %>% 
#   melt()
# 
# ggplot(melted_7_4_het_F, aes(x = variable, y = value)) +
#   geom_violin()
# 
# fixation_7_4_plot <- ggplot(melted_7_4_het_F, aes(x = value, fill = variable)) +
#   geom_histogram(binwidth = 0.5) +
#   scale_fill_nejm() +
#   theme_Publication()
# 
# melted_8_2_het_F <- Line_8_2_het_F %>% 
#   rownames_to_column() %>% 
#   melt()
# 
# ggplot(melted_8_2_het_F, aes(x = variable, y = value)) +
#   geom_violin()
# 
# fixation_8_2_plot <- ggplot(melted_8_2_het_F, aes(x = value, fill = variable)) +
#   geom_histogram(binwidth = 0.5) +
#   scale_fill_nejm() +
#   theme_Publication()
# 
# melted_8_4_het_F <- Line_8_4_het_F %>% 
#   rownames_to_column() %>% 
#   melt()
# 
# ggplot(melted_8_4_het_F, aes(x = variable, y = value)) +
#   geom_violin()
# 
# fixation_8_4_plot <- ggplot(melted_8_4_het_F, aes(x = value, fill = variable)) +
#   geom_histogram(binwidth = 0.5) +
#   scale_fill_nejm() +
#   theme_Publication()
# 
# melted_10_2_het_F <- Line_10_2_het_F %>% 
#   rownames_to_column() %>% 
#   melt()
# 
# ggplot(melted_10_2_het_F, aes(x = variable, y = value)) +
#   geom_violin()
# 
# fixation_10_2_plot <- ggplot(melted_10_2_het_F, aes(x = value, fill = variable)) +
#   geom_histogram(binwidth = 0.5) +
#   scale_fill_nejm() +
#   theme_Publication()
# 
# melted_10_5_het_F <- Line_10_5_het_F %>% 
#   rownames_to_column() %>% 
#   melt()
# 
# ggplot(melted_10_5_het_F, aes(x = variable, y = value)) +
#   geom_violin()
# 
# fixation_10_5_plot <- ggplot(melted_10_5_het_F, aes(x = value, fill = variable)) +
#   geom_histogram(binwidth = 0.5) +
#   scale_fill_nejm() +
#   theme_Publication()
# 
# melted_13_3_het_F <- Line_13_3_het_F %>% 
#   rownames_to_column() %>% 
#   melt()
# 
# ggplot(melted_13_3_het_F, aes(x = variable, y = value)) +
#   geom_violin()
# 
# fixation_13_3_plot <- ggplot(melted_13_3_het_F, aes(x = value, fill = variable)) +
#   geom_histogram(binwidth = 0.5) +
#   scale_fill_nejm() +
#   theme_Publication()
# 
# melted_13_4_het_F <- Line_13_4_het_F %>% 
#   rownames_to_column() %>% 
#   melt()
# 
# ggplot(melted_13_4_het_F, aes(x = variable, y = value)) +
#   geom_violin()
# 
# fixation_13_4_plot <- ggplot(melted_13_4_het_F, aes(x = value, fill = variable)) +
#   geom_histogram(binwidth = 0.5) +
#   scale_fill_nejm() +
#   theme_Publication()
# 
# melted_16_1_het_F <- Line_16_1_het_F %>% 
#   rownames_to_column() %>% 
#   melt()
# 
# ggplot(melted_16_1_het_F, aes(x = variable, y = value)) +
#   geom_violin()
# 
# fixation_16_1_plot <- ggplot(melted_16_1_het_F, aes(x = value, fill = variable)) +
#   geom_histogram(binwidth = 0.5) +
#   scale_fill_nejm() +
#   theme_Publication()
# 
# melted_16_5_het_F <- Line_16_5_het_F %>% 
#   rownames_to_column() %>% 
#   melt()
# 
# ggplot(melted_16_5_het_F, aes(x = variable, y = value)) +
#   geom_violin()
# 
# fixation_16_5_plot <- ggplot(melted_16_5_het_F, aes(x = value, fill = variable)) +
#   geom_histogram(binwidth = 0.5) +
#   scale_fill_nejm() +
#   theme_Publication()
# 
# melted_17_2_het_F <- Line_17_2_het_F %>% 
#   rownames_to_column() %>% 
#   melt()
# 
# ggplot(melted_17_2_het_F, aes(x = variable, y = value)) +
#   geom_violin()
# 
# fixation_17_2_plot <- ggplot(melted_17_2_het_F, aes(x = value, fill = variable)) +
#   geom_histogram(binwidth = 0.5) +
#   scale_fill_nejm() +
#   theme_Publication()
# 
# melted_17_5_het_F <- Line_17_5_het_F %>% 
#   rownames_to_column() %>% 
#   melt()
# 
# ggplot(melted_17_5_het_F, aes(x = variable, y = value)) +
#   geom_violin()
# 
# fixation_17_5_plot <- ggplot(melted_17_5_het_F, aes(x = value, fill = variable)) +
#   geom_histogram(binwidth = 0.5) +
#   scale_fill_nejm() +
#   theme_Publication()
# 
# melted_19_2_het_F <- Line_19_2_het_F %>% 
#   rownames_to_column() %>% 
#   melt()
# 
# ggplot(melted_19_2_het_F, aes(x = variable, y = value)) +
#   geom_violin()
# 
# fixation_19_2_plot <- ggplot(melted_19_2_het_F, aes(x = value, fill = variable)) +
#   geom_histogram(binwidth = 0.5) +
#   scale_fill_nejm() +
#   theme_Publication()
# 
# melted_19_5_het_F <- Line_19_5_het_F %>% 
#   rownames_to_column() %>% 
#   melt()
# 
# ggplot(melted_19_5_het_F, aes(x = variable, y = value)) +
#   geom_violin()
# 
# fixation_19_5_plot <- ggplot(melted_19_5_het_F, aes(x = value, fill = variable)) +
#   geom_histogram(binwidth = 0.5) +
#   scale_fill_nejm() +
#   theme_Publication()
# 
# melted_20_1_het_F <- Line_20_1_het_F %>% 
#   rownames_to_column() %>% 
#   melt()
# 
# ggplot(melted_20_1_het_F, aes(x = variable, y = value)) +
#   geom_violin()
# 
# fixation_20_1_plot <- ggplot(melted_20_1_het_F, aes(x = value, fill = variable)) +
#   geom_histogram(binwidth = 0.5) +
#   scale_fill_nejm() +
#   theme_Publication()
# 
# melted_20_4_het_F <- Line_20_4_het_F %>% 
#   rownames_to_column() %>% 
#   melt()
# 
# ggplot(melted_20_4_het_F, aes(x = variable, y = value)) +
#   geom_violin()
# 
# fixation_20_4_plot <- ggplot(melted_20_4_het_F, aes(x = value, fill = variable)) +
#   geom_histogram(binwidth = 0.5) +
#   scale_fill_nejm() +
#   theme_Publication()
# 
# melted_21_2_het_F <- Line_21_2_het_F %>% 
#   rownames_to_column() %>% 
#   melt()
# 
# ggplot(melted_21_2_het_F, aes(x = variable, y = value)) +
#   geom_violin()
# 
# fixation_21_2_plot <- ggplot(melted_21_2_het_F, aes(x = value, fill = variable)) +
#   geom_histogram(binwidth = 0.5) +
#   scale_fill_nejm() +
#   theme_Publication()
# 
# melted_21_6_het_F <- Line_21_6_het_F %>% 
#   rownames_to_column() %>% 
#   melt()
# 
# ggplot(melted_21_6_het_F, aes(x = variable, y = value)) +
#   geom_violin()
# 
# fixation_21_6_plot <- ggplot(melted_21_6_het_F, aes(x = value, fill = variable)) +
#   geom_histogram(binwidth = 0.5) +
#   scale_fill_nejm() +
#   theme_Publication()
# 
# melted_23_2_het_F <- Line_23_2_het_F %>% 
#   rownames_to_column() %>% 
#   melt()
# 
# ggplot(melted_23_2_het_F, aes(x = variable, y = value)) +
#   geom_violin()
# 
# fixation_23_2_plot <- ggplot(melted_23_2_het_F, aes(x = value, fill = variable)) +
#   geom_histogram(binwidth = 0.5) +
#   scale_fill_nejm() +
#   theme_Publication()
# 
# melted_23_4_het_F <- Line_23_4_het_F %>% 
#   rownames_to_column() %>% 
#   melt()
# 
# ggplot(melted_23_4_het_F, aes(x = variable, y = value)) +
#   geom_violin()
# 
# fixation_23_4_plot <- ggplot(melted_23_4_het_F, aes(x = value, fill = variable)) +
#   geom_histogram(binwidth = 0.5) +
#   scale_fill_nejm() +
#   theme_Publication()
# 
# melted_26_1_het_F <- Line_26_1_het_F %>% 
#   rownames_to_column() %>% 
#   melt()
# 
# ggplot(melted_26_1_het_F, aes(x = variable, y = value)) +
#   geom_violin()
# 
# fixation_26_1_plot <- ggplot(melted_26_1_het_F, aes(x = value, fill = variable)) +
#   geom_histogram(binwidth = 0.5) +
#   scale_fill_nejm() +
#   theme_Publication()
# 
# melted_26_4_het_F <- Line_26_4_het_F %>% 
#   rownames_to_column() %>% 
#   melt()
# 
# ggplot(melted_26_4_het_F, aes(x = variable, y = value)) +
#   geom_violin()
# 
# fixation_26_4_plot <- ggplot(melted_26_4_het_F, aes(x = value, fill = variable)) +
#   geom_histogram(binwidth = 0.5) +
#   scale_fill_nejm() +
#   theme_Publication()
# 
# melted_27_2_het_F <- Line_27_2_het_F %>% 
#   rownames_to_column() %>% 
#   melt()
# 
# ggplot(melted_27_2_het_F, aes(x = variable, y = value)) +
#   geom_violin()
# 
# fixation_27_2_plot <- ggplot(melted_27_2_het_F, aes(x = value, fill = variable)) +
#   geom_histogram(binwidth = 0.5) +
#   scale_fill_nejm() +
#   theme_Publication()
# 
# melted_29_2_het_F <- Line_29_2_het_F %>% 
#   rownames_to_column() %>% 
#   melt()
# 
# ggplot(melted_29_2_het_F, aes(x = variable, y = value)) +
#   geom_violin()
# 
# fixation_29_2_plot <- ggplot(melted_29_2_het_F, aes(x = value, fill = variable)) +
#   geom_histogram(binwidth = 0.5) +
#   scale_fill_nejm() +
#   theme_Publication()
# 
# melted_29_4_het_F <- Line_29_4_het_F %>% 
#   rownames_to_column() %>% 
#   melt()
# 
# ggplot(melted_29_4_het_F, aes(x = variable, y = value)) +
#   geom_violin()
# 
# fixation_29_4_plot <- ggplot(melted_29_4_het_F, aes(x = value, fill = variable)) +
#   geom_histogram(binwidth = 0.5) +
#   scale_fill_nejm() +
#   theme_Publication()
# 
# fixation_plots <- ggarrange(fixation_1_1_plot, fixation_6_1_plot, fixation_6_4_plot, fixation_7_2_plot, fixation_7_4_plot, 
#                             fixation_8_2_plot, fixation_8_4_plot, fixation_13_3_plot, fixation_13_4_plot, fixation_16_1_plot, 
#                             fixation_16_5_plot, fixation_17_2_plot, fixation_17_5_plot, fixation_19_2_plot, fixation_19_5_plot, 
#                             fixation_20_1_plot, fixation_20_4_plot, fixation_21_2_plot, fixation_21_6_plot, fixation_23_2_plot, 
#                             fixation_23_4_plot, fixation_26_1_plot, fixation_26_4_plot, fixation_27_2_plot, fixation_29_2_plot, 
#                             fixation_29_4_plot)
# 
# fixation_plots

### With all loci #####################################################################################
### FS gen ##############################################################################################
F_1_1_with_snp_names <- Line_1_1_corrected %>% 
  rownames_to_column() %>% 
  select(1:2)

# F_1_4_with_snp_names <- Line_1_4_corrected %>% 
#   rownames_to_column() %>% 
#   select(1:2)

F_6_1_with_snp_names <- Line_6_1_corrected %>% 
  rownames_to_column() %>% 
  select(1:2)

F_6_4_with_snp_names <- Line_6_4_corrected %>% 
  rownames_to_column() %>% 
  select(1:2)

F_7_2_with_snp_names <- Line_7_2_corrected %>% 
  rownames_to_column() %>% 
  select(1:2)

F_7_4_with_snp_names <- Line_7_4_corrected %>% 
  rownames_to_column() %>% 
  select(1:2)

F_8_2_with_snp_names <- Line_8_2_corrected %>% 
  rownames_to_column() %>% 
  select(1:2)

F_8_4_with_snp_names <- Line_8_4_corrected %>% 
  rownames_to_column() %>% 
  select(1:2)

F_10_2_with_snp_names <- Line_10_2_corrected %>%
  rownames_to_column() %>%
  select(1:2)

F_10_5_with_snp_names <- Line_10_5_corrected %>%
  rownames_to_column() %>%
  select(1:2)

F_13_3_with_snp_names <- Line_13_3_corrected %>% 
  rownames_to_column() %>% 
  select(1:2)

F_13_4_with_snp_names <- Line_13_4_corrected %>% 
  rownames_to_column() %>% 
  select(1:2)

F_16_1_with_snp_names <- Line_16_1_corrected %>% 
  rownames_to_column() %>% 
  select(1:2)

F_16_5_with_snp_names <- Line_16_5_corrected %>% 
  rownames_to_column() %>% 
  select(1:2)

F_17_2_with_snp_names <- Line_17_2_corrected %>% 
  rownames_to_column() %>% 
  select(1:2)

F_17_5_with_snp_names <- Line_17_5_corrected %>% 
  rownames_to_column() %>% 
  select(1:2)

F_19_2_with_snp_names <- Line_19_2_corrected %>% 
  rownames_to_column() %>% 
  select(1:2)

F_19_5_with_snp_names <- Line_19_5_corrected %>% 
  rownames_to_column() %>% 
  select(1:2)

F_20_1_with_snp_names <- Line_20_1_corrected %>% 
  rownames_to_column() %>% 
  select(1:2)

F_20_4_with_snp_names <- Line_20_4_corrected %>% 
  rownames_to_column() %>% 
  select(1:2)

F_21_2_with_snp_names <- Line_21_2_corrected %>% 
  rownames_to_column() %>% 
  select(1:2)

F_21_6_with_snp_names <- Line_21_6_corrected %>% 
  rownames_to_column() %>% 
  select(1:2)

F_23_2_with_snp_names <- Line_23_2_corrected %>%
  rownames_to_column() %>%
  select(1:2)

F_23_4_with_snp_names <- Line_23_4_corrected %>%
  rownames_to_column() %>%
  select(1:2)

F_26_1_with_snp_names <- Line_26_1_corrected %>% 
  rownames_to_column() %>% 
  select(1:2)

F_26_4_with_snp_names <- Line_26_4_corrected %>% 
  rownames_to_column() %>% 
  select(1:2)

F_27_2_with_snp_names <- Line_27_2_corrected %>%
  rownames_to_column() %>%
  select(1:2)

F_29_2_with_snp_names <- Line_29_2_corrected %>% 
  rownames_to_column() %>% 
  select(1:2)

F_29_4_with_snp_names <- Line_29_4_corrected %>% 
  rownames_to_column() %>% 
  select(1:2)


F_lines <- merge(F_1_1_with_snp_names, F_6_1_with_snp_names, by = "rowname", all = T)
F_lines <- merge(F_lines, F_6_4_with_snp_names, by = "rowname", all = T)
F_lines <- merge(F_lines, F_7_2_with_snp_names, by = "rowname", all = T)
F_lines <- merge(F_lines, F_7_4_with_snp_names, by = "rowname", all = T)
F_lines <- merge(F_lines, F_8_2_with_snp_names, by = "rowname", all = T)
F_lines <- merge(F_lines, F_8_4_with_snp_names, by = "rowname", all = T)
F_lines <- merge(F_lines, F_10_2_with_snp_names, by = "rowname", all = T)
F_lines <- merge(F_lines, F_10_5_with_snp_names, by = "rowname", all = T)
F_lines <- merge(F_lines, F_13_3_with_snp_names, by = "rowname", all = T)
F_lines <- merge(F_lines, F_13_4_with_snp_names, by = "rowname", all = T)
F_lines <- merge(F_lines, F_16_1_with_snp_names, by = "rowname", all = T)
F_lines <- merge(F_lines, F_16_5_with_snp_names, by = "rowname", all = T)
F_lines <- merge(F_lines, F_17_2_with_snp_names, by = "rowname", all = T)
F_lines <- merge(F_lines, F_17_5_with_snp_names, by = "rowname", all = T)
F_lines <- merge(F_lines, F_19_2_with_snp_names, by = "rowname", all = T)
F_lines <- merge(F_lines, F_19_5_with_snp_names, by = "rowname", all = T)
F_lines <- merge(F_lines, F_20_1_with_snp_names, by = "rowname", all = T)
F_lines <- merge(F_lines, F_20_4_with_snp_names, by = "rowname", all = T)
F_lines <- merge(F_lines, F_21_2_with_snp_names, by = "rowname", all = T)
F_lines <- merge(F_lines, F_21_6_with_snp_names, by = "rowname", all = T)
F_lines <- merge(F_lines, F_23_2_with_snp_names, by = "rowname", all = T)
F_lines <- merge(F_lines, F_23_4_with_snp_names, by = "rowname", all = T)
F_lines <- merge(F_lines, F_26_1_with_snp_names, by = "rowname", all = T)
F_lines <- merge(F_lines, F_26_4_with_snp_names, by = "rowname", all = T)
F_lines <- merge(F_lines, F_27_2_with_snp_names, by = "rowname", all = T)
F_lines <- merge(F_lines, F_29_2_with_snp_names, by = "rowname", all = T)
F_lines <- merge(F_lines, F_29_4_with_snp_names, by = "rowname", all = T)

# Minor allele frequencies
F_lines <- F_lines %>% 
  column_to_rownames()

n0 <- apply(F_lines==0,1,sum,na.rm=T)
n1 <- apply(F_lines==1,1,sum,na.rm=T)
n2 <- apply(F_lines==2,1,sum,na.rm=T)

n <- n0 + n1 + n2

p <- ((2*n0)+n1)/(2*n)
q <- 1 - p
maf <- pmin(p, q)
mgf <- apply(cbind(n0,n1,n2),1,min) / n

# allele_freqs_F <- Propfunc(F_lines)
allele_freqs_F <- data.frame(SNP = row.names(F_lines), frq = maf)

ggplot(allele_freqs_F, aes(x = frq)) +
  geom_density()

### S1 gen ##############################################################################################
S1_1_1_with_snp_names <- Line_1_1_corrected %>% 
  rownames_to_column() %>% 
  select(1, 3)

# S1_1_4_with_snp_names <- Line_1_4_corrected %>% 
#   rownames_to_column() %>% 
#   select(1, 3)

S1_6_1_with_snp_names <- Line_6_1_corrected %>% 
  rownames_to_column() %>% 
  select(1, 3)

S1_6_4_with_snp_names <- Line_6_4_corrected %>% 
  rownames_to_column() %>% 
  select(1, 3)

S1_7_2_with_snp_names <- Line_7_2_corrected %>% 
  rownames_to_column() %>% 
  select(1, 3)

S1_7_4_with_snp_names <- Line_7_4_corrected %>% 
  rownames_to_column() %>% 
  select(1, 3)

S1_8_2_with_snp_names <- Line_8_2_corrected %>% 
  rownames_to_column() %>% 
  select(1, 3)

S1_8_4_with_snp_names <- Line_8_4_corrected %>% 
  rownames_to_column() %>% 
  select(1, 3)

S1_10_2_with_snp_names <- Line_10_2_corrected %>%
  rownames_to_column() %>%
  select(1, 3)

S1_10_5_with_snp_names <- Line_10_5_corrected %>%
  rownames_to_column() %>%
  select(1, 3)

S1_13_3_with_snp_names <- Line_13_3_corrected %>% 
  rownames_to_column() %>% 
  select(1, 3)

S1_13_4_with_snp_names <- Line_13_4_corrected %>% 
  rownames_to_column() %>% 
  select(1, 3)

S1_16_1_with_snp_names <- Line_16_1_corrected %>% 
  rownames_to_column() %>% 
  select(1, 3)

S1_16_5_with_snp_names <- Line_16_5_corrected %>% 
  rownames_to_column() %>% 
  select(1, 3)

S1_17_2_with_snp_names <- Line_17_2_corrected %>% 
  rownames_to_column() %>% 
  select(1, 3)

S1_17_5_with_snp_names <- Line_17_5_corrected %>% 
  rownames_to_column() %>% 
  select(1, 3)

S1_19_2_with_snp_names <- Line_19_2_corrected %>% 
  rownames_to_column() %>% 
  select(1, 3)

S1_19_5_with_snp_names <- Line_19_5_corrected %>% 
  rownames_to_column() %>% 
  select(1, 3)

S1_20_1_with_snp_names <- Line_20_1_corrected %>% 
  rownames_to_column() %>% 
  select(1, 3)

S1_20_4_with_snp_names <- Line_20_4_corrected %>% 
  rownames_to_column() %>% 
  select(1, 3)

S1_21_2_with_snp_names <- Line_21_2_corrected %>% 
  rownames_to_column() %>% 
  select(1, 3)

S1_21_6_with_snp_names <- Line_21_6_corrected %>% 
  rownames_to_column() %>% 
  select(1, 3)

S1_23_2_with_snp_names <- Line_23_2_corrected %>% 
  rownames_to_column() %>% 
  select(1, 3)

S1_23_4_with_snp_names <- Line_23_4_corrected %>% 
  rownames_to_column() %>% 
  select(1, 3)

S1_26_1_with_snp_names <- Line_26_1_corrected %>% 
  rownames_to_column() %>% 
  select(1, 3)

S1_26_1_with_snp_names <- Line_26_1_corrected %>% 
  rownames_to_column() %>% 
  select(1, 3)

S1_26_4_with_snp_names <- Line_26_4_corrected %>% 
  rownames_to_column() %>% 
  select(1, 3)

S1_27_2_with_snp_names <- Line_27_2_corrected %>%
  rownames_to_column() %>%
  select(1, 3)

S1_29_2_with_snp_names <- Line_29_2_corrected %>% 
  rownames_to_column() %>% 
  select(1, 3)

S1_29_4_with_snp_names <- Line_29_4_corrected %>% 
  rownames_to_column() %>% 
  select(1, 3)


S1_lines <- merge(S1_1_1_with_snp_names, S1_6_1_with_snp_names, by = "rowname", all = T)
S1_lines <- merge(S1_lines, S1_6_4_with_snp_names, by = "rowname", all = T)
S1_lines <- merge(S1_lines, S1_7_2_with_snp_names, by = "rowname", all = T)
S1_lines <- merge(S1_lines, S1_7_4_with_snp_names, by = "rowname", all = T)
S1_lines <- merge(S1_lines, S1_8_2_with_snp_names, by = "rowname", all = T)
S1_lines <- merge(S1_lines, S1_8_4_with_snp_names, by = "rowname", all = T)
S1_lines <- merge(S1_lines, S1_10_2_with_snp_names, by = "rowname", all = T)
S1_lines <- merge(S1_lines, S1_10_5_with_snp_names, by = "rowname", all = T)
S1_lines <- merge(S1_lines, S1_13_3_with_snp_names, by = "rowname", all = T)
S1_lines <- merge(S1_lines, S1_13_4_with_snp_names, by = "rowname", all = T)
S1_lines <- merge(S1_lines, S1_16_1_with_snp_names, by = "rowname", all = T)
S1_lines <- merge(S1_lines, S1_16_5_with_snp_names, by = "rowname", all = T)
S1_lines <- merge(S1_lines, S1_17_2_with_snp_names, by = "rowname", all = T)
S1_lines <- merge(S1_lines, S1_17_5_with_snp_names, by = "rowname", all = T)
S1_lines <- merge(S1_lines, S1_19_2_with_snp_names, by = "rowname", all = T)
S1_lines <- merge(S1_lines, S1_19_5_with_snp_names, by = "rowname", all = T)
S1_lines <- merge(S1_lines, S1_20_1_with_snp_names, by = "rowname", all = T)
S1_lines <- merge(S1_lines, S1_20_4_with_snp_names, by = "rowname", all = T)
S1_lines <- merge(S1_lines, S1_21_2_with_snp_names, by = "rowname", all = T)
S1_lines <- merge(S1_lines, S1_21_6_with_snp_names, by = "rowname", all = T)
S1_lines <- merge(S1_lines, S1_23_2_with_snp_names, by = "rowname", all = T)
S1_lines <- merge(S1_lines, S1_23_4_with_snp_names, by = "rowname", all = T)
S1_lines <- merge(S1_lines, S1_26_1_with_snp_names, by = "rowname", all = T)
S1_lines <- merge(S1_lines, S1_26_4_with_snp_names, by = "rowname", all = T)
S1_lines <- merge(S1_lines, S1_27_2_with_snp_names, by = "rowname", all = T)
S1_lines <- merge(S1_lines, S1_29_2_with_snp_names, by = "rowname", all = T)
S1_lines <- merge(S1_lines, S1_29_4_with_snp_names, by = "rowname", all = T)

# Minor allele freqs
S1_lines <- S1_lines %>% 
  column_to_rownames()

n0 <- apply(S1_lines==0,1,sum,na.rm=T)
n1 <- apply(S1_lines==1,1,sum,na.rm=T)
n2 <- apply(S1_lines==2,1,sum,na.rm=T)

n <- n0 + n1 + n2

p <- ((2*n0)+n1)/(2*n)
q <- 1 - p
maf <- pmin(p, q)
mgf <- apply(cbind(n0,n1,n2),1,min) / n

# allele_freqs_S1 <- Propfunc(S1_lines)
allele_freqs_S1 <- data.frame(SNP = rownames(S1_lines), frq = maf)

ggplot(allele_freqs_S1, aes(x = frq)) +
  geom_histogram(binwidth = 0.01)

### S2 gen ##############################################################################################
S2_1_1_with_snp_names <- Line_1_1_corrected %>% 
  rownames_to_column() %>% 
  select(1, 4)

# S2_1_4_with_snp_names <- Line_1_4_corrected %>% 
#   rownames_to_column() %>% 
#   select(1, 4)

S2_6_1_with_snp_names <- Line_6_1_corrected %>% 
  rownames_to_column() %>% 
  select(1, 4)

S2_6_4_with_snp_names <- Line_6_4_corrected %>% 
  rownames_to_column() %>% 
  select(1, 4)

S2_7_2_with_snp_names <- Line_7_2_corrected %>% 
  rownames_to_column() %>% 
  select(1, 4)

S2_7_4_with_snp_names <- Line_7_4_corrected %>% 
  rownames_to_column() %>% 
  select(1, 4)

S2_8_2_with_snp_names <- Line_8_2_corrected %>% 
  rownames_to_column() %>% 
  select(1, 4)

S2_8_4_with_snp_names <- Line_8_4_corrected %>% 
  rownames_to_column() %>% 
  select(1, 4)

S2_10_2_with_snp_names <- Line_10_2_corrected %>%
  rownames_to_column() %>%
  select(1, 4)

S2_10_5_with_snp_names <- Line_10_5_corrected %>%
  rownames_to_column() %>%
  select(1, 4)

S2_13_3_with_snp_names <- Line_13_3_corrected %>% 
  rownames_to_column() %>% 
  select(1, 4)

S2_13_4_with_snp_names <- Line_13_4_corrected %>% 
  rownames_to_column() %>% 
  select(1, 4)

S2_16_1_with_snp_names <- Line_16_1_corrected %>% 
  rownames_to_column() %>% 
  select(1, 4)

S2_16_5_with_snp_names <- Line_16_5_corrected %>% 
  rownames_to_column() %>% 
  select(1, 4)

S2_17_2_with_snp_names <- Line_17_2_corrected %>% 
  rownames_to_column() %>% 
  select(1, 4)

S2_17_5_with_snp_names <- Line_17_5_corrected %>% 
  rownames_to_column() %>% 
  select(1, 4)

S2_19_2_with_snp_names <- Line_19_2_corrected %>% 
  rownames_to_column() %>% 
  select(1, 4)

S2_19_5_with_snp_names <- Line_19_5_corrected %>% 
  rownames_to_column() %>% 
  select(1, 4)

S2_20_1_with_snp_names <- Line_20_1_corrected %>% 
  rownames_to_column() %>% 
  select(1, 4)

S2_20_4_with_snp_names <- Line_20_4_corrected %>% 
  rownames_to_column() %>% 
  select(1, 4)

S2_21_2_with_snp_names <- Line_21_2_corrected %>% 
  rownames_to_column() %>% 
  select(1, 4)

S2_21_6_with_snp_names <- Line_21_6_corrected %>% 
  rownames_to_column() %>% 
  select(1, 4)

S2_23_2_with_snp_names <- Line_23_2_corrected %>% 
  rownames_to_column() %>% 
  select(1, 4)

S2_23_4_with_snp_names <- Line_23_4_corrected %>% 
  rownames_to_column() %>% 
  select(1, 4)

S2_26_1_with_snp_names <- Line_26_1_corrected %>% 
  rownames_to_column() %>% 
  select(1, 4)

S2_26_1_with_snp_names <- Line_26_1_corrected %>% 
  rownames_to_column() %>% 
  select(1, 4)

S2_26_4_with_snp_names <- Line_26_4_corrected %>% 
  rownames_to_column() %>% 
  select(1, 4)

S2_27_2_with_snp_names <- Line_27_2_corrected %>%
  rownames_to_column() %>%
  select(1, 4)

S2_29_2_with_snp_names <- Line_29_2_corrected %>% 
  rownames_to_column() %>% 
  select(1, 4)

S2_29_4_with_snp_names <- Line_29_4_corrected %>% 
  rownames_to_column() %>% 
  select(1, 4)


S2_lines <- merge(S2_1_1_with_snp_names, S2_6_1_with_snp_names, by = "rowname", all = T)
S2_lines <- merge(S2_lines, S2_6_4_with_snp_names, by = "rowname", all = T)
S2_lines <- merge(S2_lines, S2_7_2_with_snp_names, by = "rowname", all = T)
S2_lines <- merge(S2_lines, S2_7_4_with_snp_names, by = "rowname", all = T)
S2_lines <- merge(S2_lines, S2_8_2_with_snp_names, by = "rowname", all = T)
S2_lines <- merge(S2_lines, S2_8_4_with_snp_names, by = "rowname", all = T)
S2_lines <- merge(S2_lines, S2_10_2_with_snp_names, by = "rowname", all = T)
S2_lines <- merge(S2_lines, S2_10_5_with_snp_names, by = "rowname", all = T)
S2_lines <- merge(S2_lines, S2_13_3_with_snp_names, by = "rowname", all = T)
S2_lines <- merge(S2_lines, S2_13_4_with_snp_names, by = "rowname", all = T)
S2_lines <- merge(S2_lines, S2_16_1_with_snp_names, by = "rowname", all = T)
S2_lines <- merge(S2_lines, S2_16_5_with_snp_names, by = "rowname", all = T)
S2_lines <- merge(S2_lines, S2_17_2_with_snp_names, by = "rowname", all = T)
S2_lines <- merge(S2_lines, S2_17_5_with_snp_names, by = "rowname", all = T)
S2_lines <- merge(S2_lines, S2_19_2_with_snp_names, by = "rowname", all = T)
S2_lines <- merge(S2_lines, S2_19_5_with_snp_names, by = "rowname", all = T)
S2_lines <- merge(S2_lines, S2_20_1_with_snp_names, by = "rowname", all = T)
S2_lines <- merge(S2_lines, S2_20_4_with_snp_names, by = "rowname", all = T)
S2_lines <- merge(S2_lines, S2_21_2_with_snp_names, by = "rowname", all = T)
S2_lines <- merge(S2_lines, S2_21_6_with_snp_names, by = "rowname", all = T)
S2_lines <- merge(S2_lines, S2_23_2_with_snp_names, by = "rowname", all = T)
S2_lines <- merge(S2_lines, S2_23_4_with_snp_names, by = "rowname", all = T)
S2_lines <- merge(S2_lines, S2_26_1_with_snp_names, by = "rowname", all = T)
S2_lines <- merge(S2_lines, S2_26_4_with_snp_names, by = "rowname", all = T)
S2_lines <- merge(S2_lines, S2_27_2_with_snp_names, by = "rowname", all = T)
S2_lines <- merge(S2_lines, S2_29_2_with_snp_names, by = "rowname", all = T)
S2_lines <- merge(S2_lines, S2_29_4_with_snp_names, by = "rowname", all = T)

# Minor allele freqs
S2_lines <- S2_lines %>% 
  column_to_rownames()

n0 <- apply(S2_lines==0,1,sum,na.rm=T)
n1 <- apply(S2_lines==1,1,sum,na.rm=T)
n2 <- apply(S2_lines==2,1,sum,na.rm=T)

n <- n0 + n1 + n2

p <- ((2*n0)+n1)/(2*n)
q <- 1 - p
maf <- pmin(p, q)
mgf <- apply(cbind(n0,n1,n2),1,min) / n

#allele_freqs_S2 <- Propfunc(S2_lines)
allele_freqs_S2 <- data.frame(SNP = rownames(S2_lines), frq = maf)

ggplot(allele_freqs_S2, aes(x = frq)) +
  geom_density()

### S3 gen ##############################################################################################
S3_1_1_with_snp_names <- Line_1_1_corrected %>% 
  rownames_to_column() %>% 
  select(1, 5)

# S3_1_4_with_snp_names <- Line_1_4_corrected %>% 
#   rownames_to_column() %>% 
#   select(1, 5)

S3_6_1_with_snp_names <- Line_6_1_corrected %>% 
  rownames_to_column() %>% 
  select(1, 5)

S3_6_4_with_snp_names <- Line_6_4_corrected %>% 
  rownames_to_column() %>% 
  select(1, 5)

S3_7_2_with_snp_names <- Line_7_2_corrected %>% 
  rownames_to_column() %>% 
  select(1, 5)

S3_7_4_with_snp_names <- Line_7_4_corrected %>% 
  rownames_to_column() %>% 
  select(1, 5)

S3_8_2_with_snp_names <- Line_8_2_corrected %>% 
  rownames_to_column() %>% 
  select(1, 5)

S3_8_4_with_snp_names <- Line_8_4_corrected %>% 
  rownames_to_column() %>% 
  select(1, 5)

S3_10_2_with_snp_names <- Line_10_2_corrected %>%
  rownames_to_column() %>%
  select(1, 5)

S3_10_5_with_snp_names <- Line_10_5_corrected %>%
  rownames_to_column() %>%
  select(1, 5)

S3_13_3_with_snp_names <- Line_13_3_corrected %>% 
  rownames_to_column() %>% 
  select(1, 5)

S3_13_4_with_snp_names <- Line_13_4_corrected %>% 
  rownames_to_column() %>% 
  select(1, 5)

S3_16_1_with_snp_names <- Line_16_1_corrected %>% 
  rownames_to_column() %>% 
  select(1, 5)

S3_16_5_with_snp_names <- Line_16_5_corrected %>% 
  rownames_to_column() %>% 
  select(1, 5)

S3_17_2_with_snp_names <- Line_17_2_corrected %>% 
  rownames_to_column() %>% 
  select(1, 5)

S3_17_5_with_snp_names <- Line_17_5_corrected %>% 
  rownames_to_column() %>% 
  select(1, 5)

S3_19_2_with_snp_names <- Line_19_2_corrected %>% 
  rownames_to_column() %>% 
  select(1, 5)

S3_19_5_with_snp_names <- Line_19_5_corrected %>% 
  rownames_to_column() %>% 
  select(1, 5)

S3_20_1_with_snp_names <- Line_20_1_corrected %>% 
  rownames_to_column() %>% 
  select(1, 5)

S3_20_4_with_snp_names <- Line_20_4_corrected %>% 
  rownames_to_column() %>% 
  select(1, 5)

S3_21_2_with_snp_names <- Line_21_2_corrected %>% 
  rownames_to_column() %>% 
  select(1, 5)

S3_21_6_with_snp_names <- Line_21_6_corrected %>% 
  rownames_to_column() %>% 
  select(1, 5)

S3_23_2_with_snp_names <- Line_23_2_corrected %>% 
  rownames_to_column() %>% 
  select(1, 5)

S3_23_4_with_snp_names <- Line_23_4_corrected %>% 
  rownames_to_column() %>% 
  select(1, 5)

S3_26_1_with_snp_names <- Line_26_1_corrected %>% 
  rownames_to_column() %>% 
  select(1, 5)

S3_26_1_with_snp_names <- Line_26_1_corrected %>% 
  rownames_to_column() %>% 
  select(1, 5)

S3_26_4_with_snp_names <- Line_26_4_corrected %>% 
  rownames_to_column() %>% 
  select(1, 5)

S3_27_2_with_snp_names <- Line_27_2_corrected %>%
  rownames_to_column() %>%
  select(1, 5)

S3_29_2_with_snp_names <- Line_29_2_corrected %>% 
  rownames_to_column() %>% 
  select(1, 5)

S3_29_4_with_snp_names <- Line_29_4_corrected %>% 
  rownames_to_column() %>% 
  select(1, 5)


S3_lines <- merge(S3_1_1_with_snp_names, S3_6_1_with_snp_names, by = "rowname", all = T)
S3_lines <- merge(S3_lines, S3_6_4_with_snp_names, by = "rowname", all = T)
S3_lines <- merge(S3_lines, S3_7_2_with_snp_names, by = "rowname", all = T)
S3_lines <- merge(S3_lines, S3_7_4_with_snp_names, by = "rowname", all = T)
S3_lines <- merge(S3_lines, S3_8_2_with_snp_names, by = "rowname", all = T)
S3_lines <- merge(S3_lines, S3_8_4_with_snp_names, by = "rowname", all = T)
S3_lines <- merge(S3_lines, S3_10_2_with_snp_names, by = "rowname", all = T)
S3_lines <- merge(S3_lines, S3_10_5_with_snp_names, by = "rowname", all = T)
S3_lines <- merge(S3_lines, S3_13_3_with_snp_names, by = "rowname", all = T)
S3_lines <- merge(S3_lines, S3_13_4_with_snp_names, by = "rowname", all = T)
S3_lines <- merge(S3_lines, S3_16_1_with_snp_names, by = "rowname", all = T)
S3_lines <- merge(S3_lines, S3_16_5_with_snp_names, by = "rowname", all = T)
S3_lines <- merge(S3_lines, S3_17_2_with_snp_names, by = "rowname", all = T)
S3_lines <- merge(S3_lines, S3_17_5_with_snp_names, by = "rowname", all = T)
S3_lines <- merge(S3_lines, S3_19_2_with_snp_names, by = "rowname", all = T)
S3_lines <- merge(S3_lines, S3_19_5_with_snp_names, by = "rowname", all = T)
S3_lines <- merge(S3_lines, S3_20_1_with_snp_names, by = "rowname", all = T)
S3_lines <- merge(S3_lines, S3_20_4_with_snp_names, by = "rowname", all = T)
S3_lines <- merge(S3_lines, S3_21_2_with_snp_names, by = "rowname", all = T)
S3_lines <- merge(S3_lines, S3_21_6_with_snp_names, by = "rowname", all = T)
S3_lines <- merge(S3_lines, S3_23_2_with_snp_names, by = "rowname", all = T)
S3_lines <- merge(S3_lines, S3_23_4_with_snp_names, by = "rowname", all = T)
S3_lines <- merge(S3_lines, S3_26_1_with_snp_names, by = "rowname", all = T)
S3_lines <- merge(S3_lines, S3_26_4_with_snp_names, by = "rowname", all = T)
S3_lines <- merge(S3_lines, S3_27_2_with_snp_names, by = "rowname", all = T)
S3_lines <- merge(S3_lines, S3_29_2_with_snp_names, by = "rowname", all = T)
S3_lines <- merge(S3_lines, S3_29_4_with_snp_names, by = "rowname", all = T)

# Minor allele freqs
S3_lines <- S3_lines %>% 
  column_to_rownames()

n0 <- apply(S3_lines==0,1,sum,na.rm=T)
n1 <- apply(S3_lines==1,1,sum,na.rm=T)
n2 <- apply(S3_lines==2,1,sum,na.rm=T)

n <- n0 + n1 + n2

p <- ((2*n0)+n1)/(2*n)
q <- 1 - p
maf <- pmin(p, q)
mgf <- apply(cbind(n0,n1,n2),1,min) / n

#allele_freqs_S3 <- Propfunc(S3_lines)
allele_freqs_S3 <- data.frame(SNP = rownames(S3_lines), frq = maf)

ggplot(allele_freqs_S3, aes(x = frq)) +
  geom_density()

### S4 gen ##############################################################################################
S4_1_1_with_snp_names <- Line_1_1_corrected %>% 
  rownames_to_column() %>% 
  select(1, 6)

# S4_1_4_with_snp_names <- Line_1_4_corrected %>% 
#   rownames_to_column() %>% 
#   select(1, 6)

S4_6_1_with_snp_names <- Line_6_1_corrected %>% 
  rownames_to_column() %>% 
  select(1, 6)

S4_6_4_with_snp_names <- Line_6_4_corrected %>% 
  rownames_to_column() %>% 
  select(1, 6)

S4_7_2_with_snp_names <- Line_7_2_corrected %>% 
  rownames_to_column() %>% 
  select(1, 6)

S4_7_4_with_snp_names <- Line_7_4_corrected %>% 
  rownames_to_column() %>% 
  select(1, 6)

S4_8_2_with_snp_names <- Line_8_2_corrected %>% 
  rownames_to_column() %>% 
  select(1, 6)

S4_8_4_with_snp_names <- Line_8_4_corrected %>% 
  rownames_to_column() %>% 
  select(1, 6)

S4_10_2_with_snp_names <- Line_10_2_corrected %>%
  rownames_to_column() %>%
  select(1, 6)

S4_10_5_with_snp_names <- Line_10_5_corrected %>%
  rownames_to_column() %>%
  select(1, 6)

S4_13_3_with_snp_names <- Line_13_3_corrected %>% 
  rownames_to_column() %>% 
  select(1, 6)

S4_13_4_with_snp_names <- Line_13_4_corrected %>% 
  rownames_to_column() %>% 
  select(1, 6)

S4_16_1_with_snp_names <- Line_16_1_corrected %>% 
  rownames_to_column() %>% 
  select(1, 6)

S4_16_5_with_snp_names <- Line_16_5_corrected %>% 
  rownames_to_column() %>% 
  select(1, 6)

S4_17_2_with_snp_names <- Line_17_2_corrected %>% 
  rownames_to_column() %>% 
  select(1, 6)

S4_17_5_with_snp_names <- Line_17_5_corrected %>% 
  rownames_to_column() %>% 
  select(1, 6)

S4_19_2_with_snp_names <- Line_19_2_corrected %>% 
  rownames_to_column() %>% 
  select(1, 6)

S4_19_5_with_snp_names <- Line_19_5_corrected %>% 
  rownames_to_column() %>% 
  select(1, 6)

S4_20_1_with_snp_names <- Line_20_1_corrected %>% 
  rownames_to_column() %>% 
  select(1, 6)

S4_20_4_with_snp_names <- Line_20_4_corrected %>% 
  rownames_to_column() %>% 
  select(1, 6)

S4_21_2_with_snp_names <- Line_21_2_corrected %>% 
  rownames_to_column() %>% 
  select(1, 6)

S4_21_6_with_snp_names <- Line_21_6_corrected %>% 
  rownames_to_column() %>% 
  select(1, 6)

S4_23_2_with_snp_names <- Line_23_2_corrected %>% 
  rownames_to_column() %>% 
  select(1, 6)

S4_23_4_with_snp_names <- Line_23_4_corrected %>% 
  rownames_to_column() %>% 
  select(1, 6)

S4_26_1_with_snp_names <- Line_26_1_corrected %>% 
  rownames_to_column() %>% 
  select(1, 6)

S4_26_4_with_snp_names <- Line_26_4_corrected %>% 
  rownames_to_column() %>% 
  select(1, 6)

S4_27_2_with_snp_names <- Line_27_2_corrected %>%
  rownames_to_column() %>%
  select(1, 6)

S4_29_2_with_snp_names <- Line_29_2_corrected %>% 
  rownames_to_column() %>% 
  select(1, 6)

S4_29_4_with_snp_names <- Line_29_4_corrected %>% 
  rownames_to_column() %>% 
  select(1, 6)


S4_lines <- merge(S4_1_1_with_snp_names, S4_6_1_with_snp_names, by = "rowname", all = T)
S4_lines <- merge(S4_lines, S4_6_4_with_snp_names, by = "rowname", all = T)
S4_lines <- merge(S4_lines, S4_7_2_with_snp_names, by = "rowname", all = T)
S4_lines <- merge(S4_lines, S4_7_4_with_snp_names, by = "rowname", all = T)
S4_lines <- merge(S4_lines, S4_8_2_with_snp_names, by = "rowname", all = T)
S4_lines <- merge(S4_lines, S4_8_4_with_snp_names, by = "rowname", all = T)
S4_lines <- merge(S4_lines, S4_10_2_with_snp_names, by = "rowname", all = T)
S4_lines <- merge(S4_lines, S4_10_5_with_snp_names, by = "rowname", all = T)
S4_lines <- merge(S4_lines, S4_13_3_with_snp_names, by = "rowname", all = T)
S4_lines <- merge(S4_lines, S4_13_4_with_snp_names, by = "rowname", all = T)
S4_lines <- merge(S4_lines, S4_16_1_with_snp_names, by = "rowname", all = T)
S4_lines <- merge(S4_lines, S4_16_5_with_snp_names, by = "rowname", all = T)
S4_lines <- merge(S4_lines, S4_17_2_with_snp_names, by = "rowname", all = T)
S4_lines <- merge(S4_lines, S4_17_5_with_snp_names, by = "rowname", all = T)
S4_lines <- merge(S4_lines, S4_19_2_with_snp_names, by = "rowname", all = T)
S4_lines <- merge(S4_lines, S4_19_5_with_snp_names, by = "rowname", all = T)
S4_lines <- merge(S4_lines, S4_20_1_with_snp_names, by = "rowname", all = T)
S4_lines <- merge(S4_lines, S4_20_4_with_snp_names, by = "rowname", all = T)
S4_lines <- merge(S4_lines, S4_21_2_with_snp_names, by = "rowname", all = T)
S4_lines <- merge(S4_lines, S4_21_6_with_snp_names, by = "rowname", all = T)
S4_lines <- merge(S4_lines, S4_23_2_with_snp_names, by = "rowname", all = T)
S4_lines <- merge(S4_lines, S4_23_4_with_snp_names, by = "rowname", all = T)
S4_lines <- merge(S4_lines, S4_26_1_with_snp_names, by = "rowname", all = T)
S4_lines <- merge(S4_lines, S4_26_4_with_snp_names, by = "rowname", all = T)
S4_lines <- merge(S4_lines, S4_27_2_with_snp_names, by = "rowname", all = T)
S4_lines <- merge(S4_lines, S4_29_2_with_snp_names, by = "rowname", all = T)
S4_lines <- merge(S4_lines, S4_29_4_with_snp_names, by = "rowname", all = T)

# Minor allele freqs
S4_lines <- S4_lines %>% 
  column_to_rownames()

n0 <- apply(S4_lines==0,1,sum,na.rm=T)
n1 <- apply(S4_lines==1,1,sum,na.rm=T)
n2 <- apply(S4_lines==2,1,sum,na.rm=T)

n <- n0 + n1 + n2

p <- ((2*n0)+n1)/(2*n)
q <- 1 - p
maf <- pmin(p, q)
mgf <- apply(cbind(n0,n1,n2),1,min) / n

#allele_freqs_S4 <- Propfunc(S4_lines)
allele_freqs_S4 <- data.frame(SNP = rownames(S4_lines), frq = maf)

ggplot(allele_freqs_S4, aes(x = frq)) +
  geom_histogram(binwidth = 0.01)

### S5 gen ##############################################################################################
S5_1_1_with_snp_names <- Line_1_1_corrected %>% 
  rownames_to_column() %>% 
  select(1, 7)

# S5_1_4_with_snp_names <- Line_1_4_corrected %>% 
#   rownames_to_column() %>% 
#   select(1, 7)

S5_6_4_with_snp_names <- Line_6_4_corrected %>% 
  rownames_to_column() %>% 
  select(1, 7)

S5_10_2_with_snp_names <- Line_10_2_corrected %>%
  rownames_to_column() %>%
  select(1, 7)

S5_16_5_with_snp_names <- Line_16_5_corrected %>% 
  rownames_to_column() %>% 
  select(1, 7)

S5_17_2_with_snp_names <- Line_17_2_corrected %>% 
  rownames_to_column() %>% 
  select(1, 7)

S5_17_5_with_snp_names <- Line_17_5_corrected %>% 
  rownames_to_column() %>% 
  select(1, 7)

S5_21_2_with_snp_names <- Line_21_2_corrected %>% 
  rownames_to_column() %>% 
  select(1, 7)

S5_21_6_with_snp_names <- Line_21_6_corrected %>% 
  rownames_to_column() %>% 
  select(1, 7)

S5_23_2_with_snp_names <- Line_23_2_corrected %>% 
  rownames_to_column() %>% 
  select(1, 7)

S5_29_2_with_snp_names <- Line_29_2_corrected %>% 
  rownames_to_column() %>% 
  select(1, 7)

S5_29_4_with_snp_names <- Line_29_4_corrected %>% 
  rownames_to_column() %>% 
  select(1, 7)


S5_lines <- merge(S5_1_1_with_snp_names, S5_6_4_with_snp_names, by = "rowname", all = T)
S5_lines <- merge(S5_lines, S5_10_2_with_snp_names, by = "rowname", all = T)
S5_lines <- merge(S5_lines, S5_17_2_with_snp_names, by = "rowname", all = T)
S5_lines <- merge(S5_lines, S5_17_5_with_snp_names, by = "rowname", all = T)
S5_lines <- merge(S5_lines, S5_21_2_with_snp_names, by = "rowname", all = T)
S5_lines <- merge(S5_lines, S5_21_6_with_snp_names, by = "rowname", all = T)
S5_lines <- merge(S5_lines, S5_23_2_with_snp_names, by = "rowname", all = T)
S5_lines <- merge(S5_lines, S5_29_2_with_snp_names, by = "rowname", all = T)
S5_lines <- merge(S5_lines, S5_29_4_with_snp_names, by = "rowname", all = T)

# Minor allele freqs
S5_lines <- S5_lines %>% 
  column_to_rownames()

#### Combine all gens (all) ########################
gt_1_1 <- data.frame(SNP = F_1_1_with_snp_names$rowname, `1_1-F` = F_1_1_with_snp_names$`1_1-F`, 
                     `111-S1` = S1_1_1_with_snp_names$`111-S1`, `111-2-S2` = S2_1_1_with_snp_names$`111-2-S2`, 
                     `111-22-S3` = S3_1_1_with_snp_names$`111-22-S3`, `111-221-S4` = S4_1_1_with_snp_names$`111-221-S4`, 
                     `111-221-S5` = S5_1_1_with_snp_names$`111-221-S5`, check.names = F)


gt_6_1 <- data.frame(SNP = F_6_1_with_snp_names$rowname, `6_1-F` = F_6_1_with_snp_names$`6_1-F`, 
                     `611-S1` = S1_6_1_with_snp_names$`611-S1`, `611-1-S2` = S2_6_1_with_snp_names$`611-1-S2`, 
                     `611-11-S3` = S3_6_1_with_snp_names$`611-11-S3`, `611-111-S4` = S4_6_1_with_snp_names$`611-111-S4`, check.names = F)


gt_6_4 <- data.frame(SNP = F_6_4_with_snp_names$rowname, `6_4-F` = F_6_4_with_snp_names$`6_4-F`, 
                     `646-S1` = S1_6_4_with_snp_names$`646-S1`, `646-4-S2` = S2_6_4_with_snp_names$`646-4-S2`, 
                     `646-45-S3` = S3_6_4_with_snp_names$`646-45-S3`, `646-454-S4` = S4_6_4_with_snp_names$`646-454-S4`, 
                     `646-454-S5` = S5_6_4_with_snp_names$`646-454-S5`, check.names = F)


gt_7_2 <- data.frame(SNP = F_7_2_with_snp_names$rowname, `7_2-F` = F_7_2_with_snp_names$`7_2-F`, 
                     `721-S1` = S1_7_2_with_snp_names$`721-S1`, `721-2-S2` = S2_7_2_with_snp_names$`721-2-S2`, 
                     `721-21-S3` = S3_7_2_with_snp_names$`721-21-S3`, `721-211-S4` = S4_7_2_with_snp_names$`721-211-S4`, check.names = F)


gt_7_4 <- data.frame(SNP = F_7_4_with_snp_names$rowname, `7_4-F` = F_7_4_with_snp_names$`7_4-F`, 
                     `745-S1` = S1_7_4_with_snp_names$`745-S1`, `745-4-S2` = S2_7_4_with_snp_names$`745-4-S2`, 
                     `745-44-S3` = S3_7_4_with_snp_names$`745-44-S3`, `745-444-S4` = S4_7_4_with_snp_names$`745-444-S4`, check.names = F)


gt_8_2 <- data.frame(SNP = F_8_2_with_snp_names$rowname, `8_2-F` = F_8_2_with_snp_names$`8_2-F`, 
                     `821-S1` = S1_8_2_with_snp_names$`821-S1`, `821-1-S2` = S2_8_2_with_snp_names$`821-1-S2`, 
                     `821-11-S3` = S3_8_2_with_snp_names$`821-11-S3`, `821-111-S4` = S4_8_2_with_snp_names$`821-111-S4`, check.names = F)


gt_8_4 <- data.frame(SNP = F_8_4_with_snp_names$rowname, `8_4-F` = F_8_4_with_snp_names$`8_4-F`, 
                     `845-S1` = S1_8_4_with_snp_names$`845-S1`, `845-4-S2` = S2_8_4_with_snp_names$`845-4-S2`, 
                     `845-44-S3` = S3_8_4_with_snp_names$`845-44-S3`, `845-444-S4` = S4_8_4_with_snp_names$`845-444-S4`, check.names = F)


gt_10_2 <- data.frame(SNP = F_10_2_with_snp_names$rowname, `10_2-F` = F_10_2_with_snp_names$`10_2-F`, 
                      `1022-S1` = S1_10_2_with_snp_names$`1022-S1`, `1022-1-S2` = S2_10_2_with_snp_names$`1022-1-S2`, 
                      `1022-11-S3` = S3_10_2_with_snp_names$`1022-11-S3`, `1022-113-S4` = S4_10_2_with_snp_names$`1022-113-S4`,
                      `1022-113-S5` = S5_10_2_with_snp_names$`1022-113-S5`, check.names = F)


gt_10_5 <- data.frame(SNP = F_10_5_with_snp_names$rowname, `10_5-F` = F_10_5_with_snp_names$`10_5-F`, 
                      `1056-S1` = S1_10_5_with_snp_names$`1056-S1`, `1056-4-S2` = S2_10_5_with_snp_names$`1056-4-S2`, 
                      `1056-45-S3` = S3_10_5_with_snp_names$`1056-45-S3`, `1056-454-S4` = S4_10_5_with_snp_names$`1056-454-S4`, check.names = F)


gt_13_3 <- data.frame(SNP = F_13_3_with_snp_names$rowname, `13_3-F` = F_13_3_with_snp_names$`13_3-F`, 
                      `1332-S1` = S1_13_3_with_snp_names$`1332-S1`, `1332-1-S2` = S2_13_3_with_snp_names$`1332-1-S2`, 
                      `1332-11-S3` = S3_13_3_with_snp_names$`1332-11-S3`, `1332-111-S4` = S4_13_3_with_snp_names$`1332-111-S4`, check.names = F)


gt_13_4 <- data.frame(SNP = F_13_4_with_snp_names$rowname, `13_4-F` = F_13_4_with_snp_names$`13_4-F`, 
                      `1345-S1` = S1_13_4_with_snp_names$`1345-S1`, `1345-5-S2` = S2_13_4_with_snp_names$`1345-5-S2`, 
                      `1345-54-S3` = S3_13_4_with_snp_names$`1345-54-S3`, `1345-544-S4` = S4_13_4_with_snp_names$`1345-544-S4`, check.names = F)


gt_16_1 <- data.frame(SNP = F_16_1_with_snp_names$rowname, `16_1-F` = F_16_1_with_snp_names$`16_1-F`, 
                      `1611-S1` = S1_16_1_with_snp_names$`1611-S1`, `1611-2-S2` = S2_16_1_with_snp_names$`1611-2-S2`, 
                      `1611-21-S3` = S3_16_1_with_snp_names$`1611-21-S3`, `1611-211-S4` = S4_16_1_with_snp_names$`1611-211-S4`, check.names = F)


gt_16_5 <- data.frame(SNP = F_16_5_with_snp_names$rowname, `16_5-F` = F_16_5_with_snp_names$`16_5-F`, 
                      `1654-S1` = S1_16_5_with_snp_names$`1654-S1`, `1654-4-S2` = S2_16_5_with_snp_names$`1654-4-S2`, 
                      `1654-45-S3` = S3_16_5_with_snp_names$`1654-45-S3`, `1654-454-S4` = S4_16_5_with_snp_names$`1654-454-S4`, 
                      `1654-454-S5` = S5_16_5_with_snp_names$`1654-454-S5`, check.names = F)


gt_17_2 <- data.frame(SNP = F_17_2_with_snp_names$rowname, `17_2-F` = F_17_2_with_snp_names$`17_2-F`, 
                      `1721-S1` = S1_17_2_with_snp_names$`1721-S1`, `1721-1-S2` = S2_17_2_with_snp_names$`1721-1-S2`, 
                      `1721-11-S3` = S3_17_2_with_snp_names$`1721-11-S3`, `1721-111-S4` = S4_17_2_with_snp_names$`1721-111-S4`, 
                      `1721-111-S5` = S5_17_2_with_snp_names$`1721-111-S5`, check.names = F)


gt_17_5 <- data.frame(SNP = F_17_5_with_snp_names$rowname, `17_5-F` = F_17_5_with_snp_names$`17_5-F`, 
                      `1755-S1` = S1_17_5_with_snp_names$`1755-S1`, `1755-5-S2` = S2_17_5_with_snp_names$`1755-5-S2`, 
                      `1755-54-S3` = S3_17_5_with_snp_names$`1755-54-S3`, `1755-545-S4` = S4_17_5_with_snp_names$`1755-545-S4`, 
                      `1755-545-S5` = S5_17_5_with_snp_names$`1755-545-S5`, check.names = F)


gt_19_2 <- data.frame(SNP = F_19_2_with_snp_names$rowname, `19_2-F` = F_19_2_with_snp_names$`19_2-F`, 
                      `1922-S1` = S1_19_2_with_snp_names$`1922-S1`, `1922-1-S2` = S2_19_2_with_snp_names$`1922-1-S2`, 
                      `1922-11-S3` = S3_19_2_with_snp_names$`1922-11-S3`, `1922-111-S4` = S4_19_2_with_snp_names$`1922-111-S4`, check.names = F)


gt_19_5 <- data.frame(SNP = F_19_5_with_snp_names$rowname, `19_5-F` = F_19_5_with_snp_names$`19_5-F`, 
                      `1955-S1` = S1_19_5_with_snp_names$`1955-S1`, `1955-5-S2` = S2_19_5_with_snp_names$`1955-5-S2`, 
                      `1955-54-S3` = S3_19_5_with_snp_names$`1955-54-S3`, `1955-544-S4` = S4_19_5_with_snp_names$`1955-544-S4`, check.names = F)


gt_20_1 <- data.frame(SNP = F_20_1_with_snp_names$rowname, `20_1-F` = F_20_1_with_snp_names$`20_1-F`, 
                      `2013-S1` = S1_20_1_with_snp_names$`2013-S1`, `2013-1-S2` = S2_20_1_with_snp_names$`2013-1-S2`, 
                      `2013-13-S3` = S3_20_1_with_snp_names$`2013-13-S3`, `2013-131-S4` = S4_20_1_with_snp_names$`2013-131-S4`, check.names = F)


gt_20_4 <- data.frame(SNP = F_20_4_with_snp_names$rowname, `20_4-F` = F_20_4_with_snp_names$`20_4-F`, 
                      `2045-S1` = S1_20_4_with_snp_names$`2045-S1`, `2045-5-S2` = S2_20_4_with_snp_names$`2045-5-S2`, 
                      `2045-54-S3` = S3_20_4_with_snp_names$`2045-54-S3`, `2045-544-S4` = S4_20_4_with_snp_names$`2045-544-S4`, check.names = F)


gt_21_2 <- data.frame(SNP = F_21_2_with_snp_names$rowname, `21_2-F` = F_21_2_with_snp_names$`21_2-F`, 
                      `2121-S1` = S1_21_2_with_snp_names$`2121-S1`, `2121-1-S2` = S2_21_2_with_snp_names$`2121-1-S2`, 
                      `2121-11-S3` = S3_21_2_with_snp_names$`2121-11-S3`, `2121-111-S4` = S4_21_2_with_snp_names$`2121-111-S4`, 
                      `2121-111-S5` = S5_21_2_with_snp_names$`2121-111-S5`, check.names = F)


gt_21_6 <- data.frame(SNP = F_21_6_with_snp_names$rowname, `21_6-F` = F_21_6_with_snp_names$`21_6-F`, 
                      `2165-S1` = S1_21_6_with_snp_names$`2165-S1`, `2165-4-S2` = S2_21_6_with_snp_names$`2165-4-S2`, 
                      `2165-44-S3` = S3_21_6_with_snp_names$`2165-44-S3`, `2165-444-S4` = S4_21_6_with_snp_names$`2165-444-S4`, 
                      `2165-444-S5` = S5_21_6_with_snp_names$`2165-444-S5`, check.names = F)


gt_23_2 <- data.frame(SNP = F_23_2_with_snp_names$rowname, `23_2-F` = F_23_2_with_snp_names$`23_2-F`, 
                      `2323-S1` = S1_23_2_with_snp_names$`2323-S1`, `2323-2-S2` = S2_23_2_with_snp_names$`2323-2-S2`, 
                      `2323-21-S3` = S3_23_2_with_snp_names$`2323-21-S3`, `2323-211-S4` = S4_23_2_with_snp_names$`2323-211-S4`, 
                      `2323-211-S5` = S5_23_2_with_snp_names$`2323-211-S5`, check.names = F)


gt_23_4 <- data.frame(SNP = F_23_4_with_snp_names$rowname, `23_4-F` = F_23_4_with_snp_names$`23_4-F`, 
                      `2344-S1` = S1_23_4_with_snp_names$`2344-S1`, `2344-4-S2` = S2_23_4_with_snp_names$`2344-4-S2`, 
                      `2344-46-S3` = S3_23_4_with_snp_names$`2344-46-S3`, `2344-464-S4` = S4_23_4_with_snp_names$`2344-464-S4`, check.names = F)


gt_26_1 <- data.frame(SNP = F_26_1_with_snp_names$rowname, `26_1-F` = F_26_1_with_snp_names$`26_1-F`, 
                      `2612-S1` = S1_26_1_with_snp_names$`2612-S1`, `2612-1-S2` = S2_26_1_with_snp_names$`2612-1-S2`, 
                      `2612-12-S3` = S3_26_1_with_snp_names$`2612-12-S3`, `2612-121-S4` = S4_26_1_with_snp_names$`2612-121-S4`, check.names = F)


gt_26_4 <- data.frame(SNP = F_26_4_with_snp_names$rowname, `26_4-F` = F_26_4_with_snp_names$`26_4-F`, 
                      `2645-S1` = S1_26_4_with_snp_names$`2645-S1`, `2645-4-S2` = S2_26_4_with_snp_names$`2645-4-S2`, 
                      `2645-46-S3` = S3_26_4_with_snp_names$`2645-46-S3`, `2645-464-S4` = S4_26_4_with_snp_names$`2645-464-S4`, check.names = F)


gt_27_2 <- data.frame(SNP = F_27_2_with_snp_names$rowname, `27_2-F` = F_27_2_with_snp_names$`27_2-F`, 
                      `2721-S1` = S1_27_2_with_snp_names$`2721-S1`, `2721-1-S2` = S2_27_2_with_snp_names$`2721-1-S2`, 
                      `2721-12-S3` = S3_27_2_with_snp_names$`2721-12-S3`, `2721-121-S4` = S4_27_2_with_snp_names$`2721-121-S4`, check.names = F)


gt_29_2 <- data.frame(SNP = F_29_2_with_snp_names$rowname, `29_2-F` = F_29_2_with_snp_names$`29_2-F`, 
                      `2923-S1` = S1_29_2_with_snp_names$`2923-S1`, `2923-1-S2` = S2_29_2_with_snp_names$`2923-1-S2`, 
                      `2923-12-S3` = S3_29_2_with_snp_names$`2923-12-S3`, `2923-121-S4` = S4_29_2_with_snp_names$`2923-121-S4`, 
                      `2923-121-S5` = S5_29_2_with_snp_names$`2923-121-S5`, check.names = F)


gt_29_4 <- data.frame(SNP = F_29_4_with_snp_names$rowname, `29_4-F` = F_29_4_with_snp_names$`29_4-F`, 
                      `2944-S1` = S1_29_4_with_snp_names$`2944-S1`, `2944-5-S2` = S2_29_4_with_snp_names$`2944-5-S2`, 
                      `2944-56-S3` = S3_29_4_with_snp_names$`2944-56-S3`, `2944-565-S4` = S4_29_4_with_snp_names$`2944-565-S4`, 
                      `2944-565-S5` = S5_29_4_with_snp_names$`2944-565-S5`, check.names = F)


# Convert SNP to character
gt_1_1$SNP <- as.character(gt_1_1$SNP)
gt_6_1$SNP <- as.character(gt_6_1$SNP)
gt_6_4$SNP <- as.character(gt_6_4$SNP)
gt_7_2$SNP <- as.character(gt_7_2$SNP)
gt_7_4$SNP <- as.character(gt_7_4$SNP)
gt_8_2$SNP <- as.character(gt_8_2$SNP)
gt_8_4$SNP <- as.character(gt_8_4$SNP)
gt_10_2$SNP <- as.character(gt_10_2$SNP)
gt_10_5$SNP <- as.character(gt_10_5$SNP)
gt_13_3$SNP <- as.character(gt_13_3$SNP)
gt_13_4$SNP <- as.character(gt_13_4$SNP)
gt_16_1$SNP <- as.character(gt_16_1$SNP)
gt_16_5$SNP <- as.character(gt_16_5$SNP)
gt_17_2$SNP <- as.character(gt_17_2$SNP)
gt_17_5$SNP <- as.character(gt_17_5$SNP)
gt_19_2$SNP <- as.character(gt_19_2$SNP)
gt_19_5$SNP <- as.character(gt_19_5$SNP)
gt_20_1$SNP <- as.character(gt_20_1$SNP)
gt_20_4$SNP <- as.character(gt_20_4$SNP)
gt_21_2$SNP <- as.character(gt_21_2$SNP)
gt_21_6$SNP <- as.character(gt_21_6$SNP)
gt_23_2$SNP <- as.character(gt_23_2$SNP)
gt_23_4$SNP <- as.character(gt_23_4$SNP)
gt_26_1$SNP <- as.character(gt_26_1$SNP)
gt_26_4$SNP <- as.character(gt_26_4$SNP)
gt_27_2$SNP <- as.character(gt_27_2$SNP)
gt_29_2$SNP <- as.character(gt_29_2$SNP)
gt_29_4$SNP <- as.character(gt_29_4$SNP)

# Get fixed ref, alt, not fixed
# 1_1
gt_1_1_fixed_ref <- gt_1_1 %>% 
  mutate(fixed = if_else((`1_1-F` == 0 | `111-S1` == 0 | `111-2-S2` == 0 | `111-22-S3` == 0 | 
                            `111-221-S4` == 0 | `111-221-S5` == 0), "fixed_ref", 
                         if_else((`1_1-F` == 2 | `111-S1` == 2 | `111-2-S2` == 2 | `111-22-S3` == 2 | 
                                    `111-221-S4` == 2 | `111-221-S5` == 2), "fixed_alt", "not_fixed"))) %>% 
  filter(fixed == "fixed_ref")

gt_1_1_fixed_alt <- gt_1_1 %>% 
  mutate(fixed = if_else((`1_1-F` == 0 | `111-S1` == 0 | `111-2-S2` == 0 | `111-22-S3` == 0 | 
                            `111-221-S4` == 0 | `111-221-S5` == 0), "fixed_ref", 
                         if_else((`1_1-F` == 2 | `111-S1` == 2 | `111-2-S2` == 2 | `111-22-S3` == 2 | 
                                    `111-221-S4` == 2 | `111-221-S5` == 2), "fixed_alt", "not_fixed"))) %>% 
  filter(fixed == "fixed_alt")

gt_1_1_not_fixed <- gt_1_1 %>% 
  mutate(fixed = if_else((`1_1-F` == 0 | `111-S1` == 0 | `111-2-S2` == 0 | `111-22-S3` == 0 | 
                            `111-221-S4` == 0 | `111-221-S5` == 0), "fixed_ref", 
                         if_else((`1_1-F` == 2 | `111-S1` == 2 | `111-2-S2` == 2 | `111-22-S3` == 2 | 
                                    `111-221-S4` == 2 | `111-221-S5` == 2), "fixed_alt", "not_fixed"))) %>% 
  filter(fixed == "not_fixed")

# 6_1
gt_6_1_fixed_ref <- gt_6_1 %>% 
  mutate(fixed = if_else((`6_1-F` == 0 | `611-S1` == 0 | `611-1-S2` == 0 | `611-11-S3` == 0 | 
                            `611-111-S4` == 0), "fixed_ref", 
                         if_else((`6_1-F` == 2 | `611-S1` == 2 | `611-1-S2` == 2 | `611-11-S3` == 2 | 
                                    `611-111-S4` == 2), "fixed_alt", "not_fixed"))) %>% 
  filter(fixed == "fixed_ref")

gt_6_1_fixed_alt <- gt_6_1 %>% 
  mutate(fixed = if_else((`6_1-F` == 0 | `611-S1` == 0 | `611-1-S2` == 0 | `611-11-S3` == 0 | 
                            `611-111-S4` == 0), "fixed_ref", 
                         if_else((`6_1-F` == 2 | `611-S1` == 2 | `611-1-S2` == 2 | `611-11-S3` == 2 | 
                                    `611-111-S4` == 2), "fixed_alt", "not_fixed"))) %>% 
  filter(fixed == "fixed_alt")

gt_6_1_not_fixed <- gt_6_1 %>% 
  mutate(fixed = if_else((`6_1-F` == 0 | `611-S1` == 0 | `611-1-S2` == 0 | `611-11-S3` == 0 | 
                            `611-111-S4` == 0), "fixed_ref", 
                         if_else((`6_1-F` == 2 | `611-S1` == 2 | `611-1-S2` == 2 | `611-11-S3` == 2 | 
                                    `611-111-S4` == 2), "fixed_alt", "not_fixed"))) %>% 
  filter(fixed == "not_fixed")

# 6_4
gt_6_4_fixed_ref <- gt_6_4 %>% 
  mutate(fixed = if_else((`6_4-F` == 0 | `646-S1` == 0 | `646-4-S2` == 0 | `646-45-S3` == 0 | 
                            `646-454-S4` == 0 | `646-454-S5` == 0), "fixed_ref", 
                         if_else((`6_4-F` == 2 | `646-S1` == 2 | `646-4-S2` == 2 | `646-45-S3` == 2 | 
                                    `646-454-S4` == 2 | `646-454-S4` == 2), "fixed_alt", "not_fixed"))) %>% 
  filter(fixed == "fixed_ref")

gt_6_4_fixed_alt <- gt_6_4 %>% 
  mutate(fixed = if_else((`6_4-F` == 0 | `646-S1` == 0 | `646-4-S2` == 0 | `646-45-S3` == 0 | 
                            `646-454-S4` == 0 | `646-454-S5` == 0), "fixed_ref", 
                         if_else((`6_4-F` == 2 | `646-S1` == 2 | `646-4-S2` == 2 | `646-45-S3` == 2 | 
                                    `646-454-S4` == 2 | `646-454-S4` == 2), "fixed_alt", "not_fixed"))) %>% 
  filter(fixed == "fixed_alt")

gt_6_4_not_fixed <- gt_6_4 %>% 
  mutate(fixed = if_else((`6_4-F` == 0 | `646-S1` == 0 | `646-4-S2` == 0 | `646-45-S3` == 0 | 
                            `646-454-S4` == 0 | `646-454-S5` == 0), "fixed_ref", 
                         if_else((`6_4-F` == 2 | `646-S1` == 2 | `646-4-S2` == 2 | `646-45-S3` == 2 | 
                                    `646-454-S4` == 2 | `646-454-S4` == 2), "fixed_alt", "not_fixed"))) %>% 
  filter(fixed == "not_fixed")

# 7_2
gt_7_2_fixed_ref <- gt_7_2 %>% 
  mutate(fixed = if_else((`7_2-F` == 0 | `721-S1` == 0 | `721-2-S2` == 0 | `721-21-S3` == 0 | 
                            `721-211-S4` == 0), "fixed_ref", 
                         if_else((`7_2-F` == 2 | `721-S1` == 2 | `721-2-S2` == 2 | `721-21-S3` == 2 | 
                                    `721-211-S4` == 2), "fixed_alt", "not_fixed"))) %>% 
  filter(fixed == "fixed_ref")

gt_7_2_fixed_alt <- gt_7_2 %>% 
  mutate(fixed = if_else((`7_2-F` == 0 | `721-S1` == 0 | `721-2-S2` == 0 | `721-21-S3` == 0 | 
                            `721-211-S4` == 0), "fixed_ref", 
                         if_else((`7_2-F` == 2 | `721-S1` == 2 | `721-2-S2` == 2 | `721-21-S3` == 2 | 
                                    `721-211-S4` == 2), "fixed_alt", "not_fixed"))) %>% 
  filter(fixed == "fixed_alt")

gt_7_2_not_fixed <- gt_7_2 %>% 
  mutate(fixed = if_else((`7_2-F` == 0 | `721-S1` == 0 | `721-2-S2` == 0 | `721-21-S3` == 0 | 
                            `721-211-S4` == 0), "fixed_ref", 
                         if_else((`7_2-F` == 2 | `721-S1` == 2 | `721-2-S2` == 2 | `721-21-S3` == 2 | 
                                    `721-211-S4` == 2), "fixed_alt", "not_fixed"))) %>% 
  filter(fixed == "not_fixed")

# 7_4
gt_7_4_fixed_ref <- gt_7_4 %>% 
  mutate(fixed = if_else((`7_4-F` == 0 | `745-S1` == 0 | `745-4-S2` == 0 | `745-44-S3` == 0 | 
                            `745-444-S4` == 0), "fixed_ref", 
                         if_else((`7_4-F` == 2 | `745-S1` == 2 | `745-4-S2` == 2 | `745-44-S3` == 2 | 
                                    `745-444-S4` == 2), "fixed_alt", "not_fixed"))) %>% 
  filter(fixed == "fixed_ref")

gt_7_4_fixed_alt <- gt_7_4 %>% 
  mutate(fixed = if_else((`7_4-F` == 0 | `745-S1` == 0 | `745-4-S2` == 0 | `745-44-S3` == 0 | 
                            `745-444-S4` == 0), "fixed_ref", 
                         if_else((`7_4-F` == 2 | `745-S1` == 2 | `745-4-S2` == 2 | `745-44-S3` == 2 | 
                                    `745-444-S4` == 2), "fixed_alt", "not_fixed"))) %>% 
  filter(fixed == "fixed_alt")

gt_7_4_not_fixed <- gt_7_4 %>% 
  mutate(fixed = if_else((`7_4-F` == 0 | `745-S1` == 0 | `745-4-S2` == 0 | `745-44-S3` == 0 | 
                            `745-444-S4` == 0), "fixed_ref", 
                         if_else((`7_4-F` == 2 | `745-S1` == 2 | `745-4-S2` == 2 | `745-44-S3` == 2 | 
                                    `745-444-S4` == 2), "fixed_alt", "not_fixed"))) %>% 
  filter(fixed == "not_fixed")

# 8_2
gt_8_2_fixed_ref <- gt_8_2 %>% 
  mutate(fixed = if_else((`8_2-F` == 0 | `821-S1` == 0 | `821-1-S2` == 0 | `821-11-S3` == 0 | 
                            `821-111-S4` == 0), "fixed_ref", 
                         if_else((`8_2-F` == 2 | `821-S1` == 2 | `821-1-S2` == 2 | `821-11-S3` == 2 | 
                                    `821-111-S4` == 2), "fixed_alt", "not_fixed"))) %>% 
  filter(fixed == "fixed_ref")

gt_8_2_fixed_alt <- gt_8_2 %>% 
  mutate(fixed = if_else((`8_2-F` == 0 | `821-S1` == 0 | `821-1-S2` == 0 | `821-11-S3` == 0 | 
                            `821-111-S4` == 0), "fixed_ref", 
                         if_else((`8_2-F` == 2 | `821-S1` == 2 | `821-1-S2` == 2 | `821-11-S3` == 2 | 
                                    `821-111-S4` == 2), "fixed_alt", "not_fixed"))) %>% 
  filter(fixed == "fixed_alt")

gt_8_2_not_fixed <- gt_8_2 %>% 
  mutate(fixed = if_else((`8_2-F` == 0 | `821-S1` == 0 | `821-1-S2` == 0 | `821-11-S3` == 0 | 
                            `821-111-S4` == 0), "fixed_ref", 
                         if_else((`8_2-F` == 2 | `821-S1` == 2 | `821-1-S2` == 2 | `821-11-S3` == 2 | 
                                    `821-111-S4` == 2), "fixed_alt", "not_fixed"))) %>% 
  filter(fixed == "not_fixed")

# 8_4
gt_8_4_fixed_ref <- gt_8_4 %>% 
  mutate(fixed = if_else((`8_4-F` == 0 | `845-S1` == 0 | `845-4-S2` == 0 | `845-44-S3` == 0 | 
                            `845-444-S4` == 0), "fixed_ref", 
                         if_else((`8_4-F` == 2 | `845-S1` == 2 | `845-4-S2` == 2 | `845-44-S3` == 2 | 
                                    `845-444-S4` == 2), "fixed_alt", "not_fixed"))) %>% 
  filter(fixed == "fixed_ref")

gt_8_4_fixed_alt <- gt_8_4 %>% 
  mutate(fixed = if_else((`8_4-F` == 0 | `845-S1` == 0 | `845-4-S2` == 0 | `845-44-S3` == 0 | 
                            `845-444-S4` == 0), "fixed_ref", 
                         if_else((`8_4-F` == 2 | `845-S1` == 2 | `845-4-S2` == 2 | `845-44-S3` == 2 | 
                                    `845-444-S4` == 2), "fixed_alt", "not_fixed"))) %>% 
  filter(fixed == "fixed_alt")

gt_8_4_not_fixed <- gt_8_4 %>% 
  mutate(fixed = if_else((`8_4-F` == 0 | `845-S1` == 0 | `845-4-S2` == 0 | `845-44-S3` == 0 | 
                            `845-444-S4` == 0), "fixed_ref", 
                         if_else((`8_4-F` == 2 | `845-S1` == 2 | `845-4-S2` == 2 | `845-44-S3` == 2 | 
                                    `845-444-S4` == 2), "fixed_alt", "not_fixed"))) %>% 
  filter(fixed == "not_fixed")

# 10_2
gt_10_2_fixed_ref <- gt_10_2 %>% 
  mutate(fixed = if_else((`10_2-F` == 0 | `1022-S1` == 0 | `1022-1-S2` == 0 | `1022-11-S3` == 0 | 
                            `1022-113-S4` == 0 | `1022-113-S5` == 0), "fixed_ref", 
                         if_else((`10_2-F` == 2 | `1022-S1` == 2 | `1022-1-S2` == 2 | `1022-11-S3` == 2 | 
                                    `1022-113-S4` == 2 | `1022-113-S5` == 2), "fixed_alt", "not_fixed"))) %>% 
  filter(fixed == "fixed_ref")

gt_10_2_fixed_alt <- gt_10_2 %>% 
  mutate(fixed = if_else((`10_2-F` == 0 | `1022-S1` == 0 | `1022-1-S2` == 0 | `1022-11-S3` == 0 | 
                            `1022-113-S4` == 0 | `1022-113-S5` == 0), "fixed_ref", 
                         if_else((`10_2-F` == 2 | `1022-S1` == 2 | `1022-1-S2` == 2 | `1022-11-S3` == 2 | 
                                    `1022-113-S4` == 2 | `1022-113-S5` == 2), "fixed_alt", "not_fixed"))) %>% 
  filter(fixed == "fixed_alt")

gt_10_2_not_fixed <- gt_10_2 %>% 
  mutate(fixed = if_else((`10_2-F` == 0 | `1022-S1` == 0 | `1022-1-S2` == 0 | `1022-11-S3` == 0 | 
                            `1022-113-S4` == 0 | `1022-113-S5` == 0), "fixed_ref", 
                         if_else((`10_2-F` == 2 | `1022-S1` == 2 | `1022-1-S2` == 2 | `1022-11-S3` == 2 | 
                                    `1022-113-S4` == 2 | `1022-113-S5` == 2), "fixed_alt", "not_fixed"))) %>%
  filter(fixed == "not_fixed")

# 10_5
gt_10_5_fixed_ref <- gt_10_5 %>% 
  mutate(fixed = if_else((`10_5-F` == 0 | `1056-S1` == 0 | `1056-4-S2` == 0 | `1056-45-S3` == 0 | 
                            `1056-454-S4` == 0), "fixed_ref", 
                         if_else((`10_5-F` == 2 | `1056-S1` == 2 | `1056-4-S2` == 2 | `1056-45-S3` == 2 | 
                                    `1056-454-S4` == 2), "fixed_alt", "not_fixed"))) %>% 
  filter(fixed == "fixed_ref")

gt_10_5_fixed_alt <- gt_10_5 %>% 
  mutate(fixed = if_else((`10_5-F` == 0 | `1056-S1` == 0 | `1056-4-S2` == 0 | `1056-45-S3` == 0 | 
                            `1056-454-S4` == 0), "fixed_ref", 
                         if_else((`10_5-F` == 2 | `1056-S1` == 2 | `1056-4-S2` == 2 | `1056-45-S3` == 2 | 
                                    `1056-454-S4` == 2), "fixed_alt", "not_fixed"))) %>% 
  filter(fixed == "fixed_alt")

gt_10_5_not_fixed <- gt_10_5 %>% 
  mutate(fixed = if_else((`10_5-F` == 0 | `1056-S1` == 0 | `1056-4-S2` == 0 | `1056-45-S3` == 0 | 
                            `1056-454-S4` == 0), "fixed_ref", 
                         if_else((`10_5-F` == 2 | `1056-S1` == 2 | `1056-4-S2` == 2 | `1056-45-S3` == 2 | 
                                    `1056-454-S4` == 2), "fixed_alt", "not_fixed"))) %>% 
  filter(fixed == "not_fixed")

# 13_3
gt_13_3_fixed_ref <- gt_13_3 %>% 
  mutate(fixed = if_else((`13_3-F` == 0 | `1332-S1` == 0 | `1332-1-S2` == 0 | `1332-11-S3` == 0 | 
                            `1332-111-S4` == 0), "fixed_ref", 
                         if_else((`13_3-F` == 2 | `1332-S1` == 2 | `1332-1-S2` == 2 | `1332-11-S3` == 2 | 
                                    `1332-111-S4` == 2), "fixed_alt", "not_fixed"))) %>% 
  filter(fixed == "fixed_ref")

gt_13_3_fixed_alt <- gt_13_3 %>% 
  mutate(fixed = if_else((`13_3-F` == 0 | `1332-S1` == 0 | `1332-1-S2` == 0 | `1332-11-S3` == 0 | 
                            `1332-111-S4` == 0), "fixed_ref", 
                         if_else((`13_3-F` == 2 | `1332-S1` == 2 | `1332-1-S2` == 2 | `1332-11-S3` == 2 | 
                                    `1332-111-S4` == 2), "fixed_alt", "not_fixed"))) %>% 
  filter(fixed == "fixed_alt")

gt_13_3_not_fixed <- gt_13_3 %>% 
  mutate(fixed = if_else((`13_3-F` == 0 | `1332-S1` == 0 | `1332-1-S2` == 0 | `1332-11-S3` == 0 | 
                            `1332-111-S4` == 0), "fixed_ref", 
                         if_else((`13_3-F` == 2 | `1332-S1` == 2 | `1332-1-S2` == 2 | `1332-11-S3` == 2 | 
                                    `1332-111-S4` == 2), "fixed_alt", "not_fixed"))) %>% 
  filter(fixed == "not_fixed")

# 13_4
gt_13_4_fixed_ref <- gt_13_4 %>% 
  mutate(fixed = if_else((`13_4-F` == 0 | `1345-S1` == 0 | `1345-5-S2` == 0 | `1345-54-S3` == 0 | 
                            `1345-544-S4` == 0), "fixed_ref", 
                         if_else((`13_4-F` == 2 | `1345-S1` == 2 | `1345-5-S2` == 2 | `1345-54-S3` == 2 | 
                                    `1345-544-S4` == 2), "fixed_alt", "not_fixed"))) %>% 
  filter(fixed == "fixed_ref")

gt_13_4_fixed_alt <- gt_13_4 %>% 
  mutate(fixed = if_else((`13_4-F` == 0 | `1345-S1` == 0 | `1345-5-S2` == 0 | `1345-54-S3` == 0 | 
                            `1345-544-S4` == 0), "fixed_ref", 
                         if_else((`13_4-F` == 2 | `1345-S1` == 2 | `1345-5-S2` == 2 | `1345-54-S3` == 2 | 
                                    `1345-544-S4` == 2), "fixed_alt", "not_fixed"))) %>% 
  filter(fixed == "fixed_alt")

gt_13_4_not_fixed <- gt_13_4 %>% 
  mutate(fixed = if_else((`13_4-F` == 0 | `1345-S1` == 0 | `1345-5-S2` == 0 | `1345-54-S3` == 0 | 
                            `1345-544-S4` == 0), "fixed_ref", 
                         if_else((`13_4-F` == 2 | `1345-S1` == 2 | `1345-5-S2` == 2 | `1345-54-S3` == 2 | 
                                    `1345-544-S4` == 2), "fixed_alt", "not_fixed"))) %>% 
  filter(fixed == "not_fixed")

# 16_1
gt_16_1_fixed_ref <- gt_16_1 %>% 
  mutate(fixed = if_else((`16_1-F` == 0 | `1611-S1` == 0 | `1611-2-S2` == 0 | `1611-21-S3` == 0 | 
                            `1611-211-S4` == 0), "fixed_ref", 
                         if_else((`16_1-F` == 2 | `1611-S1` == 2 | `1611-2-S2` == 2 | `1611-21-S3` == 2 | 
                                    `1611-211-S4` == 2), "fixed_alt", "not_fixed"))) %>% 
  filter(fixed == "fixed_ref")

gt_16_1_fixed_alt <- gt_16_1 %>% 
  mutate(fixed = if_else((`16_1-F` == 0 | `1611-S1` == 0 | `1611-2-S2` == 0 | `1611-21-S3` == 0 | 
                            `1611-211-S4` == 0), "fixed_ref", 
                         if_else((`16_1-F` == 2 | `1611-S1` == 2 | `1611-2-S2` == 2 | `1611-21-S3` == 2 | 
                                    `1611-211-S4` == 2), "fixed_alt", "not_fixed"))) %>% 
  filter(fixed == "fixed_alt")

gt_16_1_not_fixed <- gt_16_1 %>% 
  mutate(fixed = if_else((`16_1-F` == 0 | `1611-S1` == 0 | `1611-2-S2` == 0 | `1611-21-S3` == 0 | 
                            `1611-211-S4` == 0), "fixed_ref", 
                         if_else((`16_1-F` == 2 | `1611-S1` == 2 | `1611-2-S2` == 2 | `1611-21-S3` == 2 | 
                                    `1611-211-S4` == 2), "fixed_alt", "not_fixed"))) %>% 
  filter(fixed == "not_fixed")

# 16_5
gt_16_5_fixed_ref <- gt_16_5 %>% 
  mutate(fixed = if_else((`16_5-F` == 0 | `1654-S1` == 0 | `1654-4-S2` == 0 | `1654-45-S3` == 0 | 
                            `1654-454-S4` == 0 | `1654-454-S5` == 0), "fixed_ref", 
                         if_else((`16_5-F` == 2 | `1654-S1` == 2 | `1654-4-S2` == 2 | `1654-45-S3` == 2 | 
                                    `1654-454-S4` == 2 | `1654-454-S5` == 2), "fixed_alt", "not_fixed"))) %>% 
  filter(fixed == "fixed_ref")

gt_16_5_fixed_alt <- gt_16_5 %>% 
  mutate(fixed = if_else((`16_5-F` == 0 | `1654-S1` == 0 | `1654-4-S2` == 0 | `1654-45-S3` == 0 | 
                            `1654-454-S4` == 0 | `1654-454-S5` == 0), "fixed_ref", 
                         if_else((`16_5-F` == 2 | `1654-S1` == 2 | `1654-4-S2` == 2 | `1654-45-S3` == 2 | 
                                    `1654-454-S4` == 2 | `1654-454-S5` == 2), "fixed_alt", "not_fixed"))) %>% 
  filter(fixed == "fixed_alt")

gt_16_5_not_fixed <- gt_16_5 %>% 
  mutate(fixed = if_else((`16_5-F` == 0 | `1654-S1` == 0 | `1654-4-S2` == 0 | `1654-45-S3` == 0 | 
                            `1654-454-S4` == 0 | `1654-454-S5` == 0), "fixed_ref", 
                         if_else((`16_5-F` == 2 | `1654-S1` == 2 | `1654-4-S2` == 2 | `1654-45-S3` == 2 | 
                                    `1654-454-S4` == 2 | `1654-454-S5` == 2), "fixed_alt", "not_fixed"))) %>%
  filter(fixed == "not_fixed")

# 17_2
gt_17_2_fixed_ref <- gt_17_2 %>% 
  mutate(fixed = if_else((`17_2-F` == 0 | `1721-S1` == 0 | `1721-1-S2` == 0 | `1721-11-S3` == 0 | 
                            `1721-111-S4` == 0 | `1721-111-S5` == 0), "fixed_ref", 
                         if_else((`17_2-F` == 2 | `1721-S1` == 2 | `1721-1-S2` == 2 | `1721-11-S3` == 2 | 
                                    `1721-111-S4` == 2 | `1721-111-S5` == 2), "fixed_alt", "not_fixed"))) %>% 
  filter(fixed == "fixed_ref")

gt_17_2_fixed_alt <- gt_17_2 %>% 
  mutate(fixed = if_else((`17_2-F` == 0 | `1721-S1` == 0 | `1721-1-S2` == 0 | `1721-11-S3` == 0 | 
                            `1721-111-S4` == 0 | `1721-111-S5` == 0), "fixed_ref", 
                         if_else((`17_2-F` == 2 | `1721-S1` == 2 | `1721-1-S2` == 2 | `1721-11-S3` == 2 | 
                                    `1721-111-S4` == 2 | `1721-111-S5` == 2), "fixed_alt", "not_fixed"))) %>% 
  filter(fixed == "fixed_alt")

gt_17_2_not_fixed <- gt_17_2 %>% 
  mutate(fixed = if_else((`17_2-F` == 0 | `1721-S1` == 0 | `1721-1-S2` == 0 | `1721-11-S3` == 0 | 
                            `1721-111-S4` == 0 | `1721-111-S5` == 0), "fixed_ref", 
                         if_else((`17_2-F` == 2 | `1721-S1` == 2 | `1721-1-S2` == 2 | `1721-11-S3` == 2 | 
                                    `1721-111-S4` == 2 | `1721-111-S5` == 2), "fixed_alt", "not_fixed"))) %>% 
  filter(fixed == "not_fixed")

# 17_5
gt_17_5_fixed_ref <- gt_17_5 %>% 
  mutate(fixed = if_else((`17_5-F` == 0 | `1755-S1` == 0 | `1755-5-S2` == 0 | `1755-54-S3` == 0 | 
                            `1755-545-S4` == 0 | `1755-545-S5` == 0), "fixed_ref", 
                         if_else((`17_5-F` == 2 | `1755-S1` == 2 | `1755-5-S2` == 2 | `1755-54-S3` == 2 | 
                                    `1755-545-S4` == 2 | `1755-545-S5` == 2), "fixed_alt", "not_fixed"))) %>% 
  filter(fixed == "fixed_ref")

gt_17_5_fixed_alt <- gt_17_5 %>% 
  mutate(fixed = if_else((`17_5-F` == 0 | `1755-S1` == 0 | `1755-5-S2` == 0 | `1755-54-S3` == 0 | 
                            `1755-545-S4` == 0 | `1755-545-S5` == 0), "fixed_ref", 
                         if_else((`17_5-F` == 2 | `1755-S1` == 2 | `1755-5-S2` == 2 | `1755-54-S3` == 2 | 
                                    `1755-545-S4` == 2 | `1755-545-S5` == 2), "fixed_alt", "not_fixed"))) %>% 
  filter(fixed == "fixed_alt")

gt_17_5_not_fixed <- gt_17_5 %>% 
  mutate(fixed = if_else((`17_5-F` == 0 | `1755-S1` == 0 | `1755-5-S2` == 0 | `1755-54-S3` == 0 | 
                            `1755-545-S4` == 0 | `1755-545-S5` == 0), "fixed_ref", 
                         if_else((`17_5-F` == 2 | `1755-S1` == 2 | `1755-5-S2` == 2 | `1755-54-S3` == 2 | 
                                    `1755-545-S4` == 2 | `1755-545-S5` == 2), "fixed_alt", "not_fixed"))) %>% 
  filter(fixed == "not_fixed")

# 19_2
gt_19_2_fixed_ref <- gt_19_2 %>% 
  mutate(fixed = if_else((`19_2-F` == 0 | `1922-S1` == 0 | `1922-1-S2` == 0 | `1922-11-S3` == 0 | 
                            `1922-111-S4` == 0), "fixed_ref", 
                         if_else((`19_2-F` == 2 | `1922-S1` == 2 | `1922-1-S2` == 2 | `1922-11-S3` == 2 | 
                                    `1922-111-S4` == 2), "fixed_alt", "not_fixed"))) %>% 
  filter(fixed == "fixed_ref")

gt_19_2_fixed_alt <- gt_19_2 %>% 
  mutate(fixed = if_else((`19_2-F` == 0 | `1922-S1` == 0 | `1922-1-S2` == 0 | `1922-11-S3` == 0 | 
                            `1922-111-S4` == 0), "fixed_ref", 
                         if_else((`19_2-F` == 2 | `1922-S1` == 2 | `1922-1-S2` == 2 | `1922-11-S3` == 2 | 
                                    `1922-111-S4` == 2), "fixed_alt", "not_fixed"))) %>% 
  filter(fixed == "fixed_alt")

gt_19_2_not_fixed <- gt_19_2 %>% 
  mutate(fixed = if_else((`19_2-F` == 0 | `1922-S1` == 0 | `1922-1-S2` == 0 | `1922-11-S3` == 0 | 
                            `1922-111-S4` == 0), "fixed_ref", 
                         if_else((`19_2-F` == 2 | `1922-S1` == 2 | `1922-1-S2` == 2 | `1922-11-S3` == 2 | 
                                    `1922-111-S4` == 2), "fixed_alt", "not_fixed"))) %>% 
  filter(fixed == "not_fixed")

# 19_5
gt_19_5_fixed_ref <- gt_19_5 %>% 
  mutate(fixed = if_else((`19_5-F` == 0 | `1955-S1` == 0 | `1955-5-S2` == 0 | `1955-54-S3` == 0 | 
                            `1955-544-S4` == 0), "fixed_ref", 
                         if_else((`19_5-F` == 2 | `1955-S1` == 2 | `1955-5-S2` == 2 | `1955-54-S3` == 2 | 
                                    `1955-544-S4` == 2), "fixed_alt", "not_fixed"))) %>% 
  filter(fixed == "fixed_ref")

gt_19_5_fixed_alt <- gt_19_5 %>% 
  mutate(fixed = if_else((`19_5-F` == 0 | `1955-S1` == 0 | `1955-5-S2` == 0 | `1955-54-S3` == 0 | 
                            `1955-544-S4` == 0), "fixed_ref", 
                         if_else((`19_5-F` == 2 | `1955-S1` == 2 | `1955-5-S2` == 2 | `1955-54-S3` == 2 | 
                                    `1955-544-S4` == 2), "fixed_alt", "not_fixed"))) %>% 
  filter(fixed == "fixed_alt")

gt_19_5_not_fixed <- gt_19_5 %>% 
  mutate(fixed = if_else((`19_5-F` == 0 | `1955-S1` == 0 | `1955-5-S2` == 0 | `1955-54-S3` == 0 | 
                            `1955-544-S4` == 0), "fixed_ref", 
                         if_else((`19_5-F` == 2 | `1955-S1` == 2 | `1955-5-S2` == 2 | `1955-54-S3` == 2 | 
                                    `1955-544-S4` == 2), "fixed_alt", "not_fixed"))) %>% 
  filter(fixed == "not_fixed")

# 20_1
gt_20_1_fixed_ref <- gt_20_1 %>% 
  mutate(fixed = if_else((`20_1-F` == 0 | `2013-S1` == 0 | `2013-1-S2` == 0 | `2013-13-S3` == 0 | 
                            `2013-131-S4` == 0), "fixed_ref", 
                         if_else((`20_1-F` == 2 | `2013-S1` == 2 | `2013-1-S2` == 2 | `2013-13-S3` == 2 | 
                                    `2013-131-S4` == 2), "fixed_alt", "not_fixed"))) %>% 
  filter(fixed == "fixed_ref")

gt_20_1_fixed_alt <- gt_20_1 %>% 
  mutate(fixed = if_else((`20_1-F` == 0 | `2013-S1` == 0 | `2013-1-S2` == 0 | `2013-13-S3` == 0 | 
                            `2013-131-S4` == 0), "fixed_ref", 
                         if_else((`20_1-F` == 2 | `2013-S1` == 2 | `2013-1-S2` == 2 | `2013-13-S3` == 2 | 
                                    `2013-131-S4` == 2), "fixed_alt", "not_fixed"))) %>% 
  filter(fixed == "fixed_alt")

gt_20_1_not_fixed <- gt_20_1 %>% 
  mutate(fixed = if_else((`20_1-F` == 0 | `2013-S1` == 0 | `2013-1-S2` == 0 | `2013-13-S3` == 0 | 
                            `2013-131-S4` == 0), "fixed_ref", 
                         if_else((`20_1-F` == 2 | `2013-S1` == 2 | `2013-1-S2` == 2 | `2013-13-S3` == 2 | 
                                    `2013-131-S4` == 2), "fixed_alt", "not_fixed"))) %>% 
  filter(fixed == "not_fixed")

# 20_4
gt_20_4_fixed_ref <- gt_20_4 %>% 
  mutate(fixed = if_else((`20_4-F` == 0 | `2045-S1` == 0 | `2045-5-S2` == 0 | `2045-54-S3` == 0 | 
                            `2045-544-S4` == 0), "fixed_ref", 
                         if_else((`20_4-F` == 2 | `2045-S1` == 2 | `2045-5-S2` == 2 | `2045-54-S3` == 2 | 
                                    `2045-544-S4` == 2), "fixed_alt", "not_fixed"))) %>% 
  filter(fixed == "fixed_ref")

gt_20_4_fixed_alt <- gt_20_4 %>% 
  mutate(fixed = if_else((`20_4-F` == 0 | `2045-S1` == 0 | `2045-5-S2` == 0 | `2045-54-S3` == 0 | 
                            `2045-544-S4` == 0), "fixed_ref", 
                         if_else((`20_4-F` == 2 | `2045-S1` == 2 | `2045-5-S2` == 2 | `2045-54-S3` == 2 | 
                                    `2045-544-S4` == 2), "fixed_alt", "not_fixed"))) %>% 
  filter(fixed == "fixed_alt")

gt_20_4_not_fixed <- gt_20_4 %>% 
  mutate(fixed = if_else((`20_4-F` == 0 | `2045-S1` == 0 | `2045-5-S2` == 0 | `2045-54-S3` == 0 | 
                            `2045-544-S4` == 0), "fixed_ref", 
                         if_else((`20_4-F` == 2 | `2045-S1` == 2 | `2045-5-S2` == 2 | `2045-54-S3` == 2 | 
                                    `2045-544-S4` == 2), "fixed_alt", "not_fixed"))) %>% 
  filter(fixed == "not_fixed")

# 21_2
gt_21_2_fixed_ref <- gt_21_2 %>% 
  mutate(fixed = if_else((`21_2-F` == 0 | `2121-S1` == 0 | `2121-1-S2` == 0 | `2121-11-S3` == 0 | 
                            `2121-111-S4` == 0 | `2121-111-S5` == 0), "fixed_ref", 
                         if_else((`21_2-F` == 2 | `2121-S1` == 2 | `2121-1-S2` == 2 | `2121-11-S3` == 2 | 
                                    `2121-111-S4` == 2 | `2121-111-S5` == 2), "fixed_alt", "not_fixed"))) %>% 
  filter(fixed == "fixed_ref")

gt_21_2_fixed_alt <- gt_21_2 %>% 
  mutate(fixed = if_else((`21_2-F` == 0 | `2121-S1` == 0 | `2121-1-S2` == 0 | `2121-11-S3` == 0 | 
                            `2121-111-S4` == 0 | `2121-111-S5` == 0), "fixed_ref", 
                         if_else((`21_2-F` == 2 | `2121-S1` == 2 | `2121-1-S2` == 2 | `2121-11-S3` == 2 | 
                                    `2121-111-S4` == 2 | `2121-111-S5` == 2), "fixed_alt", "not_fixed"))) %>% 
  filter(fixed == "fixed_alt")

gt_21_2_not_fixed <- gt_21_2 %>% 
  mutate(fixed = if_else((`21_2-F` == 0 | `2121-S1` == 0 | `2121-1-S2` == 0 | `2121-11-S3` == 0 | 
                            `2121-111-S4` == 0 | `2121-111-S5` == 0), "fixed_ref", 
                         if_else((`21_2-F` == 2 | `2121-S1` == 2 | `2121-1-S2` == 2 | `2121-11-S3` == 2 | 
                                    `2121-111-S4` == 2 | `2121-111-S5` == 2), "fixed_alt", "not_fixed"))) %>% 
  filter(fixed == "not_fixed")

# 21_6
gt_21_6_fixed_ref <- gt_21_6 %>% 
  mutate(fixed = if_else((`21_6-F` == 0 | `2165-S1` == 0 | `2165-4-S2` == 0 | `2165-44-S3` == 0 | 
                            `2165-444-S4` == 0 | `2165-444-S5` == 0), "fixed_ref", 
                         if_else((`21_6-F` == 2 | `2165-S1` == 2 | `2165-4-S2` == 2 | `2165-44-S3` == 2 | 
                                    `2165-444-S4` == 2 | `2165-444-S5` == 2), "fixed_alt", "not_fixed"))) %>%  
  filter(fixed == "fixed_ref")

gt_21_6_fixed_alt <- gt_21_6 %>% 
  mutate(fixed = if_else((`21_6-F` == 0 | `2165-S1` == 0 | `2165-4-S2` == 0 | `2165-44-S3` == 0 | 
                            `2165-444-S4` == 0 | `2165-444-S5` == 0), "fixed_ref", 
                         if_else((`21_6-F` == 2 | `2165-S1` == 2 | `2165-4-S2` == 2 | `2165-44-S3` == 2 | 
                                    `2165-444-S4` == 2 | `2165-444-S5` == 2), "fixed_alt", "not_fixed"))) %>% 
  filter(fixed == "fixed_alt")

gt_21_6_not_fixed <- gt_21_6 %>% 
  mutate(fixed = if_else((`21_6-F` == 0 | `2165-S1` == 0 | `2165-4-S2` == 0 | `2165-44-S3` == 0 | 
                            `2165-444-S4` == 0 | `2165-444-S5` == 0), "fixed_ref", 
                         if_else((`21_6-F` == 2 | `2165-S1` == 2 | `2165-4-S2` == 2 | `2165-44-S3` == 2 | 
                                    `2165-444-S4` == 2 | `2165-444-S5` == 2), "fixed_alt", "not_fixed"))) %>% 
  filter(fixed == "not_fixed")

# 23_2
gt_23_2_fixed_ref <- gt_23_2 %>% 
  mutate(fixed = if_else((`23_2-F` == 0 | `2323-S1` == 0 | `2323-2-S2` == 0 | `2323-21-S3` == 0 | 
                            `2323-211-S4` == 0 | `2323-211-S5` == 0), "fixed_ref", 
                         if_else((`23_2-F` == 2 | `2323-S1` == 2 | `2323-2-S2` == 2 | `2323-21-S3` == 2 | 
                                    `2323-211-S4` == 2 | `2323-211-S5` == 2), "fixed_alt", "not_fixed"))) %>% 
  filter(fixed == "fixed_ref")

gt_23_2_fixed_alt <- gt_23_2 %>% 
  mutate(fixed = if_else((`23_2-F` == 0 | `2323-S1` == 0 | `2323-2-S2` == 0 | `2323-21-S3` == 0 | 
                            `2323-211-S4` == 0 | `2323-211-S5` == 0), "fixed_ref", 
                         if_else((`23_2-F` == 2 | `2323-S1` == 2 | `2323-2-S2` == 2 | `2323-21-S3` == 2 | 
                                    `2323-211-S4` == 2 | `2323-211-S5` == 2), "fixed_alt", "not_fixed"))) %>% 
  filter(fixed == "fixed_alt")

gt_23_2_not_fixed <- gt_23_2 %>% 
  mutate(fixed = if_else((`23_2-F` == 0 | `2323-S1` == 0 | `2323-2-S2` == 0 | `2323-21-S3` == 0 | 
                            `2323-211-S4` == 0 | `2323-211-S5` == 0), "fixed_ref", 
                         if_else((`23_2-F` == 2 | `2323-S1` == 2 | `2323-2-S2` == 2 | `2323-21-S3` == 2 | 
                                    `2323-211-S4` == 2 | `2323-211-S5` == 2), "fixed_alt", "not_fixed"))) %>% 
  filter(fixed == "not_fixed")

# 23_4
gt_23_4_fixed_ref <- gt_23_4 %>% 
  mutate(fixed = if_else((`23_4-F` == 0 | `2344-S1` == 0 | `2344-4-S2` == 0 | `2344-46-S3` == 0 | 
                            `2344-464-S4` == 0), "fixed_ref", 
                         if_else((`23_4-F` == 2 | `2344-S1` == 2 | `2344-4-S2` == 2 | `2344-46-S3` == 2 | 
                                    `2344-464-S4` == 2), "fixed_alt", "not_fixed"))) %>%
  filter(fixed == "fixed_ref")

gt_23_4_fixed_alt <- gt_23_4 %>% 
  mutate(fixed = if_else((`23_4-F` == 0 | `2344-S1` == 0 | `2344-4-S2` == 0 | `2344-46-S3` == 0 | 
                            `2344-464-S4` == 0), "fixed_ref", 
                         if_else((`23_4-F` == 2 | `2344-S1` == 2 | `2344-4-S2` == 2 | `2344-46-S3` == 2 | 
                                    `2344-464-S4` == 2), "fixed_alt", "not_fixed"))) %>%  
  filter(fixed == "fixed_alt")

gt_23_4_not_fixed <- gt_23_4 %>% 
  mutate(fixed = if_else((`23_4-F` == 0 | `2344-S1` == 0 | `2344-4-S2` == 0 | `2344-46-S3` == 0 | 
                            `2344-464-S4` == 0), "fixed_ref", 
                         if_else((`23_4-F` == 2 | `2344-S1` == 2 | `2344-4-S2` == 2 | `2344-46-S3` == 2 | 
                                    `2344-464-S4` == 2), "fixed_alt", "not_fixed"))) %>%
  filter(fixed == "not_fixed")

# 26_1
gt_26_1_fixed_ref <- gt_26_1 %>% 
  mutate(fixed = if_else((`26_1-F` == 0 | `2612-S1` == 0 | `2612-1-S2` == 0 | `2612-12-S3` == 0 | 
                            `2612-121-S4` == 0), "fixed_ref", 
                         if_else((`26_1-F` == 2 | `2612-S1` == 2 | `2612-1-S2` == 2 | `2612-12-S3` == 2 | 
                                    `2612-121-S4` == 2), "fixed_alt", "not_fixed"))) %>% 
  filter(fixed == "fixed_ref")

gt_26_1_fixed_alt <- gt_26_1 %>% 
  mutate(fixed = if_else((`26_1-F` == 0 | `2612-S1` == 0 | `2612-1-S2` == 0 | `2612-12-S3` == 0 | 
                            `2612-121-S4` == 0), "fixed_ref", 
                         if_else((`26_1-F` == 2 | `2612-S1` == 2 | `2612-1-S2` == 2 | `2612-12-S3` == 2 | 
                                    `2612-121-S4` == 2), "fixed_alt", "not_fixed"))) %>% 
  filter(fixed == "fixed_alt")

gt_26_1_not_fixed <- gt_26_1 %>% 
  mutate(fixed = if_else((`26_1-F` == 0 | `2612-S1` == 0 | `2612-1-S2` == 0 | `2612-12-S3` == 0 | 
                            `2612-121-S4` == 0), "fixed_ref", 
                         if_else((`26_1-F` == 2 | `2612-S1` == 2 | `2612-1-S2` == 2 | `2612-12-S3` == 2 | 
                                    `2612-121-S4` == 2), "fixed_alt", "not_fixed"))) %>% 
  filter(fixed == "not_fixed")

# 26_4
gt_26_4_fixed_ref <- gt_26_4 %>% 
  mutate(fixed = if_else((`26_4-F` == 0 | `2645-S1` == 0 | `2645-4-S2` == 0 | `2645-46-S3` == 0 | 
                            `2645-464-S4` == 0), "fixed_ref", 
                         if_else((`26_4-F` == 2 | `2645-S1` == 2 | `2645-4-S2` == 2 | `2645-46-S3` == 2 | 
                                    `2645-464-S4` == 2), "fixed_alt", "not_fixed"))) %>%
  filter(fixed == "fixed_ref")

gt_26_4_fixed_alt <- gt_26_4 %>% 
  mutate(fixed = if_else((`26_4-F` == 0 | `2645-S1` == 0 | `2645-4-S2` == 0 | `2645-46-S3` == 0 | 
                            `2645-464-S4` == 0), "fixed_ref", 
                         if_else((`26_4-F` == 2 | `2645-S1` == 2 | `2645-4-S2` == 2 | `2645-46-S3` == 2 | 
                                    `2645-464-S4` == 2), "fixed_alt", "not_fixed"))) %>%
  filter(fixed == "fixed_alt")

gt_26_4_not_fixed <- gt_26_4 %>% 
  mutate(fixed = if_else((`26_4-F` == 0 | `2645-S1` == 0 | `2645-4-S2` == 0 | `2645-46-S3` == 0 | 
                            `2645-464-S4` == 0), "fixed_ref", 
                         if_else((`26_4-F` == 2 | `2645-S1` == 2 | `2645-4-S2` == 2 | `2645-46-S3` == 2 | 
                                    `2645-464-S4` == 2), "fixed_alt", "not_fixed"))) %>% 
  filter(fixed == "not_fixed")

# 27_2
gt_27_2_fixed_ref <- gt_27_2 %>% 
  mutate(fixed = if_else((`27_2-F` == 0 | `2721-S1` == 0 | `2721-1-S2` == 0 | `2721-12-S3` == 0 | 
                            `2721-121-S4` == 0), "fixed_ref", 
                         if_else((`27_2-F` == 2 | `2721-S1` == 2 | `2721-1-S2` == 2 | `2721-12-S3` == 2 | 
                                    `2721-121-S4` == 2), "fixed_alt", "not_fixed"))) %>% 
  filter(fixed == "fixed_ref")

gt_27_2_fixed_alt <- gt_27_2 %>% 
  mutate(fixed = if_else((`27_2-F` == 0 | `2721-S1` == 0 | `2721-1-S2` == 0 | `2721-12-S3` == 0 | 
                            `2721-121-S4` == 0), "fixed_ref", 
                         if_else((`27_2-F` == 2 | `2721-S1` == 2 | `2721-1-S2` == 2 | `2721-12-S3` == 2 | 
                                    `2721-121-S4` == 2), "fixed_alt", "not_fixed"))) %>% 
  filter(fixed == "fixed_alt")

gt_27_2_not_fixed <- gt_27_2 %>% 
  mutate(fixed = if_else((`27_2-F` == 0 | `2721-S1` == 0 | `2721-1-S2` == 0 | `2721-12-S3` == 0 | 
                            `2721-121-S4` == 0), "fixed_ref", 
                         if_else((`27_2-F` == 2 | `2721-S1` == 2 | `2721-1-S2` == 2 | `2721-12-S3` == 2 | 
                                    `2721-121-S4` == 2), "fixed_alt", "not_fixed"))) %>% 
  filter(fixed == "not_fixed")

# 29_2
gt_29_2_fixed_ref <- gt_29_2 %>% 
  mutate(fixed = if_else((`29_2-F` == 0 | `2923-S1` == 0 | `2923-1-S2` == 0 | `2923-12-S3` == 0 | 
                            `2923-121-S4` == 0 | `2923-121-S5` == 0), "fixed_ref", 
                         if_else((`29_2-F` == 2 | `2923-S1` == 2 | `2923-1-S2` == 2 | `2923-12-S3` == 2 | 
                                    `2923-121-S4` == 2 | `2923-121-S5` == 2), "fixed_alt", "not_fixed"))) %>% 
  filter(fixed == "fixed_ref")

gt_29_2_fixed_alt <- gt_29_2 %>% 
  mutate(fixed = if_else((`29_2-F` == 0 | `2923-S1` == 0 | `2923-1-S2` == 0 | `2923-12-S3` == 0 | 
                            `2923-121-S4` == 0 | `2923-121-S5` == 0), "fixed_ref", 
                         if_else((`29_2-F` == 2 | `2923-S1` == 2 | `2923-1-S2` == 2 | `2923-12-S3` == 2 | 
                                    `2923-121-S4` == 2 | `2923-121-S5` == 2), "fixed_alt", "not_fixed"))) %>% 
  filter(fixed == "fixed_alt")

gt_29_2_not_fixed <- gt_29_2 %>% 
  mutate(fixed = if_else((`29_2-F` == 0 | `2923-S1` == 0 | `2923-1-S2` == 0 | `2923-12-S3` == 0 | 
                            `2923-121-S4` == 0 | `2923-121-S5` == 0), "fixed_ref", 
                         if_else((`29_2-F` == 2 | `2923-S1` == 2 | `2923-1-S2` == 2 | `2923-12-S3` == 2 | 
                                    `2923-121-S4` == 2 | `2923-121-S5` == 2), "fixed_alt", "not_fixed"))) %>% 
  filter(fixed == "not_fixed")

# 29_4
gt_29_4_fixed_ref <- gt_29_4 %>% 
  mutate(fixed = if_else((`29_4-F` == 0 | `2944-S1` == 0 | `2944-5-S2` == 0 | `2944-56-S3` == 0 | 
                            `2944-565-S4` == 0 | `2944-565-S5` == 0), "fixed_ref", 
                         if_else((`29_4-F` == 2 | `2944-S1` == 2 | `2944-5-S2` == 2 | `2944-56-S3` == 2 | 
                                    `2944-565-S4` == 2 | `2944-565-S5` == 2), "fixed_alt", "not_fixed"))) %>% 
  filter(fixed == "fixed_ref")

gt_29_4_fixed_alt <- gt_29_4 %>% 
  mutate(fixed = if_else((`29_4-F` == 0 | `2944-S1` == 0 | `2944-5-S2` == 0 | `2944-56-S3` == 0 | 
                            `2944-565-S4` == 0 | `2944-565-S5` == 0), "fixed_ref", 
                         if_else((`29_4-F` == 2 | `2944-S1` == 2 | `2944-5-S2` == 2 | `2944-56-S3` == 2 | 
                                    `2944-565-S4` == 2 | `2944-565-S5` == 2), "fixed_alt", "not_fixed"))) %>%
  filter(fixed == "fixed_alt")

gt_29_4_not_fixed <- gt_29_4 %>% 
  mutate(fixed = if_else((`29_4-F` == 0 | `2944-S1` == 0 | `2944-5-S2` == 0 | `2944-56-S3` == 0 | 
                            `2944-565-S4` == 0 | `2944-565-S5` == 0), "fixed_ref", 
                         if_else((`29_4-F` == 2 | `2944-S1` == 2 | `2944-5-S2` == 2 | `2944-56-S3` == 2 | 
                                    `2944-565-S4` == 2 | `2944-565-S5` == 2), "fixed_alt", "not_fixed"))) %>%
  filter(fixed == "not_fixed")

# ### All fixed at gen S1 (unused) #####
# gt_1_1_fixed_S1_ref <- gt_1_1 %>% 
#   filter(`111-S1` == 0)
# 
# gt_6_1_fixed_S1_ref <- gt_6_1 %>% 
#   filter(`611-S1` == 0)
# 
# gt_6_4_fixed_S1_ref <- gt_6_4 %>% 
#   filter(`646-S1` == 0)
# 
# gt_7_2_fixed_S1_ref <- gt_7_2 %>% 
#   filter(`721-S1` == 0)
# 
# gt_7_4_fixed_S1_ref <- gt_7_4 %>% 
#   filter(`745-S1` == 0)
# 
# gt_8_2_fixed_S1_ref <- gt_8_2 %>% 
#   filter(`821-S1` == 0)
# 
# gt_8_4_fixed_S1_ref <- gt_8_4 %>% 
#   filter(`845-S1` == 0)
# 
# gt_10_2_fixed_S1_ref <- gt_10_2 %>% 
#   filter(`1022-S1` == 0)
# 
# gt_10_5_fixed_S1_ref <- gt_10_5 %>% 
#   filter(`1056-S1` == 0)
# 
# gt_13_3_fixed_S1_ref <- gt_13_3 %>% 
#   filter(`1332-S1` == 0)
# 
# gt_13_4_fixed_S1_ref <- gt_13_4 %>% 
#   filter(`1345-S1` == 0)
# 
# gt_16_1_fixed_S1_ref <- gt_16_1 %>% 
#   filter(`1611-S1` == 0)
# 
# gt_16_5_fixed_S1_ref <- gt_16_5 %>% 
#   filter(`1654-S1` == 0)
# 
# gt_17_2_fixed_S1_ref <- gt_17_2 %>% 
#   filter(`1721-S1` == 0)
# 
# gt_17_5_fixed_S1_ref <- gt_17_5 %>% 
#   filter(`1755-S1` == 0)
# 
# gt_19_2_fixed_S1_ref <- gt_19_2 %>% 
#   filter(`1922-S1` == 0)
# 
# gt_19_5_fixed_S1_ref <- gt_19_5 %>% 
#   filter(`1955-S1` == 0)
# 
# gt_20_1_fixed_S1_ref <- gt_20_1 %>% 
#   filter(`2013-S1` == 0)
# 
# gt_20_4_fixed_S1_ref <- gt_20_4 %>% 
#   filter(`2045-S1` == 0)
# 
# gt_21_2_fixed_S1_ref <- gt_21_2 %>% 
#   filter(`2121-S1` == 0)
# 
# gt_21_6_fixed_S1_ref <- gt_21_6 %>% 
#   filter(`2165-S1` == 0)
# 
# gt_26_1_fixed_S1_ref <- gt_26_1 %>% 
#   filter(`2612-S1` == 0)
# 
# gt_26_4_fixed_S1_ref <- gt_26_4 %>% 
#   filter(`2645-S1` == 0)
# 
# gt_27_2_fixed_S1_ref <- gt_27_2 %>% 
#   filter(`2721-S1` == 0)
# 
# gt_29_2_fixed_S1_ref <- gt_29_2 %>% 
#   filter(`2923-S1` == 0)
# 
# gt_29_4_fixed_S1_ref <- gt_29_4 %>% 
#   filter(`2944-S1` == 0)
# 
# ### All fixed at gen S1 alt#####
# gt_1_1_fixed_S1_alt <- gt_1_1 %>% 
#   filter(`111-S1` == 2)
# 
# gt_6_1_fixed_S1_alt <- gt_6_1 %>% 
#   filter(`611-S1` == 2)
# 
# gt_6_4_fixed_S1_alt <- gt_6_4 %>% 
#   filter(`646-S1` == 2)
# 
# gt_7_2_fixed_S1_alt <- gt_7_2 %>% 
#   filter(`721-S1` == 2)
# 
# gt_7_4_fixed_S1_alt <- gt_7_4 %>% 
#   filter(`745-S1` == 2)
# 
# gt_8_2_fixed_S1_alt <- gt_8_2 %>% 
#   filter(`821-S1` == 2)
# 
# gt_8_4_fixed_S1_alt <- gt_8_4 %>% 
#   filter(`845-S1` == 2)
# 
# gt_10_2_fixed_S1_alt <- gt_10_2 %>% 
#   filter(`1022-S1` == 2)
# 
# gt_10_5_fixed_S1_alt <- gt_10_5 %>% 
#   filter(`1056-S1` == 2)
# 
# gt_13_3_fixed_S1_alt <- gt_13_3 %>% 
#   filter(`1332-S1` == 2)
# 
# gt_13_4_fixed_S1_alt <- gt_13_4 %>% 
#   filter(`1345-S1` == 2)
# 
# gt_16_1_fixed_S1_alt <- gt_16_1 %>% 
#   filter(`1611-S1` == 2)
# 
# gt_16_5_fixed_S1_alt <- gt_16_5 %>% 
#   filter(`1654-S1` == 2)
# 
# gt_17_2_fixed_S1_alt <- gt_17_2 %>% 
#   filter(`1721-S1` == 2)
# 
# gt_17_5_fixed_S1_alt <- gt_17_5 %>% 
#   filter(`1755-S1` == 2)
# 
# gt_19_2_fixed_S1_alt <- gt_19_2 %>% 
#   filter(`1922-S1` == 2)
# 
# gt_19_5_fixed_S1_alt <- gt_19_5 %>% 
#   filter(`1955-S1` == 2)
# 
# gt_20_1_fixed_S1_alt <- gt_20_1 %>% 
#   filter(`2013-S1` == 2)
# 
# gt_20_4_fixed_S1_alt <- gt_20_4 %>% 
#   filter(`2045-S1` == 2)
# 
# gt_21_2_fixed_S1_alt <- gt_21_2 %>% 
#   filter(`2121-S1` == 2)
# 
# gt_21_6_fixed_S1_alt <- gt_21_6 %>% 
#   filter(`2165-S1` == 2)
# 
# gt_26_1_fixed_S1_alt <- gt_26_1 %>% 
#   filter(`2612-S1` == 2)
# 
# gt_26_4_fixed_S1_alt <- gt_26_4 %>% 
#   filter(`2645-S1` == 2)
# 
# gt_27_2_fixed_S1_alt <- gt_27_2 %>% 
#   filter(`2721-S1` == 2)
# 
# gt_29_2_fixed_S1_alt <- gt_29_2 %>% 
#   filter(`2923-S1` == 2)
# 
# gt_29_4_fixed_S1_alt <- gt_29_4 %>% 
#   filter(`2944-S1` == 2)
# 
# ### All not fixed at gen S1 alt#####
# gt_1_1_not_fixed_S1 <- gt_1_1 %>% 
#   filter(`111-S1` == 1)
# 
# gt_6_1_not_fixed_S1 <- gt_6_1 %>% 
#   filter(`611-S1` == 1)
# 
# gt_6_4_not_fixed_S1 <- gt_6_4 %>% 
#   filter(`646-S1` == 1)
# 
# gt_7_2_not_fixed_S1 <- gt_7_2 %>% 
#   filter(`721-S1` == 1)
# 
# gt_7_4_not_fixed_S1 <- gt_7_4 %>% 
#   filter(`745-S1` == 1)
# 
# gt_8_2_not_fixed_S1 <- gt_8_2 %>% 
#   filter(`821-S1` == 1)
# 
# gt_8_4_not_fixed_S1 <- gt_8_4 %>% 
#   filter(`845-S1` == 1)
# 
# gt_10_2_not_fixed_S1 <- gt_10_2 %>% 
#   filter(`1022-S1` == 1)
# 
# gt_10_5_not_fixed_S1 <- gt_10_5 %>% 
#   filter(`1056-S1` == 1)
# 
# gt_13_3_not_fixed_S1 <- gt_13_3 %>% 
#   filter(`1332-S1` == 1)
# 
# gt_13_4_not_fixed_S1 <- gt_13_4 %>% 
#   filter(`1345-S1` == 1)
# 
# gt_16_1_not_fixed_S1 <- gt_16_1 %>% 
#   filter(`1611-S1` == 1)
# 
# gt_16_5_not_fixed_S1 <- gt_16_5 %>% 
#   filter(`1654-S1` == 1)
# 
# gt_17_2_not_fixed_S1 <- gt_17_2 %>% 
#   filter(`1721-S1` == 1)
# 
# gt_17_5_not_fixed_S1 <- gt_17_5 %>% 
#   filter(`1755-S1` == 1)
# 
# gt_19_2_not_fixed_S1 <- gt_19_2 %>% 
#   filter(`1922-S1` == 1)
# 
# gt_19_5_not_fixed_S1 <- gt_19_5 %>% 
#   filter(`1955-S1` == 1)
# 
# gt_20_1_not_fixed_S1 <- gt_20_1 %>% 
#   filter(`2013-S1` == 1)
# 
# gt_20_4_not_fixed_S1 <- gt_20_4 %>% 
#   filter(`2045-S1` == 1)
# 
# gt_21_2_not_fixed_S1 <- gt_21_2 %>% 
#   filter(`2121-S1` == 1)
# 
# gt_21_6_not_fixed_S1 <- gt_21_6 %>% 
#   filter(`2165-S1` == 1)
# 
# gt_26_1_not_fixed_S1 <- gt_26_1 %>% 
#   filter(`2612-S1` == 1)
# 
# gt_26_4_not_fixed_S1 <- gt_26_4 %>% 
#   filter(`2645-S1` == 1)
# 
# gt_27_2_not_fixed_S1 <- gt_27_2 %>% 
#   filter(`2721-S1` == 1)
# 
# gt_29_2_not_fixed_S1 <- gt_29_2 %>% 
#   filter(`2923-S1` == 1)
# 
# gt_29_4_not_fixed_S1 <- gt_29_4 %>% 
#   filter(`2944-S1` == 1)


gt_all <- merge(gt_1_1, gt_6_1, by = "SNP", all = T)
gt_all <- merge(gt_all, gt_6_4, by = "SNP", all = T)
gt_all <- merge(gt_all, gt_7_2, by = "SNP", all = T)
gt_all <- merge(gt_all, gt_7_4, by = "SNP", all = T)
gt_all <- merge(gt_all, gt_8_2, by = "SNP", all = T)
gt_all <- merge(gt_all, gt_8_4, by = "SNP", all = T)
gt_all <- merge(gt_all, gt_10_2, by = "SNP", all = T)
gt_all <- merge(gt_all, gt_10_5, by = "SNP", all = T)
gt_all <- merge(gt_all, gt_13_3, by = "SNP", all = T)
gt_all <- merge(gt_all, gt_13_4, by = "SNP", all = T)
gt_all <- merge(gt_all, gt_16_1, by = "SNP", all = T)
gt_all <- merge(gt_all, gt_16_5, by = "SNP", all = T)
gt_all <- merge(gt_all, gt_17_2, by = "SNP", all = T)
gt_all <- merge(gt_all, gt_17_5, by = "SNP", all = T)
gt_all <- merge(gt_all, gt_19_2, by = "SNP", all = T)
gt_all <- merge(gt_all, gt_19_5, by = "SNP", all = T)
gt_all <- merge(gt_all, gt_20_1, by = "SNP", all = T)
gt_all <- merge(gt_all, gt_20_4, by = "SNP", all = T)
gt_all <- merge(gt_all, gt_21_2, by = "SNP", all = T)
gt_all <- merge(gt_all, gt_21_6, by = "SNP", all = T)
gt_all <- merge(gt_all, gt_23_2, by = "SNP", all = T)
gt_all <- merge(gt_all, gt_23_4, by = "SNP", all = T)
gt_all <- merge(gt_all, gt_26_1, by = "SNP", all = T)
gt_all <- merge(gt_all, gt_26_4, by = "SNP", all = T)
gt_all <- merge(gt_all, gt_27_2, by = "SNP", all = T)
gt_all <- merge(gt_all, gt_29_2, by = "SNP", all = T)
gt_all <- merge(gt_all, gt_29_4, by = "SNP", all = T)

gt_all_F <- data.frame(SNP = gt_all$SNP, select_at(gt_all, vars(ends_with("F"))), check.names = F)
gt_all_S1 <- data.frame(SNP = gt_all$SNP, select_at(gt_all, vars(ends_with("S1"))), check.names = F)
gt_all_S2 <- data.frame(SNP = gt_all$SNP, select_at(gt_all, vars(ends_with("S2"))), check.names = F)
gt_all_S3 <- data.frame(SNP = gt_all$SNP, select_at(gt_all, vars(ends_with("S3"))), check.names = F)
gt_all_S4 <- data.frame(SNP = gt_all$SNP, select_at(gt_all, vars(ends_with("S4"))), check.names = F)
gt_all_S5 <- data.frame(SNP = gt_all$SNP, select_at(gt_all, vars(ends_with("S5"))), check.names = F)

F_lines_012_matrix <- gt_all_F %>% 
  filter(SNP %in% all_snp_ids$V1)

S1_lines_012_matrix <- gt_all_S1 %>% 
  filter(SNP %in% all_snp_ids$V1) 
  
S2_lines_012_matrix <- gt_all_S2 %>% 
  filter(SNP %in% all_snp_ids$V1) 
  
S3_lines_012_matrix <- gt_all_S3 %>% 
  filter(SNP %in% all_snp_ids$V1)
  
S4_lines_012_matrix <- gt_all_S4 %>% 
  filter(SNP %in% all_snp_ids$V1)
  
S5_lines_012_matrix <- gt_all_S5 %>% 
  filter(SNP %in% all_snp_ids$V1) 
  
all_lines_012 <- merge(F_lines_012_matrix, S1_lines_012_matrix, by = "SNP")
all_lines_012 <- merge(all_lines_012, S2_lines_012_matrix, by = "SNP")
all_lines_012 <- merge(all_lines_012, S3_lines_012_matrix, by = "SNP")
all_lines_012 <- merge(all_lines_012, S4_lines_012_matrix, by = "SNP")
all_lines_012 <- merge(all_lines_012, S5_lines_012_matrix, by = "SNP")

# write.table(all_lines_012, "all_lines_012_matrix.txt", row.names = F, col.names = T, quote = F)


# write.table(gt_all_F, "FS_lines_012.txt", row.names = F, quote = F)
# write.table(gt_all_S1, "S1_lines_012.txt", row.names = F, quote = F)
# write.table(gt_all_S2, "S2_lines_012.txt", row.names = F, quote = F)
# write.table(gt_all_S3, "S3_lines_012.txt", row.names = F, quote = F)
# write.table(gt_all_S4, "S4_lines_012.txt", row.names = F, quote = F)
# write.table(gt_all_S5, "S5_lines_012.txt", row.names = F, quote = F)

### Corrected allele matrices for pop gen, heterozygosity #####
FS_lines_allele_matrix <- read.csv("FS_lines_allele_matrix.csv", check.names = F, row.names = 1)
S1_lines_allele_matrix <- read.csv("S1_lines_allele_matrix.csv", check.names = F, row.names = 1)
S2_lines_allele_matrix <- read.csv("S2_lines_allele_matrix.csv", check.names = F, row.names = 1)
S3_lines_allele_matrix <- read.csv("S3_lines_allele_matrix.csv", check.names = F, row.names = 1)
S4_lines_allele_matrix <- read.csv("S4_lines_allele_matrix.csv", check.names = F, row.names = 1)
S5_lines_allele_matrix <- read.csv("S5_lines_allele_matrix.csv", check.names = F, row.names = 1)


### Filter for final SNP list
FS_lines_allele_matrix_all_filtered <- FS_lines_allele_matrix %>%
  rownames_to_column() %>%
  filter(rowname %in% all_snp_ids$V1) %>%
  column_to_rownames()

S1_lines_allele_matrix_all_filtered <- S1_lines_allele_matrix %>%
  rownames_to_column() %>%
  filter(rowname %in% all_snp_ids$V1) %>%
  column_to_rownames()

S2_lines_allele_matrix_all_filtered <- S2_lines_allele_matrix %>%
  rownames_to_column() %>%
  filter(rowname %in% all_snp_ids$V1) %>%
  column_to_rownames()

S3_lines_allele_matrix_all_filtered <- S3_lines_allele_matrix %>%
  rownames_to_column() %>%
  filter(rowname %in% all_snp_ids$V1) %>%
  column_to_rownames()

S4_lines_allele_matrix_all_filtered <- S4_lines_allele_matrix %>%
  rownames_to_column() %>%
  filter(rowname %in% all_snp_ids$V1) %>%
  column_to_rownames()

S5_lines_allele_matrix_all_filtered <- S5_lines_allele_matrix %>%
  rownames_to_column() %>%
  filter(rowname %in% all_snp_ids$V1) %>%
  column_to_rownames()

FS_lines_allele_matrix_all_filtered_t <- t(FS_lines_allele_matrix_all_filtered)
S1_lines_allele_matrix_all_filtered_t <- t(S1_lines_allele_matrix_all_filtered)
S2_lines_allele_matrix_all_filtered_t <- t(S2_lines_allele_matrix_all_filtered)
S3_lines_allele_matrix_all_filtered_t <- t(S3_lines_allele_matrix_all_filtered)
S4_lines_allele_matrix_all_filtered_t <- t(S4_lines_allele_matrix_all_filtered)
S5_lines_allele_matrix_all_filtered_t <- t(S5_lines_allele_matrix_all_filtered)

all_allele_matrix <- rbind(FS_lines_allele_matrix_all_filtered_t, S1_lines_allele_matrix_all_filtered_t, S2_lines_allele_matrix_all_filtered_t, S3_lines_allele_matrix_all_filtered_t, S4_lines_allele_matrix_all_filtered_t, S5_lines_allele_matrix_all_filtered_t)
# all_allele_matrix_neg_fis_filter <- rbind(FS_lines_allele_matrix_neg_fis_filter_t, S1_lines_allele_matrix_neg_fis_filter_t, 
#                                               S2_lines_allele_matrix_neg_fis_filter_t, S3_lines_allele_matrix_neg_fis_filter_t, 
#                                               S4_lines_allele_matrix_neg_fis_filter_t, S5_lines_allele_matrix_neg_fis_filter_t)

# Convert to genind
FS_S5_genind <- df2genind(all_allele_matrix, sep = "/", NA.char = "NA")
FS_lines_genind <- FS_S5_genind[1:28]
S1_lines_genind <- FS_S5_genind[29:56]
S2_lines_genind <- FS_S5_genind[57:84]
S3_lines_genind <- FS_S5_genind[85:112]
S4_lines_genind <- FS_S5_genind[113:140]
S5_lines_genind <- FS_S5_genind[141:151]

# Get summary for heterozygosity
FS_lines_div <- summary(FS_lines_genind)
S1_lines_div <- summary(S1_lines_genind)
S2_lines_div <- summary(S2_lines_genind)
S3_lines_div <- summary(S3_lines_genind)
S4_lines_div <- summary(S4_lines_genind)
S5_lines_div <- summary(S5_lines_genind)

### Getting expected het from FS observed het ################
generation_FS_S5_hobs <- data.frame(FS_hobs = FS_lines_div$Hobs,
                                    S1_hobs = S1_lines_div$Hobs,
                                    S2_hobs = S2_lines_div$Hobs,
                                    S3_hobs = S3_lines_div$Hobs,
                                    S4_hobs = S4_lines_div$Hobs,
                                    S5_hobs = S5_lines_div$Hobs)

colnames(generation_FS_S5_hobs) <- c("FS", "S1", "S2", "S3", "S4", "S5")
generation_FS_S5_hobs <- rownames_to_column(generation_FS_S5_hobs)
melted_generation_FS_S5_hobs <- melt(generation_FS_S5_hobs)
# 

# Add expectations of halving het every gen
expectations_halving <- melted_generation_FS_S5_hobs %>%
  filter(variable == "FS") %>%
  mutate(S1 = (value / 2), S2 = (value / 4), S3 = (value / 8), S4 = (value/ 16), S5 = (value / 32)) %>%
  rename(FS = value) %>%
  select(rowname, FS, S1, S2, S3, S4, S5)
# 
melted_expectations_halving <- melt(expectations_halving)

# write.table(melted_expectations_halving, "melted_hexp_halving_all_snps.txt", quote = F, row.names = F)
# write.table(melted_generation_FS_S5_hobs, "melted_generation_FS_S5_hobs_all_snps.txt", quote = F, row.names = F)

# Read in simplified hets with no expected het for FS - combined outside of R due to memory issues
generation_heterozygosities_obs_exp_halving <- read.table("observed_expected_het_all_snps.txt", header = F)
FS_S5_hets <- generation_heterozygosities_obs_exp_halving %>% 
  rename(Heterozygosity = V4)

# Observed hets for each gen
FS_S5_hets %>% 
  filter(Heterozygosity == "Observed") %>% 
  group_by(V2) %>% 
  summarise(mean = mean(V3), median = median(V3), sd = sd(V3))

FS_S5_hets %>% 
  filter(Heterozygosity == "Expected") %>% 
  group_by(V2) %>% 
  summarise(mean = mean(V3), median = median(V3), sd = sd(V3))
# summarise_all(list(~mean(.),~sd(.), ~list(c(summary(.))))) %>% 
# unnest_wider(list)

FS_het <- FS_S5_hets %>% 
  filter(Heterozygosity == "Observed", V2 == "FS")

S1_het <- FS_S5_hets %>% 
  filter(Heterozygosity == "Observed", V2 == "S1")

S2_het <- FS_S5_hets %>% 
  filter(Heterozygosity == "Observed", V2 == "S2")

S3_het <- FS_S5_hets %>% 
  filter(Heterozygosity == "Observed", V2 == "S3")

S4_het <- FS_S5_hets %>% 
  filter(Heterozygosity == "Observed", V2 == "S4")

S5_het <- FS_S5_hets %>% 
  filter(Heterozygosity == "Observed", V2 == "S5")

### Plots for Figure 4A ####
het_boxplot_exp_halving <- ggplot(FS_S5_hets, aes(x = V2, y = V3, fill = Heterozygosity)) +
  geom_boxplot(outlier.alpha = 0.3) +
  theme_Publication() +
  scale_fill_grey(start = 0.35) +
  scale_y_continuous(name = "Heterozygosity", breaks = c(0, 0.2, 0.4, 0.6, 0.8, 1)) +
  xlab("Generation") +
  scale_x_discrete("Generation", breaks = unique(FS_S5_hets$V2),
                   labels = c("FS",
                              "S1",
                              "S2",
                              "S3",
                              "S4",
                              "S5")) +
  theme(legend.title = element_blank(), legend.position = "top")
# theme(axis.text.x = element_text(vjust = -4))
# 
# het_violinplot_exp_halving <- ggplot(generation_heterozygosities_exp_halving, aes(x = V2, y = V3)) +
#   geom_violin(aes(color = V4), trim = F, position = position_dodge(0.8), scale = "count", width = 1.1) +
#   geom_boxplot(aes(color = V4), width = 0.05, position = position_dodge(0.8), outlier.alpha = 0.3) +
#   # stat_summary(fun = mean, geom="point", shape=23, size=2, position = position_dodge(0.9)) +
#   theme_Publication() +
#   scale_color_nejm() +
#   scale_y_continuous(name = "Heterozygosity", breaks = c(0, 0.2, 0.4, 0.6, 0.8, 1)) +
#   xlab("Generation")
# 
# het_violinplot_exp_halving <- ggplot(FS_S5_hets, aes(x = V2, y = V3)) +
#   geom_violin(aes(color = Heterozygosity), trim = T, position = position_dodge(1), scale = "width") +
#   geom_boxplot(aes(color = Heterozygosity), width = 0.15, position = position_dodge(1), outlier.alpha = 0.3) +
#   # stat_summary(fun = mean, geom="point", shape=23, size=2, position = position_dodge(0.9)) +
#   theme_Publication() +
#   scale_color_nejm() +
#   scale_y_continuous(name = "Heterozygosity", breaks = c(0, 0.2, 0.4, 0.6, 0.8, 1)) +
#   xlab("Generation") +
#   scale_x_discrete("Generation", breaks = unique(FS_S5_hets$V2),
#                    labels = c("FS (n = 28)",
#                               "S1 (n = 28)",
#                               "S2 (n = 28)",
#                               "S3 (n = 28)",
#                               "S4 (n = 28)",
#                               "S5 (n = 11)"))

het_boxplot_exp_halving
# het_violinplot_exp_halving

# ggsave("het_boxplot_exp_halving_exp_from_obs_legend_top_all_snps.tiff", het_boxplot_exp_halving, width = 7, height = 6)
# ggsave("het_violinplot_scale_width_S5.svg", het_violinplot_exp_halving, width = 10, height = 7)


## Wilcoxon test for difference of means ####
generation_heterozygosities_obs <- generation_heterozygosities_obs_exp_halving %>% 
  filter(V4 == "Observed")

generation_heterozygosities_exp <- generation_heterozygosities_obs_exp_halving %>% 
  filter(V4 == "Expected")

FS_obs <- generation_heterozygosities_obs %>% 
  filter(V2 == "FS")

summary(generation_heterozygosities_exp$V2)

het_two_cols <- read.table("observed_expected_het_all_snps_two_cols.txt", header = T)

FS <- het_two_cols %>% 
  filter(Generation == "FS")

mean(FS$Observed)
mean(FS$Expected)

S1 <- het_two_cols %>% 
  filter(Generation == "S1")

summary(S1$Observed)
summary(S1$Expected)

wilcox.test(S1$Observed, S1$Expected, paired = T, alternative = "less", digits.rank = 7)
mood.test(S1$Obs, S1$Exp, paired = T, alternative = "greater")

S2 <- het_two_cols %>% 
  filter(Generation == "S2")

mean(S2$Observed)
mean(S2$Expected)
summary(S2$Observed)
summary(S2$Expected)

wilcox.test(S2$Observed, S2$Expected, paired = T, alternative = "less", digits.rank = 7)
mood.test(S2$Obs, S2$Exp, paired = T, alternative = "greater")

S3 <- het_two_cols %>% 
  filter(Generation == "S3")

summary(S3$Observed)
summary(S3$Expected)

wilcox.test(S3$Observed, S3$Expected, paired = T, alternative = "less", digits.rank = 7)

S4 <- het_two_cols %>% 
  filter(Generation == "S4")

summary(S4$Observed)
summary(S4$Expected)

wilcox.test(S4$Observed, S4$Expected, paired = T, alternative = "less", digits.rank = 7)

S5 <- het_two_cols %>% 
  filter(Generation == "S5")

summary(S5$Observed)
summary(S5$Expected)

wilcox.test(S5$Observed, S5$Expected, paired = T, alternative = "less", digits.rank = 7)

### Sign test for difference of medians ####
#FS
FS_het <- generation_heterozygosities_obs_exp_halving %>% 
  filter(V2 =="FS")

FS_het %>% 
  group_by(V4) %>%
  summarise(across(V3, list(mean = mean, median = median)))

bxp <- ggpaired(FS_het, x = "V4", y = "V3", 
                order = c("Observed", "Expected"),
                ylab = "Het", xlab = "group")
# bxp

FS_test <- FS_het %>% 
  sign_test(V3 ~ V4) %>% 
  add_significance()
FS_test

#S1
S1_het <- generation_heterozygosities_obs_exp_halving %>% 
  filter(V2 =="S1")#, V4 == "Observed")

S1_het %>% 
  group_by(V4) %>%
  summarise(across(V3, list(mean = mean, median = median)))

bxp <- ggpaired(S1_het, x = "V4", y = "V3", 
                order = c("Observed", "Expected"),
                ylab = "Het", xlab = "group")
# bxp

S1_test <- S1_het %>%
  pairwise_sign_test(V3 ~ V4, detailed = T, ref.group = "Observed", alternative = "less") %>% #1, mu = 0.143) %>%
  add_significance()
S1_test


het_two_cols %>% 
  filter(Generation == "S1", !(Observed == Expected)) %>% 
  count(Observed > Expected)
  
p <- dbinom(0:7338, 14856, 0.5)
sum(p.adjust(p = p, method = "holm"))

#S2
S2_het <- generation_heterozygosities_obs_exp_halving %>% 
  filter(V2 =="S2")#, V4 == "Observed")

S2_het %>% 
  group_by(V4) %>%
  summarise(across(V3, list(mean = mean, median = median)))

bxp <- ggpaired(S2_het, x = "V4", y = "V3", 
                order = c("Observed", "Expected"),
                ylab = "Het", xlab = "group")
# bxp

S2_test <- S2_het %>% 
  pairwise_sign_test(V3 ~ V4, detailed = T, ref.group = "Observed", alternative = "less") %>% #1, mu = 0.0714) %>% 
  add_significance()
S2_test

#S3
S3_het <- generation_heterozygosities_obs_exp_halving %>% 
  filter(V2 =="S3")#, V4 == "Observed")

S3_het %>% 
  group_by(V4) %>%
  summarise(across(V3, list(mean = mean, median = median)))

bxp <- ggpaired(S3_het, x = "V4", y = "V3", 
                order = c("Observed", "Expected"),
                ylab = "Het", xlab = "group")
# bxp

S3_test <- S3_het %>% 
  pairwise_sign_test(V3 ~ V4, ref.group = "Observed", alternative = "less", detailed = T) %>% #1, mu = 0.0357) %>% 
  add_significance()
S3_test

#S4
S4_het <- generation_heterozygosities_obs_exp_halving %>% 
  filter(V2 =="S4")#, V4 == "Observed")

S4_het %>% 
  group_by(V4) %>%
  summarise(across(V3, list(mean = mean, median = median)))

bxp <- ggpaired(S4_het, x = "V4", y = "V3", 
                order = c("Observed", "Expected"),
                ylab = "Het", xlab = "group")
# bxp

S4_test <- S4_het %>%  
  pairwise_sign_test(V3 ~ V4, ref.group = "Observed", alternative = "less", detailed = T) %>% #,1, mu = 0.0179) %>% 
  add_significance()
S4_test

#S5
S5_het <- generation_heterozygosities_obs_exp_halving %>% 
  filter(V2 =="S5")#, V4 == "Observed")

S5_het %>% 
  group_by(V4) %>%
  summarise(across(V3, list(mean = mean, median = median)))

bxp <- ggpaired(S5_het, x = "V4", y = "V3", 
                order = c("Observed", "Expected"),
                ylab = "Het", xlab = "group")
# bxp

S5_test <- S5_het %>% 
  pairwise_sign_test(V3 ~ V4, ref.group = "Observed", alternative = "less", detailed = T) %>% 
  add_significance()
S5_test


### Heterozygosities in diverse pop #####
RWP_vcf <- read.vcfR("rwp.all_filtered_snps.vcf")
RWP_genind <- vcfR2genind(RWP_vcf)
RWP_div <- summary(RWP_genind)

summary(RWP_div$Hobs)
sd(RWP_div$Hobs)

### With just het F loci #####################################################################################
### FS gen ##############################################################################################
F_1_1_with_snp_names <- Line_1_1_het_F %>% 
  rownames_to_column() %>% 
  select(1:2)

# F_1_4_with_snp_names <- Line_1_4_het_F %>% 
#   rownames_to_column() %>% 
#   select(1:2)

F_6_1_with_snp_names <- Line_6_1_het_F %>% 
  rownames_to_column() %>% 
  select(1:2)

F_6_4_with_snp_names <- Line_6_4_het_F %>% 
  rownames_to_column() %>% 
  select(1:2)

F_7_2_with_snp_names <- Line_7_2_het_F %>% 
  rownames_to_column() %>% 
  select(1:2)

F_7_4_with_snp_names <- Line_7_4_het_F %>% 
  rownames_to_column() %>% 
  select(1:2)

F_8_2_with_snp_names <- Line_8_2_het_F %>% 
  rownames_to_column() %>% 
  select(1:2)

F_8_4_with_snp_names <- Line_8_4_het_F %>% 
  rownames_to_column() %>% 
  select(1:2)

F_10_2_with_snp_names <- Line_10_2_het_F %>%
  rownames_to_column() %>%
  select(1:2)

F_10_5_with_snp_names <- Line_10_5_het_F %>%
  rownames_to_column() %>%
  select(1:2)

F_13_3_with_snp_names <- Line_13_3_het_F %>% 
  rownames_to_column() %>% 
  select(1:2)

F_13_4_with_snp_names <- Line_13_4_het_F %>% 
  rownames_to_column() %>% 
  select(1:2)

F_16_1_with_snp_names <- Line_16_1_het_F %>% 
  rownames_to_column() %>% 
  select(1:2)

F_16_5_with_snp_names <- Line_16_5_het_F %>% 
  rownames_to_column() %>% 
  select(1:2)

F_17_2_with_snp_names <- Line_17_2_het_F %>% 
  rownames_to_column() %>% 
  select(1:2)

F_17_5_with_snp_names <- Line_17_5_het_F %>% 
  rownames_to_column() %>% 
  select(1:2)

F_19_2_with_snp_names <- Line_19_2_het_F %>% 
  rownames_to_column() %>% 
  select(1:2)

F_19_5_with_snp_names <- Line_19_5_het_F %>% 
  rownames_to_column() %>% 
  select(1:2)

F_20_1_with_snp_names <- Line_20_1_het_F %>% 
  rownames_to_column() %>% 
  select(1:2)

F_20_4_with_snp_names <- Line_20_4_het_F %>% 
  rownames_to_column() %>% 
  select(1:2)

F_21_2_with_snp_names <- Line_21_2_het_F %>% 
  rownames_to_column() %>% 
  select(1:2)

F_21_6_with_snp_names <- Line_21_6_het_F %>% 
  rownames_to_column() %>% 
  select(1:2)

F_23_2_with_snp_names <- Line_23_2_het_F %>%
  rownames_to_column() %>%
  select(1:2)

F_23_4_with_snp_names <- Line_23_4_het_F %>%
  rownames_to_column() %>%
  select(1:2)

F_26_1_with_snp_names <- Line_26_1_het_F %>% 
  rownames_to_column() %>% 
  select(1:2)

F_26_4_with_snp_names <- Line_26_4_het_F %>% 
  rownames_to_column() %>% 
  select(1:2)

F_27_2_with_snp_names <- Line_27_2_het_F %>%
  rownames_to_column() %>%
  select(1:2)

F_29_2_with_snp_names <- Line_29_2_het_F %>% 
  rownames_to_column() %>% 
  select(1:2)

F_29_4_with_snp_names <- Line_29_4_het_F %>% 
  rownames_to_column() %>% 
  select(1:2)


F_lines <- merge(F_1_1_with_snp_names, F_6_1_with_snp_names, by = "rowname", all = T)
F_lines <- merge(F_lines, F_6_4_with_snp_names, by = "rowname", all = T)
F_lines <- merge(F_lines, F_7_2_with_snp_names, by = "rowname", all = T)
F_lines <- merge(F_lines, F_7_4_with_snp_names, by = "rowname", all = T)
F_lines <- merge(F_lines, F_8_2_with_snp_names, by = "rowname", all = T)
F_lines <- merge(F_lines, F_8_4_with_snp_names, by = "rowname", all = T)
F_lines <- merge(F_lines, F_10_2_with_snp_names, by = "rowname", all = T)
F_lines <- merge(F_lines, F_10_5_with_snp_names, by = "rowname", all = T)
F_lines <- merge(F_lines, F_13_3_with_snp_names, by = "rowname", all = T)
F_lines <- merge(F_lines, F_13_4_with_snp_names, by = "rowname", all = T)
F_lines <- merge(F_lines, F_16_1_with_snp_names, by = "rowname", all = T)
F_lines <- merge(F_lines, F_16_5_with_snp_names, by = "rowname", all = T)
F_lines <- merge(F_lines, F_17_2_with_snp_names, by = "rowname", all = T)
F_lines <- merge(F_lines, F_17_5_with_snp_names, by = "rowname", all = T)
F_lines <- merge(F_lines, F_19_2_with_snp_names, by = "rowname", all = T)
F_lines <- merge(F_lines, F_19_5_with_snp_names, by = "rowname", all = T)
F_lines <- merge(F_lines, F_20_1_with_snp_names, by = "rowname", all = T)
F_lines <- merge(F_lines, F_20_4_with_snp_names, by = "rowname", all = T)
F_lines <- merge(F_lines, F_21_2_with_snp_names, by = "rowname", all = T)
F_lines <- merge(F_lines, F_21_6_with_snp_names, by = "rowname", all = T)
F_lines <- merge(F_lines, F_23_2_with_snp_names, by = "rowname", all = T)
F_lines <- merge(F_lines, F_23_4_with_snp_names, by = "rowname", all = T)
F_lines <- merge(F_lines, F_26_1_with_snp_names, by = "rowname", all = T)
F_lines <- merge(F_lines, F_26_4_with_snp_names, by = "rowname", all = T)
F_lines <- merge(F_lines, F_27_2_with_snp_names, by = "rowname", all = T)
F_lines <- merge(F_lines, F_29_2_with_snp_names, by = "rowname", all = T)
F_lines <- merge(F_lines, F_29_4_with_snp_names, by = "rowname", all = T)

# Minor allele frequencies
F_lines <- F_lines %>% 
  column_to_rownames()

n0 <- apply(F_lines==0,1,sum,na.rm=T)
n1 <- apply(F_lines==1,1,sum,na.rm=T)
n2 <- apply(F_lines==2,1,sum,na.rm=T)

n <- n0 + n1 + n2

p <- ((2*n0)+n1)/(2*n)
q <- 1 - p
maf <- pmin(p, q)
mgf <- apply(cbind(n0,n1,n2),1,min) / n

# allele_freqs_F <- Propfunc(F_lines)
allele_freqs_F <- data.frame(SNP = row.names(F_lines), frq = maf)

ggplot(allele_freqs_F, aes(x = frq)) +
  geom_density()

### S1 gen ##############################################################################################
S1_1_1_with_snp_names <- Line_1_1_het_F %>% 
  rownames_to_column() %>% 
  select(1, 3)

# S1_1_4_with_snp_names <- Line_1_4_het_F %>% 
#   rownames_to_column() %>% 
#   select(1, 3)

S1_6_1_with_snp_names <- Line_6_1_het_F %>% 
  rownames_to_column() %>% 
  select(1, 3)

S1_6_4_with_snp_names <- Line_6_4_het_F %>% 
  rownames_to_column() %>% 
  select(1, 3)

S1_7_2_with_snp_names <- Line_7_2_het_F %>% 
  rownames_to_column() %>% 
  select(1, 3)

S1_7_4_with_snp_names <- Line_7_4_het_F %>% 
  rownames_to_column() %>% 
  select(1, 3)

S1_8_2_with_snp_names <- Line_8_2_het_F %>% 
  rownames_to_column() %>% 
  select(1, 3)

S1_8_4_with_snp_names <- Line_8_4_het_F %>% 
  rownames_to_column() %>% 
  select(1, 3)

S1_10_2_with_snp_names <- Line_10_2_het_F %>%
  rownames_to_column() %>%
  select(1, 3)

S1_10_5_with_snp_names <- Line_10_5_het_F %>%
  rownames_to_column() %>%
  select(1, 3)

S1_13_3_with_snp_names <- Line_13_3_het_F %>% 
  rownames_to_column() %>% 
  select(1, 3)

S1_13_4_with_snp_names <- Line_13_4_het_F %>% 
  rownames_to_column() %>% 
  select(1, 3)

S1_16_1_with_snp_names <- Line_16_1_het_F %>% 
  rownames_to_column() %>% 
  select(1, 3)

S1_16_5_with_snp_names <- Line_16_5_het_F %>% 
  rownames_to_column() %>% 
  select(1, 3)

S1_17_2_with_snp_names <- Line_17_2_het_F %>% 
  rownames_to_column() %>% 
  select(1, 3)

S1_17_5_with_snp_names <- Line_17_5_het_F %>% 
  rownames_to_column() %>% 
  select(1, 3)

S1_19_2_with_snp_names <- Line_19_2_het_F %>% 
  rownames_to_column() %>% 
  select(1, 3)

S1_19_5_with_snp_names <- Line_19_5_het_F %>% 
  rownames_to_column() %>% 
  select(1, 3)

S1_20_1_with_snp_names <- Line_20_1_het_F %>% 
  rownames_to_column() %>% 
  select(1, 3)

S1_20_4_with_snp_names <- Line_20_4_het_F %>% 
  rownames_to_column() %>% 
  select(1, 3)

S1_21_2_with_snp_names <- Line_21_2_het_F %>% 
  rownames_to_column() %>% 
  select(1, 3)

S1_21_6_with_snp_names <- Line_21_6_het_F %>% 
  rownames_to_column() %>% 
  select(1, 3)

S1_23_2_with_snp_names <- Line_23_2_het_F %>% 
  rownames_to_column() %>% 
  select(1, 3)
 
S1_23_4_with_snp_names <- Line_23_4_het_F %>% 
  rownames_to_column() %>% 
  select(1, 3)

S1_26_1_with_snp_names <- Line_26_1_het_F %>% 
  rownames_to_column() %>% 
  select(1, 3)

S1_26_1_with_snp_names <- Line_26_1_het_F %>% 
  rownames_to_column() %>% 
  select(1, 3)

S1_26_4_with_snp_names <- Line_26_4_het_F %>% 
  rownames_to_column() %>% 
  select(1, 3)

S1_27_2_with_snp_names <- Line_27_2_het_F %>%
  rownames_to_column() %>%
  select(1, 3)

S1_29_2_with_snp_names <- Line_29_2_het_F %>% 
  rownames_to_column() %>% 
  select(1, 3)

S1_29_4_with_snp_names <- Line_29_4_het_F %>% 
  rownames_to_column() %>% 
  select(1, 3)


S1_lines <- merge(S1_1_1_with_snp_names, S1_6_1_with_snp_names, by = "rowname", all = T)
S1_lines <- merge(S1_lines, S1_6_4_with_snp_names, by = "rowname", all = T)
S1_lines <- merge(S1_lines, S1_7_2_with_snp_names, by = "rowname", all = T)
S1_lines <- merge(S1_lines, S1_7_4_with_snp_names, by = "rowname", all = T)
S1_lines <- merge(S1_lines, S1_8_2_with_snp_names, by = "rowname", all = T)
S1_lines <- merge(S1_lines, S1_8_4_with_snp_names, by = "rowname", all = T)
S1_lines <- merge(S1_lines, S1_10_2_with_snp_names, by = "rowname", all = T)
S1_lines <- merge(S1_lines, S1_10_5_with_snp_names, by = "rowname", all = T)
S1_lines <- merge(S1_lines, S1_13_3_with_snp_names, by = "rowname", all = T)
S1_lines <- merge(S1_lines, S1_13_4_with_snp_names, by = "rowname", all = T)
S1_lines <- merge(S1_lines, S1_16_1_with_snp_names, by = "rowname", all = T)
S1_lines <- merge(S1_lines, S1_16_5_with_snp_names, by = "rowname", all = T)
S1_lines <- merge(S1_lines, S1_17_2_with_snp_names, by = "rowname", all = T)
S1_lines <- merge(S1_lines, S1_17_5_with_snp_names, by = "rowname", all = T)
S1_lines <- merge(S1_lines, S1_19_2_with_snp_names, by = "rowname", all = T)
S1_lines <- merge(S1_lines, S1_19_5_with_snp_names, by = "rowname", all = T)
S1_lines <- merge(S1_lines, S1_20_1_with_snp_names, by = "rowname", all = T)
S1_lines <- merge(S1_lines, S1_20_4_with_snp_names, by = "rowname", all = T)
S1_lines <- merge(S1_lines, S1_21_2_with_snp_names, by = "rowname", all = T)
S1_lines <- merge(S1_lines, S1_21_6_with_snp_names, by = "rowname", all = T)
S1_lines <- merge(S1_lines, S1_23_2_with_snp_names, by = "rowname", all = T)
S1_lines <- merge(S1_lines, S1_23_4_with_snp_names, by = "rowname", all = T)
S1_lines <- merge(S1_lines, S1_26_1_with_snp_names, by = "rowname", all = T)
S1_lines <- merge(S1_lines, S1_26_4_with_snp_names, by = "rowname", all = T)
S1_lines <- merge(S1_lines, S1_27_2_with_snp_names, by = "rowname", all = T)
S1_lines <- merge(S1_lines, S1_29_2_with_snp_names, by = "rowname", all = T)
S1_lines <- merge(S1_lines, S1_29_4_with_snp_names, by = "rowname", all = T)

# Minor allele freqs
S1_lines <- S1_lines %>% 
  column_to_rownames()

n0 <- apply(S1_lines==0,1,sum,na.rm=T)
n1 <- apply(S1_lines==1,1,sum,na.rm=T)
n2 <- apply(S1_lines==2,1,sum,na.rm=T)

n <- n0 + n1 + n2

p <- ((2*n0)+n1)/(2*n)
q <- 1 - p
maf <- pmin(p, q)
mgf <- apply(cbind(n0,n1,n2),1,min) / n

# allele_freqs_S1 <- Propfunc(S1_lines)
allele_freqs_S1 <- data.frame(SNP = rownames(S1_lines), frq = maf)

ggplot(allele_freqs_S1, aes(x = frq)) +
  geom_histogram(binwidth = 0.01)

### S2 gen ##############################################################################################
S2_1_1_with_snp_names <- Line_1_1_het_F %>% 
  rownames_to_column() %>% 
  select(1, 4)

# S2_1_4_with_snp_names <- Line_1_4_het_F %>% 
#   rownames_to_column() %>% 
#   select(1, 4)

S2_6_1_with_snp_names <- Line_6_1_het_F %>% 
  rownames_to_column() %>% 
  select(1, 4)

S2_6_4_with_snp_names <- Line_6_4_het_F %>% 
  rownames_to_column() %>% 
  select(1, 4)

S2_7_2_with_snp_names <- Line_7_2_het_F %>% 
  rownames_to_column() %>% 
  select(1, 4)

S2_7_4_with_snp_names <- Line_7_4_het_F %>% 
  rownames_to_column() %>% 
  select(1, 4)

S2_8_2_with_snp_names <- Line_8_2_het_F %>% 
  rownames_to_column() %>% 
  select(1, 4)

S2_8_4_with_snp_names <- Line_8_4_het_F %>% 
  rownames_to_column() %>% 
  select(1, 4)

S2_10_2_with_snp_names <- Line_10_2_het_F %>%
  rownames_to_column() %>%
  select(1, 4)

S2_10_5_with_snp_names <- Line_10_5_het_F %>%
  rownames_to_column() %>%
  select(1, 4)

S2_13_3_with_snp_names <- Line_13_3_het_F %>% 
  rownames_to_column() %>% 
  select(1, 4)

S2_13_4_with_snp_names <- Line_13_4_het_F %>% 
  rownames_to_column() %>% 
  select(1, 4)

S2_16_1_with_snp_names <- Line_16_1_het_F %>% 
  rownames_to_column() %>% 
  select(1, 4)

S2_16_5_with_snp_names <- Line_16_5_het_F %>% 
  rownames_to_column() %>% 
  select(1, 4)

S2_17_2_with_snp_names <- Line_17_2_het_F %>% 
  rownames_to_column() %>% 
  select(1, 4)

S2_17_5_with_snp_names <- Line_17_5_het_F %>% 
  rownames_to_column() %>% 
  select(1, 4)

S2_19_2_with_snp_names <- Line_19_2_het_F %>% 
  rownames_to_column() %>% 
  select(1, 4)

S2_19_5_with_snp_names <- Line_19_5_het_F %>% 
  rownames_to_column() %>% 
  select(1, 4)

S2_20_1_with_snp_names <- Line_20_1_het_F %>% 
  rownames_to_column() %>% 
  select(1, 4)

S2_20_4_with_snp_names <- Line_20_4_het_F %>% 
  rownames_to_column() %>% 
  select(1, 4)

S2_21_2_with_snp_names <- Line_21_2_het_F %>% 
  rownames_to_column() %>% 
  select(1, 4)

S2_21_6_with_snp_names <- Line_21_6_het_F %>% 
  rownames_to_column() %>% 
  select(1, 4)

S2_23_2_with_snp_names <- Line_23_2_het_F %>% 
  rownames_to_column() %>% 
  select(1, 4)

S2_23_4_with_snp_names <- Line_23_4_het_F %>% 
  rownames_to_column() %>% 
  select(1, 4)

S2_26_1_with_snp_names <- Line_26_1_het_F %>% 
  rownames_to_column() %>% 
  select(1, 4)

S2_26_1_with_snp_names <- Line_26_1_het_F %>% 
  rownames_to_column() %>% 
  select(1, 4)

S2_26_4_with_snp_names <- Line_26_4_het_F %>% 
  rownames_to_column() %>% 
  select(1, 4)

S2_27_2_with_snp_names <- Line_27_2_het_F %>%
  rownames_to_column() %>%
  select(1, 4)

S2_29_2_with_snp_names <- Line_29_2_het_F %>% 
  rownames_to_column() %>% 
  select(1, 4)

S2_29_4_with_snp_names <- Line_29_4_het_F %>% 
  rownames_to_column() %>% 
  select(1, 4)


S2_lines <- merge(S2_1_1_with_snp_names, S2_6_1_with_snp_names, by = "rowname", all = T)
S2_lines <- merge(S2_lines, S2_6_4_with_snp_names, by = "rowname", all = T)
S2_lines <- merge(S2_lines, S2_7_2_with_snp_names, by = "rowname", all = T)
S2_lines <- merge(S2_lines, S2_7_4_with_snp_names, by = "rowname", all = T)
S2_lines <- merge(S2_lines, S2_8_2_with_snp_names, by = "rowname", all = T)
S2_lines <- merge(S2_lines, S2_8_4_with_snp_names, by = "rowname", all = T)
S2_lines <- merge(S2_lines, S2_10_2_with_snp_names, by = "rowname", all = T)
S2_lines <- merge(S2_lines, S2_10_5_with_snp_names, by = "rowname", all = T)
S2_lines <- merge(S2_lines, S2_13_3_with_snp_names, by = "rowname", all = T)
S2_lines <- merge(S2_lines, S2_13_4_with_snp_names, by = "rowname", all = T)
S2_lines <- merge(S2_lines, S2_16_1_with_snp_names, by = "rowname", all = T)
S2_lines <- merge(S2_lines, S2_16_5_with_snp_names, by = "rowname", all = T)
S2_lines <- merge(S2_lines, S2_17_2_with_snp_names, by = "rowname", all = T)
S2_lines <- merge(S2_lines, S2_17_5_with_snp_names, by = "rowname", all = T)
S2_lines <- merge(S2_lines, S2_19_2_with_snp_names, by = "rowname", all = T)
S2_lines <- merge(S2_lines, S2_19_5_with_snp_names, by = "rowname", all = T)
S2_lines <- merge(S2_lines, S2_20_1_with_snp_names, by = "rowname", all = T)
S2_lines <- merge(S2_lines, S2_20_4_with_snp_names, by = "rowname", all = T)
S2_lines <- merge(S2_lines, S2_21_2_with_snp_names, by = "rowname", all = T)
S2_lines <- merge(S2_lines, S2_21_6_with_snp_names, by = "rowname", all = T)
S2_lines <- merge(S2_lines, S2_23_2_with_snp_names, by = "rowname", all = T)
S2_lines <- merge(S2_lines, S2_23_4_with_snp_names, by = "rowname", all = T)
S2_lines <- merge(S2_lines, S2_26_1_with_snp_names, by = "rowname", all = T)
S2_lines <- merge(S2_lines, S2_26_4_with_snp_names, by = "rowname", all = T)
S2_lines <- merge(S2_lines, S2_27_2_with_snp_names, by = "rowname", all = T)
S2_lines <- merge(S2_lines, S2_29_2_with_snp_names, by = "rowname", all = T)
S2_lines <- merge(S2_lines, S2_29_4_with_snp_names, by = "rowname", all = T)

# Minor allele freqs
S2_lines <- S2_lines %>% 
  column_to_rownames()

n0 <- apply(S2_lines==0,1,sum,na.rm=T)
n1 <- apply(S2_lines==1,1,sum,na.rm=T)
n2 <- apply(S2_lines==2,1,sum,na.rm=T)

n <- n0 + n1 + n2

p <- ((2*n0)+n1)/(2*n)
q <- 1 - p
maf <- pmin(p, q)
mgf <- apply(cbind(n0,n1,n2),1,min) / n

#allele_freqs_S2 <- Propfunc(S2_lines)
allele_freqs_S2 <- data.frame(SNP = rownames(S2_lines), frq = maf)

ggplot(allele_freqs_S2, aes(x = frq)) +
  geom_density()

### S3 gen ##############################################################################################
S3_1_1_with_snp_names <- Line_1_1_het_F %>% 
  rownames_to_column() %>% 
  select(1, 5)

# S3_1_4_with_snp_names <- Line_1_4_het_F %>% 
#   rownames_to_column() %>% 
#   select(1, 5)

S3_6_1_with_snp_names <- Line_6_1_het_F %>% 
  rownames_to_column() %>% 
  select(1, 5)

S3_6_4_with_snp_names <- Line_6_4_het_F %>% 
  rownames_to_column() %>% 
  select(1, 5)

S3_7_2_with_snp_names <- Line_7_2_het_F %>% 
  rownames_to_column() %>% 
  select(1, 5)

S3_7_4_with_snp_names <- Line_7_4_het_F %>% 
  rownames_to_column() %>% 
  select(1, 5)

S3_8_2_with_snp_names <- Line_8_2_het_F %>% 
  rownames_to_column() %>% 
  select(1, 5)

S3_8_4_with_snp_names <- Line_8_4_het_F %>% 
  rownames_to_column() %>% 
  select(1, 5)

S3_10_2_with_snp_names <- Line_10_2_het_F %>%
  rownames_to_column() %>%
  select(1, 5)

S3_10_5_with_snp_names <- Line_10_5_het_F %>%
  rownames_to_column() %>%
  select(1, 5)

S3_13_3_with_snp_names <- Line_13_3_het_F %>% 
  rownames_to_column() %>% 
  select(1, 5)

S3_13_4_with_snp_names <- Line_13_4_het_F %>% 
  rownames_to_column() %>% 
  select(1, 5)

S3_16_1_with_snp_names <- Line_16_1_het_F %>% 
  rownames_to_column() %>% 
  select(1, 5)

S3_16_5_with_snp_names <- Line_16_5_het_F %>% 
  rownames_to_column() %>% 
  select(1, 5)

S3_17_2_with_snp_names <- Line_17_2_het_F %>% 
  rownames_to_column() %>% 
  select(1, 5)

S3_17_5_with_snp_names <- Line_17_5_het_F %>% 
  rownames_to_column() %>% 
  select(1, 5)

S3_19_2_with_snp_names <- Line_19_2_het_F %>% 
  rownames_to_column() %>% 
  select(1, 5)

S3_19_5_with_snp_names <- Line_19_5_het_F %>% 
  rownames_to_column() %>% 
  select(1, 5)

S3_20_1_with_snp_names <- Line_20_1_het_F %>% 
  rownames_to_column() %>% 
  select(1, 5)

S3_20_4_with_snp_names <- Line_20_4_het_F %>% 
  rownames_to_column() %>% 
  select(1, 5)

S3_21_2_with_snp_names <- Line_21_2_het_F %>% 
  rownames_to_column() %>% 
  select(1, 5)

S3_21_6_with_snp_names <- Line_21_6_het_F %>% 
  rownames_to_column() %>% 
  select(1, 5)

S3_23_2_with_snp_names <- Line_23_2_het_F %>% 
  rownames_to_column() %>% 
  select(1, 5)
 
S3_23_4_with_snp_names <- Line_23_4_het_F %>% 
  rownames_to_column() %>% 
  select(1, 5)

S3_26_1_with_snp_names <- Line_26_1_het_F %>% 
  rownames_to_column() %>% 
  select(1, 5)

S3_26_1_with_snp_names <- Line_26_1_het_F %>% 
  rownames_to_column() %>% 
  select(1, 5)

S3_26_4_with_snp_names <- Line_26_4_het_F %>% 
  rownames_to_column() %>% 
  select(1, 5)

S3_27_2_with_snp_names <- Line_27_2_het_F %>%
  rownames_to_column() %>%
  select(1, 5)

S3_29_2_with_snp_names <- Line_29_2_het_F %>% 
  rownames_to_column() %>% 
  select(1, 5)

S3_29_4_with_snp_names <- Line_29_4_het_F %>% 
  rownames_to_column() %>% 
  select(1, 5)


S3_lines <- merge(S3_1_1_with_snp_names, S3_6_1_with_snp_names, by = "rowname", all = T)
S3_lines <- merge(S3_lines, S3_6_4_with_snp_names, by = "rowname", all = T)
S3_lines <- merge(S3_lines, S3_7_2_with_snp_names, by = "rowname", all = T)
S3_lines <- merge(S3_lines, S3_7_4_with_snp_names, by = "rowname", all = T)
S3_lines <- merge(S3_lines, S3_8_2_with_snp_names, by = "rowname", all = T)
S3_lines <- merge(S3_lines, S3_8_4_with_snp_names, by = "rowname", all = T)
S3_lines <- merge(S3_lines, S3_10_2_with_snp_names, by = "rowname", all = T)
S3_lines <- merge(S3_lines, S3_10_5_with_snp_names, by = "rowname", all = T)
S3_lines <- merge(S3_lines, S3_13_3_with_snp_names, by = "rowname", all = T)
S3_lines <- merge(S3_lines, S3_13_4_with_snp_names, by = "rowname", all = T)
S3_lines <- merge(S3_lines, S3_16_1_with_snp_names, by = "rowname", all = T)
S3_lines <- merge(S3_lines, S3_16_5_with_snp_names, by = "rowname", all = T)
S3_lines <- merge(S3_lines, S3_17_2_with_snp_names, by = "rowname", all = T)
S3_lines <- merge(S3_lines, S3_17_5_with_snp_names, by = "rowname", all = T)
S3_lines <- merge(S3_lines, S3_19_2_with_snp_names, by = "rowname", all = T)
S3_lines <- merge(S3_lines, S3_19_5_with_snp_names, by = "rowname", all = T)
S3_lines <- merge(S3_lines, S3_20_1_with_snp_names, by = "rowname", all = T)
S3_lines <- merge(S3_lines, S3_20_4_with_snp_names, by = "rowname", all = T)
S3_lines <- merge(S3_lines, S3_21_2_with_snp_names, by = "rowname", all = T)
S3_lines <- merge(S3_lines, S3_21_6_with_snp_names, by = "rowname", all = T)
S3_lines <- merge(S3_lines, S3_23_2_with_snp_names, by = "rowname", all = T)
S3_lines <- merge(S3_lines, S3_23_4_with_snp_names, by = "rowname", all = T)
S3_lines <- merge(S3_lines, S3_26_1_with_snp_names, by = "rowname", all = T)
S3_lines <- merge(S3_lines, S3_26_4_with_snp_names, by = "rowname", all = T)
S3_lines <- merge(S3_lines, S3_27_2_with_snp_names, by = "rowname", all = T)
S3_lines <- merge(S3_lines, S3_29_2_with_snp_names, by = "rowname", all = T)
S3_lines <- merge(S3_lines, S3_29_4_with_snp_names, by = "rowname", all = T)

# Minor allele freqs
S3_lines <- S3_lines %>% 
  column_to_rownames()

n0 <- apply(S3_lines==0,1,sum,na.rm=T)
n1 <- apply(S3_lines==1,1,sum,na.rm=T)
n2 <- apply(S3_lines==2,1,sum,na.rm=T)

n <- n0 + n1 + n2

p <- ((2*n0)+n1)/(2*n)
q <- 1 - p
maf <- pmin(p, q)
mgf <- apply(cbind(n0,n1,n2),1,min) / n

#allele_freqs_S3 <- Propfunc(S3_lines)
allele_freqs_S3 <- data.frame(SNP = rownames(S3_lines), frq = maf)

ggplot(allele_freqs_S3, aes(x = frq)) +
  geom_density()

### S4 gen ##############################################################################################
S4_1_1_with_snp_names <- Line_1_1_het_F %>% 
  rownames_to_column() %>% 
  select(1, 6)

# S4_1_4_with_snp_names <- Line_1_4_het_F %>% 
#   rownames_to_column() %>% 
#   select(1, 6)

S4_6_1_with_snp_names <- Line_6_1_het_F %>% 
  rownames_to_column() %>% 
  select(1, 6)

S4_6_4_with_snp_names <- Line_6_4_het_F %>% 
  rownames_to_column() %>% 
  select(1, 6)

S4_7_2_with_snp_names <- Line_7_2_het_F %>% 
  rownames_to_column() %>% 
  select(1, 6)

S4_7_4_with_snp_names <- Line_7_4_het_F %>% 
  rownames_to_column() %>% 
  select(1, 6)

S4_8_2_with_snp_names <- Line_8_2_het_F %>% 
  rownames_to_column() %>% 
  select(1, 6)

S4_8_4_with_snp_names <- Line_8_4_het_F %>% 
  rownames_to_column() %>% 
  select(1, 6)

S4_10_2_with_snp_names <- Line_10_2_het_F %>%
  rownames_to_column() %>%
  select(1, 6)

S4_10_5_with_snp_names <- Line_10_5_het_F %>%
  rownames_to_column() %>%
  select(1, 6)

S4_13_3_with_snp_names <- Line_13_3_het_F %>% 
  rownames_to_column() %>% 
  select(1, 6)

S4_13_4_with_snp_names <- Line_13_4_het_F %>% 
  rownames_to_column() %>% 
  select(1, 6)

S4_16_1_with_snp_names <- Line_16_1_het_F %>% 
  rownames_to_column() %>% 
  select(1, 6)

S4_16_5_with_snp_names <- Line_16_5_het_F %>% 
  rownames_to_column() %>% 
  select(1, 6)

S4_17_2_with_snp_names <- Line_17_2_het_F %>% 
  rownames_to_column() %>% 
  select(1, 6)

S4_17_5_with_snp_names <- Line_17_5_het_F %>% 
  rownames_to_column() %>% 
  select(1, 6)

S4_19_2_with_snp_names <- Line_19_2_het_F %>% 
  rownames_to_column() %>% 
  select(1, 6)

S4_19_5_with_snp_names <- Line_19_5_het_F %>% 
  rownames_to_column() %>% 
  select(1, 6)

S4_20_1_with_snp_names <- Line_20_1_het_F %>% 
  rownames_to_column() %>% 
  select(1, 6)

S4_20_4_with_snp_names <- Line_20_4_het_F %>% 
  rownames_to_column() %>% 
  select(1, 6)

S4_21_2_with_snp_names <- Line_21_2_het_F %>% 
  rownames_to_column() %>% 
  select(1, 6)

S4_21_6_with_snp_names <- Line_21_6_het_F %>% 
  rownames_to_column() %>% 
  select(1, 6)

S4_23_2_with_snp_names <- Line_23_2_het_F %>% 
  rownames_to_column() %>% 
  select(1, 6)
 
S4_23_4_with_snp_names <- Line_23_4_het_F %>% 
  rownames_to_column() %>% 
  select(1, 6)

S4_26_1_with_snp_names <- Line_26_1_het_F %>% 
  rownames_to_column() %>% 
  select(1, 6)

S4_26_4_with_snp_names <- Line_26_4_het_F %>% 
  rownames_to_column() %>% 
  select(1, 6)

S4_27_2_with_snp_names <- Line_27_2_het_F %>%
  rownames_to_column() %>%
  select(1, 6)

S4_29_2_with_snp_names <- Line_29_2_het_F %>% 
  rownames_to_column() %>% 
  select(1, 6)

S4_29_4_with_snp_names <- Line_29_4_het_F %>% 
  rownames_to_column() %>% 
  select(1, 6)


S4_lines <- merge(S4_1_1_with_snp_names, S4_6_1_with_snp_names, by = "rowname", all = T)
S4_lines <- merge(S4_lines, S4_6_4_with_snp_names, by = "rowname", all = T)
S4_lines <- merge(S4_lines, S4_7_2_with_snp_names, by = "rowname", all = T)
S4_lines <- merge(S4_lines, S4_7_4_with_snp_names, by = "rowname", all = T)
S4_lines <- merge(S4_lines, S4_8_2_with_snp_names, by = "rowname", all = T)
S4_lines <- merge(S4_lines, S4_8_4_with_snp_names, by = "rowname", all = T)
S4_lines <- merge(S4_lines, S4_10_2_with_snp_names, by = "rowname", all = T)
S4_lines <- merge(S4_lines, S4_10_5_with_snp_names, by = "rowname", all = T)
S4_lines <- merge(S4_lines, S4_13_3_with_snp_names, by = "rowname", all = T)
S4_lines <- merge(S4_lines, S4_13_4_with_snp_names, by = "rowname", all = T)
S4_lines <- merge(S4_lines, S4_16_1_with_snp_names, by = "rowname", all = T)
S4_lines <- merge(S4_lines, S4_16_5_with_snp_names, by = "rowname", all = T)
S4_lines <- merge(S4_lines, S4_17_2_with_snp_names, by = "rowname", all = T)
S4_lines <- merge(S4_lines, S4_17_5_with_snp_names, by = "rowname", all = T)
S4_lines <- merge(S4_lines, S4_19_2_with_snp_names, by = "rowname", all = T)
S4_lines <- merge(S4_lines, S4_19_5_with_snp_names, by = "rowname", all = T)
S4_lines <- merge(S4_lines, S4_20_1_with_snp_names, by = "rowname", all = T)
S4_lines <- merge(S4_lines, S4_20_4_with_snp_names, by = "rowname", all = T)
S4_lines <- merge(S4_lines, S4_21_2_with_snp_names, by = "rowname", all = T)
S4_lines <- merge(S4_lines, S4_21_6_with_snp_names, by = "rowname", all = T)
S4_lines <- merge(S4_lines, S4_23_2_with_snp_names, by = "rowname", all = T)
S4_lines <- merge(S4_lines, S4_23_4_with_snp_names, by = "rowname", all = T)
S4_lines <- merge(S4_lines, S4_26_1_with_snp_names, by = "rowname", all = T)
S4_lines <- merge(S4_lines, S4_26_4_with_snp_names, by = "rowname", all = T)
S4_lines <- merge(S4_lines, S4_27_2_with_snp_names, by = "rowname", all = T)
S4_lines <- merge(S4_lines, S4_29_2_with_snp_names, by = "rowname", all = T)
S4_lines <- merge(S4_lines, S4_29_4_with_snp_names, by = "rowname", all = T)

# Minor allele freqs
S4_lines <- S4_lines %>% 
  column_to_rownames()

n0 <- apply(S4_lines==0,1,sum,na.rm=T)
n1 <- apply(S4_lines==1,1,sum,na.rm=T)
n2 <- apply(S4_lines==2,1,sum,na.rm=T)

n <- n0 + n1 + n2

p <- ((2*n0)+n1)/(2*n)
q <- 1 - p
maf <- pmin(p, q)
mgf <- apply(cbind(n0,n1,n2),1,min) / n

#allele_freqs_S4 <- Propfunc(S4_lines)
allele_freqs_S4 <- data.frame(SNP = rownames(S4_lines), frq = maf)

ggplot(allele_freqs_S4, aes(x = frq)) +
  geom_histogram(binwidth = 0.01)

### S5 gen ##############################################################################################
S5_1_1_with_snp_names <- Line_1_1_het_F %>% 
  rownames_to_column() %>% 
  select(1, 7)

# S5_1_4_with_snp_names <- Line_1_4_het_F %>% 
#   rownames_to_column() %>% 
#   select(1, 7)

S5_6_4_with_snp_names <- Line_6_4_het_F %>% 
  rownames_to_column() %>% 
  select(1, 7)

S5_10_2_with_snp_names <- Line_10_2_het_F %>%
  rownames_to_column() %>%
  select(1, 7)

S5_16_5_with_snp_names <- Line_16_5_het_F %>% 
  rownames_to_column() %>% 
  select(1, 7)

S5_17_2_with_snp_names <- Line_17_2_het_F %>% 
  rownames_to_column() %>% 
  select(1, 7)

S5_17_5_with_snp_names <- Line_17_5_het_F %>% 
  rownames_to_column() %>% 
  select(1, 7)

S5_21_2_with_snp_names <- Line_21_2_het_F %>% 
  rownames_to_column() %>% 
  select(1, 7)

S5_21_6_with_snp_names <- Line_21_6_het_F %>% 
  rownames_to_column() %>% 
  select(1, 7)

S5_23_2_with_snp_names <- Line_23_2_het_F %>% 
  rownames_to_column() %>% 
  select(1, 7)

S5_29_2_with_snp_names <- Line_29_2_het_F %>% 
  rownames_to_column() %>% 
  select(1, 7)

S5_29_4_with_snp_names <- Line_29_4_het_F %>% 
  rownames_to_column() %>% 
  select(1, 7)


S5_lines <- merge(S5_1_1_with_snp_names, S5_6_4_with_snp_names, by = "rowname", all = T)
S5_lines <- merge(S5_lines, S5_10_2_with_snp_names, by = "rowname", all = T)
S5_lines <- merge(S5_lines, S5_17_2_with_snp_names, by = "rowname", all = T)
S5_lines <- merge(S5_lines, S5_17_5_with_snp_names, by = "rowname", all = T)
S5_lines <- merge(S5_lines, S5_21_2_with_snp_names, by = "rowname", all = T)
S5_lines <- merge(S5_lines, S5_21_6_with_snp_names, by = "rowname", all = T)
S5_lines <- merge(S5_lines, S5_23_2_with_snp_names, by = "rowname", all = T)
S5_lines <- merge(S5_lines, S5_29_2_with_snp_names, by = "rowname", all = T)
S5_lines <- merge(S5_lines, S5_29_4_with_snp_names, by = "rowname", all = T)

# Minor allele freqs
S5_lines <- S5_lines %>% 
  column_to_rownames()


#### Combine all gens het F ########################
gt_1_1 <- data.frame(SNP = F_1_1_with_snp_names$rowname, `1_1-F` = F_1_1_with_snp_names$`1_1-F`, 
                     `111-S1` = S1_1_1_with_snp_names$`111-S1`, `111-2-S2` = S2_1_1_with_snp_names$`111-2-S2`, 
                     `111-22-S3` = S3_1_1_with_snp_names$`111-22-S3`, `111-221-S4` = S4_1_1_with_snp_names$`111-221-S4`, 
                     `111-221-S5` = S5_1_1_with_snp_names$`111-221-S5`, check.names = F)


gt_6_1 <- data.frame(SNP = F_6_1_with_snp_names$rowname, `6_1-F` = F_6_1_with_snp_names$`6_1-F`, 
                     `611-S1` = S1_6_1_with_snp_names$`611-S1`, `611-1-S2` = S2_6_1_with_snp_names$`611-1-S2`, 
                     `611-11-S3` = S3_6_1_with_snp_names$`611-11-S3`, `611-111-S4` = S4_6_1_with_snp_names$`611-111-S4`, check.names = F)


gt_6_4 <- data.frame(SNP = F_6_4_with_snp_names$rowname, `6_4-F` = F_6_4_with_snp_names$`6_4-F`, 
                     `646-S1` = S1_6_4_with_snp_names$`646-S1`, `646-4-S2` = S2_6_4_with_snp_names$`646-4-S2`, 
                     `646-45-S3` = S3_6_4_with_snp_names$`646-45-S3`, `646-454-S4` = S4_6_4_with_snp_names$`646-454-S4`, 
                     `646-454-S5` = S5_6_4_with_snp_names$`646-454-S5`, check.names = F)


gt_7_2 <- data.frame(SNP = F_7_2_with_snp_names$rowname, `7_2-F` = F_7_2_with_snp_names$`7_2-F`, 
                     `721-S1` = S1_7_2_with_snp_names$`721-S1`, `721-2-S2` = S2_7_2_with_snp_names$`721-2-S2`, 
                     `721-21-S3` = S3_7_2_with_snp_names$`721-21-S3`, `721-211-S4` = S4_7_2_with_snp_names$`721-211-S4`, check.names = F)


gt_7_4 <- data.frame(SNP = F_7_4_with_snp_names$rowname, `7_4-F` = F_7_4_with_snp_names$`7_4-F`, 
                     `745-S1` = S1_7_4_with_snp_names$`745-S1`, `745-4-S2` = S2_7_4_with_snp_names$`745-4-S2`, 
                     `745-44-S3` = S3_7_4_with_snp_names$`745-44-S3`, `745-444-S4` = S4_7_4_with_snp_names$`745-444-S4`, check.names = F)


gt_8_2 <- data.frame(SNP = F_8_2_with_snp_names$rowname, `8_2-F` = F_8_2_with_snp_names$`8_2-F`, 
                     `821-S1` = S1_8_2_with_snp_names$`821-S1`, `821-1-S2` = S2_8_2_with_snp_names$`821-1-S2`, 
                     `821-11-S3` = S3_8_2_with_snp_names$`821-11-S3`, `821-111-S4` = S4_8_2_with_snp_names$`821-111-S4`, check.names = F)


gt_8_4 <- data.frame(SNP = F_8_4_with_snp_names$rowname, `8_4-F` = F_8_4_with_snp_names$`8_4-F`, 
                     `845-S1` = S1_8_4_with_snp_names$`845-S1`, `845-4-S2` = S2_8_4_with_snp_names$`845-4-S2`, 
                     `845-44-S3` = S3_8_4_with_snp_names$`845-44-S3`, `845-444-S4` = S4_8_4_with_snp_names$`845-444-S4`, check.names = F)


gt_10_2 <- data.frame(SNP = F_10_2_with_snp_names$rowname, `10_2-F` = F_10_2_with_snp_names$`10_2-F`, 
                      `1022-S1` = S1_10_2_with_snp_names$`1022-S1`, `1022-1-S2` = S2_10_2_with_snp_names$`1022-1-S2`, 
                      `1022-11-S3` = S3_10_2_with_snp_names$`1022-11-S3`, `1022-113-S4` = S4_10_2_with_snp_names$`1022-113-S4`,
                      `1022-113-S5` = S5_10_2_with_snp_names$`1022-113-S5`, check.names = F)


gt_10_5 <- data.frame(SNP = F_10_5_with_snp_names$rowname, `10_5-F` = F_10_5_with_snp_names$`10_5-F`, 
                      `1056-S1` = S1_10_5_with_snp_names$`1056-S1`, `1056-4-S2` = S2_10_5_with_snp_names$`1056-4-S2`, 
                      `1056-45-S3` = S3_10_5_with_snp_names$`1056-45-S3`, `1056-454-S4` = S4_10_5_with_snp_names$`1056-454-S4`, check.names = F)


gt_13_3 <- data.frame(SNP = F_13_3_with_snp_names$rowname, `13_3-F` = F_13_3_with_snp_names$`13_3-F`, 
                      `1332-S1` = S1_13_3_with_snp_names$`1332-S1`, `1332-1-S2` = S2_13_3_with_snp_names$`1332-1-S2`, 
                      `1332-11-S3` = S3_13_3_with_snp_names$`1332-11-S3`, `1332-111-S4` = S4_13_3_with_snp_names$`1332-111-S4`, check.names = F)


gt_13_4 <- data.frame(SNP = F_13_4_with_snp_names$rowname, `13_4-F` = F_13_4_with_snp_names$`13_4-F`, 
                      `1345-S1` = S1_13_4_with_snp_names$`1345-S1`, `1345-5-S2` = S2_13_4_with_snp_names$`1345-5-S2`, 
                      `1345-54-S3` = S3_13_4_with_snp_names$`1345-54-S3`, `1345-544-S4` = S4_13_4_with_snp_names$`1345-544-S4`, check.names = F)


gt_16_1 <- data.frame(SNP = F_16_1_with_snp_names$rowname, `16_1-F` = F_16_1_with_snp_names$`16_1-F`, 
                      `1611-S1` = S1_16_1_with_snp_names$`1611-S1`, `1611-2-S2` = S2_16_1_with_snp_names$`1611-2-S2`, 
                      `1611-21-S3` = S3_16_1_with_snp_names$`1611-21-S3`, `1611-211-S4` = S4_16_1_with_snp_names$`1611-211-S4`, check.names = F)


gt_16_5 <- data.frame(SNP = F_16_5_with_snp_names$rowname, `16_5-F` = F_16_5_with_snp_names$`16_5-F`, 
                      `1654-S1` = S1_16_5_with_snp_names$`1654-S1`, `1654-4-S2` = S2_16_5_with_snp_names$`1654-4-S2`, 
                      `1654-45-S3` = S3_16_5_with_snp_names$`1654-45-S3`, `1654-454-S4` = S4_16_5_with_snp_names$`1654-454-S4`, 
                      `1654-454-S5` = S5_16_5_with_snp_names$`1654-454-S5`, check.names = F)


gt_17_2 <- data.frame(SNP = F_17_2_with_snp_names$rowname, `17_2-F` = F_17_2_with_snp_names$`17_2-F`, 
                      `1721-S1` = S1_17_2_with_snp_names$`1721-S1`, `1721-1-S2` = S2_17_2_with_snp_names$`1721-1-S2`, 
                      `1721-11-S3` = S3_17_2_with_snp_names$`1721-11-S3`, `1721-111-S4` = S4_17_2_with_snp_names$`1721-111-S4`, 
                      `1721-111-S5` = S5_17_2_with_snp_names$`1721-111-S5`, check.names = F)


gt_17_5 <- data.frame(SNP = F_17_5_with_snp_names$rowname, `17_5-F` = F_17_5_with_snp_names$`17_5-F`, 
                      `1755-S1` = S1_17_5_with_snp_names$`1755-S1`, `1755-5-S2` = S2_17_5_with_snp_names$`1755-5-S2`, 
                      `1755-54-S3` = S3_17_5_with_snp_names$`1755-54-S3`, `1755-545-S4` = S4_17_5_with_snp_names$`1755-545-S4`, 
                      `1755-545-S5` = S5_17_5_with_snp_names$`1755-545-S5`, check.names = F)


gt_19_2 <- data.frame(SNP = F_19_2_with_snp_names$rowname, `19_2-F` = F_19_2_with_snp_names$`19_2-F`, 
                      `1922-S1` = S1_19_2_with_snp_names$`1922-S1`, `1922-1-S2` = S2_19_2_with_snp_names$`1922-1-S2`, 
                      `1922-11-S3` = S3_19_2_with_snp_names$`1922-11-S3`, `1922-111-S4` = S4_19_2_with_snp_names$`1922-111-S4`, check.names = F)


gt_19_5 <- data.frame(SNP = F_19_5_with_snp_names$rowname, `19_5-F` = F_19_5_with_snp_names$`19_5-F`, 
                      `1955-S1` = S1_19_5_with_snp_names$`1955-S1`, `1955-5-S2` = S2_19_5_with_snp_names$`1955-5-S2`, 
                      `1955-54-S3` = S3_19_5_with_snp_names$`1955-54-S3`, `1955-544-S4` = S4_19_5_with_snp_names$`1955-544-S4`, check.names = F)


gt_20_1 <- data.frame(SNP = F_20_1_with_snp_names$rowname, `20_1-F` = F_20_1_with_snp_names$`20_1-F`, 
                      `2013-S1` = S1_20_1_with_snp_names$`2013-S1`, `2013-1-S2` = S2_20_1_with_snp_names$`2013-1-S2`, 
                      `2013-13-S3` = S3_20_1_with_snp_names$`2013-13-S3`, `2013-131-S4` = S4_20_1_with_snp_names$`2013-131-S4`, check.names = F)


gt_20_4 <- data.frame(SNP = F_20_4_with_snp_names$rowname, `20_4-F` = F_20_4_with_snp_names$`20_4-F`, 
                      `2045-S1` = S1_20_4_with_snp_names$`2045-S1`, `2045-5-S2` = S2_20_4_with_snp_names$`2045-5-S2`, 
                      `2045-54-S3` = S3_20_4_with_snp_names$`2045-54-S3`, `2045-544-S4` = S4_20_4_with_snp_names$`2045-544-S4`, check.names = F)


gt_21_2 <- data.frame(SNP = F_21_2_with_snp_names$rowname, `21_2-F` = F_21_2_with_snp_names$`21_2-F`, 
                      `2121-S1` = S1_21_2_with_snp_names$`2121-S1`, `2121-1-S2` = S2_21_2_with_snp_names$`2121-1-S2`, 
                      `2121-11-S3` = S3_21_2_with_snp_names$`2121-11-S3`, `2121-111-S4` = S4_21_2_with_snp_names$`2121-111-S4`, 
                      `2121-111-S5` = S5_21_2_with_snp_names$`2121-111-S5`, check.names = F)


gt_21_6 <- data.frame(SNP = F_21_6_with_snp_names$rowname, `21_6-F` = F_21_6_with_snp_names$`21_6-F`, 
                      `2165-S1` = S1_21_6_with_snp_names$`2165-S1`, `2165-4-S2` = S2_21_6_with_snp_names$`2165-4-S2`, 
                      `2165-44-S3` = S3_21_6_with_snp_names$`2165-44-S3`, `2165-444-S4` = S4_21_6_with_snp_names$`2165-444-S4`, 
                      `2165-444-S5` = S5_21_6_with_snp_names$`2165-444-S5`, check.names = F)


gt_23_2 <- data.frame(SNP = F_23_2_with_snp_names$rowname, `23_2-F` = F_23_2_with_snp_names$`23_2-F`, 
                      `2323-S1` = S1_23_2_with_snp_names$`2323-S1`, `2323-2-S2` = S2_23_2_with_snp_names$`2323-2-S2`, 
                      `2323-21-S3` = S3_23_2_with_snp_names$`2323-21-S3`, `2323-211-S4` = S4_23_2_with_snp_names$`2323-211-S4`, 
                      `2323-211-S5` = S5_23_2_with_snp_names$`2323-211-S5`, check.names = F)


gt_23_4 <- data.frame(SNP = F_23_4_with_snp_names$rowname, `23_4-F` = F_23_4_with_snp_names$`23_4-F`, 
                      `2344-S1` = S1_23_4_with_snp_names$`2344-S1`, `2344-4-S2` = S2_23_4_with_snp_names$`2344-4-S2`, 
                      `2344-46-S3` = S3_23_4_with_snp_names$`2344-46-S3`, `2344-464-S4` = S4_23_4_with_snp_names$`2344-464-S4`, check.names = F)


gt_26_1 <- data.frame(SNP = F_26_1_with_snp_names$rowname, `26_1-F` = F_26_1_with_snp_names$`26_1-F`, 
                      `2612-S1` = S1_26_1_with_snp_names$`2612-S1`, `2612-1-S2` = S2_26_1_with_snp_names$`2612-1-S2`, 
                      `2612-12-S3` = S3_26_1_with_snp_names$`2612-12-S3`, `2612-121-S4` = S4_26_1_with_snp_names$`2612-121-S4`, check.names = F)


gt_26_4 <- data.frame(SNP = F_26_4_with_snp_names$rowname, `26_4-F` = F_26_4_with_snp_names$`26_4-F`, 
                      `2645-S1` = S1_26_4_with_snp_names$`2645-S1`, `2645-4-S2` = S2_26_4_with_snp_names$`2645-4-S2`, 
                      `2645-46-S3` = S3_26_4_with_snp_names$`2645-46-S3`, `2645-464-S4` = S4_26_4_with_snp_names$`2645-464-S4`, check.names = F)


gt_27_2 <- data.frame(SNP = F_27_2_with_snp_names$rowname, `27_2-F` = F_27_2_with_snp_names$`27_2-F`, 
                      `2721-S1` = S1_27_2_with_snp_names$`2721-S1`, `2721-1-S2` = S2_27_2_with_snp_names$`2721-1-S2`, 
                      `2721-12-S3` = S3_27_2_with_snp_names$`2721-12-S3`, `2721-121-S4` = S4_27_2_with_snp_names$`2721-121-S4`, check.names = F)


gt_29_2 <- data.frame(SNP = F_29_2_with_snp_names$rowname, `29_2-F` = F_29_2_with_snp_names$`29_2-F`, 
                      `2923-S1` = S1_29_2_with_snp_names$`2923-S1`, `2923-1-S2` = S2_29_2_with_snp_names$`2923-1-S2`, 
                      `2923-12-S3` = S3_29_2_with_snp_names$`2923-12-S3`, `2923-121-S4` = S4_29_2_with_snp_names$`2923-121-S4`, 
                      `2923-121-S5` = S5_29_2_with_snp_names$`2923-121-S5`, check.names = F)


gt_29_4 <- data.frame(SNP = F_29_4_with_snp_names$rowname, `29_4-F` = F_29_4_with_snp_names$`29_4-F`, 
                      `2944-S1` = S1_29_4_with_snp_names$`2944-S1`, `2944-5-S2` = S2_29_4_with_snp_names$`2944-5-S2`, 
                      `2944-56-S3` = S3_29_4_with_snp_names$`2944-56-S3`, `2944-565-S4` = S4_29_4_with_snp_names$`2944-565-S4`, 
                      `2944-565-S5` = S5_29_4_with_snp_names$`2944-565-S5`, check.names = F)


# Convert SNP to character
gt_1_1$SNP <- as.character(gt_1_1$SNP)
gt_6_1$SNP <- as.character(gt_6_1$SNP)
gt_6_4$SNP <- as.character(gt_6_4$SNP)
gt_7_2$SNP <- as.character(gt_7_2$SNP)
gt_7_4$SNP <- as.character(gt_7_4$SNP)
gt_8_2$SNP <- as.character(gt_8_2$SNP)
gt_8_4$SNP <- as.character(gt_8_4$SNP)
gt_10_2$SNP <- as.character(gt_10_2$SNP)
gt_10_5$SNP <- as.character(gt_10_5$SNP)
gt_13_3$SNP <- as.character(gt_13_3$SNP)
gt_13_4$SNP <- as.character(gt_13_4$SNP)
gt_16_1$SNP <- as.character(gt_16_1$SNP)
gt_16_5$SNP <- as.character(gt_16_5$SNP)
gt_17_2$SNP <- as.character(gt_17_2$SNP)
gt_17_5$SNP <- as.character(gt_17_5$SNP)
gt_19_2$SNP <- as.character(gt_19_2$SNP)
gt_19_5$SNP <- as.character(gt_19_5$SNP)
gt_20_1$SNP <- as.character(gt_20_1$SNP)
gt_20_4$SNP <- as.character(gt_20_4$SNP)
gt_21_2$SNP <- as.character(gt_21_2$SNP)
gt_21_6$SNP <- as.character(gt_21_6$SNP)
gt_23_2$SNP <- as.character(gt_23_2$SNP)
gt_23_4$SNP <- as.character(gt_23_4$SNP)
gt_26_1$SNP <- as.character(gt_26_1$SNP)
gt_26_4$SNP <- as.character(gt_26_4$SNP)
gt_27_2$SNP <- as.character(gt_27_2$SNP)
gt_29_2$SNP <- as.character(gt_29_2$SNP)
gt_29_4$SNP <- as.character(gt_29_4$SNP)

# Get fixed ref, alt, not fixed
# 1_1
gt_1_1_fixed_ref <- gt_1_1 %>% 
  mutate(fixed = if_else((`1_1-F` == 0 | `111-S1` == 0 | `111-2-S2` == 0 | `111-22-S3` == 0 | 
                            `111-221-S4` == 0 | `111-221-S5` == 0), "fixed_ref", 
                         if_else((`1_1-F` == 2 | `111-S1` == 2 | `111-2-S2` == 2 | `111-22-S3` == 2 | 
                                    `111-221-S4` == 2 | `111-221-S5` == 2), "fixed_alt", "not_fixed"))) %>% 
  filter(fixed == "fixed_ref")

gt_1_1_fixed_alt <- gt_1_1 %>% 
  mutate(fixed = if_else((`1_1-F` == 0 | `111-S1` == 0 | `111-2-S2` == 0 | `111-22-S3` == 0 | 
                            `111-221-S4` == 0 | `111-221-S5` == 0), "fixed_ref", 
                         if_else((`1_1-F` == 2 | `111-S1` == 2 | `111-2-S2` == 2 | `111-22-S3` == 2 | 
                                    `111-221-S4` == 2 | `111-221-S5` == 2), "fixed_alt", "not_fixed"))) %>% 
  filter(fixed == "fixed_alt")

gt_1_1_not_fixed <- gt_1_1 %>% 
  mutate(fixed = if_else((`1_1-F` == 0 | `111-S1` == 0 | `111-2-S2` == 0 | `111-22-S3` == 0 | 
                            `111-221-S4` == 0 | `111-221-S5` == 0), "fixed_ref", 
                         if_else((`1_1-F` == 2 | `111-S1` == 2 | `111-2-S2` == 2 | `111-22-S3` == 2 | 
                                    `111-221-S4` == 2 | `111-221-S5` == 2), "fixed_alt", "not_fixed"))) %>% 
  filter(fixed == "not_fixed")

# 6_1
gt_6_1_fixed_ref <- gt_6_1 %>% 
  mutate(fixed = if_else((`6_1-F` == 0 | `611-S1` == 0 | `611-1-S2` == 0 | `611-11-S3` == 0 | 
                            `611-111-S4` == 0), "fixed_ref", 
                         if_else((`6_1-F` == 2 | `611-S1` == 2 | `611-1-S2` == 2 | `611-11-S3` == 2 | 
                                    `611-111-S4` == 2), "fixed_alt", "not_fixed"))) %>% 
  filter(fixed == "fixed_ref")

gt_6_1_fixed_alt <- gt_6_1 %>% 
  mutate(fixed = if_else((`6_1-F` == 0 | `611-S1` == 0 | `611-1-S2` == 0 | `611-11-S3` == 0 | 
                            `611-111-S4` == 0), "fixed_ref", 
                         if_else((`6_1-F` == 2 | `611-S1` == 2 | `611-1-S2` == 2 | `611-11-S3` == 2 | 
                                    `611-111-S4` == 2), "fixed_alt", "not_fixed"))) %>% 
  filter(fixed == "fixed_alt")

gt_6_1_not_fixed <- gt_6_1 %>% 
  mutate(fixed = if_else((`6_1-F` == 0 | `611-S1` == 0 | `611-1-S2` == 0 | `611-11-S3` == 0 | 
                            `611-111-S4` == 0), "fixed_ref", 
                         if_else((`6_1-F` == 2 | `611-S1` == 2 | `611-1-S2` == 2 | `611-11-S3` == 2 | 
                                    `611-111-S4` == 2), "fixed_alt", "not_fixed"))) %>% 
  filter(fixed == "not_fixed")

# 6_4
gt_6_4_fixed_ref <- gt_6_4 %>% 
  mutate(fixed = if_else((`6_4-F` == 0 | `646-S1` == 0 | `646-4-S2` == 0 | `646-45-S3` == 0 | 
                            `646-454-S4` == 0 | `646-454-S5` == 0), "fixed_ref", 
                         if_else((`6_4-F` == 2 | `646-S1` == 2 | `646-4-S2` == 2 | `646-45-S3` == 2 | 
                                    `646-454-S4` == 2 | `646-454-S4` == 2), "fixed_alt", "not_fixed"))) %>% 
  filter(fixed == "fixed_ref")

gt_6_4_fixed_alt <- gt_6_4 %>% 
  mutate(fixed = if_else((`6_4-F` == 0 | `646-S1` == 0 | `646-4-S2` == 0 | `646-45-S3` == 0 | 
                            `646-454-S4` == 0 | `646-454-S5` == 0), "fixed_ref", 
                         if_else((`6_4-F` == 2 | `646-S1` == 2 | `646-4-S2` == 2 | `646-45-S3` == 2 | 
                                    `646-454-S4` == 2 | `646-454-S4` == 2), "fixed_alt", "not_fixed"))) %>% 
  filter(fixed == "fixed_alt")

gt_6_4_not_fixed <- gt_6_4 %>% 
  mutate(fixed = if_else((`6_4-F` == 0 | `646-S1` == 0 | `646-4-S2` == 0 | `646-45-S3` == 0 | 
                            `646-454-S4` == 0 | `646-454-S5` == 0), "fixed_ref", 
                         if_else((`6_4-F` == 2 | `646-S1` == 2 | `646-4-S2` == 2 | `646-45-S3` == 2 | 
                                    `646-454-S4` == 2 | `646-454-S4` == 2), "fixed_alt", "not_fixed"))) %>% 
  filter(fixed == "not_fixed")

# 7_2
gt_7_2_fixed_ref <- gt_7_2 %>% 
  mutate(fixed = if_else((`7_2-F` == 0 | `721-S1` == 0 | `721-2-S2` == 0 | `721-21-S3` == 0 | 
                            `721-211-S4` == 0), "fixed_ref", 
                         if_else((`7_2-F` == 2 | `721-S1` == 2 | `721-2-S2` == 2 | `721-21-S3` == 2 | 
                                    `721-211-S4` == 2), "fixed_alt", "not_fixed"))) %>% 
  filter(fixed == "fixed_ref")

gt_7_2_fixed_alt <- gt_7_2 %>% 
  mutate(fixed = if_else((`7_2-F` == 0 | `721-S1` == 0 | `721-2-S2` == 0 | `721-21-S3` == 0 | 
                            `721-211-S4` == 0), "fixed_ref", 
                         if_else((`7_2-F` == 2 | `721-S1` == 2 | `721-2-S2` == 2 | `721-21-S3` == 2 | 
                                    `721-211-S4` == 2), "fixed_alt", "not_fixed"))) %>% 
  filter(fixed == "fixed_alt")

gt_7_2_not_fixed <- gt_7_2 %>% 
  mutate(fixed = if_else((`7_2-F` == 0 | `721-S1` == 0 | `721-2-S2` == 0 | `721-21-S3` == 0 | 
                            `721-211-S4` == 0), "fixed_ref", 
                         if_else((`7_2-F` == 2 | `721-S1` == 2 | `721-2-S2` == 2 | `721-21-S3` == 2 | 
                                    `721-211-S4` == 2), "fixed_alt", "not_fixed"))) %>% 
  filter(fixed == "not_fixed")

# 7_4
gt_7_4_fixed_ref <- gt_7_4 %>% 
  mutate(fixed = if_else((`7_4-F` == 0 | `745-S1` == 0 | `745-4-S2` == 0 | `745-44-S3` == 0 | 
                            `745-444-S4` == 0), "fixed_ref", 
                         if_else((`7_4-F` == 2 | `745-S1` == 2 | `745-4-S2` == 2 | `745-44-S3` == 2 | 
                                    `745-444-S4` == 2), "fixed_alt", "not_fixed"))) %>% 
  filter(fixed == "fixed_ref")

gt_7_4_fixed_alt <- gt_7_4 %>% 
  mutate(fixed = if_else((`7_4-F` == 0 | `745-S1` == 0 | `745-4-S2` == 0 | `745-44-S3` == 0 | 
                            `745-444-S4` == 0), "fixed_ref", 
                         if_else((`7_4-F` == 2 | `745-S1` == 2 | `745-4-S2` == 2 | `745-44-S3` == 2 | 
                                    `745-444-S4` == 2), "fixed_alt", "not_fixed"))) %>% 
  filter(fixed == "fixed_alt")

gt_7_4_not_fixed <- gt_7_4 %>% 
  mutate(fixed = if_else((`7_4-F` == 0 | `745-S1` == 0 | `745-4-S2` == 0 | `745-44-S3` == 0 | 
                            `745-444-S4` == 0), "fixed_ref", 
                         if_else((`7_4-F` == 2 | `745-S1` == 2 | `745-4-S2` == 2 | `745-44-S3` == 2 | 
                                    `745-444-S4` == 2), "fixed_alt", "not_fixed"))) %>% 
  filter(fixed == "not_fixed")

# 8_2
gt_8_2_fixed_ref <- gt_8_2 %>% 
  mutate(fixed = if_else((`8_2-F` == 0 | `821-S1` == 0 | `821-1-S2` == 0 | `821-11-S3` == 0 | 
                            `821-111-S4` == 0), "fixed_ref", 
                         if_else((`8_2-F` == 2 | `821-S1` == 2 | `821-1-S2` == 2 | `821-11-S3` == 2 | 
                                    `821-111-S4` == 2), "fixed_alt", "not_fixed"))) %>% 
  filter(fixed == "fixed_ref")

gt_8_2_fixed_alt <- gt_8_2 %>% 
  mutate(fixed = if_else((`8_2-F` == 0 | `821-S1` == 0 | `821-1-S2` == 0 | `821-11-S3` == 0 | 
                            `821-111-S4` == 0), "fixed_ref", 
                         if_else((`8_2-F` == 2 | `821-S1` == 2 | `821-1-S2` == 2 | `821-11-S3` == 2 | 
                                    `821-111-S4` == 2), "fixed_alt", "not_fixed"))) %>% 
  filter(fixed == "fixed_alt")

gt_8_2_not_fixed <- gt_8_2 %>% 
  mutate(fixed = if_else((`8_2-F` == 0 | `821-S1` == 0 | `821-1-S2` == 0 | `821-11-S3` == 0 | 
                            `821-111-S4` == 0), "fixed_ref", 
                         if_else((`8_2-F` == 2 | `821-S1` == 2 | `821-1-S2` == 2 | `821-11-S3` == 2 | 
                                    `821-111-S4` == 2), "fixed_alt", "not_fixed"))) %>% 
  filter(fixed == "not_fixed")

# 8_4
gt_8_4_fixed_ref <- gt_8_4 %>% 
  mutate(fixed = if_else((`8_4-F` == 0 | `845-S1` == 0 | `845-4-S2` == 0 | `845-44-S3` == 0 | 
                            `845-444-S4` == 0), "fixed_ref", 
                         if_else((`8_4-F` == 2 | `845-S1` == 2 | `845-4-S2` == 2 | `845-44-S3` == 2 | 
                                    `845-444-S4` == 2), "fixed_alt", "not_fixed"))) %>% 
  filter(fixed == "fixed_ref")

gt_8_4_fixed_alt <- gt_8_4 %>% 
  mutate(fixed = if_else((`8_4-F` == 0 | `845-S1` == 0 | `845-4-S2` == 0 | `845-44-S3` == 0 | 
                            `845-444-S4` == 0), "fixed_ref", 
                         if_else((`8_4-F` == 2 | `845-S1` == 2 | `845-4-S2` == 2 | `845-44-S3` == 2 | 
                                    `845-444-S4` == 2), "fixed_alt", "not_fixed"))) %>% 
  filter(fixed == "fixed_alt")

gt_8_4_not_fixed <- gt_8_4 %>% 
  mutate(fixed = if_else((`8_4-F` == 0 | `845-S1` == 0 | `845-4-S2` == 0 | `845-44-S3` == 0 | 
                            `845-444-S4` == 0), "fixed_ref", 
                         if_else((`8_4-F` == 2 | `845-S1` == 2 | `845-4-S2` == 2 | `845-44-S3` == 2 | 
                                    `845-444-S4` == 2), "fixed_alt", "not_fixed"))) %>% 
  filter(fixed == "not_fixed")

# 10_2
gt_10_2_fixed_ref <- gt_10_2 %>% 
  mutate(fixed = if_else((`10_2-F` == 0 | `1022-S1` == 0 | `1022-1-S2` == 0 | `1022-11-S3` == 0 | 
                            `1022-113-S4` == 0 | `1022-113-S5` == 0), "fixed_ref", 
                         if_else((`10_2-F` == 2 | `1022-S1` == 2 | `1022-1-S2` == 2 | `1022-11-S3` == 2 | 
                                    `1022-113-S4` == 2 | `1022-113-S5` == 2), "fixed_alt", "not_fixed"))) %>% 
  filter(fixed == "fixed_ref")

gt_10_2_fixed_alt <- gt_10_2 %>% 
  mutate(fixed = if_else((`10_2-F` == 0 | `1022-S1` == 0 | `1022-1-S2` == 0 | `1022-11-S3` == 0 | 
                            `1022-113-S4` == 0 | `1022-113-S5` == 0), "fixed_ref", 
                         if_else((`10_2-F` == 2 | `1022-S1` == 2 | `1022-1-S2` == 2 | `1022-11-S3` == 2 | 
                                    `1022-113-S4` == 2 | `1022-113-S5` == 2), "fixed_alt", "not_fixed"))) %>% 
  filter(fixed == "fixed_alt")

gt_10_2_not_fixed <- gt_10_2 %>% 
  mutate(fixed = if_else((`10_2-F` == 0 | `1022-S1` == 0 | `1022-1-S2` == 0 | `1022-11-S3` == 0 | 
                            `1022-113-S4` == 0 | `1022-113-S5` == 0), "fixed_ref", 
                         if_else((`10_2-F` == 2 | `1022-S1` == 2 | `1022-1-S2` == 2 | `1022-11-S3` == 2 | 
                                    `1022-113-S4` == 2 | `1022-113-S5` == 2), "fixed_alt", "not_fixed"))) %>%
  filter(fixed == "not_fixed")

# 10_5
gt_10_5_fixed_ref <- gt_10_5 %>% 
  mutate(fixed = if_else((`10_5-F` == 0 | `1056-S1` == 0 | `1056-4-S2` == 0 | `1056-45-S3` == 0 | 
                            `1056-454-S4` == 0), "fixed_ref", 
                         if_else((`10_5-F` == 2 | `1056-S1` == 2 | `1056-4-S2` == 2 | `1056-45-S3` == 2 | 
                                    `1056-454-S4` == 2), "fixed_alt", "not_fixed"))) %>% 
  filter(fixed == "fixed_ref")

gt_10_5_fixed_alt <- gt_10_5 %>% 
  mutate(fixed = if_else((`10_5-F` == 0 | `1056-S1` == 0 | `1056-4-S2` == 0 | `1056-45-S3` == 0 | 
                            `1056-454-S4` == 0), "fixed_ref", 
                         if_else((`10_5-F` == 2 | `1056-S1` == 2 | `1056-4-S2` == 2 | `1056-45-S3` == 2 | 
                                    `1056-454-S4` == 2), "fixed_alt", "not_fixed"))) %>% 
  filter(fixed == "fixed_alt")

gt_10_5_not_fixed <- gt_10_5 %>% 
  mutate(fixed = if_else((`10_5-F` == 0 | `1056-S1` == 0 | `1056-4-S2` == 0 | `1056-45-S3` == 0 | 
                            `1056-454-S4` == 0), "fixed_ref", 
                         if_else((`10_5-F` == 2 | `1056-S1` == 2 | `1056-4-S2` == 2 | `1056-45-S3` == 2 | 
                                    `1056-454-S4` == 2), "fixed_alt", "not_fixed"))) %>% 
  filter(fixed == "not_fixed")

# 13_3
gt_13_3_fixed_ref <- gt_13_3 %>% 
  mutate(fixed = if_else((`13_3-F` == 0 | `1332-S1` == 0 | `1332-1-S2` == 0 | `1332-11-S3` == 0 | 
                            `1332-111-S4` == 0), "fixed_ref", 
                         if_else((`13_3-F` == 2 | `1332-S1` == 2 | `1332-1-S2` == 2 | `1332-11-S3` == 2 | 
                                    `1332-111-S4` == 2), "fixed_alt", "not_fixed"))) %>% 
  filter(fixed == "fixed_ref")

gt_13_3_fixed_alt <- gt_13_3 %>% 
  mutate(fixed = if_else((`13_3-F` == 0 | `1332-S1` == 0 | `1332-1-S2` == 0 | `1332-11-S3` == 0 | 
                            `1332-111-S4` == 0), "fixed_ref", 
                         if_else((`13_3-F` == 2 | `1332-S1` == 2 | `1332-1-S2` == 2 | `1332-11-S3` == 2 | 
                                    `1332-111-S4` == 2), "fixed_alt", "not_fixed"))) %>% 
  filter(fixed == "fixed_alt")

gt_13_3_not_fixed <- gt_13_3 %>% 
  mutate(fixed = if_else((`13_3-F` == 0 | `1332-S1` == 0 | `1332-1-S2` == 0 | `1332-11-S3` == 0 | 
                            `1332-111-S4` == 0), "fixed_ref", 
                         if_else((`13_3-F` == 2 | `1332-S1` == 2 | `1332-1-S2` == 2 | `1332-11-S3` == 2 | 
                                    `1332-111-S4` == 2), "fixed_alt", "not_fixed"))) %>% 
  filter(fixed == "not_fixed")

# 13_4
gt_13_4_fixed_ref <- gt_13_4 %>% 
  mutate(fixed = if_else((`13_4-F` == 0 | `1345-S1` == 0 | `1345-5-S2` == 0 | `1345-54-S3` == 0 | 
                            `1345-544-S4` == 0), "fixed_ref", 
                         if_else((`13_4-F` == 2 | `1345-S1` == 2 | `1345-5-S2` == 2 | `1345-54-S3` == 2 | 
                                    `1345-544-S4` == 2), "fixed_alt", "not_fixed"))) %>% 
  filter(fixed == "fixed_ref")

gt_13_4_fixed_alt <- gt_13_4 %>% 
  mutate(fixed = if_else((`13_4-F` == 0 | `1345-S1` == 0 | `1345-5-S2` == 0 | `1345-54-S3` == 0 | 
                            `1345-544-S4` == 0), "fixed_ref", 
                         if_else((`13_4-F` == 2 | `1345-S1` == 2 | `1345-5-S2` == 2 | `1345-54-S3` == 2 | 
                                    `1345-544-S4` == 2), "fixed_alt", "not_fixed"))) %>% 
  filter(fixed == "fixed_alt")

gt_13_4_not_fixed <- gt_13_4 %>% 
  mutate(fixed = if_else((`13_4-F` == 0 | `1345-S1` == 0 | `1345-5-S2` == 0 | `1345-54-S3` == 0 | 
                            `1345-544-S4` == 0), "fixed_ref", 
                         if_else((`13_4-F` == 2 | `1345-S1` == 2 | `1345-5-S2` == 2 | `1345-54-S3` == 2 | 
                                    `1345-544-S4` == 2), "fixed_alt", "not_fixed"))) %>% 
  filter(fixed == "not_fixed")

# 16_1
gt_16_1_fixed_ref <- gt_16_1 %>% 
  mutate(fixed = if_else((`16_1-F` == 0 | `1611-S1` == 0 | `1611-2-S2` == 0 | `1611-21-S3` == 0 | 
                            `1611-211-S4` == 0), "fixed_ref", 
                         if_else((`16_1-F` == 2 | `1611-S1` == 2 | `1611-2-S2` == 2 | `1611-21-S3` == 2 | 
                                    `1611-211-S4` == 2), "fixed_alt", "not_fixed"))) %>% 
  filter(fixed == "fixed_ref")

gt_16_1_fixed_alt <- gt_16_1 %>% 
  mutate(fixed = if_else((`16_1-F` == 0 | `1611-S1` == 0 | `1611-2-S2` == 0 | `1611-21-S3` == 0 | 
                            `1611-211-S4` == 0), "fixed_ref", 
                         if_else((`16_1-F` == 2 | `1611-S1` == 2 | `1611-2-S2` == 2 | `1611-21-S3` == 2 | 
                                    `1611-211-S4` == 2), "fixed_alt", "not_fixed"))) %>% 
  filter(fixed == "fixed_alt")

gt_16_1_not_fixed <- gt_16_1 %>% 
  mutate(fixed = if_else((`16_1-F` == 0 | `1611-S1` == 0 | `1611-2-S2` == 0 | `1611-21-S3` == 0 | 
                            `1611-211-S4` == 0), "fixed_ref", 
                         if_else((`16_1-F` == 2 | `1611-S1` == 2 | `1611-2-S2` == 2 | `1611-21-S3` == 2 | 
                                    `1611-211-S4` == 2), "fixed_alt", "not_fixed"))) %>% 
  filter(fixed == "not_fixed")

# 16_5
gt_16_5_fixed_ref <- gt_16_5 %>% 
  mutate(fixed = if_else((`16_5-F` == 0 | `1654-S1` == 0 | `1654-4-S2` == 0 | `1654-45-S3` == 0 | 
                            `1654-454-S4` == 0 | `1654-454-S5` == 0), "fixed_ref", 
                         if_else((`16_5-F` == 2 | `1654-S1` == 2 | `1654-4-S2` == 2 | `1654-45-S3` == 2 | 
                                    `1654-454-S4` == 2 | `1654-454-S5` == 2), "fixed_alt", "not_fixed"))) %>% 
  filter(fixed == "fixed_ref")

gt_16_5_fixed_alt <- gt_16_5 %>% 
  mutate(fixed = if_else((`16_5-F` == 0 | `1654-S1` == 0 | `1654-4-S2` == 0 | `1654-45-S3` == 0 | 
                            `1654-454-S4` == 0 | `1654-454-S5` == 0), "fixed_ref", 
                         if_else((`16_5-F` == 2 | `1654-S1` == 2 | `1654-4-S2` == 2 | `1654-45-S3` == 2 | 
                                    `1654-454-S4` == 2 | `1654-454-S5` == 2), "fixed_alt", "not_fixed"))) %>% 
  filter(fixed == "fixed_alt")

gt_16_5_not_fixed <- gt_16_5 %>% 
  mutate(fixed = if_else((`16_5-F` == 0 | `1654-S1` == 0 | `1654-4-S2` == 0 | `1654-45-S3` == 0 | 
                            `1654-454-S4` == 0 | `1654-454-S5` == 0), "fixed_ref", 
                         if_else((`16_5-F` == 2 | `1654-S1` == 2 | `1654-4-S2` == 2 | `1654-45-S3` == 2 | 
                                    `1654-454-S4` == 2 | `1654-454-S5` == 2), "fixed_alt", "not_fixed"))) %>%
  filter(fixed == "not_fixed")

# 17_2
gt_17_2_fixed_ref <- gt_17_2 %>% 
  mutate(fixed = if_else((`17_2-F` == 0 | `1721-S1` == 0 | `1721-1-S2` == 0 | `1721-11-S3` == 0 | 
                            `1721-111-S4` == 0 | `1721-111-S5` == 0), "fixed_ref", 
                         if_else((`17_2-F` == 2 | `1721-S1` == 2 | `1721-1-S2` == 2 | `1721-11-S3` == 2 | 
                                    `1721-111-S4` == 2 | `1721-111-S5` == 2), "fixed_alt", "not_fixed"))) %>% 
  filter(fixed == "fixed_ref")

gt_17_2_fixed_alt <- gt_17_2 %>% 
  mutate(fixed = if_else((`17_2-F` == 0 | `1721-S1` == 0 | `1721-1-S2` == 0 | `1721-11-S3` == 0 | 
                            `1721-111-S4` == 0 | `1721-111-S5` == 0), "fixed_ref", 
                         if_else((`17_2-F` == 2 | `1721-S1` == 2 | `1721-1-S2` == 2 | `1721-11-S3` == 2 | 
                                    `1721-111-S4` == 2 | `1721-111-S5` == 2), "fixed_alt", "not_fixed"))) %>% 
  filter(fixed == "fixed_alt")

gt_17_2_not_fixed <- gt_17_2 %>% 
  mutate(fixed = if_else((`17_2-F` == 0 | `1721-S1` == 0 | `1721-1-S2` == 0 | `1721-11-S3` == 0 | 
                            `1721-111-S4` == 0 | `1721-111-S5` == 0), "fixed_ref", 
                         if_else((`17_2-F` == 2 | `1721-S1` == 2 | `1721-1-S2` == 2 | `1721-11-S3` == 2 | 
                                    `1721-111-S4` == 2 | `1721-111-S5` == 2), "fixed_alt", "not_fixed"))) %>% 
  filter(fixed == "not_fixed")

# 17_5
gt_17_5_fixed_ref <- gt_17_5 %>% 
  mutate(fixed = if_else((`17_5-F` == 0 | `1755-S1` == 0 | `1755-5-S2` == 0 | `1755-54-S3` == 0 | 
                            `1755-545-S4` == 0 | `1755-545-S5` == 0), "fixed_ref", 
                         if_else((`17_5-F` == 2 | `1755-S1` == 2 | `1755-5-S2` == 2 | `1755-54-S3` == 2 | 
                                    `1755-545-S4` == 2 | `1755-545-S5` == 2), "fixed_alt", "not_fixed"))) %>% 
  filter(fixed == "fixed_ref")

gt_17_5_fixed_alt <- gt_17_5 %>% 
  mutate(fixed = if_else((`17_5-F` == 0 | `1755-S1` == 0 | `1755-5-S2` == 0 | `1755-54-S3` == 0 | 
                            `1755-545-S4` == 0 | `1755-545-S5` == 0), "fixed_ref", 
                         if_else((`17_5-F` == 2 | `1755-S1` == 2 | `1755-5-S2` == 2 | `1755-54-S3` == 2 | 
                                    `1755-545-S4` == 2 | `1755-545-S5` == 2), "fixed_alt", "not_fixed"))) %>% 
  filter(fixed == "fixed_alt")

gt_17_5_not_fixed <- gt_17_5 %>% 
  mutate(fixed = if_else((`17_5-F` == 0 | `1755-S1` == 0 | `1755-5-S2` == 0 | `1755-54-S3` == 0 | 
                            `1755-545-S4` == 0 | `1755-545-S5` == 0), "fixed_ref", 
                         if_else((`17_5-F` == 2 | `1755-S1` == 2 | `1755-5-S2` == 2 | `1755-54-S3` == 2 | 
                                    `1755-545-S4` == 2 | `1755-545-S5` == 2), "fixed_alt", "not_fixed"))) %>% 
  filter(fixed == "not_fixed")

# 19_2
gt_19_2_fixed_ref <- gt_19_2 %>% 
  mutate(fixed = if_else((`19_2-F` == 0 | `1922-S1` == 0 | `1922-1-S2` == 0 | `1922-11-S3` == 0 | 
                            `1922-111-S4` == 0), "fixed_ref", 
                         if_else((`19_2-F` == 2 | `1922-S1` == 2 | `1922-1-S2` == 2 | `1922-11-S3` == 2 | 
                                    `1922-111-S4` == 2), "fixed_alt", "not_fixed"))) %>% 
  filter(fixed == "fixed_ref")

gt_19_2_fixed_alt <- gt_19_2 %>% 
  mutate(fixed = if_else((`19_2-F` == 0 | `1922-S1` == 0 | `1922-1-S2` == 0 | `1922-11-S3` == 0 | 
                            `1922-111-S4` == 0), "fixed_ref", 
                         if_else((`19_2-F` == 2 | `1922-S1` == 2 | `1922-1-S2` == 2 | `1922-11-S3` == 2 | 
                                    `1922-111-S4` == 2), "fixed_alt", "not_fixed"))) %>% 
  filter(fixed == "fixed_alt")

gt_19_2_not_fixed <- gt_19_2 %>% 
  mutate(fixed = if_else((`19_2-F` == 0 | `1922-S1` == 0 | `1922-1-S2` == 0 | `1922-11-S3` == 0 | 
                            `1922-111-S4` == 0), "fixed_ref", 
                         if_else((`19_2-F` == 2 | `1922-S1` == 2 | `1922-1-S2` == 2 | `1922-11-S3` == 2 | 
                                    `1922-111-S4` == 2), "fixed_alt", "not_fixed"))) %>% 
  filter(fixed == "not_fixed")

# 19_5
gt_19_5_fixed_ref <- gt_19_5 %>% 
  mutate(fixed = if_else((`19_5-F` == 0 | `1955-S1` == 0 | `1955-5-S2` == 0 | `1955-54-S3` == 0 | 
                            `1955-544-S4` == 0), "fixed_ref", 
                         if_else((`19_5-F` == 2 | `1955-S1` == 2 | `1955-5-S2` == 2 | `1955-54-S3` == 2 | 
                                    `1955-544-S4` == 2), "fixed_alt", "not_fixed"))) %>% 
  filter(fixed == "fixed_ref")

gt_19_5_fixed_alt <- gt_19_5 %>% 
  mutate(fixed = if_else((`19_5-F` == 0 | `1955-S1` == 0 | `1955-5-S2` == 0 | `1955-54-S3` == 0 | 
                            `1955-544-S4` == 0), "fixed_ref", 
                         if_else((`19_5-F` == 2 | `1955-S1` == 2 | `1955-5-S2` == 2 | `1955-54-S3` == 2 | 
                                    `1955-544-S4` == 2), "fixed_alt", "not_fixed"))) %>% 
  filter(fixed == "fixed_alt")

gt_19_5_not_fixed <- gt_19_5 %>% 
  mutate(fixed = if_else((`19_5-F` == 0 | `1955-S1` == 0 | `1955-5-S2` == 0 | `1955-54-S3` == 0 | 
                            `1955-544-S4` == 0), "fixed_ref", 
                         if_else((`19_5-F` == 2 | `1955-S1` == 2 | `1955-5-S2` == 2 | `1955-54-S3` == 2 | 
                                    `1955-544-S4` == 2), "fixed_alt", "not_fixed"))) %>% 
  filter(fixed == "not_fixed")

# 20_1
gt_20_1_fixed_ref <- gt_20_1 %>% 
  mutate(fixed = if_else((`20_1-F` == 0 | `2013-S1` == 0 | `2013-1-S2` == 0 | `2013-13-S3` == 0 | 
                            `2013-131-S4` == 0), "fixed_ref", 
                         if_else((`20_1-F` == 2 | `2013-S1` == 2 | `2013-1-S2` == 2 | `2013-13-S3` == 2 | 
                                    `2013-131-S4` == 2), "fixed_alt", "not_fixed"))) %>% 
  filter(fixed == "fixed_ref")

gt_20_1_fixed_alt <- gt_20_1 %>% 
  mutate(fixed = if_else((`20_1-F` == 0 | `2013-S1` == 0 | `2013-1-S2` == 0 | `2013-13-S3` == 0 | 
                            `2013-131-S4` == 0), "fixed_ref", 
                         if_else((`20_1-F` == 2 | `2013-S1` == 2 | `2013-1-S2` == 2 | `2013-13-S3` == 2 | 
                                    `2013-131-S4` == 2), "fixed_alt", "not_fixed"))) %>% 
  filter(fixed == "fixed_alt")

gt_20_1_not_fixed <- gt_20_1 %>% 
  mutate(fixed = if_else((`20_1-F` == 0 | `2013-S1` == 0 | `2013-1-S2` == 0 | `2013-13-S3` == 0 | 
                            `2013-131-S4` == 0), "fixed_ref", 
                         if_else((`20_1-F` == 2 | `2013-S1` == 2 | `2013-1-S2` == 2 | `2013-13-S3` == 2 | 
                                    `2013-131-S4` == 2), "fixed_alt", "not_fixed"))) %>% 
  filter(fixed == "not_fixed")

# 20_4
gt_20_4_fixed_ref <- gt_20_4 %>% 
  mutate(fixed = if_else((`20_4-F` == 0 | `2045-S1` == 0 | `2045-5-S2` == 0 | `2045-54-S3` == 0 | 
                            `2045-544-S4` == 0), "fixed_ref", 
                         if_else((`20_4-F` == 2 | `2045-S1` == 2 | `2045-5-S2` == 2 | `2045-54-S3` == 2 | 
                                    `2045-544-S4` == 2), "fixed_alt", "not_fixed"))) %>% 
  filter(fixed == "fixed_ref")

gt_20_4_fixed_alt <- gt_20_4 %>% 
  mutate(fixed = if_else((`20_4-F` == 0 | `2045-S1` == 0 | `2045-5-S2` == 0 | `2045-54-S3` == 0 | 
                            `2045-544-S4` == 0), "fixed_ref", 
                         if_else((`20_4-F` == 2 | `2045-S1` == 2 | `2045-5-S2` == 2 | `2045-54-S3` == 2 | 
                                    `2045-544-S4` == 2), "fixed_alt", "not_fixed"))) %>% 
  filter(fixed == "fixed_alt")

gt_20_4_not_fixed <- gt_20_4 %>% 
  mutate(fixed = if_else((`20_4-F` == 0 | `2045-S1` == 0 | `2045-5-S2` == 0 | `2045-54-S3` == 0 | 
                            `2045-544-S4` == 0), "fixed_ref", 
                         if_else((`20_4-F` == 2 | `2045-S1` == 2 | `2045-5-S2` == 2 | `2045-54-S3` == 2 | 
                                    `2045-544-S4` == 2), "fixed_alt", "not_fixed"))) %>% 
  filter(fixed == "not_fixed")

# 21_2
gt_21_2_fixed_ref <- gt_21_2 %>% 
  mutate(fixed = if_else((`21_2-F` == 0 | `2121-S1` == 0 | `2121-1-S2` == 0 | `2121-11-S3` == 0 | 
                            `2121-111-S4` == 0 | `2121-111-S5` == 0), "fixed_ref", 
                         if_else((`21_2-F` == 2 | `2121-S1` == 2 | `2121-1-S2` == 2 | `2121-11-S3` == 2 | 
                                    `2121-111-S4` == 2 | `2121-111-S5` == 2), "fixed_alt", "not_fixed"))) %>% 
  filter(fixed == "fixed_ref")

gt_21_2_fixed_alt <- gt_21_2 %>% 
  mutate(fixed = if_else((`21_2-F` == 0 | `2121-S1` == 0 | `2121-1-S2` == 0 | `2121-11-S3` == 0 | 
                            `2121-111-S4` == 0 | `2121-111-S5` == 0), "fixed_ref", 
                         if_else((`21_2-F` == 2 | `2121-S1` == 2 | `2121-1-S2` == 2 | `2121-11-S3` == 2 | 
                                    `2121-111-S4` == 2 | `2121-111-S5` == 2), "fixed_alt", "not_fixed"))) %>% 
  filter(fixed == "fixed_alt")

gt_21_2_not_fixed <- gt_21_2 %>% 
  mutate(fixed = if_else((`21_2-F` == 0 | `2121-S1` == 0 | `2121-1-S2` == 0 | `2121-11-S3` == 0 | 
                            `2121-111-S4` == 0 | `2121-111-S5` == 0), "fixed_ref", 
                         if_else((`21_2-F` == 2 | `2121-S1` == 2 | `2121-1-S2` == 2 | `2121-11-S3` == 2 | 
                                    `2121-111-S4` == 2 | `2121-111-S5` == 2), "fixed_alt", "not_fixed"))) %>% 
  filter(fixed == "not_fixed")

# 21_6
gt_21_6_fixed_ref <- gt_21_6 %>% 
  mutate(fixed = if_else((`21_6-F` == 0 | `2165-S1` == 0 | `2165-4-S2` == 0 | `2165-44-S3` == 0 | 
                            `2165-444-S4` == 0 | `2165-444-S5` == 0), "fixed_ref", 
                         if_else((`21_6-F` == 2 | `2165-S1` == 2 | `2165-4-S2` == 2 | `2165-44-S3` == 2 | 
                                    `2165-444-S4` == 2 | `2165-444-S5` == 2), "fixed_alt", "not_fixed"))) %>%  
  filter(fixed == "fixed_ref")

gt_21_6_fixed_alt <- gt_21_6 %>% 
  mutate(fixed = if_else((`21_6-F` == 0 | `2165-S1` == 0 | `2165-4-S2` == 0 | `2165-44-S3` == 0 | 
                            `2165-444-S4` == 0 | `2165-444-S5` == 0), "fixed_ref", 
                         if_else((`21_6-F` == 2 | `2165-S1` == 2 | `2165-4-S2` == 2 | `2165-44-S3` == 2 | 
                                    `2165-444-S4` == 2 | `2165-444-S5` == 2), "fixed_alt", "not_fixed"))) %>% 
  filter(fixed == "fixed_alt")

gt_21_6_not_fixed <- gt_21_6 %>% 
  mutate(fixed = if_else((`21_6-F` == 0 | `2165-S1` == 0 | `2165-4-S2` == 0 | `2165-44-S3` == 0 | 
                            `2165-444-S4` == 0 | `2165-444-S5` == 0), "fixed_ref", 
                         if_else((`21_6-F` == 2 | `2165-S1` == 2 | `2165-4-S2` == 2 | `2165-44-S3` == 2 | 
                                    `2165-444-S4` == 2 | `2165-444-S5` == 2), "fixed_alt", "not_fixed"))) %>% 
  filter(fixed == "not_fixed")

# 23_2
gt_23_2_fixed_ref <- gt_23_2 %>% 
  mutate(fixed = if_else((`23_2-F` == 0 | `2323-S1` == 0 | `2323-2-S2` == 0 | `2323-21-S3` == 0 | 
                            `2323-211-S4` == 0 | `2323-211-S5` == 0), "fixed_ref", 
                         if_else((`23_2-F` == 2 | `2323-S1` == 2 | `2323-2-S2` == 2 | `2323-21-S3` == 2 | 
                                    `2323-211-S4` == 2 | `2323-211-S5` == 2), "fixed_alt", "not_fixed"))) %>% 
  filter(fixed == "fixed_ref")

gt_23_2_fixed_alt <- gt_23_2 %>% 
  mutate(fixed = if_else((`23_2-F` == 0 | `2323-S1` == 0 | `2323-2-S2` == 0 | `2323-21-S3` == 0 | 
                            `2323-211-S4` == 0 | `2323-211-S5` == 0), "fixed_ref", 
                         if_else((`23_2-F` == 2 | `2323-S1` == 2 | `2323-2-S2` == 2 | `2323-21-S3` == 2 | 
                                    `2323-211-S4` == 2 | `2323-211-S5` == 2), "fixed_alt", "not_fixed"))) %>% 
  filter(fixed == "fixed_alt")

gt_23_2_not_fixed <- gt_23_2 %>% 
  mutate(fixed = if_else((`23_2-F` == 0 | `2323-S1` == 0 | `2323-2-S2` == 0 | `2323-21-S3` == 0 | 
                            `2323-211-S4` == 0 | `2323-211-S5` == 0), "fixed_ref", 
                         if_else((`23_2-F` == 2 | `2323-S1` == 2 | `2323-2-S2` == 2 | `2323-21-S3` == 2 | 
                                    `2323-211-S4` == 2 | `2323-211-S5` == 2), "fixed_alt", "not_fixed"))) %>% 
  filter(fixed == "not_fixed")

# 23_4
gt_23_4_fixed_ref <- gt_23_4 %>% 
  mutate(fixed = if_else((`23_4-F` == 0 | `2344-S1` == 0 | `2344-4-S2` == 0 | `2344-46-S3` == 0 | 
                            `2344-464-S4` == 0), "fixed_ref", 
                         if_else((`23_4-F` == 2 | `2344-S1` == 2 | `2344-4-S2` == 2 | `2344-46-S3` == 2 | 
                                    `2344-464-S4` == 2), "fixed_alt", "not_fixed"))) %>%
  filter(fixed == "fixed_ref")

gt_23_4_fixed_alt <- gt_23_4 %>% 
  mutate(fixed = if_else((`23_4-F` == 0 | `2344-S1` == 0 | `2344-4-S2` == 0 | `2344-46-S3` == 0 | 
                            `2344-464-S4` == 0), "fixed_ref", 
                         if_else((`23_4-F` == 2 | `2344-S1` == 2 | `2344-4-S2` == 2 | `2344-46-S3` == 2 | 
                                    `2344-464-S4` == 2), "fixed_alt", "not_fixed"))) %>%  
  filter(fixed == "fixed_alt")

gt_23_4_not_fixed <- gt_23_4 %>% 
  mutate(fixed = if_else((`23_4-F` == 0 | `2344-S1` == 0 | `2344-4-S2` == 0 | `2344-46-S3` == 0 | 
                            `2344-464-S4` == 0), "fixed_ref", 
                         if_else((`23_4-F` == 2 | `2344-S1` == 2 | `2344-4-S2` == 2 | `2344-46-S3` == 2 | 
                                    `2344-464-S4` == 2), "fixed_alt", "not_fixed"))) %>%
  filter(fixed == "not_fixed")

# 26_1
gt_26_1_fixed_ref <- gt_26_1 %>% 
  mutate(fixed = if_else((`26_1-F` == 0 | `2612-S1` == 0 | `2612-1-S2` == 0 | `2612-12-S3` == 0 | 
                            `2612-121-S4` == 0), "fixed_ref", 
                         if_else((`26_1-F` == 2 | `2612-S1` == 2 | `2612-1-S2` == 2 | `2612-12-S3` == 2 | 
                                    `2612-121-S4` == 2), "fixed_alt", "not_fixed"))) %>% 
  filter(fixed == "fixed_ref")

gt_26_1_fixed_alt <- gt_26_1 %>% 
  mutate(fixed = if_else((`26_1-F` == 0 | `2612-S1` == 0 | `2612-1-S2` == 0 | `2612-12-S3` == 0 | 
                            `2612-121-S4` == 0), "fixed_ref", 
                         if_else((`26_1-F` == 2 | `2612-S1` == 2 | `2612-1-S2` == 2 | `2612-12-S3` == 2 | 
                                    `2612-121-S4` == 2), "fixed_alt", "not_fixed"))) %>% 
  filter(fixed == "fixed_alt")

gt_26_1_not_fixed <- gt_26_1 %>% 
  mutate(fixed = if_else((`26_1-F` == 0 | `2612-S1` == 0 | `2612-1-S2` == 0 | `2612-12-S3` == 0 | 
                            `2612-121-S4` == 0), "fixed_ref", 
                         if_else((`26_1-F` == 2 | `2612-S1` == 2 | `2612-1-S2` == 2 | `2612-12-S3` == 2 | 
                                    `2612-121-S4` == 2), "fixed_alt", "not_fixed"))) %>% 
  filter(fixed == "not_fixed")

# 26_4
gt_26_4_fixed_ref <- gt_26_4 %>% 
  mutate(fixed = if_else((`26_4-F` == 0 | `2645-S1` == 0 | `2645-4-S2` == 0 | `2645-46-S3` == 0 | 
                            `2645-464-S4` == 0), "fixed_ref", 
                         if_else((`26_4-F` == 2 | `2645-S1` == 2 | `2645-4-S2` == 2 | `2645-46-S3` == 2 | 
                                    `2645-464-S4` == 2), "fixed_alt", "not_fixed"))) %>%
  filter(fixed == "fixed_ref")

gt_26_4_fixed_alt <- gt_26_4 %>% 
  mutate(fixed = if_else((`26_4-F` == 0 | `2645-S1` == 0 | `2645-4-S2` == 0 | `2645-46-S3` == 0 | 
                            `2645-464-S4` == 0), "fixed_ref", 
                         if_else((`26_4-F` == 2 | `2645-S1` == 2 | `2645-4-S2` == 2 | `2645-46-S3` == 2 | 
                                    `2645-464-S4` == 2), "fixed_alt", "not_fixed"))) %>%
  filter(fixed == "fixed_alt")

gt_26_4_not_fixed <- gt_26_4 %>% 
  mutate(fixed = if_else((`26_4-F` == 0 | `2645-S1` == 0 | `2645-4-S2` == 0 | `2645-46-S3` == 0 | 
                            `2645-464-S4` == 0), "fixed_ref", 
                         if_else((`26_4-F` == 2 | `2645-S1` == 2 | `2645-4-S2` == 2 | `2645-46-S3` == 2 | 
                                    `2645-464-S4` == 2), "fixed_alt", "not_fixed"))) %>% 
  filter(fixed == "not_fixed")

# 27_2
gt_27_2_fixed_ref <- gt_27_2 %>% 
  mutate(fixed = if_else((`27_2-F` == 0 | `2721-S1` == 0 | `2721-1-S2` == 0 | `2721-12-S3` == 0 | 
                            `2721-121-S4` == 0), "fixed_ref", 
                         if_else((`27_2-F` == 2 | `2721-S1` == 2 | `2721-1-S2` == 2 | `2721-12-S3` == 2 | 
                                    `2721-121-S4` == 2), "fixed_alt", "not_fixed"))) %>% 
  filter(fixed == "fixed_ref")

gt_27_2_fixed_alt <- gt_27_2 %>% 
  mutate(fixed = if_else((`27_2-F` == 0 | `2721-S1` == 0 | `2721-1-S2` == 0 | `2721-12-S3` == 0 | 
                            `2721-121-S4` == 0), "fixed_ref", 
                         if_else((`27_2-F` == 2 | `2721-S1` == 2 | `2721-1-S2` == 2 | `2721-12-S3` == 2 | 
                                    `2721-121-S4` == 2), "fixed_alt", "not_fixed"))) %>% 
  filter(fixed == "fixed_alt")

gt_27_2_not_fixed <- gt_27_2 %>% 
  mutate(fixed = if_else((`27_2-F` == 0 | `2721-S1` == 0 | `2721-1-S2` == 0 | `2721-12-S3` == 0 | 
                            `2721-121-S4` == 0), "fixed_ref", 
                         if_else((`27_2-F` == 2 | `2721-S1` == 2 | `2721-1-S2` == 2 | `2721-12-S3` == 2 | 
                                    `2721-121-S4` == 2), "fixed_alt", "not_fixed"))) %>% 
  filter(fixed == "not_fixed")

# 29_2
gt_29_2_fixed_ref <- gt_29_2 %>% 
  mutate(fixed = if_else((`29_2-F` == 0 | `2923-S1` == 0 | `2923-1-S2` == 0 | `2923-12-S3` == 0 | 
                            `2923-121-S4` == 0 | `2923-121-S5` == 0), "fixed_ref", 
                         if_else((`29_2-F` == 2 | `2923-S1` == 2 | `2923-1-S2` == 2 | `2923-12-S3` == 2 | 
                                    `2923-121-S4` == 2 | `2923-121-S5` == 2), "fixed_alt", "not_fixed"))) %>% 
  filter(fixed == "fixed_ref")

gt_29_2_fixed_alt <- gt_29_2 %>% 
  mutate(fixed = if_else((`29_2-F` == 0 | `2923-S1` == 0 | `2923-1-S2` == 0 | `2923-12-S3` == 0 | 
                            `2923-121-S4` == 0 | `2923-121-S5` == 0), "fixed_ref", 
                         if_else((`29_2-F` == 2 | `2923-S1` == 2 | `2923-1-S2` == 2 | `2923-12-S3` == 2 | 
                                    `2923-121-S4` == 2 | `2923-121-S5` == 2), "fixed_alt", "not_fixed"))) %>% 
  filter(fixed == "fixed_alt")

gt_29_2_not_fixed <- gt_29_2 %>% 
  mutate(fixed = if_else((`29_2-F` == 0 | `2923-S1` == 0 | `2923-1-S2` == 0 | `2923-12-S3` == 0 | 
                            `2923-121-S4` == 0 | `2923-121-S5` == 0), "fixed_ref", 
                         if_else((`29_2-F` == 2 | `2923-S1` == 2 | `2923-1-S2` == 2 | `2923-12-S3` == 2 | 
                                    `2923-121-S4` == 2 | `2923-121-S5` == 2), "fixed_alt", "not_fixed"))) %>% 
  filter(fixed == "not_fixed")

# 29_4
gt_29_4_fixed_ref <- gt_29_4 %>% 
  mutate(fixed = if_else((`29_4-F` == 0 | `2944-S1` == 0 | `2944-5-S2` == 0 | `2944-56-S3` == 0 | 
                            `2944-565-S4` == 0 | `2944-565-S5` == 0), "fixed_ref", 
                         if_else((`29_4-F` == 2 | `2944-S1` == 2 | `2944-5-S2` == 2 | `2944-56-S3` == 2 | 
                                    `2944-565-S4` == 2 | `2944-565-S5` == 2), "fixed_alt", "not_fixed"))) %>% 
  filter(fixed == "fixed_ref")

gt_29_4_fixed_alt <- gt_29_4 %>% 
  mutate(fixed = if_else((`29_4-F` == 0 | `2944-S1` == 0 | `2944-5-S2` == 0 | `2944-56-S3` == 0 | 
                            `2944-565-S4` == 0 | `2944-565-S5` == 0), "fixed_ref", 
                         if_else((`29_4-F` == 2 | `2944-S1` == 2 | `2944-5-S2` == 2 | `2944-56-S3` == 2 | 
                                    `2944-565-S4` == 2 | `2944-565-S5` == 2), "fixed_alt", "not_fixed"))) %>%
  filter(fixed == "fixed_alt")

gt_29_4_not_fixed <- gt_29_4 %>% 
  mutate(fixed = if_else((`29_4-F` == 0 | `2944-S1` == 0 | `2944-5-S2` == 0 | `2944-56-S3` == 0 | 
                            `2944-565-S4` == 0 | `2944-565-S5` == 0), "fixed_ref", 
                         if_else((`29_4-F` == 2 | `2944-S1` == 2 | `2944-5-S2` == 2 | `2944-56-S3` == 2 | 
                                    `2944-565-S4` == 2 | `2944-565-S5` == 2), "fixed_alt", "not_fixed"))) %>%
  filter(fixed == "not_fixed")

# ### All fixed at gen S1 (unused) #####
# gt_1_1_fixed_S1_ref <- gt_1_1 %>% 
#   filter(`111-S1` == 0)
# 
# gt_6_1_fixed_S1_ref <- gt_6_1 %>% 
#   filter(`611-S1` == 0)
# 
# gt_6_4_fixed_S1_ref <- gt_6_4 %>% 
#   filter(`646-S1` == 0)
# 
# gt_7_2_fixed_S1_ref <- gt_7_2 %>% 
#   filter(`721-S1` == 0)
# 
# gt_7_4_fixed_S1_ref <- gt_7_4 %>% 
#   filter(`745-S1` == 0)
# 
# gt_8_2_fixed_S1_ref <- gt_8_2 %>% 
#   filter(`821-S1` == 0)
# 
# gt_8_4_fixed_S1_ref <- gt_8_4 %>% 
#   filter(`845-S1` == 0)
# 
# gt_10_2_fixed_S1_ref <- gt_10_2 %>% 
#   filter(`1022-S1` == 0)
# 
# gt_10_5_fixed_S1_ref <- gt_10_5 %>% 
#   filter(`1056-S1` == 0)
# 
# gt_13_3_fixed_S1_ref <- gt_13_3 %>% 
#   filter(`1332-S1` == 0)
# 
# gt_13_4_fixed_S1_ref <- gt_13_4 %>% 
#   filter(`1345-S1` == 0)
# 
# gt_16_1_fixed_S1_ref <- gt_16_1 %>% 
#   filter(`1611-S1` == 0)
# 
# gt_16_5_fixed_S1_ref <- gt_16_5 %>% 
#   filter(`1654-S1` == 0)
# 
# gt_17_2_fixed_S1_ref <- gt_17_2 %>% 
#   filter(`1721-S1` == 0)
# 
# gt_17_5_fixed_S1_ref <- gt_17_5 %>% 
#   filter(`1755-S1` == 0)
# 
# gt_19_2_fixed_S1_ref <- gt_19_2 %>% 
#   filter(`1922-S1` == 0)
# 
# gt_19_5_fixed_S1_ref <- gt_19_5 %>% 
#   filter(`1955-S1` == 0)
# 
# gt_20_1_fixed_S1_ref <- gt_20_1 %>% 
#   filter(`2013-S1` == 0)
# 
# gt_20_4_fixed_S1_ref <- gt_20_4 %>% 
#   filter(`2045-S1` == 0)
# 
# gt_21_2_fixed_S1_ref <- gt_21_2 %>% 
#   filter(`2121-S1` == 0)
# 
# gt_21_6_fixed_S1_ref <- gt_21_6 %>% 
#   filter(`2165-S1` == 0)
# 
# gt_26_1_fixed_S1_ref <- gt_26_1 %>% 
#   filter(`2612-S1` == 0)
# 
# gt_26_4_fixed_S1_ref <- gt_26_4 %>% 
#   filter(`2645-S1` == 0)
# 
# gt_27_2_fixed_S1_ref <- gt_27_2 %>% 
#   filter(`2721-S1` == 0)
# 
# gt_29_2_fixed_S1_ref <- gt_29_2 %>% 
#   filter(`2923-S1` == 0)
# 
# gt_29_4_fixed_S1_ref <- gt_29_4 %>% 
#   filter(`2944-S1` == 0)
# 
# ### All fixed at gen S1 alt#####
# gt_1_1_fixed_S1_alt <- gt_1_1 %>% 
#   filter(`111-S1` == 2)
# 
# gt_6_1_fixed_S1_alt <- gt_6_1 %>% 
#   filter(`611-S1` == 2)
# 
# gt_6_4_fixed_S1_alt <- gt_6_4 %>% 
#   filter(`646-S1` == 2)
# 
# gt_7_2_fixed_S1_alt <- gt_7_2 %>% 
#   filter(`721-S1` == 2)
# 
# gt_7_4_fixed_S1_alt <- gt_7_4 %>% 
#   filter(`745-S1` == 2)
# 
# gt_8_2_fixed_S1_alt <- gt_8_2 %>% 
#   filter(`821-S1` == 2)
# 
# gt_8_4_fixed_S1_alt <- gt_8_4 %>% 
#   filter(`845-S1` == 2)
# 
# gt_10_2_fixed_S1_alt <- gt_10_2 %>% 
#   filter(`1022-S1` == 2)
# 
# gt_10_5_fixed_S1_alt <- gt_10_5 %>% 
#   filter(`1056-S1` == 2)
# 
# gt_13_3_fixed_S1_alt <- gt_13_3 %>% 
#   filter(`1332-S1` == 2)
# 
# gt_13_4_fixed_S1_alt <- gt_13_4 %>% 
#   filter(`1345-S1` == 2)
# 
# gt_16_1_fixed_S1_alt <- gt_16_1 %>% 
#   filter(`1611-S1` == 2)
# 
# gt_16_5_fixed_S1_alt <- gt_16_5 %>% 
#   filter(`1654-S1` == 2)
# 
# gt_17_2_fixed_S1_alt <- gt_17_2 %>% 
#   filter(`1721-S1` == 2)
# 
# gt_17_5_fixed_S1_alt <- gt_17_5 %>% 
#   filter(`1755-S1` == 2)
# 
# gt_19_2_fixed_S1_alt <- gt_19_2 %>% 
#   filter(`1922-S1` == 2)
# 
# gt_19_5_fixed_S1_alt <- gt_19_5 %>% 
#   filter(`1955-S1` == 2)
# 
# gt_20_1_fixed_S1_alt <- gt_20_1 %>% 
#   filter(`2013-S1` == 2)
# 
# gt_20_4_fixed_S1_alt <- gt_20_4 %>% 
#   filter(`2045-S1` == 2)
# 
# gt_21_2_fixed_S1_alt <- gt_21_2 %>% 
#   filter(`2121-S1` == 2)
# 
# gt_21_6_fixed_S1_alt <- gt_21_6 %>% 
#   filter(`2165-S1` == 2)
# 
# gt_26_1_fixed_S1_alt <- gt_26_1 %>% 
#   filter(`2612-S1` == 2)
# 
# gt_26_4_fixed_S1_alt <- gt_26_4 %>% 
#   filter(`2645-S1` == 2)
# 
# gt_27_2_fixed_S1_alt <- gt_27_2 %>% 
#   filter(`2721-S1` == 2)
# 
# gt_29_2_fixed_S1_alt <- gt_29_2 %>% 
#   filter(`2923-S1` == 2)
# 
# gt_29_4_fixed_S1_alt <- gt_29_4 %>% 
#   filter(`2944-S1` == 2)
# 
# ### All not fixed at gen S1 alt#####
# gt_1_1_not_fixed_S1 <- gt_1_1 %>% 
#   filter(`111-S1` == 1)
# 
# gt_6_1_not_fixed_S1 <- gt_6_1 %>% 
#   filter(`611-S1` == 1)
# 
# gt_6_4_not_fixed_S1 <- gt_6_4 %>% 
#   filter(`646-S1` == 1)
# 
# gt_7_2_not_fixed_S1 <- gt_7_2 %>% 
#   filter(`721-S1` == 1)
# 
# gt_7_4_not_fixed_S1 <- gt_7_4 %>% 
#   filter(`745-S1` == 1)
# 
# gt_8_2_not_fixed_S1 <- gt_8_2 %>% 
#   filter(`821-S1` == 1)
# 
# gt_8_4_not_fixed_S1 <- gt_8_4 %>% 
#   filter(`845-S1` == 1)
# 
# gt_10_2_not_fixed_S1 <- gt_10_2 %>% 
#   filter(`1022-S1` == 1)
# 
# gt_10_5_not_fixed_S1 <- gt_10_5 %>% 
#   filter(`1056-S1` == 1)
# 
# gt_13_3_not_fixed_S1 <- gt_13_3 %>% 
#   filter(`1332-S1` == 1)
# 
# gt_13_4_not_fixed_S1 <- gt_13_4 %>% 
#   filter(`1345-S1` == 1)
# 
# gt_16_1_not_fixed_S1 <- gt_16_1 %>% 
#   filter(`1611-S1` == 1)
# 
# gt_16_5_not_fixed_S1 <- gt_16_5 %>% 
#   filter(`1654-S1` == 1)
# 
# gt_17_2_not_fixed_S1 <- gt_17_2 %>% 
#   filter(`1721-S1` == 1)
# 
# gt_17_5_not_fixed_S1 <- gt_17_5 %>% 
#   filter(`1755-S1` == 1)
# 
# gt_19_2_not_fixed_S1 <- gt_19_2 %>% 
#   filter(`1922-S1` == 1)
# 
# gt_19_5_not_fixed_S1 <- gt_19_5 %>% 
#   filter(`1955-S1` == 1)
# 
# gt_20_1_not_fixed_S1 <- gt_20_1 %>% 
#   filter(`2013-S1` == 1)
# 
# gt_20_4_not_fixed_S1 <- gt_20_4 %>% 
#   filter(`2045-S1` == 1)
# 
# gt_21_2_not_fixed_S1 <- gt_21_2 %>% 
#   filter(`2121-S1` == 1)
# 
# gt_21_6_not_fixed_S1 <- gt_21_6 %>% 
#   filter(`2165-S1` == 1)
# 
# gt_26_1_not_fixed_S1 <- gt_26_1 %>% 
#   filter(`2612-S1` == 1)
# 
# gt_26_4_not_fixed_S1 <- gt_26_4 %>% 
#   filter(`2645-S1` == 1)
# 
# gt_27_2_not_fixed_S1 <- gt_27_2 %>% 
#   filter(`2721-S1` == 1)
# 
# gt_29_2_not_fixed_S1 <- gt_29_2 %>% 
#   filter(`2923-S1` == 1)
# 
# gt_29_4_not_fixed_S1 <- gt_29_4 %>% 
#   filter(`2944-S1` == 1)


gt_all <- merge(gt_1_1, gt_6_1, by = "SNP", all = T)
gt_all <- merge(gt_all, gt_6_4, by = "SNP", all = T)
gt_all <- merge(gt_all, gt_7_2, by = "SNP", all = T)
gt_all <- merge(gt_all, gt_7_4, by = "SNP", all = T)
gt_all <- merge(gt_all, gt_8_2, by = "SNP", all = T)
gt_all <- merge(gt_all, gt_8_4, by = "SNP", all = T)
gt_all <- merge(gt_all, gt_10_2, by = "SNP", all = T)
gt_all <- merge(gt_all, gt_10_5, by = "SNP", all = T)
gt_all <- merge(gt_all, gt_13_3, by = "SNP", all = T)
gt_all <- merge(gt_all, gt_13_4, by = "SNP", all = T)
gt_all <- merge(gt_all, gt_16_1, by = "SNP", all = T)
gt_all <- merge(gt_all, gt_16_5, by = "SNP", all = T)
gt_all <- merge(gt_all, gt_17_2, by = "SNP", all = T)
gt_all <- merge(gt_all, gt_17_5, by = "SNP", all = T)
gt_all <- merge(gt_all, gt_19_2, by = "SNP", all = T)
gt_all <- merge(gt_all, gt_19_5, by = "SNP", all = T)
gt_all <- merge(gt_all, gt_20_1, by = "SNP", all = T)
gt_all <- merge(gt_all, gt_20_4, by = "SNP", all = T)
gt_all <- merge(gt_all, gt_21_2, by = "SNP", all = T)
gt_all <- merge(gt_all, gt_21_6, by = "SNP", all = T)
gt_all <- merge(gt_all, gt_23_2, by = "SNP", all = T)
gt_all <- merge(gt_all, gt_23_4, by = "SNP", all = T)
gt_all <- merge(gt_all, gt_26_1, by = "SNP", all = T)
gt_all <- merge(gt_all, gt_26_4, by = "SNP", all = T)
gt_all <- merge(gt_all, gt_27_2, by = "SNP", all = T)
gt_all <- merge(gt_all, gt_29_2, by = "SNP", all = T)
gt_all <- merge(gt_all, gt_29_4, by = "SNP", all = T)

gt_all_F <- data.frame(SNP = gt_all$SNP, select_at(gt_all, vars(ends_with("F"))), check.names = F)
gt_all_S1 <- data.frame(SNP = gt_all$SNP, select_at(gt_all, vars(ends_with("S1"))), check.names = F)
gt_all_S2 <- data.frame(SNP = gt_all$SNP, select_at(gt_all, vars(ends_with("S2"))), check.names = F)
gt_all_S3 <- data.frame(SNP = gt_all$SNP, select_at(gt_all, vars(ends_with("S3"))), check.names = F)
gt_all_S4 <- data.frame(SNP = gt_all$SNP, select_at(gt_all, vars(ends_with("S4"))), check.names = F)
gt_all_S5 <- data.frame(SNP = gt_all$SNP, select_at(gt_all, vars(ends_with("S5"))), check.names = F)

# write.table(gt_all_F, "F_lines.txt", row.names = F, quote = F)
# write.table(gt_all_S1, "S1_lines.txt", row.names = F, quote = F)
# write.table(gt_all_S2, "S2_lines.txt", row.names = F, quote = F)
# write.table(gt_all_S3, "S3_lines.txt", row.names = F, quote = F)
# write.table(gt_all_S4, "S4_lines.txt", row.names = F, quote = F)
# write.table(gt_all_S5, "S5_lines.txt", row.names = F, quote = F)

# F_with_obs_het <- gt_all_F %>% 
#   mutate(het = sum)
# 
# fixed_ref_all <- merge(gt_1_1_fixed_ref, gt_6_1_fixed_ref, by = "SNP", all = F)
# fixed_ref_all <- merge(fixed_ref_all, gt_6_4_fixed_ref, by = "SNP", all = F)
# fixed_ref_all <- merge(fixed_ref_all, gt_7_4_fixed_ref, by = "SNP", all = F)
# fixed_ref_all <- merge(fixed_ref_all, gt_8_2_fixed_ref, by = "SNP", all = F)
# fixed_ref_all <- merge(fixed_ref_all, gt_8_4_fixed_ref, by = "SNP", all = F)
# fixed_ref_all <- merge(fixed_ref_all, gt_13_4_fixed_ref, by = "SNP", all = F)
# fixed_ref_all <- merge(fixed_ref_all, gt_16_1_fixed_ref, by = "SNP", all = F)
# fixed_ref_all <- merge(fixed_ref_all, gt_17_2_fixed_ref, by = "SNP", all = F)
# fixed_ref_all <- merge(fixed_ref_all, gt_17_5_fixed_ref, by = "SNP", all = F)
# fixed_ref_all <- merge(fixed_ref_all, gt_19_5_fixed_ref, by = "SNP", all = F)
# fixed_ref_all <- merge(fixed_ref_all, gt_20_1_fixed_ref, by = "SNP", all = F)
# fixed_ref_all <- merge(fixed_ref_all, gt_20_4_fixed_ref, by = "SNP", all = F)
# fixed_ref_all <- merge(fixed_ref_all, gt_21_2_fixed_ref, by = "SNP", all = F)
# fixed_ref_all <- merge(fixed_ref_all, gt_21_6_fixed_ref, by = "SNP", all = F)
# fixed_ref_all <- merge(fixed_ref_all, gt_23_2_fixed_ref, by = "SNP", all = F)
# fixed_ref_all <- merge(fixed_ref_all, gt_23_4_fixed_ref, by = "SNP", all = F)
# fixed_ref_all <- merge(fixed_ref_all, gt_26_4_fixed_ref, by = "SNP", all = F)
# fixed_ref_all <- merge(fixed_ref_all, gt_29_2_fixed_ref, by = "SNP", all = F)
# fixed_ref_all <- merge(fixed_ref_all, gt_29_4_fixed_ref, by = "SNP", all = F)
# 
# 
# fixed_alt_all <- merge(gt_1_1_fixed_alt, gt_6_1_fixed_alt, by = "SNP", all = F)
# fixed_alt_all <- merge(fixed_alt_all, gt_6_4_fixed_alt, by = "SNP", all = F)
# fixed_alt_all <- merge(fixed_alt_all, gt_7_4_fixed_alt, by = "SNP", all = F)
# 
# not_fixed_all <- merge(gt_1_1_not_fixed, gt_6_1_not_fixed, by = "SNP", all = F)
# not_fixed_all <- merge(not_fixed_all, gt_6_4_not_fixed, by = "SNP", all = F)
# not_fixed_all <- merge(not_fixed_all, gt_7_4_not_fixed, by = "SNP", all = F)
# not_fixed_all <- merge(not_fixed_all, gt_8_2_not_fixed, by = "SNP", all = F)
# not_fixed_all <- merge(not_fixed_all, gt_8_4_not_fixed, by = "SNP", all = F)
# not_fixed_all <- merge(not_fixed_all, gt_13_4_not_fixed, by = "SNP", all = F)
# not_fixed_all <- merge(not_fixed_all, gt_16_1_not_fixed, by = "SNP", all = F)
# not_fixed_all <- merge(not_fixed_all, gt_17_2_not_fixed, by = "SNP", all = F)
# not_fixed_all <- merge(not_fixed_all, gt_17_5_not_fixed, by = "SNP", all = F)
# not_fixed_all <- merge(not_fixed_all, gt_19_5_not_fixed, by = "SNP", all = F)
# not_fixed_all <- merge(not_fixed_all, gt_20_1_not_fixed, by = "SNP", all = F)
# not_fixed_all <- merge(not_fixed_all, gt_20_4_not_fixed, by = "SNP", all = F)
# not_fixed_all <- merge(not_fixed_all, gt_21_2_not_fixed, by = "SNP", all = F)
# not_fixed_all <- merge(not_fixed_all, gt_21_6_not_fixed, by = "SNP", all = F)
# not_fixed_all <- merge(not_fixed_all, gt_23_2_not_fixed, by = "SNP", all = F)
# not_fixed_all <- merge(not_fixed_all, gt_23_4_not_fixed, by = "SNP", all = F)
# not_fixed_all <- merge(not_fixed_all, gt_26_4_not_fixed, by = "SNP", all = F)
# not_fixed_all <- merge(not_fixed_all, gt_29_2_not_fixed, by = "SNP", all = F)
# not_fixed_all <- merge(not_fixed_all, gt_29_4_not_fixed, by = "SNP", all = F)

# write.table(not_fixed_all$SNP, "heterozygous_snps_all_lines.txt", quote = F, row.names = F, col.names = F)

### Saving overlap of fixation tables ########################
ref_fixed_overlap_table <- data.frame(table(sort(c(gt_1_1_fixed_ref$SNP, gt_6_1_fixed_ref$SNP, gt_6_4_fixed_ref$SNP, 
                                                   gt_7_2_fixed_ref$SNP, gt_7_4_fixed_ref$SNP, gt_8_2_fixed_ref$SNP,
                                                   gt_8_4_fixed_ref$SNP, gt_10_2_fixed_ref$SNP, gt_10_5_fixed_ref$SNP,
                                                   gt_13_3_fixed_ref$SNP, gt_13_4_fixed_ref$SNP, gt_16_1_fixed_ref$SNP, 
                                                   gt_17_2_fixed_ref$SNP, gt_17_5_fixed_ref$SNP, gt_19_2_fixed_ref$SNP, 
                                                   gt_19_5_fixed_ref$SNP, gt_20_1_fixed_ref$SNP, gt_20_4_fixed_ref$SNP,
                                                   gt_21_2_fixed_ref$SNP, gt_21_6_fixed_ref$SNP,
                                                   gt_23_2_fixed_ref$SNP, gt_23_4_fixed_ref$SNP,
                                                   gt_26_1_fixed_ref$SNP, gt_26_4_fixed_ref$SNP, gt_27_2_fixed_ref$SNP,
                                                   gt_29_2_fixed_ref$SNP, gt_29_4_fixed_ref$SNP))))

# write.table(ref_fixed_overlap_table, "ref_fixed_overlap_table.txt", quote = F)

# ref_fixed_S1_overlap_table <- data.frame(table(sort(c(gt_1_1_fixed_S1_ref$SNP, gt_6_1_fixed_S1_ref$SNP, gt_6_4_fixed_S1_ref$SNP, 
#                                                       gt_7_2_fixed_S1_ref$SNP, gt_7_4_fixed_S1_ref$SNP, gt_8_2_fixed_S1_ref$SNP,
#                                                       gt_8_4_fixed_S1_ref$SNP, gt_10_2_fixed_S1_ref$SNP, gt_10_5_fixed_S1_ref$SNP,
#                                                       gt_13_3_fixed_S1_ref$SNP, gt_13_4_fixed_S1_ref$SNP, gt_16_1_fixed_S1_ref$SNP, 
#                                                       gt_17_2_fixed_S1_ref$SNP, gt_17_5_fixed_S1_ref$SNP, gt_19_2_fixed_S1_ref$SNP, 
#                                                       gt_19_5_fixed_S1_ref$SNP, gt_20_1_fixed_S1_ref$SNP, gt_20_4_fixed_S1_ref$SNP,
#                                                       gt_21_2_fixed_S1_ref$SNP, gt_21_6_fixed_S1_ref$SNP,
#                                                       gt_23_2_fixed_S1_ref$SNP, gt_23_4_fixed_S1_ref$SNP,
#                                                       gt_26_1_fixed_S1_ref$SNP, gt_26_4_fixed_S1_ref$SNP, gt_27_2_fixed_S1_ref$SNP,
#                                                       gt_29_2_fixed_S1_ref$SNP, gt_29_4_fixed_S1_ref$SNP))))


alt_fixed_overlap_table <- data.frame(table(sort(c(gt_1_1_fixed_alt$SNP, gt_6_1_fixed_alt$SNP, gt_6_4_fixed_alt$SNP, 
                                                   gt_7_2_fixed_alt$SNP, gt_7_4_fixed_alt$SNP, gt_8_2_fixed_alt$SNP,
                                                   gt_8_4_fixed_alt$SNP, gt_10_2_fixed_alt$SNP, gt_10_5_fixed_alt$SNP,
                                                   gt_13_3_fixed_alt$SNP, gt_13_4_fixed_alt$SNP, gt_16_1_fixed_alt$SNP, 
                                                   gt_17_2_fixed_alt$SNP, gt_17_5_fixed_alt$SNP, gt_19_2_fixed_alt$SNP, 
                                                   gt_19_5_fixed_alt$SNP, gt_20_1_fixed_alt$SNP, gt_20_4_fixed_alt$SNP,
                                                   gt_21_2_fixed_alt$SNP, gt_21_6_fixed_alt$SNP,
                                                   gt_23_2_fixed_alt$SNP, gt_23_4_fixed_alt$SNP,
                                                   gt_26_1_fixed_alt$SNP, gt_26_4_fixed_alt$SNP, gt_27_2_fixed_alt$SNP,
                                                   gt_29_2_fixed_alt$SNP, gt_29_4_fixed_alt$SNP))))

# write.table(alt_fixed_overlap_table, "alt_fixed_overlap_table.txt", quote = F)

# alt_fixed_S1_overlap_table <- data.frame(table(sort(c(gt_1_1_fixed_S1_alt$SNP, gt_6_1_fixed_S1_alt$SNP, gt_6_4_fixed_S1_alt$SNP, 
#                                                       gt_7_2_fixed_S1_alt$SNP, gt_7_4_fixed_S1_alt$SNP, gt_8_2_fixed_S1_alt$SNP,
#                                                       gt_8_4_fixed_S1_alt$SNP, gt_10_2_fixed_S1_alt$SNP, gt_10_5_fixed_S1_alt$SNP,
#                                                       gt_13_3_fixed_S1_alt$SNP, gt_13_4_fixed_S1_alt$SNP, gt_16_1_fixed_S1_alt$SNP, 
#                                                       gt_17_2_fixed_S1_alt$SNP, gt_17_5_fixed_S1_alt$SNP, gt_19_2_fixed_S1_alt$SNP, 
#                                                       gt_19_5_fixed_S1_alt$SNP, gt_20_1_fixed_S1_alt$SNP, gt_20_4_fixed_S1_alt$SNP,
#                                                       gt_21_2_fixed_S1_alt$SNP, gt_21_6_fixed_S1_alt$SNP,
#                                                       gt_23_2_fixed_S1_alt$SNP, gt_23_4_fixed_S1_alt$SNP,
#                                                       gt_26_1_fixed_S1_alt$SNP, gt_26_4_fixed_S1_alt$SNP, gt_27_2_fixed_S1_alt$SNP,
                                                      # gt_29_2_fixed_S1_alt$SNP, gt_29_4_fixed_S1_alt$SNP))))


not_fixed_overlap_table <- data.frame(table(sort(c(gt_1_1_not_fixed$SNP, gt_6_1_not_fixed$SNP, gt_6_4_not_fixed$SNP, 
                                                   gt_7_2_not_fixed$SNP, gt_7_4_not_fixed$SNP, gt_8_2_not_fixed$SNP,
                                                   gt_8_4_not_fixed$SNP, gt_10_2_not_fixed$SNP, gt_10_5_not_fixed$SNP,
                                                   gt_13_3_not_fixed$SNP, gt_13_4_not_fixed$SNP, gt_16_1_not_fixed$SNP, 
                                                   gt_17_2_not_fixed$SNP, gt_17_5_not_fixed$SNP, gt_19_2_not_fixed$SNP, 
                                                   gt_19_5_not_fixed$SNP, gt_20_1_not_fixed$SNP, gt_20_4_not_fixed$SNP,
                                                   gt_21_2_not_fixed$SNP, gt_21_6_not_fixed$SNP,
                                                   gt_23_2_not_fixed$SNP, gt_23_4_not_fixed$SNP,
                                                   gt_26_1_not_fixed$SNP, gt_26_4_not_fixed$SNP, gt_27_2_not_fixed$SNP,
                                                   gt_29_2_not_fixed$SNP, gt_29_4_not_fixed$SNP))))

# write.table(not_fixed_overlap_table, "not_fixed_overlap_table.txt", quote = F)

# not_fixed_S1_overlap_table <- data.frame(table(sort(c(gt_1_1_not_fixed_S1$SNP, gt_6_1_not_fixed_S1$SNP, gt_6_4_not_fixed_S1$SNP, 
#                                                       gt_7_2_not_fixed_S1$SNP, gt_7_4_not_fixed_S1$SNP, gt_8_2_not_fixed_S1$SNP,
#                                                       gt_8_4_not_fixed_S1$SNP, gt_10_2_not_fixed_S1$SNP, gt_10_5_not_fixed_S1$SNP,
#                                                       gt_13_3_not_fixed_S1$SNP, gt_13_4_not_fixed_S1$SNP, gt_16_1_not_fixed_S1$SNP, 
#                                                       gt_17_2_not_fixed_S1$SNP, gt_17_5_not_fixed_S1$SNP, gt_19_2_not_fixed_S1$SNP, 
#                                                       gt_19_5_not_fixed_S1$SNP, gt_20_1_not_fixed_S1$SNP, gt_20_4_not_fixed_S1$SNP,
#                                                       gt_21_2_not_fixed_S1$SNP, gt_21_6_not_fixed_S1$SNP,
#                                                       gt_23_2_not_fixed_S1$SNP, gt_23_4_not_fixed_S1$SNP,
#                                                       gt_26_1_not_fixed_S1$SNP, gt_26_4_not_fixed_S1$SNP, gt_27_2_not_fixed_S1$SNP,
#                                                       gt_29_2_not_fixed_S1$SNP, gt_29_4_not_fixed_S1$SNP))))


### Chi-squared analysis #######################
# Creating table of SNPs and fixed/notfixed lines
S4_fate_table <- merge(not_fixed_overlap_table, ref_fixed_overlap_table, by = "Var1", all = T)
S4_fate_table <- merge(S4_fate_table, alt_fixed_overlap_table, by = "Var1", all = T)

S4_fate_table <- S4_fate_table %>% 
  rename(SNP = Var1, not_fixed = Freq.x, ref_fixed = Freq.y, alt_fixed = Freq)

# missing SNPs
missing_snps <- allele_freqs_F %>% 
  filter(!(SNP %in% S4_fate_table$SNP)) %>%
  select(SNP) %>% 
  mutate(not_fixed = rep(NA, 1), ref_fixed = rep(NA, 1), alt_fixed = rep(NA, 1))

S4_fate_table <- rbind(S4_fate_table, missing_snps)

S4_fate_table[is.na(S4_fate_table)] <- 0

# Calculate proportions (not really needed)
S4_fate_table_props <- S4_fate_table %>% 
  mutate(total = (not_fixed + ref_fixed + alt_fixed)) %>% 
  mutate(not_fixed_prop = not_fixed/(not_fixed + ref_fixed + alt_fixed)) %>% 
  mutate(ref_fixed_prop = ref_fixed/(not_fixed + ref_fixed + alt_fixed)) %>%
  mutate(alt_fixed_prop = alt_fixed/(not_fixed + ref_fixed + alt_fixed))

# Do chi-squared test for all three 
S4_fate_table_props_for_chisq <- S4_fate_table_props %>% 
  # mutate(not_fixed_exp = rep(0.03125, 44099), ref_fixed_exp = rep(0.484375, 44099), alt_fixed_exp = rep(0.484375, 44099)) %>%
  mutate(not_fixed_exp = 0.0625 * total, ref_fixed_exp = 0.46875 * total, alt_fixed_exp = 0.46875 * total) %>%
  mutate(chi_sq = (((not_fixed - not_fixed_exp)^2/not_fixed_exp) + ((ref_fixed - ref_fixed_exp)^2/ref_fixed_exp) + ((alt_fixed - alt_fixed_exp)^2/alt_fixed_exp))) %>%
  mutate(p.value = pchisq(chi_sq, 2, lower.tail = F)) %>%
  mutate(p.adj.BH = p.adjust(p.value, "BH"), p.adj.bonf = p.adjust(p.value, "bonferroni"), p.adj.holm = p.adjust(p.value, "holm"))


# write.csv(S4_fate_table_props_for_chisq, "S4_fate_table_chisq_analysis_filtered_snps.csv", row.names = F)

# outlier results
S4_fate_table_outlier <- S4_fate_table_props_for_chisq %>% 
  filter(total >= 3) %>% 
  filter(p.adj.BH < 0.05)

# write.csv(S4_fate_table_outlier, "S4_fate_table_outlier.csv", row.names = F)

### Analyses for individual lines (unused) ####
# Line_1_1_genind <- FS_S5_genind[c(1, 21, 41, 61, 81, 101)]
# 
# Line_1_1_F_genind <- FS_S5_genind[1]
# Line_1_1_S1_genind <- FS_S5_genind[29]
# Line_1_1_S2_genind <- FS_S5_genind[57]
# Line_1_1_S3_genind <- FS_S5_genind[85]
# Line_1_1_S4_genind <- FS_S5_genind[113]
# Line_1_1_S5_genind <- FS_S5_genind[141]
# 
# Line_6_1_F_genind <- FS_S5_genind[2]
# Line_6_1_S1_genind <- FS_S5_genind[30]
# Line_6_1_S2_genind <- FS_S5_genind[58]
# Line_6_1_S3_genind <- FS_S5_genind[86]
# Line_6_1_S4_genind <- FS_S5_genind[114]
# 
# Line_6_4_F_genind <- FS_S5_genind[3]
# Line_6_4_S1_genind <- FS_S5_genind[31]
# Line_6_4_S2_genind <- FS_S5_genind[59]
# Line_6_4_S3_genind <- FS_S5_genind[87]
# Line_6_4_S4_genind <- FS_S5_genind[115]
# Line_6_4_S5_genind <- FS_S5_genind[142]
# 
# Line_7_2_F_genind <- FS_S5_genind[4]
# Line_7_2_S1_genind <- FS_S5_genind[32]
# Line_7_2_S2_genind <- FS_S5_genind[60]
# Line_7_2_S3_genind <- FS_S5_genind[88]
# Line_7_2_S4_genind <- FS_S5_genind[116]
# 
# Line_7_4_F_genind <- FS_S5_genind[5]
# Line_7_4_S1_genind <- FS_S5_genind[33]
# Line_7_4_S2_genind <- FS_S5_genind[61]
# Line_7_4_S3_genind <- FS_S5_genind[89]
# Line_7_4_S4_genind <- FS_S5_genind[117]
# 
# Line_8_2_F_genind <- FS_S5_genind[6]
# Line_8_2_S1_genind <- FS_S5_genind[34]
# Line_8_2_S2_genind <- FS_S5_genind[62]
# Line_8_2_S3_genind <- FS_S5_genind[90]
# Line_8_2_S4_genind <- FS_S5_genind[118]
# 
# Line_8_4_F_genind <- FS_S5_genind[7]
# Line_8_4_S1_genind <- FS_S5_genind[35]
# Line_8_4_S2_genind <- FS_S5_genind[63]
# Line_8_4_S3_genind <- FS_S5_genind[91]
# Line_8_4_S4_genind <- FS_S5_genind[119]
# 
# Line_10_2_F_genind <- FS_S5_genind[8]
# Line_10_2_S1_genind <- FS_S5_genind[36]
# Line_10_2_S2_genind <- FS_S5_genind[64]
# Line_10_2_S3_genind <- FS_S5_genind[92]
# Line_10_2_S4_genind <- FS_S5_genind[120]
# Line_10_2_S5_genind <- FS_S5_genind[143]
# 
# Line_10_5_F_genind <- FS_S5_genind[9]
# Line_10_5_S1_genind <- FS_S5_genind[37]
# Line_10_5_S2_genind <- FS_S5_genind[65]
# Line_10_5_S3_genind <- FS_S5_genind[93]
# Line_10_5_S4_genind <- FS_S5_genind[121]
# 
# Line_13_3_F_genind <- FS_S5_genind[10]
# Line_13_3_S1_genind <- FS_S5_genind[38]
# Line_13_3_S2_genind <- FS_S5_genind[66]
# Line_13_3_S3_genind <- FS_S5_genind[94]
# Line_13_3_S4_genind <- FS_S5_genind[122]
# 
# Line_13_4_F_genind <- FS_S5_genind[11]
# Line_13_4_S1_genind <- FS_S5_genind[39]
# Line_13_4_S2_genind <- FS_S5_genind[67]
# Line_13_4_S3_genind <- FS_S5_genind[95]
# Line_13_4_S4_genind <- FS_S5_genind[123]
# 
# Line_16_1_F_genind <- FS_S5_genind[12]
# Line_16_1_S1_genind <- FS_S5_genind[40]
# Line_16_1_S2_genind <- FS_S5_genind[68]
# Line_16_1_S3_genind <- FS_S5_genind[96]
# Line_16_1_S4_genind <- FS_S5_genind[124]
# 
# Line_16_5_F_genind <- FS_S5_genind[13]
# Line_16_5_S1_genind <- FS_S5_genind[41]
# Line_16_5_S2_genind <- FS_S5_genind[69]
# Line_16_5_S3_genind <- FS_S5_genind[97]
# Line_16_5_S4_genind <- FS_S5_genind[125]
# Line_16_5_S5_genind <- FS_S5_genind[144]
# 
# Line_17_2_F_genind <- FS_S5_genind[14]
# Line_17_2_S1_genind <- FS_S5_genind[42]
# Line_17_2_S2_genind <- FS_S5_genind[70]
# Line_17_2_S3_genind <- FS_S5_genind[98]
# Line_17_2_S4_genind <- FS_S5_genind[126]
# Line_17_2_S5_genind <- FS_S5_genind[145]
# 
# Line_17_5_F_genind <- FS_S5_genind[15]
# Line_17_5_S1_genind <- FS_S5_genind[43]
# Line_17_5_S2_genind <- FS_S5_genind[71]
# Line_17_5_S3_genind <- FS_S5_genind[99]
# Line_17_5_S4_genind <- FS_S5_genind[127]
# Line_17_5_S5_genind <- FS_S5_genind[146]
# 
# Line_19_2_F_genind <- FS_S5_genind[16]
# Line_19_2_S1_genind <- FS_S5_genind[44]
# Line_19_2_S2_genind <- FS_S5_genind[72]
# Line_19_2_S3_genind <- FS_S5_genind[100]
# Line_19_2_S4_genind <- FS_S5_genind[128]
# 
# Line_19_5_F_genind <- FS_S5_genind[17]
# Line_19_5_S1_genind <- FS_S5_genind[45]
# Line_19_5_S2_genind <- FS_S5_genind[73]
# Line_19_5_S3_genind <- FS_S5_genind[101]
# Line_19_5_S4_genind <- FS_S5_genind[129]
# 
# Line_20_1_F_genind <- FS_S5_genind[18]
# Line_20_1_S1_genind <- FS_S5_genind[46]
# Line_20_1_S2_genind <- FS_S5_genind[74]
# Line_20_1_S3_genind <- FS_S5_genind[102]
# Line_20_1_S4_genind <- FS_S5_genind[130]
# 
# Line_20_4_F_genind <- FS_S5_genind[19]
# Line_20_4_S1_genind <- FS_S5_genind[47]
# Line_20_4_S2_genind <- FS_S5_genind[75]
# Line_20_4_S3_genind <- FS_S5_genind[103]
# Line_20_4_S4_genind <- FS_S5_genind[131]
# 
# Line_21_2_F_genind <- FS_S5_genind[20]
# Line_21_2_S1_genind <- FS_S5_genind[48]
# Line_21_2_S2_genind <- FS_S5_genind[76]
# Line_21_2_S3_genind <- FS_S5_genind[104]
# Line_21_2_S4_genind <- FS_S5_genind[132]
# Line_21_2_S5_genind <- FS_S5_genind[147]
# 
# Line_21_6_F_genind <- FS_S5_genind[21]
# Line_21_6_S1_genind <- FS_S5_genind[49]
# Line_21_6_S2_genind <- FS_S5_genind[77]
# Line_21_6_S3_genind <- FS_S5_genind[105]
# Line_21_6_S4_genind <- FS_S5_genind[133]
# Line_21_6_S5_genind <- FS_S5_genind[148]
# 
# Line_23_2_F_genind <- FS_S5_genind[22]
# Line_23_2_S1_genind <- FS_S5_genind[50]
# Line_23_2_S2_genind <- FS_S5_genind[78]
# Line_23_2_S3_genind <- FS_S5_genind[106]
# Line_23_2_S4_genind <- FS_S5_genind[134]
# Line_23_2_S5_genind <- FS_S5_genind[149]
# 
# Line_23_4_F_genind <- FS_S5_genind[23]
# Line_23_4_S1_genind <- FS_S5_genind[51]
# Line_23_4_S2_genind <- FS_S5_genind[79]
# Line_23_4_S3_genind <- FS_S5_genind[107]
# Line_23_4_S4_genind <- FS_S5_genind[135]
# 
# Line_26_1_F_genind <- FS_S5_genind[24]
# Line_26_1_S1_genind <- FS_S5_genind[52]
# Line_26_1_S2_genind <- FS_S5_genind[80]
# Line_26_1_S3_genind <- FS_S5_genind[108]
# Line_26_1_S4_genind <- FS_S5_genind[136]
# 
# Line_26_4_F_genind <- FS_S5_genind[25]
# Line_26_4_S1_genind <- FS_S5_genind[53]
# Line_26_4_S2_genind <- FS_S5_genind[81]
# Line_26_4_S3_genind <- FS_S5_genind[109]
# Line_26_4_S4_genind <- FS_S5_genind[137]
# 
# Line_27_2_F_genind <- FS_S5_genind[26]
# Line_27_2_S1_genind <- FS_S5_genind[54]
# Line_27_2_S2_genind <- FS_S5_genind[82]
# Line_27_2_S3_genind <- FS_S5_genind[110]
# Line_27_2_S4_genind <- FS_S5_genind[138]
# 
# Line_29_2_F_genind <- FS_S5_genind[27]
# Line_29_2_S1_genind <- FS_S5_genind[55]
# Line_29_2_S2_genind <- FS_S5_genind[83]
# Line_29_2_S3_genind <- FS_S5_genind[111]
# Line_29_2_S4_genind <- FS_S5_genind[139]
# Line_29_2_S5_genind <- FS_S5_genind[150]
# 
# Line_29_4_F_genind <- FS_S5_genind[28]
# Line_29_4_S1_genind <- FS_S5_genind[56]
# Line_29_4_S2_genind <- FS_S5_genind[84]
# Line_29_4_S3_genind <- FS_S5_genind[112]
# Line_29_4_S4_genind <- FS_S5_genind[140]
# Line_29_4_S5_genind <- FS_S5_genind[151]
# 
# 
# # FS_S4_clusters <- find.clusters(FS_S4_genind)
# # FS_S4_dapc <- dapc(FS_S4_genind, FS_S4_clusters$grp)
# # scatter(FS_S4_dapc)
# # 
# # FS_S5_stats <- genind2hierfstat(FS_S5_genind, pop = popmap_generations$V2)
# # FS_S4_stats_FS_lines <- genind2hierfstat(FS_S4_genind, pop = popmap_FS_lines$V2)
# # FS_S5_stats_parental_lines <- genind2hierfstat(FS_S4_genind, pop = popmap_parental_lines$V2)
# # 
# # # Genotype PCA
# # x <- indpca(FS_S5_stats)
# # 
# # pdf(file = "FS-S5_generations_PCA_small.pdf")
# # plot(x, cex = 0.2)
# # dev.off()
# # 
# # y <- indpca(FS_S5_stats_FS_lines)
# # 
# # pdf(file = "FS-S5_lines_PCA_small.pdf")
# # plot(y, cex = 0.2)
# # dev.off()
# # 
# # z <- indpca(FS_S5_stats_parental_lines)
# # 
# # pdf(file = "FS-S5_parental_lines_PCA_small.pdf")
# # plot(z, cex = 0.2)
# # dev.off()
# 
# # Get summary for heterozygosity
# 
# Line_1_1_F_div <- summary(Line_1_1_F_genind)
# Line_1_1_S1_div <- summary(Line_1_1_S1_genind)
# Line_1_1_S2_div <- summary(Line_1_1_S2_genind)
# Line_1_1_S3_div <- summary(Line_1_1_S3_genind)
# Line_1_1_S4_div <- summary(Line_1_1_S4_genind)
# Line_1_1_S5_div <- summary(Line_1_1_S5_genind)
# 
# Line_6_1_F_div <- summary(Line_6_1_F_genind)
# Line_6_1_S1_div <- summary(Line_6_1_S1_genind)
# Line_6_1_S2_div <- summary(Line_6_1_S2_genind)
# Line_6_1_S3_div <- summary(Line_6_1_S3_genind)
# Line_6_1_S4_div <- summary(Line_6_1_S4_genind)
# 
# Line_6_4_F_div <- summary(Line_6_4_F_genind)
# Line_6_4_S1_div <- summary(Line_6_4_S1_genind)
# Line_6_4_S2_div <- summary(Line_6_4_S2_genind)
# Line_6_4_S3_div <- summary(Line_6_4_S3_genind)
# Line_6_4_S4_div <- summary(Line_6_4_S4_genind)
# Line_6_4_S5_div <- summary(Line_6_4_S5_genind)
# 
# Line_7_2_F_div <- summary(Line_7_2_F_genind)
# Line_7_2_S1_div <- summary(Line_7_2_S1_genind)
# Line_7_2_S2_div <- summary(Line_7_2_S2_genind)
# Line_7_2_S3_div <- summary(Line_7_2_S3_genind)
# Line_7_2_S4_div <- summary(Line_7_2_S4_genind)
# 
# Line_7_4_F_div <- summary(Line_7_4_F_genind)
# Line_7_4_S1_div <- summary(Line_7_4_S1_genind)
# Line_7_4_S2_div <- summary(Line_7_4_S2_genind)
# Line_7_4_S3_div <- summary(Line_7_4_S3_genind)
# Line_7_4_S4_div <- summary(Line_7_4_S4_genind)
# 
# Line_8_2_F_div <- summary(Line_8_2_F_genind)
# Line_8_2_S1_div <- summary(Line_8_2_S1_genind)
# Line_8_2_S2_div <- summary(Line_8_2_S2_genind)
# Line_8_2_S3_div <- summary(Line_8_2_S3_genind)
# Line_8_2_S4_div <- summary(Line_8_2_S4_genind)
# 
# Line_8_4_F_div <- summary(Line_8_4_F_genind)
# Line_8_4_S1_div <- summary(Line_8_4_S1_genind)
# Line_8_4_S2_div <- summary(Line_8_4_S2_genind)
# Line_8_4_S3_div <- summary(Line_8_4_S3_genind)
# Line_8_4_S4_div <- summary(Line_8_4_S4_genind)
# 
# Line_10_2_F_div <- summary(Line_10_2_F_genind)
# Line_10_2_S1_div <- summary(Line_10_2_S1_genind)
# Line_10_2_S2_div <- summary(Line_10_2_S2_genind)
# Line_10_2_S3_div <- summary(Line_10_2_S3_genind)
# Line_10_2_S4_div <- summary(Line_10_2_S4_genind)
# Line_10_2_S5_div <- summary(Line_10_2_S5_genind)
# 
# Line_10_5_F_div <- summary(Line_10_5_F_genind)
# Line_10_5_S1_div <- summary(Line_10_5_S1_genind)
# Line_10_5_S2_div <- summary(Line_10_5_S2_genind)
# Line_10_5_S3_div <- summary(Line_10_5_S3_genind)
# Line_10_5_S4_div <- summary(Line_10_5_S4_genind)
# 
# Line_13_3_F_div <- summary(Line_13_3_F_genind)
# Line_13_3_S1_div <- summary(Line_13_3_S1_genind)
# Line_13_3_S2_div <- summary(Line_13_3_S2_genind)
# Line_13_3_S3_div <- summary(Line_13_3_S3_genind)
# Line_13_3_S4_div <- summary(Line_13_3_S4_genind)
# 
# Line_13_4_F_div <- summary(Line_13_4_F_genind)
# Line_13_4_S1_div <- summary(Line_13_4_S1_genind)
# Line_13_4_S2_div <- summary(Line_13_4_S2_genind)
# Line_13_4_S3_div <- summary(Line_13_4_S3_genind)
# Line_13_4_S4_div <- summary(Line_13_4_S4_genind)
# 
# Line_16_1_F_div <- summary(Line_16_1_F_genind)
# Line_16_1_S1_div <- summary(Line_16_1_S1_genind)
# Line_16_1_S2_div <- summary(Line_16_1_S2_genind)
# Line_16_1_S3_div <- summary(Line_16_1_S3_genind)
# Line_16_1_S4_div <- summary(Line_16_1_S4_genind)
# 
# Line_16_5_F_div <- summary(Line_16_5_F_genind)
# Line_16_5_S1_div <- summary(Line_16_5_S1_genind)
# Line_16_5_S2_div <- summary(Line_16_5_S2_genind)
# Line_16_5_S3_div <- summary(Line_16_5_S3_genind)
# Line_16_5_S4_div <- summary(Line_16_5_S4_genind)
# Line_16_5_S5_div <- summary(Line_16_5_S5_genind)
# 
# Line_17_2_F_div <- summary(Line_17_2_F_genind)
# Line_17_2_S1_div <- summary(Line_17_2_S1_genind)
# Line_17_2_S2_div <- summary(Line_17_2_S2_genind)
# Line_17_2_S3_div <- summary(Line_17_2_S3_genind)
# Line_17_2_S4_div <- summary(Line_17_2_S4_genind)
# Line_17_2_S5_div <- summary(Line_17_2_S5_genind)
# 
# Line_17_5_F_div <- summary(Line_17_5_F_genind)
# Line_17_5_S1_div <- summary(Line_17_5_S1_genind)
# Line_17_5_S2_div <- summary(Line_17_5_S2_genind)
# Line_17_5_S3_div <- summary(Line_17_5_S3_genind)
# Line_17_5_S4_div <- summary(Line_17_5_S4_genind)
# Line_17_5_S5_div <- summary(Line_17_5_S5_genind)
# 
# Line_19_2_F_div <- summary(Line_19_2_F_genind)
# Line_19_2_S1_div <- summary(Line_19_2_S1_genind)
# Line_19_2_S2_div <- summary(Line_19_2_S2_genind)
# Line_19_2_S3_div <- summary(Line_19_2_S3_genind)
# Line_19_2_S4_div <- summary(Line_19_2_S4_genind)
# 
# Line_19_5_F_div <- summary(Line_19_5_F_genind)
# Line_19_5_S1_div <- summary(Line_19_5_S1_genind)
# Line_19_5_S2_div <- summary(Line_19_5_S2_genind)
# Line_19_5_S3_div <- summary(Line_19_5_S3_genind)
# Line_19_5_S4_div <- summary(Line_19_5_S4_genind)
# 
# Line_20_1_F_div <- summary(Line_20_1_F_genind)
# Line_20_1_S1_div <- summary(Line_20_1_S1_genind)
# Line_20_1_S2_div <- summary(Line_20_1_S2_genind)
# Line_20_1_S3_div <- summary(Line_20_1_S3_genind)
# Line_20_1_S4_div <- summary(Line_20_1_S4_genind)
# 
# Line_20_4_F_div <- summary(Line_20_4_F_genind)
# Line_20_4_S1_div <- summary(Line_20_4_S1_genind)
# Line_20_4_S2_div <- summary(Line_20_4_S2_genind)
# Line_20_4_S3_div <- summary(Line_20_4_S3_genind)
# Line_20_4_S4_div <- summary(Line_20_4_S4_genind)
# 
# Line_21_2_F_div <- summary(Line_21_2_F_genind)
# Line_21_2_S1_div <- summary(Line_21_2_S1_genind)
# Line_21_2_S2_div <- summary(Line_21_2_S2_genind)
# Line_21_2_S3_div <- summary(Line_21_2_S3_genind)
# Line_21_2_S4_div <- summary(Line_21_2_S4_genind)
# Line_21_2_S5_div <- summary(Line_21_2_S5_genind)
# 
# Line_21_6_F_div <- summary(Line_21_6_F_genind)
# Line_21_6_S1_div <- summary(Line_21_6_S1_genind)
# Line_21_6_S2_div <- summary(Line_21_6_S2_genind)
# Line_21_6_S3_div <- summary(Line_21_6_S3_genind)
# Line_21_6_S4_div <- summary(Line_21_6_S4_genind)
# Line_21_6_S5_div <- summary(Line_21_6_S5_genind)
# 
# Line_23_2_F_div <- summary(Line_23_2_F_genind)
# Line_23_2_S1_div <- summary(Line_23_2_S1_genind)
# Line_23_2_S2_div <- summary(Line_23_2_S2_genind)
# Line_23_2_S3_div <- summary(Line_23_2_S3_genind)
# Line_23_2_S4_div <- summary(Line_23_2_S4_genind)
# Line_23_2_S5_div <- summary(Line_23_2_S5_genind)
# 
# Line_23_4_F_div <- summary(Line_23_4_F_genind)
# Line_23_4_S1_div <- summary(Line_23_4_S1_genind)
# Line_23_4_S2_div <- summary(Line_23_4_S2_genind)
# Line_23_4_S3_div <- summary(Line_23_4_S3_genind)
# Line_23_4_S4_div <- summary(Line_23_4_S4_genind)
# 
# Line_26_1_F_div <- summary(Line_26_1_F_genind)
# Line_26_1_S1_div <- summary(Line_26_1_S1_genind)
# Line_26_1_S2_div <- summary(Line_26_1_S2_genind)
# Line_26_1_S3_div <- summary(Line_26_1_S3_genind)
# Line_26_1_S4_div <- summary(Line_26_1_S4_genind)
# 
# Line_26_4_F_div <- summary(Line_26_4_F_genind)
# Line_26_4_S1_div <- summary(Line_26_4_S1_genind)
# Line_26_4_S2_div <- summary(Line_26_4_S2_genind)
# Line_26_4_S3_div <- summary(Line_26_4_S3_genind)
# Line_26_4_S4_div <- summary(Line_26_4_S4_genind)
# 
# Line_27_2_F_div <- summary(Line_27_2_F_genind)
# Line_27_2_S1_div <- summary(Line_27_2_S1_genind)
# Line_27_2_S2_div <- summary(Line_27_2_S2_genind)
# Line_27_2_S3_div <- summary(Line_27_2_S3_genind)
# Line_27_2_S4_div <- summary(Line_27_2_S4_genind)
# 
# Line_29_2_F_div <- summary(Line_29_2_F_genind)
# Line_29_2_S1_div <- summary(Line_29_2_S1_genind)
# Line_29_2_S2_div <- summary(Line_29_2_S2_genind)
# Line_29_2_S3_div <- summary(Line_29_2_S3_genind)
# Line_29_2_S4_div <- summary(Line_29_2_S4_genind)
# Line_29_2_S5_div <- summary(Line_29_2_S5_genind)
# 
# Line_29_4_F_div <- summary(Line_29_4_F_genind)
# Line_29_4_S1_div <- summary(Line_29_4_S1_genind)
# Line_29_4_S2_div <- summary(Line_29_4_S2_genind)
# Line_29_4_S3_div <- summary(Line_29_4_S3_genind)
# Line_29_4_S4_div <- summary(Line_29_4_S4_genind)
# Line_29_4_S5_div <- summary(Line_29_4_S5_genind)
# 
# mean(F_lines_div$Hobs)
# mean(F_lines_div$Hexp)
# 
# mean(S1_lines_div$Hobs)
# mean(S1_lines_div$Hexp)
# 
# mean(S2_lines_div$Hobs)
# mean(S2_lines_div$Hexp)
# 
# mean(S3_lines_div$Hobs)
# mean(S3_lines_div$Hexp)
# 
# mean(S4_lines_div$Hobs)
# mean(S4_lines_div$Hexp)
# 
# mean(S5_lines_div$Hobs)
# mean(S5_lines_div$Hexp)
# 
# # write.table(data.frame(F_lines_div$Hobs, F_lines_div$Hexp), "F_lines_Het.txt", quote = F)
# # write.table(data.frame(S1_lines_div$Hobs, S1_lines_div$Hexp), "S1_lines_Het.txt", quote = F)
# # write.table(data.frame(S2_lines_div$Hobs, S2_lines_div$Hexp), "S2_lines_Het.txt", quote = F)
# # write.table(data.frame(S3_lines_div$Hobs, S3_lines_div$Hexp), "S3_lines_Het.txt", quote = F)
# # write.table(data.frame(S4_lines_div$Hobs, S4_lines_div$Hexp), "S4_lines_Het.txt", quote = F)
# # write.table(data.frame(S5_lines_div$Hobs, S5_lines_div$Hexp), "S5_lines_Het.txt", quote = F)
# 
# # Combine generations for figure
# # Line 1_1
# generation_hobs_Line_1_1 <- data.frame(F_Hobs = Line_1_1_F_div$Hobs,
#                                        S1_Hobs = Line_1_1_S1_div$Hobs,
#                                        S2_Hobs = Line_1_1_S2_div$Hobs,
#                                        S3_Hobs = Line_1_1_S3_div$Hobs,
#                                        S4_Hobs = Line_1_1_S4_div$Hobs,
#                                        S5_Hobs = Line_1_1_S5_div$Hobs)
# 
# colnames(generation_hobs_Line_1_1) <- c("F", "S1", "S2", "S3", "S4", "S5")
# # generation_hobs <- rownames_to_column(generation_hobs_Line_1_1)
# melted_generation_hobs_Line_1_1 <- melt(generation_hobs_Line_1_1)
# 
# generation_hobs_Line_1_1_summary <- data.frame(FS = mean(generation_hobs_Line_1_1$F, na.rm = T), 
#                                                S1 = mean(generation_hobs_Line_1_1$S1, na.rm = T),
#                                                S2 = mean(generation_hobs_Line_1_1$S2, na.rm = T),
#                                                S3 = mean(generation_hobs_Line_1_1$S3, na.rm = T),
#                                                S4 = mean(generation_hobs_Line_1_1$S4, na.rm = T),
#                                                S5 = mean(generation_hobs_Line_1_1$S5, na.rm = T),
#                                                H = "Observed")
# 
# melted_generation_hobs_Line_1_1_summary <- melt(generation_hobs_Line_1_1_summary)
# 
# ggplot(melted_generation_hobs_Line_1_1, aes(x = value, fill = variable)) +
#   geom_density(alpha = 0.3) +
#   theme_Publication()
# 
# ggplot(melted_generation_hobs_Line_1_1_summary, aes(x = variable, y = value)) +
#   geom_point() +
#   scale_y_continuous(breaks = c(0.1, 0.2, 0.3, 0.4, 0.5 ,0.6, 0.7, 0.8, 0.9, 1)) +
#   ylim(c(0,1))
# 
# expectations_halving_Line_1_1 <- melted_generation_hobs_Line_1_1 %>% 
#   filter(variable == "F") %>% 
#   mutate(S1 = (value / 2), S2 = (value / 4), S3 = (value / 8), S4 = (value/ 16), S5 = (value / 32)) %>% 
#   rename(F = value) %>% 
#   select(F, S1, S2, S3, S4, S5)
# 
# expectations_halving_Line_1_1_summary <- data.frame(FS = mean(expectations_halving_Line_1_1$F, na.rm = T), 
#                                                     S1 = mean(expectations_halving_Line_1_1$S1, na.rm = T),
#                                                     S2 = mean(expectations_halving_Line_1_1$S2, na.rm = T),
#                                                     S3 = mean(expectations_halving_Line_1_1$S3, na.rm = T),
#                                                     S4 = mean(expectations_halving_Line_1_1$S4, na.rm = T),
#                                                     S5 = mean(expectations_halving_Line_1_1$S5, na.rm = T),
#                                                     H = "Expected")
# expectations_halving_Line_1_1_summary_melted <- melt(expectations_halving_Line_1_1_summary)
# 
# obs_exp_1_1 <- rbind(melted_generation_hobs_Line_1_1_summary, expectations_halving_Line_1_1_summary_melted)
# 
# obs_exp_1_1_plot <- ggplot(obs_exp_1_1, aes(x = variable, y = value, colour = H)) +
#   geom_point(size = 3) +
#   scale_y_continuous(breaks = c(0.1, 0.2, 0.3, 0.4, 0.5 ,0.6, 0.7, 0.8, 0.9, 1)) +
#   ylim(c(0,1)) +
#   scale_colour_nejm() +
#   ylab("Heterozygosity") +
#   xlab("Generation") +
#   theme_Publication() +
#   theme(legend.position = "none") +
#   ggtitle("Line 1_1")
# 
# 
# wilcox.test(generation_hobs_Line_1_1$S1, expectations_halving_Line_1_1$S1, paired = T)
# wilcox.test(generation_hobs_Line_1_1$S2, expectations_halving_Line_1_1$S2, paired = T)
# wilcox.test(generation_hobs_Line_1_1$S3, expectations_halving_Line_1_1$S3, paired = T)
# wilcox.test(generation_hobs_Line_1_1$S4, expectations_halving_Line_1_1$S4, paired = T)
# wilcox.test(generation_hobs_Line_1_1$S5, expectations_halving_Line_1_1$S5, paired = T)
# 
# write.table(generation_hobs_Line_1_1, "Line_1_1_hobs.txt", col.names = T, row.names = T, quote = F)
# write.table(expectations_halving_Line_1_1, "Line_1_1_hexp.txt", col.names = T, row.names = F, quote = F)
# 
# # Line 6_1
# generation_hobs_Line_6_1 <- data.frame(F_Hobs = Line_6_1_F_div$Hobs,
#                                        S1_Hobs = Line_6_1_S1_div$Hobs,
#                                        S2_Hobs = Line_6_1_S2_div$Hobs,
#                                        S3_Hobs = Line_6_1_S3_div$Hobs,
#                                        S4_Hobs = Line_6_1_S4_div$Hobs)
# 
# colnames(generation_hobs_Line_6_1) <- c("F", "S1", "S2", "S3", "S4")
# # generation_hobs <- rownames_to_column(generation_hobs_Line_6_1)
# melted_generation_hobs_Line_6_1 <- melt(generation_hobs_Line_6_1)
# 
# generation_hobs_Line_6_1_summary <- data.frame(FS = mean(generation_hobs_Line_6_1$F, na.rm = T), 
#                                                S1 = mean(generation_hobs_Line_6_1$S1, na.rm = T),
#                                                S2 = mean(generation_hobs_Line_6_1$S2, na.rm = T),
#                                                S3 = mean(generation_hobs_Line_6_1$S3, na.rm = T),
#                                                S4 = mean(generation_hobs_Line_6_1$S4, na.rm = T),
#                                                H = "Observed")
# 
# melted_generation_hobs_Line_6_1_summary <- melt(generation_hobs_Line_6_1_summary)
# 
# ggplot(melted_generation_hobs_Line_6_1, aes(x = value, fill = variable)) +
#   geom_density(alpha = 0.3) +
#   theme_Publication()
# 
# ggplot(melted_generation_hobs_Line_6_1_summary, aes(x = variable, y = value)) +
#   geom_point() +
#   scale_y_continuous(breaks = c(0.1, 0.2, 0.3, 0.4, 0.5 ,0.6, 0.7, 0.8, 0.9, 1)) +
#   ylim(c(0,1))
# 
# expectations_halving_Line_6_1 <- melted_generation_hobs_Line_6_1 %>% 
#   filter(variable == "F") %>% 
#   mutate(S1 = (value / 2), S2 = (value / 4), S3 = (value / 8), S4 = (value/ 16)) %>% 
#   rename(F = value) %>% 
#   select(F, S1, S2, S3, S4)
# 
# expectations_halving_Line_6_1_summary <- data.frame(FS = mean(expectations_halving_Line_6_1$F, na.rm = T), 
#                                                     S1 = mean(expectations_halving_Line_6_1$S1, na.rm = T),
#                                                     S2 = mean(expectations_halving_Line_6_1$S2, na.rm = T),
#                                                     S3 = mean(expectations_halving_Line_6_1$S3, na.rm = T),
#                                                     S4 = mean(expectations_halving_Line_6_1$S4, na.rm = T),
#                                                     H = "Expected")
# expectations_halving_Line_6_1_summary_melted <- melt(expectations_halving_Line_6_1_summary)
# 
# obs_exp_6_1 <- rbind(expectations_halving_Line_6_1_summary_melted, melted_generation_hobs_Line_6_1_summary)
# 
# obs_exp_6_1_plot <- ggplot(obs_exp_6_1, aes(x = variable, y = value, colour = H)) +
#   geom_point(size = 3) +
#   scale_y_continuous(breaks = c(0.1, 0.2, 0.3, 0.4, 0.5 ,0.6, 0.7, 0.8, 0.9, 1)) +
#   ylim(c(0,1)) +
#   scale_colour_nejm() +
#   ylab("Heterozygosity") +
#   xlab("Generation") +
#   theme_Publication() +
#   theme(legend.position = "none") +
#   ggtitle("Line 6_1")
# 
# wilcox.test(generation_hobs_Line_6_1$S1, expectations_halving_Line_6_1$S1, paired = T)
# wilcox.test(generation_hobs_Line_6_1$S2, expectations_halving_Line_6_1$S2, paired = T)
# wilcox.test(generation_hobs_Line_6_1$S3, expectations_halving_Line_6_1$S3, paired = T)
# wilcox.test(generation_hobs_Line_6_1$S4, expectations_halving_Line_6_1$S4, paired = T)
# 
# write.table(generation_hobs_Line_6_1, "Line_6_1_hobs.txt", col.names = T, row.names = T, quote = F)
# write.table(expectations_halving_Line_6_1, "Line_6_1_hexp.txt", col.names = T, row.names = F, quote = F)
# 
# # Line 6_4
# generation_hobs_Line_6_4 <- data.frame(F_Hobs = Line_6_4_F_div$Hobs,
#                                        S1_Hobs = Line_6_4_S1_div$Hobs,
#                                        S2_Hobs = Line_6_4_S2_div$Hobs,
#                                        S3_Hobs = Line_6_4_S3_div$Hobs,
#                                        S4_Hobs = Line_6_4_S4_div$Hobs,
#                                        S5_Hobs = Line_6_4_S5_div$Hobs)
# 
# colnames(generation_hobs_Line_6_4) <- c("F", "S1", "S2", "S3", "S4", "S5")
# # generation_hobs <- rownames_to_column(generation_hobs_Line_6_4)
# melted_generation_hobs_Line_6_4 <- melt(generation_hobs_Line_6_4)
# 
# generation_hobs_Line_6_4_summary <- data.frame(FS = mean(generation_hobs_Line_6_4$F, na.rm = T), 
#                                                S1 = mean(generation_hobs_Line_6_4$S1, na.rm = T),
#                                                S2 = mean(generation_hobs_Line_6_4$S2, na.rm = T),
#                                                S3 = mean(generation_hobs_Line_6_4$S3, na.rm = T),
#                                                S4 = mean(generation_hobs_Line_6_4$S4, na.rm = T),
#                                                S5 = mean(generation_hobs_Line_6_4$S5, na.rm = T),
#                                                H = "Observed")
# 
# melted_generation_hobs_Line_6_4_summary <- melt(generation_hobs_Line_6_4_summary)
# 
# ggplot(melted_generation_hobs_Line_6_4, aes(x = value, fill = variable)) +
#   geom_density(alpha = 0.3) +
#   theme_Publication()
# 
# ggplot(melted_generation_hobs_Line_6_4_summary, aes(x = variable, y = value)) +
#   geom_point() +
#   scale_y_continuous(breaks = c(0.1, 0.2, 0.3, 0.4, 0.5 ,0.6, 0.7, 0.8, 0.9, 1)) +
#   ylim(c(0,1))
# 
# expectations_halving_Line_6_4 <- melted_generation_hobs_Line_6_4 %>% 
#   filter(variable == "F") %>% 
#   mutate(S1 = (value / 2), S2 = (value / 4), S3 = (value / 8), S4 = (value/ 16), S5 = (value / 32)) %>% 
#   rename(F = value) %>% 
#   select(F, S1, S2, S3, S4, S5)
# 
# expectations_halving_Line_6_4_summary <- data.frame(FS = mean(expectations_halving_Line_6_4$F, na.rm = T), 
#                                                     S1 = mean(expectations_halving_Line_6_4$S1, na.rm = T),
#                                                     S2 = mean(expectations_halving_Line_6_4$S2, na.rm = T),
#                                                     S3 = mean(expectations_halving_Line_6_4$S3, na.rm = T),
#                                                     S4 = mean(expectations_halving_Line_6_4$S4, na.rm = T),
#                                                     S5 = mean(expectations_halving_Line_6_4$S5, na.rm = T),
#                                                     H = "Expected")
# expectations_halving_Line_6_4_summary_melted <- melt(expectations_halving_Line_6_4_summary)
# 
# obs_exp_6_4 <- rbind(expectations_halving_Line_6_4_summary_melted, melted_generation_hobs_Line_6_4_summary)
# 
# obs_exp_6_4_plot <- ggplot(obs_exp_6_4, aes(x = variable, y = value, colour = H)) +
#   geom_point(size = 3) +
#   scale_y_continuous(breaks = c(0.1, 0.2, 0.3, 0.4, 0.5 ,0.6, 0.7, 0.8, 0.9, 1)) +
#   ylim(c(0,1)) +
#   scale_colour_nejm() +
#   ylab("Heterozygosity") +
#   xlab("Generation") +
#   theme_Publication() +
#   theme(legend.position = "none") +
#   ggtitle("Line 6_4")
# 
# wilcox.test(generation_hobs_Line_6_4$S1, expectations_halving_Line_6_4$S1, paired = T)
# wilcox.test(generation_hobs_Line_6_4$S2, expectations_halving_Line_6_4$S2, paired = T)
# wilcox.test(generation_hobs_Line_6_4$S3, expectations_halving_Line_6_4$S3, paired = T)
# wilcox.test(generation_hobs_Line_6_4$S4, expectations_halving_Line_6_4$S4, paired = T)
# wilcox.test(generation_hobs_Line_6_4$S5, expectations_halving_Line_6_4$S5, paired = T)
# 
# write.table(generation_hobs_Line_6_4, "Line_6_4_hobs.txt", col.names = T, row.names = T, quote = F)
# write.table(expectations_halving_Line_6_4, "Line_6_4_hexp.txt", col.names = T, row.names = F, quote = F)
# 
# # Line 7_2
# generation_hobs_Line_7_2 <- data.frame(F_Hobs = Line_7_2_F_div$Hobs,
#                                        S1_Hobs = Line_7_2_S1_div$Hobs,
#                                        S2_Hobs = Line_7_2_S2_div$Hobs,
#                                        S3_Hobs = Line_7_2_S3_div$Hobs,
#                                        S4_Hobs = Line_7_2_S4_div$Hobs)
# 
# colnames(generation_hobs_Line_7_2) <- c("F", "S1", "S2", "S3", "S4")
# # generation_hobs <- rownames_to_column(generation_hobs_Line_7_2)
# melted_generation_hobs_Line_7_2 <- melt(generation_hobs_Line_7_2)
# 
# generation_hobs_Line_7_2_summary <- data.frame(FS = mean(generation_hobs_Line_7_2$F, na.rm = T), 
#                                                S1 = mean(generation_hobs_Line_7_2$S1, na.rm = T),
#                                                S2 = mean(generation_hobs_Line_7_2$S2, na.rm = T),
#                                                S3 = mean(generation_hobs_Line_7_2$S3, na.rm = T),
#                                                S4 = mean(generation_hobs_Line_7_2$S4, na.rm = T),
#                                                H = "Observed")
# 
# melted_generation_hobs_Line_7_2_summary <- melt(generation_hobs_Line_7_2_summary)
# 
# ggplot(melted_generation_hobs_Line_7_2, aes(x = value, fill = variable)) +
#   geom_density(alpha = 0.3) +
#   theme_Publication()
# 
# ggplot(melted_generation_hobs_Line_7_2_summary, aes(x = variable, y = value)) +
#   geom_point() +
#   scale_y_continuous(breaks = c(0.1, 0.2, 0.3, 0.4, 0.5 ,0.6, 0.7, 0.8, 0.9, 1)) +
#   ylim(c(0,1))
# 
# expectations_halving_Line_7_2 <- melted_generation_hobs_Line_7_2 %>% 
#   filter(variable == "F") %>% 
#   mutate(S1 = (value / 2), S2 = (value / 4), S3 = (value / 8), S4 = (value/ 16)) %>% 
#   rename(F = value) %>% 
#   select(F, S1, S2, S3, S4)
# 
# expectations_halving_Line_7_2_summary <- data.frame(FS = mean(expectations_halving_Line_7_2$F, na.rm = T), 
#                                                     S1 = mean(expectations_halving_Line_7_2$S1, na.rm = T),
#                                                     S2 = mean(expectations_halving_Line_7_2$S2, na.rm = T),
#                                                     S3 = mean(expectations_halving_Line_7_2$S3, na.rm = T),
#                                                     S4 = mean(expectations_halving_Line_7_2$S4, na.rm = T),
#                                                     H = "Expected")
# expectations_halving_Line_7_2_summary_melted <- melt(expectations_halving_Line_7_2_summary)
# 
# obs_exp_7_2 <- rbind(expectations_halving_Line_7_2_summary_melted, melted_generation_hobs_Line_7_2_summary)
# 
# obs_exp_7_2_plot <- ggplot(obs_exp_7_2, aes(x = variable, y = value, colour = H)) +
#   geom_point(size = 3) +
#   scale_y_continuous(breaks = c(0.1, 0.2, 0.3, 0.4, 0.5 ,0.6, 0.7, 0.8, 0.9, 1)) +
#   ylim(c(0,1)) +
#   scale_colour_nejm() +
#   ylab("Heterozygosity") +
#   xlab("Generation") +
#   theme_Publication() +
#   theme(legend.position = "none") +
#   ggtitle("Line 7_2")
# 
# wilcox.test(generation_hobs_Line_7_2$S1, expectations_halving_Line_7_2$S1, paired = T)
# wilcox.test(generation_hobs_Line_7_2$S2, expectations_halving_Line_7_2$S2, paired = T)
# wilcox.test(generation_hobs_Line_7_2$S3, expectations_halving_Line_7_2$S3, paired = T)
# wilcox.test(generation_hobs_Line_7_2$S4, expectations_halving_Line_7_2$S4, paired = T)
# 
# write.table(generation_hobs_Line_7_2, "Line_7_2_hobs.txt", col.names = T, row.names = T, quote = F)
# write.table(expectations_halving_Line_7_2, "Line_7_2_hexp.txt", col.names = T, row.names = F, quote = F)
# 
# # Line 7_4
# generation_hobs_Line_7_4 <- data.frame(F_Hobs = Line_7_4_F_div$Hobs,
#                                        S1_Hobs = Line_7_4_S1_div$Hobs,
#                                        S2_Hobs = Line_7_4_S2_div$Hobs,
#                                        S3_Hobs = Line_7_4_S3_div$Hobs,
#                                        S4_Hobs = Line_7_4_S4_div$Hobs)
# 
# colnames(generation_hobs_Line_7_4) <- c("F", "S1", "S2", "S3", "S4")
# # generation_hobs <- rownames_to_column(generation_hobs_Line_7_4)
# melted_generation_hobs_Line_7_4 <- melt(generation_hobs_Line_7_4)
# 
# generation_hobs_Line_7_4_summary <- data.frame(FS = mean(generation_hobs_Line_7_4$F, na.rm = T), 
#                                                S1 = mean(generation_hobs_Line_7_4$S1, na.rm = T),
#                                                S2 = mean(generation_hobs_Line_7_4$S2, na.rm = T),
#                                                S3 = mean(generation_hobs_Line_7_4$S3, na.rm = T),
#                                                S4 = mean(generation_hobs_Line_7_4$S4, na.rm = T),
#                                                H = "Observed")
# 
# melted_generation_hobs_Line_7_4_summary <- melt(generation_hobs_Line_7_4_summary)
# 
# ggplot(melted_generation_hobs_Line_7_4, aes(x = value, fill = variable)) +
#   geom_density(alpha = 0.3) +
#   theme_Publication()
# 
# ggplot(melted_generation_hobs_Line_7_4_summary, aes(x = variable, y = value)) +
#   geom_point() +
#   scale_y_continuous(breaks = c(0.1, 0.2, 0.3, 0.4, 0.5 ,0.6, 0.7, 0.8, 0.9, 1)) +
#   ylim(c(0,1))
# 
# expectations_halving_Line_7_4 <- melted_generation_hobs_Line_7_4 %>% 
#   filter(variable == "F") %>% 
#   mutate(S1 = (value / 2), S2 = (value / 4), S3 = (value / 8), S4 = (value/ 16)) %>% 
#   rename(F = value) %>% 
#   select(F, S1, S2, S3, S4)
# 
# expectations_halving_Line_7_4_summary <- data.frame(FS = mean(expectations_halving_Line_7_4$F, na.rm = T), 
#                                                     S1 = mean(expectations_halving_Line_7_4$S1, na.rm = T),
#                                                     S2 = mean(expectations_halving_Line_7_4$S2, na.rm = T),
#                                                     S3 = mean(expectations_halving_Line_7_4$S3, na.rm = T),
#                                                     S4 = mean(expectations_halving_Line_7_4$S4, na.rm = T),
#                                                     H = "Expected")
# expectations_halving_Line_7_4_summary_melted <- melt(expectations_halving_Line_7_4_summary)
# 
# obs_exp_7_4 <- rbind(expectations_halving_Line_7_4_summary_melted, melted_generation_hobs_Line_7_4_summary)
# 
# obs_exp_7_4_plot <- ggplot(obs_exp_7_4, aes(x = variable, y = value, colour = H)) +
#   geom_point(size = 3) +
#   scale_y_continuous(breaks = c(0.1, 0.2, 0.3, 0.4, 0.5 ,0.6, 0.7, 0.8, 0.9, 1)) +
#   ylim(c(0,1)) +
#   scale_colour_nejm() +
#   ylab("Heterozygosity") +
#   xlab("Generation") +
#   theme_Publication() +
#   theme(legend.position = "none") +
#   ggtitle("Line 7_4")
# 
# wilcox.test(generation_hobs_Line_7_4$S1, expectations_halving_Line_7_4$S1, paired = T)
# wilcox.test(generation_hobs_Line_7_4$S2, expectations_halving_Line_7_4$S2, paired = T)
# wilcox.test(generation_hobs_Line_7_4$S3, expectations_halving_Line_7_4$S3, paired = T)
# wilcox.test(generation_hobs_Line_7_4$S4, expectations_halving_Line_7_4$S4, paired = T)
# 
# write.table(generation_hobs_Line_7_4, "Line_7_4_hobs.txt", col.names = T, row.names = T, quote = F)
# write.table(expectations_halving_Line_7_4, "Line_7_4_hexp.txt", col.names = T, row.names = F, quote = F)
# 
# # Line 8_2
# generation_hobs_Line_8_2 <- data.frame(F_Hobs = Line_8_2_F_div$Hobs,
#                                        S1_Hobs = Line_8_2_S1_div$Hobs,
#                                        S2_Hobs = Line_8_2_S2_div$Hobs,
#                                        S3_Hobs = Line_8_2_S3_div$Hobs,
#                                        S4_Hobs = Line_8_2_S4_div$Hobs)
# 
# colnames(generation_hobs_Line_8_2) <- c("F", "S1", "S2", "S3", "S4")
# # generation_hobs <- rownames_to_column(generation_hobs_Line_8_2)
# melted_generation_hobs_Line_8_2 <- melt(generation_hobs_Line_8_2)
# 
# generation_hobs_Line_8_2_summary <- data.frame(FS = mean(generation_hobs_Line_8_2$F, na.rm = T), 
#                                                S1 = mean(generation_hobs_Line_8_2$S1, na.rm = T),
#                                                S2 = mean(generation_hobs_Line_8_2$S2, na.rm = T),
#                                                S3 = mean(generation_hobs_Line_8_2$S3, na.rm = T),
#                                                S4 = mean(generation_hobs_Line_8_2$S4, na.rm = T),
#                                                H = "Observed")
# 
# melted_generation_hobs_Line_8_2_summary <- melt(generation_hobs_Line_8_2_summary)
# 
# ggplot(melted_generation_hobs_Line_8_2, aes(x = value, fill = variable)) +
#   geom_density(alpha = 0.3) +
#   theme_Publication()
# 
# ggplot(melted_generation_hobs_Line_8_2_summary, aes(x = variable, y = value)) +
#   geom_point() +
#   scale_y_continuous(breaks = c(0.1, 0.2, 0.3, 0.4, 0.5 ,0.6, 0.7, 0.8, 0.9, 1)) +
#   ylim(c(0,1))
# 
# expectations_halving_Line_8_2 <- melted_generation_hobs_Line_8_2 %>% 
#   filter(variable == "F") %>% 
#   mutate(S1 = (value / 2), S2 = (value / 4), S3 = (value / 8), S4 = (value/ 16)) %>% 
#   rename(F = value) %>% 
#   select(F, S1, S2, S3, S4)
# 
# expectations_halving_Line_8_2_summary <- data.frame(FS = mean(expectations_halving_Line_8_2$F, na.rm = T), 
#                                                     S1 = mean(expectations_halving_Line_8_2$S1, na.rm = T),
#                                                     S2 = mean(expectations_halving_Line_8_2$S2, na.rm = T),
#                                                     S3 = mean(expectations_halving_Line_8_2$S3, na.rm = T),
#                                                     S4 = mean(expectations_halving_Line_8_2$S4, na.rm = T),
#                                                     H = "Expected")
# expectations_halving_Line_8_2_summary_melted <- melt(expectations_halving_Line_8_2_summary)
# 
# obs_exp_8_2 <- rbind(expectations_halving_Line_8_2_summary_melted, melted_generation_hobs_Line_8_2_summary)
# 
# obs_exp_8_2_plot <- ggplot(obs_exp_8_2, aes(x = variable, y = value, colour = H)) +
#   geom_point(size = 3) +
#   scale_y_continuous(breaks = c(0.1, 0.2, 0.3, 0.4, 0.5 ,0.6, 0.7, 0.8, 0.9, 1)) +
#   ylim(c(0,1)) +
#   scale_colour_nejm() +
#   ylab("Heterozygosity") +
#   xlab("Generation") +
#   theme_Publication() +
#   theme(legend.position = "none") +
#   ggtitle("Line 8_2")
# 
# wilcox.test(generation_hobs_Line_8_2$S1, expectations_halving_Line_8_2$S1, paired = T)
# wilcox.test(generation_hobs_Line_8_2$S2, expectations_halving_Line_8_2$S2, paired = T)
# wilcox.test(generation_hobs_Line_8_2$S3, expectations_halving_Line_8_2$S3, paired = T)
# wilcox.test(generation_hobs_Line_8_2$S4, expectations_halving_Line_8_2$S4, paired = T)
# 
# write.table(generation_hobs_Line_8_2, "Line_8_2_hobs.txt", col.names = T, row.names = T, quote = F)
# write.table(expectations_halving_Line_8_2, "Line_8_2_hexp.txt", col.names = T, row.names = F, quote = F)
# 
# # Line 8_4
# generation_hobs_Line_8_4 <- data.frame(F_Hobs = Line_8_4_F_div$Hobs,
#                                        S1_Hobs = Line_8_4_S1_div$Hobs,
#                                        S2_Hobs = Line_8_4_S2_div$Hobs,
#                                        S3_Hobs = Line_8_4_S3_div$Hobs,
#                                        S4_Hobs = Line_8_4_S4_div$Hobs)
# 
# colnames(generation_hobs_Line_8_4) <- c("F", "S1", "S2", "S3", "S4")
# # generation_hobs <- rownames_to_column(generation_hobs_Line_8_4)
# melted_generation_hobs_Line_8_4 <- melt(generation_hobs_Line_8_4)
# 
# generation_hobs_Line_8_4_summary <- data.frame(FS = mean(generation_hobs_Line_8_4$F, na.rm = T), 
#                                                S1 = mean(generation_hobs_Line_8_4$S1, na.rm = T),
#                                                S2 = mean(generation_hobs_Line_8_4$S2, na.rm = T),
#                                                S3 = mean(generation_hobs_Line_8_4$S3, na.rm = T),
#                                                S4 = mean(generation_hobs_Line_8_4$S4, na.rm = T),
#                                                H = "Observed")
# 
# melted_generation_hobs_Line_8_4_summary <- melt(generation_hobs_Line_8_4_summary)
# 
# ggplot(melted_generation_hobs_Line_8_4, aes(x = value, fill = variable)) +
#   geom_density(alpha = 0.3) +
#   theme_Publication()
# 
# ggplot(melted_generation_hobs_Line_8_4_summary, aes(x = variable, y = value)) +
#   geom_point() +
#   scale_y_continuous(breaks = c(0.1, 0.2, 0.3, 0.4, 0.5 ,0.6, 0.7, 0.8, 0.9, 1)) +
#   ylim(c(0,1))
# 
# expectations_halving_Line_8_4 <- melted_generation_hobs_Line_8_4 %>% 
#   filter(variable == "F") %>% 
#   mutate(S1 = (value / 2), S2 = (value / 4), S3 = (value / 8), S4 = (value/ 16)) %>% 
#   rename(F = value) %>% 
#   select(F, S1, S2, S3, S4)
# 
# expectations_halving_Line_8_4_summary <- data.frame(FS = mean(expectations_halving_Line_8_4$F, na.rm = T), 
#                                                     S1 = mean(expectations_halving_Line_8_4$S1, na.rm = T),
#                                                     S2 = mean(expectations_halving_Line_8_4$S2, na.rm = T),
#                                                     S3 = mean(expectations_halving_Line_8_4$S3, na.rm = T),
#                                                     S4 = mean(expectations_halving_Line_8_4$S4, na.rm = T),
#                                                     H = "Expected")
# expectations_halving_Line_8_4_summary_melted <- melt(expectations_halving_Line_8_4_summary)
# 
# obs_exp_8_4 <- rbind(expectations_halving_Line_8_4_summary_melted, melted_generation_hobs_Line_8_4_summary)
# 
# obs_exp_8_4_plot <- ggplot(obs_exp_8_4, aes(x = variable, y = value, colour = H)) +
#   geom_point(size = 3) +
#   scale_y_continuous(breaks = c(0.1, 0.2, 0.3, 0.4, 0.5 ,0.6, 0.7, 0.8, 0.9, 1)) +
#   ylim(c(0,1)) +
#   scale_colour_nejm() +
#   ylab("Heterozygosity") +
#   xlab("Generation") +
#   theme_Publication() +
#   theme(legend.position = "none") +
#   ggtitle("Line 8_4")
# 
# wilcox.test(generation_hobs_Line_8_4$S1, expectations_halving_Line_8_4$S1, paired = T)
# wilcox.test(generation_hobs_Line_8_4$S2, expectations_halving_Line_8_4$S2, paired = T)
# wilcox.test(generation_hobs_Line_8_4$S3, expectations_halving_Line_8_4$S3, paired = T)
# wilcox.test(generation_hobs_Line_8_4$S4, expectations_halving_Line_8_4$S4, paired = T)
# 
# write.table(generation_hobs_Line_8_4, "Line_8_4_hobs.txt", col.names = T, row.names = T, quote = F)
# write.table(expectations_halving_Line_8_4, "Line_8_4_hexp.txt", col.names = T, row.names = F, quote = F)
# 
# # Line 10_2
# generation_hobs_Line_10_2 <- data.frame(F_Hobs = Line_10_2_F_div$Hobs,
#                                         S1_Hobs = Line_10_2_S1_div$Hobs,
#                                         S2_Hobs = Line_10_2_S2_div$Hobs,
#                                         S3_Hobs = Line_10_2_S3_div$Hobs,
#                                         S4_Hobs = Line_10_2_S4_div$Hobs,
#                                         S5_Hobs = Line_10_2_S5_div$Hobs)
# 
# colnames(generation_hobs_Line_10_2) <- c("F", "S1", "S2", "S3", "S4", "S5")
# # generation_hobs <- rownames_to_column(generation_hobs_Line_10_2)
# melted_generation_hobs_Line_10_2 <- melt(generation_hobs_Line_10_2)
# 
# generation_hobs_Line_10_2_summary <- data.frame(FS = mean(generation_hobs_Line_10_2$F, na.rm = T), 
#                                                 S1 = mean(generation_hobs_Line_10_2$S1, na.rm = T),
#                                                 S2 = mean(generation_hobs_Line_10_2$S2, na.rm = T),
#                                                 S3 = mean(generation_hobs_Line_10_2$S3, na.rm = T),
#                                                 S4 = mean(generation_hobs_Line_10_2$S4, na.rm = T),
#                                                 S5 = mean(generation_hobs_Line_10_2$S5, na.rm = T),
#                                                 H = "Observed")
# 
# melted_generation_hobs_Line_10_2_summary <- melt(generation_hobs_Line_10_2_summary)
# 
# ggplot(melted_generation_hobs_Line_10_2, aes(x = value, fill = variable)) +
#   geom_density(alpha = 0.3) +
#   theme_Publication()
# 
# ggplot(melted_generation_hobs_Line_10_2_summary, aes(x = variable, y = value)) +
#   geom_point() +
#   scale_y_continuous(breaks = c(0.1, 0.2, 0.3, 0.4, 0.5 ,0.6, 0.7, 0.8, 0.9, 1)) +
#   ylim(c(0,1))
# 
# expectations_halving_Line_10_2 <- melted_generation_hobs_Line_10_2 %>% 
#   filter(variable == "F") %>% 
#   mutate(S1 = (value / 2), S2 = (value / 4), S3 = (value / 8), S4 = (value/ 16), S5 = (value / 32)) %>% 
#   rename(F = value) %>% 
#   select(F, S1, S2, S3, S4, S5)
# 
# expectations_halving_Line_10_2_summary <- data.frame(FS = mean(expectations_halving_Line_10_2$F, na.rm = T), 
#                                                      S1 = mean(expectations_halving_Line_10_2$S1, na.rm = T),
#                                                      S2 = mean(expectations_halving_Line_10_2$S2, na.rm = T),
#                                                      S3 = mean(expectations_halving_Line_10_2$S3, na.rm = T),
#                                                      S4 = mean(expectations_halving_Line_10_2$S4, na.rm = T),
#                                                      S5 = mean(expectations_halving_Line_10_2$S5, na.rm = T),
#                                                      H = "Expected")
# expectations_halving_Line_10_2_summary_melted <- melt(expectations_halving_Line_10_2_summary)
# 
# obs_exp_10_2 <- rbind(expectations_halving_Line_10_2_summary_melted, melted_generation_hobs_Line_10_2_summary)
# 
# obs_exp_10_2_plot <- ggplot(obs_exp_10_2, aes(x = variable, y = value, colour = H)) +
#   geom_point(size = 3) +
#   scale_y_continuous(breaks = c(0.1, 0.2, 0.3, 0.4, 0.5 ,0.6, 0.7, 0.8, 0.9, 1)) +
#   ylim(c(0,1)) +
#   scale_colour_nejm() +
#   ylab("Heterozygosity") +
#   xlab("Generation") +
#   theme_Publication() +
#   theme(legend.position = "none") +
#   ggtitle("Line 10_2")
# 
# wilcox.test(generation_hobs_Line_10_2$S1, expectations_halving_Line_10_2$S1, paired = T)
# wilcox.test(generation_hobs_Line_10_2$S2, expectations_halving_Line_10_2$S2, paired = T)
# wilcox.test(generation_hobs_Line_10_2$S3, expectations_halving_Line_10_2$S3, paired = T)
# wilcox.test(generation_hobs_Line_10_2$S4, expectations_halving_Line_10_2$S4, paired = T)
# wilcox.test(generation_hobs_Line_10_2$S5, expectations_halving_Line_10_2$S5, paired = T)
# 
# write.table(generation_hobs_Line_10_2, "Line_10_2_hobs.txt", col.names = T, row.names = T, quote = F)
# write.table(expectations_halving_Line_10_2, "Line_10_2_hexp.txt", col.names = T, row.names = F, quote = F)
# 
# # Line 10_5
# generation_hobs_Line_10_5 <- data.frame(F_Hobs = Line_10_5_F_div$Hobs,
#                                         S1_Hobs = Line_10_5_S1_div$Hobs,
#                                         S2_Hobs = Line_10_5_S2_div$Hobs,
#                                         S3_Hobs = Line_10_5_S3_div$Hobs,
#                                         S4_Hobs = Line_10_5_S4_div$Hobs)
# 
# colnames(generation_hobs_Line_10_5) <- c("F", "S1", "S2", "S3", "S4")
# # generation_hobs <- rownames_to_column(generation_hobs_Line_10_5)
# melted_generation_hobs_Line_10_5 <- melt(generation_hobs_Line_10_5)
# 
# generation_hobs_Line_10_5_summary <- data.frame(FS = mean(generation_hobs_Line_10_5$F, na.rm = T), 
#                                                 S1 = mean(generation_hobs_Line_10_5$S1, na.rm = T),
#                                                 S2 = mean(generation_hobs_Line_10_5$S2, na.rm = T),
#                                                 S3 = mean(generation_hobs_Line_10_5$S3, na.rm = T),
#                                                 S4 = mean(generation_hobs_Line_10_5$S4, na.rm = T),
#                                                 H = "Observed")
# 
# melted_generation_hobs_Line_10_5_summary <- melt(generation_hobs_Line_10_5_summary)
# 
# ggplot(melted_generation_hobs_Line_10_5, aes(x = value, fill = variable)) +
#   geom_density(alpha = 0.3) +
#   theme_Publication()
# 
# ggplot(melted_generation_hobs_Line_10_5_summary, aes(x = variable, y = value)) +
#   geom_point() +
#   scale_y_continuous(breaks = c(0.1, 0.2, 0.3, 0.4, 0.5 ,0.6, 0.7, 0.8, 0.9, 1)) +
#   ylim(c(0,1))
# 
# expectations_halving_Line_10_5 <- melted_generation_hobs_Line_10_5 %>% 
#   filter(variable == "F") %>% 
#   mutate(S1 = (value / 2), S2 = (value / 4), S3 = (value / 8), S4 = (value/ 16)) %>% 
#   rename(F = value) %>% 
#   select(F, S1, S2, S3, S4)
# 
# expectations_halving_Line_10_5_summary <- data.frame(FS = mean(expectations_halving_Line_10_5$F, na.rm = T), 
#                                                      S1 = mean(expectations_halving_Line_10_5$S1, na.rm = T),
#                                                      S2 = mean(expectations_halving_Line_10_5$S2, na.rm = T),
#                                                      S3 = mean(expectations_halving_Line_10_5$S3, na.rm = T),
#                                                      S4 = mean(expectations_halving_Line_10_5$S4, na.rm = T),
#                                                      H = "Expected")
# expectations_halving_Line_10_5_summary_melted <- melt(expectations_halving_Line_10_5_summary)
# 
# obs_exp_10_5 <- rbind(expectations_halving_Line_10_5_summary_melted, melted_generation_hobs_Line_10_5_summary)
# 
# obs_exp_10_5_plot <- ggplot(obs_exp_10_5, aes(x = variable, y = value, colour = H)) +
#   geom_point(size = 3) +
#   scale_y_continuous(breaks = c(0.1, 0.2, 0.3, 0.4, 0.5 ,0.6, 0.7, 0.8, 0.9, 1)) +
#   ylim(c(0,1)) +
#   scale_colour_nejm() +
#   ylab("Heterozygosity") +
#   xlab("Generation") +
#   theme_Publication() +
#   theme(legend.position = "none") +
#   ggtitle("Line 10_5")
# 
# wilcox.test(generation_hobs_Line_10_5$S1, expectations_halving_Line_10_5$S1, paired = T)
# wilcox.test(generation_hobs_Line_10_5$S2, expectations_halving_Line_10_5$S2, paired = T)
# wilcox.test(generation_hobs_Line_10_5$S3, expectations_halving_Line_10_5$S3, paired = T)
# wilcox.test(generation_hobs_Line_10_5$S4, expectations_halving_Line_10_5$S4, paired = T)
# 
# write.table(generation_hobs_Line_10_5, "Line_10_5_hobs.txt", col.names = T, row.names = T, quote = F)
# write.table(expectations_halving_Line_10_5, "Line_10_5_hexp.txt", col.names = T, row.names = F, quote = F)
# 
# # Line 13_3
# generation_hobs_Line_13_3 <- data.frame(F_Hobs = Line_13_3_F_div$Hobs,
#                                         S1_Hobs = Line_13_3_S1_div$Hobs,
#                                         S2_Hobs = Line_13_3_S2_div$Hobs,
#                                         S3_Hobs = Line_13_3_S3_div$Hobs,
#                                         S4_Hobs = Line_13_3_S4_div$Hobs)
# 
# colnames(generation_hobs_Line_13_3) <- c("F", "S1", "S2", "S3", "S4")
# # generation_hobs <- rownames_to_column(generation_hobs_Line_13_3)
# melted_generation_hobs_Line_13_3 <- melt(generation_hobs_Line_13_3)
# 
# generation_hobs_Line_13_3_summary <- data.frame(FS = mean(generation_hobs_Line_13_3$F, na.rm = T), 
#                                                 S1 = mean(generation_hobs_Line_13_3$S1, na.rm = T),
#                                                 S2 = mean(generation_hobs_Line_13_3$S2, na.rm = T),
#                                                 S3 = mean(generation_hobs_Line_13_3$S3, na.rm = T),
#                                                 S4 = mean(generation_hobs_Line_13_3$S4, na.rm = T),
#                                                 H = "Observed")
# 
# melted_generation_hobs_Line_13_3_summary <- melt(generation_hobs_Line_13_3_summary)
# 
# ggplot(melted_generation_hobs_Line_13_3, aes(x = value, fill = variable)) +
#   geom_density(alpha = 0.3) +
#   theme_Publication()
# 
# ggplot(melted_generation_hobs_Line_13_3_summary, aes(x = variable, y = value)) +
#   geom_point() +
#   scale_y_continuous(breaks = c(0.1, 0.2, 0.3, 0.4, 0.5 ,0.6, 0.7, 0.8, 0.9, 1)) +
#   ylim(c(0,1))
# 
# expectations_halving_Line_13_3 <- melted_generation_hobs_Line_13_3 %>% 
#   filter(variable == "F") %>% 
#   mutate(S1 = (value / 2), S2 = (value / 4), S3 = (value / 8), S4 = (value/ 16)) %>% 
#   rename(F = value) %>% 
#   select(F, S1, S2, S3, S4)
# 
# expectations_halving_Line_13_3_summary <- data.frame(FS = mean(expectations_halving_Line_13_3$F, na.rm = T), 
#                                                      S1 = mean(expectations_halving_Line_13_3$S1, na.rm = T),
#                                                      S2 = mean(expectations_halving_Line_13_3$S2, na.rm = T),
#                                                      S3 = mean(expectations_halving_Line_13_3$S3, na.rm = T),
#                                                      S4 = mean(expectations_halving_Line_13_3$S4, na.rm = T),
#                                                      H = "Expected")
# expectations_halving_Line_13_3_summary_melted <- melt(expectations_halving_Line_13_3_summary)
# 
# obs_exp_13_3 <- rbind(expectations_halving_Line_13_3_summary_melted, melted_generation_hobs_Line_13_3_summary)
# 
# obs_exp_13_3_plot <- ggplot(obs_exp_13_3, aes(x = variable, y = value, colour = H)) +
#   geom_point(size = 3) +
#   scale_y_continuous(breaks = c(0.1, 0.2, 0.3, 0.4, 0.5 ,0.6, 0.7, 0.8, 0.9, 1)) +
#   ylim(c(0,1)) +
#   scale_colour_nejm() +
#   ylab("Heterozygosity") +
#   xlab("Generation") +
#   theme_Publication() +
#   theme(legend.position = "none") +
#   ggtitle("Line 13_3")
# 
# wilcox.test(generation_hobs_Line_13_3$S1, expectations_halving_Line_13_3$S1, paired = T)
# wilcox.test(generation_hobs_Line_13_3$S2, expectations_halving_Line_13_3$S2, paired = T)
# wilcox.test(generation_hobs_Line_13_3$S3, expectations_halving_Line_13_3$S3, paired = T)
# wilcox.test(generation_hobs_Line_13_3$S4, expectations_halving_Line_13_3$S4, paired = T)
# 
# write.table(generation_hobs_Line_13_3, "Line_13_3_hobs.txt", col.names = T, row.names = T, quote = F)
# write.table(expectations_halving_Line_13_3, "Line_13_3_hexp.txt", col.names = T, row.names = F, quote = F)
# 
# # Line 13_4
# generation_hobs_Line_13_4 <- data.frame(F_Hobs = Line_13_4_F_div$Hobs,
#                                         S1_Hobs = Line_13_4_S1_div$Hobs,
#                                         S2_Hobs = Line_13_4_S2_div$Hobs,
#                                         S3_Hobs = Line_13_4_S3_div$Hobs,
#                                         S4_Hobs = Line_13_4_S4_div$Hobs)
# 
# colnames(generation_hobs_Line_13_4) <- c("F", "S1", "S2", "S3", "S4")
# # generation_hobs <- rownames_to_column(generation_hobs_Line_13_4)
# melted_generation_hobs_Line_13_4 <- melt(generation_hobs_Line_13_4)
# 
# generation_hobs_Line_13_4_summary <- data.frame(FS = mean(generation_hobs_Line_13_4$F, na.rm = T), 
#                                                 S1 = mean(generation_hobs_Line_13_4$S1, na.rm = T),
#                                                 S2 = mean(generation_hobs_Line_13_4$S2, na.rm = T),
#                                                 S3 = mean(generation_hobs_Line_13_4$S3, na.rm = T),
#                                                 S4 = mean(generation_hobs_Line_13_4$S4, na.rm = T),
#                                                 H = "Observed")
# 
# melted_generation_hobs_Line_13_4_summary <- melt(generation_hobs_Line_13_4_summary)
# 
# ggplot(melted_generation_hobs_Line_13_4, aes(x = value, fill = variable)) +
#   geom_density(alpha = 0.3) +
#   theme_Publication()
# 
# ggplot(melted_generation_hobs_Line_13_4_summary, aes(x = variable, y = value)) +
#   geom_point() +
#   scale_y_continuous(breaks = c(0.1, 0.2, 0.3, 0.4, 0.5 ,0.6, 0.7, 0.8, 0.9, 1)) +
#   ylim(c(0,1))
# 
# expectations_halving_Line_13_4 <- melted_generation_hobs_Line_13_4 %>% 
#   filter(variable == "F") %>% 
#   mutate(S1 = (value / 2), S2 = (value / 4), S3 = (value / 8), S4 = (value/ 16)) %>% 
#   rename(F = value) %>% 
#   select(F, S1, S2, S3, S4)
# 
# expectations_halving_Line_13_4_summary <- data.frame(FS = mean(expectations_halving_Line_13_4$F, na.rm = T), 
#                                                      S1 = mean(expectations_halving_Line_13_4$S1, na.rm = T),
#                                                      S2 = mean(expectations_halving_Line_13_4$S2, na.rm = T),
#                                                      S3 = mean(expectations_halving_Line_13_4$S3, na.rm = T),
#                                                      S4 = mean(expectations_halving_Line_13_4$S4, na.rm = T),
#                                                      H = "Expected")
# expectations_halving_Line_13_4_summary_melted <- melt(expectations_halving_Line_13_4_summary)
# 
# obs_exp_13_4 <- rbind(expectations_halving_Line_13_4_summary_melted, melted_generation_hobs_Line_13_4_summary)
# 
# obs_exp_13_4_plot <- ggplot(obs_exp_13_4, aes(x = variable, y = value, colour = H)) +
#   geom_point(size = 3) +
#   scale_y_continuous(breaks = c(0.1, 0.2, 0.3, 0.4, 0.5 ,0.6, 0.7, 0.8, 0.9, 1)) +
#   ylim(c(0,1)) +
#   scale_colour_nejm() +
#   ylab("Heterozygosity") +
#   xlab("Generation") +
#   theme_Publication() +
#   theme(legend.position = "none") +
#   ggtitle("Line 13_4")
# 
# wilcox.test(generation_hobs_Line_13_4$S1, expectations_halving_Line_13_4$S1, paired = T)
# wilcox.test(generation_hobs_Line_13_4$S2, expectations_halving_Line_13_4$S2, paired = T)
# wilcox.test(generation_hobs_Line_13_4$S3, expectations_halving_Line_13_4$S3, paired = T)
# wilcox.test(generation_hobs_Line_13_4$S4, expectations_halving_Line_13_4$S4, paired = T)
# 
# write.table(generation_hobs_Line_13_4, "Line_13_4_hobs.txt", col.names = T, row.names = T, quote = F)
# write.table(expectations_halving_Line_13_4, "Line_13_4_hexp.txt", col.names = T, row.names = F, quote = F)
# 
# # Line 16_1
# generation_hobs_Line_16_1 <- data.frame(F_Hobs = Line_16_1_F_div$Hobs,
#                                         S1_Hobs = Line_16_1_S1_div$Hobs,
#                                         S2_Hobs = Line_16_1_S2_div$Hobs,
#                                         S3_Hobs = Line_16_1_S3_div$Hobs,
#                                         S4_Hobs = Line_16_1_S4_div$Hobs)
# 
# colnames(generation_hobs_Line_16_1) <- c("F", "S1", "S2", "S3", "S4")
# # generation_hobs <- rownames_to_column(generation_hobs_Line_16_1)
# melted_generation_hobs_Line_16_1 <- melt(generation_hobs_Line_16_1)
# 
# generation_hobs_Line_16_1_summary <- data.frame(FS = mean(generation_hobs_Line_16_1$F, na.rm = T), 
#                                                 S1 = mean(generation_hobs_Line_16_1$S1, na.rm = T),
#                                                 S2 = mean(generation_hobs_Line_16_1$S2, na.rm = T),
#                                                 S3 = mean(generation_hobs_Line_16_1$S3, na.rm = T),
#                                                 S4 = mean(generation_hobs_Line_16_1$S4, na.rm = T),
#                                                 H = "Observed")
# 
# melted_generation_hobs_Line_16_1_summary <- melt(generation_hobs_Line_16_1_summary)
# 
# ggplot(melted_generation_hobs_Line_16_1, aes(x = value, fill = variable)) +
#   geom_density(alpha = 0.3) +
#   theme_Publication()
# 
# ggplot(melted_generation_hobs_Line_16_1_summary, aes(x = variable, y = value)) +
#   geom_point() +
#   scale_y_continuous(breaks = c(0.1, 0.2, 0.3, 0.4, 0.5 ,0.6, 0.7, 0.8, 0.9, 1)) +
#   ylim(c(0,1))
# 
# expectations_halving_Line_16_1 <- melted_generation_hobs_Line_16_1 %>% 
#   filter(variable == "F") %>% 
#   mutate(S1 = (value / 2), S2 = (value / 4), S3 = (value / 8), S4 = (value/ 16)) %>% 
#   rename(F = value) %>% 
#   select(F, S1, S2, S3, S4)
# 
# expectations_halving_Line_16_1_summary <- data.frame(FS = mean(expectations_halving_Line_16_1$F, na.rm = T), 
#                                                      S1 = mean(expectations_halving_Line_16_1$S1, na.rm = T),
#                                                      S2 = mean(expectations_halving_Line_16_1$S2, na.rm = T),
#                                                      S3 = mean(expectations_halving_Line_16_1$S3, na.rm = T),
#                                                      S4 = mean(expectations_halving_Line_16_1$S4, na.rm = T),
#                                                      H = "Expected")
# expectations_halving_Line_16_1_summary_melted <- melt(expectations_halving_Line_16_1_summary)
# 
# obs_exp_16_1 <- rbind(expectations_halving_Line_16_1_summary_melted, melted_generation_hobs_Line_16_1_summary)
# 
# obs_exp_16_1_plot <- ggplot(obs_exp_16_1, aes(x = variable, y = value, colour = H)) +
#   geom_point(size = 3) +
#   scale_y_continuous(breaks = c(0.1, 0.2, 0.3, 0.4, 0.5 ,0.6, 0.7, 0.8, 0.9, 1)) +
#   ylim(c(0,1)) +
#   scale_colour_nejm() +
#   ylab("Heterozygosity") +
#   xlab("Generation") +
#   theme_Publication() +
#   theme(legend.position = "none") +
#   ggtitle("Line 16_1")
# 
# wilcox.test(generation_hobs_Line_16_1$S1, expectations_halving_Line_16_1$S1, paired = T)
# wilcox.test(generation_hobs_Line_16_1$S2, expectations_halving_Line_16_1$S2, paired = T)
# wilcox.test(generation_hobs_Line_16_1$S3, expectations_halving_Line_16_1$S3, paired = T)
# wilcox.test(generation_hobs_Line_16_1$S4, expectations_halving_Line_16_1$S4, paired = T)
# 
# write.table(generation_hobs_Line_16_1, "Line_16_1_hobs.txt", col.names = T, row.names = T, quote = F)
# write.table(expectations_halving_Line_16_1, "Line_16_1_hexp.txt", col.names = T, row.names = F, quote = F)
# 
# # Line 16_5
# generation_hobs_Line_16_5 <- data.frame(F_Hobs = Line_16_5_F_div$Hobs,
#                                         S1_Hobs = Line_16_5_S1_div$Hobs,
#                                         S2_Hobs = Line_16_5_S2_div$Hobs,
#                                         S3_Hobs = Line_16_5_S3_div$Hobs,
#                                         S4_Hobs = Line_16_5_S4_div$Hobs,
#                                         S5_Hobs = Line_16_5_S5_div$Hobs)
# 
# colnames(generation_hobs_Line_16_5) <- c("F", "S1", "S2", "S3", "S4", "S5")
# # generation_hobs <- rownames_to_column(generation_hobs_Line_16_5)
# melted_generation_hobs_Line_16_5 <- melt(generation_hobs_Line_16_5)
# 
# generation_hobs_Line_16_5_summary <- data.frame(FS = mean(generation_hobs_Line_16_5$F, na.rm = T), 
#                                                 S1 = mean(generation_hobs_Line_16_5$S1, na.rm = T),
#                                                 S2 = mean(generation_hobs_Line_16_5$S2, na.rm = T),
#                                                 S3 = mean(generation_hobs_Line_16_5$S3, na.rm = T),
#                                                 S4 = mean(generation_hobs_Line_16_5$S4, na.rm = T),
#                                                 S5 = mean(generation_hobs_Line_16_5$S5, na.rm = T),
#                                                 H = "Observed")
# 
# melted_generation_hobs_Line_16_5_summary <- melt(generation_hobs_Line_16_5_summary)
# 
# ggplot(melted_generation_hobs_Line_16_5, aes(x = value, fill = variable)) +
#   geom_density(alpha = 0.3) +
#   theme_Publication()
# 
# ggplot(melted_generation_hobs_Line_16_5_summary, aes(x = variable, y = value)) +
#   geom_point() +
#   scale_y_continuous(breaks = c(0.1, 0.2, 0.3, 0.4, 0.5 ,0.6, 0.7, 0.8, 0.9, 1)) +
#   ylim(c(0,1))
# 
# expectations_halving_Line_16_5 <- melted_generation_hobs_Line_16_5 %>% 
#   filter(variable == "F") %>% 
#   mutate(S1 = (value / 2), S2 = (value / 4), S3 = (value / 8), S4 = (value/ 16), S5 = (value / 32)) %>% 
#   rename(F = value) %>% 
#   select(F, S1, S2, S3, S4, S5)
# 
# expectations_halving_Line_16_5_summary <- data.frame(FS = mean(expectations_halving_Line_16_5$F, na.rm = T), 
#                                                      S1 = mean(expectations_halving_Line_16_5$S1, na.rm = T),
#                                                      S2 = mean(expectations_halving_Line_16_5$S2, na.rm = T),
#                                                      S3 = mean(expectations_halving_Line_16_5$S3, na.rm = T),
#                                                      S4 = mean(expectations_halving_Line_16_5$S4, na.rm = T),
#                                                      S5 = mean(expectations_halving_Line_16_5$S5, na.rm = T),
#                                                      H = "Expected")
# expectations_halving_Line_16_5_summary_melted <- melt(expectations_halving_Line_16_5_summary)
# 
# obs_exp_16_5 <- rbind(expectations_halving_Line_16_5_summary_melted, melted_generation_hobs_Line_16_5_summary)
# 
# obs_exp_16_5_plot <- ggplot(obs_exp_16_5, aes(x = variable, y = value, colour = H)) +
#   geom_point(size = 3) +
#   scale_y_continuous(breaks = c(0.1, 0.2, 0.3, 0.4, 0.5 ,0.6, 0.7, 0.8, 0.9, 1)) +
#   ylim(c(0,1)) +
#   scale_colour_nejm() +
#   ylab("Heterozygosity") +
#   xlab("Generation") +
#   theme_Publication() +
#   theme(legend.position = "none") +
#   ggtitle("Line 16_5")
# 
# wilcox.test(generation_hobs_Line_16_5$S1, expectations_halving_Line_16_5$S1, paired = T)
# wilcox.test(generation_hobs_Line_16_5$S2, expectations_halving_Line_16_5$S2, paired = T)
# wilcox.test(generation_hobs_Line_16_5$S3, expectations_halving_Line_16_5$S3, paired = T)
# wilcox.test(generation_hobs_Line_16_5$S4, expectations_halving_Line_16_5$S4, paired = T)
# wilcox.test(generation_hobs_Line_16_5$S5, expectations_halving_Line_16_5$S5, paired = T)
# 
# write.table(generation_hobs_Line_16_5, "Line_16_5_hobs.txt", col.names = T, row.names = T, quote = F)
# write.table(expectations_halving_Line_16_5, "Line_16_5_hexp.txt", col.names = T, row.names = F, quote = F)
# 
# # Line 17_2
# generation_hobs_Line_17_2 <- data.frame(F_Hobs = Line_17_2_F_div$Hobs,
#                                         S1_Hobs = Line_17_2_S1_div$Hobs,
#                                         S2_Hobs = Line_17_2_S2_div$Hobs,
#                                         S3_Hobs = Line_17_2_S3_div$Hobs,
#                                         S4_Hobs = Line_17_2_S4_div$Hobs,
#                                         S5_Hobs = Line_17_2_S5_div$Hobs)
# 
# colnames(generation_hobs_Line_17_2) <- c("F", "S1", "S2", "S3", "S4", "S5")
# # generation_hobs <- rownames_to_column(generation_hobs_Line_17_2)
# melted_generation_hobs_Line_17_2 <- melt(generation_hobs_Line_17_2)
# 
# generation_hobs_Line_17_2_summary <- data.frame(FS = mean(generation_hobs_Line_17_2$F, na.rm = T), 
#                                                 S1 = mean(generation_hobs_Line_17_2$S1, na.rm = T),
#                                                 S2 = mean(generation_hobs_Line_17_2$S2, na.rm = T),
#                                                 S3 = mean(generation_hobs_Line_17_2$S3, na.rm = T),
#                                                 S4 = mean(generation_hobs_Line_17_2$S4, na.rm = T),
#                                                 S5 = mean(generation_hobs_Line_17_2$S5, na.rm = T),
#                                                 H = "Observed")
# 
# melted_generation_hobs_Line_17_2_summary <- melt(generation_hobs_Line_17_2_summary)
# 
# ggplot(melted_generation_hobs_Line_17_2, aes(x = value, fill = variable)) +
#   geom_density(alpha = 0.3) +
#   theme_Publication()
# 
# ggplot(melted_generation_hobs_Line_17_2_summary, aes(x = variable, y = value)) +
#   geom_point() +
#   scale_y_continuous(breaks = c(0.1, 0.2, 0.3, 0.4, 0.5 ,0.6, 0.7, 0.8, 0.9, 1)) +
#   ylim(c(0,1))
# 
# expectations_halving_Line_17_2 <- melted_generation_hobs_Line_17_2 %>% 
#   filter(variable == "F") %>% 
#   mutate(S1 = (value / 2), S2 = (value / 4), S3 = (value / 8), S4 = (value/ 16), S5 = (value / 32)) %>% 
#   rename(F = value) %>% 
#   select(F, S1, S2, S3, S4, S5)
# 
# expectations_halving_Line_17_2_summary <- data.frame(FS = mean(expectations_halving_Line_17_2$F, na.rm = T), 
#                                                      S1 = mean(expectations_halving_Line_17_2$S1, na.rm = T),
#                                                      S2 = mean(expectations_halving_Line_17_2$S2, na.rm = T),
#                                                      S3 = mean(expectations_halving_Line_17_2$S3, na.rm = T),
#                                                      S4 = mean(expectations_halving_Line_17_2$S4, na.rm = T),
#                                                      S5 = mean(expectations_halving_Line_17_2$S5, na.rm = T),
#                                                      H = "Expected")
# expectations_halving_Line_17_2_summary_melted <- melt(expectations_halving_Line_17_2_summary)
# 
# obs_exp_17_2 <- rbind(expectations_halving_Line_17_2_summary_melted, melted_generation_hobs_Line_17_2_summary)
# 
# obs_exp_17_2_plot <- ggplot(obs_exp_17_2, aes(x = variable, y = value, colour = H)) +
#   geom_point(size = 3) +
#   scale_y_continuous(breaks = c(0.1, 0.2, 0.3, 0.4, 0.5 ,0.6, 0.7, 0.8, 0.9, 1)) +
#   ylim(c(0,1)) +
#   scale_colour_nejm() +
#   ylab("Heterozygosity") +
#   xlab("Generation") +
#   theme_Publication() +
#   theme(legend.position = "none") +
#   ggtitle("Line 17_2")
# 
# wilcox.test(generation_hobs_Line_17_2$S1, expectations_halving_Line_17_2$S1, paired = T)
# wilcox.test(generation_hobs_Line_17_2$S2, expectations_halving_Line_17_2$S2, paired = T)
# wilcox.test(generation_hobs_Line_17_2$S3, expectations_halving_Line_17_2$S3, paired = T)
# wilcox.test(generation_hobs_Line_17_2$S4, expectations_halving_Line_17_2$S4, paired = T)
# wilcox.test(generation_hobs_Line_17_2$S5, expectations_halving_Line_17_2$S5, paired = T)
# 
# write.table(generation_hobs_Line_17_2, "Line_17_2_hobs.txt", col.names = T, row.names = T, quote = F)
# write.table(expectations_halving_Line_17_2, "Line_17_2_hexp.txt", col.names = T, row.names = F, quote = F)
# 
# # Line 17_5
# generation_hobs_Line_17_5 <- data.frame(F_Hobs = Line_17_5_F_div$Hobs,
#                                         S1_Hobs = Line_17_5_S1_div$Hobs,
#                                         S2_Hobs = Line_17_5_S2_div$Hobs,
#                                         S3_Hobs = Line_17_5_S3_div$Hobs,
#                                         S4_Hobs = Line_17_5_S4_div$Hobs,
#                                         S5_Hobs = Line_17_5_S5_div$Hobs)
# 
# colnames(generation_hobs_Line_17_5) <- c("F", "S1", "S2", "S3", "S4", "S5")
# # generation_hobs <- rownames_to_column(generation_hobs_Line_17_5)
# melted_generation_hobs_Line_17_5 <- melt(generation_hobs_Line_17_5)
# 
# generation_hobs_Line_17_5_summary <- data.frame(FS = mean(generation_hobs_Line_17_5$F, na.rm = T), 
#                                                 S1 = mean(generation_hobs_Line_17_5$S1, na.rm = T),
#                                                 S2 = mean(generation_hobs_Line_17_5$S2, na.rm = T),
#                                                 S3 = mean(generation_hobs_Line_17_5$S3, na.rm = T),
#                                                 S4 = mean(generation_hobs_Line_17_5$S4, na.rm = T),
#                                                 S5 = mean(generation_hobs_Line_17_5$S5, na.rm = T),
#                                                 H = "Observed")
# 
# melted_generation_hobs_Line_17_5_summary <- melt(generation_hobs_Line_17_5_summary)
# 
# ggplot(melted_generation_hobs_Line_17_5, aes(x = value, fill = variable)) +
#   geom_density(alpha = 0.3) +
#   theme_Publication()
# 
# ggplot(melted_generation_hobs_Line_17_5_summary, aes(x = variable, y = value)) +
#   geom_point() +
#   scale_y_continuous(breaks = c(0.1, 0.2, 0.3, 0.4, 0.5 ,0.6, 0.7, 0.8, 0.9, 1)) +
#   ylim(c(0,1))
# 
# expectations_halving_Line_17_5 <- melted_generation_hobs_Line_17_5 %>% 
#   filter(variable == "F") %>% 
#   mutate(S1 = (value / 2), S2 = (value / 4), S3 = (value / 8), S4 = (value/ 16), S5 = (value / 32)) %>% 
#   rename(F = value) %>% 
#   select(F, S1, S2, S3, S4, S5)
# 
# expectations_halving_Line_17_5_summary <- data.frame(FS = mean(expectations_halving_Line_17_5$F, na.rm = T), 
#                                                      S1 = mean(expectations_halving_Line_17_5$S1, na.rm = T),
#                                                      S2 = mean(expectations_halving_Line_17_5$S2, na.rm = T),
#                                                      S3 = mean(expectations_halving_Line_17_5$S3, na.rm = T),
#                                                      S4 = mean(expectations_halving_Line_17_5$S4, na.rm = T),
#                                                      S5 = mean(expectations_halving_Line_17_5$S5, na.rm = T),
#                                                      H = "Expected")
# expectations_halving_Line_17_5_summary_melted <- melt(expectations_halving_Line_17_5_summary)
# 
# obs_exp_17_5 <- rbind(expectations_halving_Line_17_5_summary_melted, melted_generation_hobs_Line_17_5_summary)
# 
# obs_exp_17_5_plot <- ggplot(obs_exp_17_5, aes(x = variable, y = value, colour = H)) +
#   geom_point(size = 3) +
#   scale_y_continuous(breaks = c(0.1, 0.2, 0.3, 0.4, 0.5 ,0.6, 0.7, 0.8, 0.9, 1)) +
#   ylim(c(0,1)) +
#   scale_colour_nejm() +
#   ylab("Heterozygosity") +
#   xlab("Generation") +
#   theme_Publication() +
#   theme(legend.position = "none") +
#   ggtitle("Line 17_5")
# 
# wilcox.test(generation_hobs_Line_17_5$S1, expectations_halving_Line_17_5$S1, paired = T)
# wilcox.test(generation_hobs_Line_17_5$S2, expectations_halving_Line_17_5$S2, paired = T)
# wilcox.test(generation_hobs_Line_17_5$S3, expectations_halving_Line_17_5$S3, paired = T)
# wilcox.test(generation_hobs_Line_17_5$S4, expectations_halving_Line_17_5$S4, paired = T)
# wilcox.test(generation_hobs_Line_17_5$S5, expectations_halving_Line_17_5$S5, paired = T)
# 
# write.table(generation_hobs_Line_17_5, "Line_17_5_hobs.txt", col.names = T, row.names = T, quote = F)
# write.table(expectations_halving_Line_17_5, "Line_17_5_hexp.txt", col.names = T, row.names = F, quote = F)
# 
# # Line 19_2
# generation_hobs_Line_19_2 <- data.frame(F_Hobs = Line_19_2_F_div$Hobs,
#                                         S1_Hobs = Line_19_2_S1_div$Hobs,
#                                         S2_Hobs = Line_19_2_S2_div$Hobs,
#                                         S3_Hobs = Line_19_2_S3_div$Hobs,
#                                         S4_Hobs = Line_19_2_S4_div$Hobs)
# 
# colnames(generation_hobs_Line_19_2) <- c("F", "S1", "S2", "S3", "S4")
# # generation_hobs <- rownames_to_column(generation_hobs_Line_19_2)
# melted_generation_hobs_Line_19_2 <- melt(generation_hobs_Line_19_2)
# 
# generation_hobs_Line_19_2_summary <- data.frame(FS = mean(generation_hobs_Line_19_2$F, na.rm = T), 
#                                                 S1 = mean(generation_hobs_Line_19_2$S1, na.rm = T),
#                                                 S2 = mean(generation_hobs_Line_19_2$S2, na.rm = T),
#                                                 S3 = mean(generation_hobs_Line_19_2$S3, na.rm = T),
#                                                 S4 = mean(generation_hobs_Line_19_2$S4, na.rm = T),
#                                                 H = "Observed")
# 
# melted_generation_hobs_Line_19_2_summary <- melt(generation_hobs_Line_19_2_summary)
# 
# ggplot(melted_generation_hobs_Line_19_2, aes(x = value, fill = variable)) +
#   geom_density(alpha = 0.3) +
#   theme_Publication()
# 
# ggplot(melted_generation_hobs_Line_19_2_summary, aes(x = variable, y = value)) +
#   geom_point() +
#   scale_y_continuous(breaks = c(0.1, 0.2, 0.3, 0.4, 0.5 ,0.6, 0.7, 0.8, 0.9, 1)) +
#   ylim(c(0,1))
# 
# expectations_halving_Line_19_2 <- melted_generation_hobs_Line_19_2 %>% 
#   filter(variable == "F") %>% 
#   mutate(S1 = (value / 2), S2 = (value / 4), S3 = (value / 8), S4 = (value/ 16)) %>% 
#   rename(F = value) %>% 
#   select(F, S1, S2, S3, S4)
# 
# expectations_halving_Line_19_2_summary <- data.frame(FS = mean(expectations_halving_Line_19_2$F, na.rm = T), 
#                                                      S1 = mean(expectations_halving_Line_19_2$S1, na.rm = T),
#                                                      S2 = mean(expectations_halving_Line_19_2$S2, na.rm = T),
#                                                      S3 = mean(expectations_halving_Line_19_2$S3, na.rm = T),
#                                                      S4 = mean(expectations_halving_Line_19_2$S4, na.rm = T),
#                                                      H = "Expected")
# expectations_halving_Line_19_2_summary_melted <- melt(expectations_halving_Line_19_2_summary)
# 
# obs_exp_19_2 <- rbind(expectations_halving_Line_19_2_summary_melted, melted_generation_hobs_Line_19_2_summary)
# 
# obs_exp_19_2_plot <- ggplot(obs_exp_19_2, aes(x = variable, y = value, colour = H)) +
#   geom_point(size = 3) +
#   scale_y_continuous(breaks = c(0.1, 0.2, 0.3, 0.4, 0.5 ,0.6, 0.7, 0.8, 0.9, 1)) +
#   ylim(c(0,1)) +
#   scale_colour_nejm() +
#   ylab("Heterozygosity") +
#   xlab("Generation") +
#   theme_Publication() +
#   theme(legend.position = "none") +
#   ggtitle("Line 19_2")
# 
# wilcox.test(generation_hobs_Line_19_2$S1, expectations_halving_Line_19_2$S1, paired = T)
# wilcox.test(generation_hobs_Line_19_2$S2, expectations_halving_Line_19_2$S2, paired = T)
# wilcox.test(generation_hobs_Line_19_2$S3, expectations_halving_Line_19_2$S3, paired = T)
# wilcox.test(generation_hobs_Line_19_2$S4, expectations_halving_Line_19_2$S4, paired = T)
# 
# write.table(generation_hobs_Line_19_2, "Line_19_2_hobs.txt", col.names = T, row.names = T, quote = F)
# write.table(expectations_halving_Line_19_2, "Line_19_2_hexp.txt", col.names = T, row.names = F, quote = F)
# 
# # Line 19_5
# generation_hobs_Line_19_5 <- data.frame(F_Hobs = Line_19_5_F_div$Hobs,
#                                         S1_Hobs = Line_19_5_S1_div$Hobs,
#                                         S2_Hobs = Line_19_5_S2_div$Hobs,
#                                         S3_Hobs = Line_19_5_S3_div$Hobs,
#                                         S4_Hobs = Line_19_5_S4_div$Hobs)
# 
# colnames(generation_hobs_Line_19_5) <- c("F", "S1", "S2", "S3", "S4")
# # generation_hobs <- rownames_to_column(generation_hobs_Line_19_5)
# melted_generation_hobs_Line_19_5 <- melt(generation_hobs_Line_19_5)
# 
# generation_hobs_Line_19_5_summary <- data.frame(FS = mean(generation_hobs_Line_19_5$F, na.rm = T), 
#                                                 S1 = mean(generation_hobs_Line_19_5$S1, na.rm = T),
#                                                 S2 = mean(generation_hobs_Line_19_5$S2, na.rm = T),
#                                                 S3 = mean(generation_hobs_Line_19_5$S3, na.rm = T),
#                                                 S4 = mean(generation_hobs_Line_19_5$S4, na.rm = T),
#                                                 H = "Observed")
# 
# melted_generation_hobs_Line_19_5_summary <- melt(generation_hobs_Line_19_5_summary)
# 
# ggplot(melted_generation_hobs_Line_19_5, aes(x = value, fill = variable)) +
#   geom_density(alpha = 0.3) +
#   theme_Publication()
# 
# ggplot(melted_generation_hobs_Line_19_5_summary, aes(x = variable, y = value)) +
#   geom_point() +
#   scale_y_continuous(breaks = c(0.1, 0.2, 0.3, 0.4, 0.5 ,0.6, 0.7, 0.8, 0.9, 1)) +
#   ylim(c(0,1))
# 
# expectations_halving_Line_19_5 <- melted_generation_hobs_Line_19_5 %>% 
#   filter(variable == "F") %>% 
#   mutate(S1 = (value / 2), S2 = (value / 4), S3 = (value / 8), S4 = (value/ 16)) %>% 
#   rename(F = value) %>% 
#   select(F, S1, S2, S3, S4)
# 
# expectations_halving_Line_19_5_summary <- data.frame(FS = mean(expectations_halving_Line_19_5$F, na.rm = T), 
#                                                      S1 = mean(expectations_halving_Line_19_5$S1, na.rm = T),
#                                                      S2 = mean(expectations_halving_Line_19_5$S2, na.rm = T),
#                                                      S3 = mean(expectations_halving_Line_19_5$S3, na.rm = T),
#                                                      S4 = mean(expectations_halving_Line_19_5$S4, na.rm = T),
#                                                      H = "Expected")
# expectations_halving_Line_19_5_summary_melted <- melt(expectations_halving_Line_19_5_summary)
# 
# obs_exp_19_5 <- rbind(expectations_halving_Line_19_5_summary_melted, melted_generation_hobs_Line_19_5_summary)
# 
# obs_exp_19_5_plot <- ggplot(obs_exp_19_5, aes(x = variable, y = value, colour = H)) +
#   geom_point(size = 3) +
#   scale_y_continuous(breaks = c(0.1, 0.2, 0.3, 0.4, 0.5 ,0.6, 0.7, 0.8, 0.9, 1)) +
#   ylim(c(0,1)) +
#   scale_colour_nejm() +
#   ylab("Heterozygosity") +
#   xlab("Generation") +
#   theme_Publication() +
#   theme(legend.position = "none") +
#   ggtitle("Line 19_5")
# 
# wilcox.test(generation_hobs_Line_19_5$S1, expectations_halving_Line_19_5$S1, paired = T)
# wilcox.test(generation_hobs_Line_19_5$S2, expectations_halving_Line_19_5$S2, paired = T)
# wilcox.test(generation_hobs_Line_19_5$S3, expectations_halving_Line_19_5$S3, paired = T)
# wilcox.test(generation_hobs_Line_19_5$S4, expectations_halving_Line_19_5$S4, paired = T)
# 
# write.table(generation_hobs_Line_19_5, "Line_19_5_hobs.txt", col.names = T, row.names = T, quote = F)
# write.table(expectations_halving_Line_19_5, "Line_19_5_hexp.txt", col.names = T, row.names = F, quote = F)
# 
# # Line 20_1
# generation_hobs_Line_20_1 <- data.frame(F_Hobs = Line_20_1_F_div$Hobs,
#                                         S1_Hobs = Line_20_1_S1_div$Hobs,
#                                         S2_Hobs = Line_20_1_S2_div$Hobs,
#                                         S3_Hobs = Line_20_1_S3_div$Hobs,
#                                         S4_Hobs = Line_20_1_S4_div$Hobs)
# 
# colnames(generation_hobs_Line_20_1) <- c("F", "S1", "S2", "S3", "S4")
# # generation_hobs <- rownames_to_column(generation_hobs_Line_20_1)
# melted_generation_hobs_Line_20_1 <- melt(generation_hobs_Line_20_1)
# 
# generation_hobs_Line_20_1_summary <- data.frame(FS = mean(generation_hobs_Line_20_1$F, na.rm = T), 
#                                                 S1 = mean(generation_hobs_Line_20_1$S1, na.rm = T),
#                                                 S2 = mean(generation_hobs_Line_20_1$S2, na.rm = T),
#                                                 S3 = mean(generation_hobs_Line_20_1$S3, na.rm = T),
#                                                 S4 = mean(generation_hobs_Line_20_1$S4, na.rm = T),
#                                                 H = "Observed")
# 
# melted_generation_hobs_Line_20_1_summary <- melt(generation_hobs_Line_20_1_summary)
# 
# ggplot(melted_generation_hobs_Line_20_1, aes(x = value, fill = variable)) +
#   geom_density(alpha = 0.3) +
#   theme_Publication()
# 
# ggplot(melted_generation_hobs_Line_20_1_summary, aes(x = variable, y = value)) +
#   geom_point() +
#   scale_y_continuous(breaks = c(0.1, 0.2, 0.3, 0.4, 0.5 ,0.6, 0.7, 0.8, 0.9, 1)) +
#   ylim(c(0,1))
# 
# expectations_halving_Line_20_1 <- melted_generation_hobs_Line_20_1 %>% 
#   filter(variable == "F") %>% 
#   mutate(S1 = (value / 2), S2 = (value / 4), S3 = (value / 8), S4 = (value/ 16)) %>% 
#   rename(F = value) %>% 
#   select(F, S1, S2, S3, S4)
# 
# expectations_halving_Line_20_1_summary <- data.frame(FS = mean(expectations_halving_Line_20_1$F, na.rm = T), 
#                                                      S1 = mean(expectations_halving_Line_20_1$S1, na.rm = T),
#                                                      S2 = mean(expectations_halving_Line_20_1$S2, na.rm = T),
#                                                      S3 = mean(expectations_halving_Line_20_1$S3, na.rm = T),
#                                                      S4 = mean(expectations_halving_Line_20_1$S4, na.rm = T),
#                                                      H = "Expected")
# expectations_halving_Line_20_1_summary_melted <- melt(expectations_halving_Line_20_1_summary)
# 
# obs_exp_20_1 <- rbind(expectations_halving_Line_20_1_summary_melted, melted_generation_hobs_Line_20_1_summary)
# 
# obs_exp_20_1_plot <- ggplot(obs_exp_20_1, aes(x = variable, y = value, colour = H)) +
#   geom_point(size = 3) +
#   scale_y_continuous(breaks = c(0.1, 0.2, 0.3, 0.4, 0.5 ,0.6, 0.7, 0.8, 0.9, 1)) +
#   ylim(c(0,1)) +
#   scale_colour_nejm() +
#   ylab("Heterozygosity") +
#   xlab("Generation") +
#   theme_Publication() +
#   theme(legend.position = "none") +
#   ggtitle("Line 20_1")
# 
# wilcox.test(generation_hobs_Line_20_1$S1, expectations_halving_Line_20_1$S1, paired = T)
# wilcox.test(generation_hobs_Line_20_1$S2, expectations_halving_Line_20_1$S2, paired = T)
# wilcox.test(generation_hobs_Line_20_1$S3, expectations_halving_Line_20_1$S3, paired = T)
# wilcox.test(generation_hobs_Line_20_1$S4, expectations_halving_Line_20_1$S4, paired = T)
# 
# write.table(generation_hobs_Line_20_1, "Line_20_1_hobs.txt", col.names = T, row.names = T, quote = F)
# write.table(expectations_halving_Line_20_1, "Line_20_1_hexp.txt", col.names = T, row.names = F, quote = F)
# 
# # Line 20_4
# generation_hobs_Line_20_4 <- data.frame(F_Hobs = Line_20_4_F_div$Hobs,
#                                         S1_Hobs = Line_20_4_S1_div$Hobs,
#                                         S2_Hobs = Line_20_4_S2_div$Hobs,
#                                         S3_Hobs = Line_20_4_S3_div$Hobs,
#                                         S4_Hobs = Line_20_4_S4_div$Hobs)
# 
# colnames(generation_hobs_Line_20_4) <- c("F", "S1", "S2", "S3", "S4")
# # generation_hobs <- rownames_to_column(generation_hobs_Line_20_4)
# melted_generation_hobs_Line_20_4 <- melt(generation_hobs_Line_20_4)
# 
# generation_hobs_Line_20_4_summary <- data.frame(FS = mean(generation_hobs_Line_20_4$F, na.rm = T), 
#                                                 S1 = mean(generation_hobs_Line_20_4$S1, na.rm = T),
#                                                 S2 = mean(generation_hobs_Line_20_4$S2, na.rm = T),
#                                                 S3 = mean(generation_hobs_Line_20_4$S3, na.rm = T),
#                                                 S4 = mean(generation_hobs_Line_20_4$S4, na.rm = T),
#                                                 H = "Observed")
# 
# melted_generation_hobs_Line_20_4_summary <- melt(generation_hobs_Line_20_4_summary)
# 
# ggplot(melted_generation_hobs_Line_20_4, aes(x = value, fill = variable)) +
#   geom_density(alpha = 0.3) +
#   theme_Publication()
# 
# ggplot(melted_generation_hobs_Line_20_4_summary, aes(x = variable, y = value)) +
#   geom_point() +
#   scale_y_continuous(breaks = c(0.1, 0.2, 0.3, 0.4, 0.5 ,0.6, 0.7, 0.8, 0.9, 1)) +
#   ylim(c(0,1))
# 
# expectations_halving_Line_20_4 <- melted_generation_hobs_Line_20_4 %>% 
#   filter(variable == "F") %>% 
#   mutate(S1 = (value / 2), S2 = (value / 4), S3 = (value / 8), S4 = (value/ 16)) %>% 
#   rename(F = value) %>% 
#   select(F, S1, S2, S3, S4)
# 
# expectations_halving_Line_20_4_summary <- data.frame(FS = mean(expectations_halving_Line_20_4$F, na.rm = T), 
#                                                      S1 = mean(expectations_halving_Line_20_4$S1, na.rm = T),
#                                                      S2 = mean(expectations_halving_Line_20_4$S2, na.rm = T),
#                                                      S3 = mean(expectations_halving_Line_20_4$S3, na.rm = T),
#                                                      S4 = mean(expectations_halving_Line_20_4$S4, na.rm = T),
#                                                      H = "Expected")
# expectations_halving_Line_20_4_summary_melted <- melt(expectations_halving_Line_20_4_summary)
# 
# obs_exp_20_4 <- rbind(expectations_halving_Line_20_4_summary_melted, melted_generation_hobs_Line_20_4_summary)
# 
# obs_exp_20_4_plot <- ggplot(obs_exp_20_4, aes(x = variable, y = value, colour = H)) +
#   geom_point(size = 3) +
#   scale_y_continuous(breaks = c(0.1, 0.2, 0.3, 0.4, 0.5 ,0.6, 0.7, 0.8, 0.9, 1)) +
#   ylim(c(0,1)) +
#   scale_colour_nejm() +
#   ylab("Heterozygosity") +
#   xlab("Generation") +
#   theme_Publication() +
#   theme(legend.position = "none") +
#   ggtitle("Line 20_4")
# 
# wilcox.test(generation_hobs_Line_20_4$S1, expectations_halving_Line_20_4$S1, paired = T)
# wilcox.test(generation_hobs_Line_20_4$S2, expectations_halving_Line_20_4$S2, paired = T)
# wilcox.test(generation_hobs_Line_20_4$S3, expectations_halving_Line_20_4$S3, paired = T)
# wilcox.test(generation_hobs_Line_20_4$S4, expectations_halving_Line_20_4$S4, paired = T)
# 
# write.table(generation_hobs_Line_20_4, "Line_20_4_hobs.txt", col.names = T, row.names = T, quote = F)
# write.table(expectations_halving_Line_20_4, "Line_20_4_hexp.txt", col.names = T, row.names = F, quote = F)
# 
# # Line 21_2
# generation_hobs_Line_21_2 <- data.frame(F_Hobs = Line_21_2_F_div$Hobs,
#                                         S1_Hobs = Line_21_2_S1_div$Hobs,
#                                         S2_Hobs = Line_21_2_S2_div$Hobs,
#                                         S3_Hobs = Line_21_2_S3_div$Hobs,
#                                         S4_Hobs = Line_21_2_S4_div$Hobs,
#                                         S5_Hobs = Line_21_2_S5_div$Hobs)
# 
# colnames(generation_hobs_Line_21_2) <- c("F", "S1", "S2", "S3", "S4", "S5")
# # generation_hobs <- rownames_to_column(generation_hobs_Line_21_2)
# melted_generation_hobs_Line_21_2 <- melt(generation_hobs_Line_21_2)
# 
# generation_hobs_Line_21_2_summary <- data.frame(FS = mean(generation_hobs_Line_21_2$F, na.rm = T), 
#                                                 S1 = mean(generation_hobs_Line_21_2$S1, na.rm = T),
#                                                 S2 = mean(generation_hobs_Line_21_2$S2, na.rm = T),
#                                                 S3 = mean(generation_hobs_Line_21_2$S3, na.rm = T),
#                                                 S4 = mean(generation_hobs_Line_21_2$S4, na.rm = T),
#                                                 S5 = mean(generation_hobs_Line_21_2$S5, na.rm = T),
#                                                 H = "Observed")
# 
# melted_generation_hobs_Line_21_2_summary <- melt(generation_hobs_Line_21_2_summary)
# 
# ggplot(melted_generation_hobs_Line_21_2, aes(x = value, fill = variable)) +
#   geom_density(alpha = 0.3) +
#   theme_Publication()
# 
# ggplot(melted_generation_hobs_Line_21_2_summary, aes(x = variable, y = value)) +
#   geom_point() +
#   scale_y_continuous(breaks = c(0.1, 0.2, 0.3, 0.4, 0.5 ,0.6, 0.7, 0.8, 0.9, 1)) +
#   ylim(c(0,1))
# 
# expectations_halving_Line_21_2 <- melted_generation_hobs_Line_21_2 %>% 
#   filter(variable == "F") %>% 
#   mutate(S1 = (value / 2), S2 = (value / 4), S3 = (value / 8), S4 = (value/ 16), S5 = (value / 32)) %>% 
#   rename(F = value) %>% 
#   select(F, S1, S2, S3, S4, S5)
# 
# expectations_halving_Line_21_2_summary <- data.frame(FS = mean(expectations_halving_Line_21_2$F, na.rm = T), 
#                                                      S1 = mean(expectations_halving_Line_21_2$S1, na.rm = T),
#                                                      S2 = mean(expectations_halving_Line_21_2$S2, na.rm = T),
#                                                      S3 = mean(expectations_halving_Line_21_2$S3, na.rm = T),
#                                                      S4 = mean(expectations_halving_Line_21_2$S4, na.rm = T),
#                                                      S5 = mean(expectations_halving_Line_21_2$S5, na.rm = T),
#                                                      H = "Expected")
# expectations_halving_Line_21_2_summary_melted <- melt(expectations_halving_Line_21_2_summary)
# 
# obs_exp_21_2 <- rbind(expectations_halving_Line_21_2_summary_melted, melted_generation_hobs_Line_21_2_summary)
# 
# obs_exp_21_2_plot <- ggplot(obs_exp_21_2, aes(x = variable, y = value, colour = H)) +
#   geom_point(size = 3) +
#   scale_y_continuous(breaks = c(0.1, 0.2, 0.3, 0.4, 0.5 ,0.6, 0.7, 0.8, 0.9, 1)) +
#   ylim(c(0,1)) +
#   scale_colour_nejm() +
#   ylab("Heterozygosity") +
#   xlab("Generation") +
#   theme_Publication() +
#   theme(legend.position = "none") +
#   ggtitle("Line 21_2")
# 
# wilcox.test(generation_hobs_Line_21_2$S1, expectations_halving_Line_21_2$S1, paired = T)
# wilcox.test(generation_hobs_Line_21_2$S2, expectations_halving_Line_21_2$S2, paired = T)
# wilcox.test(generation_hobs_Line_21_2$S3, expectations_halving_Line_21_2$S3, paired = T)
# wilcox.test(generation_hobs_Line_21_2$S4, expectations_halving_Line_21_2$S4, paired = T)
# wilcox.test(generation_hobs_Line_21_2$S5, expectations_halving_Line_21_2$S5, paired = T)
# 
# write.table(generation_hobs_Line_21_2, "Line_21_2_hobs.txt", col.names = T, row.names = T, quote = F)
# write.table(expectations_halving_Line_21_2, "Line_21_2_hexp.txt", col.names = T, row.names = F, quote = F)
# 
# # Line 21_6
# generation_hobs_Line_21_6 <- data.frame(F_Hobs = Line_21_6_F_div$Hobs,
#                                         S1_Hobs = Line_21_6_S1_div$Hobs,
#                                         S2_Hobs = Line_21_6_S2_div$Hobs,
#                                         S3_Hobs = Line_21_6_S3_div$Hobs,
#                                         S4_Hobs = Line_21_6_S4_div$Hobs,
#                                         S5_Hobs = Line_21_6_S5_div$Hobs)
# 
# colnames(generation_hobs_Line_21_6) <- c("F", "S1", "S2", "S3", "S4", "S5")
# # generation_hobs <- rownames_to_column(generation_hobs_Line_21_6)
# melted_generation_hobs_Line_21_6 <- melt(generation_hobs_Line_21_6)
# 
# generation_hobs_Line_21_6_summary <- data.frame(FS = mean(generation_hobs_Line_21_6$F, na.rm = T), 
#                                                 S1 = mean(generation_hobs_Line_21_6$S1, na.rm = T),
#                                                 S2 = mean(generation_hobs_Line_21_6$S2, na.rm = T),
#                                                 S3 = mean(generation_hobs_Line_21_6$S3, na.rm = T),
#                                                 S4 = mean(generation_hobs_Line_21_6$S4, na.rm = T),
#                                                 S5 = mean(generation_hobs_Line_21_6$S5, na.rm = T),
#                                                 H = "Observed")
# 
# melted_generation_hobs_Line_21_6_summary <- melt(generation_hobs_Line_21_6_summary)
# 
# ggplot(melted_generation_hobs_Line_21_6, aes(x = value, fill = variable)) +
#   geom_density(alpha = 0.3) +
#   theme_Publication()
# 
# ggplot(melted_generation_hobs_Line_21_6_summary, aes(x = variable, y = value)) +
#   geom_point() +
#   scale_y_continuous(breaks = c(0.1, 0.2, 0.3, 0.4, 0.5 ,0.6, 0.7, 0.8, 0.9, 1)) +
#   ylim(c(0,1))
# 
# expectations_halving_Line_21_6 <- melted_generation_hobs_Line_21_6 %>% 
#   filter(variable == "F") %>% 
#   mutate(S1 = (value / 2), S2 = (value / 4), S3 = (value / 8), S4 = (value/ 16), S5 = (value / 32)) %>% 
#   rename(F = value) %>% 
#   select(F, S1, S2, S3, S4, S5)
# 
# expectations_halving_Line_21_6_summary <- data.frame(FS = mean(expectations_halving_Line_21_6$F, na.rm = T), 
#                                                      S1 = mean(expectations_halving_Line_21_6$S1, na.rm = T),
#                                                      S2 = mean(expectations_halving_Line_21_6$S2, na.rm = T),
#                                                      S3 = mean(expectations_halving_Line_21_6$S3, na.rm = T),
#                                                      S4 = mean(expectations_halving_Line_21_6$S4, na.rm = T),
#                                                      S5 = mean(expectations_halving_Line_21_6$S5, na.rm = T),
#                                                      H = "Expected")
# expectations_halving_Line_21_6_summary_melted <- melt(expectations_halving_Line_21_6_summary)
# 
# obs_exp_21_6 <- rbind(expectations_halving_Line_21_6_summary_melted, melted_generation_hobs_Line_21_6_summary)
# 
# obs_exp_21_6_plot <- ggplot(obs_exp_21_6, aes(x = variable, y = value, colour = H)) +
#   geom_point(size = 3) +
#   scale_y_continuous(breaks = c(0.1, 0.2, 0.3, 0.4, 0.5 ,0.6, 0.7, 0.8, 0.9, 1)) +
#   ylim(c(0,1)) +
#   scale_colour_nejm() +
#   ylab("Heterozygosity") +
#   xlab("Generation") +
#   theme_Publication() +
#   theme(legend.position = "none") +
#   ggtitle("Line 21_6")
# 
# wilcox.test(generation_hobs_Line_21_6$S1, expectations_halving_Line_21_6$S1, paired = T)
# wilcox.test(generation_hobs_Line_21_6$S2, expectations_halving_Line_21_6$S2, paired = T)
# wilcox.test(generation_hobs_Line_21_6$S3, expectations_halving_Line_21_6$S3, paired = T)
# wilcox.test(generation_hobs_Line_21_6$S4, expectations_halving_Line_21_6$S4, paired = T)
# wilcox.test(generation_hobs_Line_21_6$S5, expectations_halving_Line_21_6$S5, paired = T)
# 
# write.table(generation_hobs_Line_21_6, "Line_21_6_hobs.txt", col.names = T, row.names = T, quote = F)
# write.table(expectations_halving_Line_21_6, "Line_21_6_hexp.txt", col.names = T, row.names = F, quote = F)
# 
# # Line 23_2
# generation_hobs_Line_23_2 <- data.frame(F_Hobs = Line_23_2_F_div$Hobs,
#                                         S1_Hobs = Line_23_2_S1_div$Hobs,
#                                         S2_Hobs = Line_23_2_S2_div$Hobs,
#                                         S3_Hobs = Line_23_2_S3_div$Hobs,
#                                         S4_Hobs = Line_23_2_S4_div$Hobs,
#                                         S5_Hobs = Line_23_2_S5_div$Hobs)
# 
# colnames(generation_hobs_Line_23_2) <- c("F", "S1", "S2", "S3", "S4", "S5")
# # generation_hobs <- rownames_to_column(generation_hobs_Line_23_2)
# melted_generation_hobs_Line_23_2 <- melt(generation_hobs_Line_23_2)
# 
# generation_hobs_Line_23_2_summary <- data.frame(FS = mean(generation_hobs_Line_23_2$F, na.rm = T), 
#                                                 S1 = mean(generation_hobs_Line_23_2$S1, na.rm = T),
#                                                 S2 = mean(generation_hobs_Line_23_2$S2, na.rm = T),
#                                                 S3 = mean(generation_hobs_Line_23_2$S3, na.rm = T),
#                                                 S4 = mean(generation_hobs_Line_23_2$S4, na.rm = T),
#                                                 S5 = mean(generation_hobs_Line_23_2$S5, na.rm = T),
#                                                 H = "Observed")
# 
# melted_generation_hobs_Line_23_2_summary <- melt(generation_hobs_Line_23_2_summary)
# 
# ggplot(melted_generation_hobs_Line_23_2, aes(x = value, fill = variable)) +
#   geom_density(alpha = 0.3) +
#   theme_Publication()
# 
# ggplot(melted_generation_hobs_Line_23_2_summary, aes(x = variable, y = value)) +
#   geom_point() +
#   scale_y_continuous(breaks = c(0.1, 0.2, 0.3, 0.4, 0.5 ,0.6, 0.7, 0.8, 0.9, 1)) +
#   ylim(c(0,1))
# 
# expectations_halving_Line_23_2 <- melted_generation_hobs_Line_23_2 %>% 
#   filter(variable == "F") %>% 
#   mutate(S1 = (value / 2), S2 = (value / 4), S3 = (value / 8), S4 = (value/ 16), S5 = (value / 32)) %>% 
#   rename(F = value) %>% 
#   select(F, S1, S2, S3, S4, S5)
# 
# expectations_halving_Line_23_2_summary <- data.frame(FS = mean(expectations_halving_Line_23_2$F, na.rm = T), 
#                                                      S1 = mean(expectations_halving_Line_23_2$S1, na.rm = T),
#                                                      S2 = mean(expectations_halving_Line_23_2$S2, na.rm = T),
#                                                      S3 = mean(expectations_halving_Line_23_2$S3, na.rm = T),
#                                                      S4 = mean(expectations_halving_Line_23_2$S4, na.rm = T),
#                                                      S5 = mean(expectations_halving_Line_23_2$S5, na.rm = T),
#                                                      H = "Expected")
# expectations_halving_Line_23_2_summary_melted <- melt(expectations_halving_Line_23_2_summary)
# 
# obs_exp_23_2 <- rbind(expectations_halving_Line_23_2_summary_melted, melted_generation_hobs_Line_23_2_summary)
# 
# obs_exp_23_2_plot <- ggplot(obs_exp_23_2, aes(x = variable, y = value, colour = H)) +
#   geom_point(size = 3) +
#   scale_y_continuous(breaks = c(0.1, 0.2, 0.3, 0.4, 0.5 ,0.6, 0.7, 0.8, 0.9, 1)) +
#   ylim(c(0,1)) +
#   scale_colour_nejm() +
#   ylab("Heterozygosity") +
#   xlab("Generation") +
#   theme_Publication() +
#   theme(legend.position = "none") +
#   ggtitle("Line 23_2")
# 
# wilcox.test(generation_hobs_Line_23_2$S1, expectations_halving_Line_23_2$S1, paired = T)
# wilcox.test(generation_hobs_Line_23_2$S2, expectations_halving_Line_23_2$S2, paired = T)
# wilcox.test(generation_hobs_Line_23_2$S3, expectations_halving_Line_23_2$S3, paired = T)
# wilcox.test(generation_hobs_Line_23_2$S4, expectations_halving_Line_23_2$S4, paired = T)
# wilcox.test(generation_hobs_Line_23_2$S5, expectations_halving_Line_23_2$S5, paired = T)
# 
# write.table(generation_hobs_Line_23_2, "Line_23_2_hobs.txt", col.names = T, row.names = T, quote = F)
# write.table(expectations_halving_Line_23_2, "Line_23_2_hexp.txt", col.names = T, row.names = F, quote = F)
# 
# # Line 23_4
# generation_hobs_Line_23_4 <- data.frame(F_Hobs = Line_23_4_F_div$Hobs,
#                                         S1_Hobs = Line_23_4_S1_div$Hobs,
#                                         S2_Hobs = Line_23_4_S2_div$Hobs,
#                                         S3_Hobs = Line_23_4_S3_div$Hobs,
#                                         S4_Hobs = Line_23_4_S4_div$Hobs)
# 
# colnames(generation_hobs_Line_23_4) <- c("F", "S1", "S2", "S3", "S4")
# # generation_hobs <- rownames_to_column(generation_hobs_Line_23_4)
# melted_generation_hobs_Line_23_4 <- melt(generation_hobs_Line_23_4)
# 
# generation_hobs_Line_23_4_summary <- data.frame(FS = mean(generation_hobs_Line_23_4$F, na.rm = T), 
#                                                 S1 = mean(generation_hobs_Line_23_4$S1, na.rm = T),
#                                                 S2 = mean(generation_hobs_Line_23_4$S2, na.rm = T),
#                                                 S3 = mean(generation_hobs_Line_23_4$S3, na.rm = T),
#                                                 S4 = mean(generation_hobs_Line_23_4$S4, na.rm = T),
#                                                 H = "Observed")
# 
# melted_generation_hobs_Line_23_4_summary <- melt(generation_hobs_Line_23_4_summary)
# 
# ggplot(melted_generation_hobs_Line_23_4, aes(x = value, fill = variable)) +
#   geom_density(alpha = 0.3) +
#   theme_Publication()
# 
# ggplot(melted_generation_hobs_Line_23_4_summary, aes(x = variable, y = value)) +
#   geom_point() +
#   scale_y_continuous(breaks = c(0.1, 0.2, 0.3, 0.4, 0.5 ,0.6, 0.7, 0.8, 0.9, 1)) +
#   ylim(c(0,1))
# 
# expectations_halving_Line_23_4 <- melted_generation_hobs_Line_23_4 %>% 
#   filter(variable == "F") %>% 
#   mutate(S1 = (value / 2), S2 = (value / 4), S3 = (value / 8), S4 = (value/ 16)) %>% 
#   rename(F = value) %>% 
#   select(F, S1, S2, S3, S4)
# 
# expectations_halving_Line_23_4_summary <- data.frame(FS = mean(expectations_halving_Line_23_4$F, na.rm = T), 
#                                                      S1 = mean(expectations_halving_Line_23_4$S1, na.rm = T),
#                                                      S2 = mean(expectations_halving_Line_23_4$S2, na.rm = T),
#                                                      S3 = mean(expectations_halving_Line_23_4$S3, na.rm = T),
#                                                      S4 = mean(expectations_halving_Line_23_4$S4, na.rm = T),
#                                                      H = "Expected")
# expectations_halving_Line_23_4_summary_melted <- melt(expectations_halving_Line_23_4_summary)
# 
# obs_exp_23_4 <- rbind(expectations_halving_Line_23_4_summary_melted, melted_generation_hobs_Line_23_4_summary)
# 
# obs_exp_23_4_plot <- ggplot(obs_exp_23_4, aes(x = variable, y = value, colour = H)) +
#   geom_point(size = 3) +
#   scale_y_continuous(breaks = c(0.1, 0.2, 0.3, 0.4, 0.5 ,0.6, 0.7, 0.8, 0.9, 1)) +
#   ylim(c(0,1)) +
#   scale_colour_nejm() +
#   ylab("Heterozygosity") +
#   xlab("Generation") +
#   theme_Publication() +
#   theme(legend.position = "none") +
#   ggtitle("Line 23_4")
# 
# wilcox.test(generation_hobs_Line_23_4$S1, expectations_halving_Line_23_4$S1, paired = T)
# wilcox.test(generation_hobs_Line_23_4$S2, expectations_halving_Line_23_4$S2, paired = T)
# wilcox.test(generation_hobs_Line_23_4$S3, expectations_halving_Line_23_4$S3, paired = T)
# wilcox.test(generation_hobs_Line_23_4$S4, expectations_halving_Line_23_4$S4, paired = T)
# 
# write.table(generation_hobs_Line_23_4, "Line_23_4_hobs.txt", col.names = T, row.names = T, quote = F)
# write.table(expectations_halving_Line_23_4, "Line_23_4_hexp.txt", col.names = T, row.names = F, quote = F)
# 
# # Line 26_1
# generation_hobs_Line_26_1 <- data.frame(F_Hobs = Line_26_1_F_div$Hobs,
#                                         S1_Hobs = Line_26_1_S1_div$Hobs,
#                                         S2_Hobs = Line_26_1_S2_div$Hobs,
#                                         S3_Hobs = Line_26_1_S3_div$Hobs,
#                                         S4_Hobs = Line_26_1_S4_div$Hobs)
# 
# colnames(generation_hobs_Line_26_1) <- c("F", "S1", "S2", "S3", "S4")
# # generation_hobs <- rownames_to_column(generation_hobs_Line_26_1)
# melted_generation_hobs_Line_26_1 <- melt(generation_hobs_Line_26_1)
# 
# generation_hobs_Line_26_1_summary <- data.frame(FS = mean(generation_hobs_Line_26_1$F, na.rm = T), 
#                                                 S1 = mean(generation_hobs_Line_26_1$S1, na.rm = T),
#                                                 S2 = mean(generation_hobs_Line_26_1$S2, na.rm = T),
#                                                 S3 = mean(generation_hobs_Line_26_1$S3, na.rm = T),
#                                                 S4 = mean(generation_hobs_Line_26_1$S4, na.rm = T),
#                                                 H = "Observed")
# 
# melted_generation_hobs_Line_26_1_summary <- melt(generation_hobs_Line_26_1_summary)
# 
# ggplot(melted_generation_hobs_Line_26_1, aes(x = value, fill = variable)) +
#   geom_density(alpha = 0.3) +
#   theme_Publication()
# 
# ggplot(melted_generation_hobs_Line_26_1_summary, aes(x = variable, y = value)) +
#   geom_point() +
#   scale_y_continuous(breaks = c(0.1, 0.2, 0.3, 0.4, 0.5 ,0.6, 0.7, 0.8, 0.9, 1)) +
#   ylim(c(0,1))
# 
# expectations_halving_Line_26_1 <- melted_generation_hobs_Line_26_1 %>% 
#   filter(variable == "F") %>% 
#   mutate(S1 = (value / 2), S2 = (value / 4), S3 = (value / 8), S4 = (value/ 16)) %>% 
#   rename(F = value) %>% 
#   select(F, S1, S2, S3, S4)
# 
# expectations_halving_Line_26_1_summary <- data.frame(FS = mean(expectations_halving_Line_26_1$F, na.rm = T), 
#                                                      S1 = mean(expectations_halving_Line_26_1$S1, na.rm = T),
#                                                      S2 = mean(expectations_halving_Line_26_1$S2, na.rm = T),
#                                                      S3 = mean(expectations_halving_Line_26_1$S3, na.rm = T),
#                                                      S4 = mean(expectations_halving_Line_26_1$S4, na.rm = T),
#                                                      H = "Expected")
# expectations_halving_Line_26_1_summary_melted <- melt(expectations_halving_Line_26_1_summary)
# 
# obs_exp_26_1 <- rbind(expectations_halving_Line_26_1_summary_melted, melted_generation_hobs_Line_26_1_summary)
# 
# obs_exp_26_1_plot <- ggplot(obs_exp_26_1, aes(x = variable, y = value, colour = H)) +
#   geom_point(size = 3) +
#   scale_y_continuous(breaks = c(0.1, 0.2, 0.3, 0.4, 0.5 ,0.6, 0.7, 0.8, 0.9, 1)) +
#   ylim(c(0,1)) +
#   scale_colour_nejm() +
#   ylab("Heterozygosity") +
#   xlab("Generation") +
#   theme_Publication() +
#   theme(legend.position = "none") +
#   ggtitle("Line 26_1")
# 
# wilcox.test(generation_hobs_Line_26_1$S1, expectations_halving_Line_26_1$S1, paired = T)
# wilcox.test(generation_hobs_Line_26_1$S2, expectations_halving_Line_26_1$S2, paired = T)
# wilcox.test(generation_hobs_Line_26_1$S3, expectations_halving_Line_26_1$S3, paired = T)
# wilcox.test(generation_hobs_Line_26_1$S4, expectations_halving_Line_26_1$S4, paired = T)
# 
# write.table(generation_hobs_Line_26_1, "Line_26_1_hobs.txt", col.names = T, row.names = T, quote = F)
# write.table(expectations_halving_Line_26_1, "Line_26_1_hexp.txt", col.names = T, row.names = F, quote = F)
# 
# # Line 26_4
# generation_hobs_Line_26_4 <- data.frame(F_Hobs = Line_26_4_F_div$Hobs,
#                                         S1_Hobs = Line_26_4_S1_div$Hobs,
#                                         S2_Hobs = Line_26_4_S2_div$Hobs,
#                                         S3_Hobs = Line_26_4_S3_div$Hobs,
#                                         S4_Hobs = Line_26_4_S4_div$Hobs)
# 
# colnames(generation_hobs_Line_26_4) <- c("F", "S1", "S2", "S3", "S4")
# # generation_hobs <- rownames_to_column(generation_hobs_Line_26_4)
# melted_generation_hobs_Line_26_4 <- melt(generation_hobs_Line_26_4)
# 
# generation_hobs_Line_26_4_summary <- data.frame(FS = mean(generation_hobs_Line_26_4$F, na.rm = T), 
#                                                 S1 = mean(generation_hobs_Line_26_4$S1, na.rm = T),
#                                                 S2 = mean(generation_hobs_Line_26_4$S2, na.rm = T),
#                                                 S3 = mean(generation_hobs_Line_26_4$S3, na.rm = T),
#                                                 S4 = mean(generation_hobs_Line_26_4$S4, na.rm = T),
#                                                 H = "Observed")
# 
# melted_generation_hobs_Line_26_4_summary <- melt(generation_hobs_Line_26_4_summary)
# 
# ggplot(melted_generation_hobs_Line_26_4, aes(x = value, fill = variable)) +
#   geom_density(alpha = 0.3) +
#   theme_Publication()
# 
# ggplot(melted_generation_hobs_Line_26_4_summary, aes(x = variable, y = value)) +
#   geom_point() +
#   scale_y_continuous(breaks = c(0.1, 0.2, 0.3, 0.4, 0.5 ,0.6, 0.7, 0.8, 0.9, 1)) +
#   ylim(c(0,1))
# 
# expectations_halving_Line_26_4 <- melted_generation_hobs_Line_26_4 %>% 
#   filter(variable == "F") %>% 
#   mutate(S1 = (value / 2), S2 = (value / 4), S3 = (value / 8), S4 = (value/ 16)) %>% 
#   rename(F = value) %>% 
#   select(F, S1, S2, S3, S4)
# 
# expectations_halving_Line_26_4_summary <- data.frame(FS = mean(expectations_halving_Line_26_4$F, na.rm = T), 
#                                                      S1 = mean(expectations_halving_Line_26_4$S1, na.rm = T),
#                                                      S2 = mean(expectations_halving_Line_26_4$S2, na.rm = T),
#                                                      S3 = mean(expectations_halving_Line_26_4$S3, na.rm = T),
#                                                      S4 = mean(expectations_halving_Line_26_4$S4, na.rm = T),
#                                                      H = "Expected")
# expectations_halving_Line_26_4_summary_melted <- melt(expectations_halving_Line_26_4_summary)
# 
# obs_exp_26_4 <- rbind(expectations_halving_Line_26_4_summary_melted, melted_generation_hobs_Line_26_4_summary)
# 
# obs_exp_26_4_plot <- ggplot(obs_exp_26_4, aes(x = variable, y = value, colour = H)) +
#   geom_point(size = 3) +
#   scale_y_continuous(breaks = c(0.1, 0.2, 0.3, 0.4, 0.5 ,0.6, 0.7, 0.8, 0.9, 1)) +
#   ylim(c(0,1)) +
#   scale_colour_nejm() +
#   ylab("Heterozygosity") +
#   xlab("Generation") +
#   theme_Publication() +
#   theme(legend.position = "none") +
#   ggtitle("Line 26_4")
# 
# wilcox.test(generation_hobs_Line_26_4$S1, expectations_halving_Line_26_4$S1, paired = T)
# wilcox.test(generation_hobs_Line_26_4$S2, expectations_halving_Line_26_4$S2, paired = T)
# wilcox.test(generation_hobs_Line_26_4$S3, expectations_halving_Line_26_4$S3, paired = T)
# wilcox.test(generation_hobs_Line_26_4$S4, expectations_halving_Line_26_4$S4, paired = T)
# 
# write.table(generation_hobs_Line_26_4, "Line_26_4_hobs.txt", col.names = T, row.names = T, quote = F)
# write.table(expectations_halving_Line_26_4, "Line_26_4_hexp.txt", col.names = T, row.names = F, quote = F)
# 
# # Line 27_2
# generation_hobs_Line_27_2 <- data.frame(F_Hobs = Line_27_2_F_div$Hobs,
#                                         S1_Hobs = Line_27_2_S1_div$Hobs,
#                                         S2_Hobs = Line_27_2_S2_div$Hobs,
#                                         S3_Hobs = Line_27_2_S3_div$Hobs,
#                                         S4_Hobs = Line_27_2_S4_div$Hobs)
# 
# colnames(generation_hobs_Line_27_2) <- c("F", "S1", "S2", "S3", "S4")
# # generation_hobs <- rownames_to_column(generation_hobs_Line_27_2)
# melted_generation_hobs_Line_27_2 <- melt(generation_hobs_Line_27_2)
# 
# generation_hobs_Line_27_2_summary <- data.frame(FS = mean(generation_hobs_Line_27_2$F, na.rm = T), 
#                                                 S1 = mean(generation_hobs_Line_27_2$S1, na.rm = T),
#                                                 S2 = mean(generation_hobs_Line_27_2$S2, na.rm = T),
#                                                 S3 = mean(generation_hobs_Line_27_2$S3, na.rm = T),
#                                                 S4 = mean(generation_hobs_Line_27_2$S4, na.rm = T),
#                                                 H = "Observed")
# 
# melted_generation_hobs_Line_27_2_summary <- melt(generation_hobs_Line_27_2_summary)
# 
# ggplot(melted_generation_hobs_Line_27_2, aes(x = value, fill = variable)) +
#   geom_density(alpha = 0.3) +
#   theme_Publication()
# 
# ggplot(melted_generation_hobs_Line_27_2_summary, aes(x = variable, y = value)) +
#   geom_point() +
#   scale_y_continuous(breaks = c(0.1, 0.2, 0.3, 0.4, 0.5 ,0.6, 0.7, 0.8, 0.9, 1)) +
#   ylim(c(0,1))
# 
# expectations_halving_Line_27_2 <- melted_generation_hobs_Line_27_2 %>% 
#   filter(variable == "F") %>% 
#   mutate(S1 = (value / 2), S2 = (value / 4), S3 = (value / 8), S4 = (value/ 16)) %>% 
#   rename(F = value) %>% 
#   select(F, S1, S2, S3, S4)
# 
# expectations_halving_Line_27_2_summary <- data.frame(FS = mean(expectations_halving_Line_27_2$F, na.rm = T), 
#                                                      S1 = mean(expectations_halving_Line_27_2$S1, na.rm = T),
#                                                      S2 = mean(expectations_halving_Line_27_2$S2, na.rm = T),
#                                                      S3 = mean(expectations_halving_Line_27_2$S3, na.rm = T),
#                                                      S4 = mean(expectations_halving_Line_27_2$S4, na.rm = T),
#                                                      H = "Expected")
# expectations_halving_Line_27_2_summary_melted <- melt(expectations_halving_Line_27_2_summary)
# 
# obs_exp_27_2 <- rbind(expectations_halving_Line_27_2_summary_melted, melted_generation_hobs_Line_27_2_summary)
# 
# obs_exp_27_2_plot <- ggplot(obs_exp_27_2, aes(x = variable, y = value, colour = H)) +
#   geom_point(size = 3) +
#   scale_y_continuous(breaks = c(0.1, 0.2, 0.3, 0.4, 0.5 ,0.6, 0.7, 0.8, 0.9, 1)) +
#   ylim(c(0,1)) +
#   scale_colour_nejm() +
#   ylab("Heterozygosity") +
#   xlab("Generation") +
#   theme_Publication() +
#   theme(legend.position = "none") +
#   ggtitle("Line 27_2")
# 
# wilcox.test(generation_hobs_Line_27_2$S1, expectations_halving_Line_27_2$S1, paired = T)
# wilcox.test(generation_hobs_Line_27_2$S2, expectations_halving_Line_27_2$S2, paired = T)
# wilcox.test(generation_hobs_Line_27_2$S3, expectations_halving_Line_27_2$S3, paired = T)
# wilcox.test(generation_hobs_Line_27_2$S4, expectations_halving_Line_27_2$S4, paired = T)
# 
# write.table(generation_hobs_Line_27_2, "Line_27_2_hobs.txt", col.names = T, row.names = T, quote = F)
# write.table(expectations_halving_Line_27_2, "Line_27_2_hexp.txt", col.names = T, row.names = F, quote = F)
# 
# 
# # Line 29_2
# generation_hobs_Line_29_2 <- data.frame(F_Hobs = Line_29_2_F_div$Hobs,
#                                         S1_Hobs = Line_29_2_S1_div$Hobs,
#                                         S2_Hobs = Line_29_2_S2_div$Hobs,
#                                         S3_Hobs = Line_29_2_S3_div$Hobs,
#                                         S4_Hobs = Line_29_2_S4_div$Hobs)
# 
# colnames(generation_hobs_Line_29_2) <- c("F", "S1", "S2", "S3", "S4")
# # generation_hobs <- rownames_to_column(generation_hobs_Line_29_2)
# melted_generation_hobs_Line_29_2 <- melt(generation_hobs_Line_29_2)
# 
# generation_hobs_Line_29_2_summary <- data.frame(FS = mean(generation_hobs_Line_29_2$F, na.rm = T), 
#                                                 S1 = mean(generation_hobs_Line_29_2$S1, na.rm = T),
#                                                 S2 = mean(generation_hobs_Line_29_2$S2, na.rm = T),
#                                                 S3 = mean(generation_hobs_Line_29_2$S3, na.rm = T),
#                                                 S4 = mean(generation_hobs_Line_29_2$S4, na.rm = T),
#                                                 H = "Observed")
# 
# melted_generation_hobs_Line_29_2_summary <- melt(generation_hobs_Line_29_2_summary)
# 
# ggplot(melted_generation_hobs_Line_29_2, aes(x = value, fill = variable)) +
#   geom_density(alpha = 0.3) +
#   theme_Publication()
# 
# ggplot(melted_generation_hobs_Line_29_2_summary, aes(x = variable, y = value)) +
#   geom_point() +
#   scale_y_continuous(breaks = c(0.1, 0.2, 0.3, 0.4, 0.5 ,0.6, 0.7, 0.8, 0.9, 1)) +
#   ylim(c(0,1))
# 
# expectations_halving_Line_29_2 <- melted_generation_hobs_Line_29_2 %>% 
#   filter(variable == "F") %>% 
#   mutate(S1 = (value / 2), S2 = (value / 4), S3 = (value / 8), S4 = (value/ 16)) %>% 
#   rename(F = value) %>% 
#   select(F, S1, S2, S3, S4)
# 
# expectations_halving_Line_29_2_summary <- data.frame(FS = mean(expectations_halving_Line_29_2$F, na.rm = T), 
#                                                      S1 = mean(expectations_halving_Line_29_2$S1, na.rm = T),
#                                                      S2 = mean(expectations_halving_Line_29_2$S2, na.rm = T),
#                                                      S3 = mean(expectations_halving_Line_29_2$S3, na.rm = T),
#                                                      S4 = mean(expectations_halving_Line_29_2$S4, na.rm = T),
#                                                      H = "Expected")
# expectations_halving_Line_29_2_summary_melted <- melt(expectations_halving_Line_29_2_summary)
# 
# obs_exp_29_2 <- rbind(expectations_halving_Line_29_2_summary_melted, melted_generation_hobs_Line_29_2_summary)
# 
# obs_exp_29_2_plot <- ggplot(obs_exp_29_2, aes(x = variable, y = value, colour = H)) +
#   geom_point(size = 3) +
#   scale_y_continuous(breaks = c(0.1, 0.2, 0.3, 0.4, 0.5 ,0.6, 0.7, 0.8, 0.9, 1)) +
#   ylim(c(0,1)) +
#   scale_colour_nejm() +
#   ylab("Heterozygosity") +
#   xlab("Generation") +
#   theme_Publication() +
#   theme(legend.position = "none") +
#   ggtitle("Line 29_2")
# 
# wilcox.test(generation_hobs_Line_29_2$S1, expectations_halving_Line_29_2$S1, paired = T)
# wilcox.test(generation_hobs_Line_29_2$S2, expectations_halving_Line_29_2$S2, paired = T)
# wilcox.test(generation_hobs_Line_29_2$S3, expectations_halving_Line_29_2$S3, paired = T)
# wilcox.test(generation_hobs_Line_29_2$S4, expectations_halving_Line_29_2$S4, paired = T)
# 
# write.table(generation_hobs_Line_29_2, "Line_29_2_hobs.txt", col.names = T, row.names = T, quote = F)
# write.table(expectations_halving_Line_29_2, "Line_29_2_hexp.txt", col.names = T, row.names = F, quote = F)
# 
# # Line 29_4
# generation_hobs_Line_29_4 <- data.frame(F_Hobs = Line_29_4_F_div$Hobs,
#                                         S1_Hobs = Line_29_4_S1_div$Hobs,
#                                         S2_Hobs = Line_29_4_S2_div$Hobs,
#                                         S3_Hobs = Line_29_4_S3_div$Hobs,
#                                         S4_Hobs = Line_29_4_S4_div$Hobs)
# 
# colnames(generation_hobs_Line_29_4) <- c("F", "S1", "S2", "S3", "S4")
# # generation_hobs <- rownames_to_column(generation_hobs_Line_29_4)
# melted_generation_hobs_Line_29_4 <- melt(generation_hobs_Line_29_4)
# 
# generation_hobs_Line_29_4_summary <- data.frame(FS = mean(generation_hobs_Line_29_4$F, na.rm = T), 
#                                                 S1 = mean(generation_hobs_Line_29_4$S1, na.rm = T),
#                                                 S2 = mean(generation_hobs_Line_29_4$S2, na.rm = T),
#                                                 S3 = mean(generation_hobs_Line_29_4$S3, na.rm = T),
#                                                 S4 = mean(generation_hobs_Line_29_4$S4, na.rm = T),
#                                                 H = "Observed")
# 
# melted_generation_hobs_Line_29_4_summary <- melt(generation_hobs_Line_29_4_summary)
# 
# ggplot(melted_generation_hobs_Line_29_4, aes(x = value, fill = variable)) +
#   geom_density(alpha = 0.3) +
#   theme_Publication()
# 
# ggplot(melted_generation_hobs_Line_29_4_summary, aes(x = variable, y = value)) +
#   geom_point() +
#   scale_y_continuous(breaks = c(0.1, 0.2, 0.3, 0.4, 0.5 ,0.6, 0.7, 0.8, 0.9, 1)) +
#   ylim(c(0,1))
# 
# expectations_halving_Line_29_4 <- melted_generation_hobs_Line_29_4 %>% 
#   filter(variable == "F") %>% 
#   mutate(S1 = (value / 2), S2 = (value / 4), S3 = (value / 8), S4 = (value/ 16)) %>% 
#   rename(F = value) %>% 
#   select(F, S1, S2, S3, S4)
# 
# expectations_halving_Line_29_4_summary <- data.frame(FS = mean(expectations_halving_Line_29_4$F, na.rm = T), 
#                                                      S1 = mean(expectations_halving_Line_29_4$S1, na.rm = T),
#                                                      S2 = mean(expectations_halving_Line_29_4$S2, na.rm = T),
#                                                      S3 = mean(expectations_halving_Line_29_4$S3, na.rm = T),
#                                                      S4 = mean(expectations_halving_Line_29_4$S4, na.rm = T),
#                                                      H = "Expected")
# expectations_halving_Line_29_4_summary_melted <- melt(expectations_halving_Line_29_4_summary)
# 
# obs_exp_29_4 <- rbind(expectations_halving_Line_29_4_summary_melted, melted_generation_hobs_Line_29_4_summary)
# 
# obs_exp_29_4_plot <- ggplot(obs_exp_29_4, aes(x = variable, y = value, colour = H)) +
#   geom_point(size = 3) +
#   scale_y_continuous(breaks = c(0.1, 0.2, 0.3, 0.4, 0.5 ,0.6, 0.7, 0.8, 0.9, 1)) +
#   ylim(c(0,1)) +
#   scale_colour_nejm() +
#   ylab("Heterozygosity") +
#   xlab("Generation") +
#   theme_Publication() +
#   ggtitle("Line 29_4")
# 
# wilcox.test(generation_hobs_Line_29_4$S1, expectations_halving_Line_29_4$S1, paired = T)
# wilcox.test(generation_hobs_Line_29_4$S2, expectations_halving_Line_29_4$S2, paired = T)
# wilcox.test(generation_hobs_Line_29_4$S3, expectations_halving_Line_29_4$S3, paired = T)
# wilcox.test(generation_hobs_Line_29_4$S4, expectations_halving_Line_29_4$S4, paired = T)
# 
# write.table(generation_hobs_Line_29_4, "Line_29_4_hobs.txt", col.names = T, row.names = T, quote = F)
# write.table(expectations_halving_Line_29_4, "Line_29_4_hexp.txt", col.names = T, row.names = F, quote = F)
# 
# all_line_heterozygosity_plots <- ggarrange(obs_exp_1_1_plot, obs_exp_6_1_plot, obs_exp_6_4_plot, obs_exp_7_2_plot, obs_exp_7_4_plot, 
#                                            obs_exp_8_2_plot, obs_exp_8_4_plot, obs_exp_13_3_plot, obs_exp_13_4_plot, obs_exp_16_1_plot, 
#                                            obs_exp_16_5_plot, obs_exp_17_2_plot, obs_exp_17_5_plot, obs_exp_19_2_plot, obs_exp_19_5_plot, 
#                                            obs_exp_20_1_plot, obs_exp_20_4_plot, obs_exp_21_2_plot, obs_exp_21_6_plot, obs_exp_23_2_plot, 
#                                            obs_exp_23_4_plot, obs_exp_26_1_plot, obs_exp_26_4_plot, obs_exp_27_2_plot, obs_exp_29_2_plot, 
#                                            obs_exp_29_4_plot)
# 
# ggsave("all_line_heterozygosity_plots.svg", all_line_heterozygosity_plots, width = 25, height = 20)
# 
# ### Combining lines for figure ###########
# 
# FS_S5_mean_het <- data.frame(rbind(generation_hobs_Line_1_1_summary, generation_hobs_Line_6_4_summary, generation_hobs_Line_17_2_summary,
#                                    generation_hobs_Line_17_5_summary, generation_hobs_Line_21_2_summary, generation_hobs_Line_21_6_summary, 
#                                    generation_hobs_Line_23_2_summary, generation_hobs_Line_29_2_summary, generation_hobs_Line_29_4_summary)) %>% 
#   mutate(Line = factor(c("1_1", "6_4", "17_2", "17_5", "21_2", "21_6", "23_2", "29_2", "29_4")))
# 
# melted_FS_S5_mean_het <- melt(FS_S5_mean_het)
# 
# ggplot(melted_FS_S5_mean_het, aes(x = variable, y = value, fill = Line, colour = Line, group = Line)) +
#   geom_point(size = 2) +
#   geom_line() +
#   ylim(0, 1) +
#   scale_fill_nejm() +
#   theme_Publication()


## Where are het SNPs on putative linkage groups (unused)

# het_snps <- generation_FS_S5_hobs %>%
#   filter(S4_hobs == 1) 
# 
# 
# clipr::write_clip(rownames(het_snps))
# 
# # plot het snps on LG
# WRC_LG <- read.table("~/UBC/GSAT/PhD/WRC/GS/wrc/snps/cedar/FINAL_DATA_SET/filtered_normalised/traits_snps_from_genome_paper/bayesR_maf_filter/WRC_chromosomes.txt", header = T)
# # WRC_LG$chr <- factor(WRC_LG$chr)
# 
# het_snps_in_LG <- het_snps %>% 
#   rownames_to_column() %>% 
#   filter(rowname %in% WRC_LG$SNP) %>% 
#   rename(SNP = rowname)
# 
# 
# 
# het_snps_in_LG <- merge(het_snps_in_LG,WRC_LG, by = "SNP")
# 
# het_snps_in_LG <- het_snps_in_LG %>% 
#   arrange(chr,accurate_pos)
# 
# het_snps_in_LG$SNP <- factor(het_snps_in_LG$SNP, levels=unique(het_snps_in_LG$SNP))
# 
# het_snps_in_LG <- het_snps_in_LG %>% 
#   select(SNP, chr, accurate_pos)
# 
# don_het_snps_in_LG <- het_snps_in_LG %>% 
#   
#   # Compute chromosome size
#   group_by(chr) %>% 
#   summarise(chr_len=max(accurate_pos)) %>% 
#   
#   # Calculate cumulative position of each chromosome
#   mutate(tot=cumsum(as.numeric(chr_len))-as.numeric(chr_len)) %>%
#   select(-chr_len) %>%
#   
#   # Add this info to the initial dataset
#   left_join(het_snps_in_LG, ., by=c("chr"="chr")) %>%
#   
#   # Add a cumulative position of each SNP
#   arrange(chr, accurate_pos) %>%
#   mutate(BPcum=accurate_pos+tot)
# 
# axisdf = don_het_snps_in_LG %>% group_by(chr) %>% summarize(center=( max(BPcum) + min(BPcum)) / 2 )
# 
# het_snps_in_LG_plot <- ggplot(don_het_snps_in_LG, aes(x=BPcum, y = 1)) +
#   
#   # Show all points
#   geom_point(alpha = 0.8) +
#   scale_color_nejm() +
#   
#   # custom X axis:
#   scale_x_continuous(label = axisdf$chr, breaks= axisdf$center) +
#   # scale_y_continuous(expand = c(0, 0) ) +     # remove space between plot area and x axis
#   # geom_vline(xintercept = c(888232888, 1683028619, 2539001849, 3225909214, 3875153489, 4496253278, 5139632346, 5771913333, 6413980409, 7022253572), linetype = "dashed") +
#   xlab("Linkage Group") +
#   
#   # Custom the theme:
#   theme_Publication() +
#   theme( 
#     legend.position="none",
#     panel.border = element_blank(),
#     panel.grid.major.x = element_blank(),
#     panel.grid.minor.x = element_blank()
#   )
# 
# het_snps_in_LG_plot
```
