## Supplementary material for "The western redcedar genome reveals low genetic diversity in a self-compatible conifer": Code S7: inbreeding_coefficients.R

```
rm(list = ls())

library(asreml)
library(asremlPlus)
library(nadiv)
library(synbreed)
library(sommer)
library(ggsci)
library(tidyverse)

source("publication_theme.r")

library(extrafont)
library(extrafontdb)

# RWP IB
RWP_ibc <- read.table("rwp_unlinked_mac3_snps.id.ibc", header = T)
RWP_ibc <- RWP_ibc %>% 
  mutate(Fhat3_noneg = if_else(Fhat3 < 0, 0, Fhat3))

# S lines IB
S_lines_ibc <- read.table("corrected_snps_S_lines_unlinked_mac3.id.ibc", header = T)
S_lines_ibc <- S_lines_ibc %>% 
  mutate(Generation = c(rep("FS", 28), rep("S1", 28), rep("S2", 28), rep("S3", 28), rep("S4", 28), rep("S5", 11)),
         exp = c(rep(0, 28), rep(0.5, 28), rep(0.75, 28), rep(0.875, 28), rep(0.9375, 28), rep(0.96875, 11)))

## Summary stats
S_lines_ibc %>% 
  group_by(Generation) %>% 
  summarise(mean = mean(Fhat3, na.rm = T), median = median(Fhat3, na.rm = T), sd = sd(Fhat3))

RWP_ibc %>% 
  summarise(mean = mean(Fhat3, na.rm = T), median = median(Fhat3, na.rm = T), sd = sd(Fhat3))


# Plot for Figure 4
fhat_gen_violin <- ggplot(S_lines_ibc, aes(x = Generation, y = Fhat3)) +
  geom_violin(scale = "area") +
  geom_point(aes(x = Generation, y = exp), shape = 23, fill = "red", size = 3) +
  geom_boxplot(width = 0.15) +
  theme_Publication() +
  scale_color_nejm() +
  scale_x_discrete("Generation", breaks = unique(S_lines_ibc$Generation),
                   labels = c("FS",
                              "S1",
                              "S2",
                              "S3",
                              "S4",
                              "S5")) +
  scale_y_continuous(name = expression(paste(bolditalic("F"))), breaks = c(0, 0.2, 0.4, 0.6, 0.8, 1),limits = c(0, 1))

fhat_gen_violin


ggsave("fhat3_S_violin_r2_mac_snps.tiff", fhat_gen_violin, dpi = 300, width = 7, height = 6)

# Some stats
FS_lines_ibc <- S_lines_ibc %>% 
  filter(Generation == "FS")

S1_lines_ibc <- S_lines_ibc %>% 
  filter(Generation == "S1")

S2_lines_ibc <- S_lines_ibc %>% 
  filter(Generation == "S2")

S3_lines_ibc <- S_lines_ibc %>% 
  filter(Generation == "S3")

S4_lines_ibc <- S_lines_ibc %>% 
  filter(Generation == "S4")

S5_lines_ibc <- S_lines_ibc %>% 
  filter(Generation == "S5")

t.test(FS_lines_ibc$Fhat3, mu = 0)
t.test(S1_lines_ibc$Fhat3, mu = 0.5)
t.test(S2_lines_ibc$Fhat3, mu = 0.75)
t.test(S3_lines_ibc$Fhat3, mu = 0.875)
t.test(S4_lines_ibc$Fhat3, mu = 0.9375)
t.test(S5_lines_ibc$Fhat3, mu = 0.96875)

# chisq.test(S1_lines_ibc$Fhat3_noneg, S1_lines_ibc$exp,)
# BSDA::z.test(S1_lines_ibc$Fhat3_noneg, mu =0.5, sigma.x = 0.05)
# BSDA::z.test(S2_lines_ibc$Fhat3_noneg, mu =0.75, sigma.x = 0.05)
# BSDA::z.test(S3_lines_ibc$Fhat3_noneg, mu =0.875, sigma.x = 0.05)
# BSDA::z.test(S4_lines_ibc$Fhat3_noneg, mu =0.9375, sigma.x = 0.05)
# BSDA::z.test(S5_lines_ibc$Fhat3_noneg, mu =0.96875, sigma.x = 0.05)
```
