## Supplementary material for "The western redcedar genome reveals low genetic diversity in a self-compatible conifer": Code S8: variant_effect_predictions_and_snp_annotation.R

```
rm(list=ls())

library(gtsummary)
library(reshape2)
library(ggsci)
library(tidyverse)
source("publication_theme.r")

library(extrafont)
library(extrafontdb)

RWP_vep_all <- read.table("vep_all_snps.txt", header = T, na.strings = "-", stringsAsFactors = F)
outlier_snps <- read.table("outlier_snps.txt", header = F)
RWP_maf <- read.table("RWP_maf.txt", header = T)
wrc_sco_genes <- read.table("wrc_sco_genes.txt")
final_snp_ids <- read.table("all_filtered_snp_ids.txt")

# Get SNPs in SCO genes
sco_vep <- RWP_vep_all %>% 
  filter(Gene %in% wrc_sco_genes$V1 | is.na(Feature_type))

# write.table(sco_vep$SNP, "sco_snps_all.txt", quote = F, col.names = F, row.names = F)

# Filter RWP vep

RWP_vep <- RWP_vep_all %>% 
  filter(SNP %in% final_snp_ids$V1)

# write.csv(RWP_vep, "filtered_vep.csv")

summary(levels(factor(RWP_vep$Scaffold)))

RWP_vep %>% 
  filter(!(Consequence == "upstream_gene_variant" | Consequence == "downstream_gene_variant" | Consequence == "intergenic_variant")) %>% 
  group_by(factor(Gene)) %>% 
  summarise(factor(Gene), n = n())

RWP_vep %>%
  filter(!(Consequence == "upstream_gene_variant" | Consequence == "downstream_gene_variant" | Consequence == "intergenic_variant")) %>% 
  filter(Gene %in% wrc_sco_genes$V1) %>% 
  group_by(factor(Gene))

table(RWP_vep$Consequence)/18371*100


# Getting outlier SNPs and adding MAF
outlier_snps_vep <- RWP_vep %>%
  filter(SNP %in% outlier_snps$V1) %>% 
  arrange(SNP)

maf_selected_snps <- RWP_maf %>% 
  filter(SNP %in% outlier_snps_vep$SNP) %>% 
  arrange(SNP)

outlier_snps_vep_w_maf <- outlier_snps_vep %>% 
  mutate(maf = maf_selected_snps$MAF, minor_allele = maf_selected_snps$minor_allele)

# SNP consequences from vep file
SNP_consequences_parents <- RWP_vep %>%
  select(Consequence) %>%
  group_by(Consequence) %>% 
  summarise(count_parents =n()) %>% 
  mutate(freq_parents = count_parents / sum(count_parents))

SNP_consequences_outlier <- outlier_snps_vep %>% 
  select(Consequence) %>%
  group_by(Consequence) %>% 
  summarise(count_selected =n()) %>% 
  mutate(freq_selected = count_selected / sum(count_selected))

SNP_consequences_all <- merge(SNP_consequences_parents, SNP_consequences_outlier, all = T)
SNP_consequences_all[is.na(SNP_consequences_all)] <- 0

SNP_consequences_all_freq <- SNP_consequences_all %>% 
  select(Consequence, freq_parents, freq_selected) %>% 
  arrange(desc(freq_selected))

SNP_consequences_all_freq$Consequence <- factor(SNP_consequences_all_freq$Consequence, 
                                              levels = c("splice_acceptor_variant", "splice_donor_variant", "stop_gained", "stop_lost",
                                                         "start_lost", "missense_variant", "splice_region_variant", "stop_retained_variant", "synonymous_variant", 
                                                         "5_prime_UTR_variant", "3_prime_UTR_variant", "intron_variant",
                                                         "upstream_gene_variant", "downstream_gene_variant", "intergenic_variant"))

RWP_vep$Consequence <- factor(RWP_vep$Consequence, 
                                 levels = c("splice_acceptor_variant", "splice_donor_variant", "stop_gained", "stop_lost",
                                            "start_lost", "missense_variant", "splice_region_variant", "stop_retained_variant", "synonymous_variant", 
                                            "5_prime_UTR_variant", "3_prime_UTR_variant", "intron_variant",
                                            "upstream_gene_variant", "downstream_gene_variant", "intergenic_variant"))

outlier_snps_vep$Consequence <- factor(outlier_snps_vep$Consequence, 
                                          levels = c("splice_acceptor_variant", "splice_donor_variant", "stop_gained", "stop_lost",
                                                     "start_lost", "missense_variant", "splice_region_variant", "stop_retained_variant", "synonymous_variant", 
                                                     "5_prime_UTR_variant", "3_prime_UTR_variant", "intron_variant",
                                                     "upstream_gene_variant", "downstream_gene_variant", "intergenic_variant"))

melted <- melt(SNP_consequences_all_freq)

# Plot of distribution of SNP consequences in all SNPs
vep <- ggplot(RWP_vep, aes(x = Consequence)) +
  geom_histogram(stat = "count") +
  ylab("Count") +
  coord_flip() +
  theme_Publication() +
  # theme(axis.text.x = element_text(angle = 45, vjust = 0.5, hjust = 0.5)) +
  # scale_y_continuous(name = "Frequency", breaks = c(0.05, 0.1, 0.15, 0.2, 0.25, 0.3), limits = c(0,0.3)) +
  scale_x_discrete(limits = rev(levels(melted$Consequence)), breaks = c("splice_acceptor_variant", "splice_donor_variant", "stop_gained", "stop_lost",
                                                                        "start_lost", "missense_variant", "splice_region_variant", "stop_retained_variant", "synonymous_variant", 
                                                                        "5_prime_UTR_variant", "3_prime_UTR_variant", "intron_variant",
                                                                        "upstream_gene_variant", "downstream_gene_variant", "intergenic_variant"),
                   labels = c("Splice acceptor variant", "Splice donor variant", "Stop gained", "Stop lost", "Start lost", "Missense variant", 
                              "Splice region variant", "Stop retained", "Synonymous variant", "5' UTR variant", "3' UTR variant", "Intron variant", 
                              "Upstream gene variant", "Downstream gene variant", "Intergenic variant")) 
  
vep

ggplot(SNP_consequences_parents, aes(x = Consequence, y = freq_parents)) +
  geom_bar(stat = "identity") +
  theme(axis.text.x = element_text(angle = 45, vjust = 1, hjust = 1))

# ggsave("vep.svg", vep, height = 6, width = 11)
# ggsave("S_vep.svg", S_vep, height = 6, width = 11)

### Plot for Figure S7 ####
vep_outlier <- ggplot(melted, aes (x = Consequence, y =value)) +
  geom_bar(aes(fill = variable), stat = "identity", position = position_dodge()) +
  coord_flip() +
  theme_Publication() +
  # theme(axis.text.x = element_text(angle = 45, vjust = 0.5, hjust = 0.5)) +
  scale_y_continuous(name = "Frequency", breaks = c(0.1, 0.2, 0.3, 0.4), limits = c(0,0.45)) +
  scale_x_discrete(limits = rev(levels(melted$Consequence)), breaks = c("splice_acceptor_variant", "splice_donor_variant", "stop_gained", "stop_lost",
                                                                         "start_lost", "missense_variant", "splice_region_variant", "stop_retained_variant",
                                                                         "synonymous_variant", "5_prime_UTR_variant", "3_prime_UTR_variant", "intron_variant",
                                                                         "upstream_gene_variant", "downstream_gene_variant", "intergenic_variant"),
                   labels = c("Splice acceptor variant", "Splice donor variant", "Stop gained", "Stop lost",
                                                                        "Start lost", "Missense variant", "Splice region variant", "Stop retained variant",
                                                                        "Synonymous variant", "5' UTR variant", "3' UTR variant", "Intron variant",
                                                                        "Upstream gene variant", "Downstream gene variant", "Intergenic variant")) +
  # geom_signif(comparisons = list(c("Missense variant", "5' UTR variant", "3' UTR variant", "Downstream gene variant", "Intergenic variant")),
  #             map_signif_level = T) +
  scale_fill_grey(breaks=c("freq_parents", "freq_selected"),
                    labels=c("All SNPs", "Outlier SNPs"), start = 0.35) +
  guides(y_continuous = FALSE) +
  guides(x_discrete = FALSE) +
  theme(legend.title = element_blank(), legend.position = c(0.9, 0.95))
  # geom_bracket(
  #   xmin = group1, xmax = group2,
  #   y.position = 0.05, label = "*",
  #   tip.length = 0.01, coord.flip = T)
  # geom_signif(stat="identity",
  #             data=data.frame(x=c(0.875, 1.875), xend=c(1.125, 2.125),
  #                             y=c(0.25, 0.15), annotation=c("**", "NS")),
  #             aes(x=x,xend=xend, y=y, yend=y, annotation=annotation))
  
vep_outlier


# ggsave("vep_outlier_all_snps.svg", vep_outlier, height = 9, width = 11)

### Fisher Exact Test for overrepresentation of SNP consequences ####
table_maker_func <- function(x){
  matrix(c(SNP_consequences_all[x, 2], 
           (sum(SNP_consequences_all$count_parents) - SNP_consequences_all[x, 2]),
           SNP_consequences_all[x, 4],
           (sum(SNP_consequences_all$count_selected) - SNP_consequences_all[x, 4])),
         nrow = 2,
         dimnames = list(c(paste0(SNP_consequences_all[x, 1]), "Other"), 
                         c("All", "Selected")))
}  

for(i in 1:15){
  x <- table_maker_func(i)
  y <- fisher.test(x)
  z <- data.frame(Name = row.names(x)[1], p = y$p.value)
  lapply(z, write, "FET_results_compound_effects_removed_all_snps.txt", append = T)
}


### MAF for outlier SNPs, most severe consequences #### (Not used)
# S4_fate_table <- read.csv("S4_fate_table_chisq_analysis_all_snps.csv", header = T)
# 
# outlier_fixed <- outlier_snps %>% 
#   filter(ref_fixed > not_fixed | alt_fixed > not_fixed)
#   
# outlier_fixed_vep <- outlier_snps_vep_w_maf %>% 
#   filter(SNP %in% outlier_fixed$SNP)
# 
# stop_start_splice_gained_lost_snps <- outlier_snps_vep_w_maf %>% 
#   filter(Consequence =="stop_gained" | Consequence == "stop_lost" | Consequence == "start_lost" | 
#            Consequence == "splice_acceptor_variant" | Consequence == "splice_donor_variant") %>% 
#   arrange(SNP)
# 
# S4_fate_start_stop_splice_gained_lost <- S4_fate_table %>% 
#   filter(SNP %in% stop_start_splice_gained_lost_snps$SNP) %>% 
#   arrange(SNP) %>% 
#   mutate(maf = stop_start_splice_gained_lost_snps$maf, minor_allele = stop_start_splice_gained_lost_snps$minor_allele)
# 
# Mapping outliers to LGs
LG_old_snps <- read.csv("WRC_linkage_group_SNPs_nucmer.csv")
LG_outlier_snps <- LG_old_snps %>%
  filter(SNP %in% outlier_snps$V1)

LG_outlier_snps$chr <- factor(LG_outlier_snps$chr)
LG_outlier_snps$WRC_scaffold <- factor(LG_outlier_snps$WRC_scaffold)

LG_outlier_snps <- LG_outlier_snps %>%
  arrange(chr,accurate_pos) %>%
  rename(chromosome = WRC_scaffold)

LG_outlier_snps$SNP <- factor(LG_outlier_snps$SNP, levels=unique(LG_outlier_snps$SNP))

snps_in_LG <- LG_outlier_snps %>%

  # Compute chromosome size
  group_by(chr) %>%
  summarise(chr_len=max(accurate_pos)) %>%

  # Calculate cumulative position of each chromosome
  mutate(tot=cumsum(as.numeric(chr_len))-as.numeric(chr_len)) %>%
  select(-chr_len) %>%

  # Add this info to the initial dataset
  left_join(LG_outlier_snps, ., by=c("chr"="chr")) %>%

  # Add a cumulative position of each SNP
  arrange(chr, accurate_pos) %>%
  mutate(BPcum=accurate_pos+tot) %>%
  mutate(chrom_color_group = case_when(as.numeric(chr) %% 2 != 0 ~ "even",
                                       chr == "X" ~ "even",
                                       TRUE ~ "odd" )) %>%
  mutate(chromosome = factor(chr))

chr_lens <- snps_in_LG %>%
  group_by(chr) %>%
  group_map(~ max(.x$BPcum - min(.x$BPcum)))
chr_lens <- t(data.frame(chr_lens))

chr_table_df <- data.frame(table(snps_in_LG$chr), chrom_color_group = rep_len(c("odd", "even"), 11))
chr_table_df <- chr_table_df %>%
  mutate(chr_lens = as.vector(chr_lens)) %>%
  mutate(prop_total_len = chr_lens/sum(chr_lens)) %>%
  mutate(adj_amount = (Freq/chr_lens)*1e7) %>%
  mutate(exp_amount = prop_total_len*sum(Freq))

axisdf = snps_in_LG %>% group_by(chr) %>% summarize(center=( max(BPcum) + min(BPcum)) / 2 )

LG_plot <- ggplot(chr_table_df, aes(x=Var1, y = adj_amount, fill = chrom_color_group)) +

  # Show all points
  # geom_jitter(alpha = 0.8, size = 2.5) +
  geom_bar(stat = "identity") +
  # scale_color_nejm() +
  # custom X axis:
  # scale_x_continuous(label = axisdf$chr, breaks= axisdf$center) +
  # scale_y_continuous(expand = c(0, 0) ) +     # remove space between plot area and x axis
  # geom_vline(xintercept = c(888232888, 1683028619, 2539001849, 3225909214, 3875153489, 4496253278, 5139632346, 5771913333, 6413980409, 7022253572), linetype = "dashed") +
  xlab("Linkage Group") +
  ylab("Adjusted frequency") +

  # Custom the theme:
  theme_Publication() +
  theme(
    legend.position="top",
    panel.border = element_blank(),
    panel.grid.major.x = element_blank(),
    panel.grid.minor.x = element_blank()
  ) +
  guides(fill = "none") +
  # gghighlight(avg_pi > quantile(avg_pi,prob=1-1/100, na.rm = T), keep_scales = T,
  #             unhighlighted_params = list(colour = ifelse(snps_in_LG$chrom_color_group == "even", "grey50", "black"), alpha = 0.5)) +
  scale_fill_manual(values = c("grey50", "black"))

LG_plot

# ggsave("LG_barplot_outliers.tiff", dpi = 300, width = 9, height = 6)

LG_plot <- ggplot(snps_in_LG, aes(x=BPcum, y = "", color = chrom_color_group)) +

  # Show all points
  geom_jitter(alpha = 0.5, size = 1.5, width = 0, height = 0.08) +
  # geom_bar(stat = "identity") +
  # scale_color_nejm() +
  # custom X axis:
  scale_x_continuous(label = axisdf$chr, breaks= axisdf$center) +
  # scale_y_continuous(expand = c(0, 0) ) +     # remove space between plot area and x axis
  # geom_vline(xintercept = c(888232888, 1683028619, 2539001849, 3225909214, 3875153489, 4496253278, 5139632346, 5771913333, 6413980409, 7022253572), linetype = "dashed") +
  xlab("Linkage Group") +
  ylab("") +

  # Custom the theme:
  theme_Publication() +
  theme(
    legend.position="top",
    panel.border = element_blank(),
    panel.grid.major.x = element_blank(),
    panel.grid.minor.x = element_blank(),
    axis.ticks.y = element_blank()
  ) +
  guides(color = "none") +
  # gghighlight(avg_pi > quantile(avg_pi,prob=1-1/100, na.rm = T), keep_scales = T,
  #             unhighlighted_params = list(colour = ifelse(snps_in_LG$chrom_color_group == "even", "grey50", "black"), alpha = 0.5)) +
  scale_color_manual(values = c("grey50", "black"))

LG_plot

# ggsave("LG_scatterplot_outliers.tiff", dpi = 300, width = 9, height = 3)

### JGI annotations ##########################

gene_annotations <- read.delim("gene_functions.txt", header = F, stringsAsFactors = F, sep = "\t")
gff <- read.table("Tplicatav3.1.primaryTrs.gff3.gz")

all_snp_annotations <- gene_annotations %>%
  filter(V1 %in% RWP_vep$Gene) %>% 
  rename(Gene = V1)

all_snp_annotations <- merge(RWP_vep, all_snp_annotations, by = "Gene", all = T)

# write.csv(all_snp_annotations, "all_snp_annotations.csv")

## outlier snps ###
outlier_snp_annotations <- gene_annotations %>%
  filter(V1 %in% outlier_snps_vep$Gene) %>% 
  rename(Gene = V1)

outlier_snp_annotations <- merge(outlier_snps_vep, outlier_snp_annotations, by = "Gene", all = T)  

# write.csv(outlier_snp_annotations, "outlier_snp_JGI_annotations_all_snps.csv")

### GO annotations ######
outlier_snp_GO <- outlier_snp_annotations %>% 
  filter(V3 == "GO")

# write.csv(outlier_snp_GO, "outlier_snps_GO_all_snps.csv")
outlier_snp_GO_one_snp_per_gene <- read.table("outlier_snps_GO_one_snp_per_gene.csv", sep =",", header = T)
# write.csv(table(outlier_snp_GO_one_snp_per_gene$V2), "outlier_snp_GO_ID_table.csv")
# write.csv(table(outlier_snp_GO_one_snp_per_gene$V4), "outlier_snp_GO_term_table.csv")


## all snps

all_snp_annotations <- gene_annotations %>%
  filter(V1 %in% RWP_vep$Gene) %>% 
  rename(Gene = V1)

all_snp_annotations <- merge(RWP_vep, all_snp_annotations, by = "Gene", all = T)  

all_snp_GO <- all_snp_annotations %>% 
  filter(V3 == "GO")

# write.csv(all_snp_GO, "all_snps_GO_all_snps.csv")
all_snp_GO_one_snp_per_gene <- read.csv("all_snps_GO_one_snp_per_gene.csv", sep = ",", header = T)
write.csv(table(all_snp_GO_one_snp_per_gene$V2), "all_snp_GO_ID_table.csv")
# write.csv(table(all_snp_GO_one_snp_per_gene$V4), "all_snp_GO_term_table.csv")

select_GO_counts <- read.table("outlier_snp_GO_ID_table.csv", sep = ",", header = T)
all_GO_counts <- read.table("all_snp_GO_ID_table.csv", sep = ",", header = T)

merged_GO_counts <- merge(all_GO_counts, select_GO_counts, by = "Var1", all = T)
merged_GO_counts[is.na(merged_GO_counts)] <- 0

### Fisher's test for over-representation of GO categories
table_maker_func <- function(x){
  matrix(c(merged_GO_counts[x, 2], 
           (sum(merged_GO_counts$Freq.x) - merged_GO_counts[x, 2]),
           merged_GO_counts[x, 3],
           (sum(merged_GO_counts$Freq.y) - merged_GO_counts[x, 3])),
         nrow = 2,
         dimnames = list(c(merged_GO_counts[x, 1], "Other"), 
                         c("All", "Selected")))
}  

for(i in 1:1115){
  x <- table_maker_func(i)
  y <- fisher.test(x, alternative = "less")
  z <- data.frame(Name = row.names(x)[1], p = y$p.value)
  lapply(z, write, "FET_results_GO_categories_all_snps_pvalues.txt", append = T)
}

# Get proportions of outlier and all GOs
all_GO_counts <- all_GO_counts %>% 
  mutate(prop = Freq / sum(Freq))

select_GO <- read.csv("outlier_snp_GO_ID_and_description_table.csv")
select_GO_counts <- select_GO %>% 
  mutate(prop = Freq / sum(Freq))

all_GO_counts_in_outlier <- all_GO_counts %>% 
  filter(Var1 %in% select_GO_counts$GO_ID)

all_and_outlier_GO <- data.frame(GO_ID = select_GO_counts$GO_ID, GO_description = select_GO_counts$GO_description, prop_all = all_GO_counts_in_outlier$prop, prop_outlier = select_GO_counts$prop)  
melt_all_and_outlier_GO <- melt(all_and_outlier_GO)

GO_outlier <- ggplot(melt_all_and_outlier_GO, aes (x = GO_description, y =value)) +
  geom_bar(aes(fill = variable), stat = "identity", position = position_dodge()) +
  coord_flip() +
  theme_Publication() +
  # theme(axis.text.x = element_text(angle = 45, vjust = 0.5, hjust = 0.5)) +
  scale_y_continuous(name = "Frequency", breaks = c(0.02, 0.04, 0.06, 0.08, 0.1), limits = c(0,0.10)) +
  # scale_x_discrete(limits = rev(levels(melted$Consequence)), breaks = c("splice_acceptor_variant", "splice_donor_variant", "stop_gained", "stop_lost",
  #                                                                       "start_lost", "missense_variant", "splice_region_variant", "stop_retained_variant",
  #                                                                       "synonymous_variant", "5_prime_UTR_variant", "3_prime_UTR_variant", "intron_variant",
  #                                                                       "upstream_gene_variant", "downstream_gene_variant", "intergenic_variant"),
  #                  labels = c("Splice acceptor variant", "Splice donor variant", "Stop gained", "Stop lost",
  #                             "Start lost", "Missense variant", "Splice region variant", "Stop retained variant",
  #                             "Synonymous variant", "5' UTR variant", "3' UTR variant", "Intron variant",
  #                             "Upstream gene variant", "Downstream gene variant", "Intergenic variant")) +
  # geom_signif(comparisons = list(c("Missense variant", "5' UTR variant", "3' UTR variant", "Downstream gene variant", "Intergenic variant")),
  #             map_signif_level = T) +
  scale_fill_manual(breaks=c("prop_all", "prop_outlier"),
                    labels=c("All SNPs", "Outlier SNPs"),
                    values = pal_nejm()(8)) +
  guides(y_continuous = FALSE) +
  guides(x_discrete = FALSE) +
  theme(legend.title = element_blank(), legend.position = c(0.9, 0.95))

# geom_bracket(
#   xmin = group1, xmax = group2,
#   y.position = 0.05, label = "*",
#   tip.length = 0.01, coord.flip = T)
# geom_signif(stat="identity",
#             data=data.frame(x=c(0.875, 1.875), xend=c(1.125, 2.125),
#                             y=c(0.25, 0.15), annotation=c("**", "NS")),
#             aes(x=x,xend=xend, y=y, yend=y, annotation=annotation))

GO_outlier
```
